# Docking 14 million virtual isoquinuclidines against the mu and kappa opioid receptors reveals dual antagonists-inverse agonists with reduced withdrawal effects

**DOI:** 10.1101/2025.01.09.632033

**Authors:** Seth F. Vigneron, Shohei Ohno, Joao Braz, Joseph Y. Kim, Oh Sang Kweon, Chase Webb, Christian Billesbølle, Karnika Bhardwaj, John Irwin, Aashish Manglik, Allan I. Basbaum, Jonathan A. Ellman, Brian K. Shoichet

## Abstract

Large library docking of tangible molecules has revealed potent ligands across many targets. While make-on-demand libraries now exceed 75 billion enumerated molecules, their synthetic routes are dominated by a few reaction types, reducing diversity and inevitably leaving many interesting bioactive-like chemotypes unexplored. Here, we investigate the large-scale enumeration and targeted docking of isoquinuclidines. These “natural-product-like” molecules are rare in the current libraries and are functionally congested, making them interesting as receptor probes. Using a modular, four-component reaction scheme, we built and docked a virtual library of over 14.6 million isoquinuclidines against both the µ- and *κ*-opioid receptors (MOR and KOR, respectively). Synthesis and experimental testing of 18 prioritized compounds found nine ligands with low µM affinities. Structure-based optimization revealed low- and sub-nM antagonists and inverse agonists targeting both receptors. Cryo-electron microscopy (cryoEM) structures illuminate the origins of activity on each target. In mouse behavioral studies, a potent member of the series with joint MOR-antagonist and KOR-inverse-agonist activity reversed morphine-induced analgesia, phenocopying the MOR-selective anti-overdose agent naloxone. Encouragingly, the new molecule induced less severe opioid-induced withdrawal symptoms compared to naloxone during withdrawal precipitation, and did not induce conditioned-place aversion, likely reflecting a reduction of dysphoria due to the compound’s KOR-inverse agonism. The strengths and weaknesses of bespoke library docking, and of docking for opioid receptor polypharmacology, will be considered.

## Introduction

The size of readily accessible virtual chemical libraries now exceeds 75 billion make-on-demand molecules that can be synthesized and delivered within weeks. This has expanded the space of ligands virtual screening can sample, improving the quality, hit rate, and potencies of docking-prioritized compounds^1-3^. Still, despite their size, these make-on-demand libraries do not capture the true range of chemical space accessible by modern synthetic methods, owing to their emphasis on compounds formed via well-studied two- and three-component reactions that require minimal purification. This has led to libraries dominated by molecules synthesized via amide coupling reactions (**Figure 1B-D**). Amide couplings allow for the sampling of the many diverse chemotypes decorating the amine and carboxylic acid building blocks (the two largest synthon classes contributing to the make-on-demand libraries^4^), yet biasing toward them leaves many interesting, bio-like, and newly synthetically accessible areas of chemical space unexplored^5-7^.

**Figure 1.**
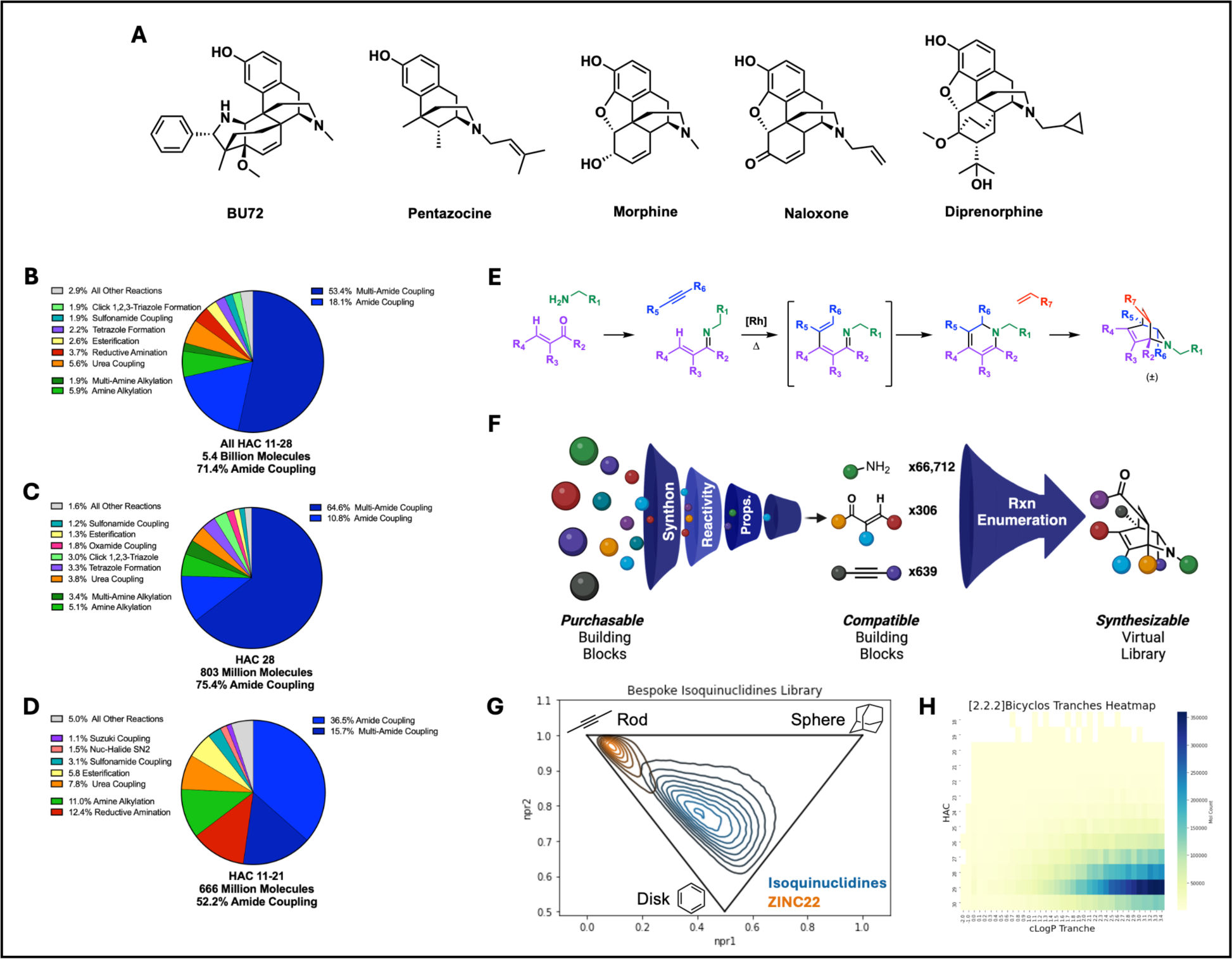
Enumeration of a tangible virtual library of [2.2.2]bicyclic isoquinuclidines from purchasable building blocks. **A.** Example structures of known ligands for MOR. **B-D** Breakdown of the reactions used to enumerate the Enamine REAL Library dominated by amide coupling reactions across different heavy atom count ranges. **E.** Synthetic route for the synthesis of [2.2.2]bicyclic isoquinuclidines from modular building blocks.^30, 31^ **F.** Cartoon diagram of the bespoke library enumeration pipeline beginning from purchasable building blocks, filtering to only those compatible with the reaction, before combining combinatorically to furnish the final tangible bicyclic compounds. A web-based tool for bespoke library enumeration around any reaction is available at ChemistryCommons. **G.** Inertial plot of 5% of the isoquinuclidine library (blue) compared to an equal number of representatives from the ZINC22 library (orange) property matching in HAC and clogP ranges. **G.** Heatmap of clogP vs HAC showing chemical properties of the library.

Physical combinatorial chemical libraries synthesized around a modular reaction scheme^8^ have revealed ligands with more complex structures, both in the identification a new classes of ligands for a target^9^ and in the finding analogs of known ligands with improved properties^10-12^. However, they are often limited in size by the practicalities of synthesis and curation^8^, leaving much of the chemical space made tangible by the scheme unincorporated.

In principle, molecular docking is well-suited to explore these under-represented spaces, as large bespoke libraries can be enumerated and screened virtually, prioritizing the best ranking molecules for synthesis and testing^13-16^. Even so, constructing these bespoke libraries is time consuming, and so to be pragmatic and potentially impactful, they should be characterized by three features: their molecules are under-sampled in standard make-on-demand libraries, bespoke library members are readily synthesizable, and they represent bio-like chemotypes.

Isoquinuclidines are among the molecules that fit these criteria (**Figure 1**). Their amine-containing [2.2.2]bicyclic scaffold is topologically complex, with high sp^3^ content and a caged core that confers more disk-to-sphere like shapes, which are rare among the ultra-large libraries^6^ that are typically more rod-like (**Figure 1**). An efficient reaction scheme gives improved access to these congested compounds, constructed via a one-pot cycloaddition of a rigidifying bridgehead to a modularly constructed dihydropyridine. This modularity and wide substrate scope make diverse isoquinuclidines with up to seven accessible points of differentiation in orthogonal directions synthetically accessible at scale. The chemically dense isoquinuclidine scaffold resembles many aminergic bioactive molecules^17^, appearing well-suited to target peptide receptors like the Mu and Kappa opioid receptors (MOR and KOR, respectively). Both MOR and KOR have large, solvent exposed, non-linear orthosteric sites known to bind to multiple ligand classes. These ligands include many caged, cationic nitrogen-containing compounds, such as diversly-decorated classical morphinians, e.g., morphine, naloxone, and buprenorphine, and those with larger or smaller ring systems, e.g., BU72, and pentazocine^18^ (**Figure 1A**). Agonists of MOR, notably morphine and fentanyl, confer almost unmatched analgesia across a wide range of pain conditions, yet have major liabilities of reinforcement, tolerance, constipation, and addiction^19, 20^. Antagonists of MOR, including naloxone, can act as reversal agents for opioid overdose, yet induce major aversive withdrawal symptoms^19, 20^. To improve side effect profiles of MOR modulating therapeutics, investigators have sought molecules that also antagonize KOR^21-23^. Blockade of KOR is thought to counter the stress-induced compulsive drug seeking involved in the pro-addictive profiles of MOR agonists, and the harsh dysphoria and aversive responses that accompany withdrawal precipitation by MOR antagonists^19, 24-27^.

Believing that isoquinuclidines might be amenable for binding to both MOR and KOR, and that their highly tunable scaffold may confer interesting pharmacology^28^, we sought to create a large library of derivatives that could be readily accessed. Based on the scope of an efficient modular synthesis, we curated a set of commercially available and reaction-compatible building blocks to enumerate a library of 14.6 million isoquinuclidines with drug-like properties^29^ (cLogP < 3.5, heavy atom count (HAC) < 30). These structures were then prepared for docking by being computationally built in three dimensional form, computing hundreds of conformations for each, along with partial atomic charges and solvation energies. This library of disk/sphere-shaped molecules **(Figure 1G)** was docked against MOR and KOR specifically seeking ligands with polypharmacology against both receptors; compounds that would either activate or deactivate MOR while simultaneously antagonizing KOR. We consider how the modularity and three-dimensionality of the library lend themselves to identifying and optimizing ligands with polypharmacology in these complex binding sites. Additionally, we examine the constraints in diversity and function from exploring only a single core scaffold in library docking versus the multiple scaffolds that are present in the larger make-on-demand libraries.

## Results

### Dominance of amide coupling in the construction of the Enamine REAL database

While isoquinuclidines are well-represented among natural products and bioactive compounds^17^, including ibogaine and dioscorine, few are found among the tangible libraries. For instance, of 5.4 billion Enamine REAL compounds, only around 95,000 fell into this class, even loosely defined. As these isoquinuclidines would be preinstalled on building blocks, this further limits their diversity, topological complexity, and potential for derivatization of decorating groups. If we represent REAL molecules by principal moments of inertia, most may be characterized as rod-like and to a lesser extent disk-like; few are sphere like (**Figure 1G**). Both the sparse representation of isoquinuclidines and the bias toward rod-like shapes among tangible molecules at least partly reflects the dominance of only a few reactions underlying the REAL set. Though almost 200 reactions contribute to creating this tangible virtual library, most library compounds are synthesized via amide-coupling (**Figure 1B-D**). Indeed, of all the reactions used to synthesize the 5.41 billion molecules, 71.4% can be classified as amide coupling reactions, with 4.04 billion of all library members containing a formed amide bond. Formation of a urea, which is closely analogous to an amide bond, contributes the third largest fraction. By nature, amide-coupling combines constitutive building blocks linearly, contributing to a bias toward rod-like compounds^6^.

### A tangible isoquinuclidine library

Given their scarcity in the tangible libraries, bio-likeness, and their dense, sphere-like topology, we built a library of synthetically-feasible isoquinuclidines, guided by a modular synthetic route^30, 31^ (**Figure 1E**). Although several reaction enumeration tools are available^15, 32-34^, these can be difficult to practically apply to large-scale libraries and are not always amenable to new reactions. Accordingly, we created a python-based bespoke library building pipeline adaptable to any chemical transformation, organizing output library members in DOCK compatible formats. Final bicyclic compounds are furnished from 4 input building block types (synthons): primary amines and anilines, α,β-unsaturated carbonyls, internal alkynes, and activated alkenes. Library enumeration occurred in two steps: building block compatibility filtering followed by reaction enumeration. Building block filtering consisted of taking all purchasable building blocks and removing those that did not pass SMARTS-based inclusion and exclusion rules specific for each synthon class (see **Supplementary Table 2**). Inclusion SMARTS filters ensured all building blocks contained only the correct reactive synthon, while exclusion SMARTS filters removed those building blocks that contained groups incompatible with the reaction; typically either those that would result in undesired side reactions, final properties, such as those with certain PAINS moieties^35^, or too many rotatable bonds. These reaction-compatible building blocks were then combinatorically enumerated into furnished isoquinuclidine library members with reaction SMARTS (See **Supplementary Table 3**). To increase the confidence in synthesis success and maintain low molecular weights, *N*-methyl acrylamide and methyl acrylate were the only activated alkenes chosen for the rigidifying bridgehead elements. To improve the drug-likeness of the final library, a final filter excluded all bicyclics with greater than 30 heavy atoms and a cLogP > 3.5. This resulted in a virtual library of 14.6 million virtual [2.2.2]bicyclic isoquinuclidines. Consistent with the congested functionality and high three-dimensionality of its molecules, analysis of the principal moments of inertia of this bespoke, tangible library was centered between sphere- and disk-like geometries, unlike property-matched representatives of the general tangible ZINC22 library (**Figure 1G**). Attempting to map our bespoke library to Smallworld^6^ (NextMove Software, https://www.nextmovesoftware.com/smallworld.html), which allows one to rapidly search in the much larger 75 billion molecule space, only 290,000 members could be indexed, indicating nearly 98% of the library contained a unique anonymous graph to any other chemical library.

### Molecular Docking of the isoquinuclidine library against the *µ*- and *k*-opioid receptors

The isoquinuclidines seemed well-suited to bind several peptide recognizing G protein coupled receptors (GPCRs), particularly the opioid receptors. While different topologically from the classic morphinan ligands of these receptors, the [2.2.2]bicyclic structure and cationic nature of the isoquinuclidines sterically and electrostatically resembled them. Accordingly, we docked the isoquinuclidine library against both MOR and KOR, seeking molecules that would act as either agonists or antagonists of the former and antagonists of the latter. In either case, molecules with MOR activity that also block KOR might have phenotypic advantages over ligands selective for either receptor individually.

Seeking KOR antagonists, we used the inactive state structure of the receptor (PDB ID 4DJH^36^) for docking. For MOR, we chose an active state structure (PDB ID 5C1M^37^), preferring agonists but knowing that docking against a particular state of a GPCR, active or inactive, could return ligands with the opposite function (i.e., agonists from docking against antagonist structures or antagonists from docking against active structures)^38-42^. For both receptors, control calculations were conducted to optimize electrostatic and desolvation boundaries, improving docking enrichment of known ligands against property matched decoys^43^ and extrema sets of molecules^44, 45^. The enrichment achieved in these control calculations were consistent with earlier campaigns against MOR. While the annotated known ligands—e.g. fentanyl, methadone, and classic morphinans—docked in geometries consistent with their experimental structures using the optimized potential grids, an initial screen of the full virtual isoquinuclidine library led to what we considered unreasonable poses. Accordingly, we further optimized the hot spots (‘matching spheres’) using the coordinates of [2.2.2]bicyclic cores from a few of the well-posed docked isoquinuclidines, biasing sampling of the core scaffold towards key recognition residues.

The full 14.6 million virtual isoquinuclidine library was docked against the optimized MOR and KOR models. For MOR, each isoquinuclidine was fit in an average 25,760 orientations, with each sampling an average of 220 conformations; a total of 5.81 trillion complexes were scored, taking a total of 14,411 core hours (about half a day on a ∼1000 core cluster). Similar sampling and timings were observed for the KOR docking screen. Because MOR activity would drive the underlying pharmacology we sought—analgesia for agonists, opioid reversal for antagonists—with KOR activity modulating side effects, our strategy for polypharmacology focused first on identifying the top compounds against MOR, and then selecting molecules that also scored highly against KOR. For each receptor, the top ranking one million compounds were filtered with LUNA interaction fingerprints^46^ removing compounds with poses that did not ion-pair with the key recognition aspartate of TM3, contained unsatisfied hydrogen bond donors, or more than three unsatisfied hydrogen bond acceptors. This left 27,915 compounds against MOR.

Of the 27,915 docked compounds passing the MOR ionic filter, 14,164 also ranked well and made the equivalent aspartate salt bridge in the KOR docking screen. With knowledge of known ligands often participating in a water-mediated hydrogen bond network to Tyr148, we further filtered the number of compounds to 2,787 (MOR) and 3,781 (KOR) that contained an oxygen or nitrogen atom within 4Å of this solvated region. The MOR docked compounds were clustered with LUNA interaction fingerprint (IFP)-based Tanimoto coefficient (Tc) > 0.35, finding 406 unique cluster heads that were then inspected manually. We ultimately prioritized 48 isoquinuclidines by visual inspection in MOR, of which 19 contained cluster members that also passed visual inspection in KOR, the “polypharmacology cohort” (**Figure 2B**). As a previous docking study of the opioid receptors described a high false negative rate for polypharmacology, in which compounds that only docked well at only one receptor were found experimentally to have good affinities for both^47^, we retained the other 29 isoquinuclidines as a separate “MOR only cohort” against which our success in predicting polypharmacolgy can be compared.

**Figure 2.**
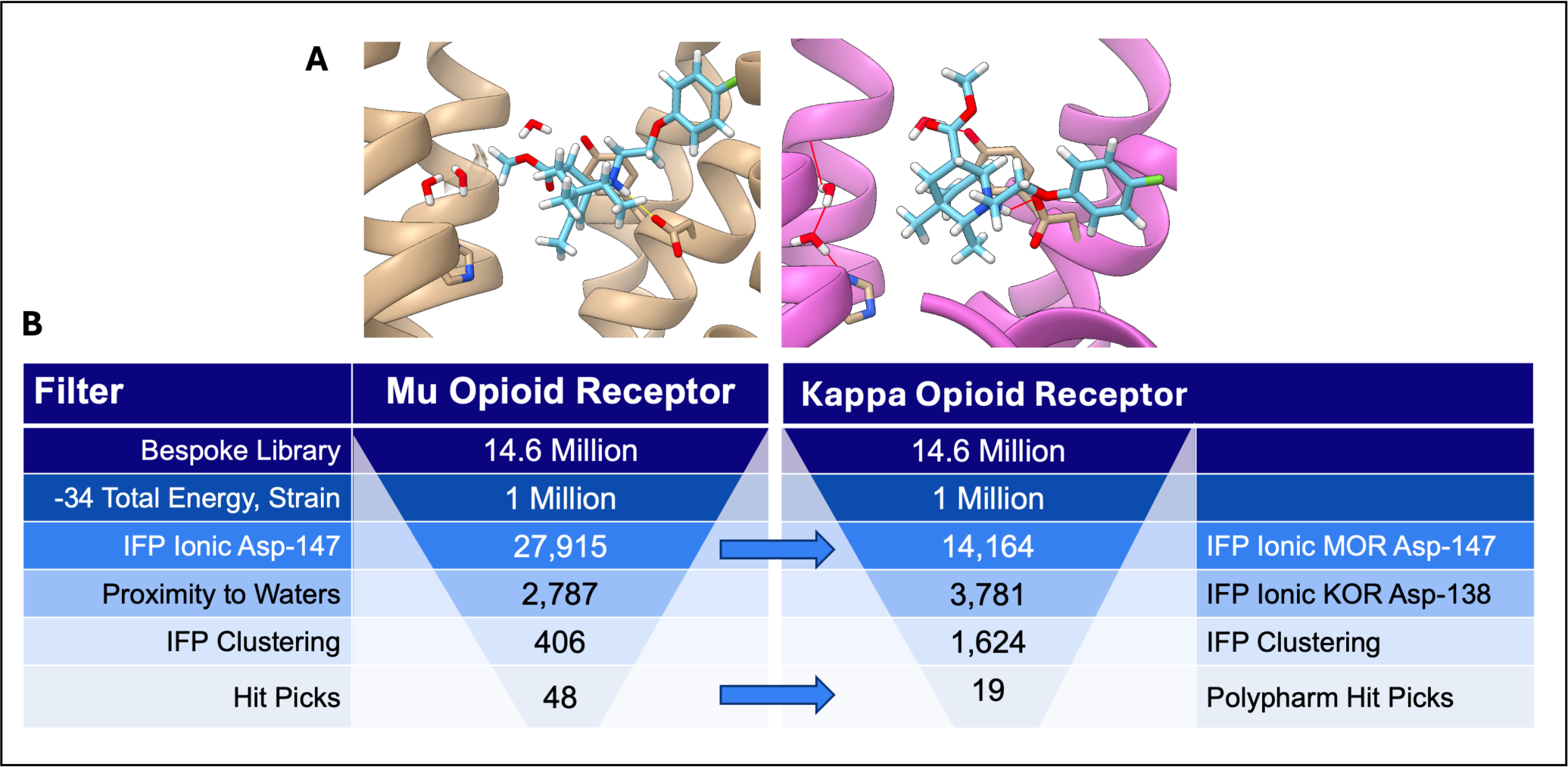
Molecular docking for polypharmacology against the Mu and Kappa Opiate receptors. **A.** Docking poses of the same ligand making similar interactions in both MOR (left) and KOR (right) **B.** Diagram overviewing the docking of 14.6 million isoquinuclidines and subsequent filtering for polypharmacology for both receptors.

### Synthesis and Experimental Testing of Prioritized Isoquinuclidines

Of the docking-prioritized virtual isoquinuclidines, eighteen were synthesized; nine from the polypharmacology cohort, which considered both MOR and KOR docking, and nine from the MOR-only cohort. Synthesis was performed as previously described^30, 31^, with first imine condensation between the chosen primary amine and α,β-unsaturated carbonyl building blocks. Dihydropyridines were formed via a one-pot Rh(I)-catalysed C–H addition of the imine to the desired alkyne building block with subsequent *in situ* electrocyclization. Without work-up or isolation, the alkene was added to the reaction mixture to fully furnish the final desired [2.2.2]bicyclic core via Diels-Alder cycloaddition. While all molecules contained the same isoquinuclidine scaffold, sidechains were diverse, resulting in bicyclics with both one or two potentially basic amines, and an array of alkyl, aromatic, and heteroaromatic groups (**Table 1)**. The amine and enone building blocks were the most frequently varied in this initial set, with most using 2-butyne as their alkyne component. This was not surprising as amines remain the largest class of purchasable building blocks and thereby impart their diversity on the *N*-substituent of the final compound. Intriguingly, exploration off the alkyne appeared to be restricted by the geometry of the opiate receptor, as the binding site is narrower in the direction orthogonal to the typical placement of the N-substituent.

**Table 1.** Initial hits efficiently synthesized from purchasable building blocks. Breakdown of initial hits displaying experimentally determined K_i_ binding affinity agains the Mu (top, blue) and Kappa (bottom, pink) opioid receptors. The modular building blocks used in the synthesis are also shown. * All initial hits were tested as mixtures of (±) enatiomers, here one representative enantiomer structure is depicted for simplicity. † K_i_ estimates calculated from the Cheng-Prusoff equation based on a single point at 33µM ligand concertation found to be near the IC50. ‡ Compounds were a mixture of pseudo-enantiomers resulting from the single enantiomer of the chiral amine input (see **SI section 5**). Here only one pseudo-enantiomer is depicted for simplicity.

| Compound | Structure * | Affinity | Amine BB | Enone BB | Alkyne BB | Acyl BB |
| --- | --- | --- | --- | --- | --- | --- |
| (+)-#33 |  | MOR<br>KOR<br>0.98μM<br>1.2μM |  |  |  |  |
| (±)-#19‡ |  | MOR<br>KOR<br>8.9μM<br>4.6μM |  |  |  |  |
| (±)-#57 |  | MOR<br>KOR<br>5.0μM<br>1.1μM |  |  |  |  |
| (±)-#61 |  | MOR<br>KOR<br>3.1μM<br>3.8μM |  |  |  |  |
| (±)-#11‡ |  | MOR<br>KOR<br>2.3μM<br>0.40μM |  |  |  |  |
| (±)-#56 |  | MOR<br>KOR<br>8.8μM<br>--- |  |  |  |  |
| (±)-#03 |  | MOR<br>KOR<br>14μM†<br>--- |  |  |  |  |
| (±)-#09 |  | MOR<br>KOR<br>14μM†<br>12μM† |  |  |  |  |
| (±)-#05 |  | MOR<br>KOR<br>16μM†<br>8.0μM |  |  |  |  |
| (±)-#001 |  | MOR<br>KOR<br>7.1μM<br>--- |  |  |  |  |

The eighteen synthesized isoquinuclidines were first tested for MOR receptor binding in single-point ^3^H-naltrexone radioligand displacement assays at a ligand concentration of 33µM. Against MOR, 9 of the 18 compounds displaced ^3^H-naltrexone to more than 50% of the DAMGO positive control, a 50% hit rate at this concentration of the isoquinuclidines (**Table 1**, **Figure 3A**). We note that isoquinuclidines containing the *N*-methyl amide bridgehead had a lower hit rate than esters at this position. Consistent with the idea that isoquinuclidines are well-suited to the peptide site of the opioid receptors, the simplified isoquinuclidine **#001**, which lacks all elaborated side chains, was also synthesized and tested, revealing an apparent *K_i_* of 7.1µM (**Table 1**). Seven of the nine MOR hits showed polypharmacology, also being hits against KOR using the same criteria; including three of the four hits from the MOR only cohort. Full dose response curves of the initial hits against both MOR and KOR revealed *K_i_* values ranging from 16 to 1 µM against both receptors (**Figure 3B,C)**. In functional live-cell GloSensor cAMP assays, all nine of the new isoquinuclidines acted as antagonists against MOR, lacking the inhibition of cAMP biosynthesis that would indicate G_i_ signal activation (**Figure 3D)**, a point to which we will return. We set out to optimize the most potent of these, compound **#33**, with binding affinities of 0.98 µM and 1.2µM to MOR and KOR respectively.

**Figure 3.**
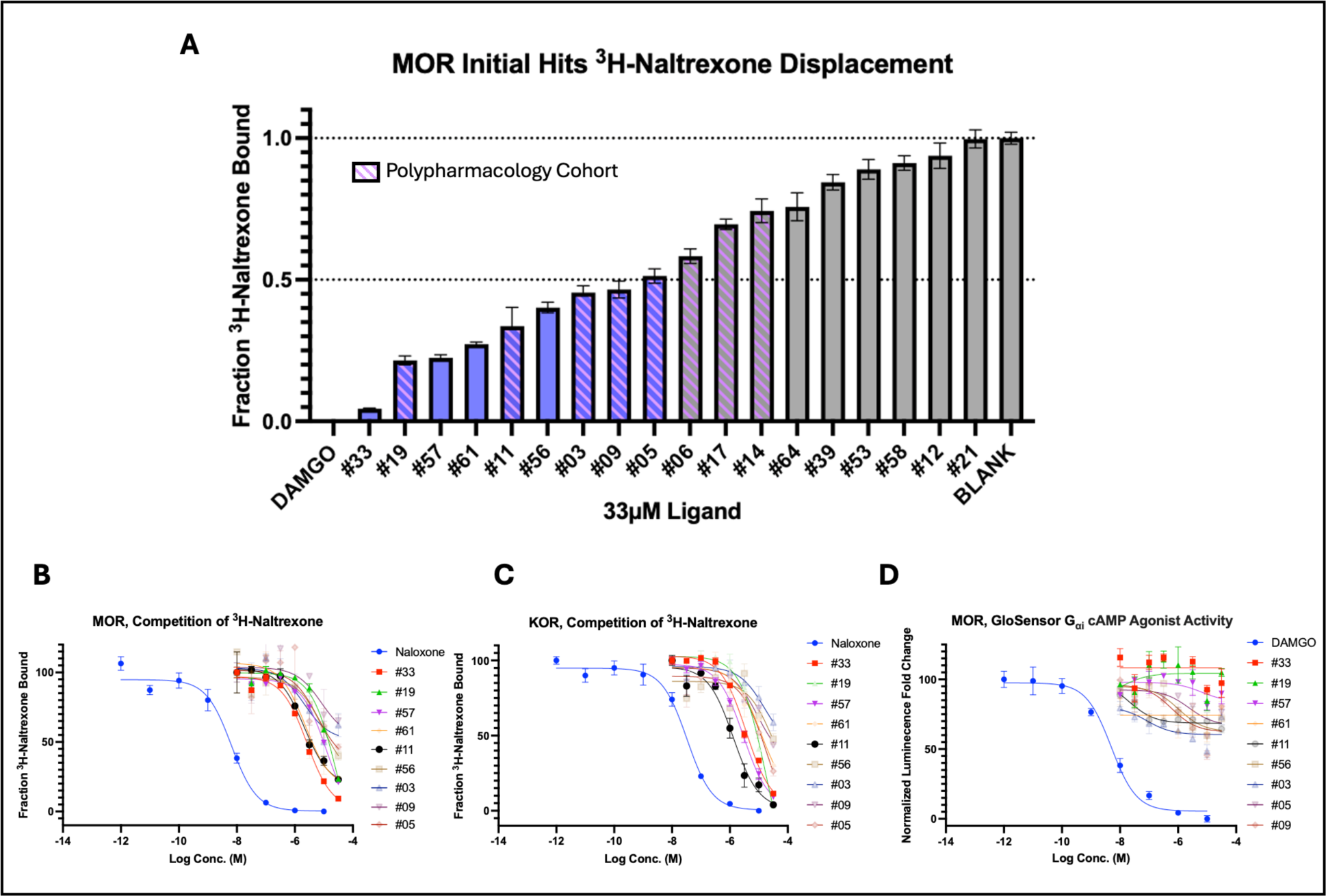
Virtual library of [2.2.2]bicyclic isoquinuclidines identifies a new opioid receptor chemotype. **A.** Single point ^3^H-naltrexone radioligand displacement assay results of initial compounds againts MOR. Ligands able to reach 50% displacement relavive to DAMGO at 33µM are considered hit compounds. **B.** Radioligand competition of ^3^H-naltrexone dose response cuve for initial hits againts MOR, DAMGO normalized. **C.** Radioligand competition of ^3^H-naltrexone dose response cuve for initial hits against KOR, salvinorin A normalized. **D.** Live cell GloSensor cAMP assay in MOR expressing HEK293T cells of initial hits showing lack of agonist activity, DAMGO normalized.

### Initial Compound Optimization & Structure Determination

In early optimization we probed each of the amine, enone, and alkene building block-derived substituents of **#33**. Fifteen analogs were synthesized and assayed for radioligand displacement against MOR (see **SI section 4**). Simplification of the amine building block by removing the auxiliary amine and reducing carbon chain length had little effect on binding, while substituting the methyl ester for a hydrogen, nitrile, acetophenone, or methylamide all greatly reduced affinity. Unexpectedly, removal of the phenol hydroxyl group of **#33**, which in classic opioid receptor ligands provides substantial affinity via interactions with an ordered water network^37, 48^, had little impact. This suggested that it was poorly placed in the site; optimization of this phenol in later rounds became central for affinity improvement.

To understand these effects and inform targeted optimization^28, 49-51^, we determined the structure of **#33** in complex with MOR by single particle cryo-EM (PDB ID: 9MQH). The antagonist structure was determined to a global nominal resolution of 3.9Å with the use of a receptor fusion complex with the nanobody Nb6M that engages with the receptor in an inactive state^52^. While not to a resolution capable of unambiguously assigning the ligand pose, with the overlay of a reasonable docked structure, four key observations could still be made: the phenol hydroxyl group is at a suboptimal angle for interaction with the water network compared to the poses of other known ligands, the methyl ester substituent is angled down towards the sodium binding site sub-pocket, it is the isoquinuclidine nitrogen, not the auxiliary nitrogen, that makes the salt bridge interaction with Asp147, and N-substituents are at a proper angle to extend towards a larger hydrophobic subpocket (**Figure 5**).

**Figure 5.**
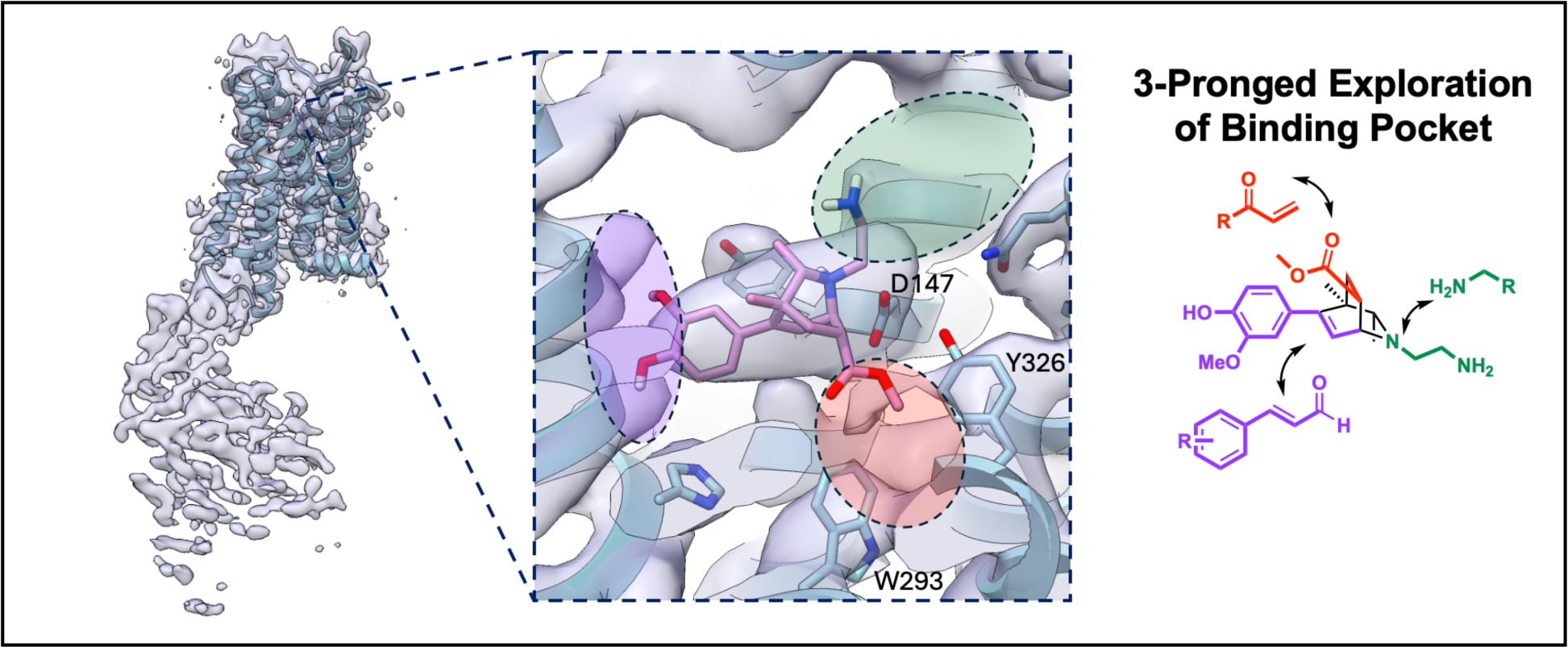
CryoEM structure with docked overlay of initial hit compound alludes to isoquinuclidne binding mode and informs pocket exploration-based optimization. Overlay of a docked pose of **#33** into the cryoEM density. The modular nature of the isoquinuclidine synthesis can be used to make targeted optimization towards specific highlighted binding site regions.

Inspired by the phenol group placement of ligands such as PZM21^39^ and BU72^37^ that we could now overlay onto our cryoEM density, we believed that a meta-substituted phenol would be better suited for water network interaction. Upon synthesis of compound **#016**, this change from para- to meta-substitution improved *K_i_* by 1.5log units down to 37nM in MOR and 7.3nM in KOR (**Table 2**). To guide the next round of compounds, we generated and docked a small set of isoquinuclidines built with our now confident phenolic enone building block, only changing the nitrogen substituent. Here we took the amine building blocks we had on hand from the synthesis of initial hits and a small number inspired by known ligands containing a hydrophobic group separated from the amine by a short linker. Seven of these compounds were synthesized (compounds **#017 - #023**) using the procedure detailed above and all were stereochemically purified by chiral column HPLC to obtain fourteen pure enantiomers (see **SI section 5**). Of these, ten were ligands with binding affinities below 50 nM against both receptors, and nine bound in the single-digit nM to high pM range (**Figure 6A**, **Table 2**). Another apparent contributor to affinity was the addition of a methyl group to the R3 position on the isoquinuclidine core, which may restrict free rotation of the phenol ring. The addition of this single methyl further improved MOR potency from 37nM to 7.4nM (**#019_E1**) for compound **#016**, and from 4.2nM to 0.91nM (**#031_E2**) for the pyrazole containing compound #**020**_**E1**. Against KOR, the new isoquinuclidines bound in the same low nM to high pM concentration range, though here the methyl group addition appeared to have no impact on the already potent KOR affinities. In GloSensor cell signaling assays, the isoquinuclidines retained the antagonism of the parent scaffolds, with all advanced compounds dose-dependently competing against the effects of known agonists DAMGO (MOR) and salvinorin A (KOR) tested at agonist EC_80_ concentrations (**Figure 6B**). In these antagonist assays, isoquinuclidine EC_50_s ranged from 2.2nM to 77nM against MOR and 16nM to 1.8µM against KOR (**Table 2**). While all compounds acted as apparently neutral antagonists against MOR, compound **#020_E1** and other analogs (**Figure 6B-C, Table 2**) acted as potent inverse agonists against KOR.

**Figure 6.**
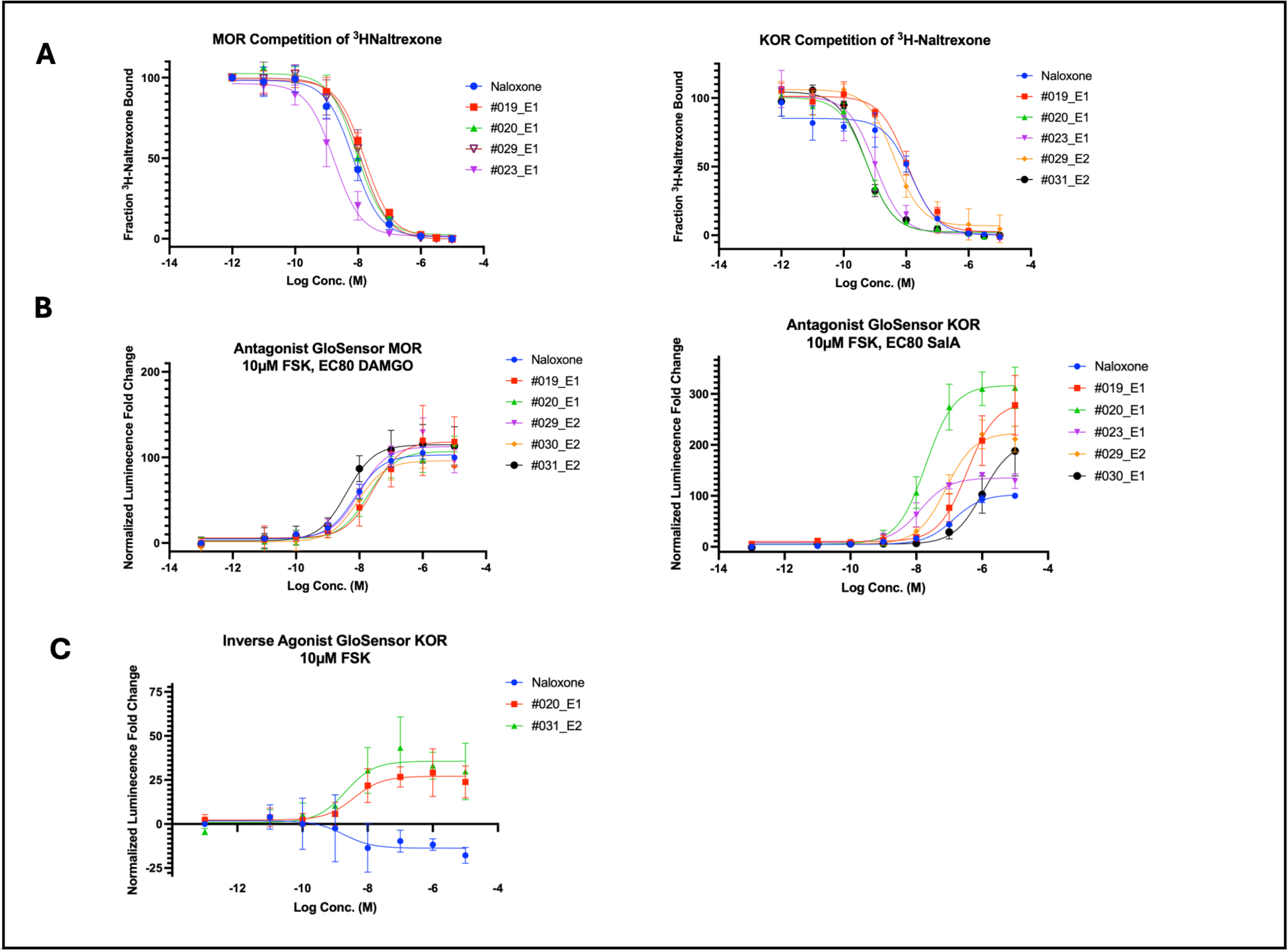
Compound optimization yields a family of potent antagonist bicyclics with polypharmacology. **A.** Radioligand displacement assay of ^3^H-naltrexone competition in MOR (left) and KOR (right) of selected advanced isoquinuclidines. **B.** Live cell GloSensor assay for cAMP in antagonist mode with HEK293T cells expressing MOR (left) and KOR (right). Cells treated with 10µM forskolin and EC80 concentrations of agonist (DAMGO for MOR, SalA for KOR). **C.** Live cell GloSensor assay in agonist mode with cells expressing KOR and treated with 10µM forskolin showing isoquinuclidines **#020_E1** and **#031_E2** blocking basal G_i_ signaling.

**Table 2.** µ- and κ-opioid polypharmacology of advanced isoquinuclidines. Breakdown of advanced isoquinuclidine enantiomers displaying their affinities and EC_50_s at both opioid receptors. E1 and E2 represent individual (+) and (-) enantiomers purified via chiral HPLC in the order of their retention times. * Whereas each enantiomer (E1 and E2) was separated and experimentally tested individually, a single representative enantiomer structure is shown for simplicity.

| Compound | Structure * | Isolated Enantiomer | MOR Ki | KOR Ki | MOR Signaling | MOR EC50 | KOR Signaling | KOR EC50 |
| --- | --- | --- | --- | --- | --- | --- | --- | --- |
| (±)-#016 |  | (±) | 37nM | 7.3nM | --- | --- | --- | --- |
| #017 |  | E1 | 1.7nM | 22nM | Antagonist | 6.0nM | Antagonist | 1800nM |
|  |  | E2 | 5.9nM | 5.5nM | Antagonist | 39nM | Antagonist | 1000nM |
| #019 |  | E1 | 7.4nM | 4.8nM | Antagonist | 32nM | Inverse Agonist | 400nM |
|  |  | E2 | 46nM | 6.9nM | Antagonist | 77nM | Inverse Agonist | 620nM |
| #020 |  | E1 | 4.2nM | 0.29nM | Antagonist | 21nM | Inverse Agonist | 20nM |
|  |  | E2 | 10nM | 1.1nM | Antagonist | 37nM | Antagonist | 380nM |
| #023 |  | E1 | 0.83nM | 0.76nM | Antagonist | 1.5nM | Antagonist | 17nM |
|  |  | E2 | 1.1nM | 0.40nM | Antagonist | 1.5nM | Inverse Agonist | 38nM |
| #028 |  | E1 | 32nM | 9.7nM | Antagonist | 8.1nM | Antagonist | 50nM |
|  |  | E2 | 34nM | 8.8nM | Antagonist | 18nM | Inverse Agonist | 380nM |
| #029 |  | E1 | 7.2nM | 1.2nM | Antagonist | 3.4nM | Antagonist | 16nM |
|  |  | E2 | 4.4nM | 2.2nM | Antagonist | 11nM | Inverse Agonist | 63nM |
| #030 |  | E1 | 3.5nM | 21nM | Antagonist | 5.3nM | Inverse Agonist | 1200nM |
|  |  | E2 | 2.9nM | 1.5nM | Antagonist | 13nM | Inverse Agonist | 230nM |
| #031 |  | E1 | 1.3nM | 2.5nM | Antagonist | 2.2nM | Inverse Agonist | 230nM |
|  |  | E2 | 0.91nM | 0.27nM | Antagonist | 3.9nM | Inverse Agonist | 42nM |
| (±)-#024 |  | (±) | 86nM | --- | --- | --- | --- | --- |
| (±)-#025 |  | (±) | 35nM | --- | --- | --- | --- | --- |

To investigate the methyl ester substituent, we synthesized five additional compounds that differed from other advanced isoquinuclidines only at this bridgehead. Compounds **#024, #025**, and **#030** replace the methyl ester with a methyl ketone, ethyl ketone, and hydrogen respectively, while **#028** and **#029** introduced aromatic heterocycles: 1,3,4-oxadiazole and 1,2,4-triazine. These analogs were for the most part synthesized following the same protocol detailed above (see **SI section 5**). Compounds **#028**, **#029**, and **#030** were additionally stereochemically purified by chiral column HPLC to isolate the six individual enantiomers for testing. Replacement of the methyl ester for an ethyl ketone reduced MOR binding affinity four-fold, and with a methyl ketone, around seven-fold compared to the parent **(±)-#019**. The more major substitutions for a hydrogen, 1,3,4-oxadiazole or 1,2,4-triazine also reduced binding affinity to a similar degree from the more potent parent **#023**, but each still maintained near-single to single digit nM affinities for both receptors. Together, these results show that while the methyl ester is valuable for binding, substitution at that bridgehead is tolerated by both receptors and high affinity can be maintained through compensation by other sidechains.

### Structure Determination of an Optimized Isoquinuclidine Bound to MOR and to KOR

To provide a structural basis for the binding and signaling of these molecules, we determined the cryoEM structures of MOR and KOR in complex with compound **#020_E1**, which has sub-25nM antagonism and inverse agonism against MOR and KOR, respectively, and is even more potent by binding. Using a universal nanobody and Fab strategy reported previously for inactive GPCRs^52^, we obtained cryoEM structures of **#020_E1** bound to both human MOR and human KOR. MOR and KOR were individually expressed in Expi293 cells and purified to homogeneity in detergent micelles in the presence of **#020_E1**. To add additional density to assist in particle alignment during data processing, prior to cryoEM grid preparation, receptors were complexed with the universal nanobody Nb6M, a Fab fragment specific for nanobodies (NabFab), and a Fab-specific nanobody (Anti-Fab Nb) that provided additional stability. As Nb6M is specific for the 3rd intracellular loop of KOR, we additionally mutated two residues in MOR ICL3 to enable binding. We obtained global nominal resolutions for the MOR and KOR complexes of 3.3Å and 3.2Å, respectively, with MOR resolved as a monomer and KOR as an anti-parallel heterodimer (MOR PDB ID: 9MQI; KOR PDB ID: 9MQK). Further local refinement around the transmembrane domains resulted in improved resolutions of 3.2Å for MOR and 3.0Å for KOR (MOR PDB ID: 9MQJ; KOR PDB ID: 9MQL). In these structures, both MOR and KOR are in an inactive state, based on observing both the orientation of transmembrane domains and conserved motifs of class A GPCRs being comparable to past inactive state structures (PDB 7UL4 ^52^, PDB 4DJH^36^, PDB 6VI4^53^).

We resolved clear ligand density in the orthosteric binding pocket for both receptors, allowing unambiguous modeling of compound **#020_E1** (**Figure 7A**). The isoquinuclidine adopted similar poses and interactions in both MOR and KOR, including predicted hydrogen bonds between the phenol and the ordered water network around residues Tyr^3.33^ and His^6.52^, and the methyl ester situated between Trp^6.48^ (Trp295 and Trp287 in MOR and KOR, respectively) and Tyr^7.42^ (Tyr328 and Tyr320 in MOR and KOR respectively). Intriguingly, deviations from traditional ligand binding modes were observed in the conserved salt bridge between the isoquinuclidine nitrogen of **#020_E1** and Asp^3.32^ (Asp149 and Asp138 in MOR and KOR, respectively). Instead, the bicyclic core appears to occlude the typical conformer of this key recognition aspartate, forcing it to angle away from the center of the binding site to 4.1Å away from the isoquinuclidine cationic nitrogen (**Figure 7B)**. While this conformation resembles that adopted in a recent structure of MOR bound to an antagonistic extracellular nanobody^54^, it is rare in MOR-drug complexes, and was not represented in our rigid receptor docking model, potentially incurring false negatives in our virtual screen from isoquinucildine cores clashing with Asp^3.32^. The outward angle of this aspartate further pulls down Gln^2.60^ (Gln126 and Gln115 in MOR and KOR respectively) and engages in a triangular, bidentate hydrogen bonding network with Tyr^7.42^ (**Figure 7C**). This interaction network may lead to the antagonistic effect of **#020_E1**, as any outward swinging of TM6 to initiate G-protein signaling would require its disruption.

**Fig.7:**
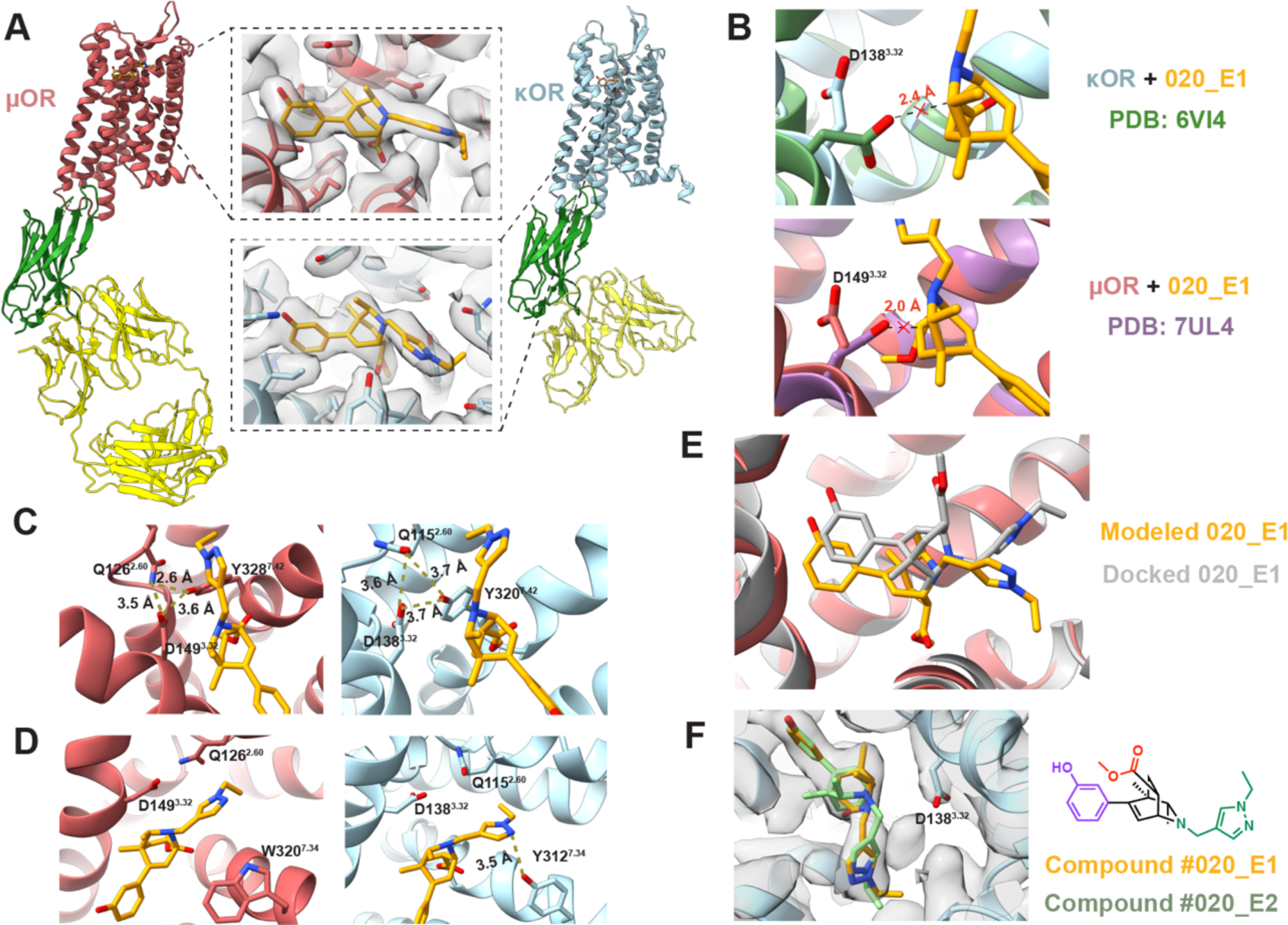
CryoEM inactive state structures of both the MOR and KOR bound to #020_E1. **A.** CryoEM model of MOR (red) and KOR (blue) bound with inactive-state specific nanobody Nb6M (green) and nanobody-binding antibody fragment NabFab (yellow). Insets for both receptors show compound **#020_E1** modelled into the map density of the orthosteric pocket. **B.** Overlay of the cryoEM model with other inactive opioid receptors (Above – KOR + JDTic from PDB 6VI4, Below - MOR + alvimopan from PDB 7UL4) demonstrating the steric clash between D^3.32^ and the bicyclic moiety of **#020_E1**. **C.** Models from both receptors having its D^3.32^ residue engaged in a triangular ionic network with Q^2.60^ and Y^7.42^, further sequestering D^3.32^ away from the ligand. **D.** Insertion of the pyrazole group from **#020_E1** into a rare subpocket, forming an additional salt-bridge with Y^7.34^ in KOR, not seen in MOR due to its replacement by W^7.34^. **E.** Comparison of the modeled **#020_E1** in MOR against the compound docked into the model from PDB 7UL4, revealing different ligand poses. **F.** Model of both enantiomers (alternative conformer in green) fit into the ligand density in KOR, with the chemical structure of **#020_E1** shown on the right for reference.

We had designed **#020_E1** to extend its pyrazole ring towards the hydrophobic subpocket typically occupied by the aromatic rings of other known ligands, e.g. BU72, PZM21, and fentanyl^55^ (PDB IDs 5C1M, 7SBF, 8EF5 respectively), however, the movement of Gln^2.60^ hindered this access, diverting the pyrazole into a rarely seen subpocket between TM4 and TM5 (**Figure 7E**). Although observed once before in an active state structure of MOR stabilized by the selective agonist mitragynine pseudoindoxyl (MP)^28^, itself an unusual indole alkaloid supporting congested functional groups, to our knowledge, no previous ligand has been shown to occupy this site in KOR. Compared to **#020_E1**’s pose in MOR, the pyrazole extends further into the ‘MP pocket’ of KOR and makes an additional hydrogen bond with Y^7.34^. The importance of this residue in ligand recognition has only recently been recognized at the structural level with cryoEM structures of KOR bound with the agonists nalfurafine^49^ or U-50,488H^56^. In MOR, this residue is replaced by a tryptophan, with no equivalent interaction being observed. This may explain the improved affinity observed for KOR versus MOR despite the similar binding mode adopted in both receptors.

The docking pose of compound **#020_E1** within the MOR orthosteric site differed from the experimental pose, with the ligand flipped by nearly 180° (**Figure 7D)**. In the docking model, the methyl ester points up towards the extracellular opening of the binding site, while in the experimental structure it angles downwards toward the center of the receptor in a tight hydrophobic pocket. This discrepancy can be explained by the residue conformations of the inactive receptor structure (PDB 7UL4^52^) that was used for docking. Here, Gln^2.60^ blocks the MP pocket, requiring the pyrazole to extend laterally towards the traditional hydrophobic site and, to accommodate this, forcing the bicyclic core to rotate, placing the methyl ester upwards. Despite these differences, **#020_E1** docks to occupy the same site it does in the cryoEM structure, and its key recognition groups, including the cationic nitrogen and its phenol, interact with the same key residues and waters, reflecting their placement along the axis of rotation of the molecule. Interestingly, modeling both **#020_E1** and its enantiomer, **#020_E2**, into the refined cryoEM density reveal they can both form similar and reasonable poses (**Figure 7F**), with **#020_E2** capable of making the same experimentally observed interactions as **#020_E1**, differing only in the exact angle and rotation of the core isoquinuclidine. This may explain why the (+) and (-) enantiomers of many of the advanced isoquinuclidines bind with similar affinities in the typically highly stereoselective opioid receptors.

### Opioid Withdrawal Pharmacology *in vivo*

The combination of MOR antagonism, conferring an ability to reverse opioid effects, with strong KOR inverse agonism, reversing the dysphoric effects of withdrawal^57-59^, suggested assessing **#020_E1** as an opioid overdose reversal agent, akin to naloxone (Narcan), but potentially with fewer of naloxone’s aversive side effects^60-62^. In pharmacokinetic studies of several of the advanced isoquinuclidines, compound **#020_E1** had among the best overall exposure in the CSF, a proxy for brain free fraction, and retained good coverage for over an hour at 10 mg/kg dosing (see **SI section 7)**. Accordingly, we investigated this molecule’s ability to reverse the analgesic effects of morphine in mice compared to naloxone. In an acute heat nociception assay, a 20mg/kg i.p. dose of morphine significantly increased tail flick latency, a spinal cord reflex that correlates with other pain behaviors. When 30 mg/kg of **#20_E1** was concurrently administered, the morphine-induced analgesia was fully blocked, comparable to the effect of 10mg/kg of naloxone (**Figure 8A)**.

**Figure 8.**
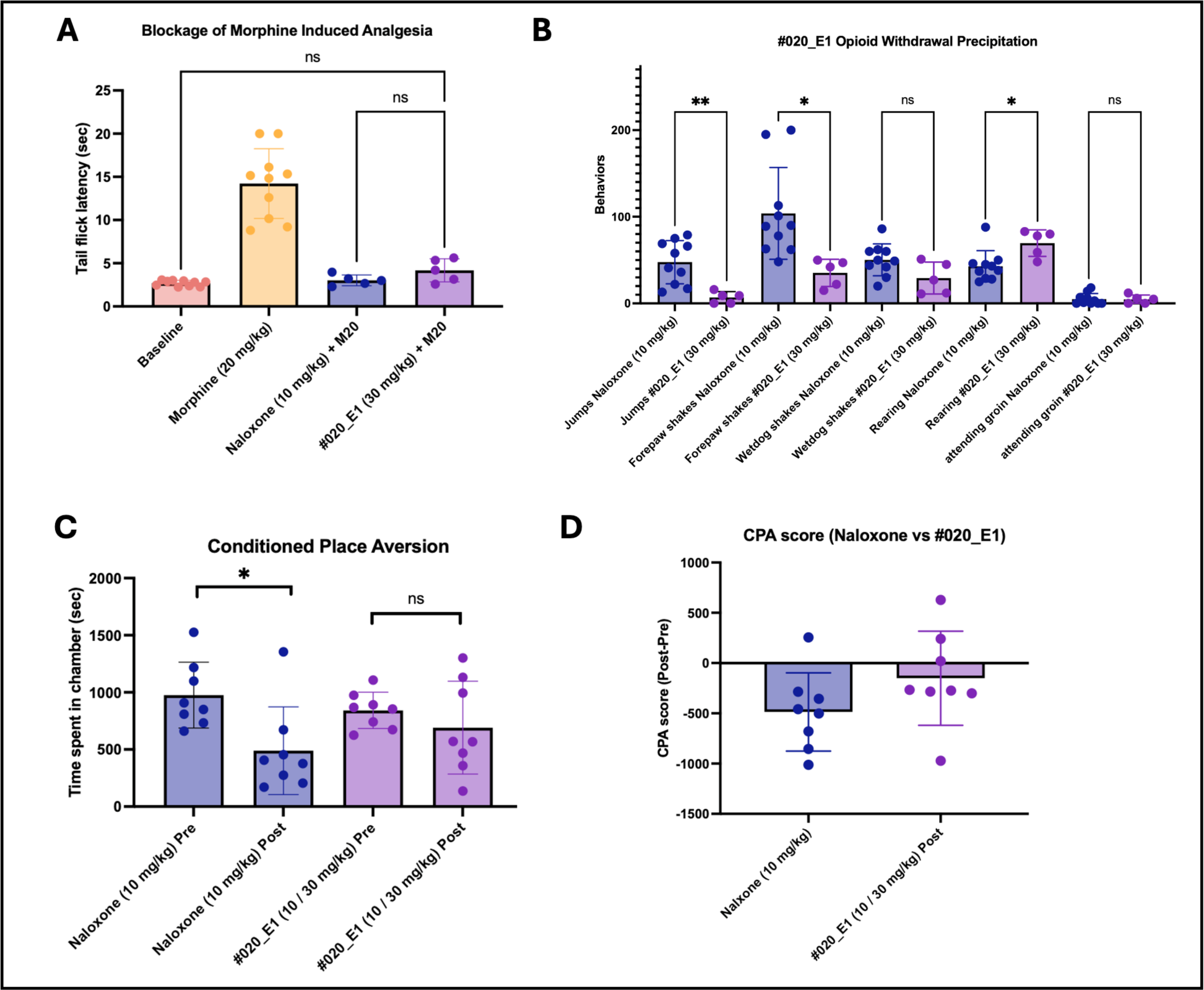
Isoquinuclidine 020_E1 reverses morphine *in vivo* with reduced withdrawal in vivo. **A.** Administration of **#020_E1** blocks morphine-induced analgesia, returning tail flick latency to baseline levels. **B.** Precipitation of opioid withdrawal with **#020_E1** produces fewer withdrawal symptoms than mice administered naloxone for withdrawal precipitation. **C.** Morphine-tolerant mice administered naloxone show a marked decrease in time spent in the antagonist-paired chamber compared to pre-treatment. This avoidance to the antagonist chamber is not observed for mice receiving **#020_E1**. **D.** CPA score, calculated by subtracting the time spent in the drug-paired compartment prior to antagonist treatment, shows that mice administered naloxone avoided the treatment compartment to a greater extent than did the mice that received **#020_E1**, which showed no side preference.

To examine the ability of **#20_E1** to precipitate opioid withdrawal, and the resultant symptoms, we generated morphine tolerant mice following a previously reported protocol^63^. Briefly, we injected mice twice a day for four days with escalating doses of morphine (10 to 75 mg/kg). On day five, opioid withdrawal was induced by administering a single dose of morphine (20 mg/kg) followed 1h later by naloxone (positive control; 10 mg/kg) or **#020_E1** (30 mg/kg). As expected, naloxone induced behaviors consistent with aversive withdrawal, including increased jumping, wet-dog shakes, rearing, and forepaw shakes. Conversely, precipitated withdrawal by **#020_E1** was associated with significantly fewer of these stress associated phenotypes, particularly a notable reduction in escape jumps (**Figure 8B**). Next, to test whether the reduction in opioid withdrawal symptoms was associated with a decreased aversive state, we used a modified conditioned place preference assay in which mice learn to associate one chamber of the apparatus with opioid withdrawal precipitated by an antagonist (either naloxone or **#020_E1**). If mice spend less time in the reversal drug paired chamber, then the compound is considered to be aversive, perhaps dysphoric. As expected, after conditioning, morphine tolerant mice spent significantly less time in the chamber in which opioid withdrawal was precipitated with naloxone. Encouragingly, mice injected with the isoquinuclidine exhibited no significant aversion for the withdrawal precipitation chamber (**Figure 8C, D)**. Taken together, these studies suggest that **#020_E1** is effective at blocking morphine’s activity, and in doing so it induces less severe opiate withdrawal symptoms than does naloxone, presumably due to the isoquinuclidine’s potent KOR inverse-agonism.

## Discussion

The advent of make-on-demand libraries, now exceeding 75 billion molecules, has vastly expanded the range of molecules readily available to the scientific community. However, this growth has been biased towards compounds synthesized via amide coupling reactions, resulting in a predominantly linear library that leaves many complex and bioactive scaffolds underexplored. Here, we sought to investigate one class of such under-represented molecules, [2.2.2]bicyiclic isoquinuclidines, which seemed by topology, physical properties, and derivatization centers well-suited to probing the opioid receptors. Four key observations emerged: **First**, the expansion of the isoquinuclidine reaction generated a virtual library of novel structures. As with a previous tetrahydropyridine bespoke library^14^, none of the isoquinuclidines had an equivalent in the general-purpose library, and 98% had unprecedented anonymous graphs in the Smallworld database, indicating no other indexed molecule contained the same topology. **Second,** the 50% hit rate observed for this library outperformed previous docking screens against the opioid receptors^23, 39, 47^, and is high by docking standards ^37, 64-67^, supporting the value of docking these bespoke libraries against binding sites for which their scaffolds are well-suited.

**Third**, the modular nature of the isoquinuclidine core combined with cryoEM structure determination allowed for targeted chemical modifications to parent structures that much improved potency. Synthesizing only 31 additional isoquinuclidines from an initial low µM hit resulted in a family of single-digit nM to sub-nM ligands to both MOR and KOR stabilizing rare receptor inactive confirmations. **Fourth,** the polypharmacology of compound **#020_E1**, combining potent MOR antagonism and KOR inverse agonism, conferred reversal of opioid analgesia that was comparable to a high dose of naloxone, but was associated with fewer of its aversive effects associated with opioid withdrawal—a phenotype much sought for new overdose medications.

Several cautions merit airing. Although we filtered building blocks for compatibility with the isoquinuclidine synthesis scheme (**Figure 1F**), our SMARTS-based patterns struggled to capture aspects such as strain of the Diels Alder transition state, with many compounds requiring reaction condition optimization, increasing cost and time. Additionally, the synthesis of certain analogs valuable for SAR hypothesis testing, such as **#028**, and **#030**, fell outside the scope of the general reaction scheme, requiring the time-consuming development of new synthetic routes. Overall, 49 isoquinuclidines were synthesized for this campaign. While this led to potent and efficacious leads, this number is a fraction of what is possible to test in a screen of the standard make-on-demand library. In the latter, testing 500 initial docking hits is plausible^2, 3, 65, 68^, as is making dozens of optimization analogs for each of the more active molecules^3, 40, 69^. The limited experimental scale of our bespoke library approach affected our ability to optimize hit compounds with a similar breadth that may have been able to improve pharmacokinetic liabilities and broaden the range of signaling modalities. While we were interested in both MOR agonists and antagonists, only antagonists were found. This result is counter to our experience when testing multiple scaffolds emerging from the make-on-demand libraries, which often reveal multiple ligand functions against a receptor. This finding may reflect a constraint intrinsic to the family of phenolic isoquinuclidines, as supported by the cryoEM structures determined here, and may be a feature commonly encountered with libraries based around a single scaffold. With bespoke libraries, tested molecules are fewer and the timelines longer, but the quality of the compounds is often better: purity is higher, stereochemically resolved compounds are typically explored, and the wrong compound are not synthesized, something that, while rare, can occur in make-on-demand campaigns^3^. It must also be admitted that we did not have a method that would ensure that isoquinuclidines were appropriate for the opioid receptors, instead relying on gross apparent compatibility (shape, derivatization vectors, cationic nature). An unbiased method to identify targets for which a bespoke library or chemistry would be well suited would increase the impact of this approach, particularly when a scaffold has little precedence in compound databases.

These cautions should not obscure the main findings of this study. The enumeration and docking of this 14.6 million molecule bespoke library explored an underrepresented region of chemical space, revealing a class of potent opioid antagonists and inverse agonists. The highly three-dimensional and congested isoquinuclidine scaffold was well suited to probe the non-linear MOR and KOR binding sites. This study illustrates how reaction enumeration, combined with molecular docking, can identify bioactive molecules from regions of chemical space that would otherwise be left out of lead discovery campaigns. Other scaffolds and targets may be well-suited to this computational, structure-based approach, bringing cutting-edge synthetic chemistry to new areas of biology.

## Methods

### Enamine REAL Reaction Counting

The reaction codes of all compounds for the public 2024-03 release of the Enamine REAL catalog were curated and each occurrence of a specific reaction code was counted. These codes were then manually categorized into reaction classes. For example, amide couplings performed with different activating reagents have unique reaction codes but are all included under the “Amide Coupling” class. Different reaction codes that specify the use of subsequent 1-pot deprotections were also categorized under the reaction class of the main coupling reaction. In the cases of a multi-step reaction where more than one main coupling reaction was performed, ie for a 1-pot Amide Coupling and Click 1,2,3-triazole formation, these reactions were counted for both reaction types, ie +1 to the count for “Amide Coupling” and +1 to the count for “Click 1,2,3-triazole formation” for each single occurrence of that reaction code. Reaction codes used for each reaction class can be found in **Supplementary Table 1**.

### Isoquinuclidine Virtual Library Generation

Purchasable building blocks were obtained via SMARTS-based queries of the ZINC22 in-stock building blocks catalog and separated by synthon class, finding in 228,305 amines and anilines, 35,637 enones, and 6,344 internal alkynes. These compounds were then filtered by discarding those compounds with substructure matches to exclusion SMARTs filters using RDkit v2018.09. A full list of the SMARTs patterns used for filtering each synthon class is listed in **Supplementary Table 1**. Building blocks were also removed if their heavy atom count (HAC) exceeded 15. This left 66,712 nucleophilic amines, 306 enones, and 639 internal alkynes to be used in combinatorial reaction enumeration with both N-methyl acrylamide and methyl acrylate as the activated alkene components. All combinations of one building block from each synthon class were combined using the list of reaction SMARTs in Supplementary Table 2. Both the (+) and (-) enantiomers of each synthon combination were generated. Furnished isoquinuclidines were then assessed prior to becoming a library member, removing compounds with HACs > 30, cLogPs > 3.5, and more than 7 rotatable bonds. Ligands were built using the ZINC22 ligand building pipeline^70^ with the only modification being to allow increased sampling of nitrogen stereochemistry interconversions to 3 per ligand using MN-AM Corina software.

### Library Analysis

Rdkit v2018.09 was used for all molecule property calculations including heavy atom count, cLogP calculation. Rdkit v2018.09 was also used to determine normalized principal moments of inertia determination (npr1 and npr2 values) for a random ∼5% of the overall isoquinuclidine library, a total of 683,997 compounds. These were then split into tranches according to their HAC and cLogP. For each ligand, a random ZINC22 molecule from the equivalent HAC-cLogP tranche was also obtained and normalized principal moments of inertia calculated. The distribution of npr1-npr2 values for both the isoquinuclidine library and ZINC22 representatives were visualized using the gaussian kernel density estimate (KDE) function within the Seaborn v0.10.1 python package. The Scott method was used for KDE bandwidth estimation.

### Receptor Model Preparation

Receptors were prepared for docking with DOCK Blaster (https://blaster.docking.org) using the active state structure of the murine µ-opioid receptor structure (PDB: 5C1M) in complex with the agonist BU72, and with the inactive state structure of the human k-opioid receptor structure (PDB: 4DJH) in complex with the antagonist JDTic. A total of 45 binding hot spots (spheres) ^44, 71^ were used based on the binding pose of the receptors’ respective complexed ligand. For MOR, waters were modeled based on high occupancy in MD simulations and precedence amongst class A GPCRs. Parameters from the united-atom AMBER force filed were used to assign partial charges for all receptor atoms. Molecular docking grids used for determining the energy contributions of each term in the DOCK3.8 scoring function were precalculated using an grid-based version of the AMBER force-field for the van der Waals component, and were calculated using the Poisson–Boltzmann method QNIFFT^72^ for the electrostatics component. Context-dependent ligand desolvation grids were generated via an adapted version of the generalized Born method^73^. Prepared receptor grids were evaluated and subsequently optimized based on their ability to prioritize a set of known ligands from decoys molecules with similar chemical properties of the known ligands, yet with topologically different structures, as generated with the DUD-EZ approach^45^.

### Radioligand Binding Assay

Radioligand competition assays were performed using membrane preparations of HEK293T cells transiently expressing either human Mu opioid receptor or Kappa opioid receptor. ^3^H-naltrexone radioligand affinities were measured for each membrane preparation in saturation experiments accounting for non-specific binding. Assays were performed in 96 well V-PP coated plates. Each well contained 200µL of homogenous membrane solution in binding buffer (50mM HEPES, 0.1mM EDTA, 10mM MgCl_2_, 0.1% BSA, pH 7.5), 25µL of 10x ligand dilution, and 25µL of 10x ^3^H-naltrexone, both diluted in binding buffer. Final radioligand concentration was run at 1.2x its K_d_ measured for the membrane preparation used. The assay plate was sealed, protected from light, and incubated for 2h at room temperature. The reaction wells were then vacuum filtered through 0.125% PEI-soaked PerkinElmer glass fiber filtermats followed by five washes with cold wash buffer (50mM HEPES, pH 7.5). Filtermats were then dried, and ^3^H-naltrexone counts were measured via scintillation using MeltiLex B/HS wax scintillant on a Perkin Elmer BetaMax scintillation counter. Results were analyzed on GraphPad Prism v.10, normalizing to the naloxone curve present on each plate and using the ‘on-site fit K_i_’ nonlinear regression equation.

### GloSensor cAMP Accumulation Assay

To assess G_i/o_-mediated cAMP production inhibition, live HEK293T cells at ∼70% confluency in a 10cm tissue culture dish were transiently transfected with a ThermoFischer Lipofectamine system at a 1:5 ratio of human MOR and Promega pGloSensor-22F split-luciferase cAMP biosensor. For KOR, this ratio was 1:4. After at least 22h post transfection, cells were washed with phosphate buffered saline and a small volume of trypsin added to dissociate the cells from the cell culture plate. Cells were suspended in warmed CO_2_-Independent media and pelleted by centrifugation at 275 RCF for 5min. Cells were then resuspended in 10.5mL of warmed CO_2_-Independent media and 50µL was added to each well of two 96 well white, flat bottom, tissue culture treated plates. To each well, 40µL of a warmed 10x Beetle luciferin solution in CO_2_-Independent media was added. Plates were loosely capped and incubated at 37°C for 1h before being moved to room temperature for an additional 1h. Baseline luminescence was then recorded on a ClarioSTAR microbeta plate reader. 5µL of 20x ligand dilutions in room temperature CO_2_-Independent media were added to the wells and allowed to equilibrate for 5min prior to the addition of 5µL of forskolin (FSK) and agonist (DAMGO for MOR and SalA for KOR) diluted in room temperature CO_2_-Independent media (10µM FSK and EC_80_ agonist final concentrations). Immediately following the FSK addition, luminescence measurements were monitored for 12min. Luminescence fold change from baseline for each well was calculated for the 10min luminescence measurement. Data was normalized to the naloxone control on each plate and analyzed using GraphPad Prism v.10.

### X-ray Structure for Ligand Stereochemistry

Low-temperature diffraction data (ω-scans) were collected on a Rigaku MicroMax-007HF diffractometer coupled to a Dectris Pilatus3R detector with Mo Kα (λ = 0.71073 Å) for the structure of 007c-24016. The diffraction images were processed and scaled using Rigaku Oxford Diffraction software (CrysAlisPro; Rigaku OD: The Woodlands, TX, 2015). The structure was solved with SHELXT and was refined against F2 on all data by full-matrix least squares with SHELXL (Sheldrick, G. M. Acta Cryst. 2008, A64, 112–122). A solvent mask was calculated, and 40 electrons were found in a volume of 130 Å^3^ in 1 void per unit cell. This is consistent with the presence of 1[C6H14] per formula unit which account for 100 electrons per unit cell. All non-hydrogen atoms were refined anisotropically. Hydrogen atoms were included in the model at geometrically calculated positions and refined using a riding model. The isotropic displacement parameters of all hydrogen atoms were fixed to 1.2 times the U value of the atoms to which they are linked (1.5 times for methyl and alcohol groups). The full numbering scheme of compound 007c-24016 can be found in the full details of the X-ray structure determination (CIF), which is included as Supporting Information (see SI Section 6). CCDC number XXXXXX (007c-24016) contains the supplementary crystallographic data for this paper. These data can be obtained free of charge from The Cambridge Crystallographic Data Center via www.ccdc.cam.ac.uk/data_request/cif

### Expression and Purification of MOR and KOR complexes

Gene constructs for human MOR and KOR were codon-optimized and synthesized (Twist Biosciences) into pCDNA3.1-zeo-tetO vectors with an N-terminal signal FLAG tag. The MOR construct was mutated at two positions at the third intracellular loop (M266K and L271R) by site-directed mutagenesis to enable binding to the KOR-specific nanobody Nb6M for structure determination. Expi293F Inducible cells (ThermoFisher Scientific), which stably express the tetracycline repressor gene and maintained in Expi293 Expression Medium (ThermoFisher Scientific) supplemented with 50 µg/mL of Blasticidin (Invivogen), were transfected with these constructs at 1 µg/mL using the ExpiFectamine 293 Transfection kit (ThermoFisher Scientific) according to manufacturer’s instructions at a cell density of 3 x 10^6^ cells/mL without Blasticidin. Transfected cells were supplemented 18-24 post-transfection with enhancer 1 and enhancer 2 from the ExpiFectamine 293 Transfection kit, 1 µM of Naloxone Hydrochloride, and in the case of MOR-transfected cells, 2 µg/mL of Doxycycline Hyclate. Cells were harvested 48-72 hours post-transfection by centrifugation at 4000xg for 10-20 minutes and freezing the cell pellet at -80°C.

Nb6M, NabFab, and anti-Fab Nb were expressed and purified as previously described, snap-frozen in liquid nitrogen, and stored at -80°C until complexing with the receptors^52, 74^.

For MOR complexed with compound **#33**, cell pellets from a 400 mL culture were thawed on the day of purification in a room-temperature water bath and lysed for 10 minutes at 4°C with 20 mM HEPES pH 7.5, 1 mM EDTA, 1 protease inhibitor cocktail tablet (ThermoFisher Scientific), and 10 µM of compound **#33**. Cells were spun down at 16,000 rpm for 15 minutes at 4°C (JA-18 rotor, Beckman Coulter) to pellet the cell membranes. Cell membranes were dounced in a tight glass homogenizer until no pellets were visible, then subsequently solubilized for 1 hour at 4°C with 20 mM HEPES pH 7.5, 300 mM NaCl, 1% dodecyl maltoside (DDM, Anatrace), 0.1% cholesterol hemissucinate (CHS, Anatrace), 0.3% 3-[(3-cholamidopropyl)dimethylammonio]-1-propanesulfonate (CHAPS), 2 mM MgCl_2_, 5 mM ATP, 100 µM tris (2-carboxyethyl) phosphine (TCEP), 1 protease inhibitor tablet, and 10 µM compound **#33**. Solubilized membranes were spun down at 16,000 rpm for 20 minutes at 4°C (JA-18 rotor) to clarify the solution. The clarified membrane solution was supplemented with CaCl_2_ to a final concentration of 2 mM then mixed with homemade M1-FLAG resin pre-equilibrated with 2 column volumes (CVs) ATP wash buffer consisting of 20 mM HEPES pH 7.5, 300mM NaCl, 0.1% DDM, 0.01% CHS, 0.03% CHAPS, 2mM CaCl_2_, 2mM MgCl_2_, 5mM adenosine triphosphate (ATP), and 10 µM compound **#33**. The receptor + resin solution was gently rotated for at least 1 hour at 4°C. The resin was loaded onto a Kimble® Flex-Column® (2 mL loading capacity, DWK Life Sciences) and washed with 10 CVs of ATP wash buffer and 10 CVs of low-DDM buffer consisting of 20 mM HEPES pH 7.5, 150 mM NaCl, 0.1% DDM, 0.01% CHS, 0.03% CHAPS, 2 mM CaCl_2_, and 10 µM compound **#33**. DDM was then gradually exchanged for glyco-diosgenin (GDN, Anatrace) with 5 CVs of GDN exchange buffer 1 (1:1 ratio of low-DDM buffer and base-GDN exchange buffer, consisting of 20 mM HEPES pH 7.5, 150 mM NaCl, 0.4% GDN, 0.04% CHS, 2mM CaCl_2_, and 10µM compound **#33**), 5 CVs of GDN exchange buffer 2 (1:3 ratio of low-DDM buffer and GDN exchange buffer), 5 CVs of GDN exchange buffer 3 (1:7 ratio of low-DDM buffer and GDN exchange buffer), and 1 CV of base-GDN exchange buffer. The resin was then washed with 10 CVs of low-GDN wash buffer (20mM HEPES pH 7.5, 150mM NaCl, 0.04% GDN, 0.004% CHS, 2mM CaCl_2_, and 10µM compound **#33**) and eluted with 3 CVs of elution buffer (20mM HEPES pH 7.5, 150mM NaCl, 0.01% GDN, 0.001% CHS, 5mM EDTA, 10µM compound **#33**, 0.2 mg/mL FLAG peptide). Elution fractions were then pooled, concentrated with a 50 kDa cutoff Amicon® concentrator (Sigma Aldrich), and purified by size exclusion chromatography (SEC, ӒKTA pure™, Cytiva) in a Superdex 200 Increase 10/300 GL column (GE Healthcare) equilibrated with 20 mM HEPES pH 7.5, 150mM NaCl, 0.02% GDN, 0.002% CHS, and 10 µM compound **#33**. Purified receptors were then pooled, concentrated, quantified, and then complexed overnight with 30 µM compound **#33** and 1.5x molar excess of purified Nb6M, NabFab, and anti-Fab Nb. The purified receptor complex then underwent a final SEC purification in a Superdex 200 Increase 10/300 GL column equilibrated with 20 mM HEPES pH 7.5, 150mM NaCl, 0.01% GDN, 0.001% CHS, and 10 µM compound **#33**.

For MOR and KOR complexed with compound **#020_E1**, cell pellets from 100 mL and 200 mL culture respectively were thawed on the day of purification in a room-temperature water bath and lysed for 10 minutes at 4°C with 20 mM HEPES pH 7.5, 5 mM MgCl_2_, 100 µM TCEP, 1 protease inhibitor cocktail tablet, 2 µL of Benzonase® Nuclease (Sigma Aldrich) and 10 µM of compound **#020_E1**. Cells were spun down at 14,000 rpm for 15 minutes at 4°C (JA 25.50 rotor, Beckman Coulter) to pellet the cell membranes. Cell membranes were dounced with 10 strokes in a tight glass homogenizer, then subsequently solubilized for 1 hour at 4°C with 1% lauryl maltose neopentyl glycol (LMNG, Anatrace), 0.1% CHS, 250 mM NaCl, 50 mM HEPES pH 7.5, 1 mM MgCl_2_, 1 protease inhibitor cocktail tablet, 10 µM of compound **#020_E1**, 100 µM TCEP, and 2 µL of Benzonase® Nuclease. Solubilized membranes were spun down at 14,000 rpm for 30 minutes at 4°C (JA 25.50 rotor) to clarify the solution. The clarified membrane solution was supplemented with CaCl_2_ to a final concentration of 5 mM then mixed with homemade M1-FLAG resin pre-equilibrated with 2 CVs FLAG wash buffer consisting of 250 mM NaCl, 20 mM HEPES pH 7.5, 10 µM compound **#020_E1**, 2 mM CaCl_2_, 0.1% LMNG, and 0.01% CHS. The receptor + resin solution was gently rotated for at least 1 hour at 4°C, then centrifuged at 300 rpm (Sorvall Legend XTR, ThermoFisher Scientific) for 3 minutes at 4°C to gently recover the resin. The resin was loaded onto a Kimble® Flex-Column® and washed with 20 CVs of FLAG wash buffer. Protein was eluted with 3-5 CVs of FLAG elution buffer consisting of 150 mM NaCl, 20 mM HEPES pH 7.5, 10 µM compound **#020_E1**, 1 mM EDTA, 0.2 mg/mL FLAG peptide, 0.1% LMNG, and 0.01% CHS. Purified receptor was concentrated with a 50 kDa cutoff Amicon® concentrator. Purified Nb6M, NabFab, and anti-Fab Nb was added to the receptor at a 2:2:2:1 molar ratio and complexed for 1 hour at 4°C. Receptor complex was further purified SEC in a Superdex 200 Increase 10/300 GL column equilibrated with 150 mM NaCl, 20 mM HEPES pH 7.5, 30 µM compound **#020_E1**, 0.001% LMNG, 0.0001% CHS, and 0.00033% Glyco-Diosgenin.

### CryoEM sample preparation and data collection

Fractions from monodisperse peaks in the SEC profile containing purified complexes of MOR with **#33** were collected and concentrated to 3-6 mg/mL with a 50 kDa cutoff Amicon® concentrator and used immediately for cryo-EM sample preparation. 300 mesh UltrAuFoil R 1.2/1.3 gold grids (Quantifoil) were glow-discharged in an EMS 700 Glow Discharge system (Electron Microscopy Sciences). 3 µL of sample was applied to the glow-discharged grid in a Vitrobot Mark IV vitrification system (ThermoFisher Scientific) cooled to 4°C with 100% relative humidity. After a 10 second wait time, grids were blotted for 1.5 seconds with Whatman® No. 1 filter paper (Sigma-Aldrich) and then plunge-frozen in liquid ethane. Grids were clipped into Autogrids (ThermoFisher Scientific) and stored in liquid nitrogen until cryoEM data collection.

Fractions from monodisperse peaks in the SEC profile containing purified complexes of MOR and KOR with **#020_E1** were collected and concentrated to 3 mg/mL and 8 mg/mL respectively with a 50 kDa cutoff Amicon® concentrator and used immediately for cryo-EM sample preparation. 300 mesh UltrAuFoil R 1.2/1.3 gold grids were glow-discharged in an EMS 700 Glow Discharge system. 3 µL of sample was applied to the glow-discharged grid in a Vitrobot Mark IV vitrification system (ThermoFisher Scientific) cooled to 4°C with 100% relative humidity. After a 10 second wait time, grids were blotted for 1.5 to 3.0 seconds with Whatman® No. 1 filter paper (Sigma-Aldrich) and then plunge-frozen in liquid ethane. Grids were clipped into Autogrids (ThermoFisher Scientific) and stored in liquid nitrogen until cryoEM data collection.

Clipped MOR + **#33** grids were loaded into a Titan Krios G3 microscope (ThermoFisher Scientific) set at 300 kV equipped with a BioQuantum energy filter set at 20 eV slit width and a K3 direct electron detector camera. After atlasing the grids and screening, single-particle cryoEM data was collected in dose fractionation mode on SerialEM by multi-shot data acquisition with fringe-free imaging (FFI). X-frame movies were collected at a defocus range of -0.9 to -2.0 µm at a pixel size of 0.835 Å (nominal magnification of 105,000x) in counting mode at an exposure rate of 16 e^-^/pix/s for a total exposure time of 2.0 seconds and a total electron exposure of 45.8 e^-^/Å^2^. In total, 7,766 movies were collected over two sessions, with on-the-fly motion correction and alignment being done through MotionCor2 in Scipion.

Clipped KOR + **#020_E1** grids were loaded into a Talos Arctica microscope (ThermoFisher Scientific) set at 200 kV equipped with a BioQuantum energy filter (Gatan Inc.) set at 20 eV slit width and a K3 direct electron detector camera (Gatan Inc.). After atlasing the grids and screening, single-particle cryoEM data was collected in dose fractionation mode on SerialEM by a 3x3 multi-shot data acquisition with fringe-free imaging (FFI) at 2 shots per hole. 75-frame movies were collected at a defocus range of -0.8 to -2.1 µm at a pixel size of 0.865Å (nominal magnification of 45,000x) in counting mode at an exposure rate of 16 e^-^/pix/s for a total exposure time of 3.0 seconds and a total electron exposure of 64.1 e^-^/Å^2^. In total, 4,967 movies were collected, with on-the-fly motion correction and alignment being done through MotionCor2 in Scipion.

Clipped MOR + **#020_E1** grids were loaded into a Titan Krios G3 microscope set at 300 kV equipped with a BioQuantum energy filter set at 20 eV slit width and a K3 direct electron detector camera. After atlasing the grids and screening, single-particle cryoEM data was collected in dose fractionation mode on SerialEM by a 5x5 multi-shot data acquisition with fringe-free imaging (FFI) at 3 shots per hole. 80-frame movies were collected at a defocus range of -0.8 to -2.1 µm at a pixel size of 0.8189 Å (nominal magnification of 105,000x) in counting mode at an exposure rate of 16 e^-^/pix/s for a total exposure time of 2.0 seconds and a total electron exposure of 47.7 e^-^/Å^2^. In total, 9,877 movies were collected, with on-the-fly motion correction and alignment being done through MotionCor2 in Scipion.

### CryoEM data processing

All 7,764 motion-corrected micrographs from the MOR + **#33** dataset collected over two sessions were imported into cryoSPARC v4.0.3 (Structura Biotechnology Inc.) and estimated for its contrast transfer function (CTF) by patch CTF estimation. Micrographs were curated based on a CTF fit resolution range of 2.5 -10 Å, leaving a total of 6,674 micrographs for further processing. From a subset of 2,409 micrographs, blob picker was initially used to pick 1,923,561 particles at a minimum/maximum diameter of 80-180 Å. 1,442,738 particles were extracted at a box size of 360 pixels, binned to 96 pixels. A series of 2D classification and particle re-extraction were undertaken with gradual box unbinning until the Fabs and nanobodies were visible across several classes at several orientations. 72,817 particles from classes with visible Fabs and nanobodies were then extracted at a box size of 400 pixels and used to create an ab-initio 3D model from a single class. This model then underwent non uniform-refinement and then used to produce 2D templates for template picking and to produce a model for heterogeneous refinement. Template picking was performed at a particle diameter of 160 Å to pick 7,498,887 particles from the original CTF-curated micrographs. 6,519,032 particles were extracted at a box size of 360 pixels, binned by 4 to 90 pixels. Particles underwent one round of 2D classification before going through several rounds of heterogeneous refinement, homogeneous refinement, and ab-initio reconstruction. Particles belonging to good 3D classes were iteratively extracted with gradual box unbinning. The best class containing 334,011 particles underwent non-uniform refinement and then further refined by local refinement using a TM-specific mask. 3D classification was then used with a class similarity of 0.5 and hard classification at a 3.5 Å target resolution across 6 classes to separate out finer features without pose realignment. The best class, containing a total of 40,722 particles, was selected for non-uniform refinement to 3.98 Å and local refinement and sharpening using a TM-specific mask to a final resolution of 3.9 Å. The final MOR EM map was seen as a monomer with the majority of the nanobodies and Fab fragment clearly visible.

All 4,967 motion-corrected micrographs from the KOR + **#020_E1** dataset were imported into cryoSPARC v4.0.3 and estimated for its contrast transfer function (CTF) by patch CTF estimation. Micrographs were curated based on a maximum CTF fit resolution of 4 Å, leaving a total of 2,678 micrographs for further processing. Blob picker was initially used to pick 930,768 particles at a minimum/maximum diameter of 150-300 Å. 808,182 particles were extracted at a box size of 256 pixels, binned by 4 to 64 pixels. A series of 2D classification and particle re-extraction were undertaken until transmembrane (TM) helices were well visible across several classes at several orientations. 165,141 particles from classes with clearly resolved TM helices were then extracted at a box size of 360 pixels and used to create an Ab-initio 3D model at a resolution range of 7-9 Å from 8 classes. The best two 3D models were selected to produce 2D templates for template picking and to serve as good classes for heterogeneous refinement. Template picking was done at a particle diameter of 180 Å to pick 2,056,565 particles from the original CTF-curated micrographs. 1,797,537 particles were extracted at a box size of 360 pixels and classified by heterogeneous refinement with the two good classes and two junk classes. The best class containing 835,214 particles was selected for non-uniform refinement, and then further refined by local refinement using a TM-specific mask. 3D classification was then used with a class similarity of 0.1 and hard classification at a 3.3 Å target resolution across 8 classes to separate out finer features without pose realignment. The best class, containing 98,518 particles, was selected for non-uniform refinement to 3.18 Å and local refinement and sharpening using a TM-specific mask to a final resolution of 2.96 Å. The final KOR EM map was seen as an antiparallel heterodimer with the Fab fragment and part of the nanobody clearly visible on one of the monomeric subunits; efforts to isolate the monomeric form of KOR were unsuccessful.

All 9,877 motion-corrected micrographs from the MOR + **#020_E1** dataset were imported into cryoSPARC v4.0.3 and estimated for its contrast transfer function (CTF) by patch CTF estimation. Micrographs were curated based on a maximum CTF fit resolution of 4 Å, leaving a total of 8,433 micrographs for further processing. Blob picker was initially used to pick 6,878,984 particles at a minimum/maximum diameter of 90-360 Å. 5,814,596 particles were extracted at a box size of 360 pixels, binned by 4 to 90 pixels. A series of 2D classification and particle re-extraction were undertaken with gradual box unbinning until transmembrane (TM) helices were well visible across several classes at several orientations. 162,842 particles from classes with clearly resolved TM helices were then extracted at a box size of 360 pixels and used to create an ab-initio 3D model at a resolution range of 7-9 Å from 8 classes. The best 3D model was selected to produce 2D templates for template picking and to produce a model for heterogeneous refinement. Template picking was performed at a particle diameter of 180 Å to pick 5,949,666 particles from the original CTF-curated micrographs. 5,259,530 particles were extracted at a box size of 360 pixels binned by 4 to 90 pixels. Several rounds of heterogeneous refinement through a combination of good classes and junk classes were performed with particles being extracted from the good 3D classes with gradual box unbinning. The best class containing 322,546 particles was selected for non-uniform refinement, and then further refined by local refinement using a TM-specific mask. 3D classification was then used with a class similarity of 0.1 and hard classification at a 3.0 Å target resolution across 3 classes to separate out finer features without pose realignment. The best two classes, containing a total of 259,816 particles, was selected for non-uniform refinement to 3.30 Å and local refinement and sharpening using a TM-specific mask to a final resolution of 3.23 Å. The final MOR EM map was seen as a monomer with the majority of the nanobodies and Fab fragment clearly visible.

### CryoEM model building and refinement

The locally-refined, sharpened map for MOR + **#33** was modeled by initially docking a cryoEM inactive structure of mouse MOR bound to alvimopan (PDB 7UL4) in UCSF ChimeraX. The alvimopan was removed, and residues at the N-terminus and the C-terminus with poor fit into the density map were truncated, and additionally certain residues for MOR were mutated to match the human variant and to account for the mutations necessary to bind Nb6M. The model was then subject to a rough refinement using ISOLDE and subjected to real space refinements through Phenix and visually inspected in COOT. Geometrical validations and model-to-map FSC were performed by MolProbity. The structure was deposited into the PDB with PDB ID 9MQH. Due to the lower resolution of the final map, we could not unambiguously model in compound **#33** into the ligand density. As an alternative, compound **#33** was docked into the MOR structure by Maestro (Schrödinger LLC.).

The locally-refined, sharpened map for KOR + **#020_E1** was modeled by initially docking an AlphaFold-derived model (AF-P41145-F1) while the locally-refined, sharpened map for MOR + **#020_E1** was modeled by initially docking a cryoEM inactive structure of mouse MOR bound to alvimopan (PDB 7UL4) in UCSF ChimeraX. The alvimopan for the MOR structure was removed, and residues at the N-terminus and the C-terminus with poor fit into the density map were truncated. Certain residues for both MOR and KOR were mutated to match the human variant and to account for the mutations necessary to bind Nb6M. The model was then subject to a rough refinement using ISOLDE. Coordinates and restraints for the ligand **#020_E1** were generated by eLBOW through Phenix and then manually fit into the putative ligand density by COOT. Four different stereochemical geometries **#020_E1** were tested, and the best one was selected based on its fit to the EM map density, known past interactions between the receptor and its ligands, and minimization of steric clashes. The model was then subjected to iterative rounds of real space refinements through Phenix and manual refinements in COOT, with geometrical validations and model-to-map FSC being performed by MolProbity. 3D anisotropy was analyzed by PyEM and 3DFSC, and all models were visualized using ChimeraX. While separate models for the non-locally refined density maps resolving the nanobodies and Fabs were created with models of Nb6M (PDB 7UL3) and NabFab (PDB 7PIJ), all high-detailed structural analysis was performed on the models that were derived from the sharpened, locally refined map for both receptors to ensure the highest accuracy possible for the ligand. The full structures (MOR PDB ID: 9MQI; KOR PDB ID: 9MQK) and locally refined structures (MOR PDB ID: 9MQJ; KOR PDB ID: 9MQL) were deposited into the PDB.

### *In Vivo* Behavioral Studies

Animal experiments were approved by the UCSF Institutional Animal Care and Use Committee and were conducted in accordance with the NIH Guide for the Care and Use of Laboratory animals (protocol #AN195657). Adult (8-10 weeks old) male C56BL/6 mice (strain # 664) were purchased from the Jackson Laboratory. Mice were housed in cages on a standard 12:12 hour light/dark cycle with food and water ad libitum. For all behavioral tests, the experimenter was always blind to treatment. To measure thermal sensitivity, 30 minutes after the compound’s injection, we recorded the latency for the tail’s mouse to flick when immersed into a 50oC water bath.

### Conditioned Place Aversion

To determine if **#020_E1** was inherently aversive we modified a previously described conditioned place paradigm (Juarez-Salinas et al., 2018). Briefly, mice were habituated to the test apparatus, on two consecutive days, and their preference for each chamber recorded for 30 minutes (Pretest). Mice were then made tolerant to increasing doses of morphine over four days (see below). On the 5th days, tolerant mice were injected with either naloxone or **#020_E1** and immediately placed in their preferred chamber for 30 min (conditioning day). On the sixth day (test day), mice were placed back in the apparatus where they were allowed to roam freely between the three chambers and we recorded the time spent in each chamber for 30 minutes. To calculate the CPA score, we subtracted the time spent in each chamber of the box on the pretest day from that of the test day (CPA score = Test - Pretest).

### Opioid Withdrawal Symptoms Assay

To investigate the ability of the novel opioid antagonists to precipitate opioid withdrawal symptoms, we first generated mice tolerant to morphine, as previously described (Wilson et al., 2021; Singh et al., 2023). Briefly, mice received 8 escalating doses of morphine (IP) over 4 days (10, 15, 20, 30, 50, 60, 70 and 75 mg/kg; twice daily, i.p.). On the 5th day, mice received a single dose of morphine (20 mg/kg, i.p.), followed 1h later by a single dose of naloxone or the novel antagonists and were immediately video recorded for the next 20 minutes. To document withdrawal, we scored the number of naloxone-precipitated jumps, forepaw shakes, wet dog shakes and rearing over the next 20 minutes, as well as the latency for the first jump.

## Supporting information

SI section

NMR spectra

## ASSOCIATED CONTENT

### Supporting information

Supporting Information is available free of charge on the ACS Publication website.

- Supplemental tables and figures describing reaction codes and SMARTS patterns used for building block filtering and reaction enumeration, and biochemistry of purified MOR and KOR used for cryoEM
- General synthetic procedures
- Complete Synthetic methods for initial screening hits (**#03-#64**) and analogs (**#001-#036**)
- X-ray crystallography of compound **#020_E1**
- Pharmacokinetic Report of compound **#020_E1**
- CryoEM Validation Statistics
- NMR spectra of all synthesized compounds

### Author information

#### Corresponding Authors

**John J. Irwin** – Department of Pharmaceutical Chemistry, University of California, San Francisco, 94158, United States.

**Aashish Manglik** - Department of Pharmaceutical Chemistry, University of California, San Francisco, 94158, United States.

**Allan I. Basbaum** - Department of Anatomy, University of California, San Francisco, 94158, United States.

**Jonathan A. Ellman** - Department of Chemistry, Yale University, New Haven, Connecticut 06520, United States.

**Brian K. Shoichet** - Department of Pharmaceutical Chemistry, University of California, San Francisco, 94158, United States.

#### Authors

**Seth F. Vigneron** - Department of Pharmaceutical Chemistry, University of California, San Francisco, 94158, United States.

**Shohei Ohno** - Department of Chemistry, Yale University, New Haven, Connecticut 06520, United States.

**Joao Braz** - Department of Anatomy, University of California, San Francisco, 94158, United States.

**Joseph Y. Kim** - Department of Pharmaceutical Chemistry, University of California, San Francisco, 94158, United States.

**Oh Sang Kweon** - Department of Chemistry, Yale University, New Haven, Connecticut 06520, United States.

**Chase Webb** - Department of Pharmaceutical Chemistry, University of California, San Francisco, 94158, United States.

**Christian Billesbølle** - Department of Pharmaceutical Chemistry, University of California, San Francisco, 94158, United States.

**Karnika Bhardwaj** - Department of Anatomy, University of California, San Francisco, 94158, United States.

#### Author Contributions

S.V., guided by J.J.I., constructed the library of 14.6 isoquinuclidines as advised for synthetic accessibility by J.A.E. and J. Kweon. S.V. docked them to structures of the MOR and KOR, using energy potential grids initially generated by C.W.. Working with B.K.S., S.V. prioritized high ranking isoquinuclidines for synthesis by S.O. and J. Kweon, and tested them experimentally in both ligand-displacement and functional assays. J.Y.K. and S.V. also biochemically purified the MOR protein that was used for structure determination by J.Y.K., advised by A.M. and C.B.. J.Y.K. also purified the KOR protein and determined its structure, advised by A.M. and C.B.. J.B. performed all *in vivo* mouse experiments with input from A.B.. J.A.E. & B.K.S. conceived of and supervised the project and reviewed data. S.V., B.K.S. & J.A.E. wrote the paper with input from the other authors.

#### Data availability

The cryoEM structures of #020_E1 in complex with MOR and KOR have been deposited in the PDB. MOR-#020_E1 PDB IDs: full structure 9MQI, locally refined structure 9MQJ. KOR-#020_E1 PDB IDs: full structure 9MQK, locally refined structure 9MQL. The cryoEM structure of #33 in complex with MOR has also been deposited into the PDB with ID 9MQH.

#### Notes

The authors declare the following competing financial interest(s): A patent has been submitted for the chemical space surrounding the presented isoquinuclidines. BKS is co-founder of BlueDolphin LLC, Epiodyne Inc, and Deep Apple Therapeutics, Inc., and serves on the SRB of Genentech, the SAB of Schrodinger LLC, and the SAB of Vilya Therapeutics. A.M. is a co-founder of Epiodyne Inc and Stipple Bio and serves on the SAB for Alkermes, Septerna, and Abalone. No other authors declare competing interests.

## Abbreviation Used

MOR: Mu Opioid Receptor (µ-Opioid Receptor)
KOR: Kappa Opioid Receptor (κ-Opioid Receptor)
GPCR: G protein coupled receptor
HAC: Heavy Atom Count
CryoEM: Cryo-electron microscopy
IFP: Interaction Fingerprint
Tc: Tanimoto Coefficient
Fab: Fragment Antigen-Binding region
NabFab: Nanobody-targeting Fab fragment
Anti-Fab Nb: Fab-targeting Nanobody
MP: mitragynine pseudoindoxyl
i.p.: Intraperitoneal

