## Supplementary material for "Docking 14 million virtual isoquinuclidines against the mu and kappa opioid receptors reveals dual antagonists-inverse agonists with reduced withdrawal effects": SI section

### 1. Supplementary Tables and Figures

| Reaction Class | Reaction Codes | Reaction Class | Reaction Codes |
|---|---|---|---|
| Amide Coupling | 11, 22, 58, 527, 1606, 1626, 2740, 188690, 240690, 269946, 270006, 270062, 270112, 271144, 272270, 272430, 272610, 273012, 273290, 273450, 273652, 273692, 274079, 274614, 274860, 276010, 276750, 280130, 280190 | Acrylamide Furnishing | 273270 |
| Multi-Amide Coupling | 274552, 275592 | Purine Formation | 275054 |
| Urea Coupling | 68, 487, 512, 2430, 2554, 2708, 272164, 273390, 273392, 273452, 273454, 273458, 273464, 273574, 274080 | Pyrrole Formation | 274092 |
| Reductive Amination | 207, 269862, 270004, 270302, 271302, 271304, 272690, 273456, 274370, 275532, 275550, 279530 | Imine Formation | 4 |
| Esterification | 1458, 276436, 279392 | Condensation Product | 6, 28, 46 |
| Carbamate Coupling | 274081, 282070 | Oxazolidinone Formation | 10 |
| Amine Alkylation | 27, 38, 44, 61, 88, 2230, 269956, 269982, 270034, 270122, 270166, 275550, 280170, 280178, 280530, 281630, 282490 | Biginelli Product | 12, 276390, 276950 |
| Multi-Amine Alkylation | 282030 | Enamine Formation | 17, 570 |
| Sulfonamide Coupling | 20, 40, 1606, 196680, 232682, 270084, 270188, 271082, 273578, 274078, 278578, 280530, 282270 | Mannich Product | 26 |
| Multi-Sulfonamid Coupling | 271242 | Thiodiazine Formation | 41 |
| Sulfonic Esterification | 87 | Quinoline Formation | 42 |
| Oxamide Coupling | 2718, 271948, 271949, 273460, 273462, 273492, 273494, 273496, 273498 | Cyclic Hydrazine Formation | 49, 52, 279312 |
| Suzuki Coupling | 271570, 273584, 273712, 275652 | Benzofurane Formation | 55 |
| Thiazole Formation | 1, 2, 3, 24, 43, 50, 272150, 274972, 276096 | Benzoimidazole Formation | 60 |
| Oxadiazole Formation | 265764, 270036, 270196, 270690, 271722, 271362, 274072 | Sulfide Product | 62 |
| Oxadiazoline Formation | 274990 | Epoxide Opening or Michael Adduct | 51, 63, 273610, 273790, 277110 |
| 1,2,4-Triazole Formation | 270942, 274052, 276090, 276630, 276670, 276910, 279370 | Tetrahydroquinazolone Formation | 66 |
| Click 1,2,3-Triazole Formation | 273910, 274090, 274860, 274952, 274972, 275090, 275530, 275534, 275536, 276010, 280190, 282512 | Thiourea Formation | 8 |
| Nuc-Halide SN2 | 7, 34, 270344, 272692, 272710, 273580, 273654, 280172 | Cyclic Thiourea Formation | 248 |
| Piperazindione Formation | 276530 | Pyrimidine-4-one | 708 |
| Tetrazole Formation | 275630, 277030, 280130, 280172, 280178, 281630, 282490, 280170 | Cyclic Imide Formation | 1070 |
| Pyrrolidinone Formation | 279390 | Sulfoxide Product | 1498, 265282 |
| Dihydrouracyl Formation | 276550, 277130 | Sulfone Product | 1500, 2714 |
| Aryldiazepinone Formation | 276770 | Aminoquinazoline Formation | 1982 |
| Diazepindione_Formation | 279770 | Guanidine Product | 58668, 264724, 264822, 275310, 276492 |
| Bargellini Reaction | 276110, 277030 | Activated Methylene Product | 194680 |
| Hydrotoin Formation | 272104, 272126, 272212 | Uracyl Formation | 270192 |
| Hydantoinic Acid Formation | 272230, 272310, 272330 | Thiazolidinone-thione Formation | 571 |

**Supplementary Table 1.** Full list of reaction codes used to synthesize the Enamine REAL make-on-demand library categorized into their respective reaction classes.

| Synthon Class | Inclusion SMARTS | Exclusion SMARTS |
|---|---|---|
| **Applied to all Synthons** | n/a | [SH1] [CX4]S S~S S[F,Cl,Br,I] C[Br,I] [CX4]Cl N[F,Cl,Br,I] [B] [*]=C=[*] N=[CX3,cX3] [Si] N#[CX1] [N,n]~[O,o,s] [H]C([H])=C([H])[CX4,c] [H]C([H])=C([CX4,c,Cl])[CX4,c] [Cl]C([H])=C([H])[CX4,c,Cl] [CX4]([O,N,S,n,+n])[O,N,n,S] [OH,NH2]cc[OH,NH2] [OH,NH2]caac[OH,NH2] NC#N [C]OS(=O)=O [N,n][N,n][C,c]=O [O,N]C=C P[H] [P+] [C-] [c-] N1=NC1 O1CC1 [C,c](=S)[N,n] [F,Cl,Br,I]C=O C=CC=C C(=O)C(=O) C(=O)CC(=O) C(=O)C=CC(=O) C(=O)([CH])C=CC(=O) [NX3,OX2]CCC=O [NX3,OX2]CC=CC=O [H]C([CX4,c,O,N,n,S,s])=O [CX4,c]C(=O)[CX4,c] N=[+N]=[-N] [+n] C(=O)C(F)(F)[H] [CX4,c][NH1][CX4,c] |
| **Amines** | [C][N]([H])[H] [C]N([H])C(OC(C)(C)C)=O | [c,O,N,n,S][N]([H])[H] [CH0][N]([H])[H] C=CC[N]([H])[H] [CX4]N([H])[CX4] [H]N([H])CC(=O)[NX3&H1] [nX2,o,s][c,n]C[N]([H])[H] c(=O)[c,n]C[N]([H])[H] c(O)[c,n]C[N]([H])[H] O=CN([H])[H] [C,c](=O)[C,c]=[C,c] C#C |
| **Anilines** | [c][N]([H])[H] [c]N([H])C(OC(C)(C)C)=O | [C,O,N,n][N]([H])[H] [CX4]N([H])[CX4] [H]N([H])cc(=O) O=CN([H])[H] [H]N([H])c([n])[n,cH0&R1] [H]N([H])c([cH0,NH0])[cH0,NH0] [C,c](=O)[C,c]=[C,c] C#C [H]N(c1ncncn1)[H] [H]N(c1nnncc1)[H] [H]N(c1cnnnc1)[H] [H]N(c1nnccn1)[H] [H]N(c1nncnc1)[H] [H]N(c1cnncn1)[H] [nX3]aac(=[S,O,N]) [nX3]c(=[S,O,N])cN [nX3]c(=[S,O,N])cN [H]N([H])ca[nX3]c(=[S,O,N]) [CX4][NH1]c |
| **Enone** | [C]C(=O)C([C,a])=C([H])[H] [H]C(=O)C([C,a])=C([H])[H] [C]C(=O)/C=C([H])/[C,a,F] [H]C(=O)/C=C([H])/[C,a,F] | C(=O)\C=C([H])/[C,a] [C,c,O,N,n][N]([H])[H] [C;R](=O)/[C;R]=[C;R]([H])/[C,a] C=CC(C=C)=O C(=O)CC=C C#C [C]C(=O)/C([H])=C([H])/[C] [a]C(=O)C=C [CX4&H0]C(=O)C=C C(=O)C([CX4&H0,Cl,F,CX2])=C C(=O)C=C([CX4&H0,Cl,CX2]) C(=O)C=Cnn [C;r4](=O)C=C |
| **Internal Alkyne** | C#C | [C,c,O,N,n][N]([H])[H] [C,c](=O)[C,c]=[C,c] C#C[H] C#C[CX4&H0,O,N,S,F,Cl] C#CC(=O)[C,c] C#CC(=O)[H] C#C[CX4][OH,NX4&H0] C#C[CH2]OC=O C#CC=C C(=O)C#CC(=O) C#CC#N [NX3]CC#N C(=O)C#N C(=O)CC#N |

**Supplementary Table 2.** Inclusion and Exclusion SMARTS patterns used for each synthon type for reaction compatibility filtering of building blocks.

| Alkyne Pattern | Reaction SMARTS Used |
|---|---|
| [*:5]C#C[*:6] | [*:1][N]([H])[H].[*:2]C(=O)C([*:3])=C([H])[*:4].[*:5]C#C[*:6].[H]C([H])=C([H])[*:7]>>[*:1]N1[C@H]([*:6])[C@]2([*:5])C([*:4])=C([*:3])[C@]1([C@@H]([*:7])C2)[*:2]'<br>'[*:1][N]([H])[H].[*:2]C(=O)C([*:3])=C([H])[*:4].[*:5]C#C[*:6].[H]C([H])=C([H])[*:7]>>[*:1]N1[C@@]([*:2])([C@H]([*:7])C2)C([*:3])=C([*:4])[C@]2([*:5])[C@@H]1[*:6]' |
| [a:5]C#C[A:6] | [*:1][N]([H])[H].[*:2]C(=O)C([*:3])=C([H])[*:4].[*:5]C#C[*:6].[H]C([H])=C([H])[*:7]>>[*:1]N1[C@H]([*:5])[C@]2([*:6])C([*:4])=C([*:3])[C@]1([C@@H]([*:7])C2)[*:2]'<br>[*:1][N]([H])[H].[*:2]C(=O)C([*:3])=C([H])[*:4].[*:5]C#C[*:6].[H]C([H])=C([H])[*:7]>>[*:1]N1[C@@]([*:2])([C@H]([*:7])C2)C([*:3])=C([*:4])[C@]2([*:6])[C@@H]1[*:5]' |
| [*:5]C(C#C[A:6])=O | [*:1][N]([H])[H].[*:2]C(=O)C([*:3])=C([H])[*:4].[a:5]C#C[A:6].[H]C([H])=C([H])[*:7]>>[*:1]N1[C@H]([A:6])[C@]2([a:5])C([*:4])=C([*:3])[C@]1([C@@H]([*:7])C2)[*:2]'<br>'[*:1][N]([H])[H].[*:2]C(=O)C([*:3])=C([H])[*:4].[a:5]C#C[A:6].[H]C([H])=C([H])[*:7]>>[*:1]N1[C@@]([*:2])([C@H]([*:7])C2)C([*:3])=C([*:4])[C@]2([a:5])[C@@H]1[A:6]' |
| [CX3:6]=CC#C[CX4:5] | [*:1][N]([H])[H].[*:2]C(=O)C([*:3])=C([H])[*:4].[*:5]C(C#C[A:6])=O.[H]C([H])=C([H])[*:7]>>[*:1]N1[C@H]([*:6])[C@]2(C([*:5])=O)C([*:4])=C([*:3])[C@]1([C@@H]([*:7])C2)[*:2]'<br>'[*:1][N]([H])[H].[*:2]C(=O)C([*:3])=C([H])[*:4].[*:5]C(C#C[A:6])=O.[H]C([H])=C([H])[*:7]>>[*:1]N1[C@@]([*:2])([C@H]([*:7])C2)C([*:3])=C([*:4])[C@]2(C([*:5])=O)[C@@H]1[A:6]' |
| [*:5]C#C[*:6] | [*:1][N]([H])[H].[*:2]C(=O)C([*:3])=C([H])[*:4].[CX3:6]=C([H])C#C[CX4:5].[H]C([H])=C([H])[*:7]>>[*:1]N1[C@H](C=[CX3:6])[C@]2([CX4:5])C([*:4])=C([*:3])[C@]1([C@@H]([*:7])C2)[*:2]'<br>'[*:1][N]([H])[H].[*:2]C(=O)C([*:3])=C([H])[*:4].[CX3:6]=C([H])C#C[CX4:5].[H]C([H])=C([H])[*:7]>>[*:1]N1[C@@]([*:2])([C@H]([*:7])C2)C([*:3])=C([*:4])[C@]2([CX4:5])[C@@H]1C=[CX3:6]' |

**Supplementary Table 3. Reaction SMARTS used to combine building blocks into final isoquinuclidines.** The substituents on the internal alkyne building block dictate it's regiochemistry in the final product. For every combination of 1 amine, enone, alkyne and alkene, both the plus and minus enantiomers were generated with the regiochemistry dictated by the alkyne.

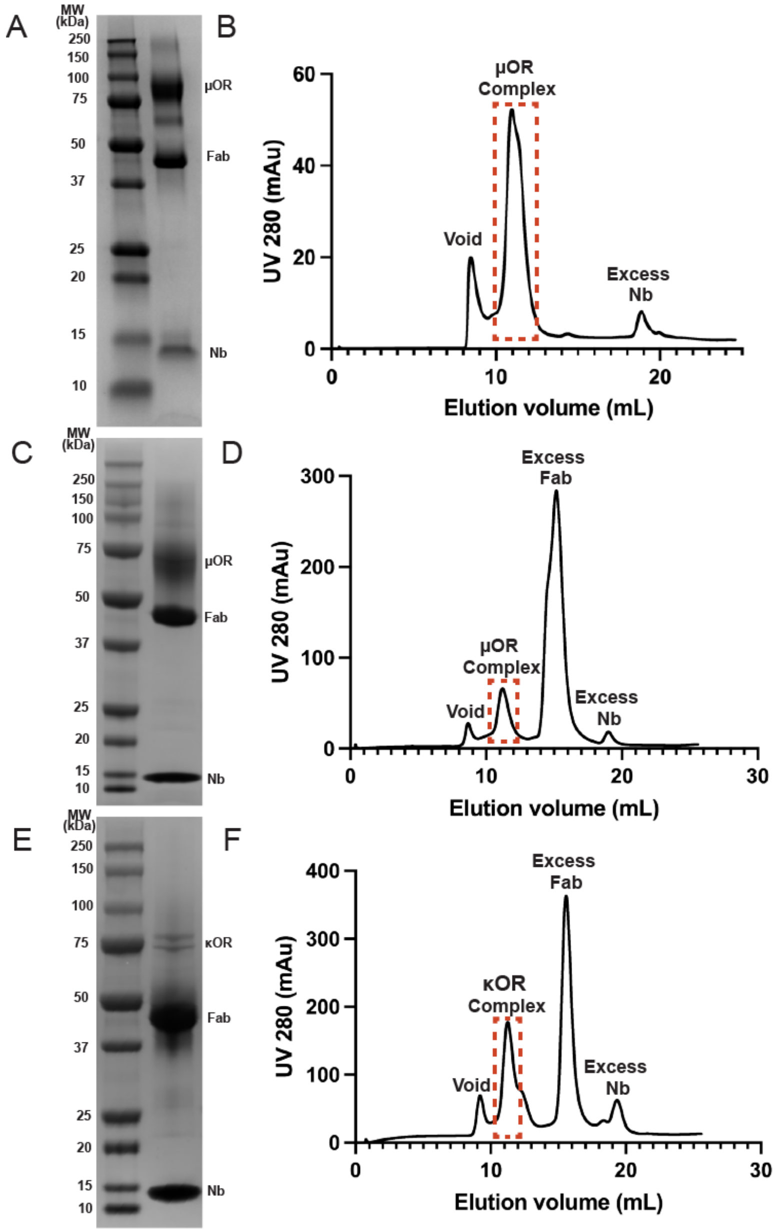

**Supplemental Figure 1: Biochemistry of purified MOR and KOR used for cryoEM.** **A.** Cropped SDS-PAGE of MOR complexed with compound **#33**, Nb6M, NabFab, and Anti-Fab Nanobody. The MOR was mutated at the ICL3 to enable binding to Nb6M. **B.** Final size-exclusion chromatogram (SEC) profile of the MOR complex. The fractionated peaks that corresponds to the MOR complex (dotted red box) were immediately collected and concentrated prior to cryoEM sample preparation. **C.** Cropped SDS-PAGE of MOR complexed with compound **#020_E1**, Nb6M, NabFab, and Anti-Fab Nanobody. **D.** Final size-exclusion chromatogram (SEC) profile of the MOR complex. The fractionated peaks that corresponds to the MOR complex (dotted red box) were immediately collected and concentrated prior to cryoEM sample preparation. **E.** Cropped SDS-PAGE of KOR complexed with compound **#020_E1**, Nb6M, NabFab, and Anti-Fab Nanobody. **F.** Final SEC profile of the KOR complex. The fractionated peaks that corresponds to the KOR complex (dotted red box) were immediately collected and concentrated prior to cryoEM sample preparation.

#### 2. SCHEME S1. General Synthesis Sequence

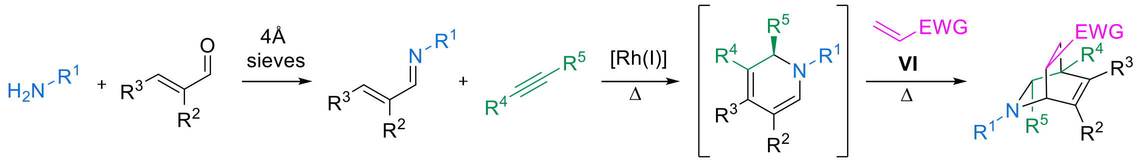

I II III IV ( )-V ( )-VII

#### 3. GENERAL PROCEDURES

##### Preparation of Rhodium Catalyst Stock Solution

$[RhCl(coe)_2]_2$ was purchased from Strem and stored inside a $N_2$-filled inert atmosphere glovebox at -25 ºC. Stock solutions of the rhodium catalyst were prepared in the glovebox. For the preparation of a 100 mM solution in THF, a 5 mL glass vial was charged with $[RhCl(coe)_2]_2$ (100 mg, 0.139 mmol) and p-$Me_2N$-$C_6H_4$-$PEt_2$ (58 mg, 0.278 mmol) in anhydrous THF (1.4 mL). Stock solutions, when stored in a –25 ºC freezer inside a $N_2$-filled glovebox, could be used for several weeks without loss of catalytic activity.

##### General Procedure A for the synthesis of imines III

To a flame-dried round-bottom flask was added a magnetic stir bar, amine **I** (1 equiv), enal **II** (1-5 equiv), dry THF (0.5 M), and 20 vol% of molecular sieves 4 Å. The reaction solution was stirred under $N_2$ at rt for 12 h. Upon reaction completion, the reaction mixture was filtered through a pad of celite. The filtrate was concentrated in vacuo to provide the desired imine **III**, which was stored at –20 °C in a nitrogen filled glovebox and taken on to the next step without further purification.

##### General Procedure B for the synthesis of DHPs V

Inside a $N_2$-filled inert atmosphere glovebox, imine **III** (1.0 equiv) was dissolved in anhydrous THF (0.5 M total concentration), and the solution was transferred to a flame-dried sealable glass vessel equipped with a magnetic stir bar. A stock solution of Rh catalyst (see General Information) at the catalyst loading indicated was added to the tube, followed by alkyne **IV** (2-5 equiv). A small portion (~0.4 mL) of this reaction mixture was transferred to an oven-dried J. Young NMR tube that was equipped with a flamesealed melting point capillary containing benzene-$d_6$ for reaction monitoring. The J. Young NMR tube and glass vessel were then sealed, removed from the glovebox, and heated at the indicated temperature. The reaction progress was monitored by $^1H$ NMR spectroscopy based on the disappearance of the imine **III** signals and the increasing resonances of the dihydropyridine **V**. Upon reaction completion, this crude DHP solution was used for the synthesis of isoquinuclidines **VII** (General Procedure C).

**General Procedure C for the synthesis of isoquinuclidines VII from DHPs V via the Diels–Alder reaction**

To the reaction vessel including the crude DHP **V** solution was added the desired alkene **VI** (10 equiv). The reaction vessel was then sealed, and the mixture was stirred at the indicated temperature. After reaction completion, the reaction vessel was opened, and the solvent was removed in vacuo. The resulting residue was purified by column chromatography on silica gel or preparative thin-layer chromatography to afford the desired isoquinuclidine **VII**.

**General Procedure D for the cleavage of acid-labile protecting groups from isoquinuclidines VII**

To an oven-dried round-bottom flask equipped with a magnetic stir bar was added the isoquinuclidine **VII** (1.0 equiv) in $CH_2Cl_2$ (0.1 M). The reaction mixture was cooled to 0 °C, and trifluoroacetic acid (10.0 equiv) was added. The ice bath was removed and the reaction mixture was stirred at room temperature until the reaction was complete as monitored by thin-layer chromatography. The reaction mixture was then diluted with $CH_2Cl_2$ and saturated $NaHCO_3$ (aq) and extracted three times with $CH_2Cl_2$. The combined organic layers were washed with brine, dried with $Na_2SO_4$, filtered and concentrated in vacuo. The crude material was purified by the indicated chromatographic method to afford the desired product.

**General Procedure E for the cleavage of *p*-methoxybenzyl group on nitrogen**

To a flame-dried round-bottom flask equipped with a magnetic stir bar was added isoquinuclidine **VII** (1.0 equiv) and dichloroethane (0.2 M). The reaction mixture was cooled to 0 °C, and 1-chloroethyl chloroformate (5 equiv) was added. The mixture warmed to room temperature and was stirred at rt until complete as monitored by thin-layer chromatography. The reaction mixture was then concentrated in vacuo, and the crude material was purified by silica gel chromatography with only pure fractions isolated to afford the carbamate intermediate. To a flame-dried round-bottom flask was added the purified intermediate followed by excess MeOH, and the reaction solution was stirred at 40 °C for 12 h. Upon reaction completion, the crude material was purified by the indicated chromatographic method to afford the desired product.

#### 4. Initial Virtual Screening Hits (#03-#64): Synthesis Procedures and Analytical Data

**Synthesis of #03**

・**Synthesis of Aldehyde**

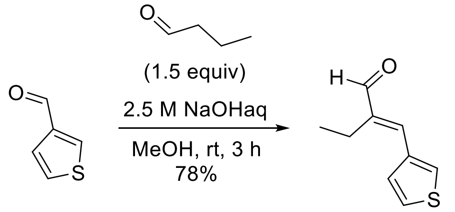

To a round-bottom flask was added a magnetic stir bar, thiophene-3-carbaldehyde (1 g, 8.92 mmol, 1 equiv) and butanal (1.21 mL, 13.4 mmol, 1.5 equiv) dissolved in MeOH (9 mL). To this solution was then added sodium hydroxide (2.5 M aq., 3.56 mL, 1.0 equiv) dropwise at 0 °C. The reaction mixture was stirred at room temperature for 3 h. After the reaction was complete as monitored by thin-layer chromatography, the mixture was then diluted with water, extracted with $Et_2O$, washed with brine, dried with $Na_2SO_4$ and filtered. The organic layer was concentrated and the crude residue was purified via silica gel chromatography (gradient of 0-100% EtOAc in hexane) to give 1.15 g of desired aldehyde in 78% yield. $^1$H NMR (400 MHz, $CDCl_3$) δ 9.48 (s, 1H), 7.60 (dd, *J* = 3.0, 1.3 Hz, 1H), 7.40 (dd, *J* = 5.1, 2.9 Hz, 1H), 7.32 (dd, *J* = 5.1, 1.3 Hz, 1H), 7.16 (s, 1H), 2.57 (q, *J* = 7.6 Hz, 2H), 1.11 (t, *J* = 7.5 Hz, 3H).

・**Synthesis of Imine**

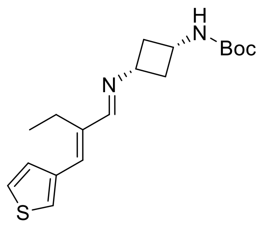

General procedure A was followed using above obtained aldehyde (200 mg, 1.20 mmol, 1.0 equiv), *cis*-*tert*-butyl *N*-(3-aminocyclobutyl)carbamate (246 mg, 1.32 mmol, 1.1 equiv), molecular sieves 4 Å, and THF (2.4 mL). Imine (370 mg, 99% yield) was obtained as a white solid. $^1$H NMR (400 MHz, $CD_3OD$) δ 7.82 (d, *J* = 1.3 Hz, 1H), 7.52 (d, *J* = 2.4 Hz, 1H), 7.44 (dd, *J* = 5.1, 2.9 Hz, 1H), 7.26 (dd, *J* = 5.0, 1.3 Hz, 1H), 6.81 (s, 1H), 3.86 (p, *J* = 8.4 Hz, 1H), 3.74 (p, *J* = 7.9 Hz, 1H), 2.75 – 2.52 (m, 4H), 1.99 (qd, *J* = 9.0, 2.8 Hz, 2H), 1.42 (s, 9H), 1.11 (t, *J* = 7.5 Hz, 3H).

・**Key Reaction**

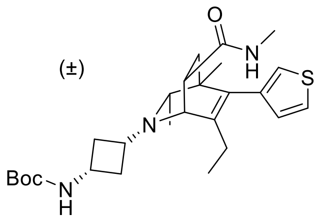

General procedure B was followed using above obtained imine (80 mg, 0.240 mmol, 1.0 equiv) and 2-butyne (94 µL, 1.20 mmol, 5 equiv) in THF (0.24 mL). Rh catalyst (10 mol %, 0.24 mL, 0.024 mmol, 100 mM in THF) was added, and the reaction was carried out at 65 °C for 12 h to produce the DHP intermediate. Following general procedure C, *N*-methyl acryamide (204 mg, 2.40 mmol, 10 equiv) was added to the solution of DHP intermediate, and the reaction was carried out at 90 °C for 16 h. The resulting crude material was purified by column chromatography (5% MeOH/$CH_2Cl_2$ + 1% $NH_4OH$) to afford the desired product (26 mg, 23% yield from imine) as a white solid. $^1$H NMR (400 MHz, $CD_3OD$) δ 7.34 (dd, *J* = 4.9, 2.9 Hz, 1H), 7.00 (dd, *J* = 2.9, 1.2 Hz, 1H), 6.87 (dd, *J* = 4.9, 1.2 Hz, 1H), 3.74 (p, *J* = 8.5 Hz, 1H), 3.45 (d, *J* = 3.0 Hz, 1H), 3.02 (dt, *J* = 8.9, 4.1 Hz, 1H), 2.94 (p, *J* = 8.5, 7.9 Hz, 1H), 2.66 (s, 3H), 2.65 – 2.57 (m, 1H), 2.53 – 2.41 (m, 1H), 2.19 (q, *J* = 6.3 Hz, 1H), 1.96 – 1.85 (m, 3H), 1.74 (q, *J* = 9.7 Hz, 1H), 1.49-1.34 (m, 11H), 0.97 (d, *J* = 6.2 Hz, 3H), 0.84 (t, *J* = 7.6 Hz, 3H), 0.82 (s, 3H).

・**Deprotection**

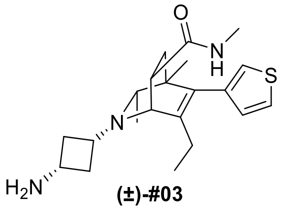

**(±)-#03**

**Final Compound #03** General Procedure D was followed using protected isoquinuclidine (26 mg, 0.055 mmol, 1.00 equiv) in $CH_2Cl_2$ (1.0 mL) and was reacted with trifluoroacetic acid (41 µL, 0.55 mmol, 10.0 equiv) for 12 h. The crude product was purified by preparative thin-layer chromatography (10% MeOH/$CH_2Cl_2$ + 1% $NH_4OH$) to yield the racemic isoquinuclidine (16 mg, 78% yield) as a pale-yellow oil. $^1$H NMR (400 MHz, $CD_3OD$) δ 7.35 (dd, *J* = 4.9, 3.0 Hz, 1H), 7.00 (dd, *J* = 3.0, 1.2 Hz, 1H), 6.87 (dd, *J* = 4.9, 1.2 Hz, 1H), 3.47 (d, *J* = 3.0 Hz, 1H), 3.12 (p, *J* = 8.3 Hz, 1H), 3.01 (dt, *J* = 8.7, 4.0 Hz, 1H), 2.91 (p, *J* = 7.9 Hz, 1H), 2.68 – 2.59 (m, 1H), 2.66 (s, 3H), 2.55 – 2.33 (m, 1H), 2.18 (q, *J* = 6.3 Hz, 1H), 2.09 – 1.76 (m, 4H), 1.66 (q, *J* = 9.6 Hz, 1H), 1.39 (dd, *J* = 12.8, 9.4 Hz, 1H), 0.98 (d, *J* = 6.2 Hz, 3H), 0.84 (t, *J* = 7.8 Hz, 3H), 0.82 (s, 3H). $^{13}$C NMR (101 MHz, $CD_3OD$) δ 174.8, 141.2, 138.5, 134.8, 128.5, 124.1, 121.8, 62.9, 56.7, 50.2, 41.5, 40.4, 39.3, 38.7, 38.3, 35.3, 25.3, 24.5, 20.6, 19.9, 11.4. HRMS (ESI+, m/z) calcd for $C_{21}H_{31}N_3OS$ $(M+H)^+$ : 374.2261, found: 374.2248.

**Synthesis of #05**

・**Synthesis of Imine**

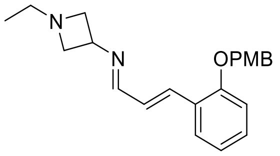

General procedure A was followed using (E)-3-(2-((4 methoxybenzyl)oxy)phenyl)acrylaldehyde (121 mg, 0.449 mmol, 0.9 equiv), 1-ethylazetidin-3-amine (50 mg, 0.50 mmol, 1.0 equiv), molecular sieves 4 Å, and THF (1.0 mL). Imine (175 mg, 99% yield) was obtained as pale-yellow oil. $^1$H NMR (400 MHz, $CDCl_3$) δ 7.96 (d, J = 8.9 Hz, 1H), 7.55 (dd, J = 8.1, 1.7 Hz, 1H), 7.37 – 7.33 (m, 2H), 7.28 – 7.24 (m, 2H), 7.19 – 7.15 (m, 1H), 6.97 – 6.91 (m, 4H), 5.05 (s, 2H), 4.23 (p, J = 6.7 Hz, 1H), 3.83 (m, 5H), 3.17 (t, J = 7.3 Hz, 2H), 2.62 (q, J = 7.2 Hz, 2H), 1.04 (t, J = 7.2 Hz, 3H).

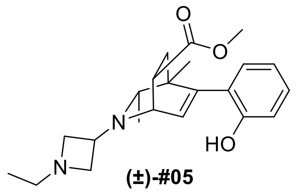

**(±)-#05**

**Final Compound #05** Following General Procedure B for the formation of 1,2-dihydropyridines, the reaction was set up with above obtained imine (200 mg, 0.571 mmol, 1.00 equiv), Rh stock solution (857 µL, 85.7 µmol, 15 mol %), 2-butyne (447 µL, 5.71 mmol, 10.0 equiv), and a reaction time of 20 h at 90 ºC. Isoquinuclidine formation was conducted according to General Procedure C with methyl acrylate (517 µL, 5.71 mmol, 10.0 equiv) for 16 h at 90 ºC. After reaction completion, the solvent was removed in vacuo and the crude isoquinuclidine product was carried forward to deprotection without purification. Following General Procedure D, $CH_2Cl_2$ (4.76 mL) was added to the crude protected isoquinuclidine and was reacted with trifluoroacetic acid (437 µL, 5.71 mmol, 10.0 equiv) for 10 h. The crude product was purified by silica gel chromatography (5% MeOH/$CH_2Cl_2$ + 1% $NH_4OH$) to yield two isomers of the final product. Isomer 1 (50 mg, 24% yield over three steps) and Isomer 2 (12 mg, 5.7% yield over three steps) were isolated as pale-yellow oil.

Isomer 1

$^1$H NMR (500 MHz, $CDCl_3$) δ 7.16 (td, $J$ = 7.7, 1.7 Hz, 1H), 7.00 (dd, $J$ = 7.6, 1.7 Hz, 1H), 6.89 – 6.84 (m, 2H), 6.55 (d, $J$ = 6.4 Hz, 1H), 3.90 (dd, $J$ = 6.4, 1.8 Hz, 1H), 3.76 (s, 3H), 3.52 – 3.47 (m, 3H), 2.87 – 2.84 (m, 1H), 2.73 (m, 1H), 2.64 – 2.57 (m, 2H), 2.48 (m, 2H), 2.18 (dd, $J$ = 13.3, 5.7 Hz, 1H), 1.78 (dd, $J$ = 13.3, 11.3 Hz, 1H), 0.99 – 0.95 (m, 6H), 0.91 (d, $J$ = 6.4 Hz, 3H). $^{13}$C NMR (126

MHz, $CDCl_3$) δ 174.5, 152.9, 143.8, 133.8, 129.5, 128.6, 126.1, 119.8, 115.3, 63.8, 61.7, 59.8, 53.6, 52.0, 51.9, 51.8, 45.3, 41.8, 34.8, 20.6, 18.5, 12.4. HRMS (ESI+, m/z) $[M+H]^+$ calcd: 371.2329, found: 371.2323

Isomer 2

$^1$H NMR (500 MHz, $CDCl_3$) δ 7.17 (td, *J* = 7.7, 1.7 Hz, 1H), 6.98 (dd, *J* = 7.6, 1.7 Hz, 1H), 6.93 (d, *J* = 8.1 Hz, 1H), 6.84 (d, *J* = 7.3 Hz, 1H), 6.25 (d, *J* = 6.4 Hz, 1H), 3.74 – 3.69 (m, 5H), 3.61 (dd, *J* = 6.4, 3.3 Hz, 1H), 3.54 – 3.49 (m, 1H), 3.24 (m, 1H), 2.99 (t, *J* = 7.0 Hz, 1H), 2.92 (t, *J* = 6.1, 4.9 Hz, 1H), 2.58 (m, 2H), 2.20 (q, *J* = 6.4 Hz, 1H), 1.88 (dd, *J* = 13.6, 3.6 Hz, 1H), 1.58 (dd, *J* = 13.5, 9.2 Hz, 1H), 1.04 – 1.00 (m, 6H), 0.90 (s, 3H). $^{13}$C NMR (126 MHz, $CDCl_3$) δ 177.3, 154.0, 144.4, 130.5, 128.8, 128.8, 119.3, 116.3, 113.8, 77.3, 77.0, 76.8, 63.1, 60.3, 59.0, 53.0, 52.7, 52.1, 39.3, 36.0, 20.9, 20.2, 12.0. HRMS (ESI+, m/z) $[M+H]^+$ calcd: 371.2329, found: 371.2324

**Synthesis of #06**

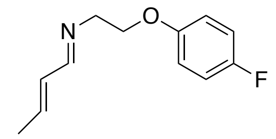

General Procedure A was followed using 2-(4-fluorophenoxy)ethan-1-amine (100 mg, 0.644 mmol, 1.00 equiv), *trans*-crotonaldehyde (0.26 mL, 3.22 mmol, 5.00 equiv), $K_2CO_3$ (31.2 mg, 0.225 mmol, 0.35 equiv), and THF (1.29 mL) for 4 h at rt. The product imine was obtained as pale-yellow oil (133 mg, 100%). $^1$H NMR (400 MHz, $CDCl_3$) δ 7.92 (m, 1H), 6.96 – 6.92 (m, 2H), 6.86 – 6.82 (m, 2H), 6.25 (m, 2H), 4.15 (t, *J* = 5.6 Hz, 2H), 3.79 (t, *J* = 5.6 Hz, 2H), 1.89 (d, *J* = 5.1 Hz, 3H).

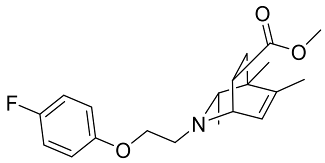

**(±)-#06**

**Final Compound #06** Following General Procedure B for the formation of 1,2-dihydropyridines, the reaction was set up with above obtained imine (50.0 mg, 0.241 mmol, 1.00 equiv), Rh stock solution (241 µL, 24.1 µmol, 10 mol %), 2-butyne (189 µL, 2.41 mmol, 10.0 equiv), and a reaction time of 20 h at 70 ºC. Isoquinuclidine formation was conducted according to the General Procedure C with methyl acrylate (218 µL, 2.41 mmol, 10.0 equiv) for 16 h at 90 ºC. The product after the Diels-Alder reaction was purified by silica gel chromatography (50% EtOAc/hexanes + 1% $Et_3N$) to afford the isoquinuclidine product (20 mg, 24% yield over two steps) as pale-yellow oil. $^1$H NMR (400 MHz, $CDCl_3$) δ 6.94 (t, *J* = 8.7 Hz, 2H), 6.82 (m, 2H), 6.01 (d, *J* = 6.0 Hz, 1H), 4.06 (t, *J* = 6.3 Hz, 2H), 3.76 (m, 1H), 3.62 (s, 3H), 3.18 (m, 1H), 2.99 (m, 2H), 2.10 (m, 1H), 1.74 (m, 4H), 1.45 (dd, *J* = 13.0, 9.5

Hz, 1H), 1.08 (s, 3H), 0.85 (d, *J* = 6.3 Hz, 3H). $^{13}$C NMR (126 MHz, $CDCl_3$) δ 161.7, 158.2(d, *J* = 5.1 Hz), 156.3, 154.8-154.6 (m), 124.9, 115.9, 115.7, 115.4, 115.4, 68.2, 65.0, 54.9, 53.2, 51.8, 40.3, 37.5, 35.4, 19.8, 19.7, 18.8. $^{19}$F NMR (376 MHz, $CDCl_3$) δ -124.08. HRMS (ESI+, m/z) $[M+H]^+$ calcd: 348.1969, found: 348.1966

**#09 (mixture of diastereomers)**

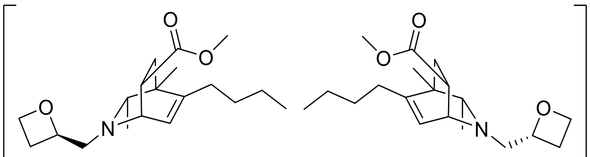

**#09**
mixture of diastereomers

**Final Compound #09** Following General Procedure B for the formation of 1,2-dihydropyridines, the reaction was set up with (1*E*,2*E*)-*N*-(((*R*)-oxetan-2-yl)methyl)hept-2-en-1-imine (175 mg, 0.965 mmol, 1.00 equiv), Rh stock solution (965 µL, 96.5 µmol, 10 mol %), 2-butyne (755 µL, 9.65 mmol, 10.0 equiv), and a reaction time of 14 h at 70 ºC. Isoquinuclidine formation was conducted according to the General Procedure C with methyl acrylate (874 µL, 9.65 mmol, 10.0 equiv) for 16 h at 90 ºC. The product after the Diels-Alder reaction was purified by silica gel chromatography (8% MeOH/$CH_2Cl_2$ + 1% $NH_4OH$) to afford the isoquinuclidine product (86 mg, 28% yield over two steps) as a mixture of diastereomers.

$^1$H NMR (500 MHz, $CDCl_3$) δ 5.93 (d, *J* = 6.4 Hz, 1H), 4.99 – 4.90 (m, 1H), 4.65 (m, 1H), 4.51 (m, 1H), 3.68 (dd, *J* = 6.6, 3.2 Hz, 1H), 3.61 (s, 3H), 3.15 (m, 1H), 2.99 (dd, *J* = 13.0, 7.1 Hz, 1H), 2.94 – 2.76 (m, 1H), 2.74 – 2.63 (m, 2H), 2.44 – 2.29 (m, 1H), 2.08 – 1.91 (m, 3H), 1.72 (dd, *J* = 12.9, 4.7 Hz, 1H), 1.40 (m, 3H), 1.31 (m, 3H), 1.07 (s, 3H), 0.88 (t, *J* = 7.1 Hz, 3H), 0.77 (d, *J* = 6.0 Hz, 3H). $^{13}$C NMR (126 MHz, $CDCl_3$) δ 175.1, 174.9, 147.5, 146.9, 123.7, 123.1, 82.1, 81.4, 77.3, 77.0, 76.8, 68.3, 68.3, 64.8, 64.3, 61.9, 61.0, 56.0, 54.6, 51.7, 40.5, 40.3, 37.8, 37.5, 35.9, 35.4, 31.3, 29.3, 29.3, 26.5, 26.3, 22.5, 22.5, 19.9, 19.7, 19.5, 14.0. HRMS (ESI+, m/z) $[M+H]^+$ calcd: 322.2377, found: 322.2368

**Synthesis of #11**

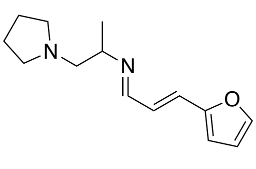

General Procedure A was followed using 1-(pyrrolidin-1-yl)propan-2-amine (200 mg, 1.56 mmol, 1.00 equiv), (*E*)-3-(furan-2-yl)acrylaldehyde (171 mg, 1.40 mmol, 0.9 equiv), 3 Å MS (780 mg), and THF (3.12 mL) for 4 h at 80 ºC. The product imine was obtained as pale-yellow oil (362 mg, 100%). $^1$H

NMR (400 MHz, $CDCl_3$) δ 7.98 (d, *J* = 8.8 Hz, 1H), 7.43 (s, 1H), 6.84 – 6.67 (m, 2H), 6.48 – 6.40 (m, 2H), 3.40 (m, 1H), 2.64 – 2.58 (m, 2H), 2.51 (m, 4H), 1.73 (m, 4H), 1.20 (d, *J* = 6.4 Hz, 3H).

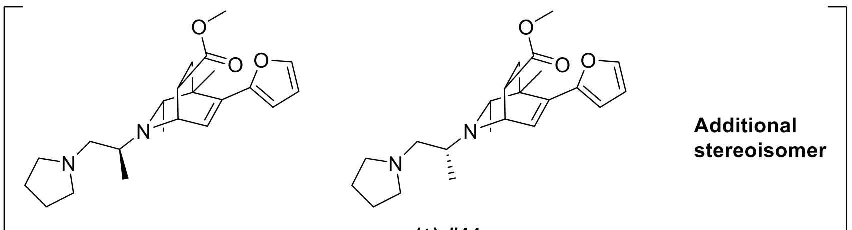

**(±)-#11**
3 isomers not assigned to specific structures

**Final Compound #11** Following General Procedure B for the formation of 1,2-dihydropyridines, the reaction was set up with above obtained imine (300 mg, 1.29 mmol, 1.00 equiv), Rh stock solution (1.94 mL, 0.194 mmol, 15 mol %), 2-butyne (1.01 mL, 12.9 mmol, 10.0 equiv), and a reaction time of 20 h at 90 ºC. Isoquinuclidine formation was conducted according to the General Procedure C with methyl acrylate (1.17 mL, 12.9 mmol, 10.0 equiv) for 16 h at 90 ºC. The product after the Diels-Alder reaction was purified by silica gel chromatography (5% MeOH/$CH_2Cl_2$ + 1% $NH_4OH$) to afford three isomers of the isoquinuclidine product. Isomer 1 (135 mg, 28% yield over two steps), Isomer 2 (135 mg, 28% yield over two steps), and Isomer 3 (17 mg, 3.5% yield over two steps) were isolated as pale-yellow oil.

Isomer 1:

$^1$H NMR (500 MHz, $CDCl_3$) δ 7.32 (d, *J* = 1.7 Hz, 1H), 6.59 (d, *J* = 6.6 Hz, 1H), 6.31 (m, 2H), 4.17 (dd, *J* = 6.7, 3.1 Hz, 1H), 3.62 (m, 4H), 3.23 (dt, *J* = 9.9, 3.8 Hz, 1H), 2.80 (m, 1H), 2.65 (dd, *J* = 12.1, 5.6 Hz, 1H), 2.54 – 2.46 (m, 4H), 2.35 (m, 1H), 1.95 (dd, *J* = 13.1, 4.3 Hz, 1H), 1.74 (m, 4H), 1.55 (dd, *J* = 13.2, 9.8 Hz, 1H), 1.35 (s, 3H), 1.19 (d, *J* = 6.4 Hz, 3H), 0.89 (d, *J* = 6.0 Hz, 3H). $^{13}$C NMR (126 MHz, $CDCl_3$) δ 174.7, 152.8, 141.4, 137.1, 127.8, 110.7, 107.1, 62.8, 62.0, 55.3, 54.7, 51.7, 50.9, 40.7, 39.5, 35.9, 23.5, 22.2, 20.9, 20.1. HRMS (ESI+, m/z) $[M+H]^+$ calcd: 373.2486, found: 373.2477

Isomer 2:

$^1$H NMR (500 MHz, $CDCl_3$) δ 7.32 (m, 1H), 6.56 (d, *J* = 6.7 Hz, 1H), 6.31 (m, 2H), 4.13 (dd, *J* = 7.0, 3.3 Hz, 1H), 3.62 (m, 4H), 3.15 (dt, *J* = 9.7, 3.7 Hz, 1H), 2.60 – 2.45 (m, 6H), 2.27 (m, 1H), 1.98 (dd, *J* = 13.1, 4.1 Hz, 1H), 1.76 (m, 4H), 1.52 (dd, *J* = 13.2, 9.6 Hz, 1H), 1.36 (s, 3H), 1.26 (d, *J* = 6.2 Hz, 3H), 0.90 (d, *J* = 6.0 Hz, 3H). $^{13}$C NMR (126 MHz, $CDCl_3$) δ 174.5, 152.6, 141.5, 137.6, 127.0, 110.7, 107.3, 63.7, 61.9, 54.9, 54.7, 51.8, 50.7, 40.7, 35.9, 35.3, 23.5, 22.2, 21.3, 17.5. HRMS (ESI+, m/z) $[M+H]^+$ calcd: 373.2486, found: 373.2491

Isomer 3:

$^1$H NMR (500 MHz, $CDCl_3$) δ 7.34 (d, *J* = 1.8 Hz, 1H), 6.76 (d, *J* = 6.2 Hz, 1H), 6.35 (dd, *J* = 3.3, 1.8 Hz, 1H), 6.30 (d, *J* = 3.4 Hz, 1H), 3.87 (dd, *J* = 6.3, 2.0 Hz, 1H), 3.70 (s, 3H), 3.12 (q, *J* = 6.4 Hz, 1H), 2.97 (m, 1H), 2.59 – 2.41 (m, 7H), 2.13 (dd, *J* = 13.1, 5.4 Hz, 1H), 1.80 – 1.72 (m, 4H), 1.55 (dd, *J* = 13.1, 10.9 Hz, 1H), 1.37 (s, 3H), 1.04 (d, *J* = 6.6 Hz, 3H), 0.86 (d, *J* = 6.3 Hz, 3H). $^{13}$C NMR (126 MHz, $CDCl_3$) δ 174.6, 153.0, 141.3, 135.6, 130.2, 110.9, 106.8, 63.3, 54.4, 51.6, 51.2, 50.8, 46.5, 41.6, 34.5, 29.7, 23.4, 22.1, 19.0, 15.7. HRMS (ESI+, m/z) $[M+H]^+$ calcd: 373.2486, found: 373.2491

**Synthesis of #12**

・**Synthesis of Amine**

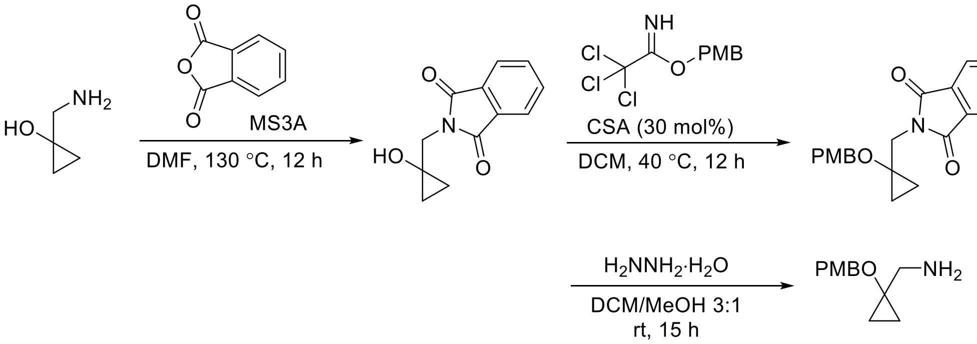

To a flame-dried round-bottom flask was added a magnetic stir bar, 1-(aminomethyl)cyclopropanol (1.0 g, 11.5 mmol, 1.0 equiv) and phthalic anhydride (1.7 g, 11.5 mmol, 1.0 equiv) dissolved in DMF (23 mL). To this solution was then added molecular sieves 4 Å. The reaction mixture was stirred at 130 °C for 12 h. After the reaction was complete as monitored by thin-layer chromatography, the mixture was concentrated and the crude residue was purified via silica gel chromatography (gradient of 0-100% EtOAc in hexane +1% $Et_3N$) to give 2.2 g of the amine protected compound in 88% yield. To a stirred solution of the above obtained compound (1.0 g, 4.6 mmol, 1.0 equiv) in DCM (23 mL) was added 4-methoxybenzyl-2,2,2-trichloroacetimidate (1431 mg, 5.06 mmol, 1.1 equiv) and 10-camphorsulfonic acid (321 mg, 1.38 mmol, 0.3 equiv). The reaction mixture was stirred at 40 °C for 12 h. After the reaction was complete as monitored by thin-layer chromatography, the mixture was then diluted with saturated $NaHCO_3$ (aq), extracted with DCM, washed with brine, dried with $Na_2SO_4$ and filtered. The organic layer was concentrated and the crude residue was purified via silica gel chromatography (gradient of 0-100% EtOAc in hexane) to give 1.3 g of alcohol protected compound in 84% yield. To a stirred solution of the above obtained compound (1.3 g, 3.85 mmol, 1.0 equiv) in MeOH/DCM 1:3 (39 mL) was added hydrazine monohydrate (2.9 g, 57.8 mmol, 15 equiv) and the reaction mixture was stirred at room temperature for 15 h. After the reaction was complete as monitored by thin-layer chromatography, the mixture was concentrated and the crude residue was purified via silica gel chromatography (10% MeOH/$CH_2Cl_2$ + 1% $NH_4OH$) to give 320 mg of desired amine in 40% yield. $^{1}$H NMR (500 MHz, $CDCl_3$) δ 7.23 (d, *J* = 8.2 Hz, 2H), 6.86 (d, *J* = 8.2 Hz, 2H), 4.45 (s, 2H), 3.79 (s, 3H), 2.80 (br, 2H), 0.89 (t, *J* = 6.0 Hz, 2H), 0.54 (t, *J* = 6.0 Hz, 2H).

・**Synthesis of Imine**

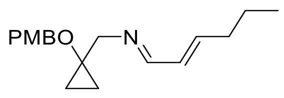

General procedure A was followed using above obtained amine (150 mg, 0.724 mmol, 1.0 equiv), *trans*-2-hexenal (71 mg, 0.724 mmol, 1.0 equiv), molecular sieves 4 Å, and DCM (1.5 mL). Imine (180 mg, 95% yield) was obtained as a pale-yellow oil. $^{1}$H NMR (400 MHz, $CDCl_3$) δ 7.82 (d, *J* = 8.5 Hz, 1H), 7.18 (d, *J* = 8.5 Hz, 2H), 6.81 (d, *J* = 8.6 Hz, 2H), 6.30 – 6.09 (m, 2H), 4.50 (s, 2H), 3.77 (s, 3H), 3.67 (s, 2H), 2.17 (q, *J* = 7.3 Hz, 2H), 1.47 (h, *J* = 7.3 Hz, 2H), 0.96 – 0.88 (m, 5H), 0.68 – 0.63 (dd, *J* = 7.2, 5.2 Hz, 2H).

・**Key Reaction**

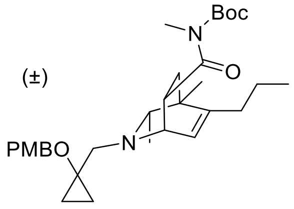

General procedure B was followed using above obtained imine (120 mg, 0.418 mmol, 1.0 equiv) and 2-butyne (164 µL, 2.09 mmol, 5 equiv) in THF (0.42 mL). Rh catalyst (10 mol %, 0.42 mL, 0.042 mmol, 100 mM in THF) was added, and the reaction was carried out at 65 °C for 12 h to produce the DHP intermediate. Following general procedure C, Boc-protected *N*-methyl acrylamide (773 mg, 4.18 mmol, 10 equiv) was added to the solution of DHP intermediate, and the reaction was carried out at 50 °C for 12 h. The resulting crude material was purified by column chromatography (30% *t*BuOMe/Pentane+1% $Et_3N$) to afford the desired product (80 mg, 36% yield from imine) as a pale-yellow oil. $^{1}$H NMR (400 MHz, $CD_3OD$) δ 7.18 (d, *J* = 8.6 Hz, 2H), 6.81 (d, *J* = 8.6 Hz, 2H), 5.92 (dt, *J* = 6.5, 1.8 Hz, 1H), 4.55 (d, *J* = 10.6 Hz, 1H), 4.48 (d, *J* = 10.6 Hz, 1H), 4.28 – 4.21 (m, 1H), 3.79 – 3.71 (m, 4H), 3.08 (d, *J* = 14.2 Hz, 1H), 3.00 (s, 3H), 2.71 (d, *J* = 14.2 Hz, 1H), 2.17 – 1.90 (m, 3H), 1.59 – 1.40 (m, 13H), 1.06 (s, 3H), 0.93 (t, *J* = 7.3 Hz, 3H), 0.87 (d, *J* = 6.2 Hz, 3H), 0.85 – 0.81 (m, 2H), 0.63 – 0.60 (m, 2H).

・**Deprotection**

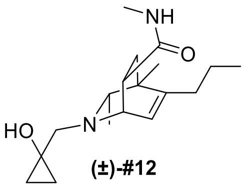

(±)-#12

**Final Compound #12** General Procedure D was followed using protected isoquinuclidine (80 mg, 0.152 mmol, 1.00 equiv) in $CH_2Cl_2$ (1.5 mL) and was reacted with trifluoroacetic acid (113 µL, 1.52 mmol, 10.0 equiv) for 12 h. The crude product was purified by preparative thin-layer chromatography (10% MeOH/$CH_2Cl_2$ + 1% $NH_4OH$) to yield the racemic isoquinuclidine (28 mg, 60% yield) as a pale-yellow oil. $^1$H NMR (400 MHz, $CD_3OD$) δ 5.95 (dt, *J* = 6.5, 1.9 Hz, 1H), 3.70 (dd, *J* = 6.5, 2.7 Hz, 1H), 3.00 (ddd, *J* = 9.5, 5.2, 2.7 Hz, 1H), 2.88 (d, *J* = 13.4 Hz, 1H), 2.65 (s, 3H), 2.58 (d, *J* = 13.4 Hz, 1H), 2.16 – 1.92 (m, 3H), 1.59 (dd, *J* = 12.7, 5.2 Hz, 1H), 1.55 – 1.38 (m, 3H), 1.09 (s, 3H), 0.95 (t, *J* = 7.3 Hz, 3H), 0.87 (d, *J* = 6.2 Hz, 3H), 0.71 – 0.69 (m, 2H), 0.60 – 0.51 (m, 2H). $^{13}$C NMR (101 MHz, $CD_3OD$) δ 175.7, 146.5, 123.4, 64.9, 60.9, 55.6, 52.9, 40.3, 37.9, 36.0, 33.4, 25.1, 20.1, 18.6, 18.2, 12.9, 11.5, 11.3. HRMS (ESI+, m/z) calcd for $C_{18}H_{31}N_2O_2$ $(M+H)^+$ : 307.2380, found: 307.2375.

**Synthesis of #14**

・**Synthesis of Imine**

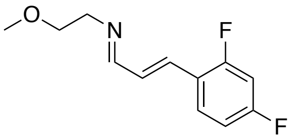

General procedure A was followed using 2-methoxyethan-1-amine (100 mg, 1.33 mmol, 1.00 equiv), (*E*)-3-(2,4-difluorophenyl)acrylaldehyde (224 mg, 1.33 mmol, 1.00 equiv), $K_2CO_3$ (64 mg, 0.466 mmol, 0.35 equiv), and THF (2.66 mL) for 4 h at rt. The product imine was obtained as pale-yellow oil (198 mg, 66%). $^1$H NMR (400 MHz, $CDCl_3$) δ 8.06 (m, 1H), 7.51 (m, 1H), 7.06 (m, 1H), 6.96 – 6.79 (m, 3H), 3.71 (m, 2H), 3.69 – 3.64 (m, 2H), 3.38 (s, 3H).

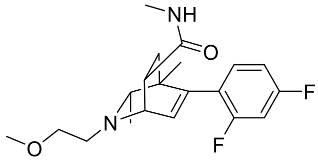

(±)-#14

**Final Compound #14** Following General Procedure B for the formation of 1,2-dihydropyridines, the reaction was set up with above obtained imine (40.0 mg, 0.178 mmol, 1.00 equiv), Rh stock solution (178 µL, 17.8 µmol, 10 mol %), 2-butyne (139 µL, 1.78 mmol, 10.0 equiv), and a reaction time of 20 h at 70 ºC. Isoquinuclidine formation was conducted according to the General Procedure C with N-methylacrylamide (151 mg, 1.78 mmol, 10.0 equiv) for 16 h at 90 ºC. The product after the Diels-

Alder reaction was purified by silica gel chromatography (50% EtOAc/hexanes + 1% $Et_3N$) to afford the isoquinuclidine product as pale-yellow oil (13.3 mg, 21% yield over two steps). $^1H$ NMR (400 MHz, $CDCl_3$) δ 7.16 (td, *J* = 8.4, 6.5 Hz, 1H), 6.79 (m, 2H), 6.26 (d, *J* = 6.4 Hz, 1H), 5.67 (s, 1H), 3.80 (dd, *J* = 6.6, 2.7 Hz, 1H), 3.58 (m, 2H), 3.36 (s, 3H), 3.16 (m, 1H), 2.86 (m, 2H), 2.75 (d, *J* = 4.8 Hz, 3H), 2.08 (q, *J* = 6.4 Hz, 1H), 1.88 (dd, *J* = 13.1, 5.2 Hz, 1H), 1.52 (dd, *J* = 13.0, 9.5 Hz, 1H), 0.95 (d, *J* = 6.2 Hz, 3H), 0.92 (s, 3H). $^{13}C$ NMR (126 MHz, $CDCl_3$) δ 174.3, 163.2 (d, *J* = 11.8 Hz), 161.3 (d, *J* = 11.9 Hz), 160.7 (d, *J* = 12.4 Hz), 142.5, 132.1(dd, *J* = 9.3, 5.6 Hz), 131.6, 111.1 (dd, *J* = 20.9, 3.4 Hz), 103.79 – 103.28 (m), 72.5, 65.1, 58.9, 55.4, 53.1, 41.2, 38.7, 36.8, 26.5, 20.2 (d, *J* = 2.6 Hz), 19.3. $^{19}F$ NMR (376 MHz, $CDCl_3$) δ -107.9 (m), -111.6 (m). HRMS (ESI+, m/z) $[M+H]^+$ calcd: 365.2035, found: 365.2033

**Synthesis of #17**

・**Synthesis of Amine**

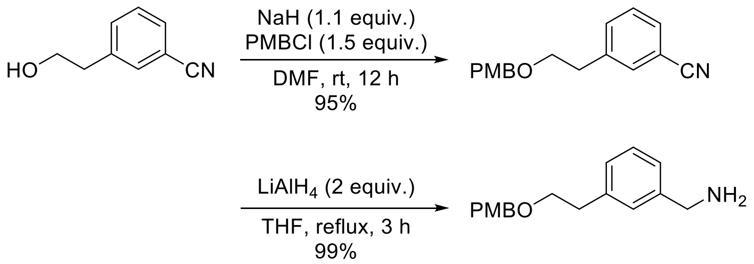

To a flame-dried round-bottom flask was added a magnetic stir bar, 3-(2-hydroxyethyl)benzonitrile (580 mg, 3.94 mmol, 1 equiv) and 4-methoxybenzyl chloride (926 mg, 5.91 mmol, 1.5 equiv) dissolved in DMF (8 mL). To this solution was then added sodium hydride (109 mg, 4.73 mmol, 1.2 equiv). The reaction mixture was stirred at room temperature for 12 h. After the reaction was complete as monitored by thin-layer chromatography, the mixture was then diluted with water, extracted with ethyl acetate, washed with brine, dried with $Na_2SO_4$ and filtered. The organic layer was concentrated and the crude residue was purified via silica gel chromatography (gradient of 0-100% EtOAc in hexane +1% $Et_3N$) to give 1001 mg of the protected alcohol in 95% yield. To a stirred solution of the above obtained compound in THF (37 mL) was added $LiAlH_4$ (284 mg, 7.48 mmol, 2 equiv), and the reaction mixture was heated under reflux for 12 h. After the reaction was complete as monitored by thin-layer chromatography, the mixture was then quenched by careful addition of 1N NaOH and then $H_2O$, extracted with ethyl acetate, washed with brine, dried with $Na_2SO_4$ and filtered. The organic layer was concentrated and the crude residue was purified via silica gel chromatography (5% $MeOH/CH_2Cl_2$ + 1% $NH_4OH$) to give 1005 mg of the amine in 99% yield. $^1H$ NMR (400 MHz, $CDCl_3$) δ 7.30 – 7.18 (m, 3H), 7.14 (d, *J* = 7.8 Hz, 2H), 7.09 (d, *J* = 7.1 Hz, 1H), 6.85 (d, *J* = 8.6 Hz, 2H), 4.44 (s, 2H), 3.82 (s, 2H), 3.79 (s, 3H), 3.65 (t, *J* = 7.2 Hz, 2H), 2.90 (t, *J* = 7.2 Hz, 2H).

・**Synthesis of Imine**

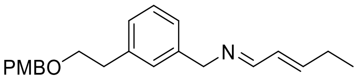

General procedure A was followed using above obtained amine (200 mg, 0.737 mmol, 1.0 equiv), *trans*-2-pentenal (0.36 mL, 3.69 mmol, 5.0 equiv), molecular sieves 4 Å, and THF (1.5 mL). Imine (225 mg, 91% yield) was obtained as a pale-yellow oil. $^1$H NMR (400 MHz, $CDCl_3$) δ 8.02 – 7.89 (m, 1H), 7.23 – 7.18 (m, 3H), 7.16 – 7.02 (m, 3H), 6.85 (d, *J* = 8.6 Hz, 2H), 6.32 – 6.21 (m, 2H), 4.57 (s, 2H), 4.43 (s, 2H), 3.78 (s, 3H), 3.64 (t, *J* = 7.3 Hz, 2H), 2.89 (t, *J* = 7.3 Hz, 2H), 2.22 (dq, *J* = 11.4, 7.3 Hz, 2H), 1.05 (t, *J* = 7.4 Hz, 3H).

・**Key Reaction**

(±)

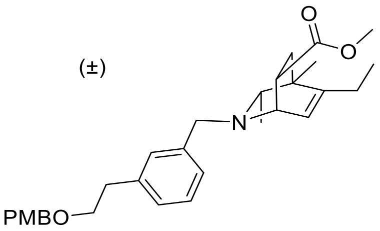

General procedure B was followed using above obtained imine (220 mg, 0.652 mmol, 1.0 equiv) and 2-butyne (256 µL, 3.26 mmol, 5 equiv) in THF (0.65 mL). Rh catalyst (10 mol %, 0.65 mL, 0.065 mmol, 100 mM in THF) was added, and the reaction was carried out at 65 °C for 12 h to produce the DHP intermediate. Following general procedure C, methyl acrylate (0.59 mL, 6.52 mmol, 10 equiv) was added to the solution of DHP intermediate, and the reaction was carried out at 50 °C for 12 h. The resulting crude material was purified by column chromatography (30% *t*BuOMe/Pentane+1% $Et_3N$) to afford the desired product (65 mg, 21% yield from imine) as a pale-yellow oil. $^1$H NMR (400 MHz, $CD_3OD$) δ 7.29 – 7.06 (m, 6H), 6.83 (d, *J* = 8.7 Hz, 2H), 5.81 (dt, *J* = 6.5, 2.0 Hz, 1H), 4.41 (s, 2H), 3.75 (s, 3H), 3.74 – 3.56 (m, 3H), 3.56 (s, 3H), 3.36 – 3.29 (m, 1H), 2.86 (t, *J* = 6.9 Hz, 2H), 2.22 – 1.94 (m, 3H), 1.70 (dd, *J* = 12.9, 4.7 Hz, 1H), 1.52 (dd, *J* = 12.9, 9.4 Hz, 1H), 1.08 (s, 3H), 0.99 (t, *J* = 7.3 Hz, 3H), 0.66 (d, *J* = 6.2 Hz, 3H).

・**Deprotection**

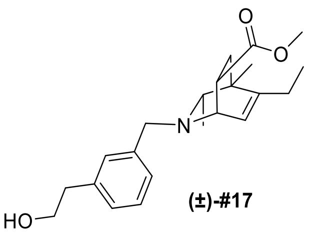

(±)-#17

**Final Compound #17** General Procedure D was followed using protected isoquinuclidine (32 mg, 0.067 mmol, 1.00 equiv) in $CH_2Cl_2$ (1.0 mL) and was reacted with trifluoroacetic acid (50 µL, 0.67

mmol, 10.0 equiv) for 12 h. The crude product was purified by preparative thin-layer chromatography (10% MeOH/$CH_2Cl_2$ + 1% $NH_4OH$) to yield the racemic isoquinuclidine (22 mg, 92% yield) as a pale-yellow oil. $^1H$ NMR (400 MHz, $CD_3OD$) δ 7.30 – 7.17 (m, 3H), 7.12 (dt, $J$ = 6.6, 2.1 Hz, 1H), 5.82 (dt, $J$ = 6.5, 2.0 Hz, 1H), 3.80 – 3.68 (m, 3H), 3.63 – 3.51 (m, 2H), 3.57 (s, 3H), 3.35 (ddd, $J$ = 9.1, 4.5, 2.9 Hz, 1H), 2.80 (t, $J$ = 7.2 Hz, 2H), 2.23 – 1.93 (m, 3H), 1.71 (dd, $J$ = 12.9, 4.7 Hz, 1H), 1.52 (dd, $J$ = 12.9, 9.4 Hz, 1H), 1.09 (s, 3H), 0.99 (t, $J$ = 7.3 Hz, 3H), 0.68 (d, $J$ = 6.2 Hz, 3H). $^{13}C$ NMR (101 MHz, $CD_3OD$) δ 175.3, 149.0, 138.8, 138.5, 130.2, 127.9, 127.6, 127.3, 122.3, 64.1, 62.9, 57.3, 52.8, 50.8, 40.3, 38.8, 36.6, 35.4, 23.9, 18.4, 17.9, 10.4. HRMS (ESI+, m/z) calcd for $C_{22}H_{31}NO_3$ $(M+H)^+$ : 358.2377, found: 358.2365.

**Synthesis of #19**

・**Synthesis of Imine**

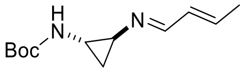

General procedure A was followed using *tert*-butyl *N*-[(1*S*)-2-aminocyclopropyl]carbamate (200 mg, 1.16 mmol, 1.0 equiv), *trans*-crotonaldehyde (0.47 mL, 5.8 mmol, 5.0 equiv), molecular sieves 4 Å, and THF (2 mL). Imine (260 mg, 99% yield) was obtained as a pale-yellow oil. $^1H$ NMR (400 MHz, $CDCl_3$) δ 8.00 – 7.93 (dd, $J$ = 5.2, 2.9 Hz, 1H), 6.16 – 6.12 (m, 2H), 2.87 – 2.79 (m, 2H), 1.85 – 1.82 (m, 3H), 1.41 (s, 9H), 1.30 – 1.17 (m, 1H), 1.07 – 1.02 (m, 1H).

・**Key reaction and Deprotection**

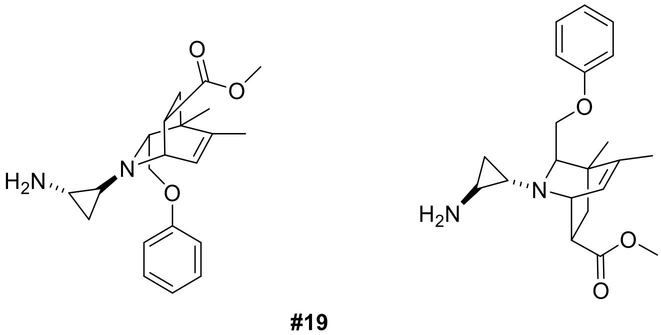

**#19**

Major and Minor diastereomers not assigned to specific structures

**Final Compound #19** General procedure B was followed using above obtained imine (200 mg, 0.892 mmol, 1.0 equiv) and 2-butyne (349 µL, 4.46 mmol, 5 equiv) in THF (0.89 mL). Rh catalyst (10 mol %, 0.89 mL, 0.089 mmol, 100 mM in THF) was added, and the reaction was carried out at 75 °C for 12 h to produce the DHP intermediate. Following general procedure C, methyl acrylate (0.8 mL, 8.92 mmol, 10 equiv) was added to the solution of DHP intermediate, and the reaction was carried out at 50 °C for 12 h. The resulting crude material was purified by preparative thin-layer chromatography

(30% *t*BuOMe/Pentane+1% $Et_3N$) to afford the mixture of diastereomers of protected product (53 mg, 13% yield from imine) as a pale-yellow oil. And General Procedure D was followed using protected isoquinuclidine (53 mg, 0.116 mmol, 1.00 equiv) in $CH_2Cl_2$ (1.2 mL) and was reacted with trifluoroacetic acid (86 µL, 1.16 mmol, 10.0 equiv) for 12 h. The crude product was purified by preparative thin-layer chromatography (10% MeOH/$CH_2Cl_2$ + 1% $NH_4OH$) to yield the diastereomers of isoquinuclidines (23 mg, 56% yield, 2:1 ratio) as a pale-yellow oil.

**Major Diastereomer**

$^1$H NMR (400 MHz, $CD_3OD$) δ 7.38 – 7.12 (m, 2H), 7.00 – 6.76 (m, 3H), 5.91 – 5.89 (m, 1H), 3.80 (dd, *J* = 10.0, 5.3 Hz, 1H), 3.67 (dd, *J* = 6.4, 3.2 Hz, 1H), 3.64 – 3.57 (m, 4H), 3.40 – 3.30 (m, 1H), 2.73 (t, *J* = 5.1 Hz, 1H), 2.50 – 2.41 (m, 1H), 1.95 – 1.86 (m, 1H), 1.80 – 1.71 (m, 4H), 1.65 (dd, *J* = 13.1, 9.4 Hz, 1H), 1.23 (s, 3H), 0.66 (ddd, *J* = 8.0, 5.6, 4.3 Hz, 1H), 0.58 (ddd, *J* = 7.7, 5.6, 4.3 Hz, 1H). $^{13}$C NMR (101 MHz, $CD_3OD$) δ 175.0, 158.7, 144.3, 129.1, 124.8, 120.3, 113.9, 69.8, 68.5, 54.2, 50.9, 44.3, 39.3, 38.1, 34.9, 32.9, 18.9, 17.1, 12.9. HRMS (ESI+, m/z) calcd for $C_{21}H_{28}N_2O_3$ $(M+H)^+$ : 357.2173, found: 357.2164.

**Minor Diastereomer**

$^1$H NMR (400 MHz, $CD_3OD$) δ 7.22 (dd, *J* = 8.7, 7.3 Hz, 2H), 6.89 – 6.84 (m, 1H), 6.82 (dd, *J* = 8.9, 1.1 Hz, 2H), 5.96– 5.94 (m, 1H), 3.82 (dd, *J* = 6.3, 3.2 Hz, 1H), 3.66 (dd, *J* = 10.1, 6.0 Hz, 1H), 3.63 (s, 3H), 3.55 (dd, *J* = 10.2, 4.2 Hz, 1H), 3.41 (dt, *J* = 9.2, 4.0 Hz ,1H), 2.63 (dd, *J* = 6.0, 4.2 Hz, 1H), 2.33 – 2.24 (m, 1H), 2.05 – 1.96 (m, 1H), 1.78 – 1.70 (m, 4H), 1.65 (dd, *J* = 13.1, 9.4 Hz, 1H), 1.21 (s, 3H), 0.94 (ddd, *J* = 8.0, 5.2, 4.1 Hz, 1H), 0.67 (dt, *J* = 7.4, 4.8 Hz, 1H). $^{13}$C NMR (101 MHz, $CD_3OD$) δ 175.1, 158.7, 144.2, 129.0, 125.0, 120.2, 113.9, 69.8, 67.9, 55.0, 50.9, 44.4, 39.4, 38.3, 34.9, 29.9, 18.9, 17.2, 16.9. HRMS (ESI+, m/z) calcd for $C_{21}H_{28}N_2O_3$ $(M+H)^+$ : 357.2173, found: 357.2184.

**Synthesis of #21**

・**Synthesis of Imine**

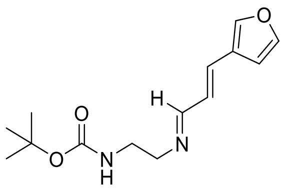

General procedure A was followed using *tert*-butyl *N*-(2-aminoethyl)carbamate (361 mg, 2.25 mmol, 1.0 equiv), (*E*)-3-(furan-3-yl)acrylaldehyde (275 mg, 2.25 mmol, 1.0 equiv), molecular sieves 4 Å, and THF (5 mL). Imine (590 mg, 99% yield) was obtained as a pale-yellow oil. $^1$H NMR (400 MHz,

$CDCl_3$) δ 7.96 (d, *J* = 9.0 Hz, 1H), 7.57 (s, 1H), 7.41 (t, *J* = 1.9 Hz, 1H), 6.87 (d, *J* = 15.9 Hz, 1H), 6.63 (dd, *J* = 15.9, 9.0 Hz, 1H), 6.59 (d, *J* = 1.9 Hz, 1H), 4.84 (br, 1H), 3.59 (t, *J* = 5.8 Hz, 2H), 3.41 (q, *J* = 5.8 Hz, 2H), 1.42 (s, 9H).

・**Key Reaction**

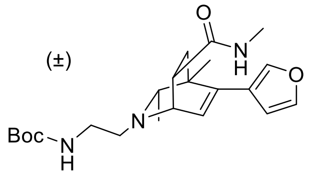

General procedure B was followed using above obtained imine (200 mg, 0.757 mmol, 1.0 equiv) and 2-butyne (296 µL, 3.78 mmol, 5 equiv) in THF (0.76 mL). Rh catalyst (10 mol %, 0.76 mL, 0.076 mmol, 100 mM in THF) was added, and the reaction was carried out at 65 °C for 12 h to produce the DHP intermediate. Following general procedure C, *N*-methyl acrylamide (644 mg, 7.57 mmol, 10 equiv) was added to the solution of DHP intermediate, and the reaction was carried out at 90 °C for 16 h. The resulting crude material was purified by column chromatography (5% MeOH/$CH_2Cl_2$ + 1% $NH_4OH$) to afford the desired product (27 mg, 9% yield from imine) as a pale-yellow oil. $^1$H NMR (400 MHz, $CD_3OD$) δ 7.46 (s, 1H), 7.41 (t, *J* = 1.8 Hz, 1H), 6.43 (d, *J* = 1.8, Hz, 1H), 6.26 (d, *J* = 6.5 Hz, 1H), 3.62 (dd, *J* = 6.5, 2.8 Hz, 1H), 3.23 – 3.07 (m, 1H), 2.81 – 2.68 (m, 4H), 2.67 (s, 3H), 2.23 – 2.08 (m, 1H), 1.81 (dd, *J* = 12.8, 4.8 Hz, 1H), 1.56 – 1.48 (m, 1H), 1.42 (s, 9H), 1.17 (s, 3H), 0.87 (d, *J* = 6.2 Hz, 3H).

・**Deprotection**

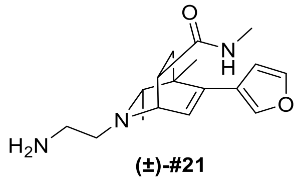

**(±)-#21**

**Final Compound #21** General Procedure D was followed using protected isoquinuclidine (27 mg, 0.067 mmol, 1.00 equiv) in $CH_2Cl_2$ (1.0 mL) and was reacted with trifluoroacetic acid (50 µL, 0.67 mmol, 10.0 equiv) for 12 h. The crude product was purified by preparative thin-layer chromatography (10% MeOH/$CH_2Cl_2$ + 1% $NH_4OH$) to yield the racemic isoquinuclidine (17.5 mg, 86% yield) as a pale-yellow oil. $^1$H NMR (500 MHz, $CD_3OD$) δ 7.47 (s, 1H), 7.42 (t, *J* = 1.8 Hz, 1H), 6.45 (d, *J* = 1.8 Hz, 1H), 6.28 (d, *J* = 6.5 Hz, 1H), 3.58 (dd, *J* = 6.5, 2.8 Hz, 1H), 3.14 (ddd, *J* = 9.5, 5.0, 2.8 Hz, 1H), 2.80 – 2.68 (m, 4H), 2.68 (s, 3H), 2.12 (q, *J* = 6.2 Hz, 1H), 1.81 (dd, *J* = 12.8, 5.0 Hz, 1H), 1.53 (dd, *J* = 12.8, 9.5 Hz, 1H), 1.19 (s, 3H), 0.88 (d, *J* = 6.2 Hz, 3H). $^{13}$C NMR (126 MHz, $CD_3OD$) δ 175.4, 142.3, 139.1, 138.4, 127.4, 124.1, 110.1, 64.6, 56.0, 55.1, 40.7, 39.7, 38.0, 35.9, 25.2, 20.9, 18.9. HRMS (ESI+, m/z) calcd for $C_{17}H_{26}N_3O_2$ $(M+H)^+$ : 304.2020, found: 304.2011.

**Synthesis of #33**

・**Synthesis of imine**

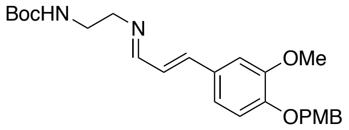

General Procedure A.1 was followed using *tert*-butyl (2-aminoethyl)carbamate (100 mg, 0.624 mmol 1.00 equiv), (*E*)-3-(3-methoxy-4-((4-methoxybenzyl)oxy)phenyl)acrylaldehyde (186 mg, 0.624 mmol, 1.00 equiv), $K_2CO_3$ (30.2 mg, 0.218 mmol, 0.35 equiv), and THF (1.25 mL) for 4 h at rt. The product imine was obtained as pale-yellow oil (245 mg, 89%). $^1$H NMR (500 MHz, $CDCl_3$) δ 8.01 (d, *J* = 8.9 Hz, 1H), 7.36 (d, *J* = 8.6 Hz, 2H), 7.05 (s, 1H), 6.96 (d, *J* = 8.3 Hz, 1H), 6.92 – 6.88 (m, 4H), 6.79 (d, *J* = 8.7 Hz, 1H), 5.10 (s, 2H), 3.89 (s, 3H), 3.80 (s, 3H), 3.61 (t, *J* = 5.7 Hz, 2H), 3.42 (m, 2H), 1.44 (s, 9H).

・**Key Reaction**

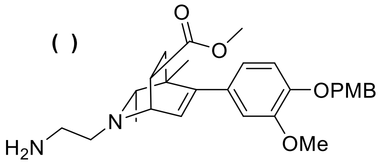

Following General Procedure B for the formation of 1,2-dihydropyridines, the reaction was set up with above obtained imine (350 mg, 0.794 mmol, 1.00 equiv), Rh stock solution (794 µL, 79.4 µmol, 10 mol %), 2-butyne (621 µL, 7.94 mmol, 10.0 equiv), and a reaction time of 16 h at 70 ºC. Isoquinuclidine formation was conducted according to the General Procedure C with methyl acrylate (720 µL, 7.94 mmol, 10.0 equiv) for 16 h at 90 ºC. The product after the Diels-Alder reaction was purified by silica gel chromatography (50% EtOAc/hexanes+1% $Et_3N$) to afford the protected isoquinuclidine as pale-yellow oil (254 mg, 55% yield over two steps). $^1$H NMR (400 MHz, $CDCl_3$) δ 7.33 (d, *J* = 8.4 Hz, 2H), 6.86 (d, *J* = 8.4 Hz, 2H), 6.80 (d, *J* = 8.1 Hz, 1H), 6.67 (d, *J* = 1.9 Hz, 1H), 6.62 (dd, *J* = 8.2, 2.0 Hz, 1H), 6.10 (d, *J* = 6.3 Hz, 1H), 5.03 (s, 2H), 3.84 (s, 3H), 3.77 (s, 3H), 3.74 (m, 1H), 3.62 (s, 3H), 3.33 – 3.24 (m, 1H), 3.22 – 3.15 (m, 2H), 2.70 (m, 5.9 Hz, 2H), 2.13 (s, 3H), 2.08 (q, *J* = 6.5 Hz, 1H), 1.95 (dd, *J* = 13.1, 4.4 Hz, 1H), 1.55 – 1.49 (m, 1H), 1.43 (s, 9H), 1.00 (s, 3H), 0.91 (d, *J* = 6.2 Hz, 3H).

・**Deprotection**

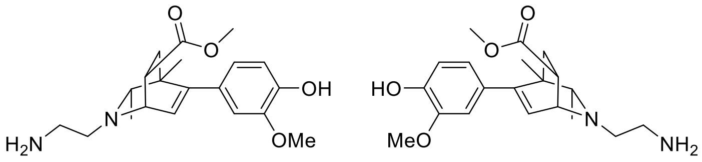

**#33**
enantiomers separated by chiral HPLC

**Final Compound #33** General Procedure D was followed using protected isoquinuclidine (254 mg, 0.437 mmol, 1.00 equiv) in $CH_2Cl_2$ (3.64 mL) and was reacted with trifluoroacetic acid (335 µL, 4.37 mmol, 10.0 equiv) for 10 h. The crude product was purified by silica gel chromatography (10% MeOH/$CH_2Cl_2$ + 1% $NH_4OH$) to yield the racemic isoquinuclidine (48.0 mg, 31% yield) as pale-yellow oil. $^1$H NMR (500 MHz, $CDCl_3$) δ 6.83 (d, *J* = 8.6 Hz, 1H), 6.65 (m, 2H), 6.12 (d, *J* = 6.3 Hz, 1H), 3.87 (s, 3H), 3.76 (dd, *J* = 6.4, 3.1 Hz, 1H), 3.64 (s, 3H), 3.25 – 3.19 (m, 1H), 3.02 (bs, 2H), 2.84 (m, 2H), 2.74 (dt, *J* = 12.2, 6.1 Hz, 1H), 2.65 (dt, *J* = 12.4, 6.3 Hz, 1H), 2.08 (q, *J* = 6.2 Hz, 1H), 1.97 (dd, *J* = 12.8, 4.2 Hz, 1H), 1.54 (dd, *J* = 13.0, 9.4 Hz, 1H), 1.02 (s, 3H), 0.94 (d, *J* = 6.2 Hz, 3H). $^{13}$C NMR (126 MHz, $CDCl_3$) δ 174.8, 148.3, 146.1, 144.9, 132.2, 127.7, 121.3, 114.0, 111.2, 65.1, 57.3, 55.9, 54.5, 51.8, 41.2, 40.6, 38.1, 36.0, 21.7, 20.5. HRMS (ESI+, m/z) $[M+H]^+$ calcd: 361.2122, found: 361.2111

**HPLC Traces**

A portion of this material was separated utilizing semi-preparative chiral HPLC (Chiralpak AD-H column, 250 x 10 mm, 10% EtOH/Hexanes + 0.1% DEA, 2.5 mL/min) to provide pure enantiomers with t = 20 min and 25 min.

Chiralpak AD-H, analytical column (250 x 4.6 mm), 10% EtOH/Hexanes+0.1% DEA, 1 mL/min, 254 nm

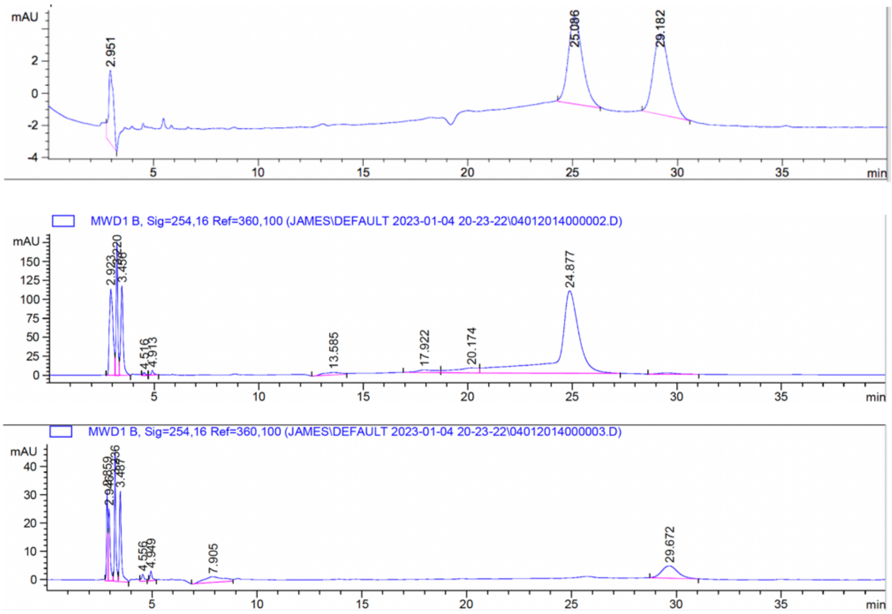

**Synthesis of #39**

・**Synthesis of Imine**

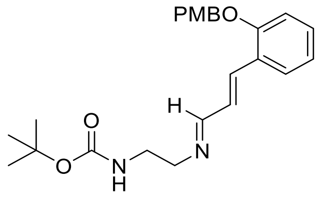

General procedure A was followed using *tert*-butyl *N*-(2-aminoethyl)carbamate (300 mg, 1.86 mmol, 1.0 equiv), (*E*)-3-[2-(4-Methoxy-benzyloxy)-phenyl]-propenal (500 mg, 1.86 mmol, 1.0 equiv), molecular sieves 4 Å, and THF (4 mL). Imine (760 mg, 99% yield) was obtained as a pale-yellow oil. $^1$H NMR (400 MHz, $CDCl_3$) δ 8.01 (d, *J* = 9.0 Hz, 1H), 7.56 (d, *J* = 8.0 Hz, 1H), 7.42 (d, *J* = 16.2 Hz, 1H), 7.34 (d, *J* = 8.6 Hz, 2H), 7.27 (ddd, *J* = 8.6, 7.6, 1.6 Hz, 1H), 6.98 – 6.90 (m, 6H), 5.05 (s, 2H), 3.81 (s, 3H), 3.59 (t, *J* = 5.8 Hz, 2H), 3.41 (q, *J* = 5.8 Hz, 2H), 1.42 (s, 9H).

・**Key Reaction**

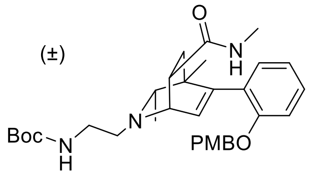

General procedure B was followed using above obtained imine (100 mg, 0.244 mmol, 1.0 equiv) and 2-butyne (95 µL, 1.22 mmol, 5 equiv) in THF (0.25 mL). Rh catalyst (10 mol %, 0.25 mL, 0.025 mmol, 100 mM in THF) was added, and the reaction was carried out at 65 °C for 12 h to produce the DHP intermediate. Following general procedure C, *N*-methyl acrylamide (207 mg, 2.44 mmol, 10 equiv) was added to the solution of DHP intermediate, and the reaction was carried out at 90 °C for 16 h. The resulting crude material was purified by column chromatography (5% $MeOH/CH_2Cl_2$ + 1% $NH_4OH$) to afford the desired product (29 mg, 22% yield from imine) as a pale-yellow oil. $^1$H NMR (400 MHz, $CD_3OD$) δ 7.37 (d, *J* = 8.6 Hz, 2H), 7.24 (ddd, *J* = 7.9, 7.3, 1.8 Hz, 1H), 7.08 (dd, *J* = 7.4, 1.8 Hz, 1H), 7.05 (d, *J* = 8.3 Hz, 1H), 6.98 – 6.84 (m, 3H), 6.10 (d, *J* = 6.4 Hz, 1H), 5.04 (d, *J* = 10.8 Hz, 1H), 4.98 (d, *J* = 10.8 Hz, 1H), 3.77 (s, 3H), 3.64 – 3.54 (m, 1H), 3.21 – 3.04 (m, 4H), 2.74 – 2.65 (m, 2H), 2.50 (s, 3H), 1.69 (dd, *J* = 13.0, 4.8 Hz, 1H), 1.49 (dd, *J* = 13.0, 9.8 Hz, 1H), 1.42 (s, 9H), 0.93 (s, 3H), 0.86 (d, *J* = 6.3 Hz, 3H).

・**Deprotection**

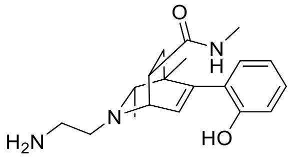

**(±)-#39**

**Final Compound #39** General Procedure D was followed using protected isoquinuclidine (29 mg, 0.053 mmol, 1.00 equiv) in $CH_2Cl_2$ (1.0 mL) and was reacted with trifluoroacetic acid (39 µL, 0.53 mmol, 10.0 equiv) for 12 h. The crude product was purified by preparative thin-layer chromatography (10% MeOH/$CH_2Cl_2$ + 1% $NH_4OH$) to yield the racemic isoquinuclidine (12 mg, 69% yield) as a pale-yellow oil. $^1$H NMR (400 MHz, $CD_3OD$) δ 7.10 (ddd, *J* = 8.5, 7.5, 1.7 Hz, 1H), 6.97 (dd, *J* = 7.5, 1.7 Hz, 1H), 6.87 – 6.73 (m, 2H), 6.14 (d, *J* = 6.4 Hz, 1H), 3.52 (dd, *J* = 6.4, 3.1 Hz, 1H), 3.26 (dt, *J* = 9.3, 3.1 Hz, 1H), 2.86 – 2.70 (m, 4H), 2.68 (s, 3H), 2.16 (q, *J* = 6.5 Hz, 1H), 1.82 (dd, *J* = 13.2, 4.0 Hz, 1H), 1.59 (dd, *J* = 13.2, 9.3 Hz, 1H), 1.00 (d, *J* = 6.2 Hz, 3H), 0.86 (s, 3H). $^{13}$C NMR (101 MHz, $CD_3OD$) δ 176.5, 154.5, 143.7, 130.2, 128.9, 128.1, 126.1, 118.7, 115.3, 64.9, 56.3, 55.1, 48.4, 41.1, 39.6, 36.3, 25.2, 19.6, 19.5. HRMS (ESI+, m/z) calcd for $C_{19}H_{28}N_3O_2$ $(M+H)^+$ : 330.2176, found: 330.2163.

**Synthesis of #53**

・**Synthesis of Amine**

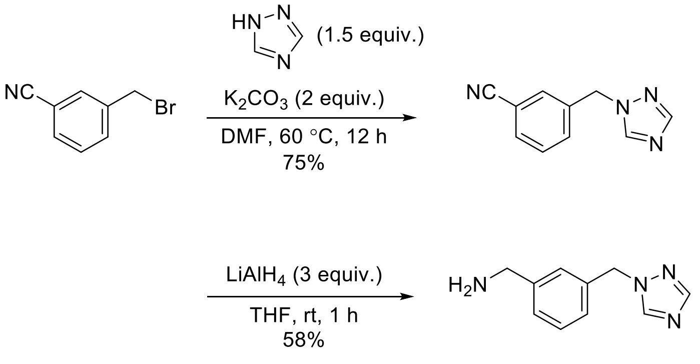

To a flame-dried round-bottom flask was added a magnetic stir bar, 3-(bromomethyl)benzonitrile (1.5 g, 7.65 mmol, 1 equiv) and 1H-1,2,4-triazole (793 mg, 11.5 mmol, 1.5 equiv) dissolved in DMF (15 mL). To this solution was then added potassium carbonate (2.1 g, 15.3 mmol, 2 equiv). The reaction mixture was stirred at 60 °C for 12 h. After the reaction was complete as monitored by thin-layer chromatography, the mixture was then diluted with water, extracted with ethyl acetate, washed with brine, dried with $Na_2SO_4$ and filtered. The organic layer was concentrated and the crude residue was purified via silica gel chromatography (gradient of 0-100% EtOAc in hexane +1% $Et_3N$) to give 1000 mg of the 3-((1H-1,2,4-triazol-1-yl)methyl)benzonitrile in 75% yield. To a stirred solution of the above obtained compound (500 mg, 2.71 mmol, 1 equiv) in THF (27 mL) was added $LiAlH_4$ (309 mg, 8.14

mmol, 3 equiv), and the reaction mixture was stirred at room temperature for 1 h. After the reaction was complete as monitored by thin-layer chromatography, the mixture was then quenched by careful addition of 1N NaOH and then $H_2O$, extracted with ethyl acetate, washed with brine, dried with $Na_2SO_4$ and filtered. The organic layer was concentrated and the crude residue was purified via silica gel chromatography (5% MeOH/$CH_2Cl_2$ + 1% $NH_4OH$) to give 295 mg of the amine in 58% yield. $^1$H NMR (400 MHz, $_3$) δ 8.04 (s, 1H), 7.95 (s, 1H), 7.36 – 7.26 (m, 2H), 7.22 (s, 1H), 7.13 (d, *J* = 7.2 Hz, 1H), 5.32 (s, 2H), 3.86 (s, 2H).

・**Synthesis of Imine**

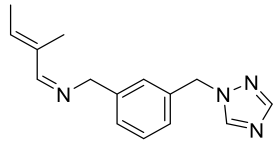

General procedure A was followed using above obtained amine (295 mg, 1.57 mmol, 1.0 equiv), *trans*-2-methyl-2-butenal (264 mg, 3.13 mmol, 2.0 equiv), molecular sieves 4 Å, and THF (3 mL). Imine (350 mg, 88% yield) was obtained as a pale-yellow oil. $^1$H NMR (400 MHz, $CDCl_3$) δ 8.02 (s, 1H), 7.94 (s, 1H), 7.89 (s, 1H), 7.31 (dd, *J* = 8.0, 7.6 Hz, 1H), 7.25 (d, 8.0 Hz, 1H), 7.17 (s, 1H), 7.11 (d, *J* = 7.6 Hz, 1H), 5.99 (q, *J* = 6.3 Hz, 1H), 5.31 (s, 2H), 4.63 (s, 2H), 1.85 (s, 3H), 1.84 (d, *J* = 6.3 Hz, 3 H).

・**Key Reaction**

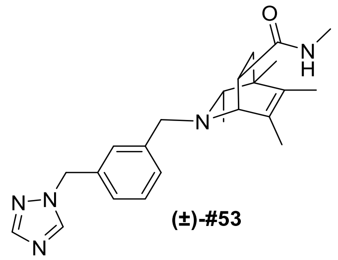

**(±)-#53**

**Final Compound #53** General procedure B was followed using above obtained imine (170 mg, 0.668 mmol, 1.0 equiv) and 2-butyne (262 µL, 3.34 mmol, 5 equiv) in THF (0.67 mL). Rh catalyst (10 mol %, 0.67 mL, 0.067 mmol, 100 mM in THF) was added, and the reaction was carried out at 65 °C for 12 h to produce the DHP intermediate. Following general procedure C, *N*-methyl acrylamide (569 mg, 6.68 mmol, 10 equiv) was added to the solution of DHP intermediate, and the reaction was carried out at 90 °C for 16 h. The resulting crude material was purified by column chromatography (10% MeOH/$CH_2Cl_2$ + 1% $NH_4OH$) to afford the desired product (24 mg, 9% yield from imine) as a pale-yellow oil. $^1$H NMR (400 MHz, $CD_3OD$) δ 8.52 (s, 1H), 7.97 (s, 1H), 7.37 – 7.34 (m, 2H), 7.30 (dd, *J* = 8.0, 7.7 Hz, 1H), 7.20 (d, *J* = 7.7 Hz, 1H), 5.40 (s, 2H), 3.70 (s, 2H), 3.15 – 3.06 (m, 2H), 2.63 (s, 3H), 2.13 (q, *J* = 6.2 Hz, 1H), 1.65 (s, 3H), 1.64 – 1.56 (m, 4H), 1.43 (dd, *J* = 12.7, 9.4 Hz, 1H), 1.04

(s, 3H), 0.56 (d, $J$ = 6.2 Hz, 3H). $^{13}$C NMR (101 MHz, $CD_3OD$) δ 175.5, 150.9, 143.6, 140.7, 135.3, 132.2, 132.1, 129.0, 128.6, 128.4, 126.5, 64.2, 60.5, 56.9, 52.8, 40.1, 37.8, 36.2, 25.1, 19.4, 18.1, 15.7, 12.4. HRMS (ESI+, m/z) calcd for $C_{23}H_{31}N_5O$ $(M+H)^+$ : 394.2601, found: 394.2582.

**Synthesis of #56**

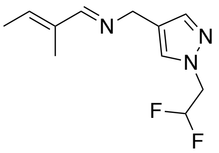

A modified version of General Procedure A was followed using the HCl salt of (1-(2,2-difluoroethyl)-1*H*-pyrazol-4-yl)methanamine (100 mg, 0.506 mmol, 1.00 equiv), (*E*)-2-methylbut-2-enal (244 µL, 2.53 mmol, 5.00 equiv), 3 Å MS (253 mg), and EtOH (1.01 mL) for 7 h at rt. The product imine was obtained as pale-yellow oil (115 mg, 100%). $^1$H NMR (500 MHz, $CDCl_3$) δ 7.87 (s, 1H), 7.45 (s, 1H), 7.34 (s, 1H), 6.17 – 5.92 (m, 2H), 4.53 (s, 2H), 4.40 (td, $J$ = 13.6, 4.3 Hz, 2H), 1.84 (m, 6H).

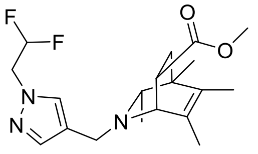

**(±)-#56**

**Final Compound #56** Following General Procedure B for the formation of 1,2-dihydropyridines, the reaction was set up with above obtained imine (120 mg, 0.528 mmol, 1.0 equiv), Rh stock solution (528 µL, 52.8 µmol, 10 mol%), 2-butyne (413 µL, 5.28 mmol, 10.0 equiv), and a reaction time of 20 h at 70 ºC. Isoquinuclidine formation was conducted according to the General Procedure C with methyl acrylate (478 µL, 5.28 mmol, 10.0 equiv) for 16 h at 70 ºC. The product after the Diels-Alder reaction was purified by silica gel chromatography (10% MeOH/$CH_2Cl_2$+1% $NH_4OH$) to afford the isoquinuclidine product as pale-yellow oil (15.5 mg, 8% yield over two steps). $^1$H NMR (500 MHz, $CDCl_3$) δ 7.54 (s, 1H), 7.44 (s, 1H), 6.06 (tt, $J$ = 55.5, 4.4 Hz, 1H), 4.43 (td, $J$ = 13.6, 4.4 Hz, 2H), 3.61 (m, 4H), 3.47 (d, $J$ = 13.4 Hz, 1H), 3.39 (s, 1H), 3.13 (m, 1H), 2.02 (m, 1H), 1.77 – 1.73 (m, 1H), 1.64 (m, 6H), 1.44 – 1.36 (m, 1H), 1.06 (s, 3H), 0.76 (d, $J$ = 6.2 Hz, 3H). $^{13}$C NMR (126 MHz, $CDCl_3$) δ 159.7, 148.0, 141.3, 130.4, 130.4, 113.4 (t, $J$ = 243.6 Hz), 64.8, 58.2, 54.0 (t, $J$ = 28.1 Hz), 51.6, 46.7, 40.3, 36.3, 20.4, 19.0, 16.6, 13.7. $^{19}$F NMR (471 MHz, $CDCl_3$) δ -122.28 (m), -122.40 (m). HRMS (ESI+, m/z) $[M+H]^+$ calcd: 368.2144, found: 368.2135

**Synthesis of #57**

・**Synthesis of Amine**

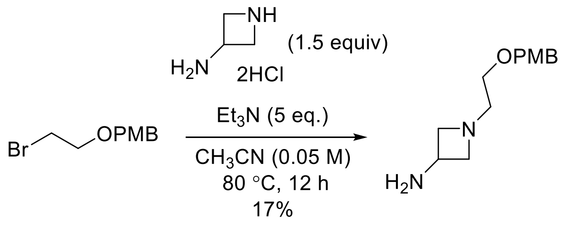

To a flame-dried round-bottom flask was added a magnetic stir bar, 1-[(2-bromoethoxy)methyl]-4-methoxy-benzene (750 mg, 3.06 mmol, 1 equiv) and azetidin-3-amine dihydrochloride (666 mg, 4.59 mmol, 1.5 equiv) dissolved in $CH_3CN$ (60 mL). To this solution was then added triethylamine (2.13 mL, 15.3 mmol, 5 equiv). The reaction mixture was stirred at 80 °C for 12 h. After the reaction was complete as monitored by thin-layer chromatography, the mixture was concentrated and the crude residue was purified via silica gel chromatography (5% MeOH/$CH_2Cl_2$ + 1% $NH_4OH$) to give 120 mg of the desired amine in 17% yield. $^1$H NMR (400 MHz, $CD_3OD$) δ 7.23 (d, $J$ = 8.5 Hz, 2H), 6.86 (d, $J$ = 8.5 Hz, 2H), 4.39 (s, 2H), 3.76 (s, 3H), 3.62 (t, $J$ = 6.8 Hz, 2H), 3.51 (p, $J$ = 6.9 Hz, 1H), 3.44 (t, $J$ = 5.5 Hz, 2H), 2.82 (t, $J$ = 6.8 Hz, 2H), 2.65 (t, $J$ = 5.5 Hz, 2H).

・**Synthesis of Imine**

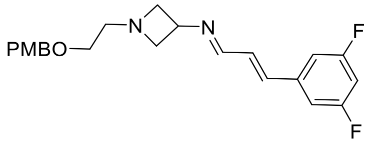

General procedure A was followed using above obtained amine (80 mg, 0.339 mmol, 1.0 equiv), (*E*)-3-(3,5-difluorophenyl)acrylaldehyde (56.9 mg, 0.339 mmol, 1.0 equiv), molecular sieves 4 Å, and THF (1 mL). Imine (130 mg, 99% yield) was obtained as a pale-yellow oil. $^1$H NMR (400 MHz, $CD_3OD$) δ 8.01 (dd, $J$ = 8.9, 1.0 Hz, 1H), 7.24 (d, $J$ = 8.6 Hz, 2H), 7.21 – 7.15 (m, 2H), 7.11 (d, $J$ = 16.0 Hz, 1H), 6.97 – 6.88 (m, 2H), 6.86 (d, $J$ = 8.7 Hz, 2H), 4.41 (s, 2H), 4.23 (p, $J$ = 6.9 Hz, 1H), 3.76 (s, 3H), 3.73 (td, $J$ = 6.8, 1.9 Hz, 2H), 3.49 (t, $J$ = 5.4 Hz, 2H), 3.22 (td, $J$ = 6.9, 1.9 Hz, 2H), 2.73 (t, $J$ = 5.4 Hz, 2H).

・**Key Reaction**

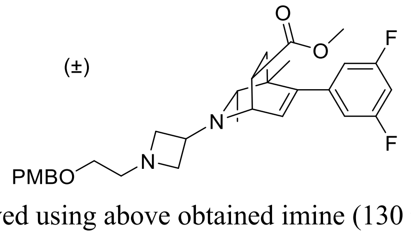

General procedure B was followed using above obtained imine (130 mg, 0.336 mmol, 1.0 equiv) and 2-butyne (132 μL, 1.68 mmol, 5 equiv) in THF (0.34 mL). Rh catalyst (10 mol %, 0.34 mL, 0.034 mmol, 100 mM in THF) was added, and the reaction was carried out at 65 °C for 12 h to produce the

DHP intermediate. Following general procedure C, methyl acrylate (0.3 mL, 3.36 mmol, 10 equiv) was added to the solution of DHP intermediate, and the reaction was carried out at 50 °C for 12 h. The resulting crude material was purified by column chromatography (30% *t*BuOMe/Pentane+1% $Et_3N$) to afford the desired product (140 mg, 79% yield from imine) as a pale-yellow oil. $^1$H NMR (400 MHz, $CD_3OD$) δ 7.25 (d, *J* = 8.6 Hz, 2H), 6.90 – 6.81 (m, 3H), 6.72 – 6.69 (m, 2H), 6.17 (d, *J* = 6.4 Hz, 1H), 4.42 (s, 2H), 3.76 (s, 3H), 3.67 (dd, *J* = 6.4, 3.2 Hz, 1H), 3.63 (s, 3H), 3.58 (t, *J* = 6.9 Hz, 2H), 3.53 – 3.43 (m, 3H), 3.16 (dt, *J* = 9.3, 3.8 Hz, 1H), 3.07 (t, *J* = 7.3 Hz, 1H), 3.00 (t, *J* = 7.7 Hz, 1H), 2.71 (t, *J* = 5.4 Hz, 2H), 2.23 (q, *J* = 6.3 Hz, 1H), 1.91 (dd, *J* = 13.2, 4.1 Hz, 1H), 1.60 (dd, *J* = 13.2, 9.4 Hz, 1H), 1.00 (s, 3H), 0.91 (d, *J* = 6.3 Hz, 3H).

・**Deprotection**

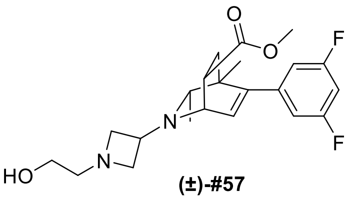

**(±)-#57**

**Final Compound #57** General Procedure D was followed using protected isoquinuclidine (90 mg, 0.171 mmol, 1.00 equiv) in $CH_2Cl_2$ (1.0 mL) and was reacted with trifluoroacetic acid (127 µL, 1.71 mmol, 10.0 equiv) for 12 h. The crude product was purified by preparative thin-layer chromatography (10% MeOH/$CH_2Cl_2$ + 1% $NH_4OH$) to yield the racemic isoquinuclidine (69 mg, 99% yield) as a pale-yellow oil. $^1$H NMR (400 MHz, cd$_3$od) δ 6.85 (tt, *J* = 9.3, 2.4 Hz, 1H), 6.71 (dd, *J* = 8.3, 2.1 Hz, 2H), 6.18 (d, *J* = 6.4 Hz, 1H), 3.69 (dd, *J* = 6.4, 3.2 Hz, 1H), 3.63 (s, 3H), 3.63 – 3.59 (m, 2H), 3.56 (t, *J* = 5.8 Hz, 2H), 3.49 (p, *J* = 7.1 Hz, 1H), 3.18 (dt, *J* = 9.3, 3.8 Hz, 1H), 3.07 (t, *J* = 7.3 Hz, 1H), 3.00 (t, *J* = 7.7 Hz, 1H), 2.64 (t, *J* = 5.8 Hz, 2H), 2.24 (q, *J* = 6.3 Hz, 1H), 1.91 (dd, *J* = 13.2, 4.1 Hz, 1H), 1.61 (dd, *J* = 13.2, 9.4 Hz, 1H), 1.01 (s, 3H), 0.92 (d, *J* = 6.3 Hz, 3H). $^{13}$C NMR (101 MHz, $CD_3OD$) δ 174.4, 162.6 (dd, *J* = 248.5, 13.1 Hz), 146.9, 143.3 (t, *J* = 9.1 Hz), 129.2, 110.8 (dd, *J* = 19.2, 7.1 Hz), 101.9 (t, *J* = 26.3 Hz), 62.8, 60.9, 60.7, 59.7, 59.5, 53.1, 51.8, 51.1, 40.8, 38.3, 35.1, 20.0, 19.5. $^{19}$F NMR (376 MHz, $CD_3OD$) δ -112.1. HRMS (ESI+, m/z) calcd for $C_{22}H_{28}F_2N_2O_3$ $(M+H)^+$ : 407.2141, found: 407.2124.

**Synthesis of #58**

・**Synthesis of Imine**

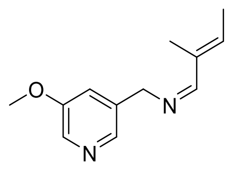

General procedure A was followed using (5-methoxy-3-pyridyl)methanamine (300 mg, 2.17 mmol, 1.0 equiv), *trans*-2-methyl-2-butenal (365 mg, 4.34 mmol, 2.0 equiv), molecular sieves 4 Å, and THF (5 mL). Imine (380 mg, 86% yield) was obtained as a pale-yellow oil. $^1$H NMR (400 MHz, $CDCl_3$) δ 8.17 (d, *J* = 2.8 Hz, 1H), 8.13 (s, 1H), 7.92 (s, 1H), 7.13 (s, 1H), 6.01 (q, *J* = 6.6 Hz, 1H), 4.63 (s, 2H), 3.83 (s, 3H), 1.85 (s, 3H), 1.84 (d, *J* = 7.7 Hz, 3H).

・**Key Reaction**

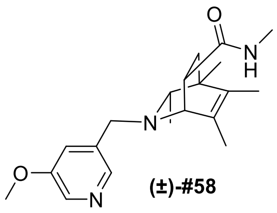

**(±)-#58**

**Final Compound #58** General procedure B was followed using above obtained imine (180 mg, 0.881 mmol, 1.0 equiv) and 2-butyne (345 µL, 4.41 mmol, 5 equiv) in THF (0.88 mL). Rh catalyst (10 mol %, 0.88 mL, 0.088 mmol, 100 mM in THF) was added, and the reaction was carried out at 65 °C for 12 h to produce the DHP intermediate. Following general procedure C, *N*-methyl acrylamide (750 mg, 8.81 mmol, 10 equiv) was added to the solution of DHP intermediate, and the reaction was carried out at 90 °C for 16 h. The resulting crude material was purified by column chromatography (10% $MeOH/CH_2Cl_2$ + 1% $NH_4OH$) to afford the desired product (70 mg, 23% yield from imine) as a white solid. $^1$H NMR (500 MHz, $CD_3OD$) δ 8.14 (s, 1H), 8.11 (s, 1H), 7.47 (s, 1H), 3.87 (s, 3H), 3.79 (d, *J* = 14.0 Hz, 1H), 3.75 (d, J = 14.0 Hz 1H), 3.18 – 3.07 (m, 2H), 2.65 (s, 3H), 2.18 (q, *J* = 6.2 Hz, 1H), 1.68 (s, 3H), 1.68 – 1.62 (m, 4H), 1.48 (dd, *J* = 12.7, 9.2 Hz, 1H), 1.08 (s, 3H), 0.66 (d, *J* = 6.2 Hz, 3H). $^{13}$C NMR (126 MHz, $CD_3OD$) δ 175.4, 156.3, 141.3, 137.4, 135.3, 132.3, 131.9, 121.6, 64.3, 60.5, 54.8, 53.9, 40.2, 37.9, 36.3, 25.1, 19.4, 18.2, 15.7, 12.4. HRMS (ESI+, m/z) calcd for $C_{20}H_{29}N_3O_2$ $(M+H)^+$ : 344.2333, found: 344.2316.

**Synthesis of #61**

・**Synthesis of Imine**

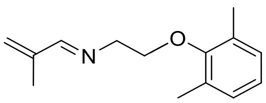

General procedure A was followed using [2-(2,6-dimethylphenoxy)ethyl]amine (250 mg, 1.39 mmol, 1.0 equiv), 2-methylprop-2-enal (486 mg, 6.93 mmol, 5.0 equiv), molecular sieves 4 Å, and THF (3 mL). Imine (310 mg, 96% yield) was obtained as a pale-yellow oil. $^{1}$H NMR (400 MHz, cdcl$_3$) δ 8.03 (s, 1H), 6.97 (d, *J* = 7.4 Hz, 2H), 6.89 (dd, *J* = 8.3, 6.6 Hz, 1H), 5.61 (s, 1H), 5.40 (s, 1H), 4.00 (t, *J* = 5.4 Hz, 2H), 3.87 (t, *J* = 5.4 Hz, 2H), 2.24 (s, 6H), 1.95 (s, 3H).

・**Key Reaction**

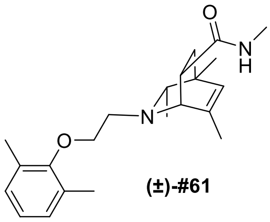

(±)-#61

**Final Compound #61** General procedure B was followed using above obtained imine (200 mg, 0.920 mmol, 1.0 equiv) and 2-butyne (361 µL, 4.60 mmol, 5 equiv) in THF (0.92 mL). Rh catalyst (10 mol %, 0.92 mL, 0.092 mmol, 100 mM in THF) was added, and the reaction was carried out at 65 °C for 12 h to produce the DHP intermediate. Following general procedure C, *N*-methyl acrylamide (783 mg, 9.20 mmol, 10 equiv) was added to the solution of DHP intermediate, and the reaction was carried out at 90 °C for 16 h. The resulting crude material was purified by column chromatography (10% MeOH/$CH_2Cl_2$ + 1% $NH_4OH$) to afford the desired product (25 mg, 8% yield from imine) as a pale-yellow oil. $^{1}$H NMR (400 MHz, $CD_3OD$) δ 6.97 (d, *J* = 7.4 Hz, 2H), 6.87 (dd, *J* = 8.2, 6.8 Hz, 1H), 5.51 (s, 1H), 3.93 (dt, *J* = 9.6, 6.3 Hz, 1H), 3.85 (dt, *J* = 9.6, 6.2 Hz, 1H), 3.44 (dd, *J* = 2.9, 1.4 Hz, 1H), 3.16 (ddd, *J* = 9.3, 4.9, 2.9 Hz, 1H), 3.09 (dt, *J* = 12.8, 6.3 Hz, 1H), 2.98 (dt, *J* = 12.9, 6.2 Hz, 1H), 2.65 (s, 3H), 2.27 (s, 6H), 2.24 (d, *J* = 2.7 Hz, 1H), 2.09 (q, *J* = 6.4 Hz, 1H), 1.76 (d, *J* = 1.7 Hz, 3H), 1.61 (dd, *J* = 12.6, 4.9 Hz, 1H), 1.44 (dd, *J* = 12.6, 9.4 Hz, 1H), 1.07 (s, 3H), 0.84 (d, *J* = 6.2 Hz, 3H). $^{13}$C NMR (101 MHz, $CD_3OD$) δ 175.2, 155.9, 140.4, 130.4, 128.9, 128.5, 123.6, 71.2, 64.8, 60.6, 53.9, 38.2, 37.7, 35.9, 25.1, 21.1, 19.1, 18.7, 15.3. HRMS (ESI+, m/z) calcd for $C_{22}H_{33}N_2O_2$ $(M+H)^+$ : 357.2537, found: 357.2541.

**Synthesis of #64**

・**Synthesis of Imine**

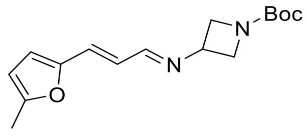

General procedure A was followed using *tert*-butyl-3-aminoazetidine-1-carboxylate (190 mg, 1.10 mmol, 1.0 equiv), (*E*)-3-(5-methyl-2-furyl)-2-propenal (150 mg, 1.10 mmol, 1.0 equiv), molecular sieves 4 Å, and THF (2 mL). Imine (320 mg, 99% yield) was obtained as a pale-yellow oil. [1]H NMR (500 MHz, $CDCl_3$) δ 7.86 (d, *J* = 7.7 Hz, 1H), 6.80 – 6.57 (m, 2H), 6.40 (d, *J* = 3.2 Hz, 1H), 6.04 (d, *J* = 3.2 Hz, 1H), 4.28 – 4.10 (m, 3H), 3.98 (d, *J* = 4.8 Hz, 2H), 2.33 (s, 3H), 1.44 (s, 9H).

・**Key Reaction**

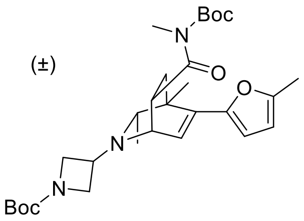

General procedure B was followed using above obtained imine (300 mg, 1.03 mmol, 1.0 equiv) and 2-butyne (405 µL, 5.17 mmol, 5 equiv) in THF (1.0 mL). Rh catalyst (20 mol %, 2.06 mL, 0.0206 mmol, 100 mM in THF) was added, and the reaction was carried out at 75 °C for 12 h to produce the DHP intermediate. Following general procedure C, Boc-protected *N*-methyl acrylamide (956 mg, 5.17 mmol, 5 equiv) was added to the solution of DHP intermediate, and the reaction was carried out at 50 °C for 12 h. The resulting crude material was purified by column chromatography (30% *t*BuOMe/Pentane+1% $Et_3N$) to afford the desired product (125 mg, 23% yield from imine) as a pale-yellow oil. [1]H NMR (400 MHz, $CD_3OD$) δ 6.34 (d, *J* = 6.6 Hz, 1H), 6.26 (d, *J* = 3.2 Hz, 1H), 5.95 (dd, *J* = 3.2, 1.3 Hz, 1H), 4.33 (dt, *J* = 9.3, 4.3 Hz, 1H), 4.16 (t, *J* = 7.2 Hz, 1H), 4.01 (t, *J* = 7.6 Hz, 1H), 3.93 (t, *J* = 5.6 Hz, 1H), 3.82 – 3.75 (m, 2H), 3.71 (dd, *J* = 6.8, 2.8 Hz, 1H), 3.04 (s, 3H), 2.27 – 2.22 (m, 4H), 2.01 (dd, *J* = 16.6, 4.3 Hz, 1H), 1.55 (s, 9H), 1.52 – 1.47 (m, 1H), 1.42 (s, 9H), 1.35 (s, 3H), 0.92 (d, *J* = 6.2 Hz, 3H).

・**Deprotection**

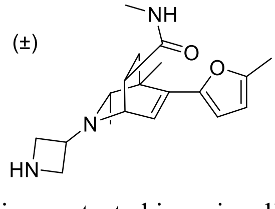

General Procedure D was followed using protected isoquinuclidine (80 mg, 0.151 mmol, 1.00 equiv) in $CH_2Cl_2$ (1.5 mL) and was reacted with trifluoroacetic acid (224 µL, 3.02 mmol, 20.0 equiv) for 12 h. The crude product was purified by preparative thin-layer chromatography (10% MeOH/$CH_2Cl_2$ + 1% $NH_4OH$) to yield the deprotected isoquinuclidine (33 mg, 51% yield) as a pale-yellow oil. [1]H NMR (400 MHz, $CD_3OD$) δ 6.47 (d, *J* = 6.6 Hz, 1H), 6.27 (d, *J* = 3.3 Hz, 1H), 5.95 (dd, *J* = 3.3, 1.1 Hz, 1H), 3.90 – 3.48 (m, 6H), 2.93 (dt, *J* = 9.4, 4.4 Hz, 1H), 2.64 (s, 3H), 2.29 – 2.17 (m, 4H), 1.80 (dd, *J* = 13.1, 4.4 Hz, 1H), 1.52 (dd, *J* = 13.1, 9.4 Hz, 1H), 1.34 (s, 3H), 0.85 (d, *J* = 6.2 Hz, 3H).

・**Addition to 3-chloro pyridazine**

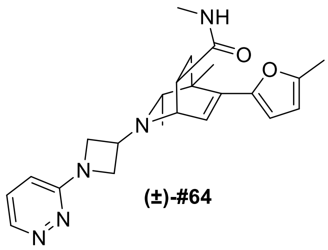

(±)-#64

**Final Compound #64** To a flame-dried round-bottom flask was added a magnetic stir bar, above obtained isoquinuclidine (33 mg, 0.1 mmol, 1.0 equiv) and 3-chloropyridazine (57 mg, 0.5 mmol, 5.0 equiv) dissolved in EtOH (1 mL). To this solution was then added *N*,*N*-diisopropylethylamine (174 μL, 1.0 mmol, 10 equiv). The reaction mixture was stirred at 90 °C for 12 h. After the reaction was complete as monitored by thin-layer chromatography, the mixture was concentrated and the crude residue was purified by preparative thin-layer chromatography (10% MeOH/$CH_2Cl_2$ + 1% $NH_4OH$) to give the racemic isoquinuclidine (21 mg, 51% yield) as a pale-yellow oil. $^1$H NMR (400 MHz, $CD_3OD$) δ 8.46 (dd, *J* = 4.5, 1.3 Hz, 1H), 7.39 (dd, *J* = 9.1, 4.5 Hz, 1H), 6.87 (dd, *J* = 9.1, 1.4 Hz, 1H), 6.49 (d, *J* = 6.6 Hz, 1H), 6.28 (d, *J* = 3.3 Hz, 1H), 5.96 (dd, *J* = 3.3, 1.1 Hz, 1H), 4.33 – 4.20 (m, 2H), 4.10 – 3.94 (m, 3H), 3.73 (dd, *J* = 6.6, 3.0 Hz, 1H), 3.03 (dt, *J* = 9.6, 4.0 Hz, 1H), 2.67 (s, 3H), 2.36 (q, *J* = 6.3 Hz, 1H), 2.24 (s, 3H), 1.87 (dd, *J* = 13.2, 4.4 Hz, 1H), 1.57 (dd, *J* = 13.1, 9.6 Hz, 1H), 1.37 (s, 3H), 0.94 (d, *J* = 6.3 Hz, 3H). $^{13}$C NMR (101 MHz, $CD_3OD$) δ 174.7, 160.9, 151.3, 150.5, 142.6, 136.7, 128.1, 124.6, 113.0, 108.5, 106.7, 62.3, 55.6, 55.5, 52.6, 52.4, 40.4, 39.1, 35.3, 25.2, 20.9, 19.3, 11.9. HRMS (ESI+, m/z) calcd for $C_{23}H_{29}N_5O_2$ $(M+H)^+$ : 408.2394, found: 408.2375.

### 5. Analogs Based upon Screening Hits (#001-#036): Synthesis Procedures and Analytical Data

**Synthesis of #001**

・**Synthesis of 0.8 M imine solution**

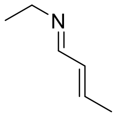

To a flame-dried vial was added a magnetic stir bar, *trans*-crotonaldehyde (0.6 mL, 7.3 mmol, 1.0 equiv) in dry THF (1.2 mL), and molecular sieves 4 Å. To this solution was then slowly added 2.0 M ethylamine in THF (7.3 mL, 14.6 mmol, 2.0 equiv) at 0 °C and the reaction solution was stirred at 0 °C for 2 h. Upon reaction completion, the 0.8 M imine solution was stored at –20 °C in a nitrogen filled glovebox and taken on to the next step without further purification.

・**Key Reaction**

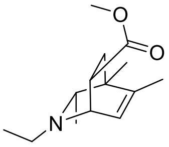

(±)-#001

**Final Compound #001** General procedure B was followed using above obtained 0.8 M solution of imine (2 mL, 1.6 mmol, 1.0 equiv) and 2-butyne (625 µL, 8.0 mmol, 5 equiv) in THF (1.6 mL). Rh catalyst (10 mol %, 1.6 mL, 0.16 mmol, 100 mM in THF) was added, and the reaction was carried out at 65 °C for 12 h to produce the DHP intermediate. Following general procedure C, methyl acrylate (1.44 mL, 16.0 mmol, 10 equiv) was added to the solution of DHP intermediate, and the reaction was carried out at 50 °C for 12 h. The resulting crude material was purified by column chromatography (30% *t*BuOMe/Pentane+1% $Et_3N$) to afford the desired product (50 mg, 13% yield from imine) as a pale-yellow oil. $^1$H NMR (400 MHz, $CD_3OD$) δ 5.95 (dd, *J* = 6.3, 1.8 Hz, 1H), 3.67 (dd, *J* = 6.3, 3.1 Hz, 1H), 3.60 (s, 3H), 3.15 (ddd, *J* = 9.5, 4.5, 3.1 Hz, 1H), 2.69 (dq, *J* = 12.2, 7.3 Hz, 1H), 2.54 (dq, *J* = 12.2, 7.3 Hz, 1H), 2.03 (q, *J* = 6.4 Hz, 1H), 1.74 (d, *J* = 1.7 Hz, 3H), 1.69 (dd, *J* = 13.0, 4.5 Hz, 1H), 1.44 (dd, *J* = 13.0, 9.5 Hz, 1H), 1.12 (t, *J* = 7.2 Hz, 3H), 1.09 (s, 3H), 0.84 (d, *J* = 6.4 Hz, 3H). $^{13}$C NMR (101 MHz, $CD_3OD$) δ 174.9, 144.4, 124.3, 64.3, 53.4, 50.9, 48.2, 39.9, 36.6, 34.5, 18.7, 18.5, 17.5, 12.7. HRMS (ESI+, m/z) calcd for $C_{14}H_{24}NO_2$ $(M+H)^+$ : 238.1802, found: 238.1798.

**Synthesis of #002**

・**Synthesis of Imine**

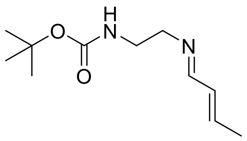

General procedure A was followed using *tert*-butyl *N*-(2-aminoethyl)carbamate (500 mg, 3.12 mmol, 1.0 equiv), *trans*-crotonaldehyde (1.29 mL, 15.6 mmol, 5.0 equiv), molecular sieves 4 Å, and THF (6 mL). Imine (662 mg, 99% yield) was obtained as a pale-yellow oil. $^1$H NMR (400 MHz, $CDCl_3$) δ 7.82 (d, $J$ = 6.9 Hz, 1H), 6.28 – 6.12 (m, 2H), 4.77 (br, 1H), 3.50 (t, $J$ = 5.8 Hz, 2H), 3.35 (q, $J$ = 5.9 Hz, 2H), 1.87 (d, $J$ = 5.1 Hz, 3H), 1.42 (s, 9H).

・**Key Reaction**

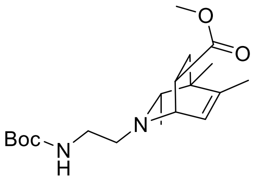

General procedure B was followed using above obtained imine (50 mg, 0.236 mmol, 1.0 equiv) and 2-butyne (93 µL, 1.18 mmol, 5 equiv) in THF (0.24 mL). Rh catalyst (10 mol %, 0.24 mL, 0.024 mmol, 100 mM in THF) was added, and the reaction was carried out at 65 °C for 12 h to produce the DHP intermediate. Following general procedure C, methyl acrylate (212 µL, 2.36 mmol, 10 equiv) was added to the solution of DHP intermediate, and the reaction was carried out at 50 °C for 16 h. The resulting crude material was purified by column chromatography (30% *t*BuOMe/Pentane+1% $Et_3N$) to afford the desired product (45 mg, 54% yield from imine) as a pale-yellow oil. $^1$H NMR (400 MHz, $CDCl_3$) δ 5.94 (dd, $J$ = 6.4, 1.9 Hz, 1H), 4.97 (s, 1H), 3.58 (s, 3H), 3.54 (dd, $J$ = 6.4, 3.1 Hz, 1H), 3.27 – 3.07 (m, 2H), 3.02 (ddd, $J$ = 9.5, 4.6, 3.1 Hz, 1H), 2.70 – 2.51 (m, 2H), 1.95 (q, $J$ = 6.2 Hz, 1H), 1.76 – 1.64 (m, 4H), 1.46 – 1.31 (m, 10H), 1.03 (s, 3H), 0.76 (d, $J$ = 6.2 Hz, 3H).

・**Deprotection**

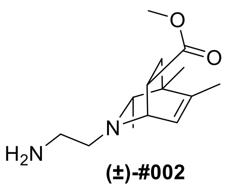

**(±)-#002**

**Final Compound #002** General Procedure D was followed using protected isoquinuclidine (45 mg, 0.128 mmol, 1.0 equiv) in $CH_2Cl_2$ (1.3 mL) and was reacted with trifluoroacetic acid (95 µL, 1.28 mmol, 10.0 equiv) for 12 h. The crude product was purified by preparative thin-layer chromatography (10% MeOH/$CH_2Cl_2$ + 1% $NH_4OH$) to yield the racemic isoquinuclidine (25 mg, 78% yield) as a pale-yellow oil. $^1$H NMR (400 MHz, $CD_3OD$) δ 5.95 (dd, $J$ = 6.3, 1.8 Hz, 1H), 3.66 – 3.52 (m, 4H),

3.16 (ddd, *J* = 9.5, 4.6, 3.0 Hz, 1H), 2.83 – 2.56 (m, 4H), 2.05 (q, *J* = 6.3 Hz, 1H), 1.73 (d, *J* = 1.8 Hz, 3H), 1.70 (dd, *J* = 13.0, 4.6 Hz, 1H), 1.45 (dd, *J* = 13.0, 9.5 Hz, 1H), 1.09 (s, 3H), 0.81 (d, *J* = 6.3 Hz, 3H). $^{13}$C NMR (101 MHz, $CD_3OD$) δ 175.0, 143.7, 124.5, 64.5, 55.6, 54.2, 50.8, 40.0, 39.3, 37.4, 34.6, 18.8, 18.7, 17.5. HRMS (ESI+, m/z) calcd for $C_{14}H_{25}N_2O_2$ $(M+H)^+$ : 253.1911, found: 253.1913.

**Synthesis of #003**

・**Key Reaction**

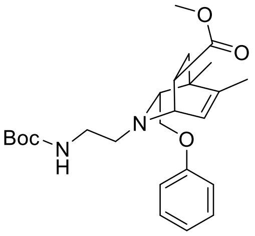

General procedure B was followed using above obtained imine (150 mg, 0.707 mmol, 1.0 equiv) and 2-butyne (277 µL, 3.53 mmol, 5 equiv) in THF (0.71 mL). Rh catalyst (10 mol %, 0.71 mL, 0.071 mmol, 100 mM in THF) was added, and the reaction was carried out at 65 °C for 12 h to produce the DHP intermediate. Following general procedure C, methyl acrylate (636 µL, 7.07 mmol, 10 equiv) was added to the solution of DHP intermediate, and the reaction was carried out at 50 °C for 16 h. The resulting crude material was purified by column chromatography (30% *t*BuOMe/Pentane+1% $Et_3N$) to afford the desired product (70 mg, 22% yield from imine) as a pale-yellow oil. $^1$H NMR (400 MHz, $CD_3OD$) δ 7.28 – 7.15 (m, 2H), 6.94 – 6.78 (m, 3H), 5.97 (dd, *J* = 6.3, 1.8 Hz, 1H), 3.73 (dd, *J* = 9.8, 5.5 Hz, 1H), 3.68 (dd, *J* = 6.4, 3.1 Hz, 1H), 3.61 (s, 3H), 3.57 – 3.49 (m, 1H), 3.25 – 3.13 (m, 3H), 2.81 – 2.70 (m, 2H), 2.39 (t, *J* = 5.3 Hz, 1H), 1.79 – 1.67 (m, 4H), 1.55 (dd, *J* = 13.1, 9.5 Hz, 1H), 1.46 – 1.32 (m, 10H), 1.20 (s, 3H).

・**Deprotection**

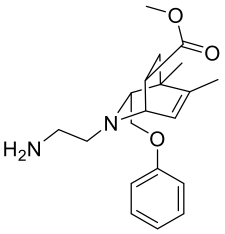

**(±)-#003**

**Final Compound #003** General Procedure D was followed using protected isoquinuclidine (70 mg, 0.157 mmol, 1.0 equiv) in $CH_2Cl_2$ (1.6 mL) and was reacted with trifluoroacetic acid (120 µL, 1.57 mmol, 10.0 equiv) for 12 h. The crude product was purified by preparative thin-layer chromatography (10% MeOH/$CH_2Cl_2$ + 1% $NH_4OH$) to yield the racemic isoquinuclidine (38 mg, 70% yield) as a pale-yellow oil. $^1$H NMR (500 MHz, $CD_3OD$) δ 7.31 – 7.17 (m, 2H), 6.96 – 6.81 (m, 3H), 6.00 (dd, *J* = 6.3, 1.9 Hz, 1H), 3.81 (dd, *J* = 9.7, 5.0 Hz, 1H), 3.71 (dd, *J* = 6.4, 3.2 Hz, 1H), 3.63 (s, 3H), 3.60 – 3.52 (m, 1H), 3.24 (dt, *J* = 9.6, 3.9 Hz, 1H), 2.92 – 2.77 (m, 4H), 2.44 (t, *J* = 5.3 Hz, 1H), 1.81 – 1.70 (m, 4H), 1.60 (dd, *J* = 13.1, 9.4 Hz, 1H), 1.24 (s, 3H). $^{13}$C NMR (126 MHz, $CD_3OD$) δ 174.9, 158.7, 144.1, 129.1, 124.9, 120.5, 114.1, 70.6, 68.1, 54.7, 53.8, 50.9, 39.2, 38.9, 37.4, 34.5, 19.0, 17.1. HRMS (ESI+, m/z) calcd for $C_{20}H_{29}N_2O_3$ $(M+H)^+$ : 345.2173, found: 345.2166

**Synthesis of #004**

・**Synthesis of 0.8 M imine solution**

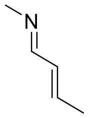

To a flame-dried vial was added a magnetic stir bar, *trans*-crotonaldehyde (0.6 mL, 7.3 mmol, 1.0 equiv) in dry THF (1.2 mL), and molecular sieves 4 Å. To this solution was then slowly added 2.0 M methylamine in THF (7.3 mL, 14.6 mmol, 2.0 equiv) at 0 °C and the reaction solution was stirred at 0 °C for 2 h. Upon reaction completion, the 0.8 M imine solution was stored at –20 °C in a nitrogen filled glovebox and taken on to the next step without further purification.

・**Key Reaction**

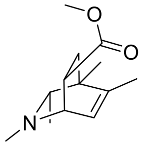

**(±)-#004**

**Final Compound #004** General procedure B was followed using above obtained 0.8 M solution of imine (3 mL, 2.4 mmol, 1.0 equiv) and 2-butyne (941 µL, 8.0 mmol, 5 equiv) in THF (0.6 mL). Rh catalyst (10 mol %, 2.4 mL, 0.24 mmol, 100 mM in THF) was added, and the reaction was carried out at 65 °C for 12 h to produce the DHP intermediate. Following general procedure C, methyl acrylate (2.16 mL, 24.0 mmol, 10 equiv) was added to the solution of DHP intermediate, and the reaction was carried out at 50 °C for 12 h. The resulting crude material was purified by column chromatography (30% *t*BuOMe/Pentane+1% $Et_3N$) to afford the desired product (42 mg, 8% yield from imine) as a pale-yellow oil. $^1$H NMR (400 MHz, $CD_3OD$) δ 5.99 (dd, *J* = 6.2, 1.8 Hz, 1H), 3.61 (s, 3H), 3.46 (dd, *J* = 6.2, 2.9 Hz, 1H), 3.21 (ddd, *J* = 9.5, 5.1, 2.9 Hz, 1H), 2.38 (s, 3H), 1.97 (q, *J* = 6.3 Hz, 1H), 1.74 (d, *J* = 1.8 Hz, 3H), 1.66 (dd, *J* = 12.9, 5.1 Hz, 1H), 1.47 (dd, *J* = 12.9, 9.5 Hz, 1H), 1.08 (s, 3H), 0.82 (d, *J* = 6.3 Hz, 3H). $^{13}$C NMR (101 MHz, $CD_3OD$) δ 174.9, 143.4, 124.9, 64.7, 56.2, 50.9, 39.9, 39.2, 36.3, 35.4, 18.5, 17.4, 16.8. HRMS (ESI+, m/z) calcd for $C_{13}H_{21}NO_2$ $(M+H)^+$ : 224.1645, found:

224.1633.

**Synthesis of #005**

・**Synthesis of Imine**

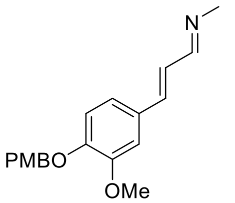

General procedure A was followed using 2.0 M methylamine in THF (0.8 mL, 1.6 mmol, 2.0 equiv), (*E*)-3-(3-methoxy-4-((4-methoxybenzyl)oxy)phenyl)acrylaldehyde (239 mg, 0.80 mmol, 1.0 equiv), molecular sieves 4 Å, and THF (1.6 mL). Imine (245 mg, 98% yield) was obtained as a yellow solid. $^{1}$H NMR (400 MHz, $CDCl_3$) δ 7.98 (d, *J* = 7.4 Hz, 1H), 7.33 (d, *J* = 8.3 Hz, 2H), 7.04 (s, 1H), 6.95 (d, *J* = 8.5 Hz, 1H), 6.89 – 6.83 (m, 5H), 5.08 (s, 2H), 3.87 (s, 3H), 3.79 (s, 3H), 3.41 (s, 3H).

・**Key Reaction**

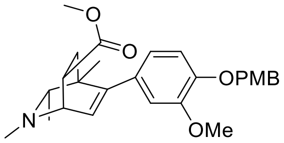

General procedure B was followed using above obtained imine (245 mg, 0.787 mmol, 1.0 equiv) and 2-butyne (308 μL, 3.93 mmol, 5 equiv) in THF (0.79 mL). Rh catalyst (10 mol %, 0.79 mL, 0.079 mmol, 100 mM in THF) was added, and the reaction was carried out at 65 °C for 12 h to produce the DHP intermediate. Following general procedure C, methyl acrylate (0.71 mL, 7.87 mmol, 10 equiv) was added to the solution of DHP intermediate, and the reaction was carried out at 50 °C for 16 h. The resulting crude material was purified by column chromatography (50% *t*BuOMe/Pentane+1% $Et_3N$) to afford the desired product (150 mg, 42% yield from imine) as a pale-yellow oil. $^{1}$H NMR (400 MHz, $CD_3OD$) δ 7.34 (d, *J* = 8.7 Hz, 2H), 6.92 (d, *J* = 8.1 Hz, 1H), 6.89 (d, *J* = 8.5 Hz, 2H), 6.70 (d, *J* = 1.9 Hz, 1H), 6.65 (dd, *J* = 8.2, 1.9 Hz, 1H), 6.12 (d, *J* = 6.3 Hz, 1H), 4.98 (s, 2H), 3.80 (s, 3H), 3.77 (s, 3H), 3.69 – 3.61 (m, 4H), 3.35 (ddd, *J* = 9.8, 4.9, 3.1 Hz, 1H), 2.46 (s, 3H), 2.07 (q, *J* = 6.3 Hz, 1H), 1.90 (dd, *J* = 13.0, 4.9 Hz, 1H), 1.60 (dd, *J* = 13.0, 9.4 Hz, 1H), 0.99 (s, 3H), 0.95 (d, *J* = 6.3 Hz, 3H).

・**Deprotection**

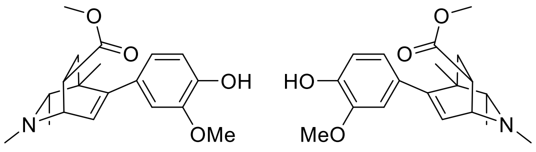

**#005**
enantiomers separated by chiral HPLC

**Final Compound #005** General Procedure D was followed using protected isoquinuclidine (30 mg, 0.066 mmol, 1.0 equiv) in $CH_2Cl_2$ (1.0 mL) and was reacted with trifluoroacetic acid (49 µL, 0.66 mmol, 10.0 equiv) for 12 h. The crude product was purified by preparative thin-layer chromatography (10% MeOH/$CH_2Cl_2$ + 1% $NH_4OH$) to yield the racemic isoquinuclidine (21 mg, 95% yield) as a pale-yellow oil. $^1H$ NMR (400 MHz, $CD_3OD$) δ 6.73 (d, $J$ = 8.1 Hz, 1H), 6.66 (d, $J$ = 1.9 Hz, 1H), 6.58 (dd, $J$ = 8.1, 2.0 Hz, 1H), 6.11 (d, $J$ = 6.3 Hz, 1H), 3.82 (s, 3H), 3.72 (dd, $J$ = 6.3, 3.0 Hz, 1H), 3.66 (s, 3H), 3.37 (ddd, $J$ = 9.4, 4.9, 3.0 Hz, 1H), 2.53 (s, 3H), 2.18 (q, $J$ = 6.4 Hz, 1H), 1.90 (dd, $J$ = 13.0, 4.9 Hz, 1H), 1.64 (dd, $J$ = 13.0, 9.4 Hz, 1H), 1.01 (s, 3H), 0.99 (d, $J$ = 6.4 Hz, 3H). $^{13}C$ NMR (101 MHz, $CD_3OD$) δ 174.5, 148.9, 146.9, 145.7, 131.1, 126.8, 120.8, 114.4, 111.7, 65.4, 56.4, 54.9, 51.1, 40.8, 39.1, 36.6, 36.1, 20.2, 16.9. HRMS (ESI+, m/z) calcd for $C_{19}H_{25}NO_4$ $(M+H)^+$ : 332.1856, found: 332.1827.

**HPLC Traces of #005**

A portion of this material was separated using semi-preparative chiral HPLC (Chiralpak IC column, 250 x 10 mm, 20% *i*PrOH/Hexanes+1% DEA, 2.5 mL/min) to provide the two enantiomers with tr = 15 min and 35 min.

Chiralpak IC, analytical column (250 x 4.6 mm), 30% *i*PrOH/Hexanes+0.1% DEA, 1 mL/min, 254 nm

**Racemic**

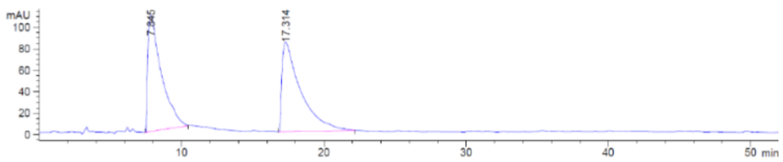

**Enantiomer 1**

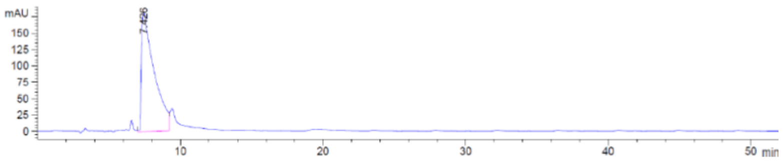

**Enantiomer 2**

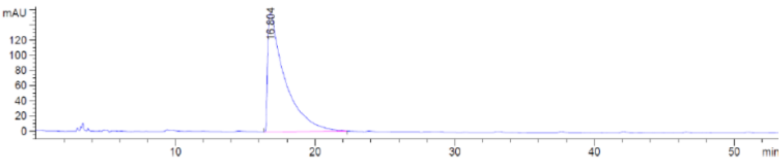

**Synthesis of #006**

・**Synthesis of Imine**

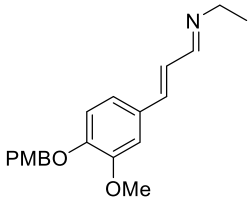

General procedure A was followed using 2.0 M ethylamine in THF (0.8 mL, 1.6 mmol, 2.0 equiv), (*E*)-3-(3-methoxy-4-((4-methoxybenzyl)oxy)phenyl)acrylaldehyde (239 mg, 0.80 mmol, 1.0 equiv), molecular sieves 4 Å, and THF (1.6 mL). Imine (248 mg, 95% yield) was obtained as a yellow solid. $^1$H NMR (400 MHz, $CDCl_3$) δ 7.99 (d, *J* = 7.4 Hz, 1H), 7.34 (d, *J* = 8.6 Hz, 2H), 7.06 (s, 1H), 6.99 – 6.82 (m, 6H), 5.09 (s, 2H), 3.87 (s, 3H), 3.79 (s, 3H), 3.55 (q, *J* = 8.5 Hz, 2H), 1.32 – 1.21 (m, 3H).

・**Key Reaction**

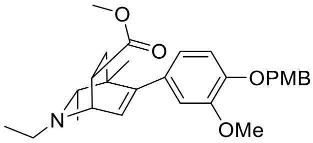

General procedure B was followed using above obtained imine (248 mg, 0.762 mmol, 1.0 equiv) and 2-butyne (299 µL, 3.81 mmol, 5 equiv) in THF (0.76 mL). Rh catalyst (10 mol %, 0.76 mL, 0.076 mmol, 100 mM in THF) was added, and the reaction was carried out at 65 °C for 12 h to produce the DHP intermediate. Following general procedure C, methyl acrylate (0.67 mL, 7.62 mmol, 10 equiv) was added to the solution of DHP intermediate, and the reaction was carried out at 50 °C for 16 h. The resulting crude material was purified by column chromatography (50% *t*BuOMe/Pentane+1% $Et_3N$) to afford the desired product (180 mg, 51% yield from imine) as a pale-yellow oil. $^1$H NMR (400 MHz, $CD_3OD$) δ 7.34 (d, *J* = 8.7 Hz, 2H), 6.92 (d, *J* = 8.3 Hz, 1H), 6.89 (d, *J* = 8.7 Hz, 2H), 6.71 (d, *J* = 2.0 Hz, 1H), 6.66 (dd, *J* = 8.3, 2.0 Hz, 1H), 6.08 (d, *J* = 6.4 Hz, 1H), 4.99 (s, 2H), 3.85 (dd, *J* = 6.4, 3.1 Hz, 1H), 3.80 (s, 3H), 3.77 (s, 3H), 3.65 (s, 3H), 2.76 (dq, *J* = 14.1, 7.2 Hz, 1H), 2.63 (dq, *J* = 14.1, 7.2 Hz, 1H), 2.15 (q, *J* = 6.4 Hz, 1H), 1.93 (dd, *J* = 13.1, 4.3 Hz, 1H), 1.58 (dd, *J* = 13.1, 9.4 Hz, 1H), 1.18 (t, *J* = 7.2 Hz, 3H), 1.01 (s, 3H), 0.98 (d, *J* = 6.4 Hz, 3H).

・**Deprotection**

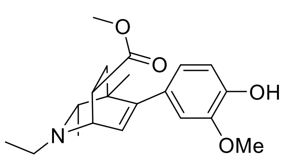

**(±)-#006**

**Final Compound #006** General Procedure D was followed using protected isoquinuclidine (30 mg, 0.064 mmol, 1.0 equiv) in $CH_2Cl_2$ (1.0 mL) and was reacted with trifluoroacetic acid (48 µL, 0.64 mmol, 10.0 equiv) for 12 h. The crude product was purified by preparative thin-layer chromatography (10% MeOH/$CH_2Cl_2$ + 1% $NH_4OH$) to yield the racemic isoquinuclidine (22 mg, 99% yield) as a pale-yellow oil.
$^1$H NMR (400 MHz, $CD_3OD$) δ 6.72 (d, *J* = 8.1 Hz, 1H), 6.66 (d, *J* = 2.0 Hz, 1H), 6.58 (dd, *J* = 8.1, 2.0 Hz, 1H), 6.07 (d, *J* = 6.4 Hz, 1H), 3.88 (dd, *J* = 6.4, 3.1 Hz, 1H), 3.82 (s, 3H), 3.65 (s, 3H), 3.34 – 3.30 (m, 1H), 2.80 (dq, *J* = 12.2, 7.2 Hz, 1H), 2.66 (dq, *J* = 12.2, 7.2 Hz, 1H), 2.19 (q, *J* = 6.4 Hz, 1H), 1.92 (dd, *J* = 13.1, 4.4 Hz, 1H), 1.59 (dd, *J* = 13.1, 9.4 Hz, 1H), 1.19 (t, *J* = 7.2 Hz, 3H), 1.02 (s, 3H), 0.99 (d, *J* = 6.4 Hz, 3H). $^{13}$C NMR (101 MHz, $CD_3OD$) δ 174.7, 149.6, 146.9, 145.6, 131.4, 126.4, 120.7, 114.3, 111.7, 64.9, 54.9, 53.6, 51.1, 48.3, 40.8, 36.9, 35.2, 20.5, 18.9, 12.6. HRMS (ESI+, m/z) calcd for $C_{20}H_{27}NO_4$ $(M+H)^+$ : 346.2013, found: 346.1993.

**Synthesis of #007**

・**Synthesis of Imine**

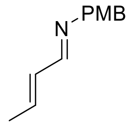

General procedure A was followed using (4-methoxyphenyl)methanamine (230 mg, 1.68 mmol, 1.0 equiv), *trans*-crotonaldehyde (277 µL, 3.35 mmol, 2.0 equiv), molecular sieves 4 Å, and THF (3.4 mL). Imine (300 mg, 95% yield) was obtained as a pale-yellow oil. $^1$H NMR (400 MHz, $CDCl_3$) δ 7.90 (d, *J* = 8.1 Hz, 1H), 7.17 (d, *J* = 8.5 Hz, 2H), 6.85 (d, *J* = 8.6 Hz, 2H), 6.31 – 6.15 (m, 2H), 4.54 (s, 2H), 3.77 (s, 3H), 1.87 (d, *J* = 5.7 Hz, 3H).

・**Key Reaction**

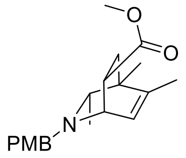

General procedure B was followed using above obtained imine (300 mg, 1.59 mmol, 1.0 equiv) and 2-butyne (621 µL, 7.93 mmol, 5 equiv) in THF (1.6 mL). Rh catalyst (10 mol %, 1.6 mL, 0.16 mmol, 100 mM in THF) was added, and the reaction was carried out at 65 °C for 12 h to produce the DHP intermediate. Following general procedure C, methyl acrylate (1.43 mL, 15.9 mmol, 10 equiv) was added to the solution of DHP intermediate, and the reaction was carried out at 50 °C for 16 h. The resulting crude material was purified by column chromatography (30% *t*BuOMe/Pentane+1% $Et_3N$)

to afford the desired product (155 mg, 30% yield from imine) as a pale-yellow oil. [1]H NMR (400 MHz, $CDCl_3$) δ 7.25 (d, *J* = 8.5 Hz, 2H), 6.83 (d, *J* = 8.5 Hz, 2H), 5.92 (dd, *J* = 6.4, 1.8 Hz, 1H), 3.78 (s, 3H), 3.66 (d, *J* = 12.7 Hz, 1H), 3.58 (s, 3H), 3.56 – 3.45 (m, 2H), 3.20 (ddd, *J* = 9.5, 4.9, 3.0 Hz, 1H), 2.07 (q, *J* = 6.2 Hz, 1H), 1.80 – 1.67 (m, 4H), 1.45 (dd, *J* = 12.8, 9.5 Hz, 1H), 1.06 (s, 3H), 0.72 (d, *J* = 6.2 Hz, 3H).

・**Deprotection of PMB on nitrogen**

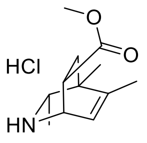

**(±)-#007**

**Final Compound #007** General Procedure E was followed using protected isoquinuclidine (75 mg, 0.228 mmol, 1.0 equiv) in DCE (1.0 mL) and was reacted with 1-chloroethyl chloroformate (123 μL, 1.14 mmol, 5.0 equiv) for 3 h. The crude product was washed with pentane to yield the HCl salt of isoquinuclidine (28 mg, 50% yield) as a white-solid. [1]H NMR (400 MHz, $CD_3OD$) δ 6.01 (dd, *J* = 6.0, 1.8 Hz, 1H), 4.35 (dd, *J* = 5.9, 2.8 Hz, 1H), 3.66 (s, 3H), 3.28 – 3.20 (m, 2H), 1.93 – 1.81 (m, 4H), 1.69 (dd, *J* = 13.4, 5.6 Hz, 1H), 1.24 (s, 3H), 1.09 (d, *J* = 6.8 Hz, 3H). [13]C NMR (101 MHz, $CD_3OD$) δ 171.6, 148.6, 119.2, 55.9, 51.4, 48.2, 39.9, 39.3, 33.8, 17.5, 17.1, 15.2. HRMS (ESI+, m/z) calcd for $C_{12}H_{19}NO_2$ $(M+H)^+$ : 210.1489, found: 210.1476.

**Synthesis of #008**

・**Synthesis of Imine**

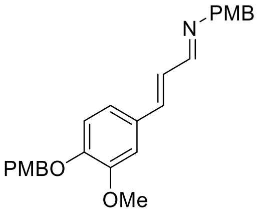

General procedure A was followed using (4-methoxyphenyl)methanamine (104 mg, 0.756 mmol, 1.1 equiv), (*E*)-3-(3-methoxy-4-((4-methoxybenzyl)oxy)phenyl)acrylaldehyde (205 mg, 0.687 mmol, 1.0 equiv), molecular sieves 4 Å, and THF (1.4 mL). Imine (280 mg, 98% yield) was obtained as a yellow solid. [1]H NMR (400 MHz, $CDCl_3$) δ 8.04 (d, *J* = 6.9 Hz, 1H), 7.34 (d, *J* = 8.6 Hz, 2H), 7.21 (d, *J* = 8.6 Hz, 2H), 7.03 (d, *J* = 2.0 Hz, 1H), 6.94 (dd, *J* = 8.3, 2.0 Hz, 1H), 6.91 – 6.84 (m, 7H), 5.09 (s, 2H), 4.63 (s, 2H), 3.86 (s, 3H), 3.78 (s, 6H).

・**Key Reaction**

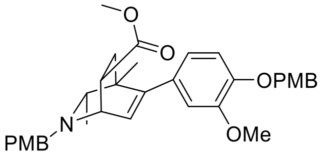

General procedure B was followed using above obtained imine (200 mg, 0.479 mmol, 1.0 equiv) and 2-butyne (188 µL, 2.40 mmol, 5 equiv) in THF (0.48 mL). Rh catalyst (10 mol %, 0.48 mL, 0.048 mmol, 100 mM in THF) was added, and the reaction was carried out at 65 °C for 12 h to produce the DHP intermediate. Following general procedure C, methyl acrylate (0.43 mL, 4.79 mmol, 10 equiv) was added to the solution of DHP intermediate, and the reaction was carried out at 50 °C for 16 h. The resulting crude material was purified by column chromatography (50% *t*BuOMe/Pentane+1% $Et_3N$) to afford the desired product (115 mg, 43% yield from imine) as a pale-yellow oil. $^1$H NMR (400 MHz, $CD_3OD$) δ 7.35 – 7.27 (m, 4H), 6.91 – 6.85 (m, 5H), 6.70 (d, *J* = 1.9 Hz, 1H), 6.63 (dd, *J* = 8.2, 1.9 Hz, 1H), 6.00 (d, *J* = 6.4 Hz, 1H), 4.96 (s, 2H), 3.80 – 3.72 (m, 10H), 3.66 (dd, *J* = 6.4, 2.9 Hz, 1H), 3.63 – 3.54 (m, 4H), 3.43 (dt, *J* = 7.8, 3.8 Hz, 1H), 2.23 (q, *J* = 6.2 Hz, 1H), 1.93 (dd, *J* = 12.9, 4.5 Hz, 1H), 1.63 (dd, *J* = 12.9, 9.4 Hz, 1H), 0.99 (s, 3H), 0.81 (d, *J* = 6.2 Hz, 3H).

**・Deprotection of PMB on nitrogen**

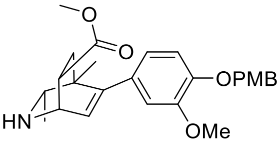

General Procedure E was followed using di-protected isoquinuclidine (25 mg, 0.045 mmol, 1.0 equiv) in DCE (1.0 mL) and was reacted with 1-chloroethyl chloroformate (24 µL, 0.224 mmol, 5.0 equiv) for 12 h. After methanol treatment according to the general procedure, the crude product was purified by preparative thin-layer chromatography (10% MeOH/$CH_2Cl_2$ + 1% $NH_4OH$) to yield the mono-protected isoquinuclidine (8.5 mg, 40% yield) as a pale-yellow oil. $^1$H NMR (400 MHz, $CDCl_3$) δ 7.35 (d, *J* = 8.7 Hz, 2H), 6.88 (d, *J* = 8.7 Hz, 2H), 6.81 (d, *J* = 8.2 Hz, 1H), 6.66 (d, *J* = 2.0 Hz, 1H), 6.62 (dd, *J* = 8.2, 2.0 Hz, 1H), 6.15 (d, *J* = 5.8 Hz, 1H), 5.05 (s, 2H), 3.92 (dd, *J* = 5.8, 2.9 Hz, 1H), 3.85 (s, 3H), 3.79 (s, 3H), 3.64 (s, 3H), 3.09 (ddd, *J* = 9.8, 5.2, 2.9 Hz, 1H), 2.82 (q, *J* = 6.4 Hz, 1H), 1.82 (dd, *J* = 12.9, 5.2 Hz, 1H), 1.70 (dd, *J* = 12.9, 9.8 Hz, 1H), 1.06 (s, 3H), 0.88 (d, *J* = 6.4 Hz, 3H).

**・Deprotection of PMB on oxygen**

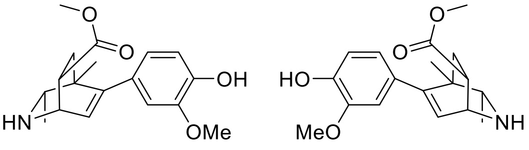

**#008**
enantiomers separated by chiral HPLC

**Final Compound #008** General Procedure D was followed using mono-protected isoquinuclidine (8.5 mg, 0.019 mmol, 1.0 equiv) in $CH_2Cl_2$ (1.0 mL) and was reacted with trifluoroacetic acid (15 µL, 0.19 mmol, 10.0 equiv) for 12 h. The crude product was purified by preparative thin-layer chromatography (10% MeOH/$CH_2Cl_2$ + 1% $NH_4OH$) to yield the racemic isoquinuclidine (6.1 mg, 99% yield) as a dark-brown oil. $^1$H NMR (400 MHz, $CD_3OD$) δ 6.72 (d, *J* = 8.0 Hz, 1H), 6.66 (d, *J* = 2.0 Hz, 1H), 6.58 (dd, *J* = 8.0, 2.0 Hz, 1H), 6.06 (d, *J* = 5.7 Hz, 1H), 3.87 (dd, *J* = 5.7, 3.0 Hz, 1H), 3.82 (s, 3H), 3.64 (s, 3H), 3.05 (ddd, *J* = 9.6, 5.0, 3.0 Hz, 1H), 2.75 (q, *J* = 6.4 Hz, 1H), 1.82 (dd, *J* = 13.0, 5.0 Hz, 1H), 1.68 (dd, *J* = 13.0, 9.6 Hz, 1H), 1.05 (s, 3H), 0.91 (d, *J* = 6.4 Hz, 3H). $^{13}$C NMR (126 MHz, $CD_3OD$) δ 174.2, 147.9, 146.9, 145.5, 131.8, 126.7, 120.9, 114.4, 111.9, 56.3, 55.0, 50.9, 49.0, 44.0, 40.3, 36.4, 19.9, 18.2. HRMS (ESI+, m/z) calcd for $C_{18}H_{23}NO_4$ $(M+H)^+$ : 318.1700, found: 318.1674.

**HPLC Traces of #008**

A portion of this material was separated using semi-preparative chiral HPLC (Chiralpak IC column, 250 x 10 mm, 20% *i*PrOH/Hexanes+1% DEA, 2.5 mL/min) to provide the two enantiomers with tr = 18.5 min and 25 min.

Chiralpak IC, analytical column (250 x 4.6 mm), 20% *i*PrOH/Hexanes+0.1% DEA, 1 mL/min, 254 nm

**Racemic**

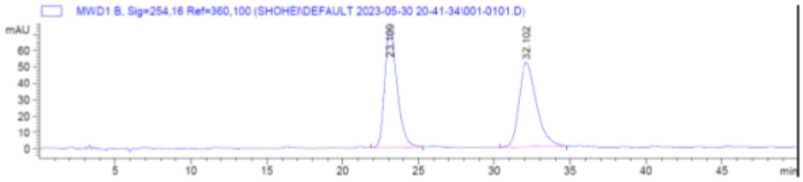

**Enantiomer 1**

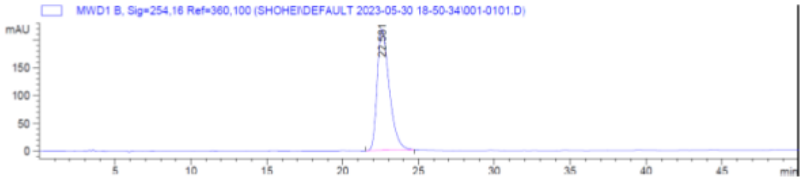

**Enantiomer 2**

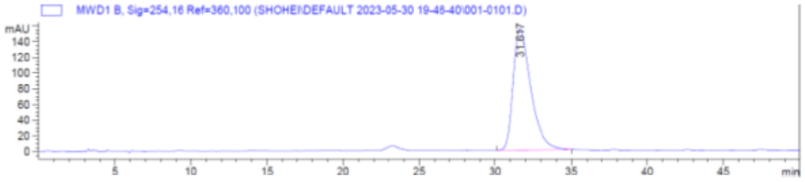

**Synthesis of #009**

・**Key Reaction**

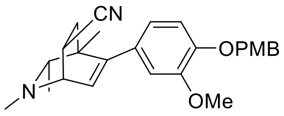

General procedure B was followed using above obtained imine (225 mg, 0.723 mmol, 1.0 equiv) and 2-butyne (283 µL, 3.61 mmol, 5 equiv) in THF (0.72 mL). Rh catalyst (10 mol %, 0.72 mL, 0.072 mmol, 100 mM in THF) was added, and the reaction was carried out at 65 °C for 12 h to produce the DHP intermediate. Following general procedure C, acrylonitrile (383 mg, 7.23 mmol, 10 equiv) was added to the solution of DHP intermediate, and the reaction was carried out at 50 °C for 16 h. The resulting crude material was purified by column chromatography (70% EtOAc/Hexane+1% $Et_3N$) to afford the desired product (133 mg, 44% yield from imine) as a pale-yellow oil. $^1H$ NMR (400 MHz, $CD_3OD$) δ 7.34 (d, *J* = 8.6 Hz, 2H), 6.95 (d, *J* = 8.2 Hz, 1H), 6.89 (d, *J* = 8.6 Hz, 2H), 6.76 (d, *J* = 2.0 Hz, 1H), 6.71 (dd, *J* = 8.2, 2.0 Hz, 1H), 6.37 (d, *J* = 6.3 Hz, 1H), 4.99 (s, 2H), 3.81 (s, 3H), 3.76 (s, 3H), 3.63 (dd, *J* = 6.3, 2.8 Hz, 1H), 3.51 (ddd, *J* = 9.7, 4.7, 2.8 Hz, 1H), 2.43 (s, 3H), 2.06 (q, *J* = 6.4 Hz, 1H), 1.94 (dd, *J* = 13.1, 9.7 Hz, 1H), 1.62 (dd, *J* = 13.1, 4.7 Hz, 1H), 1.00 (s, 3H), 0.93 (d, *J* = 6.4 Hz, 3H).

・**Deprotection**

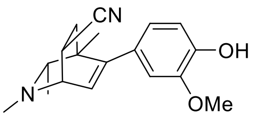

**(±)-#009**

**Final Compound #009** General Procedure D was followed using protected isoquinuclidine (42 mg, 0.787 mmol, 1.0 equiv) in $CH_2Cl_2$ (1.0 mL) and was reacted with trifluoroacetic acid (75 µL, 1.0 mmol, 10.0 equiv) for 12 h. The crude product was purified by preparative thin-layer chromatography (10% MeOH/$CH_2Cl_2$ + 1% $NH_4OH$) to yield the racemic isoquinuclidine (27 mg, 90% yield) as a pale-yellow oil. $^1H$ NMR (400 MHz, $CD_3OD$) δ 6.76 (d, *J* = 8.1 Hz, 1H), 6.71 (d, *J* = 2.0 Hz, 1H), 6.64 (dd, *J* = 8.1, 2.0 Hz, 1H), 6.35 (d, *J* = 6.4 Hz, 1H), 3.83 (s, 3H), 3.62 (dd, *J* = 6.4, 2.8 Hz, 1H), 3.50 (ddd, *J* = 9.7, 4.8, 2.8 Hz, 1H), 2.42 (s, 3H), 2.06 (q, *J* = 6.3 Hz, 1H), 1.94 (dd, *J* = 13.1, 9.7 Hz, 1H), 1.61 (dd, *J* = 13.1, 4.8 Hz, 1H), 1.01 (s, 3H), 0.93 (d, *J* = 6.3 Hz, 3H). $^{13}C$ NMR (101 MHz, $CD_3OD$) δ 149.1, 147.1, 145.9, 130.9, 127.7, 122.2, 120.8, 114.5, 111.7, 64.4, 55.7, 54.9, 40.2, 39.1, 38.6, 22.1, 19.9, 17.3. HRMS (ESI+, m/z) calcd for $C_{18}H_{23}N_2O_2$ $(M+H)^+$ : 299.1754, found: 299.1759.

**Synthesis of #010**

・**Key Reaction**

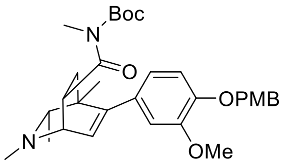

General procedure B was followed using above obtained imine (225 mg, 0.723 mmol, 1.0 equiv) and 2-butyne (283 μL, 3.61 mmol, 5 equiv) in THF (0.72 mL). Rh catalyst (10 mol %, 0.72 mL, 0.072 mmol, 100 mM in THF) was added, and the reaction was carried out at 65 °C for 12 h to produce the DHP intermediate. Following general procedure C, Boc-protected *N*-methyl acrylamide (669 mg, 3.61 mmol, 5 equiv) was added to the solution of DHP intermediate, and the reaction was carried out at 50 °C for 16 h. The resulting crude material was purified by column chromatography (50% EtOAc/Hexane+1% $Et_3N$) to afford the desired product (125 mg, 31% yield from imine) as a pale-yellow oil. $^1H$ NMR (400 MHz, $CD_3OD$) δ 7.33 (d, *J* = 8.6 Hz, 2H), 6.91 (d, *J* = 8.2 Hz, 1H), 6.89 (d, *J* = 8.6 Hz, 2H), 6.75 (d, *J* = 2.0 Hz, 1H), 6.68 (dd, *J* = 8.2, 2.0 Hz, 1H), 6.07 (d, *J* = 6.4 Hz, 1H), 4.98 (s, 2H), 4.48 (ddd, *J* = 9.3, 5.3, 2.5 Hz, 1H), 3.80 (s, 3H), 3.76 (s, 3H), 3.56 (dd, *J* = 6.4, 2.5 Hz, 1H), 3.08 (s, 3H), 2.49 (s, 3H), 2.07 (q, *J* = 6.3 Hz, 1H), 1.92 (dd, *J* = 12.6, 5.3 Hz, 1H), 1.60 (dd, *J* = 12.6, 9.3 Hz, 1H), 1.54 (s, 9H), 0.98 (s, 3H), 0.96 (d, *J* = 6.3 Hz, 3H).

・**Deprotection**

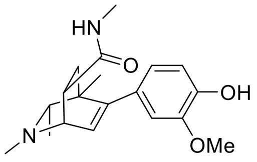

**(±)-#010**

**Final Compound #010** General Procedure D was followed using protected isoquinuclidine (43 mg, 0.078 mmol, 1.0 equiv) in $CH_2Cl_2$ (1.0 mL) and was reacted with trifluoroacetic acid (58 μL, 0.780 mmol, 10.0 equiv) for 12 h. The crude product was purified by preparative thin-layer chromatography (10% MeOH/$CH_2Cl_2$ + 1% $NH_4OH$) to yield the racemic isoquinuclidine (25 mg, 97% yield) as a pale yellow oil. $^1H$ NMR (400 MHz, $CD_3OD$) δ 6.73 (d, *J* = 1.9 Hz, 1H), 6.71 (d, *J* = 8.1 Hz, 1H), 6.62 (dd, *J* = 8.1, 1.9 Hz, 1H), 6.10 (d, *J* = 6.3 Hz, 1H), 3.82 (s, 3H), 3.46 (dd, *J* = 6.3, 2.7 Hz, 1H), 3.18 (ddd, *J* = 9.4, 5.3, 2.7 Hz, 1H), 2.69 (s, 3H), 2.46 (s, 3H), 2.03 (q, *J* = 6.3 Hz, 1H), 1.85 (dd, *J* = 12.6, 5.3 Hz, 1H), 1.52 (dd, *J* = 12.6, 9.4 Hz, 1H), 0.99 (s, 3H), 0.95 (d, *J* = 6.3 Hz, 3H). $^{13}C$ NMR (101 MHz, $CD_3OD$) δ 175.2, 147.5, 146.9, 145.5, 131.6, 127.6, 120.8, 114.3, 111.9, 64.9, 57.3, 54.9, 40.9, 39.1, 37.2, 36.8, 25.2, 20.4, 17.3. HRMS (ESI+, m/z) calcd for $C_{19}H_{27}N_2O_3$ $(M+H)^+$ : 331.2016, found: 331.2023.

**Synthesis of #011**

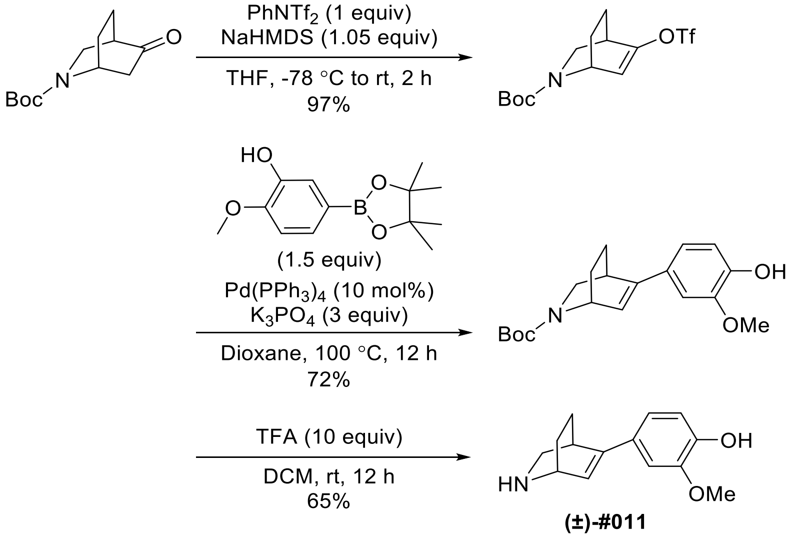

To a flame-dried round-bottom flask was added a magnetic stir bar, 2-azabicyclo[2.2.2]octane-2-carboxylic acid 5-oxo-1,1-dimethylethyl ester (900 mg, 3.99 mmol, 1 equiv) and *N*-phenyl-bis(trifluoromethanesulfonimide) (1570 mg, 4.39 mmol, 1.1 equiv) dissolved in THF (8 mL). This solution was cooled to -78 ° C and then 1.0 M sodium hexamethyldisilazane in THF (4.2 mL, 4.19 mmol, 1.05 equiv) was added dropwise over 5 min. The reaction mixture was stirred at room temperature for 2 h. After the reaction was complete as monitored by thin-layer chromatography, the mixture was then diluted with water, extracted with ethyl acetate, washed with brine, dried with $Na_2SO_4$ and filtered. The organic layer was concentrated and the crude residue was purified via silica gel chromatography (gradient of 0-50% EtOAc in hexane) to give 1390 mg of the *O*-protected compound in 97% yield. To a stirred solution of the above obtained compound (500 mg, 1.4 mmol, 1 equiv) in dioxane (7 mL) was added $K_3PO_4$ (891 mg, 4.20 mmol, 3 equiv), 4-hydroxy-3-methoxyphenylboronicacid pinacol ester (525 mg, 2.1 mmol, 1.5 equiv), $Pd(PPh_3)_4$ (162 mg, 0.14 mmol, 10 mol%) and the reaction mixture was heated under reflux for 12 h. After the reaction was complete as monitored by thin-layer chromatography, the mixture was then diluted with water, extracted with ethyl acetate, washed with brine, dried with $Na_2SO_4$ and filtered. The organic layer was concentrated and the crude residue was purified via silica gel chromatography (gradient of 0-100% EtOAc in hexane) to give 335 mg of the arylated compound in 72% yield. $^1$H NMR (400 MHz, $CD_3OD$) δ 7.00 (s, 1H), 6.90 (d, $J$ = 8.2 Hz, 1H), 6.75 (dd, $J$ = 8.3, 1.1 Hz, 1H), 6.54 (d, $J$ = 7.0 Hz, 1H), 4.75 – 4.64 (m, 1H), 3.84 (s, 3H), 3.40 – 3.21 (m, 2H), 3.04 – 2.89 (m, 1H), 2.07 – 1.91 (m, 1H), 1.82 – 1.68 (m, 1H), 1.49 – 1.39 (m, 11H).

**Final Compound #011** General Procedure D was followed using above obtained isoquinuclidine (100 mg, 0.302 mmol, 1.0 equiv) in $CH_2Cl_2$ (3.0 mL) and was reacted with trifluoroacetic acid (224 µL, 3.02 mmol, 10.0 equiv) for 12 h. The crude product was purified by preparative thin-layer chromatography (10% MeOH/$CH_2Cl_2$ + 1% $NH_4OH$) to yield the racemic isoquinuclidine (45 mg, 65% yield) as a dark-brown oil. $^1$H NMR (400 MHz, $CD_3OD$) δ 7.01 (d, *J* = 2.0 Hz, 1H), 6.91 (dd, *J* = 8.2, 2.1 Hz, 1H), 6.74 (d, *J* = 8.2 Hz, 1H), 6.53 (dd, *J* = 6.1, 1.9 Hz, 1H), 3.84 (s, 3H), 3.77 (dt, *J* = 5.9, 2.7 Hz, 1H), 3.21 (q, *J* = 2.4 Hz, 1H), 3.00 (dd, *J* = 10.5, 2.0 Hz, 1H), 2.58 (dt, *J* = 10.5, 2.6 Hz, 1H), 2.10 – 1.91 (m, 1H), 1.89 – 1.68 (m, 1H), 1.52 – 1.36 (m, 2H). $^{13}$C NMR (101 MHz, $CD_3OD$) δ 148.0, 147.2, 145.1, 128.9, 123.0, 117.7, 115.1, 108.1, 54.9, 46.7, 44.7, 31.7, 25.2, 22.6. HRMS (ESI+, m/z) calcd for $C_{14}H_{18}NO_2$ $(M+H)^+$ : 232.1332, found: 232.1309.

**Synthesis of #012**

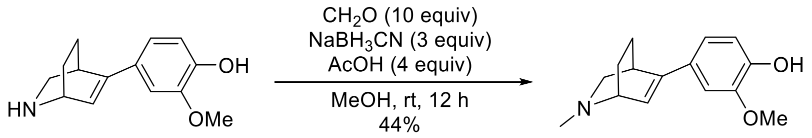

**Final Compound #012** The solution of the above obtained compound (45 mg, 0.156 mmol, 1 equiv) in MeOH (3 mL) was added AcOH (36 µL, 0.622 mmol, 4 equiv), formaldehyde (0.14 mL of 37% w/w solution, 4.67 mmol, 10 equiv), and $NaBH_3CN$ (29 mg, 0.467 mmol, 3 equiv) and the reaction mixture was stirred at room temperature for 12 h. After the reaction was complete as monitored by thin-layer chromatography, the mixture was then diluted with water, extracted with $CH_2Cl_2$, washed with brine, dried with $Na_2SO_4$ and filtered. The organic layer was concentrated and the crude residue was purified by preparative thin-layer chromatography (5% MeOH/$CH_2Cl_2$ + 1% $NH_4OH$) to yield the racemic isoquinuclidine (21 mg, 45% yield) as a white solid. $^1$H NMR (500 MHz, $CD_3OD$) δ 7.07 (s, 1H), 6.97 (dd, *J* = 8.2, 1.9 Hz, 1H), 6.78 (d, *J* = 8.2 Hz, 1H), 6.50 (dd, *J* = 5.8, 1.9 Hz, 1H), 3.88 (s, 3H), 3.76 (dt, *J* = 5.8, 2.9 Hz, 1H), 3.28 – 3.23 (m, 2H), 2.45 (s, 3H), 2.32 (dt, *J* = 11.1, 2.9 Hz, 1H), 2.17 – 2.09 (m, 1H), 1.80 – 1.68 (m, 1H), 1.55 – 1.45 (m, 1H), 1.45 – 1.34 (m, 1H). $^{13}$C NMR (126 MHz, $CD_3OD$) δ 147.9, 146.9, 146.3, 128.3, 119.7, 118.0, 114.9, 108.3, 55.3, 55.3, 55.0, 42.6, 32.2, 23.9, 20.6. HRMS (ESI+, m/z) calcd for $C_{15}H_{20}NO_2$ $(M+H)^+$ : 246.1489, found: 246.1465.

**Synthesis of #013**

・**Synthesis of Imine**

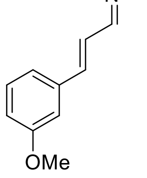

General procedure A was followed using 2.0 M methylamine in THF (1.67 mL, 3.33 mmol, 2.0 equiv),

($E$)-3-methoxycinnamaldehyde (270 mg, 1.66 mmol, 1.0 equiv), molecular sieves 4 Å, and $CH_2Cl_2$ (1.67 mL). Imine (285 mg, 98% yield) was obtained as a yellow solid. $^1$H NMR (400 MHz, $CDCl_3$) δ 8.00 (dd, $J$ = 8.3, 1.6 Hz, 1H), 7.28 (t, $J$ = 7.9 Hz, 1H), 7.08 (d, $J$ = 7.8 Hz, 1H), 7.05 – 6.95 (m, 3H), 6.89 (dd, $J$ = 8.3, 2.5 Hz, 1H), 3.81 (s, 3H), 3.44 (d, $J$ = 1.6 Hz, 3H).

・**Key Reaction**

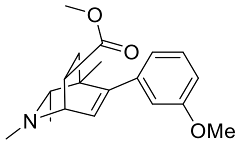

**(±)-#013**

**Final Compound #013** General procedure B was followed using above obtained imine (140 mg, 0.799 mmol, 1.0 equiv) and 2-butyne (313 µL, 3.99 mmol, 5 equiv) in THF (0.8 mL). Rh catalyst (10 mol %, 0.8 mL, 0.08 mmol, 100 mM in THF) was added, and the reaction was carried out at 65 °C for 12 h to produce the DHP intermediate. Following general procedure C, methyl acrylate (0.72 mL, 7.99 mmol, 10 equiv) was added to the solution of DHP intermediate, and the reaction was carried out at 50 °C for 16 h. The resulting crude material was purified by column chromatography (50% EtOAc/Hexane+1% $Et_3N$) to yield the racemic isoquinuclidine (72 mg, 29% yield) as a pale-yellow oil. $^1$H NMR (500 MHz, $CD_3OD$) δ 7.21 (t, $J$ = 7.9 Hz, 1H), 6.83 (dd, $J$ = 8.3, 2.6 Hz, 1H), 6.71 (d, $J$ = 7.7 Hz, 1H), 6.70 – 6.64 (m, 1H), 6.15 (d, $J$ = 6.3 Hz, 1H), 3.77 (s, 3H), 3.66 – 3.64 (m, 4H), 3.35 (ddd, $J$ = 9.5, 4.9, 2.9 Hz, 1H), 2.47 (s, 3H), 2.07 (q, $J$ = 6.3 Hz, 1H), 1.92 (dd, $J$ = 12.9, 4.9 Hz, 1H), 1.61 (dd, $J$ = 12.9, 9.5 Hz, 1H), 0.98 (s, 3H), 0.97 (d, $J$ = 6.3 Hz, 3H). $^{13}$C NMR (126 MHz, $CD_3OD$) δ 174.8, 159.3, 148.3, 141.1, 128.6, 128.1, 120.3, 113.9, 111.9, 65.1, 56.4, 54.2, 51.0, 40.7, 39.4, 36.7, 36.3, 20.3, 17.4. HRMS (ESI+, m/z) calcd for $C_{19}H_{26}NO_3$ $(M+H)^+$ : 316.1907, found: 316.1877.

**Synthesis of #014**

・**Synthesis of Imine**

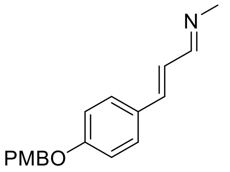

General procedure A was followed using 2.0 M methylamine in THF (0.56 mL, 1.12 mmol, 2.0 equiv), ($E$)-3-(4-(4-methoxybenzyloxy)phenyl)acrylaldehyde (150 mg, 0.559 mmol, 1.0 equiv), molecular sieves 4 Å, and $CH_2Cl_2$ (0.56 mL). Imine (150 mg, 95% yield) was obtained as a yellow solid. $^1$H NMR (400 MHz, $CDCl_3$) δ 7.97 (d, $J$ = 8.5 Hz, 1H), 7.41 (d, $J$ = 8.7 Hz, 2H), 7.34 (d, $J$ = 8.6 Hz, 2H), 7.01 – 6.86 (m, 6H), 5.00 (s, 2H), 3.80 (s, 3H), 3.40 (s, 3H).

・**Key Reaction**

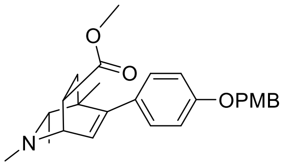

General procedure B was followed using above obtained imine (150 mg, 0.533 mmol, 1.0 equiv) and 2-butyne (209 μL, 2.67 mmol, 5 equiv) in THF (0.53 mL). Rh catalyst (10 mol %, 0.53 mL, 0.053 mmol, 100 mM in THF) was added, and the reaction was carried out at 65 °C for 12 h to produce the DHP intermediate. Following general procedure C, methyl acrylate (0.48 mL, 5.33 mmol, 10 equiv) was added to the solution of DHP intermediate, and the reaction was carried out at 50 °C for 16 h. The resulting crude material was purified by column chromatography (50% EtOAc/Hexane+1% $Et_3N$) to afford the desired product (110 mg, 49% yield from imine) as a pale-yellow oil. $^1$H NMR (400 MHz, $CD_3OD$) δ 7.33 (d, $J$ = 8.6 Hz, 2H), 7.04 (d, $J$ = 8.6 Hz, 2H), 6.90 (dd, $J$ = 8.7, 2.9 Hz, 4H), 6.10 (d, $J$ = 6.3 Hz, 1H), 4.96 (s, 2H), 3.77 (s, 3H), 3.67 – 3.61 (m, 4H), 3.37 – 3.32 (m, 1H), 2.46 (s, 3H), 2.07 (q, $J$ = 6.3 Hz, 1H), 1.88 (dd, $J$ = 12.9, 4.9 Hz, 1H), 1.60 (dd, $J$ = 12.9, 9.5 Hz, 1H), 0.98 (s, 3H), 0.95 (d, $J$ = 6.3 Hz, 3H).

・**Deprotection**

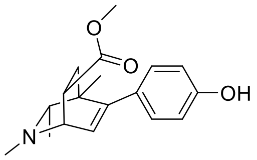

**(±)-#014**

**Final Compound #014** General Procedure D was followed using protected isoquinuclidine (60 mg, 0.142 mmol, 1.0 equiv) in $CH_2Cl_2$ (1.5 mL) and was reacted with trifluoroacetic acid (106 μL, 1.42 mmol, 10.0 equiv) for 12 h. The crude product was purified by preparative thin-layer chromatography (10% MeOH/$CH_2Cl_2$ + 1% $NH_4OH$) to yield the racemic isoquinuclidine (42 mg, 98% yield) as a pale-yellow oil. $^1$H NMR (400 MHz, $CD_3OD$) δ 6.95 (d, $J$ = 8.5 Hz, 2H), 6.70 (d, $J$ = 8.5 Hz, 2H), 6.07 (d, $J$ = 6.3 Hz, 1H), 3.64 (s, 3H), 3.61 (dd, $J$ = 6.3, 3.0 Hz, 1H), 3.36 – 3.30 (m, 1H), 2.44 (s, 3H), 2.04 (q, $J$ = 6.3 Hz, 1H), 1.87 (dd, $J$ = 12.9, 4.9 Hz, 1H), 1.58 (dd, $J$ = 12.9, 9.4 Hz, 1H), 0.97 (s, 3H), 0.94 (d, $J$ = 6.3 Hz, 3H). $^{13}$C NMR (101 MHz, $CD_3OD$) δ 174.9, 156.4, 148.4, 130.9, 129.0, 127.3, 114.3, 65.1, 56.4, 51.0, 40.8, 39.4, 36.7, 36.2, 20.4, 17.4. HRMS (ESI+, m/z) calcd for $C_{18}H_{24}NO_3$ $(M+H)^+$ : 302.1751, found: 302.1721.

**Synthesis of #015**

・**Key Reaction**

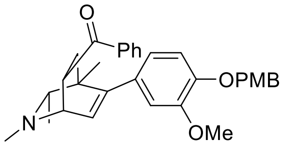

General procedure B was followed using above obtained imine (150 mg, 0.482 mmol, 1.0 equiv) and 2-butyne (189 μL, 2.41 mmol, 5 equiv) in THF (0.48 mL). Rh catalyst (10 mol %, 0.48 mL, 0.048 mmol, 100 mM in THF) was added, and the reaction was carried out at 65 °C for 12 h to produce the DHP intermediate. Following general procedure C, phenylvinyl ketone (318 mg, 2.41 mmol, 5 equiv) was added to the solution of DHP intermediate, and the reaction was carried out at 50 °C for 16 h. The resulting crude material was purified by column chromatography (50% EtOAc/Hexane+1% $Et_3N$) to afford the desired product (25 mg, 10% yield from imine) as a pale-yellow oil. $^1$H NMR (400 MHz, $CD_3OD$) δ 7.96 (d, $J$ = 7.1 Hz, 2H), 7.59 (dd, $J$ = 8.4, 6.3 Hz, 1H), 7.55 – 7.45 (m, 2H), 7.34 (d, $J$ = 8.6 Hz, 2H), 6.92 (d, $J$ = 8.4 Hz, 1H), 6.89 (d, $J$ = 8.6 Hz, 2H), 6.74 (d, $J$ = 2.0 Hz, 1H), 6.67 (dd, $J$ = 8.2, 2.0 Hz, 1H), 6.02 (d, $J$ = 6.3 Hz, 1H), 4.98 (s, 2H), 4.41 (ddd, $J$ = 8.7, 5.4, 2.5 Hz, 1H), 3.81 (s, 3H), 3.77 (s, 3H), 3.61 (dd, $J$ = 6.4, 2.5 Hz, 1H), 2.63 (s, 3H), 2.23 (q, $J$ = 6.3 Hz, 1H), 1.99 (dd, $J$ = 12.6, 5.4 Hz, 1H), 1.71 (dd, $J$ = 12.6, 9.2 Hz, 1H), 1.03 (s, 3H), 0.99 (d, $J$ = 6.3 Hz, 3H).

・**Deprotection**

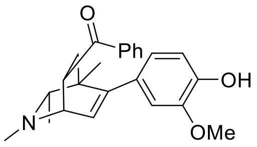

**(±)-#015**

**Final Compound #015** General Procedure D was followed using protected isoquinuclidine (25 mg, 0.05 mmol, 1.0 equiv) in $CH_2Cl_2$ (1.0 mL) and was reacted with trifluoroacetic acid (37 μL, 0.50 mmol, 10.0 equiv) for 12 h. The crude product was purified by preparative thin-layer chromatography (10% MeOH/$CH_2Cl_2$ + 1% $NH_4OH$) to yield the racemic isoquinuclidine (18 mg, 95% yield) as a pale-yellow oil. $^1$H NMR (500 MHz, $CD_3OD$) δ 7.97 (d, $J$ = 7.1 Hz, 2H), 7.68 – 7.57 (m, 1H), 7.52 (dd, $J$ = 8.4, 7.1 Hz, 2H), 6.74 (d, $J$ = 8.1 Hz, 1H), 6.70 (d, $J$ = 2.0 Hz, 1H), 6.61 (dd, $J$ = 8.1, 2.0 Hz, 1H), 6.00 (d, $J$ = 6.4 Hz, 1H), 4.47 – 4.35 (m, 1H), 3.84 (s, 3H), 3.60 (dd, $J$ = 6.4, 2.5 Hz, 1H), 2.63 (s, 3H), 2.21 (q, $J$ = 6.3 Hz, 1H), 2.01 (dd, $J$ = 12.6, 5.3 Hz, 1H), 1.69 (dd, $J$ = 12.6, 9.2 Hz, 1H), 1.04 (s, 3H), 1.00 (d, $J$ = 6.3 Hz, 3H). $^{13}$C NMR (126 MHz, $CD_3OD$) δ 200.7, 147.6, 146.9, 145.6, 136.5, 132.8, 131.4, 128.5, 127.9, 127.4, 120.8, 114.4, 111.9, 65.2, 57.1, 54.9, 41.2, 40.4, 39.3, 36.4, 20.5, 17.3. HRMS (ESI+, m/z) calcd for $C_{24}H_{28}NO_3$ $(M+H)^+$ : 378.2064, found: 378.2026.

**Synthesis of #016**

・**Synthesis of Imine**

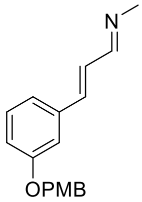

General procedure A was followed using 2.0 M methylamine in THF (0.73 mL, 1.45 mmol, 2.0 equiv), (*E*)-3-[3-(4-methoxy-benzyloxy)-phenyl]-propenal (195 mg, 0.727 mmol, 1.0 equiv), molecular sieves 4 Å, and $CH_2Cl_2$ (0.73 mL). Imine (200 mg, 98% yield) was obtained as a yellow solid. $^1$H NMR (400 MHz, $CDCl_3$) δ 8.00 (d, *J* = 7.3 Hz, 1H), 7.34 (d, *J* = 8.3 Hz, 2H), 7.29 – 7.21 (m, 1H), 7.09 – 7.01 (m, 2H), 6.95 – 6.82 (m, 5H), 4.98 (s, 2H), 3.80 (s, 3H), 3.41 (d, *J* = 1.5 Hz, 3H).

・**Key Reaction**

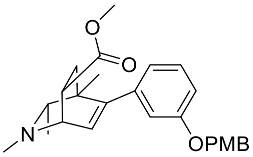

General procedure B was followed using above obtained imine (200 mg, 0.711 mmol, 1.0 equiv) and 2-butyne (279 µL, 3.55 mmol, 5 equiv) in THF (0.71 mL). Rh catalyst (10 mol %, 0.71 mL, 0.071 mmol, 100 mM in THF) was added, and the reaction was carried out at 65 °C for 12 h to produce the DHP intermediate. Following general procedure C, methyl acrylate (0.64 mL, 7.11 mmol, 10 equiv) was added to the solution of DHP intermediate, and the reaction was carried out at 50 °C for 16 h. The resulting crude material was purified by column chromatography (50% EtOAc/Hexane+1% $Et_3N$) to afford the desired product (155 mg, 52% yield from imine) as a pale-yellow oil. $^1$H NMR (400 MHz, $CD_3OD$) δ 7.32 (d, *J* = 8.6 Hz, 2H), 7.18 (t, *J* = 8.1 Hz, 1H), 6.95 – 6.82 (m, 3H), 6.74 – 6.61 (m, 2H), 6.11 (d, *J* = 6.3 Hz, 1H), 4.97 (s, 2H), 3.76 (s, 3H), 3.70 – 3.59 (m, 4H), 3.41 – 3.33 (m, 1H), 2.45 (s, 3H), 2.05 (q, *J* = 6.3 Hz, 1H), 1.88 (dd, *J* = 13.0, 4.9 Hz, 1H), 1.59 (dd, *J* = 13.0, 9.4 Hz, 1H), 0.93 (d, *J* = 6.3 Hz, 3H), 0.91 (s, 3H).

・**Deprotection**

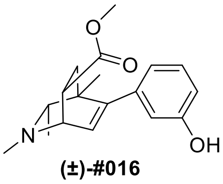

**(±)-#016**

**Final Compound #016** General Procedure D was followed using protected isoquinuclidine (30 mg,

0.071 mmol, 1.0 equiv) in $CH_2Cl_2$ (1.0 mL) and was reacted with trifluoroacetic acid (53 μL, 0.71 mmol, 10.0 equiv) for 12 h. The crude product was purified by preparative thin-layer chromatography (10% MeOH/$CH_2Cl_2$ + 1% $NH_4OH$) to yield the racemic isoquinuclidine (21 mg, 98% yield) as a pale-yellow oil. $^1$H NMR (400 MHz, $CD_3OD$) δ 7.09 (t, $J$ = 7.9 Hz, 1H), 6.72 – 6.63 (m, 1H), 6.58 (dd, $J$ = 7.2, 1.5 Hz, 2H), 6.11 (d, $J$ = 6.3 Hz, 1H), 3.69 – 3.56 (m, 4H), 3.37 – 3.30 (m, 1H), 2.45 (s, 3H), 2.05 (q, $J$ = 6.4 Hz, 1H), 1.89 (dd, $J$ = 12.9, 4.8 Hz, 1H), 1.59 (dd, $J$ = 12.9, 9.4 Hz, 1H), 0.97 (s, 3H), 0.95 (d, $J$ = 6.4 Hz, 3H). $^{13}$C NMR (101 MHz, $CD_3OD$) δ 174.8, 156.7, 148.5, 141.0, 128.5, 127.7, 119.2, 114.8, 113.7, 65.1, 56.3, 51.0, 40.7, 39.4, 36.7, 36.3, 20.3, 17.4. HRMS (ESI+, m/z) calcd for $C_{18}H_{24}NO_3$ $(M+H)^+$ : 302.1751, found: 302.1756.

**Synthesis of #017**

・**Synthesis of Imine**

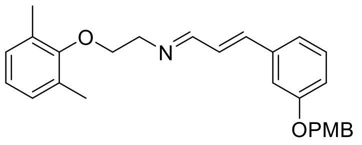

General procedure A was followed using 2-(2,6-dimethylphenoxy)ethanamine (68 mg, 0.410 mmol, 1.1 equiv), (*E*)-3-[3-(4-methoxy-benzyloxy)-phenyl]-propenal (100 mg, 0.373 mmol, 1.0 equiv), molecular sieves 4 Å, and $CH_2Cl_2$ (0.75 mL). Imine (154 mg, 99% yield) was obtained as a pale-yellow oil. $^1$H NMR (400 MHz, $CDCl_3$) δ 8.15 (d, $J$ = 6.0 Hz, 1H), 7.35 (dd, $J$ = 8.7, 2.0 Hz, 2H), 7.30 – 7.25 (m, 1H), 7.14 – 7.04 (m, 2H), 7.04 – 6.81 (m, 8H), 4.99 (s, 2H), 4.02 (t, $J$ = 5.2 Hz, 2H), 3.91 (t, $J$ = 5.2 Hz, 2H), 3.81 (s, 3H), 2.25 (s, 6H).

・**Key Reaction**

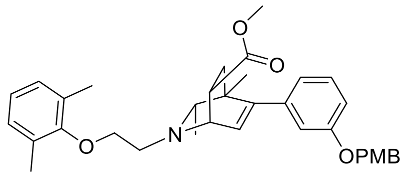

General procedure B was followed using above obtained imine (155 mg, 0.373 mmol, 1.0 equiv) and 2-butyne (146 μL, 1.87 mmol, 5 equiv) in THF (0.37 mL). Rh catalyst (10 mol %, 0.37 mL, 0.037 mmol, 100 mM in THF) was added, and the reaction was carried out at 65 °C for 12 h to produce the DHP intermediate. Following general procedure C, methyl acrylate (0.34 mL, 3.73 mmol, 10 equiv) was added to the solution of DHP intermediate, and the reaction was carried out at 50 °C for 16 h. The resulting crude material was purified by column chromatography (50% EtOAc/Hexane+1% $Et_3N$) to afford the desired product (90 mg, 43% yield from imine) as a pale-yellow oil. $^1$H NMR (400 MHz, $CD_3OD$) δ 7.32 (d, $J$ = 8.6 Hz, 2H), 7.19 (t, $J$ = 8.1 Hz, 1H), 6.98 (d, $J$ = 7.4 Hz, 2H), 6.94 – 6.82 (m,

4H), 6.76 – 6.66 (m, 2H), 6.10 (d, $J$ = 6.3 Hz, 1H), 4.98 (s, 2H), 4.03 – 3.85 (m, 3H), 3.76 (s, 3H), 3.64 (s, 3H), 3.40 (dt, $J$ = 9.5, 4.0 Hz, 1H), 3.17 – 2.99 (m, 2H), 2.28 (s, 6H), 2.24 (q, $J$ = 6.3 Hz, 1H), 1.94 (dd, $J$ = 13.1, 4.4 Hz, 1H), 1.63 (dd, $J$ = 13.1, 9.5 Hz, 1H), 0.97 (d, $J$ = 6.3 Hz, 3H), 0.95 (s, 3H).

・**Deprotection**

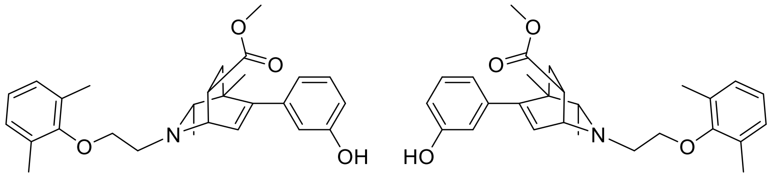

**#017**
enantiomers separated by chiral HPLC

**Final Compound #017** General Procedure D was followed using protected isoquinuclidine (90 mg, 0.162 mmol, 1.0 equiv) in $CH_2Cl_2$ (1.6 mL) and was reacted with trifluoroacetic acid (120 µL, 1.62 mmol, 10.0 equiv) for 12 h. The crude product was purified by preparative thin-layer chromatography (10% MeOH/$CH_2Cl_2$ + 1% $NH_4OH$) to yield the racemic isoquinuclidine (55 mg, 78% yield) as a pale-yellow oil. $^1$H NMR (400 MHz, $CD_3OD$) δ 7.09 (t, $J$ = 8.1 Hz, 1H), 6.97 (d, $J$ = 7.4 Hz, 2H), 6.87 (dd, $J$ = 8.1, 6.8 Hz, 1H), 6.71 – 6.64 (m, 1H), 6.59 (dd, $J$ = 6.8, 1.5 Hz, 2H), 6.10 (d, $J$ = 6.3 Hz, 1H), 4.04 – 3.82 (m, 3H), 3.63 (s, 3H), 3.39 (ddd, $J$ = 9.4, 4.4, 3.1 Hz, 1H), 3.15 – 2.99 (m, 2H), 2.28 (s, 6H), 2.23 (q, $J$ = 6.3 Hz, 1H), 1.95 (dd, $J$ = 13.0, 4.4 Hz, 1H), 1.63 (dd, $J$ = 13.0, 9.4 Hz, 1H), 1.00 (s, 3H), 0.99 (d, $J$ = 6.3 Hz, 3H). $^{13}$C NMR (126 MHz, $CD_3OD$) δ 174.9, 156.6, 155.9, 148.8, 141.2, 130.4, 129.4, 128.5, 127.4, 123.6, 119.2, 114.8, 113.6, 70.9, 65.3, 54.8, 54.1, 51.0, 40.8, 37.7, 35.7, 20.6, 19.1, 15.3. HRMS (ESI+, m/z) calcd for $C_{27}H_{34}NO_4$ $(M+H)^+$ : 436.2482, found: 436.2495.

**HPLC Traces of #017**

A portion of this material was separated using semi-preparative chiral HPLC (Chiralpak IC column, 250 x 10 mm, 10% *i*PrOH/Hexanes+1% DEA, 2.5 mL/min) to provide the two enantiomers with tr = 26.5 min and 55.5 min.

Chiralpak IC, analytical column (250 x 4.6 mm), 10% *i*PrOH/Hexanes+0.1% DEA, 1 mL/min, 254 nm

**Racemic**

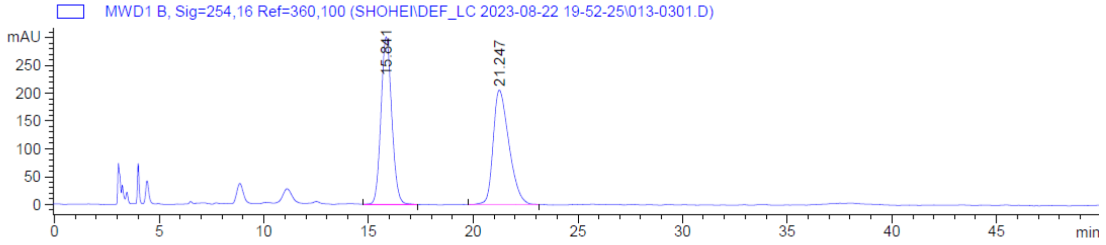

**Enantiomer 1**

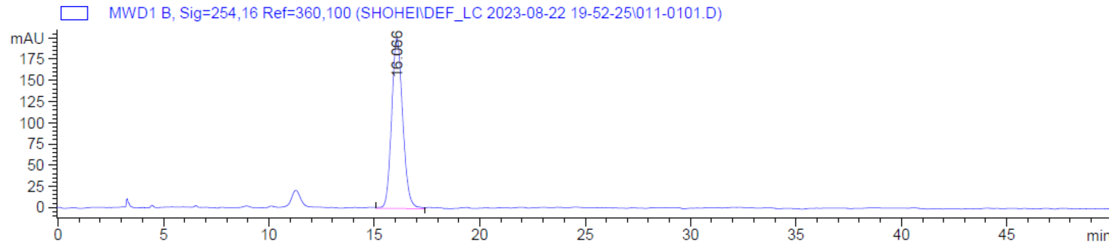

**Enantiomer 2**

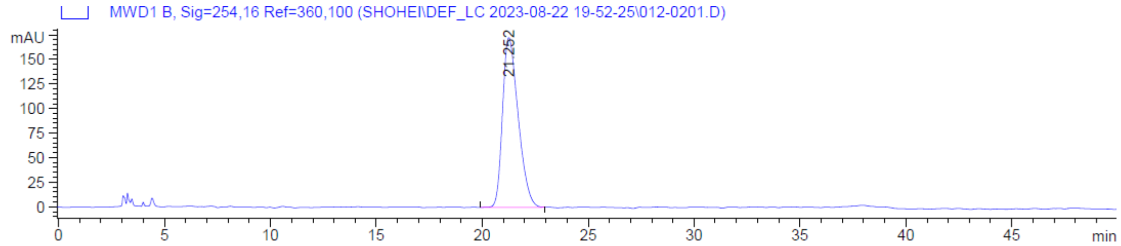

**Synthesis of #018**

・**Synthesis of Imine**

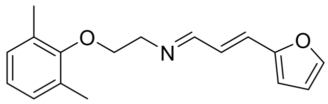

General procedure A was followed using 2-(2,6-dimethylphenoxy)ethanamine (127 mg, 0.766 mmol, 1.1 equiv), (*E*)-3-(2-furyl)propenal (85 mg, 0.696 mmol, 1.0 equiv), molecular sieves 4 Å, and THF (1.4 mL). Imine (187 mg, 99% yield) was obtained as a pale-yellow oil. $^{1}$H NMR (400 MHz, $CDCl_3$) δ 8.07 (dd, *J* = 8.1, 1.4 Hz, 1H), 7.44 (d, *J* = 1.8 Hz, 1H), 6.97 (d, *J* = 7.1 Hz, 2H), 6.89 (dd, *J* = 8.2, 6.6 Hz, 1H), 6.85 – 6.69 (m, 2H), 6.48 (d, *J* = 3.4 Hz, 1H), 6.43 (dd, *J* = 3.4, 1.8 Hz, 1H), 4.01 (t, *J* = 5.5 Hz, 2H), 3.88 (t, *J* = 5.5 Hz, 2H), 2.25 (s, 6H).

・**Key Reaction**

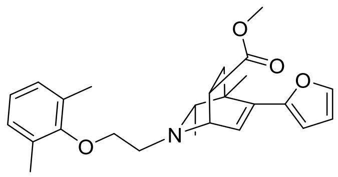

**(±)-#018**

**Final Compound #018** General procedure B was followed using above obtained imine (187 mg, 0.694 mmol, 1.0 equiv) and 2-butyne (272 µL, 3.47 mmol, 5 equiv) in THF (0.7 mL). Rh catalyst (10 mol %, 0.7 mL, 0.07 mmol, 100 mM in THF) was added, and the reaction was carried out at 65 °C for 12 h to produce the DHP intermediate. Following general procedure C, methyl acrylate (0.63 mL, 6.94 mmol, 10 equiv) was added to the solution of DHP intermediate, and the reaction was carried out at 50 °C for 16 h. The resulting crude material was purified by column chromatography (50% EtOAc/Hexane+1% $Et_3N$) to afford the desired product (101 mg, 36% yield from imine) as a pale-yellow oil. $^1$H NMR (500 MHz, $CD_3OD$) δ 7.41 (d, *J* = 1.8 Hz, 1H), 6.98 (d, *J* = 7.5 Hz, 2H), 6.88 (t, *J* = 7.5 Hz, 1H), 6.59 (d, *J* = 6.5 Hz, 1H), 6.43 – 6.31 (m, 2H), 4.02 (dd, *J* = 6.6, 3.0 Hz, 1H), 3.98 – 3.79 (m, 2H), 3.61 (s, 3H), 3.44 – 3.35 (m, 1H), 3.17 – 2.95 (m, 2H), 2.28 (s, 6H), 2.22 (q, *J* = 6.3 Hz, 1H), 1.91 (dd, *J* = 13.1, 4.7 Hz, 1H), 1.62 (dd, *J* = 13.1, 9.7 Hz, 1H), 1.34 (s, 3H), 0.89 (d, *J* = 6.3 Hz, 3H). $^{13}$C NMR (126 MHz, $CD_3OD$) δ 174.7, 155.9, 152.2, 141.6, 136.8, 130.4, 128.5, 127.9, 126.9, 123.6, 119.3, 110.6, 107.4, 71.0, 65.1, 54.3, 53.8, 51.0, 40.3, 37.1, 35.7, 20.9, 18.6, 15.3. HRMS (ESI+, m/z) calcd for $C_{25}H_{32}NO_4$ $(M+H)^+$ : 410.2326, found: 410.2337.

**Synthesis of #019**

・**Synthesis of Imine**

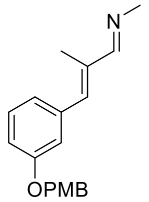

General procedure A was followed using 2.0 M methylamine in THF (3.72 mL, 7.44 mmol, 5.0 equiv), above obtained aldehyde (420 mg, 1.49 mmol, 1.0 equiv), molecular sieves 4 Å, and $CH_2Cl_2$ (3.0 mL) at 50 °C. Imine (435 mg, 99% yield) was obtained as a yellow solid. $^1$H NMR (400 MHz, $CDCl_3$) δ 7.96 (d, *J* = 1.7 Hz, 1H), 7.35 (d, *J* = 8.5 Hz, 2H), 7.28 (t, *J* = 8.1 Hz, 1H), 7.03 – 6.93 (m, 2H), 6.93 – 6.86 (m, 3H), 6.73 (s, 1H), 4.99 (s, 2H), 3.80 (s, 3H), 3.43 (s, 3H), 2.08 (s, 3H).

・**Key Reaction**

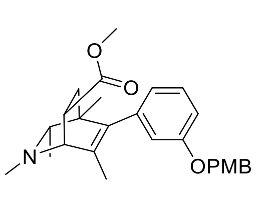

General procedure B was followed using above obtained imine (155 mg, 0.525 mmol, 1.0 equiv) and 2-butyne (206 µL, 2.62 mmol, 5 equiv) in THF (0.53 mL). Rh catalyst (10 mol %, 0.53 mL, 0.053 mmol, 100 mM in THF) was added, and the reaction was carried out at 65 °C for 12 h to produce the DHP intermediate. Following general procedure C, methyl acrylate (0.47 mL, 5.25 mmol, 10 equiv) was added to the solution of DHP intermediate, and the reaction was carried out at 50 °C for 16 h. The resulting crude material was purified by column chromatography (50% EtOAc/Hexane+1% $Et_3N$) to afford the desired product (65 mg, 28% yield from imine) as a pale-yellow oil. $^1$H NMR (400 MHz, $CD_3OD$) δ 7.31 (d, $J$ = 8.1 Hz, 2H), 7.20 (t, $J$ = 7.9 Hz, 1H), 6.96 – 6.78 (m, 3H), 6.69 – 6.52 (m, 2H), 4.97 (s, 2H), 3.75 (s, 3H), 3.66 (s, 3H), 3.45 (d, $J$ = 3.0 Hz, 1H), 3.36 – 3.30 (m, 1H), 2.46 (s, 3H), 2.03 (q, $J$ = 6.3 Hz, 1H), 1.90 (dd, $J$ = 12.9, 4.7 Hz, 1H), 1.61 – 1.45 (m, 4H), 0.94 (d, $J$ = 6.3 Hz, 3H), 0.72 (s, 3H).

・**Deprotection**

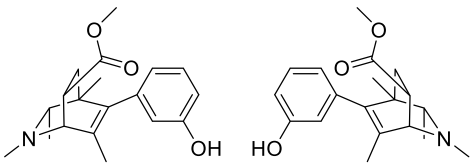

**#019**

enantiomers separated by chiral HPLC

**Final Compound #019** General Procedure D was followed using protected isoquinuclidine (65 mg, 0.149 mmol, 1.0 equiv) in $CH_2Cl_2$ (1.5 mL) and was reacted with trifluoroacetic acid (111 μL, 1.49 mmol, 10.0 equiv) for 12 h. The crude product was purified by preparative thin-layer chromatography (10% MeOH/$CH_2Cl_2$ + 1% $NH_4OH$) to yield the racemic isoquinuclidine (33 mg, 70% yield) as a pale-yellow oil. $^1$H NMR (500 MHz, $CD_3OD$) δ 7.13 (t, $J$ = 7.8 Hz, 1H), 6.67 (d, $J$ = 8.1 Hz, 1H), 6.59 – 6.42 (m, 2H), 3.68 (s, 3H), 3.48 (d, $J$ = 2.4 Hz, 1H), 3.40 – 3.34 (m, 1H), 2.49 (s, 3H), 2.07 (q, $J$ = 6.3 Hz, 1H), 1.94 (dd, $J$ = 12.9, 4.7 Hz, 1H), 1.64 – 1.53 (m, 4H), 1.00 (d, $J$ = 6.3 Hz, 3H), 0.82 (s, 3H). $^{13}$C NMR (126 MHz, $CD_3OD$, 50 °C) δ 174.7, 156.7, 139.9, 139.6, 134.6, 128.5, 120.0, 115.6, 113.3, 65.8, 61.9, 50.8, 40.6, 39.6, 36.9, 36.6, 20.3, 17.4, 16.5. HRMS (ESI+, m/z) calcd for $C_{19}H_{26}NO_3$ $(M+H)^+$ : 316.1907, found: 316.1915.

**HPLC Traces of #019**

A portion of this material was separated using semi-preparative chiral HPLC (Chiralpak IC column, 250 x 10 mm, 5% *i*PrOH/Hexanes+1% DEA, 2.5 mL/min) to provide the two enantiomers with tr = 15 min and 25.5 min.

Chiralpak IC, analytical column (250 x 4.6 mm), 5% *i*PrOH/Hexanes+0.1% DEA, 1 mL/min, 210 nm

**Racemic**

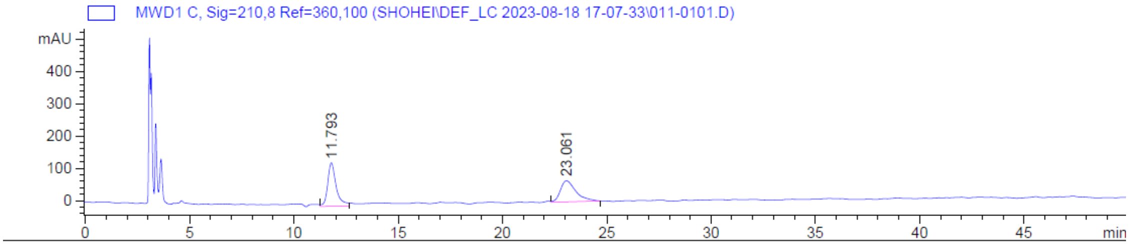

**Enantiomer 1**

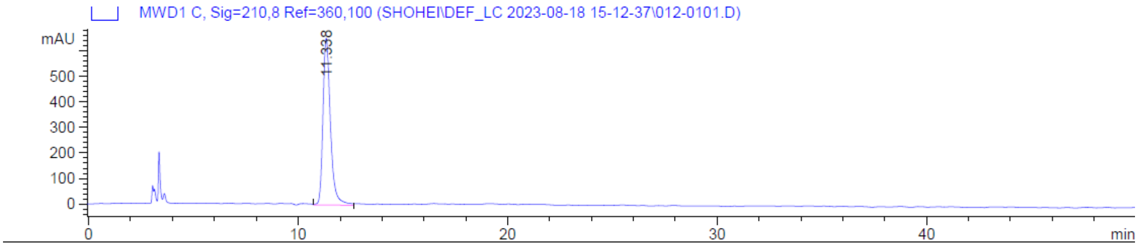

**Enantiomer 2**

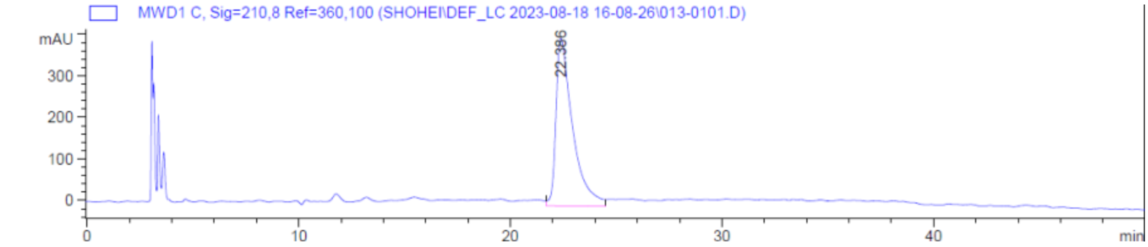

**Synthesis of #020**

・**Synthesis of Imine**

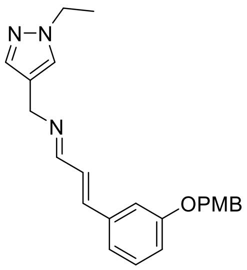

General procedure A was followed using (1-ethylpyrazol-4-yl)methanamine (52 mg, 0.418 mmol, 1.1 equiv), (*E*)-3-[3-(4-Methoxy-benzyloxy)-phenyl]-propenal (102 mg, 0.380 mmol, 1.0 equiv), molecular sieves 4 Å, and $CH_2Cl_2$ (0.76 mL). Imine (137 mg, 97% yield) was obtained as a pale-yellow oil. $^1$H NMR (400 MHz, $CDCl_3$) δ 8.07 (dd, *J* = 6.4, 1.8 Hz, 1H), 7.41 (s, 1H), 7.35 (s, 1H), 7.33 (d, *J* = 1.9 Hz, 2H), 7.27 – 7.23 (m, 1H), 7.08 – 7.02 (m, 2H), 6.95 – 6.87 (m, 5H), 4.98 (s, 2H), 4.57 (s, 2H), 4.13 (q, *J* = 7.3 Hz, 2H), 3.80 (s, 3H), 1.46 (t, *J* = 7.3 Hz, 3H).

・**Key Reaction**

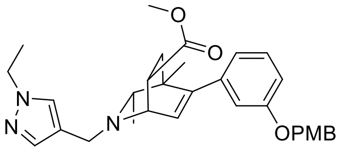

General procedure B was followed using above obtained imine (143 mg, 0.381 mmol, 1.0 equiv) and 2-butyne (149 μL, 1.90 mmol, 5 equiv) in THF (0.38 mL). Rh catalyst (10 mol %, 0.38 mL, 0.038 mmol, 100 mM in THF) was added, and the reaction was carried out at 65 °C for 12 h to produce the DHP intermediate. Following general procedure C, methyl acrylate (0.34 mL, 3.81 mmol, 10 equiv) was added to the solution of DHP intermediate, and the reaction was carried out at 50 °C for 16 h. The resulting crude material was purified by column chromatography (70% EtOAc/Hexane+1% $Et_3N$) to afford the desired product (75 mg, 38% yield from imine) as a pale-yellow oil. $^1H$ NMR (400 MHz, $CD_3OD$) δ 7.62 (s, 1H), 7.50 (s, 1H), 7.30 (d, $J$ = 8.6 Hz, 2H), 7.17 (t, $J$ = 8.1 Hz, 1H), 6.93 – 6.81 (m, 3H), 6.76 – 6.62 (m, 2H), 6.02 (d, $J$ = 6.4 Hz, 1H), 4.95 (s, 2H), 4.13 (q, $J$ = 7.3 Hz, 2H), 3.78 – 3.72 (m, 5H), 3.62 (s, 3H), 3.56 (d, $J$ = 13.3 Hz, 1H), 3.43 – 3.36 (m, 1H), 2.20 (q, $J$ = 6.3 Hz, 1H), 1.92 (dd, $J$ = 13.0, 4.5 Hz, 1H), 1.61 (dd, $J$ = 13.0, 9.3 Hz, 1H), 1.41 (t, $J$ = 7.3 Hz, 3H), 0.92 (s, 3H), 0.84 (d, $J$ = 6.3 Hz, 3H).

・**Deprotection**

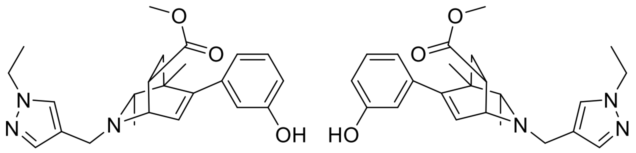

**#020**
enantiomers separated by chiral HPLC

**Final Compound #020** General Procedure D was followed using protected isoquinuclidine (75 mg, 0.145 mmol, 1.0 equiv) in $CH_2Cl_2$ (1.5 mL) and was reacted with trifluoroacetic acid (53 μL, 1.45 mmol, 10.0 equiv) for 12 h. The crude product was purified by preparative thin-layer chromatography (10% MeOH/$CH_2Cl_2$ + 1% $NH_4OH$) to yield the racemic isoquinuclidine (47 mg, 82% yield) as a white solid. $^1H$ NMR (500 MHz, $CD_3OD$) δ 7.66 (s, 1H), 7.53 (s, 1H), 7.10 (t, $J$ = 7.5 Hz, 1H), 6.68 (dd, $J$ = 8.7, 1.7 Hz, 1H), 6.65 – 6.54 (m, 2H), 6.03 (dd, $J$ = 6.4, 1.3 Hz, 1H), 4.17 (q, $J$ = 7.3 Hz, 2H), 3.82 – 3.76 (m, 2H), 3.65 (s, 3H), 3.60 (d, $J$ = 13.2 Hz, 1H), 3.46 – 3.40 (m, 1H), 2.26 (q, $J$ = 6.2 Hz, 1H), 1.96 (dd, $J$ = 13.0, 4.5 Hz, 1H), 1.66 (dd, $J$ = 13.0, 9.4 Hz, 1H), 1.44 (t, $J$ = 7.3Hz, 3H), 1.01 (s, 3H), 0.90 (d, $J$ = 6.2 Hz, 3H). $^{13}C$ NMR (126 MHz, $CD_3OD$) δ 174.9, 156.6, 149.4, 141.1, 139.5, 129.7, 128.5, 127.3, 119.2, 117.5, 114.8, 113.6, 64.4, 52.8, 51.0, 46.8, 46.4, 40.9, 36.9, 35.8, 20.4, 18.5, 14.7. HRMS (ESI+, m/z) calcd for $C_{23}H_{30}N_3O_3$ $(M+H)^+$ : 396.2282, found: 396.2291.

**HPLC Traces of #020**

A portion of this material was separated using semi-preparative chiral HPLC (Chiralpak AD-H column, 250 x 10 mm, 12% *i*PrOH/Hexanes+1% DEA, 2.5 mL/min) to provide the two enantiomers with tr = 35 min and 42 min.

Chiralpak AD-H, analytical column (250 x 4.6 mm), 12% *i*PrOH/Hexanes+0.1% DEA, 1 mL/min, 280 nm

**Racemic**

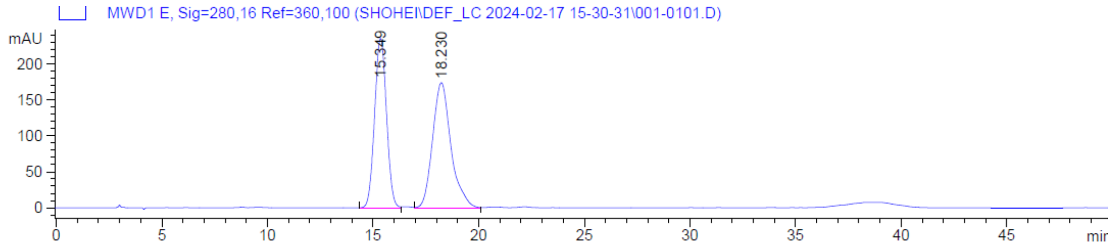

**Enantiomer 1**

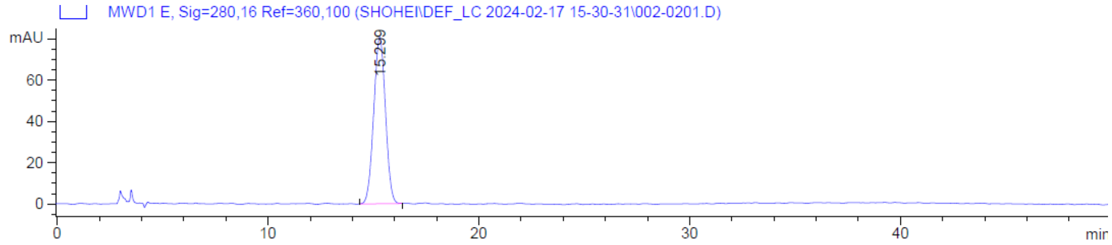

**Enantiomer 2**

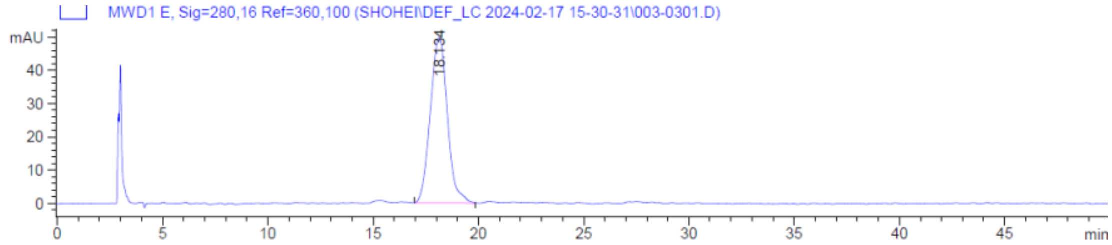

**Synthesis of #021**

・**Synthesis of Imine**

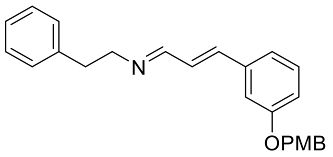

General procedure A was followed using 2-phenylethanamine (50 mg, 0.410 mmol, 1.1 equiv), (*E*)-3-[3-(4-Methoxy-benzyloxy)-phenyl]-propenal (100 mg, 0.373 mmol, 1.0 equiv), molecular sieves 4 Å, and $CH_2Cl_2$ (0.75 mL). Imine (138 mg, 99% yield) was obtained as a pale-yellow oil. $^1$H NMR (400

MHz, $CDCl_3$) δ 7.88 (dd, *J* = 6.7, 1.5 Hz, 1H), 7.34 (d, *J* = 8.7 Hz, 2H), 7.31 – 7.25 (m, 3H), 7.22 – 7.16 (m, 3H), 7.07 – 7.02 (m, 2H), 6.95 – 6.88 (m, 3H), 6.88 – 6.83 (m, 2H), 4.98 (s, 2H), 3.80 (s, 3H), 3.75 (t, *J* = 7.5 Hz, 2H), 2.96 (t, *J* = 7.5 Hz, 2H).

・**Key Reaction**

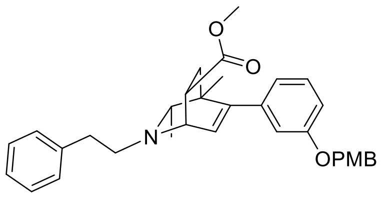

General procedure B was followed using above obtained imine (138 mg, 0.371 mmol, 1.0 equiv) and 2-butyne (146 µL, 1.86 mmol, 5 equiv) in THF (0.37 mL). Rh catalyst (10 mol %, 0.37 mL, 0.037 mmol, 100 mM in THF) was added, and the reaction was carried out at 65 °C for 12 h to produce the DHP intermediate. Following general procedure C, methyl acrylate (0.34 mL, 3.71 mmol, 10 equiv) was added to the solution of DHP intermediate, and the reaction was carried out at 50 °C for 16 h. The resulting crude material was purified by column chromatography (50% EtOAc/Hexane+1% $Et_3N$) to afford the desired product (70 mg, 37% yield from imine) as a pale-yellow oil. $^1$H NMR (400 MHz, $CD_3OD$) δ 7.31 (d, *J* = 8.6 Hz, 2H), 7.29 – 7.12 (m, 6H), 6.98 – 6.80 (m, 3H), 6.79 – 6.62 (m, 2H), 6.09 (d, *J* = 6.3 Hz, 1H), 4.97 (s, 2H), 3.88 (dd, *J* = 6.4, 3.1 Hz, 1H), 3.75 (s, 3H), 3.62 (s, 3H), 3.31 – 3.25 (m, 1H), 2.94 – 2.72 (m, 4H), 2.19 (q, *J* = 6.3 Hz, 1H), 1.91 (dd, *J* = 13.1, 4.3 Hz, 1H), 1.56 (dd, *J* = 13.1, 9.4 Hz, 1H), 0.97 (d, *J* = 6.3 Hz, 3H), 0.93 (s, 3H).

・**Deprotection**

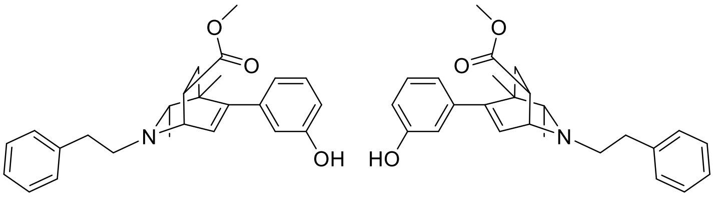

**#021**
enantiomers separated by chiral HPLC

**Final Compound #021** General Procedure D was followed using protected isoquinuclidine (70 mg, 0.137 mmol, 1.0 equiv) in $CH_2Cl_2$ (1.4 mL) and was reacted with trifluoroacetic acid (102 µL, 1.37 mmol, 10.0 equiv) for 12 h. The crude product was purified by preparative thin-layer chromatography (10% MeOH/$CH_2Cl_2$ + 1% $NH_4OH$) to yield the racemic isoquinuclidine (50 mg, 93% yield) as a pale-yellow oil. $^1$H NMR (400 MHz, $CD_3OD$) δ 7.31 – 7.19 (m, 4H), 7.21 – 7.13 (m, 1H), 7.10 (t, *J* = 8.2 Hz, 1H), 6.68 (ddd, *J* = 8.2, 2.4, 1.1 Hz, 1H), 6.64 – 6.56 (m, 2H), 6.09 (d, *J* = 6.4 Hz, 1H), 3.92 (dd, *J* = 6.4, 3.1 Hz, 1H), 3.63 (s, 3H), 3.33 – 3.27 (m, 1H), 2.94 – 2.73 (m, 4H), 2.25 (q, *J* = 6.3 Hz,

1H), 1.93 (dd, $J$ = 13.1, 4.3 Hz, 1H), 1.60 (dd, $J$ = 13.1, 9.4 Hz, 1H), 1.015 (d, $J$ = 6.3 Hz, 3H), 1.01 (s 3H). $^{13}$C NMR (101 MHz, $CD_3OD$) δ 174.7, 156.7, 149.3, 141.0, 139.7, 128.5, 128.4, 128.1, 126.9, 125.9, 119.1, 114.8, 113.7, 64.9, 56.8, 54.3, 51.0, 40.8, 37.4, 35.3, 35.2, 20.5, 19.1. HRMS (ESI+, m/z) calcd for $C_{25}H_{30}NO_3$ $(M+H)^+$ : 392.2220, found: 392.2230.

**HPLC Traces of #021**

A portion of this material was separated using semi-preparative chiral HPLC (Chiralpak IC column, 250 x 10 mm, 15% *i*PrOH/Hexanes+1% DEA, 2.5 mL/min) to provide the two enantiomers with tr = 24 min and 47 min.

Chiralpak IC, analytical column (250 x 4.6 mm), 15% *i*PrOH/Hexanes+0.1% DEA, 1 mL/min, 210 nm

**Racemic**

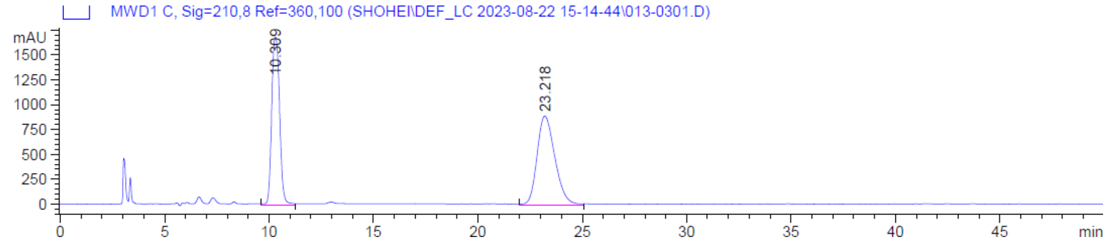

**Enantiomer 1**

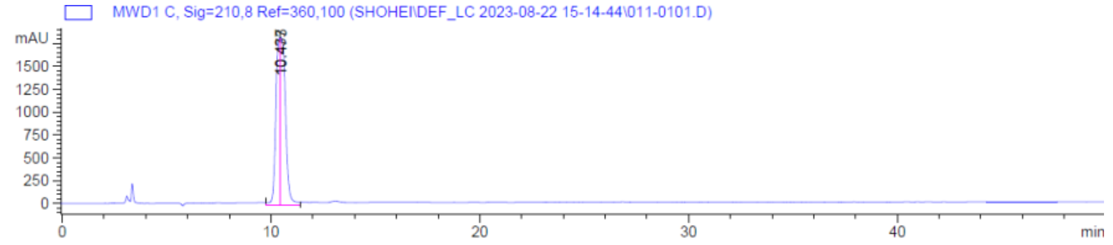

**Enantiomer 2**

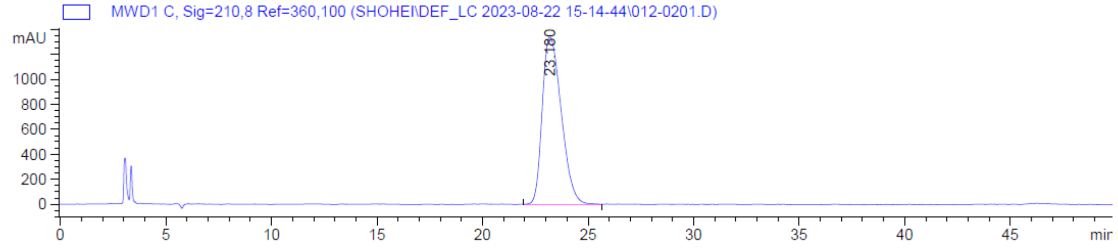

**Synthesis of #022**

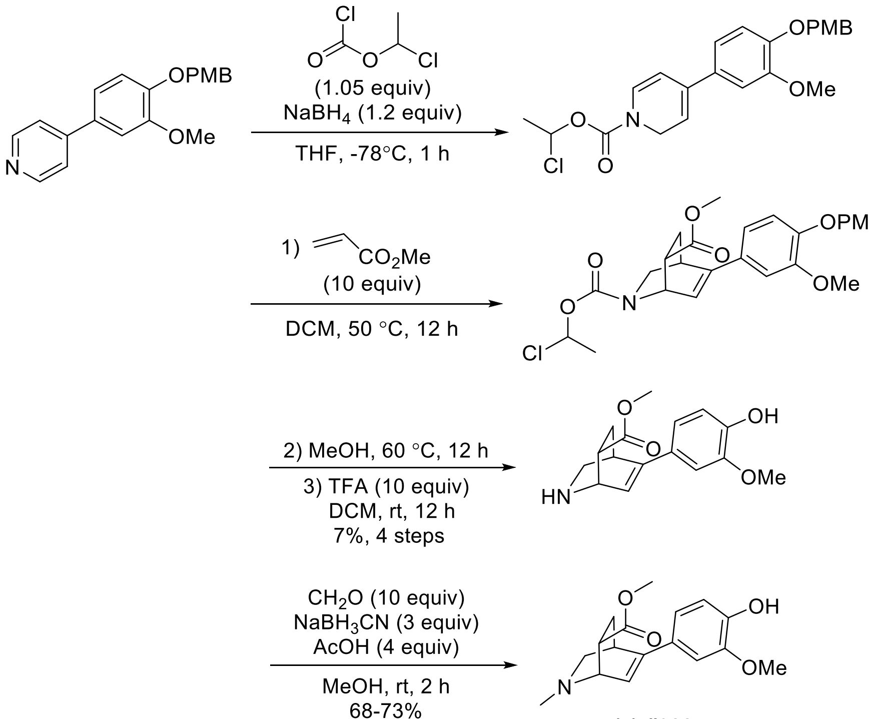

**(±)-#022**

To a flame-dried round-bottom flask was added a magnetic stir bar, 4-[3-methoxy-4-[(4-methoxyphenyl)methoxy]phenyl]pyridine (1620 mg, 5.04 mmol, 1 equiv) and $NaBH_4$ (248 mg, 6.355 mmol, 1.3 equiv) dissolved in dry THF (10 mL). This solution was cooled to -78 °C and then 1-chloroethyl chloroformate (793 mg, 5.55 mmol, 1.1 equiv) was added dropwise over 5 min. The reaction mixture was stirred at -78 ° C for 1 h. The mixture was then diluted with water, extracted with $Et_2O$, washed with brine, dried with $Na_2SO_4$ and filtered. The organic layer was concentrated and the crude residue was purified via silica gel chromatography (gradient of 0-50% EtOAc in hexane) to give 477 mg of the *N*-protected dihydropyridine in 22% yield. To a stirred solution of the above obtained compound (477 mg, 1.11 mmol, 1 equiv) in $CH_2Cl_2$ (0.1 mL) was added methyl acrylate (1.0 mL, 11.1 mmol, 10 equiv) and the reaction mixture was heated at 50 ° C for 12 h. The mixture was concentrated and the crude residue was purified via silica gel chromatography (gradient of 0-50% EtOAc in hexane) to give 170 mg of the *N*-protected isoquinuclidine in 31% yield. The solution of the above obtained compound (170 mg, 0.34 mmol, 1 equiv) in MeOH (34 mL) was heated at 60 °C for 12 h. The mixture was concentrated and diluted with $CH_2Cl_2$ (3.4 mL). Trifluoroacetic acid (253 µL, 3.40 mmol, 10.0 equiv) was added to this solution and stirred at room temperature for 12h. The mixture was concentrated and the crude residue was purified by preparative thin-layer chromatography (10% $MeOH/CH_2Cl_2$ + 1% $NH_4OH$) to yield the separate diastereomers of isoquinuclidines (93 mg, 95% combined yield, 2:1 ratio) as a dark-brown oil.

・**Major diastereomer**

$^{1}H$ NMR (400 MHz, $CD_3OD$) δ 7.03 (d, *J* = 2.0 Hz, 1H), 6.92 (dd, *J* = 8.2, 2.0 Hz, 1H), 6.75 (d, *J* = 8.2 Hz, 1H), 6.59 (dd, *J* = 6.3, 2.0 Hz, 1H), 3.96 (dd, *J* = 6.3, 2.5 Hz, 1H), 3.85 (s, 3H), 3.73 (s, 3H), 3.25 – 3.17 (m, 1H), 3.01 (dd, *J* = 10.2, 2.0 Hz, 1H), 2.62 – 2.48 (m, 2H), 2.14 (ddd, *J* = 13.0, 4.6, 2.5 Hz, 1H), 1.73 – 1.60 (m, 1H).

・**Minor diastereomer**

$^{1}H$ NMR (400 MHz, $CD_3OD$) δ 6.99 (d, *J* = 2.1 Hz, 1H), 6.91 (dd, *J* = 8.2, 2.1 Hz, 1H), 6.74 (d, *J* = 8.2 Hz, 1H), 6.42 (dd, *J* = 6.0, 2.0 Hz, 1H), 4.05 (dd, *J* = 6.0, 2.9 Hz, 1H), 3.84 (s, 3H), 3.62 (s, 3H), 3.28 – 3.24 (m, 1H), 3.09 (ddd, *J* = 9.8, 5.2, 2.9 Hz, 1H), 2.93 (dd, *J* = 10.4, 2.0 Hz, 1H), 2.51 (dt, *J* = 10.4, 2.8 Hz, 1H), 1.99 (ddd, *J* = 12.7, 9.8, 2.8 Hz, 1H), 1.87 – 1.75 (m, 1H).

**Final Compound #022**

・**Major diastereomer**

To a stirred solution of the above obtained major diastereomer (60 mg, 0.207 mmol, 1 equiv) in MeOH (2 mL) was added AcOH (48 µL, 0.830 mmol, 4 equiv), formaldehyde (0.19 mL of 37% w/w solution, 6.22 mmol, 10 equiv), $NaBH_3CN$ (39 mg, 0.622 mmol, 3 equiv) and the reaction mixture was stirred at room temperature for 12 h. After the reaction was complete as monitored by thin-layer chromatography, the mixture was then diluted with water, extracted with $CH_2Cl_2$, washed with brine, dried with $Na_2SO_4$ and filtered. The organic layer was concentrated and the crude residue was purified by preparative thin-layer chromatography (5% MeOH/$CH_2Cl_2$ + 1% $NH_4OH$) to yield the racemic isoquinuclidine (43 mg, 68% yield) as a pale-yellow oil. $^{1}H$ NMR (500 MHz, $CD_3OD$) δ 7.04 (d, *J* = 2.1 Hz, 1H), 6.93 (dd, *J* = 8.3, 2.1 Hz, 1H), 6.77 (d, *J* = 8.3 Hz, 1H), 6.50 (dd, *J* = 5.8, 1.9 Hz, 1H), 3.87 (s, 3H), 3.84 (dd, *J* = 5.8, 2.6 Hz, 1H), 3.73 (s, 3H), 3.18 (dd, *J* = 9.9, 2.3 Hz, 1H), 3.15 – 3.10 (m, 1H), 2.54 (ddd, *J* = 11.4, 4.6, 2.6 Hz, 1H), 2.25 – 2.13 (m, 4H), 1.92 (dt, *J* = 9.9, 2.6 Hz, 1H), 1.57 – 1.47 (m, 1H). $^{13}C$ NMR (126 MHz, $CD_3OD$) δ 174.7, 147.8, 146.4, 145.8, 128.9, 121.2, 117.8, 114.8, 108.2, 57.4, 55.8, 55.0, 51.1, 44.9, 44.2, 33.3, 23.7. HRMS (ESI+, m/z) calcd for $C_{17}H_{22}NO_4$ $(M+H)^+$ : 304.1543, found: 304.1529.

・**Minor diastereomer**

To a stirred solution of the above obtained minor diastereomer (33 mg, 0.114 mmol, 1 equiv) in MeOH (1.2 mL) was added AcOH (26 µL, 0.456 mmol, 4 equiv), formaldehyde (0.1 mL of 37% w/w solution, 3.42 mmol, 10 equiv), $NaBH_3CN$ (22 mg, 0.342 mmol, 3 equiv) and the reaction mixture was stirred at room temperature for 12 h. After the reaction was complete as monitored by thin-layer chromatography, the mixture was then diluted with water, extracted with $CH_2Cl_2$, washed with brine, dried with $Na_2SO_4$ and filtered. The organic layer was concentrated and the crude residue was purified by preparative thin-layer chromatography (5% MeOH/$CH_2Cl_2$ + 1% $NH_4OH$) to yield the racemic isoquinuclidine (25 mg, 72% yield) as a pale-yellow oil. $^1$H NMR (500 MHz, $CD_3OD$) δ 7.01 (d, *J* = 2.1 Hz, 1H), 6.93 (dd, *J* = 8.2, 2.1 Hz, 1H), 6.77 (d, *J* = 8.2 Hz, 1H), 6.37 (dd, *J* = 5.7, 1.9 Hz, 1H), 3.92 – 3.83 (m, 4H), 3.64 (s, 3H), 3.24 – 3.15 (m, 2H), 3.04 (dd, *J* = 10.4, 2.2 Hz, 1H), 2.33 (s, 3H), 2.10 (dt, *J* = 10.4, 2.7 Hz, 1H), 1.90 (ddd, *J* = 12.6, 9.8, 2.7 Hz, 1H), 1.86 – 1.75 (m, 1H). $^{13}$C NMR (126 MHz, $CD_3OD$) δ 173.9, 147.8, 146.6, 146.2, 128.7, 119.6, 117.9, 114.8, 108.3, 56.3, 55.0, 54.6, 51.0, 42.9, 42.6, 33.1, 25.4. HRMS (ESI+, m/z) calcd for $C_{17}H_{22}NO_4$ $(M+H)^+$ : 304.1543, found: 304.1529.

**Synthesis of #023**

・**Synthesis of Imine**

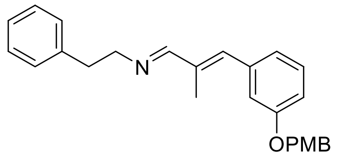

General procedure A was followed using 2-phenylethanamine (71 mg, 0.584 mmol, 1.1 equiv), above obtained aldehyde (150 mg, 0.531 mmol, 1.0 equiv), molecular sieves 4 Å, and $CH_2Cl_2$ (1.0 mL) at 50 °C. Imine (200 mg, 98% yield) was obtained as a pale-yellow oil. $^1$H NMR (400 MHz, $CDCl_3$) δ 7.84 (s, 1H), 7.35 (d, *J* = 8.6 Hz, 2H), 7.28 (m, 3H), 7.23 – 7.14 (m, 3H), 7.01 – 6.94 (m, 2H), 6.94 – 6.87 (m, 3H), 6.67 (s, 1H), 4.99 (s, 2H), 3.80 (s, 3H), 3.80 – 3.74 (m, 2H), 2.95 (t, *J* = 7.6 Hz, 2H), 2.11 (s, 3H).

・**Key Reaction**

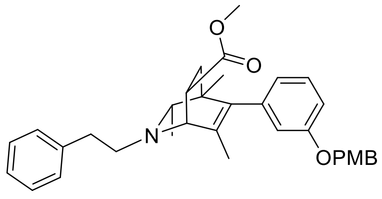

General procedure B was followed using above obtained imine (200 mg, 0.519 mmol, 1.0 equiv) and 2-butyne (203 μL, 2.59 mmol, 5 equiv) in THF (0.52 mL). Rh catalyst (10 mol %, 0.52 mL, 0.052 mmol, 100 mM in THF) was added, and the reaction was carried out at 65 °C for 12 h to produce the DHP intermediate. Following general procedure C, methyl acrylate (0.23 mL, 2.59 mmol, 5 equiv) was added to the solution of DHP intermediate, and the reaction was carried out at 50 °C for 16 h. The resulting crude material was purified by column chromatography (50% EtOAc/Hexane+1% $Et_3N$) to afford the desired product (185 mg, 68% yield from imine) as a pale-yellow oil. $^1$H NMR (400 MHz, $CD_3OD$) δ 7.35 – 7.16 (m, 8H), 6.91 – 6.84 (m, 3H), 6.60 (m, 2H), 4.99 (s, 2H), 3.80 – 3.74 (m, 4H),

3.69 – 3.61 (m, 4H), 2.93 – 2.71 (m, 4H), 2.18 (q, *J* = 6.3 Hz, 1H), 1.94 (dd, *J* = 13.0, 4.2 Hz, 1H), 1.58 – 1.46 (m, 4H), 0.98 (d, *J* = 6.2 Hz, 3H), 0.76 (s, 3H).

・**Deprotection**

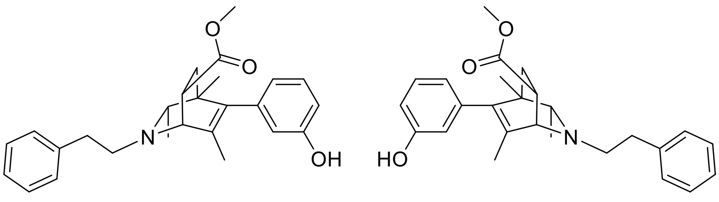

**#023**
enantiomers separated by chiral HPLC

**Final Compound #023** General Procedure D was followed using protected isoquinuclidine (100 mg, 0.190 mmol, 1.0 equiv) in $CH_2Cl_2$ (1.9 mL) and was reacted with trifluoroacetic acid (141 µL, 1.90 mmol, 10.0 equiv) for 12 h. The crude product was purified by preparative thin-layer chromatography (10% MeOH/$CH_2Cl_2$ + 1% $NH_4OH$) to yield the racemic isoquinuclidine (73 mg, 95% yield) as a white solid. $^1$H NMR (500 MHz, $CD_3OD$, 50 °C) δ 7.27 (d, *J* = 7.8 Hz, 1H), 7.26 – 7.22 (m, 3H), 7.20 – 7.14 (m, 1H), 7.12 (t, *J* = 7.8 Hz, 1H), 6.68 (dd, *J* = 7.7, 1.9 Hz, 1H), 6.52 (d, *J* = 8.2 Hz, 2H), 3.69 – 3.63 (m, 4H), 3.30 – 3.26 (m, 1H), 2.93 – 2.76 (m, 4H), 2.17 (q, *J* = 6.3 Hz, 1H), 1.96 (dd, *J* = 13.0, 4.3 Hz, 1H), 1.58 (s, 3H), 1.53 (dd, *J* = 13.0, 9.4 Hz, 1H), 1.00 (d, *J* = 6.3 Hz, 3H), 0.83 (s, 3H). $^{13}$C NMR (126 MHz, $CD_3OD$, 50 °C) δ 174.8, 156.7, 140.3, 140.1, 139.9, 134.0, 128.5, 128.4, 128.0, 125.7, 120.1, 115.7, 113.2, 65.5, 59.8, 56.6, 50.8, 40.8, 37.4, 36.1, 35.5, 20.6, 19.3, 16.5. HRMS (ESI+, m/z) calcd for $C_{26}H_{32}NO_3$ $(M+H)^+$ : 406.2377, found: 406.2390.

**HPLC Traces of #023**

A portion of this material was separated using semi-preparative chiral HPLC (Chiralpak IC column, 250 x 10 mm, 4% *i*PrOH/Hexanes+1% DEA, 2.5 mL/min) to provide the two enantiomers with tr = 15 min and 25 min.

Chiralpak IC, analytical column (250 x 4.6 mm), 4% *i*PrOH/Hexanes+0.1% DEA, 1 mL/min, 210 nm

**Racemic**

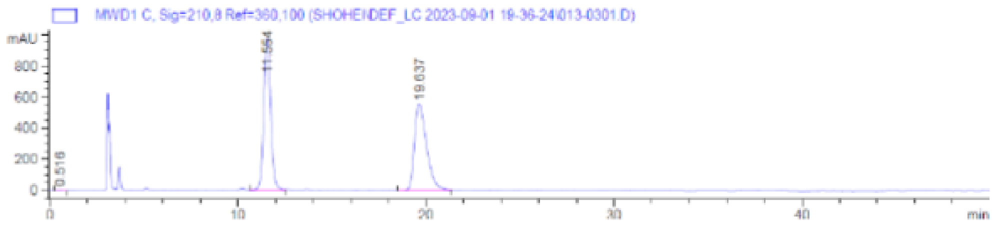

**Enantiomer 1**

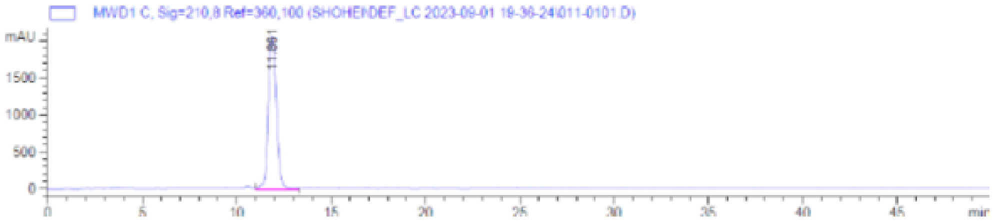

**Enantiomer 2**

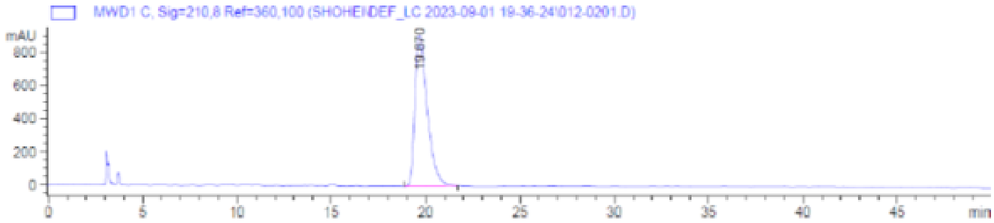

**Synthesis of #024**

・**Key Reaction**

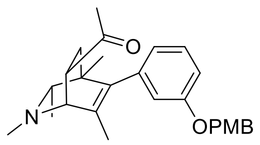

General procedure B was followed using above obtained imine (250 mg, 0.846 mmol, 1.0 equiv) and 2-butyne (332 µL, 4.23 mmol, 5 equiv) in THF (0.85 mL). Rh catalyst (10 mol %, 0.85 mL, 0.085 mmol, 100 mM in THF) was added, and the reaction was carried out at 65 °C for 12 h to produce the DHP intermediate. Following general procedure C, methyl vinyl ketone (106 µL, 1.27 mmol, 1.5 equiv) was added to the solution of DHP intermediate, and the reaction was carried out at room temperature for 12 h. The resulting crude material was purified by column chromatography (50% EtOAc/Hexane+1% $Et_3N$) to afford the desired product (101 mg, 28% yield from imine) as a pale-yellow oil. $^1H$ NMR (400 MHz, $CD_3OD$) δ 7.32 (d, $J$ = 7.9 Hz, 2H), 7.19 (t, $J$ = 8.1 Hz, 1H), 6.95 – 6.80 (m, 3H), 6.59 (d, $J$ = 7.4 Hz, 2H), 4.97 (s, 2H), 3.75 (s, 3H), 3.57 – 3.42 (m, 2H), 2.49 (s, 3H), 2.18 (s, 3H), 2.00 (q, $J$ = 6.2 Hz, 1H), 1.90 (dd, $J$ = 12.6, 5.1 Hz, 1H), 1.51 (s, 3H), 1.38 (dd, $J$ = 12.6,

9.1 Hz, 1H), 0.94 (d, $J$ = 6.3 Hz, 3H), 0.72 (s, 3H).

・**Deprotection**

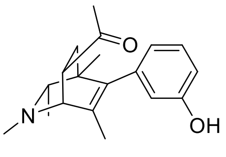

**(±)-#024**

**Final Compound #024** General Procedure D was followed using protected isoquinuclidine (101 mg, 0.241 mmol, 1.0 equiv) in $CH_2Cl_2$ (2.4 mL) and was reacted with trifluoroacetic acid (179 µL, 2.41 mmol, 10.0 equiv) for 12 h. The crude product was purified by preparative thin-layer chromatography (10% MeOH/$CH_2Cl_2$ + 1% $NH_4OH$) to yield the racemic isoquinuclidine (57 mg, 79% yield) as a white solid. $^1$H NMR (400 MHz, $CD_3OD$) δ 7.10 (t, $J$ = 7.9 Hz, 1H), 6.65 (dd, $J$ = 8.2, 1.2 Hz, 1H), 6.49 (m, 2H), 3.57 – 3.41 (m, 2H), 2.50 (s, 3H), 2.18 (s, 3H), 2.03 (q, $J$ = 6.3 Hz, 1H), 1.91 (dd, $J$ = 12.6, 5.1 Hz, 1H), 1.55 (s, 3H), 1.41 (dd, $J$ = 12.6, 8.9 Hz, 1H), 0.98 (d, $J$ = 6.3 Hz, 3H), 0.79 (s, 3H). $^{13}$C NMR (126 MHz, $CD_3OD$, 50 ℃) δ 209.2, 156.7, 139.7, 139.7, 134.4, 128.5, 120.1, 115.7, 113.2, 65.7, 61.8, 45.4, 40.8, 39.6, 35.9, 27.3, 20.4, 17.5, 16.7. HRMS (ESI+, m/z) calcd for $C_{19}H_{26}NO_2$ $(M+H)^+$ : 300.1958, found: 316.1968.

**Synthesis of #025**

・**Key Reaction**

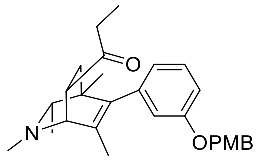

General procedure B was followed using above obtained imine (250 mg, 0.846 mmol, 1.0 equiv) and 2-butyne (332 µL, 4.23 mmol, 5 equiv) in THF (0.85 mL). Rh catalyst (10 mol %, 0.85 mL, 0.085 mmol, 100 mM in THF) was added, and the reaction was carried out at 65 °C for 12 h to produce the DHP intermediate. Following general procedure C, ethyl vinyl ketone (126 µL, 1.27 mmol, 1.5 equiv) was added to the solution of DHP intermediate, and the reaction was carried out at room temperature for 12 h. The resulting crude material was purified by column chromatography (50% EtOAc/Hexane+1% $Et_3N$) to afford the desired product (78 mg, 21% yield from imine) as a pale-yellow oil. $^1$H NMR (400 MHz, $CD_3OD$) δ 7.33 (d, $J$ = 6.8 Hz, 2H), 7.20 (t, $J$ = 7.9 Hz, 1H), 6.99 – 6.78 (m, 3H), 6.61 (d, $J$ = 7.5 Hz, 2H), 4.99 (s, 2H), 3.76 (s, 3H), 3.57 – 3.44 (m, 1H), 3.41 (d, $J$ = 2.6 Hz, 1H), 2.70 – 2.41 (m, 5H), 2.01 (q, $J$ = 6.3 Hz, 1H), 1.91 (dd, $J$ = 12.5, 5.2 Hz, 1H), 1.48 (s, 3H), 1.38 (dd, $J$ = 12.5, 9.1 Hz, 1H), 1.02 (t, $J$ = 7.3 Hz, 3H), 0.94 (d, $J$ = 6.3 Hz, 3H), 0.72 (s, 3H).

・**Deprotection**

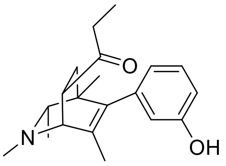

**(±)-#025**

**Final Compound #025** General Procedure D was followed using protected isoquinuclidine (78 mg, 0.180 mmol, 1.0 equiv) in $CH_2Cl_2$ (2.4 mL) and was reacted with trifluoroacetic acid (134 μL, 1.80 mmol, 10.0 equiv) for 12 h. The crude product was purified by preparative thin-layer chromatography (10% MeOH/$CH_2Cl_2$ + 1% $NH_4OH$) to yield the racemic isoquinuclidine (47 mg, 83% yield) as a white solid. $^1$H NMR (400 MHz, $CD_3OD$) δ 7.10 (t, $J$ = 8.0 Hz, 1H), 6.65 (ddd, $J$ = 8.1, 2.4, 1.1 Hz, 1H), 6.58 – 6.44 (m, 2H), 3.50 (ddd, $J$ = 8.9, 5.3, 2.7 Hz, 1H), 3.41 (d, $J$ = 2.7 Hz, 1H), 2.67 – 2.41 (m, 5H), 2.02 (q, $J$ = 6.3 Hz, 1H), 1.92 (dd, $J$ = 12.5, 5.3 Hz, 1H), 1.52 (s, 3H), 1.40 (dd, $J$ = 12.5, 8.9 Hz, 1H), 1.02 (t, $J$ = 7.3 Hz, 3H), 0.97 (d, $J$ = 6.3 Hz, 3H), 0.79 (s, 3H). $^{13}$C NMR (126 MHz, $CD_3OD$, 50 °C) δ 211.8, 156.7, 139.7, 139.5, 134.4, 128.5, 120.1, 115.7, 113.2, 65.6, 62.1, 44.3, 40.8, 39.5, 36.1, 33.9, 20.4, 17.5, 16.6, 6.8. HRMS (ESI+, m/z) calcd for $C_{20}H_{28}NO_2$ $(M+H)^+$ : 314.2115, found: 314.2124.

**Synthesis of #026**

・**Hydrogenation**

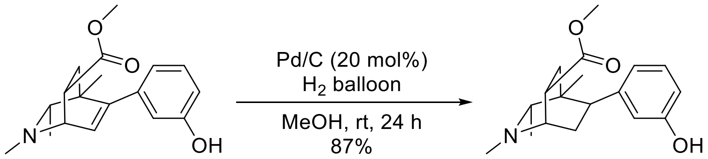

**(±)-#026**

**Final Compound #026** To a flame-dried round-bottom flask was added a magnetic stir bar, above obtained isoquinuclidine (35 mg, 0.083 mmol, 1 equiv) and 10 wt. % Pd/C (17.6 mg, 0.016 mmol, 20 mol%) dissolved in MeOH (5 mL). The headspace was purged with $H_2$ balloon and the reaction mixture was stirred at room temperature for 24 h. The reaction mixture was filtered through a pad of celite and washed with MeOH. The filtrate was concentrated and the crude residue was purified by preparative thin-layer chromatography (5% MeOH/$CH_2Cl_2$ + 1% $NH_4OH$) to yield the desired isoquinuclidine (22 mg, 87% yield) as a white solid. $^1$H NMR (500 MHz, $CDCl_3$) δ 7.15 (t, $J$ = 7.8 Hz, 1H), 6.94 (s, 1H), 6.88 (d, $J$ = 7.6 Hz, 1H), 6.73 (dd, $J$ = 8.2, 2.4 Hz, 1H), 3.78 (s, 3H), 3.15 – 3.05 (m, 1H), 3.02 – 2.89 (m, 2H), 2.52 – 2.38 (m, 4H), 2.21 – 2.08 (m, 2H), 1.78 (dd, $J$ = 13.9, 8.7 Hz, 1H),

1.20 – 1.09 (m, 4H), 0.39 (s, 3H). $^{13}$C NMR (126 MHz, $CDCl_3$) δ 175.9, 155.9, 143.9, 129.1, 121.8, 116.5, 113.6, 66.1, 56.7, 52.1, 40.7, 39.4, 36.3, 35.2, 31.4, 30.8, 22.3, 13.5. HRMS (ESI+, m/z) calcd for $C_{18}H_{26}NO_3$ $(M+H)^+$ : 304.1907, found: 304.1917.

**Synthesis of #027**

・**Synthesis of Aldehyde**

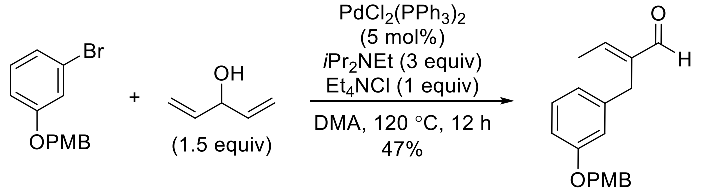

To a round-bottom flask was added a magnetic stir bar, 1-bromo-3-(4-methoxyphenyl)methoxy benzene (1.0 g, 3.41 mmol, 1 equiv) and 1,4-pentadien-3-ol (430 mg, 5.12 mmol, 1.5 equiv) dissolved in DMA (17 mL). To this solution was then added *N*,*N*-diisopropylethylamine (1.78 mL, 10.2 mmol, 3 equiv), dichlorobis(triphenylphosphine)palladium (120 mg, 0.171 mmol, 5 mol%) and tetraethylammonium chloride (565 mg, 3.41 mmol, 1 equiv). The reaction mixture was stirred at 120 °C for 12 h. After the reaction was complete, the mixture was then diluted with water, extracted with $CH_2Cl_2$, washed with brine, dried with $Na_2SO_4$ and filtered. The organic layer was concentrated and the crude residue was purified via silica gel chromatography (gradient of 0-100% EtOAc in hexane) to give 0.48 g of desired aldehyde in 47% yield. $^1$H NMR (400 MHz, $CDCl_3$) δ 9.43 (s, 1H), 7.35 – 7.30 (m, 2H), 7.17 – 7.12 (m, 1H), 6.91 – 6.87 (m, 2H), 6.79 – 6.74 (m, 3H), 6.70 (q, $J$ = 7.2 Hz, 1H), 4.92 (s, 2H), 3.80 (s, 3H), 3.59 (s, 2H), 2.00 (d, $J$ = 7.2 Hz, 3H).

・**Synthesis of Imine**

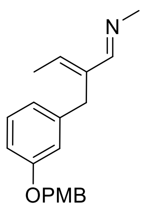

General procedure A was followed using 2.0 M methylamine in THF (1.27 mL, 2.53 mmol, 5.0 equiv), above obtained aldehyde (150 mg, 0.506 mmol, 1.0 equiv), molecular sieves 4 Å, and $CH_2Cl_2$ (1.0 mL) at 50 °C. Imine (150 mg, 96% yield) was obtained as a yellow solid. $^1$H NMR (400 MHz, $CDCl_3$) δ 7.82 (s, 1H), 7.35 – 7.30 (m, 2H), 7.13 (t, $J$ = 7.9 Hz, 1H), 6.91 – 6.86 (m, 2H), 6.83 – 6.71 (m, 3H), 6.06 (q, $J$ = 7.0 Hz, 1H), 4.92 (s, 2H), 3.80 (s, 3H), 3.72 (s, 2H), 3.33 (s, 2H), 1.82 (d, $J$ = 7.0 Hz, 3H).

・**Key Reaction**

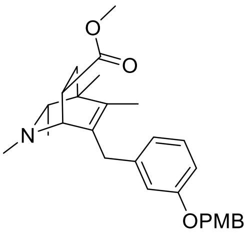

General procedure B was followed using above obtained imine (150 mg, 0.485 mmol, 1.0 equiv) and 2-butyne (190 µL, 2.42 mmol, 5 equiv) in THF (0.49 mL). Rh catalyst (10 mol %, 0.49 mL, 0.049 mmol, 100 mM in THF) was added, and the reaction was carried out at 65 °C for 12 h to produce the DHP intermediate. Following general procedure C, methyl acrylate (0.22 mL, 2.42 mmol, 5 equiv) was added to the solution of DHP intermediate, and the reaction was carried out at 50 °C for 16 h. The resulting crude material was purified by column chromatography (50% EtOAc/Hexane+1% $Et_3N$) to afford the desired product (61 mg, 28% yield from imine) as a pale-yellow oil. $^1$H NMR (400 MHz, $CDCl_3$) δ 7.35 (d, $J$ = 8.5 Hz, 2H), 7.15 (t, $J$ = 7.9 Hz, 1H), 6.99 – 6.83 (m, 3H), 6.83 – 6.70 (m, 2H), 4.97 (d, $J$ = 11.1 Hz, 1H), 4.93 (d, $J$ = 11.1 Hz, 1H), 3.80 (s, 3H), 3.56 (d, $J$ = 15.9 Hz, 1H), 3.40 (d, $J$ = 15.9 Hz, 1H), 3.31 (s, 3H), 3.07 (ddd, $J$ = 9.4, 5.5, 2.8 Hz, 1H), 2.36 (s, 3H), 1.88 (q, $J$ = 6.2 Hz, 1H), 1.80 (dd, $J$ = 12.8, 5.5 Hz, 1H), 1.39 (dd, $J$ = 12.8, 9.4 Hz, 1H), 1.09 (s, 3H), 0.84 (d, $J$ = 6.2 Hz, 3H).

・**Deprotection**

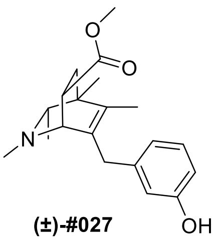

**(±)-#027**

**Final Compound #027** General Procedure D was followed using protected isoquinuclidine (45 mg, 0.100 mmol, 1.0 equiv) in $CH_2Cl_2$ (1.0 mL) and was reacted with trifluoroacetic acid (74 µL, 1.00 mmol, 10.0 equiv) for 12 h. The crude product was purified by preparative thin-layer chromatography (10% MeOH/$CH_2Cl_2$ + 1% $NH_4OH$) to yield the racemic isoquinuclidine (14 mg, 40% yield) as a pale-yellow oil. $^1$H NMR (400 MHz, $CD_3OD$) δ 7.01 (t, $J$ = 7.8 Hz, 1H), 6.66 – 6.42 (m, 3H), 3.54 (d, $J$ = 15.3 Hz, 1H), 3.36 – 3.32 (m, 1H), 3.26 (d, $J$ = 15.3 Hz, 1H), 3.13 (s, 3H), 3.06 (ddd, $J$ = 9.3, 5.8, 2.7 Hz, 1H), 2.35 (s, 3H), 1.92 (q, $J$ = 6.3 Hz, 1H), 1.80 (s, 3H), 1.77 (dd, $J$ = 12.9, 5.8 Hz, 1H), 1.38 (dd, $J$ = 12.8, 9.3 Hz, 1H), 1.11 (s, 3H), 0.83 (d, $J$ = 6.3 Hz, 3H). $^{13}$C NMR (101 MHz, $CD_3OD$) δ 174.3, 157.3, 140.8, 135.4, 135.0, 128.8, 119.7, 115.7, 112.6, 65.0, 61.1, 50.6, 40.1, 39.2, 36.3, 35.9, 35.9, 19.3, 16.8, 12.9. HRMS (ESI+, m/z) calcd for $C_{20}H_{27}NO_3$ $(M+H)^+$ : 330.2064, found: 330.2057.

**Synthesis of #028**

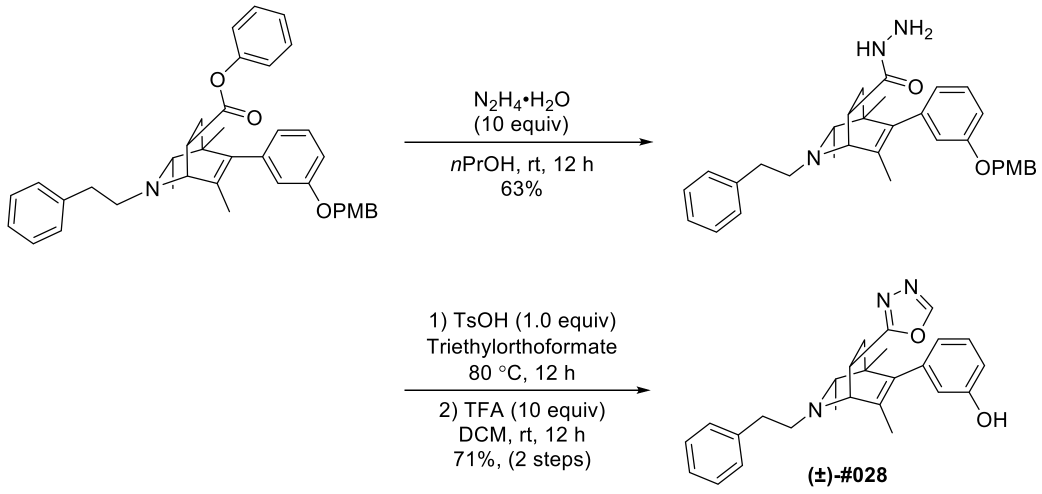

To a round-bottom flask was added a magnetic stir bar, above obtained isoquinuclidine (210 mg, 0.357 mmol, 1 equiv) dissolved in *n*PrOH (1.8 mL). To this solution was then added 65% hydrazine monohydrate (275 mg, 3.57 mmol, 10 equiv) at 0 °C. The reaction mixture was stirred at room temperature for 12 h. After the reaction was complete, the mixture was then diluted with water, extracted with $CH_2Cl_2$, dried with $Na_2SO_4$ and filtered. The organic layer was concentrated and the crude residue was purified by preparative thin-layer chromatography (10% MeOH/$CH_2Cl_2$ + 1% $NH_4OH$) to give 118 mg of amide compound in 63% yield. $^1$H NMR (400 MHz, $CD_3OD$) δ 7.34 (d, *J* = 8.2 Hz, 2H), 7.30 – 7.24 (m, 4H), 7.24 – 7.14 (m, 2H), 6.89 (d, *J* = 8.5 Hz, 2H), 6.85 (dd, *J* = 8.3, 2.4 Hz, 1H), 6.80 – 6.57 (m, 2H), 4.99 (s, 2H), 3.76 (s, 3H), 3.45 (d, *J* = 2.9 Hz, 1H), 3.16 – 3.12 (m, 1H), 2.98 – 2.72 (m, 4H), 2.20 (q, *J* = 6.3 Hz, 1H), 1.93 (dd, *J* = 12.7, 4.3 Hz, 1H), 1.57 (s, 3H), 1.47 (dd, *J* = 12.7, 9.3 Hz, 1H), 1.01 (d, *J* = 6.3 Hz, 3H), 0.77 (s, 3H).

**Final Compound #028** To a stirred solution of the above obtained compound (60 mg, 0.114 mmol, 1 equiv) in triethylorthoformate (4.6 mL) was added *p*-toluenesulfonic acid monohydrate (21.7 mg, 0.114 mmol, 1 equiv). The reaction mixture was heated at 80 °C for 12 h. After the reaction was complete, the mixture was concentrated and dissolved in $CH_2Cl_2$ (1.2 mL). This solution was added trifluoroacetic acid (85 µL, 1.14 mmol, 10.0 equiv) and stirred at room temperature for 12 h. The crude product was purified by preparative thin-layer chromatography (10% MeOH/$CH_2Cl_2$ + 1% $NH_4OH$) to yield the racemic isoquinuclidine (34 mg, 71% yield) as a pale yellow oil. $^1$H NMR (400 MHz, $CD_3OD$) δ 8.83 (s, 1H), 7.27 (d, *J* = 4.4 Hz, 4H), 7.22 – 7.06 (m, 2H), 6.66 (dd, *J* = 8.0, 2.5 Hz, 1H), 6.53 (d, *J* = 5.3 Hz, 2H), 4.03 – 3.90 (m, 1H), 3.68 (d, *J* = 3.0 Hz, 1H), 3.04 – 2.76 (m, 4H), 2.34 (q, *J* = 6.3 Hz, 1H), 2.12 (dd, *J* = 13.1, 4.5 Hz, 1H), 1.93 (dd, *J* = 13.1, 9.5 Hz, 1H), 1.37 (s, 3H), 1.04 (d, *J* = 6.2 Hz, 3H), 0.88 (s, 3H). $^{13}$C NMR (126 MHz, $CD_3OD$, 50 °C) δ 168.9, 156.8, 153.9, 140.6, 139.9, 139.6, 134.0, 128.6, 128.4, 128.0, 125.8, 115.7, 113.3, 65.2, 60.4, 56.2, 40.8, 37.5, 35.6, 29.7, 20.5,

19.3, 16.3. HRMS (ESI+, m/z) calcd for $C_{26}H_{29}N_3O_2$ $(M+H)^+$ : 416.2333, found: 416.2335.

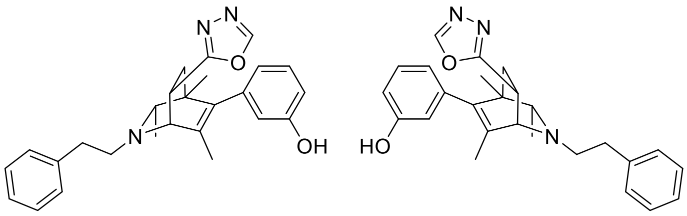

**#028**
enantiomers separated by chiral HPLC

#### HPLC Traces of #028

A portion of this material was separated using semi-preparative chiral HPLC (Chiralpak IC column, 250 x 10 mm, 10% *i*PrOH/Hexanes+1% DEA, 2.5 mL/min) to provide the two enantiomers with tr = 26 min and 39 min.

Chiralpak IC, analytical column (250 x 4.6 mm), 10% *i*PrOH/Hexanes+0.1% DEA, 1 mL/min, 210 nm

**Racemic**

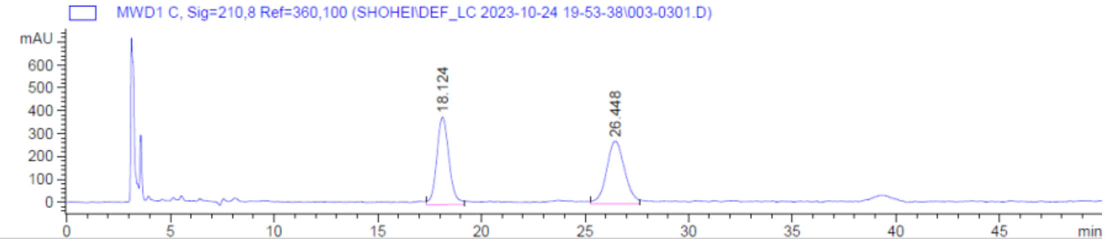

**Enantiomer 1**

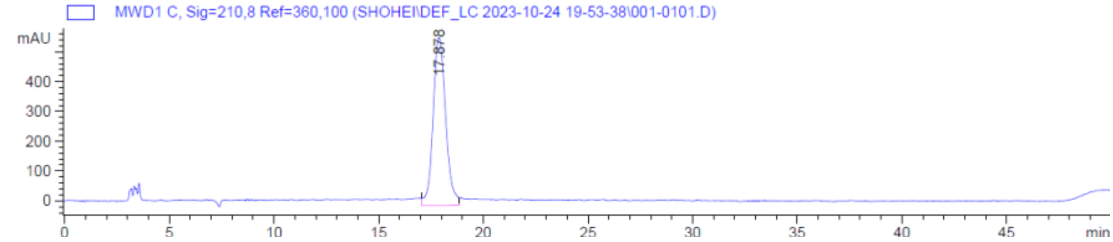

**Enantiomer 2**

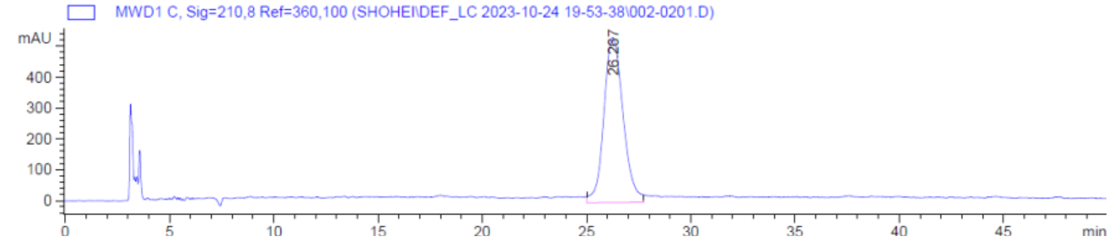

#### Synthesis of #029

・**Key Reaction**

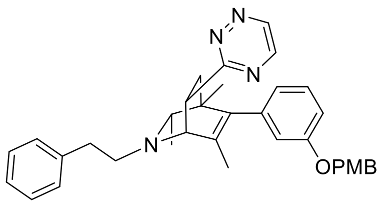

General procedure B was followed using above obtained imine (200 mg, 0.519 mmol, 1.0 equiv) and 2-butyne (203 µL, 2.59 mmol, 5 equiv) in THF (0.52 mL). Rh catalyst (10 mol %, 0.52 mL, 0.052 mmol, 100 mM in THF) was added, and the reaction was carried out at 65 °C for 12 h to produce the DHP intermediate. Following general procedure C, 3-ethenyl-1,2,4-triazine (278 mg, 2.59 mmol, 5 equiv) was added to the solution of DHP intermediate, and the reaction was carried out at 50 °C for 16 h. The resulting crude material was purified by column chromatography (50% EtOAc/Hexane+1% $Et_3N$) to afford the desired product (95 mg, 34% yield from imine) as a pale-yellow oil. $^1$H NMR (400 MHz, $CD_3OD$) δ 9.11 (d, *J* = 2.4 Hz, 1H), 8.69 (d, *J* = 2.4 Hz, 1H), 7.36 – 7.15 (m, 8H), 6.91 – 6.82 (m, 3H), 6.81 – 6.58 (m, 2H), 5.00 (s, 2H), 4.15 (dt, *J* = 9.3, 3.8 Hz, 1H), 3.81 – 3.71 (m, 4H), 3.07 – 2.79 (m, 4H), 2.57 (dd, *J* = 12.9, 4.4 Hz, 1H), 2.35 (q, *J* = 6.3 Hz, 1H), 1.76 (dd, *J* = 12.9, 9.3 Hz, 1H), 1.09 (s, 3H), 1.04 (d, *J* = 6.3 Hz, 3H), 0.84 (s, 3H).

・**Deprotection**

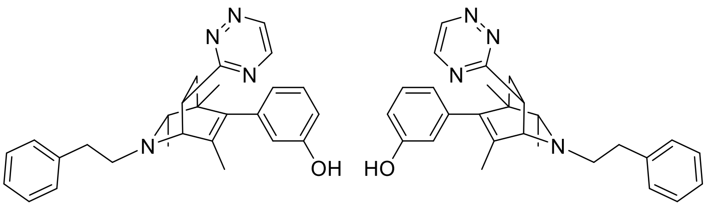

**#029**
enantiomers separated by chiral HPLC

**Final Compound #029** General Procedure D was followed using protected isoquinuclidine (95 mg, 0.174 mmol, 1.0 equiv) in $CH_2Cl_2$ (1.7 mL) and was reacted with trifluoroacetic acid (129 µL, 1.74 mmol, 10.0 equiv) for 12 h. The crude product was purified by preparative thin-layer chromatography (10% MeOH/$CH_2Cl_2$ + 1% $NH_4OH$) to yield the racemic isoquinuclidine (35 mg, 47% yield) as a pale-yellow oil. $^1$H NMR (400 MHz, $CD_3OD$) δ 9.11 (d, *J* = 2.5 Hz, 1H), 8.72 (d, *J* = 2.5 Hz, 1H), 7.34 – 7.23 (m, 4H), 7.21 – 7.14 (m, 1H), 7.11 (t, *J* = 7.7 Hz, 1H), 6.65 (dd, *J* = 8.1, 2.4 Hz, 1H), 6.62 – 6.51 (m, 2H), 4.16 (dt, *J* = 8.3, 3.8 Hz, 1H), 3.80 (d, *J* = 3.1 Hz, 1H), 3.08 – 2.80 (m, 4H), 2.57 (dd, *J* = 13.0, 4.4 Hz, 1H), 2.39 (q, *J* = 6.2 Hz, 1H), 1.78 (dd, *J* = 13.0, 9.5 Hz, 1H), 1.13 (s, 3H), 1.08 (d, *J* = 6.2 Hz, 3H), 0.91 (s, 3H). $^{13}$C NMR (126 MHz, $CD_3OD$, 50 °C) δ 170.8, 156.6, 149.3, 147.2, 140.8, 140.0, 139.9, 133.2, 128.5, 128.4, 128.0, 125.8, 120.1, 115.8, 113.1, 65.8, 62.6, 56.6, 41.2, 39.4, 36.3, 35.4, 20.7, 19.1, 16.6. HRMS (ESI+, m/z) calcd for $C_{27}H_{31}N_4O$ $(M+H)^+$ : 427.2492, found:

427.2522.

**HPLC Traces of #029**

A portion of this material was separated using semi-preparative chiral HPLC (Chiralpak AD-H column, 250 x 10 mm, 7% *i*PrOH/Hexanes+1% DEA, 2.5 mL/min) to provide the two enantiomers with tr = 44 min and 58 min.

Chiralpak AD-H, analytical column (250 x 4.6 mm), 7% *i*PrOH/Hexanes+0.1% DEA, 1 mL/min, 210 nm

**Racemic**

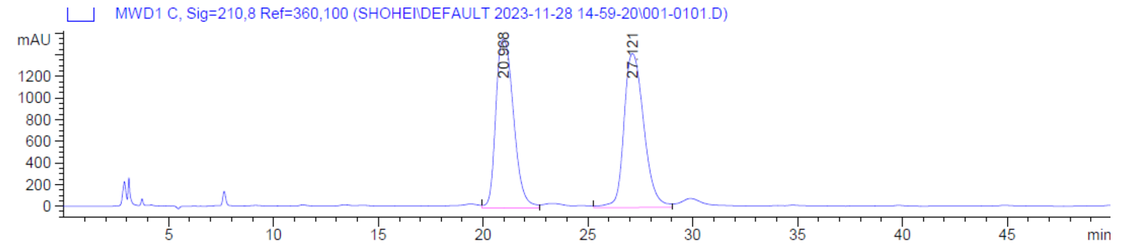

**Enantiomer 1**

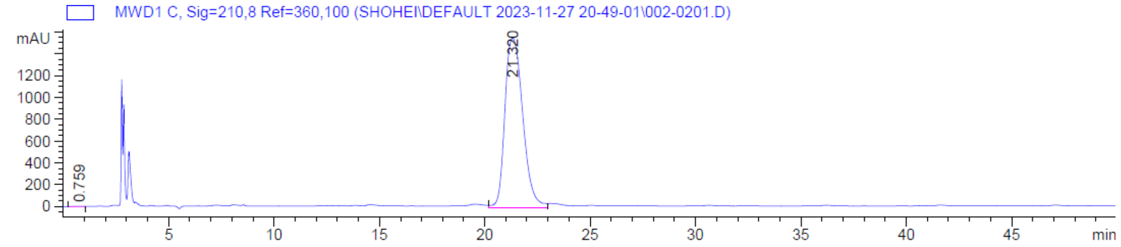

**Enantiomer 2**

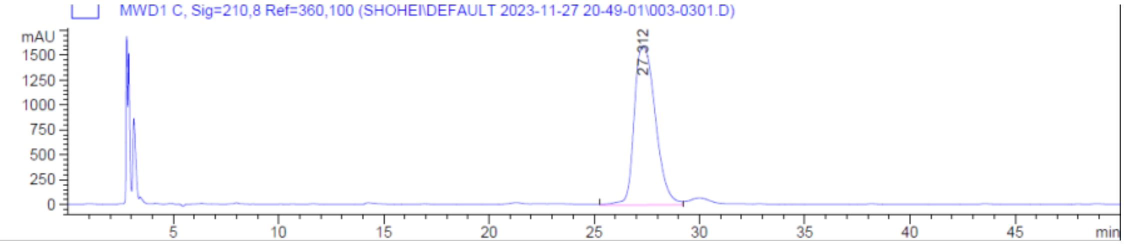

**Synthesis of #030**

・**Key Reaction**

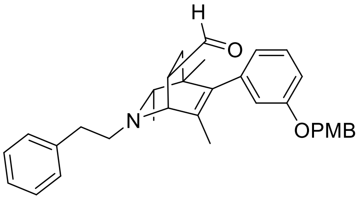

General procedure B was followed using above obtained imine (250 mg, 0.649 mmol, 1.0 equiv) and 2-butyne (254 µL, 3.24 mmol, 5 equiv) in THF (0.65 mL). Rh catalyst (10 mol %, 0.65 mL, 0.065 mmol, 100 mM in THF) was added, and the reaction was carried out at 65 °C for 12 h to produce the DHP intermediate. Following general procedure C, acrolein (217 µL, 3.24 mmol, 5 equiv) was added to the solution of DHP intermediate, and the reaction was carried out at room temperature for 12 h. The resulting crude material was purified by column chromatography (40% EtOAc/Hexane+1% $Et_3N$) to afford the desired product (90 mg, 28% yield from imine) as a pale-yellow oil. $^1$H NMR (400 MHz, $CDCl_3$) δ 9.53 (s, 1H), 7.43 – 7.17 (m, 9H), 6.90 (d, $J$ = 8.6 Hz, 2H), 6.86 (dd, $J$ = 8.2, 2.6 Hz, 1H), 6.75 – 6.51 (m, 2H), 4.97 (s, 2H), 3.80 (s, 3H), 3.69 (d, $J$ = 3.1 Hz, 1H), 3.22 (dt, $J$ = 8.6, 3.8 Hz, 1H), 2.98 – 2.77 (m, 4H), 2.15 (q, $J$ = 6.3 Hz, 1H), 1.97 (dd, $J$ = 13.0, 4.3 Hz, 1H), 1.61 (s, 3H), 1.49 (dd, $J$ = 13.0, 9.2 Hz, 1H), 0.99 (d, $J$ = 6.3 Hz, 3H), 0.84 (s, 3H).

・**Decarbonylation and deprotection**

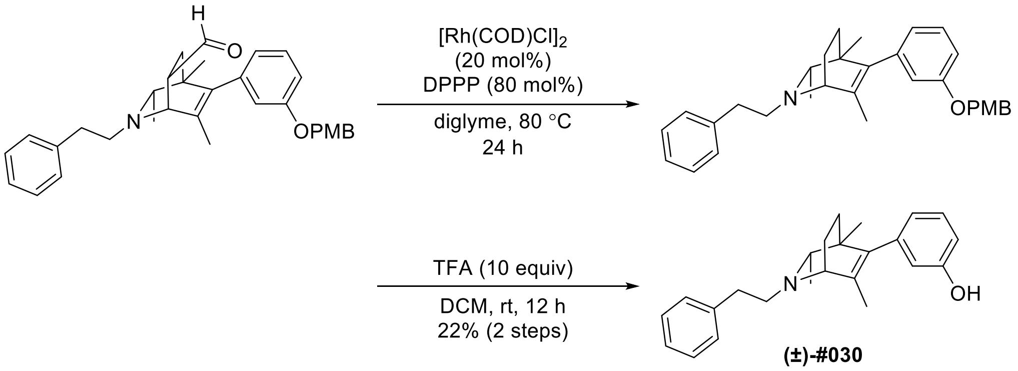

**(±)-#030**

**Final Compound #030** To a sealable glass vessel was added a magnetic stir bar, above obtained isoquinuclidine (150 mg, 0.303 mmol, 1 equiv) dissolved in diglyme (10 mL). To this solution was then added chloro(1,5-cyclooctadiene)rhodium(I) dimer (20 mol%, 30 mg, 0.061 mmol) and 1,3-bis(diphenylphosphino) propane (80 mol%, 100 mg, 0.244 mmol) at room temperature. The reaction mixture was stirred at 80 °C for 24 h. After the reaction was complete, the mixture was then diluted with water, extracted with $CH_2Cl_2$, dried with $Na_2SO_4$ and filtered. The organic layer was concentrated and the crude residue was purified by preparative thin-layer chromatography (35% EtOAc/Hexane+1% $Et_3N$) to give 45 mg of compound in 32% yield. The above obtained compound was dissolved in $CH_2Cl_2$ (1.0 mL). To this solution was added trifluoroacetic acid (65 µL, 0.877 mmol, 10.0 equiv) and the resulting solution was stirred at room temperature for 12 h. The crude product was purified by preparative thin-layer chromatography (10% MeOH/$CH_2Cl_2$ + 1% $NH_4OH$) to yield the racemic isoquinuclidine (21 mg, 69% yield) as a pale-yellow oil. $^1$H NMR (500 MHz, $CD_3OD$) δ 7.31 – 7.22 (m, 4H), 7.19 (t, *J* = 7.1 Hz, 1H), 7.14 (t, *J* = 7.9 Hz, 1H), 6.68 (dd, *J* = 8.1, 2.4 Hz, 1H), 6.63 – 6.31 (broad s, 2H), 3.42 – 3.36 (m, 1H), 2.99 – 2.70 (m, 4H), 2.36 (q, *J* = 6.3 Hz, 1H), 2.29 – 2.18 (m, 1H), 1.73 (s, 3H), 1.54 – 1.29 (m, 3H), 1.08 (d, *J* = 6.3 Hz, 3H), 0.83 (s, 3H). $^{13}$C NMR (126 MHz, $CD_3OD$) δ 156.8, 139.9, 139.8, 139.6, 136.2, 128.6, 128.3, 128.1, 125.9, 120.0, 115.6, 113.1, 66.8, 57.8, 57.2, 40.5, 34.8, 33.7, 20.9, 19.21, 19.17, 15.6. HRMS (ESI+, m/z) calcd for $C_{24}H_{30}NO$ $(M+H)^+$ : 348.2322, found: 348.2313.

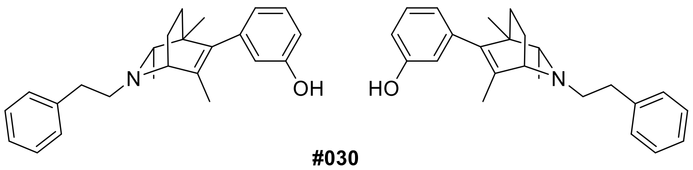

**#030**
enantiomers separated by chiral HPLC

**HPLC Traces of #030**

A portion of this material was separated using semi-preparative chiral HPLC (Chiralpak AD-H column, 250 x 10 mm, 4% *i*PrOH/Hexanes+1% DEA, 2.5 mL/min) to provide the two enantiomers with tr = 26 min and 32 min.

Chiralpak AD-H, analytical column (250 x 4.6 mm), 4% *i*PrOH/Hexanes+0.1% DEA, 1 mL/min, 280 nm

**Racemic**

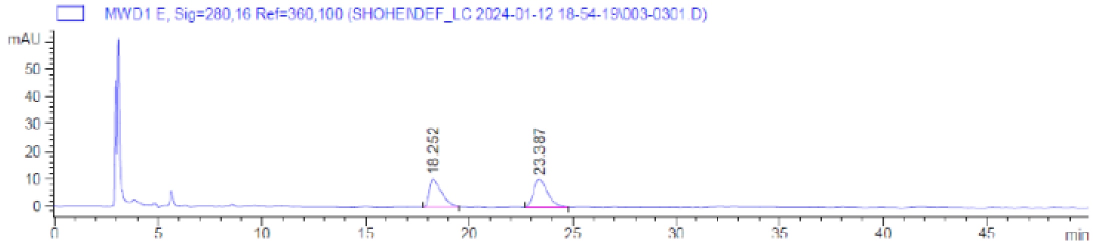

**Enantiomer 1**

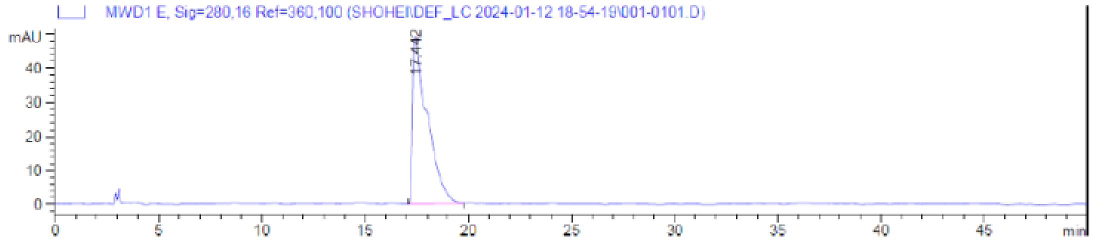

**Enantiomer 2**

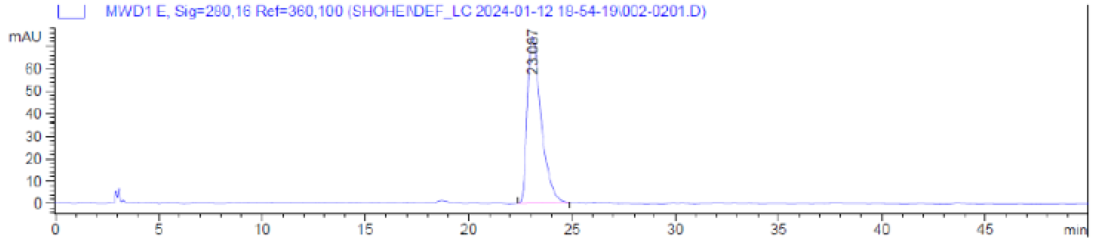

#### Synthesis of #031

##### ・Synthesis of Imine

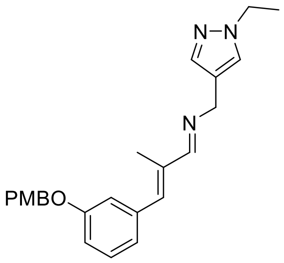

General procedure A was followed using (1-ethyl-1H-pyrazol-4-yl)methanamine (122 mg, 0.974 mmol, 1.1 equiv), above obtained aldehyde (250 mg, 0.885 mmol, 1.0 equiv), molecular sieves 4 Å, and $CH_2Cl_2$ (1.9 mL) at 50 °C. Imine (340 mg, 99% yield) was obtained as a pale-yellow oil. $^1$H NMR (400 MHz, $CDCl_3$) δ 8.01 (s, 1H), 7.46 – 7.22 (m, 6H), 7.01 – 6.75 (m, 5H), 4.99 (d, $J$ = 2.5 Hz, 2H), 4.62 (s, 2H), 4.13 (q, $J$ = 7.5 Hz, 2H), 3.80 (s, 3H), 2.13 (s, 3H), 1.46 (t, $J$ = 7.5 Hz, 3H).

##### ・Key Reaction

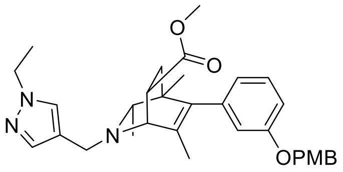

General procedure B was followed using above obtained imine (340 mg, 0.873 mmol, 1.0 equiv) and 2-butyne (342 µL, 4.36 mmol, 5 equiv) in THF (0.87 mL). Rh catalyst (10 mol %, 0.87 mL, 0.087 mmol, 100 mM in THF) was added, and the reaction was carried out at 65 °C for 12 h to produce the DHP intermediate. Following general procedure C, methyl acrylate (0.39 mL, 4.36 mmol, 5 equiv) was added to the solution of DHP intermediate, and the reaction was carried out at 50 °C for 16 h. The resulting crude material was purified by column chromatography (50% EtOAc/Hexane+1% $Et_3N$) to afford the desired product (280 mg, 61% yield from imine) as a pale-yellow oil. $^1$H NMR (400 MHz, $CD_3OD$) δ 7.63 (d, $J$ = 2.5 Hz, 1H), 7.51 (d, $J$ = 2.5 Hz, 1H), 7.31 (d, $J$ = 8.1 Hz, 2H), 7.19 (td, $J$ = 8.1, 2.4 Hz, 1H), 6.92 – 6.74 (m, 3H), 6.69 – 6.50 (m, 2H), 4.97 (s, 2H), 4.15 (qd, $J$ = 7.3, 2.4 Hz, 2H), 3.80 – 3.66 (m, 4H), 3.64 (s, 3H), 3.60 – 3.49 (m, 2H), 3.42 – 3.35 (m, 1H), 2.18 (q, $J$ = 3.9 Hz, 1H), 1.95 (dd, $J$ = 12.8, 4.4 Hz, 1H), 1.55 (dd, $J$ = 12.8, 9.2 Hz, 1H), 1.49 – 1.34 (m, 6H), 0.84 (d, $J$ = 3.9 Hz, 3H), 0.74 (s, 3H).

・**Deprotection**

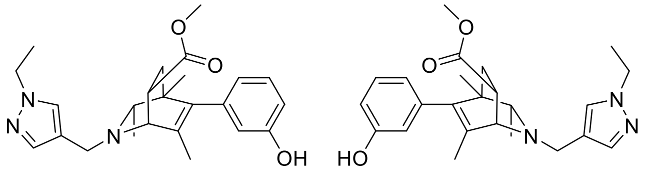

**#031**
enantiomers separated by chiral HPLC

**Final Compound #031** General Procedure D was followed using protected isoquinuclidine (150 mg, 0.283 mmol, 1.0 equiv) in $CH_2Cl_2$ (2.8 mL) and was reacted with trifluoroacetic acid (210 µL, 2.83 mmol, 10.0 equiv) for 12 h. The crude product was purified by preparative thin-layer chromatography (10% MeOH/$CH_2Cl_2$ + 1% $NH_4OH$) to yield the racemic isoquinuclidine (70 mg, 60% yield) as a white solid. $^1$H NMR (500 MHz, $CD_3OD$) δ 7.62 (s, 1H), 7.51 (s, 1H), 7.11 (t, $J$ = 7.9 Hz, 1H), 6.67 (dd, $J$ = 8.2, 3.1 Hz, 1H), 6.58 – 6.46 (m, 2H), 4.16 (q, $J$ = 7.4 Hz, 2H), 3.76 (d, $J$ = 13.4 Hz, 1H), 3.65 (s, 3H), 3.60 (d, $J$ = 13.4 Hz, 1H), 3.58 (d, $J$ = 3.0 Hz, 1H), 3.42 – 3.36 (m, 1H), 2.22 (q, $J$ = 6.2 Hz, 1H), 1.97 (dd, $J$ = 12.9, 4.6 Hz, 1H), 1.58 (dd, $J$ = 12.9, 9.4 Hz, 1H), 1.50 (s, 3H), 1.44 (t, $J$ = 7.4 Hz, 3H), 0.90 (d, $J$ = 6.2 Hz, 3H), 0.83 (s, 3H). $^{13}$C NMR (126 MHz, $CD_3OD$) δ 174.8, 156.7, 140.5, 139.8, 139.4, 134.2, 129.3, 128.5, 120.0, 118.2, 115.6, 113.2, 64.8, 58.6, 50.8, 46.4, 40.8, 36.9, 36.4, 20.5, 18.7, 16.4, 14.5. HRMS (ESI+, m/z) calcd for $C_{24}H_{32}N_3O_3$ $(M+H)^+$ : 410.2438, found: 410.2421.

**HPLC Traces of #031**

A portion of this material was separated using semi-preparative chiral HPLC (Chiralpak IC column, 250 x 10 mm, 12% *i*PrOH/Hexanes+1% DEA, 2.5 mL/min) to provide the two enantiomers with tr = 36 min and 46 min.

Chiralpak IC, analytical column (250 x 4.6 mm), 12% *i*PrOH/Hexanes+0.1% DEA, 1 mL/min, 280 nm

**Racemic**

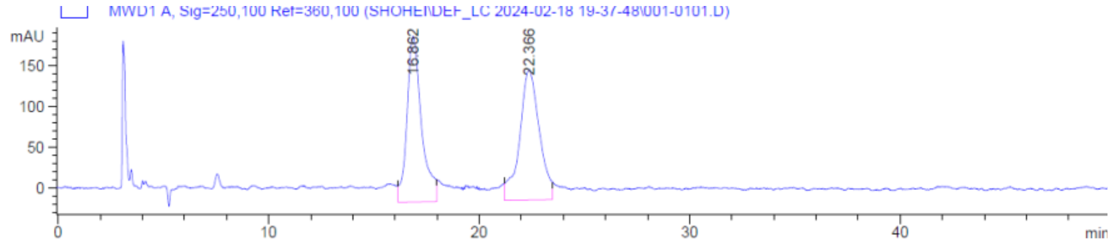

**Enantiomer 1**

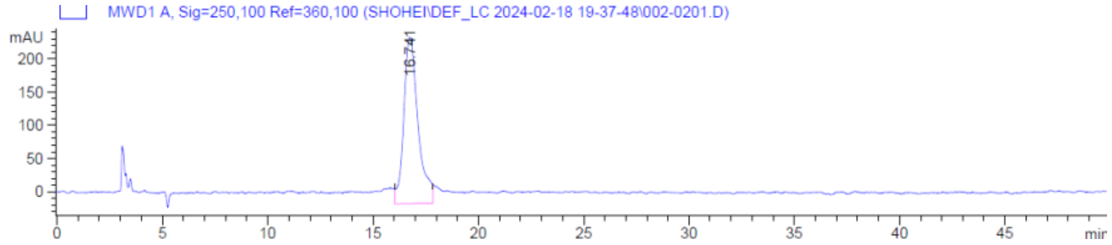

**Enantiomer 2**

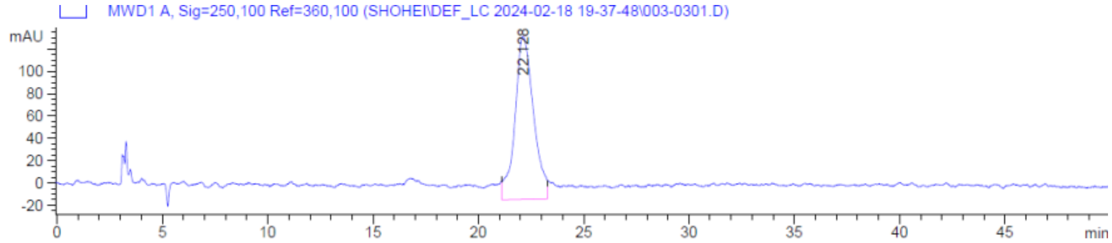

**Synthesis of #036**

・**Key Reaction**

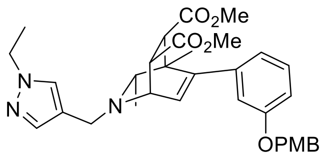

General procedure B was followed using above obtained imine (290 mg, 0.772 mmol, 1.0 equiv) and 2-butyne (303 µL, 3.86 mmol, 5 equiv) in THF (0.77 mL). Rh catalyst (10 mol %, 0.77 mL, 0.077 mmol, 100 mM in THF) was added, and the reaction was carried out at 65 °C for 12 h to produce the DHP intermediate. Following general procedure C, 1,4-dimethyl (2Z)-but-2-enedioate (167 mg, 1.16 mmol, 1.5 equiv) was added to the solution of DHP intermediate, and the reaction was carried out at 50 °C for 16 h. The resulting crude material was purified by column chromatography (70% EtOAc/Hexane+1% $Et_3N$) to afford the desired product (187 mg, 42% yield from imine) as a pale-yellow oil. $^1$H NMR (400 MHz, $CDCl_3$) δ 7.52 (s, 1H), 7.46 – 7.37 (m, 1H), 7.33 (d, *J* = 8.6 Hz, 2H), 7.19 (t, *J* = 8.0 Hz, 1H), 6.91 – 6.84 (m, 3H), 6.76 (d, *J* = 8.1 Hz, 2H), 6.15 (d, *J* = 6.5 Hz, 1H), 4.95 (s, 2H), 4.13 (q, *J* = 7.4 Hz, 2H), 3.96 – 3.83 (m, 3H), 3.79 (s, 3H), 3.74 – 3.69 (m, 4H), 3.64 (s, 3H), 3.08 (d, *J* = 5.9 Hz, 1H), 2.82 (s, 1H), 1.46 (t, *J* = 7.3 Hz, 3H), 0.97 (s, 3H), 0.83 (d, *J* = 6.2 Hz, 3H).

・**Deprotection**

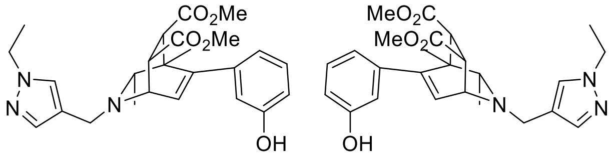

**#036**
enantiomers separated by chiral HPLC

**Final Compound #036** General Procedure D was followed using protected isoquinuclidine (187 mg, 0.326 mmol, 1.0 equiv) in $CH_2Cl_2$ (3.3 mL) and was reacted with trifluoroacetic acid (242 µL, 3.26 mmol, 10.0 equiv) for 12 h. The crude product was purified by preparative thin-layer chromatography (10% MeOH/$CH_2Cl_2$ + 1% $NH_4OH$) to yield the racemic isoquinuclidine (73 mg, 50% yield) as a white solid. $^1$H NMR (400 MHz, $CD_3OD$) δ 7.65 (s, 1H), 7.52 (s, 1H), 7.12 (t, *J* = 8.0 Hz, 1H), 6.75 – 6.64 (m, 1H), 6.64 – 6.53 (m, 2H), 6.11 (d, *J* = 6.4 Hz, 1H), 4.16 (q, *J* = 7.3 Hz, 2H), 3.98 (d, *J* = 13.3 Hz, 1H), 3.91 – 3.80 (m, 2H), 3.77 (dd, *J* = 5.9, 2.6 Hz, 1H), 3.73 (s, 3H), 3.66 (s, 3H), 3.05 (d, *J* = 5.9 Hz, 1H), 2.89 (q, *J* = 6.3 Hz, 1H), 1.43 (t, *J* = 7.3 Hz, 3H), 0.99 (s, 3H), 0.82 (d, *J* = 6.3 Hz, 3H). $^{13}$C NMR (101 MHz, $CD_3OD$) δ 173.7, 173.4, 156.8, 148.9, 140.3, 139.6, 129.7, 128.7, 119.2, 117.6, 114.9, 113.9, 56.6, 52.5, 51.4, 50.9, 50.7, 46.4, 44.4, 43.2, 41.1, 18.8, 18.4, 14.7.
HRMS (ESI+, m/z) calcd for $C_{25}H_{31}N_3O_5$ $(M+H)^+$ : 454.2336, found: 454.2315.

**HPLC Traces of #036**

A portion of this material was separated using semi-preparative chiral HPLC (Chiralpak IC column, 250 x 10 mm, 20% *i*PrOH/Hexanes+1% DEA, 2.5 mL/min) to provide the two enantiomers with tr = 16 min and 32 min.

Chiralpak IC, analytical column (250 x 4.6 mm), 20% *i*PrOH/Hexanes+0.1% DEA, 1 mL/min, 254 nm

**Racemic**

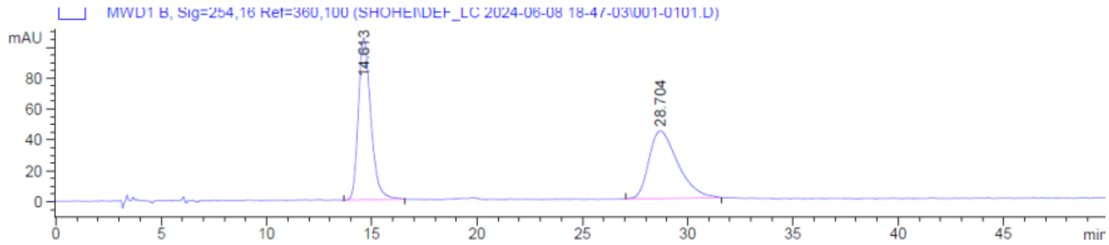

**Enantiomer 1**

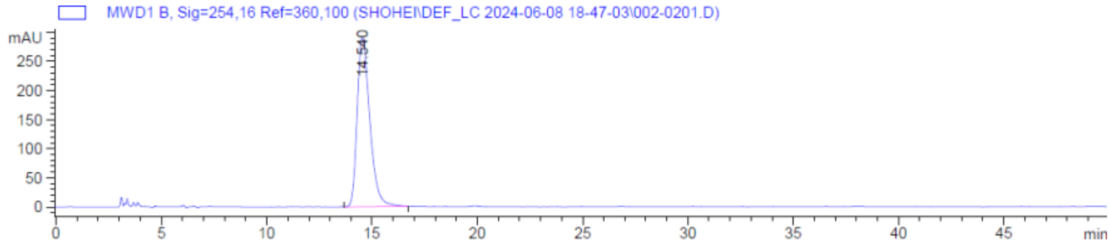

**Enantiomer 2**

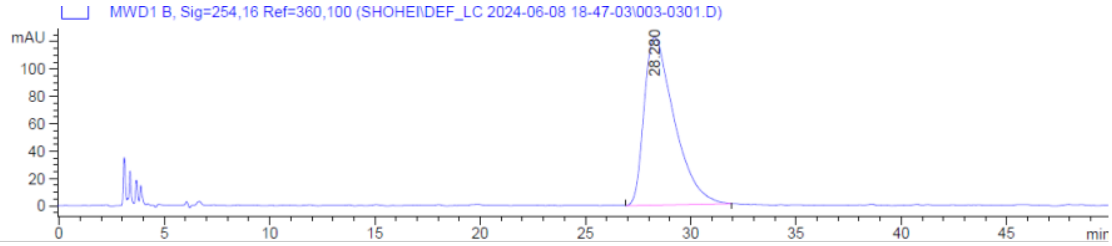

#### 6. X-ray crystallography of Compound #020_E1

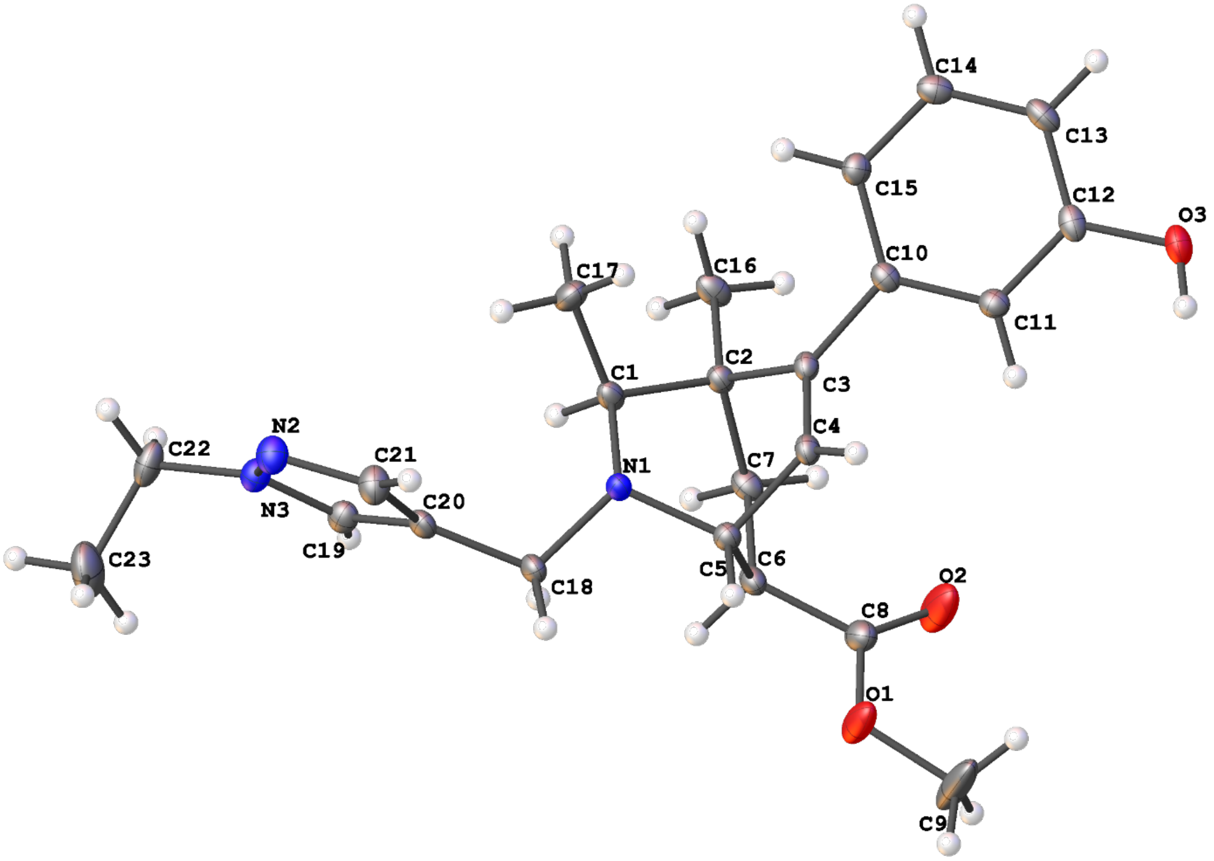

Figure 1. The complete numbering scheme of 007c-24016 with 50% thermal ellipsoid probability levels. The hydrogen atoms are shown as circles for clarity.

**Table 1 Crystal data and structure refinement for 007c-24016.**

| | |
|---|---|
| Identification code | 007c-24016 |
| Empirical formula | $C_{29}H_{43}N_3O_3$ |
| Formula weight | 395.49 |
| Temperature/K | 99.96(18) |
| Crystal system | monoclinic |
| Space group | $P2_1$ |
| a/Å | 9.9513(3) |
| b/Å | 10.1199(3) |
| c/Å | 11.5512(3) |
| α/° | 90 |
| β/° | 102.275(3) |

| | |
|---|---|
| γ/° | 90 |
| Volume/Å$^3$ | 1136.68(6) |
| Z | 2 |
| $\rho_{calc}$g/cm$^3$ | 1.156 |
| μ/mm$^{-1}$ | 0.077 |
| F(000) | 424.0 |
| Crystal size/mm$^3$ | 0.2 × 0.05 × 0.02 |
| Radiation | Mo Kα (λ = 0.71073) |
| 2Θ range for data collection/° | 5.81 to 56.56 |
| Index ranges | -13 ≤ h ≤ 13, -13 ≤ k ≤ 13, -15 ≤ l ≤ 15 |
| Reflections collected | 38297 |
| Independent reflections | 5639 [$R_{int}$ = 0.0452, $R_{sigma}$ = 0.0252] |
| Data/restraints/parameters | 5639/1/267 |
| Goodness-of-fit on F$^2$ | 1.034 |
| Final R indexes [I>=2σ (I)] | $R_1$ = 0.0372, w$R_2$ = 0.0940 |
| Final R indexes [all data] | $R_1$ = 0.0391, w$R_2$ = 0.0950 |
| Largest diff. peak/hole / e Å$^{-3}$ | 0.27/-0.19 |
| Flack parameter | 0.0(3) |

**Table 2 Fractional Atomic Coordinates (×10$^4$) and Equivalent Isotropic Displacement Parameters (Å$^2$×10$^3$) for 007c-24016. $U_{eq}$ is defined as 1/3 of the trace of the orthogonalised $U_{IJ}$ tensor.**

| Atom | *x* | *y* | *z* | U(eq) |
|---|---|---|---|---|
| O1 | 7281(2) | 1958.1(18) | 4284.3(16) | 38.5(5) |
| O2 | 7780(3) | 1501(2) | 2540.0(18) | 47.5(6) |
| O3 | 12784.4(15) | 3358.0(16) | 1321.6(14) | 22.7(3) |
| N1 | 6286.0(16) | 5875.0(16) | 3063.9(14) | 13.5(3) |
| N2 | 3548.6(18) | 9244.1(18) | 3480.9(16) | 20.3(4) |
| N3 | 2558.1(17) | 8558.5(17) | 2727.4(15) | 17.4(3) |
| C1 | 6043.5(19) | 6123(2) | 1754.6(16) | 15.0(4) |

**Table 2 Fractional Atomic Coordinates (×10[4]) and Equivalent Isotropic Displacement Parameters (Å$^2$×10$^3$) for 007c-24016. $U_{eq}$ is defined as 1/3 of the trace of the orthogonalised $U_{IJ}$ tensor.**

| Atom | *x* | *y* | *z* | U(eq) |
|---|---|---|---|---|
| C2 | 6799.6(18) | 5054.1(19) | 1161.9(16) | 14.1(3) |
| C3 | 8314.8(18) | 5042.6(18) | 1817.4(16) | 13.3(3) |
| C4 | 8454.4(18) | 4844.6(18) | 2985.5(17) | 14.6(4) |
| C5 | 7130.2(19) | 4665.8(19) | 3402.8(17) | 14.1(3) |
| C6 | 6397.3(19) | 3427.7(19) | 2746.1(17) | 15.5(4) |
| C7 | 6197.7(19) | 3691(2) | 1411.9(17) | 16.7(4) |
| C8 | 7224(2) | 2191(2) | 3139.1(19) | 21.2(4) |
| C9 | 8005(5) | 773(3) | 4755(3) | 61.9(11) |
| C10 | 9517.2(18) | 5136(2) | 1243.3(16) | 14.2(3) |
| C11 | 10577.7(19) | 4203.8(19) | 1553.6(17) | 15.0(3) |
| C12 | 11722.8(19) | 4236(2) | 1030.7(17) | 16.3(4) |
| C13 | 11804.3(19) | 5180(2) | 167.7(17) | 17.7(4) |
| C14 | 10770.1(19) | 6126(2) | -120.0(17) | 17.3(4) |
| C15 | 9638.8(19) | 6118.1(19) | 417.9(17) | 15.6(4) |
| C16 | 6511.5(19) | 5231(2) | -185.6(17) | 18.4(4) |
| C17 | 6480(2) | 7532(2) | 1536.9(19) | 21.7(4) |
| C18 | 4995.5(19) | 5830.4(19) | 3513.9(17) | 15.7(4) |
| C19 | 2916(2) | 7286(2) | 2629.0(17) | 16.8(4) |
| C20 | 4206.4(19) | 7109(2) | 3333.1(17) | 15.6(4) |
| C21 | 4549(2) | 8351(2) | 3842.7(19) | 21.4(4) |
| C22 | 1221(2) | 9162(2) | 2246.0(19) | 23.4(4) |
| C23 | 186(3) | 8854(3) | 3003(3) | 39.1(6) |

**Table 3 Anisotropic Displacement Parameters (Å$^2$×10$^3$) for 007c-24016. The Anisotropic displacement factor exponent takes the form: -$2\pi^2[h^2a^{*2}U_{11}+2hka^*b^*U_{12}+\ldots]$.**

| Atom | $U_{11}$ | $U_{22}$ | $U_{33}$ | $U_{23}$ | $U_{13}$ | $U_{12}$ |
|---|---|---|---|---|---|---|
| O1 | 63.0(12) | 29.9(9) | 29.5(9) | 11.6(7) | 25.0(8) | 28.5(9) |

**Table 3 Anisotropic Displacement Parameters (Å$^2$×10$^3$) for 007c-24016. The Anisotropic displacement factor exponent takes the form: -2π$^2$[h$^2$a*$^2$U$_{11}$+2hka*b*U$_{12}$+…].**

| Atom | U11 | U22 | U33 | U23 | U13 | U12 |
|---|---|---|---|---|---|---|
| O2 | 79.3(15) | 36.3(11) | 35.1(10) | 10.2(8) | 30.5(10) | 34.6(10) |
| O3 | 16.3(6) | 25.0(8) | 28.0(8) | -1.3(6) | 7.3(6) | 5.8(6) |
| N1 | 13.9(7) | 14.8(8) | 12.4(7) | 1.4(6) | 4.4(5) | 2.8(6) |
| N2 | 19.1(8) | 18.2(8) | 23.0(9) | -3.6(7) | 3.0(6) | 0.9(7) |
| N3 | 18.4(8) | 16.6(8) | 16.6(8) | -0.6(6) | 2.8(6) | 2.5(6) |
| C1 | 12.2(8) | 20.6(9) | 12.2(8) | 1.7(7) | 2.8(6) | 2.6(7) |
| C2 | 12.8(8) | 16.5(9) | 13.1(8) | 0.6(7) | 3.2(6) | 1.1(7) |
| C3 | 10.7(7) | 11.7(8) | 17.6(8) | -0.5(7) | 3.2(6) | 0.3(6) |
| C4 | 11.0(8) | 14.6(9) | 18.3(9) | -0.7(7) | 3.5(6) | 1.1(6) |
| C5 | 14.2(8) | 14.7(8) | 13.5(8) | 0.3(6) | 3.6(6) | 1.2(7) |
| C6 | 14.3(8) | 14.5(9) | 19.5(9) | -0.8(7) | 7.5(7) | 0.4(7) |
| C7 | 14.7(8) | 18.4(9) | 17.6(9) | -3.4(7) | 5.1(7) | -2.6(7) |
| C8 | 25.2(10) | 17.0(10) | 24.2(10) | 1.8(8) | 11.3(8) | 2.6(8) |
| C9 | 110(3) | 46.0(18) | 41.0(15) | 27.0(14) | 41.6(18) | 55(2) |
| C10 | 12.5(8) | 16.7(9) | 13.5(8) | -2.4(7) | 3.0(6) | -1.4(7) |
| C11 | 15.1(8) | 14.8(8) | 15.4(8) | -0.9(7) | 3.6(6) | -1.7(7) |
| C12 | 11.9(8) | 17.2(9) | 19.4(9) | -6.9(7) | 2.2(7) | -0.8(7) |
| C13 | 14.3(8) | 23.6(10) | 16.8(9) | -5.2(8) | 6.7(7) | -3.7(7) |
| C14 | 17.3(8) | 19.8(9) | 14.6(8) | -0.6(7) | 2.7(7) | -4.9(7) |
| C15 | 13.9(8) | 16.4(9) | 15.7(8) | -0.6(7) | 1.1(6) | 0.6(7) |
| C16 | 14.8(8) | 27.8(10) | 12.3(8) | -1.1(7) | 2.6(7) | -1.7(8) |
| C17 | 27.5(10) | 17.5(10) | 21.1(10) | 6.1(7) | 7.7(8) | 5.1(8) |
| C18 | 15.6(8) | 15.6(9) | 18.0(8) | 0.1(7) | 8.4(7) | 2.3(7) |
| C19 | 19.6(9) | 17.5(10) | 13.5(8) | -1.3(7) | 4.1(7) | 2.3(7) |
| C20 | 15.7(8) | 18.2(9) | 14.8(8) | -0.3(7) | 7.7(7) | 2.1(7) |
| C21 | 18.4(9) | 20.4(10) | 24.9(10) | -3.0(8) | 3.3(8) | 1.7(8) |
| C22 | 23.9(10) | 19.7(10) | 22.8(10) | -1.9(8) | -3.7(8) | 9.2(8) |
| C23 | 22.2(11) | 49.3(17) | 44.8(15) | -1.4(12) | 5.1(10) | 13.4(11) |

**Table 4 Bond Lengths for 007c-24016.**

| Atom | Atom | Length/Å | Atom | Atom | Length/Å |
|---|---|---|---|---|---|
| O1 | C8 | 1.333(3) | C3 | C4 | 1.342(3) |
| O1 | C9 | 1.444(3) | C3 | C10 | 1.489(2) |
| O2 | C8 | 1.198(3) | C4 | C5 | 1.507(2) |
| O3 | C12 | 1.367(2) | C5 | C6 | 1.563(3) |
| N1 | C1 | 1.501(2) | C6 | C7 | 1.535(3) |
| N1 | C5 | 1.489(2) | C6 | C8 | 1.514(3) |
| N1 | C18 | 1.485(2) | C10 | C11 | 1.404(3) |
| N2 | N3 | 1.359(2) | C10 | C15 | 1.400(3) |
| N2 | C21 | 1.343(3) | C11 | C12 | 1.399(3) |
| N3 | C19 | 1.347(3) | C12 | C13 | 1.395(3) |
| N3 | C22 | 1.462(2) | C13 | C14 | 1.393(3) |
| C1 | C2 | 1.557(3) | C14 | C15 | 1.397(3) |
| C1 | C17 | 1.526(3) | C18 | C20 | 1.505(3) |
| C2 | C3 | 1.537(2) | C19 | C20 | 1.378(3) |
| C2 | C7 | 1.555(3) | C20 | C21 | 1.399(3) |
| C2 | C16 | 1.532(3) | C22 | C23 | 1.518(4) |

**Table 5 Bond Angles for 007c-24016.**

| Atom | Atom | Atom | Angle/° | Atom | Atom | Atom | Angle/° |
|---|---|---|---|---|---|---|---|
| C8 | O1 | C9 | 115.66(19) | C7 | C6 | C5 | 107.19(15) |
| C5 | N1 | C1 | 111.30(14) | C8 | C6 | C5 | 110.46(16) |
| C18 | N1 | C1 | 113.08(14) | C8 | C6 | C7 | 113.01(16) |
| C18 | N1 | C5 | 111.38(14) | C6 | C7 | C2 | 111.59(15) |
| C21 | N2 | N3 | 104.14(17) | O1 | C8 | C6 | 110.45(17) |
| N2 | N3 | C22 | 120.54(17) | O2 | C8 | O1 | 122.9(2) |
| C19 | N3 | N2 | 111.91(16) | O2 | C8 | C6 | 126.6(2) |
| C19 | N3 | C22 | 127.05(18) | C11 | C10 | C3 | 118.36(17) |
| N1 | C1 | C2 | 109.72(14) | C15 | C10 | C3 | 122.84(17) |
| N1 | C1 | C17 | 109.38(16) | C15 | C10 | C11 | 118.79(17) |
| C17 | C1 | C2 | 113.30(16) | C12 | C11 | C10 | 120.73(18) |

**Table 5 Bond Angles for 007c-24016.**

| Atom | Atom | Atom | Angle/° | Atom | Atom | Atom | Angle/° |
|---|---|---|---|---|---|---|---|
| C3 | C2 | C1 | 107.69(14) | O3 | C12 | C11 | 122.44(18) |
| C3 | C2 | C7 | 106.15(15) | O3 | C12 | C13 | 117.49(17) |
| C7 | C2 | C1 | 107.06(14) | C13 | C12 | C11 | 120.07(17) |
| C16 | C2 | C1 | 111.22(16) | C14 | C13 | C12 | 119.24(17) |
| C16 | C2 | C3 | 116.97(15) | C13 | C14 | C15 | 120.98(18) |
| C16 | C2 | C7 | 107.21(15) | C14 | C15 | C10 | 120.08(17) |
| C4 | C3 | C2 | 112.19(16) | N1 | C18 | C20 | 112.94(15) |
| C4 | C3 | C10 | 122.42(16) | N3 | C19 | C20 | 107.74(18) |
| C10 | C3 | C2 | 125.25(16) | C19 | C20 | C18 | 125.94(18) |
| C3 | C4 | C5 | 115.40(16) | C19 | C20 | C21 | 104.15(17) |
| N1 | C5 | C4 | 107.46(15) | C21 | C20 | C18 | 129.87(18) |
| N1 | C5 | C6 | 110.63(14) | N2 | C21 | C20 | 112.05(18) |
| C4 | C5 | C6 | 106.89(15) | N3 | C22 | C23 | 111.69(19) |

**Table 6 Torsion Angles for 007c-24016.**

| A | B | C | D | Angle/° | A | B | C | D | Angle/° |
|---|---|---|---|---|---|---|---|---|---|
| O3 | C12 | C13 | C14 | -177.46(17) | C5 | C6 | C7 | C2 | 1.07(19) |
| N1 | C1 | C2 | C3 | -53.70(19) | C5 | C6 | C8 | O1 | 64.4(2) |
| N1 | C1 | C2 | C7 | 60.09(18) | C5 | C6 | C8 | O2 | -114.7(3) |
| N1 | C1 | C2 | C16 | 176.92(15) | C7 | C2 | C3 | C4 | -58.7(2) |
| N1 | C5 | C6 | C7 | 59.31(18) | C7 | C2 | C3 | C10 | 117.01(19) |
| N1 | C5 | C6 | C8 | -177.17(15) | C7 | C6 | C8 | O1 | -175.52(18) |
| N1 | C18 | C20 | C19 | 116.0(2) | C7 | C6 | C8 | O2 | 5.4(3) |
| N1 | C18 | C20 | C21 | -66.6(3) | C8 | C6 | C7 | C2 | -120.87(17) |
| N2 | N3 | C19 | C20 | -0.7(2) | C9 | O1 | C8 | O2 | -3.0(4) |
| N2 | N3 | C22 | C23 | -92.6(2) | C9 | O1 | C8 | C6 | 177.9(3) |
| N3 | N2 | C21 | C20 | -0.2(2) | C10 | C3 | C4 | C5 | -175.27(17) |
| N3 | C19 | C20 | C18 | 178.50(17) | C10 | C11 | C12 | O3 | 179.05(18) |
| N3 | C19 | C20 | C21 | 0.6(2) | C10 | C11 | C12 | C13 | -1.7(3) |
| C1 | N1 | C5 | C4 | 56.77(19) | C11 | C10 | C15 | C14 | 2.8(3) |

**Table 6 Torsion Angles for 007c-24016.**

| A | B | C | D | Angle/° | A | B | C | D | Angle/° |
|---|---|---|---|---|---|---|---|---|---|
| C1 | N1 | C5 | C6 | -59.58(18) | C11 | C12 | C13 | C14 | 3.3(3) |
| C1 | N1 | C18 | C20 | -61.5(2) | C12 | C13 | C14 | C15 | -1.8(3) |
| C1 | C2 | C3 | C4 | 55.7(2) | C13 | C14 | C15 | C10 | -1.2(3) |
| C1 | C2 | C3 | C10 | -128.60(18) | C15 | C10 | C11 | C12 | -1.3(3) |
| C1 | C2 | C7 | C6 | -59.05(18) | C16 | C2 | C3 | C4 | -178.20(18) |
| C2 | C3 | C4 | C5 | 0.5(2) | C16 | C2 | C3 | C10 | -2.5(3) |
| C2 | C3 | C10 | C11 | -130.02(19) | C16 | C2 | C7 | C6 | -178.49(15) |
| C2 | C3 | C10 | C15 | 50.5(3) | C17 | C1 | C2 | C3 | 68.86(19) |
| C3 | C2 | C7 | C6 | 55.78(19) | C17 | C1 | C2 | C7 | -177.35(15) |
| C3 | C4 | C5 | N1 | -58.7(2) | C17 | C1 | C2 | C16 | -60.5(2) |
| C3 | C4 | C5 | C6 | 60.1(2) | C18 | N1 | C1 | C2 | -128.17(16) |
| C3 | C10 | C11 | C12 | 179.22(17) | C18 | N1 | C1 | C17 | 106.97(18) |
| C3 | C10 | C15 | C14 | -177.80(18) | C18 | N1 | C5 | C4 | -176.04(15) |
| C4 | C3 | C10 | C11 | 45.2(3) | C18 | N1 | C5 | C6 | 67.61(18) |
| C4 | C3 | C10 | C15 | -134.2(2) | C18 | C20 | C21 | N2 | -178.07(18) |
| C4 | C5 | C6 | C7 | -57.39(18) | C19 | N3 | C22 | C23 | 78.7(3) |
| C4 | C5 | C6 | C8 | 66.13(19) | C19 | C20 | C21 | N2 | -0.2(2) |
| C5 | N1 | C1 | C2 | -1.9(2) | C21 | N2 | N3 | C19 | 0.5(2) |
| C5 | N1 | C1 | C17 | -126.77(16) | C21 | N2 | N3 | C22 | 173.05(19) |
| C5 | N1 | C18 | C20 | 172.25(15) | C22 | N3 | C19 | C20 | -172.63(19) |

**Table 7 Hydrogen Atom Coordinates (Å×10[4]) and Isotropic Displacement Parameters (Å[2]×10[3]) for 007c-24016.**

| Atom | *x* | *y* | *z* | U(eq) |
|---|---|---|---|---|
| H3 | 12773 | 3014 | 1981 | 34 |
| H1 | 5035 | 6041 | 1418 | 18 |
| H4 | 9328 | 4819 | 3513 | 18 |
| H5 | 7319 | 4532 | 4280 | 17 |
| H6 | 5473 | 3334 | 2947 | 19 |
| H7A | 5204 | 3668 | 1044 | 20 |

**Table 7 Hydrogen Atom Coordinates (Å×10^4) and Isotropic Displacement Parameters (Å^2×10^3) for 007c-24016.**

| Atom | *x* | *y* | *z* | U(eq) |
|---|---|---|---|---|
| H7B | 6656 | 2985 | 1046 | 20 |
| H9A | 7529 | -1 | 4353 | 93 |
| H9B | 8030 | 713 | 5606 | 93 |
| H9C | 8946 | 803 | 4626 | 93 |
| H11 | 10517 | 3544 | 2125 | 18 |
| H13 | 12557 | 5177 | -219 | 21 |
| H14 | 10835 | 6785 | -690 | 21 |
| H15 | 8951 | 6780 | 223 | 19 |
| H16A | 6861 | 6092 | -376 | 28 |
| H16B | 5518 | 5187 | -505 | 28 |
| H16C | 6972 | 4527 | -537 | 28 |
| H17A | 5969 | 8157 | 1928 | 32 |
| H17B | 6282 | 7709 | 683 | 32 |
| H17C | 7468 | 7634 | 1859 | 32 |
| H18A | 5223 | 5620 | 4370 | 19 |
| H18B | 4404 | 5112 | 3104 | 19 |
| H19 | 2376 | 6629 | 2158 | 20 |
| H21 | 5389 | 8541 | 4381 | 26 |
| H22A | 868 | 8830 | 1432 | 28 |
| H22B | 1333 | 10131 | 2202 | 28 |
| H23A | -685 | 9301 | 2673 | 59 |
| H23B | 541 | 9165 | 3814 | 59 |
| H23C | 32 | 7898 | 3010 | 59 |

**Table 8 Solvent masks information for 007c-24016.**

| Number | X | Y | Z | Volume | Electron count | Content |
|---|---|---|---|---|---|---|
| 1 | 0.183 | 0.232 | 0.436 | 65.1 | 20.4 | 1 C6H14 |
| 2 | -0.183 | 0.732 | 0.564 | 65.1 | 20.4 | 1 C6H14 |

#### 7. Pharmacokinetic Report of Compound (±)-#020

| Sample | Administration | Dose, mg/kg | Pharmacokinetic Parameters | | | | | |
|---|---|---|---|---|---|---|---|---|
| | | | Tmax, min | Cmax, ng/ml | $AUC_{0\to t\ min}$ (AUClast) ng*min/ml | $AUC_{0\to\infty}$ (AUCINF_obs) ng*min/ml | $T_{1/2}$ (HL_Lambda_z), min | $K_{el}$ (Lambda_z), $min^{-1}$ |
| Plasma | IP | 10 | 5.00 | 1410 | 39800 | 40000 | 15.7 | 0.0443 |
| Brain | | | 15.0 | 174 | 5880 | 8600 | 31.5 | 0.022 |
| SCF | | | 5.00 | 48.6 | 1460 | 2340 | 34.8 | 0.0199 |

**Table 1. Selected pharmacokinetic parameters for compound (±)-#020 in male CD-1 mice**

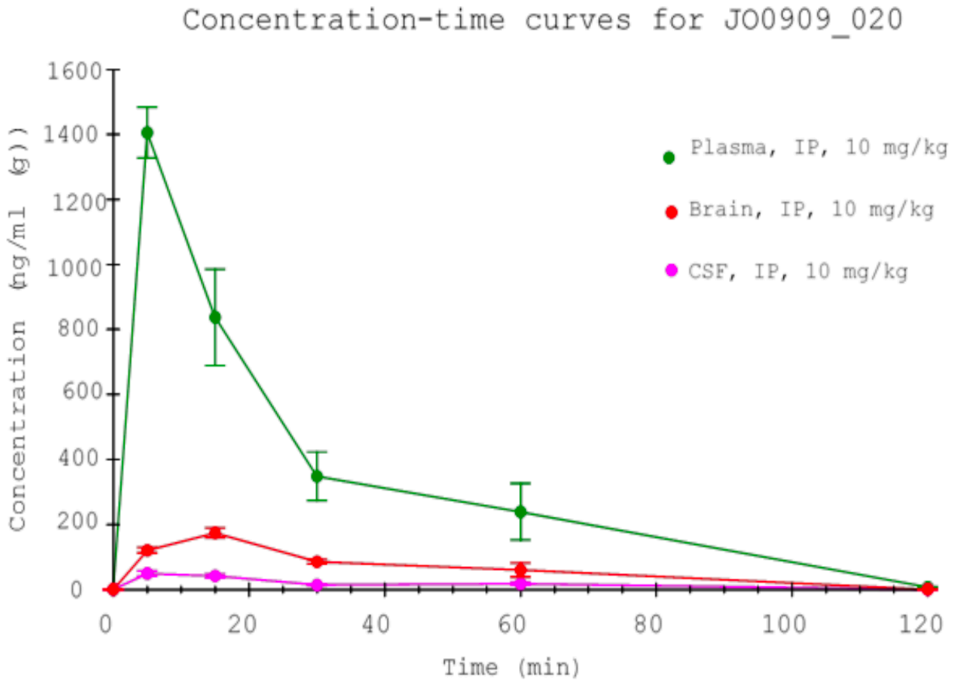

**Figure 1. Concentration-time curves for compound (±)-#020 in male CD-1 mice following IP administration**

#### 8. CryoEM Validation Statistics

| | **MOR with #33** | **MOR-Nb6M-NabFab-AntiNb Fab complex with #020_E1** | **MOR with #020_E1** | **KOR-Nb6M-NabFab-AntiNb Fab complex with #020_E1** | **KOR with #020_E1** |
|---|---|---|---|---|---|
| **Data Collection** | | | | | |
| Microscope | UCSF EM Core, Titan Krios G3i | UCSF EM Core, Titan Krios G3i | | UCSF EM Core, Talos Arctica | |
| Detector | K3 | K3 | | K3 | |
| Voltage (kV) | 300 | 300 | | 200 | |
| Magnification | 105,000x | 105,000x | | 45,000x | |
| Defocus range (µm) | 0.8 to 2.1 | 0.8 to 2.1 | | 0.8 to 2.1 | |
| Pixel size (Å) | 0.835 | 0.8189 | | 0.865 | |
| Total exposure ($e^-$/Å$^2$) | 45.8 | 47.7 | | 64.1 | |
| Frame exposure ($e^-$/Å$^2$/frame) | 0.57 | 0.596 | | 0.855 | |
| # Frames/Image | 80 | 80 | | 75 | |
| # Images | 7.766 | 9,877 | | 4,967 | |
| # Initial Particles | 1,923,561 | 5,949,666 | | 2,056,565 | |
| # Final Particles | 40,722 | 259,816 | | 98,518 | |
| Symmetry imposed | C1 | C1 | | C1 | |
| Map sharpening *B* factor (Å$^2$) | -132 | -140 | -127 | -114 | -100 |
| Map resolution (Å) | 3.90 | 3.30 | 3.23 | 3.18 | 2.96 |
| FSC threshold | 0.143 | 0.143 | 0.143 | 0.143 | 0.143 |
| **Refinement and validation** | | | | | |

| | | | | | |
|---|---|---|---|---|---|
| Initial model used (PDB) | 7UL4 | 7UL4, 7UL3, 7PIJ | 7UL4 | AF-P41145-F1(Dimer), 7UL3, 7PIJ | AF-P41145-F1 (Monomer) |
| Model resolution (Å) | 4.1 | 3.8 | 3.7 | 3.3 | 3.1 |
| FSC threshold | 0.5 | 0.5 | 0.5 | 0.5 | 0.5 |
| Model composition | | | | | |
| Chains | 1 | 5 | 2 | 7 | 2 |
| Non-hydrogen atoms | 2074 | 5654 | 2134 | 6922 | 2303 |
| Protein residues | 281 | 785 | 286 | 896 | 285 |
| Water | 0 | 0 | 0 | 0 | 0 |
| Ligands | 0 | 1 | 1 | 2 | 1 |
| *B* factors (Å$^2$) | | | | | |
| Protein (min/max/mean) | 34.66/112.17/70.96 | 3.99/148.44/65.32 | 13.63/128.54/64.06 | 31.90/236.86/116.89 | 25.49/132.65/55.20 |
| Ligand (min/max/mean) | N/A | 44.92/77.28/56.34 | 20.00/20.00/20.00 | 54.37/87.33/72.54 | 40.41/80.23/58.60 |
| R.m.s. deviations | | | | | |
| Bond lengths (Å) | 0.003 | 0.002 | 0.007 | 0.003 | 0.003 |
| Bond angles (°) | 0.588 | 0.572 | 1.081 | 0.593 | 0.533 |
| **Validation** | | | | | |
| MolProbity score | 1.97 | 1.70 | 1.51 | 2.07 | 1.66 |
| Clash score | 4.68 | 7.56 | 7.48 | 8.72 | 5.35 |
| EMRinger score | 1.35 | 1.30 | 2.26 | 1.91 | 2.45 |
| Poor rotamers (%) | 4.90 | 2.38 | 0.97 | 3.85 | 1.95 |

| | | | | | |
|---|---|---|---|---|---|
| Ramachandran plot | | | | | |
| Favored (%) | 96.75 | 97.99 | 97.52 | 97.11 | 97.15 |
| Allowed (%) | 3.25 | 2.01 | 2.48 | 2.89 | 2.85 |
| Disallowed (%) | 0 | 0 | 0 | 0 | 0 |
