## Supplementary material for "Docking 14 million virtual isoquinuclidines against the mu and kappa opioid receptors reveals dual antagonists-inverse agonists with reduced withdrawal effects": NMR spectra

#### Proton (#03)

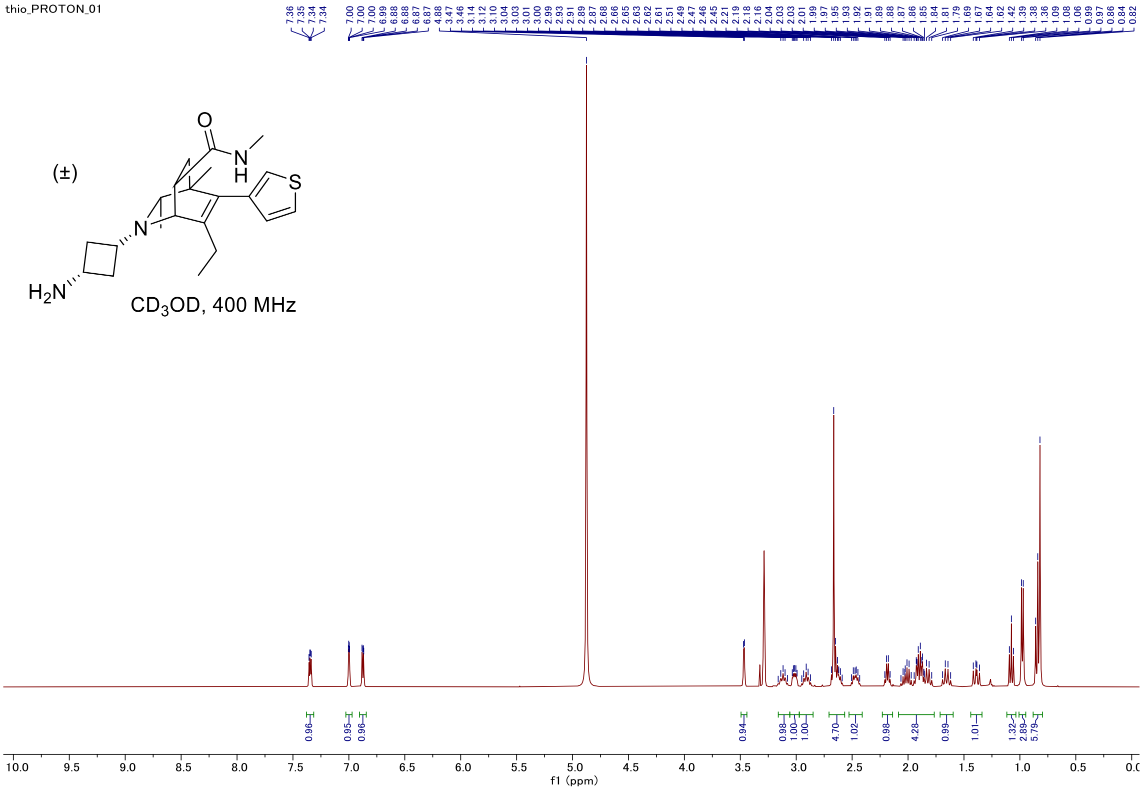


#### Carbon

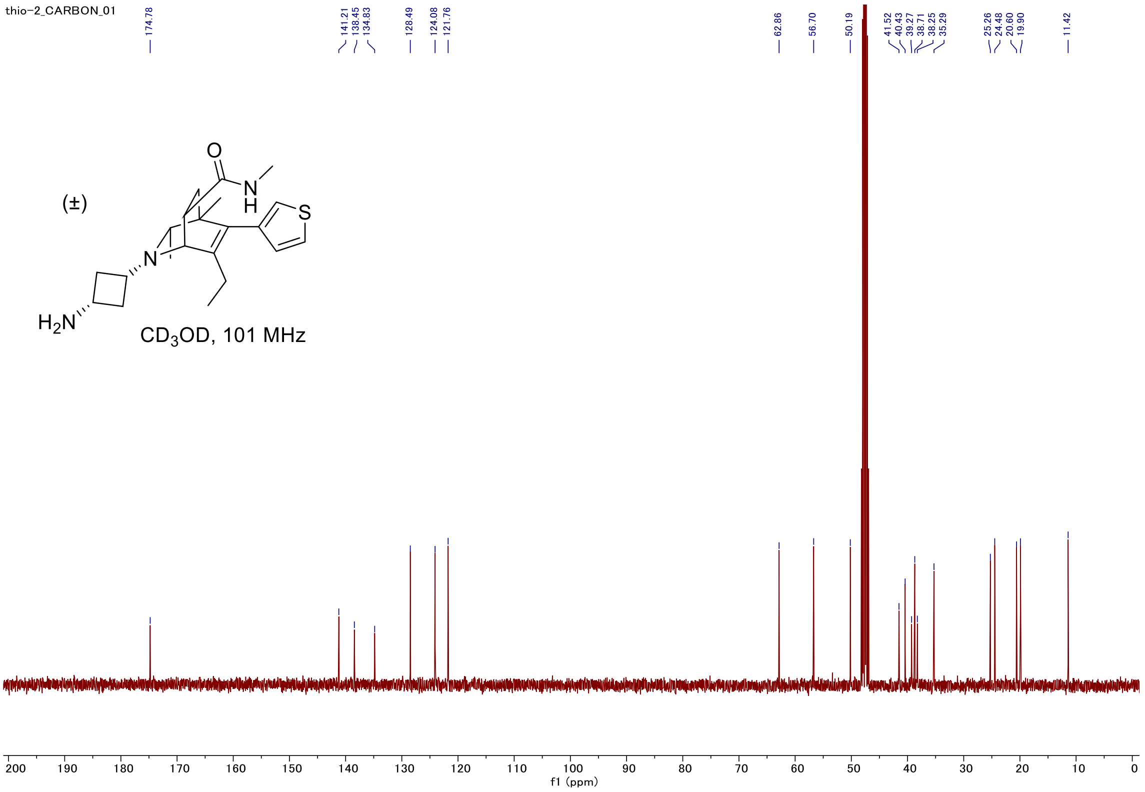

#### C-H COSY

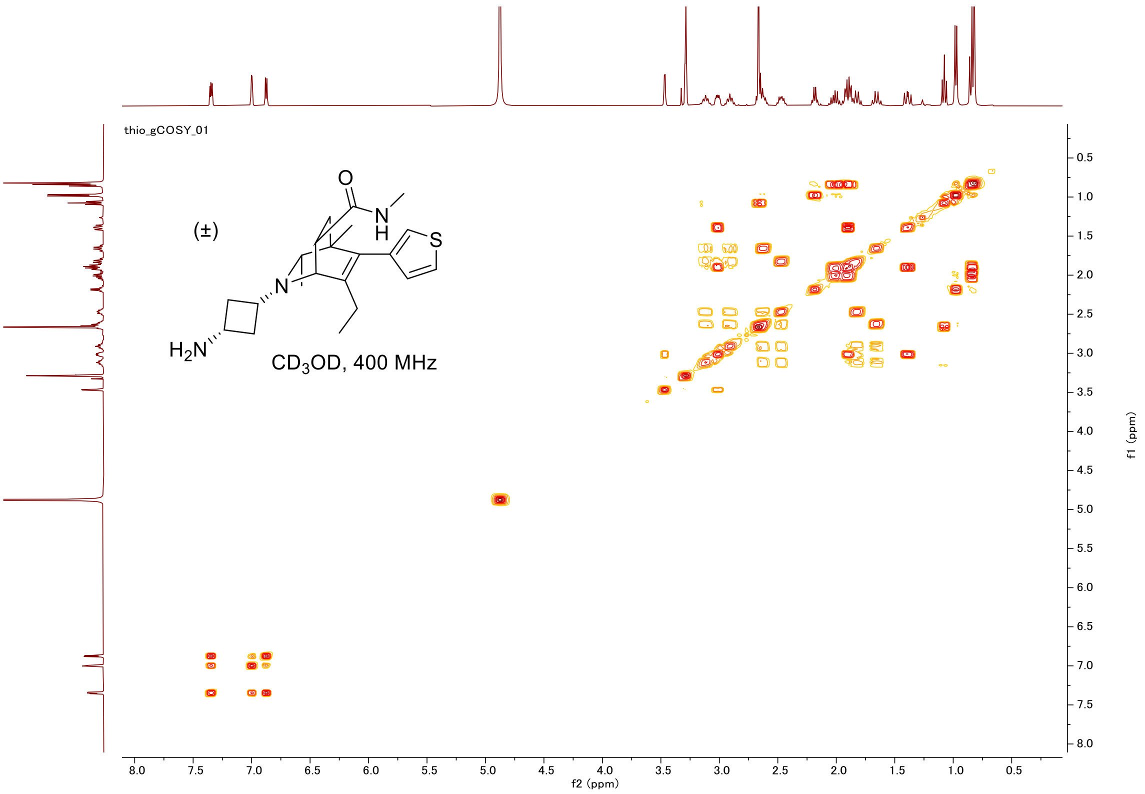

thio_gCOSY_01
(±)
O
N
H
S
N
H2N
CD3OD, 400 MHz
f2 (ppm)
f1 (ppm)


#### HSQC

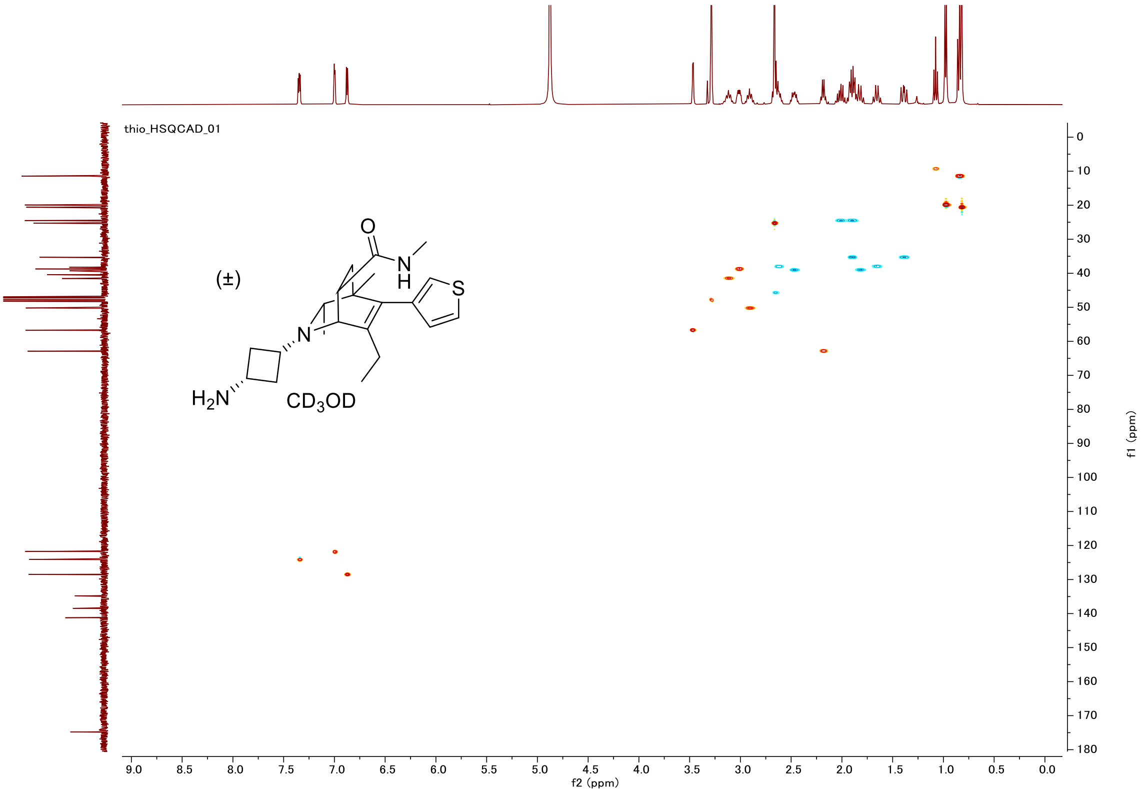

thio_HSQCAD_01
(±)
O
N
H
S
N
H2N
CD3OD
f2 (ppm)
f1 (ppm)

### Proton (#05 isomer 1)

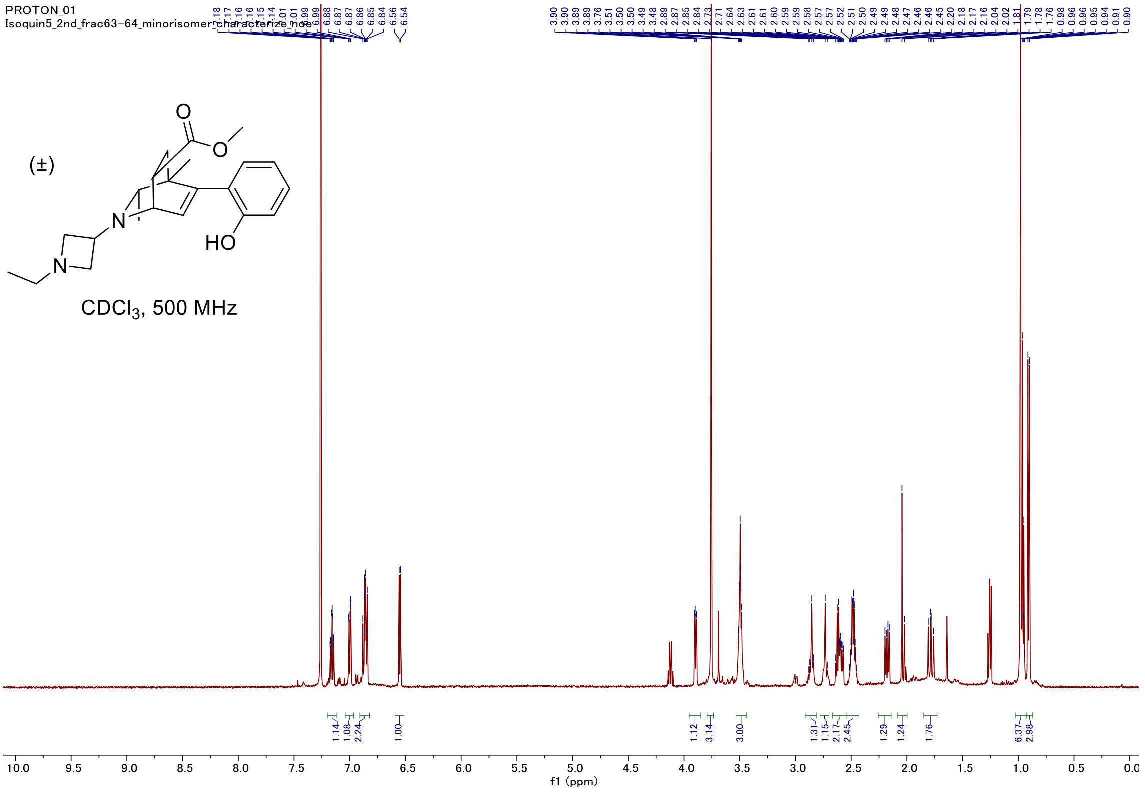


### Carbon

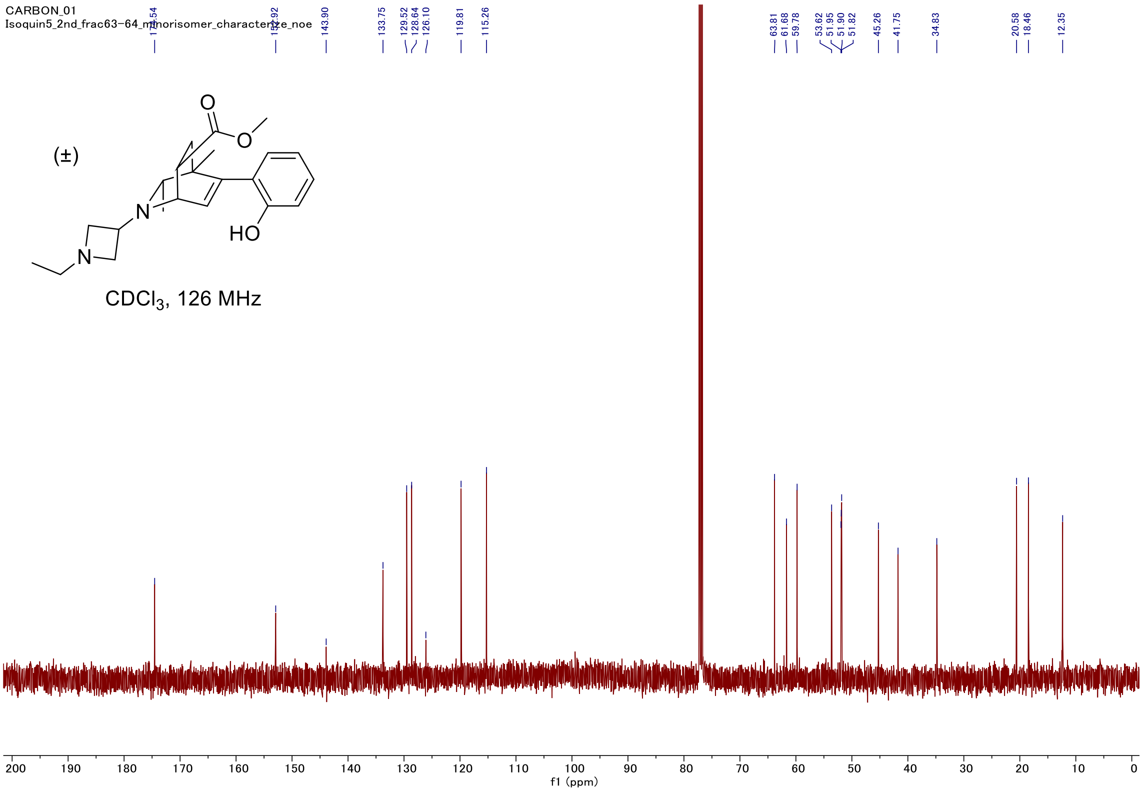

#### C-H COSY

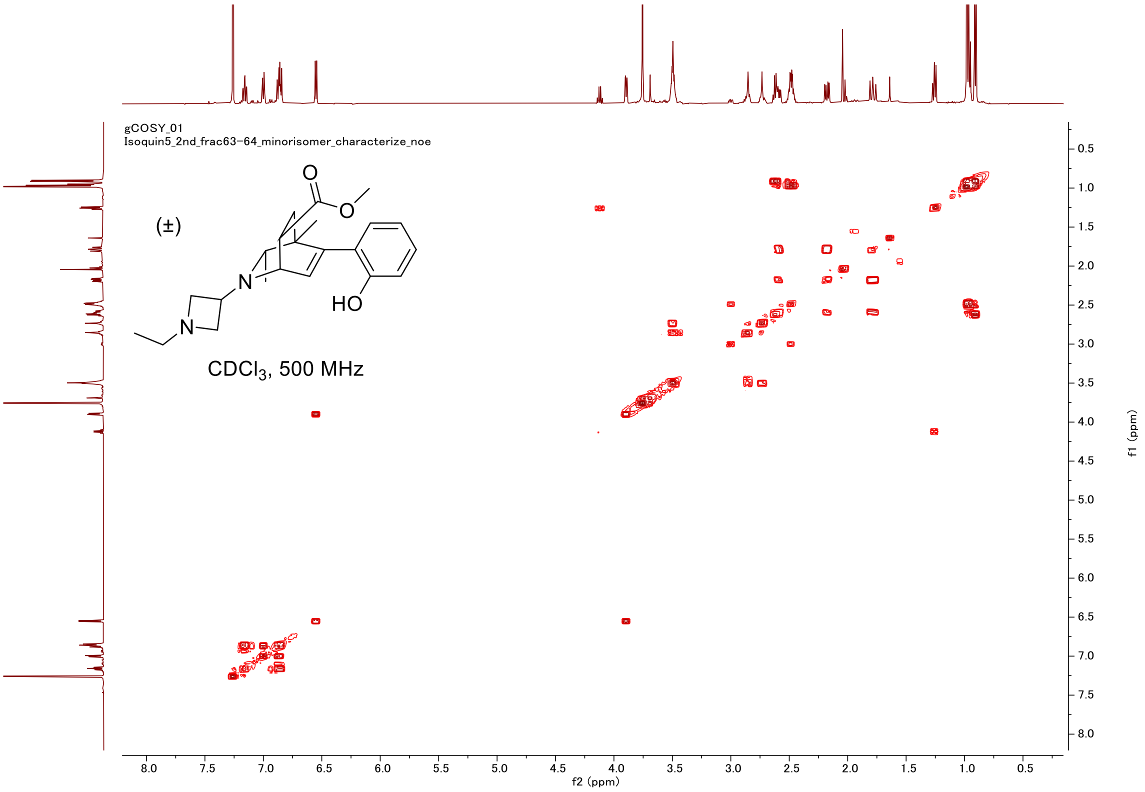
gCOSY_01
Isoquin5_2nd_frac63-64_minorisomer_characterize_noe
(±)
O
O
N
N
HO
CDCl3, 500 MHz
f2 (ppm)
f1 (ppm)

#### HSQC

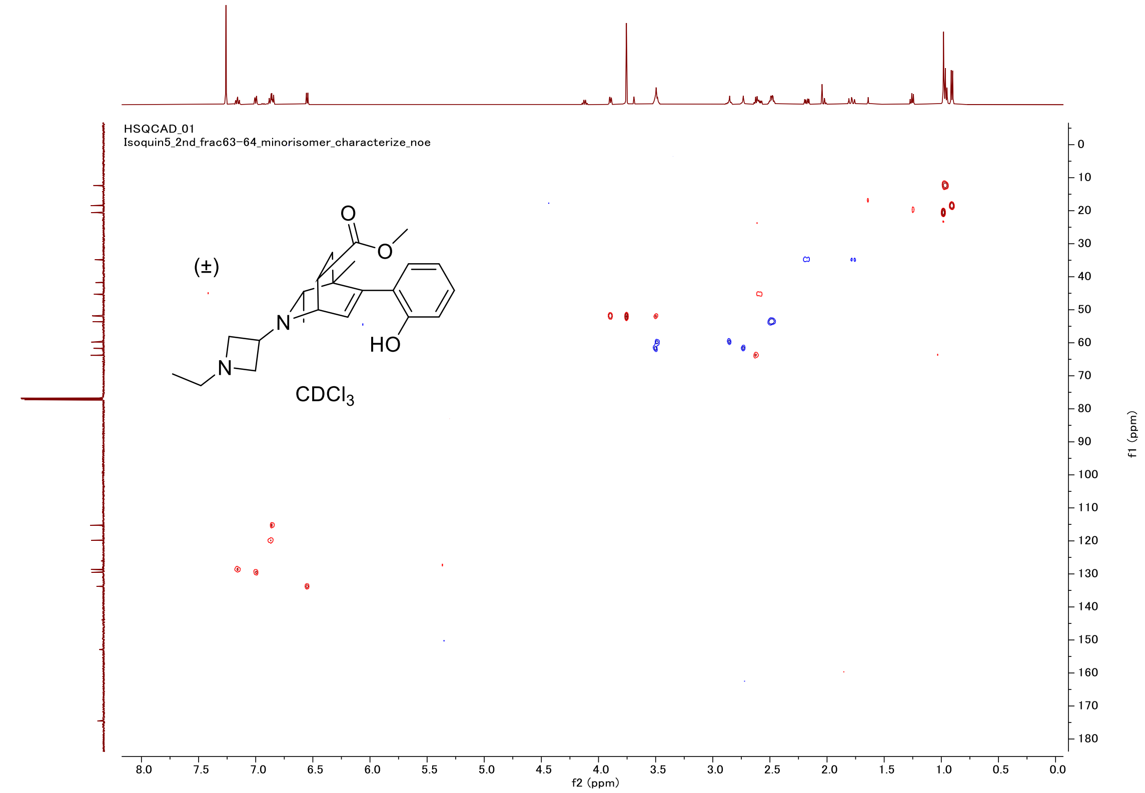
HSQCAD_01
Isoquin5_2nd_frac63-64_minorisomer_characterize_noe
(±)
O
O
N
N
HO
CDCl3
f2 (ppm)
f1 (ppm)

#### Proton (#05 isomer 2)

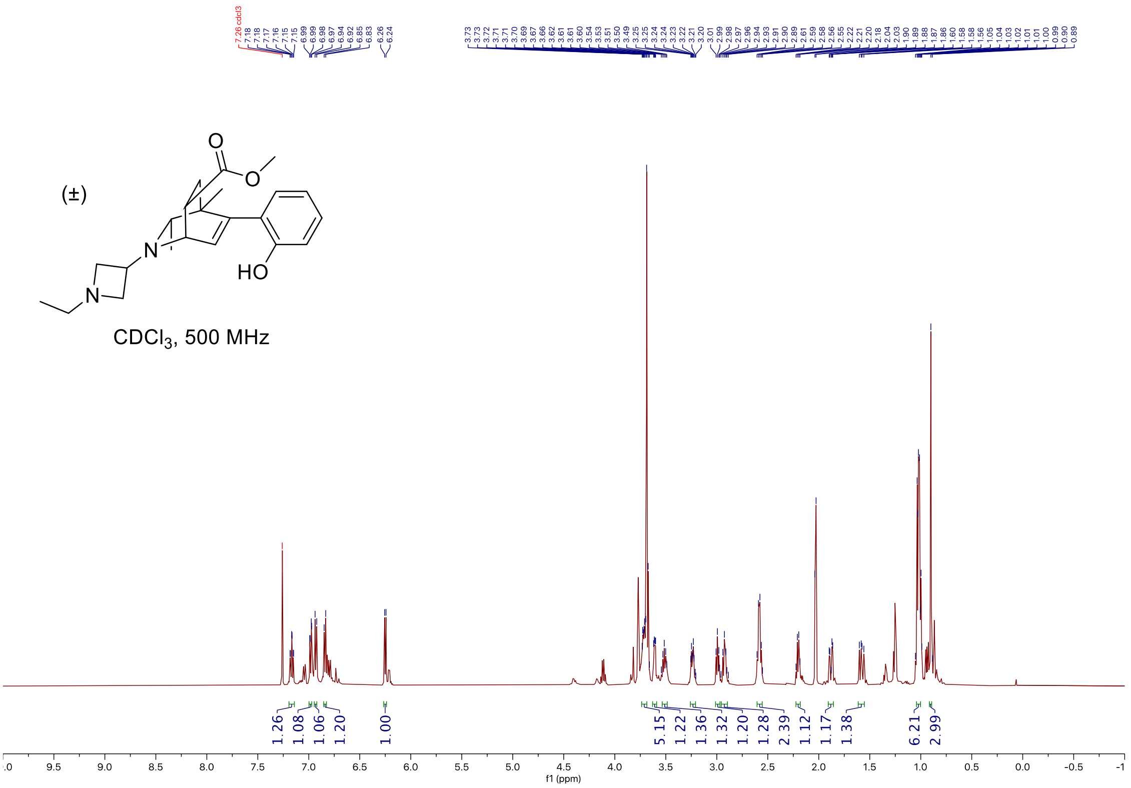


#### Carbon

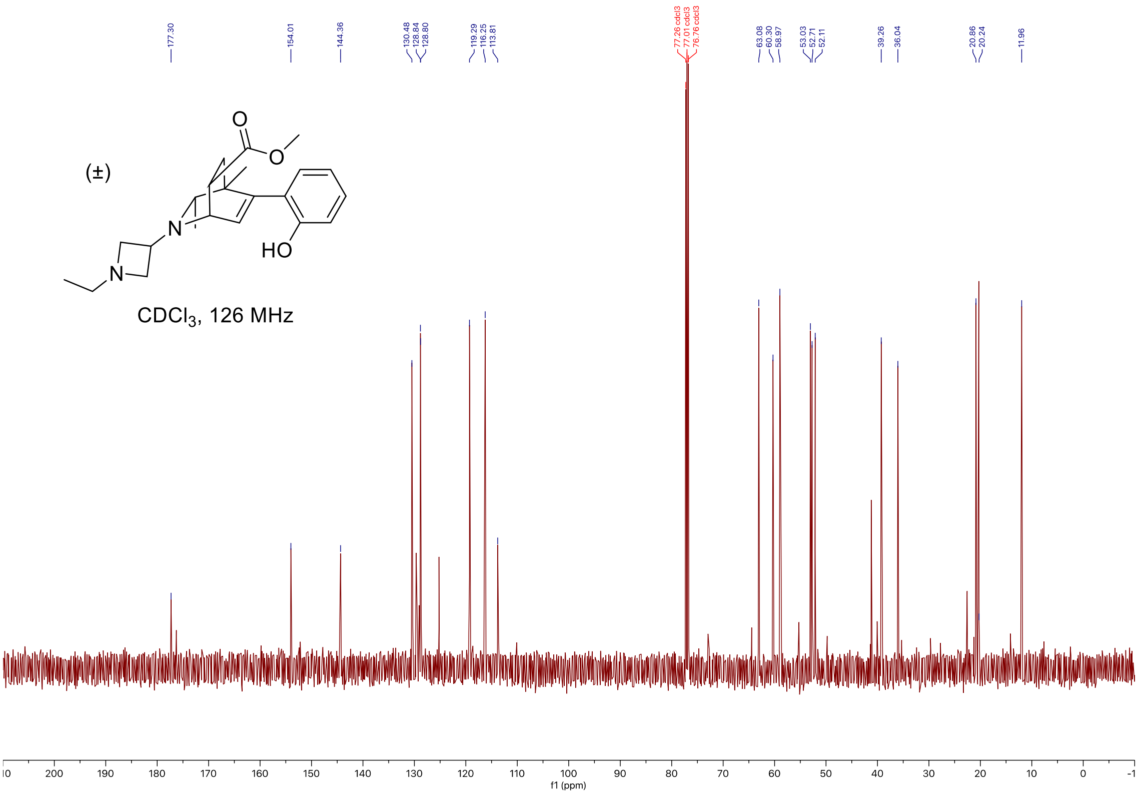

#### C-H COSY

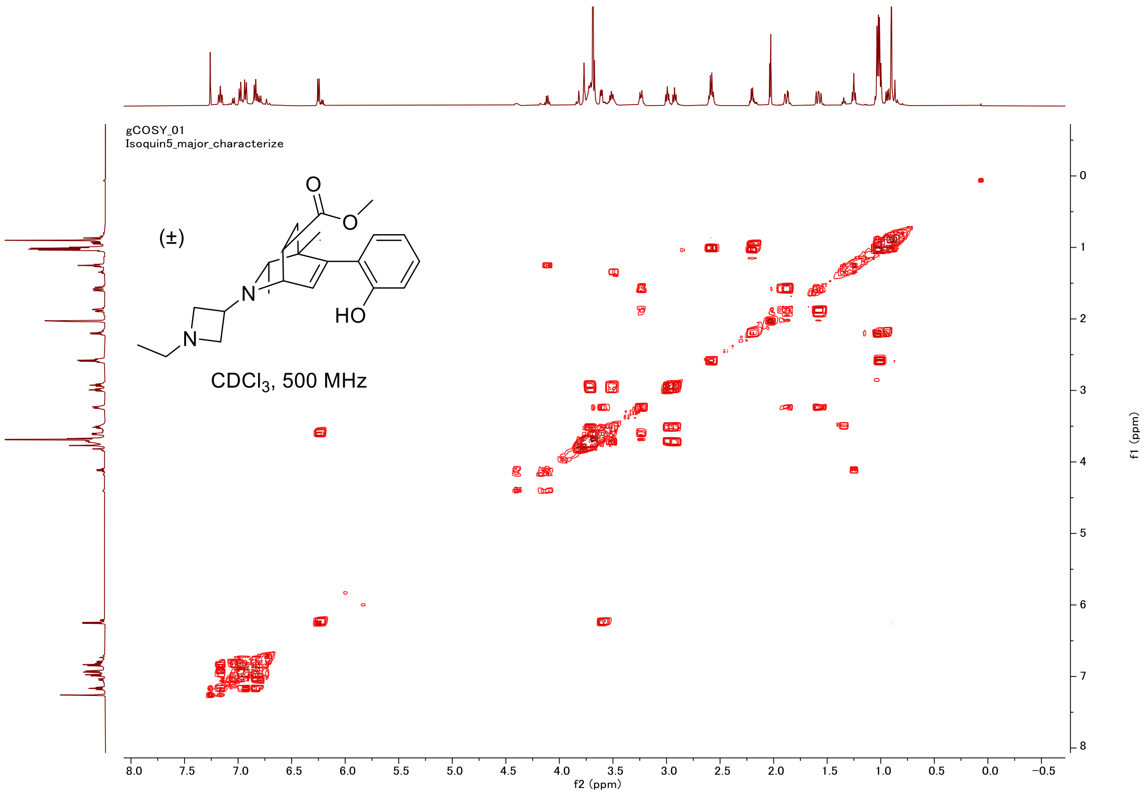
gCOSY_01
Isoquin5_major_characterize
(±)
O
O
N
N
HO
CDCl3, 500 MHz
f2 (ppm)
f1 (ppm)

#### HSQC

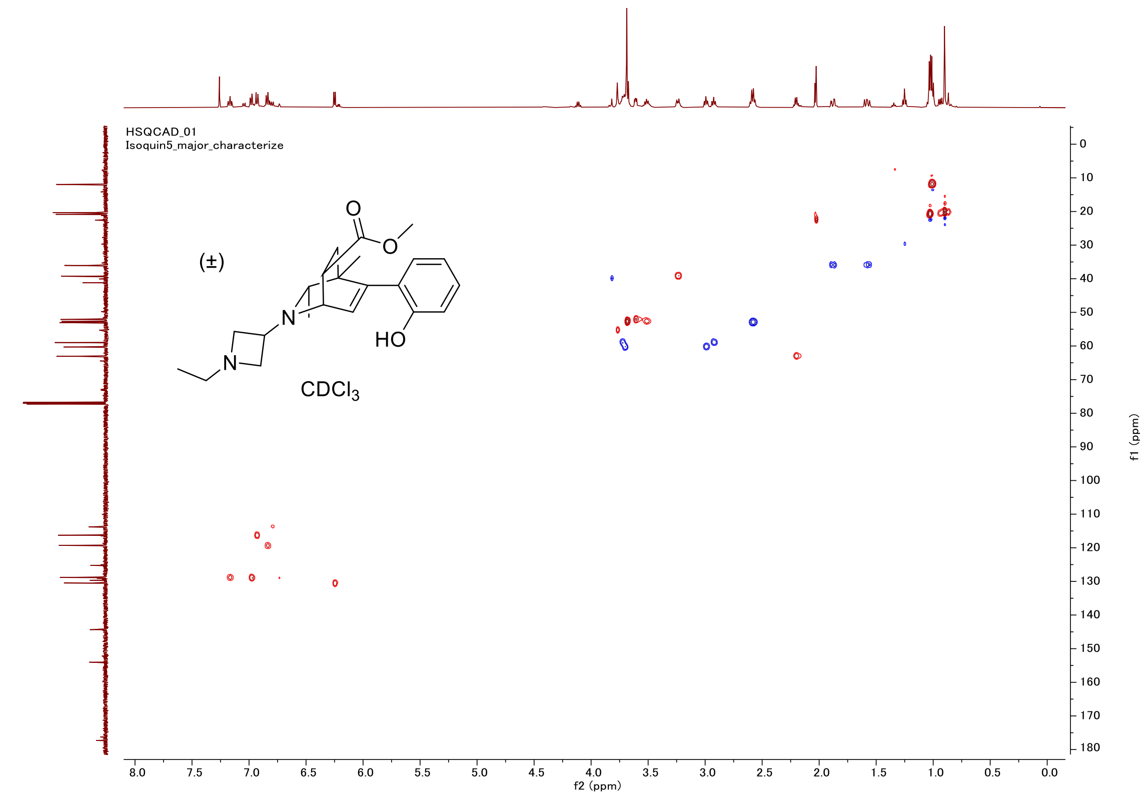
HSQCAD_01
Isoquin5_major_characterize
(±)
O
O
N
N
HO
CDCl3
f2 (ppm)
f1 (ppm)

#### Proton (#06)

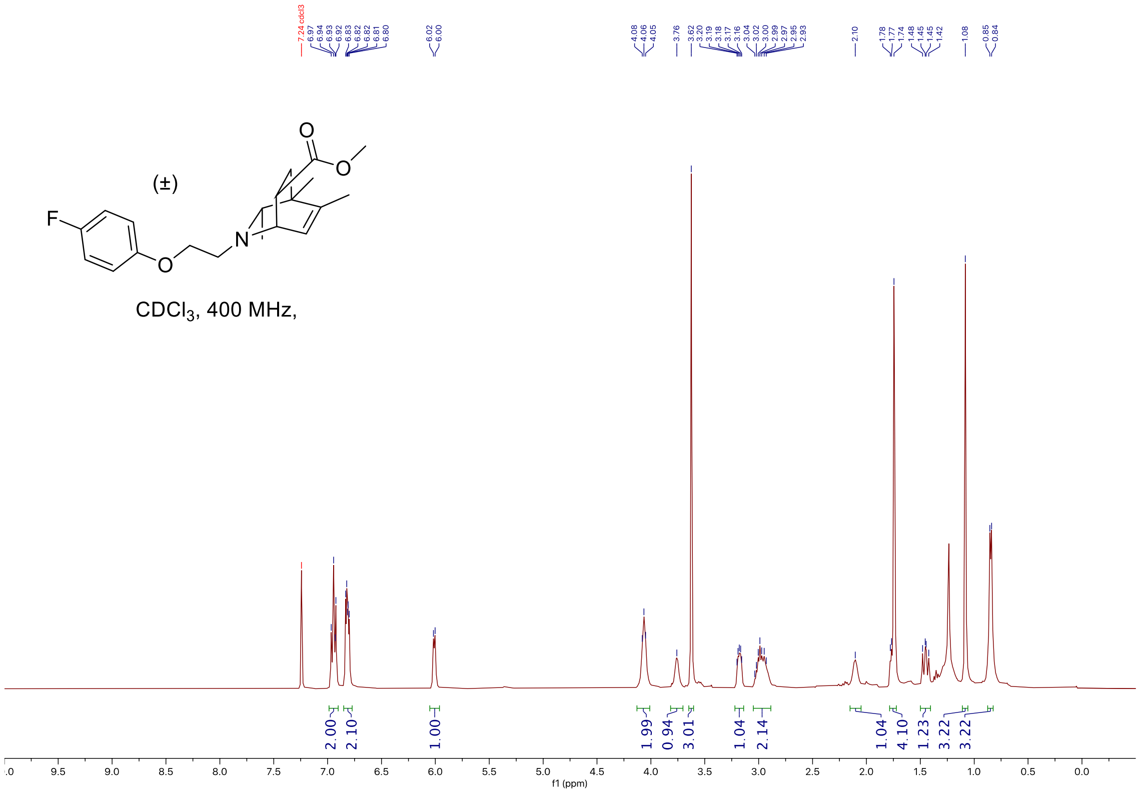


#### Carbon

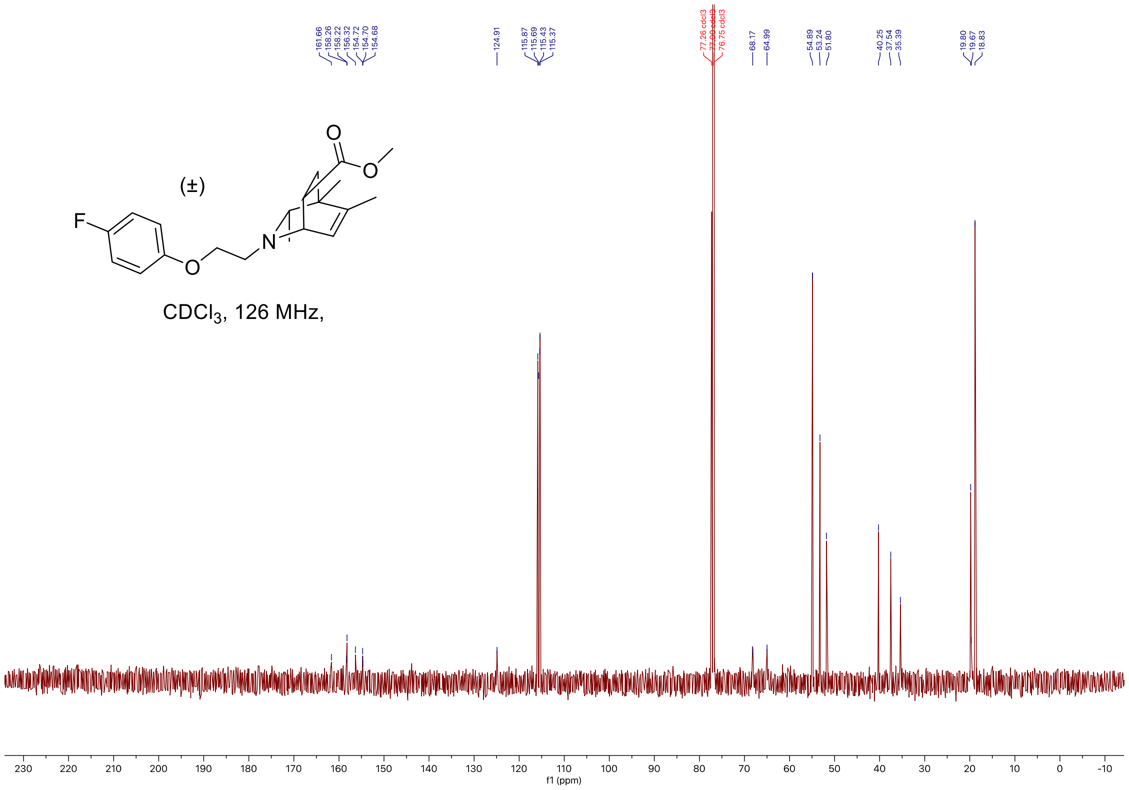

#### Fluorine

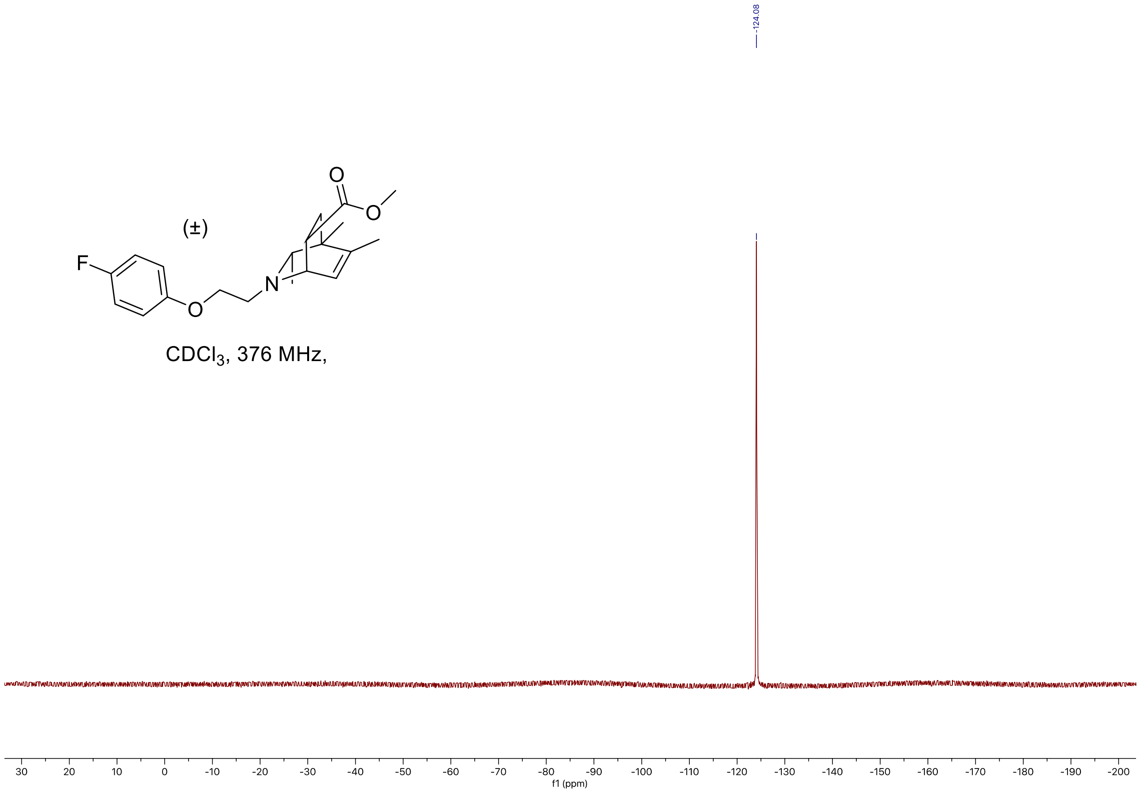


#### C-H COSY

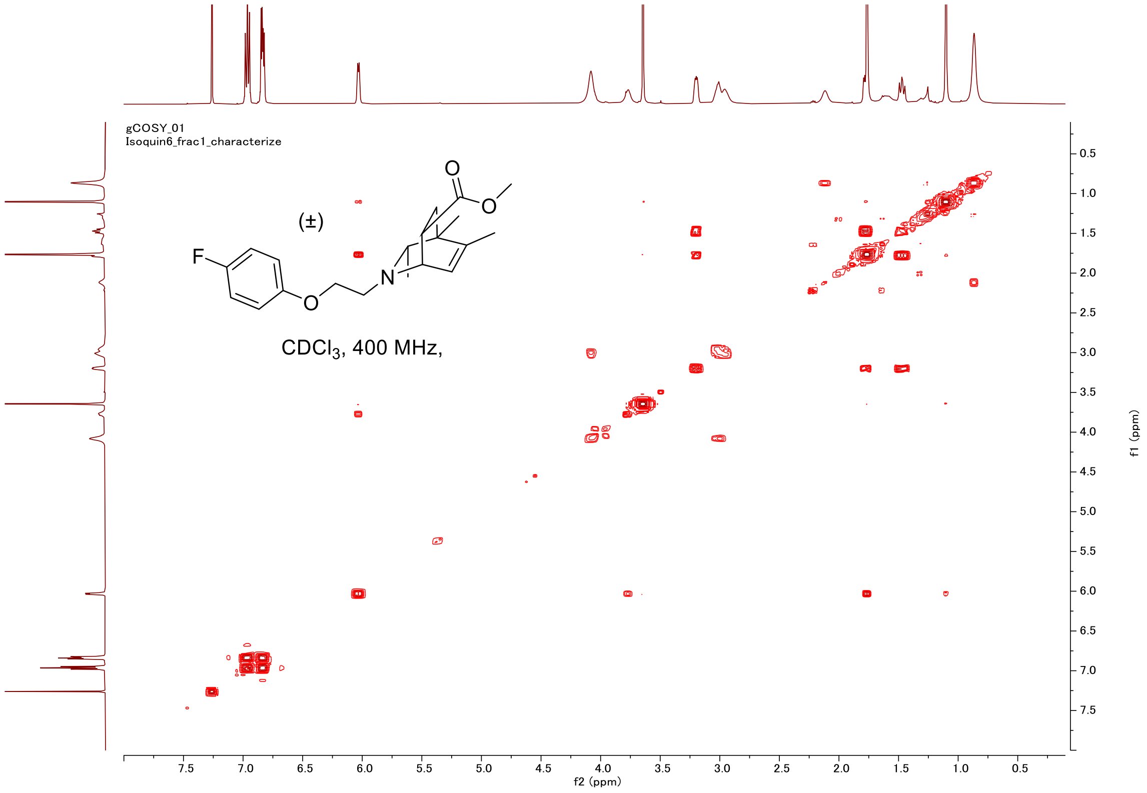

#### HSQC

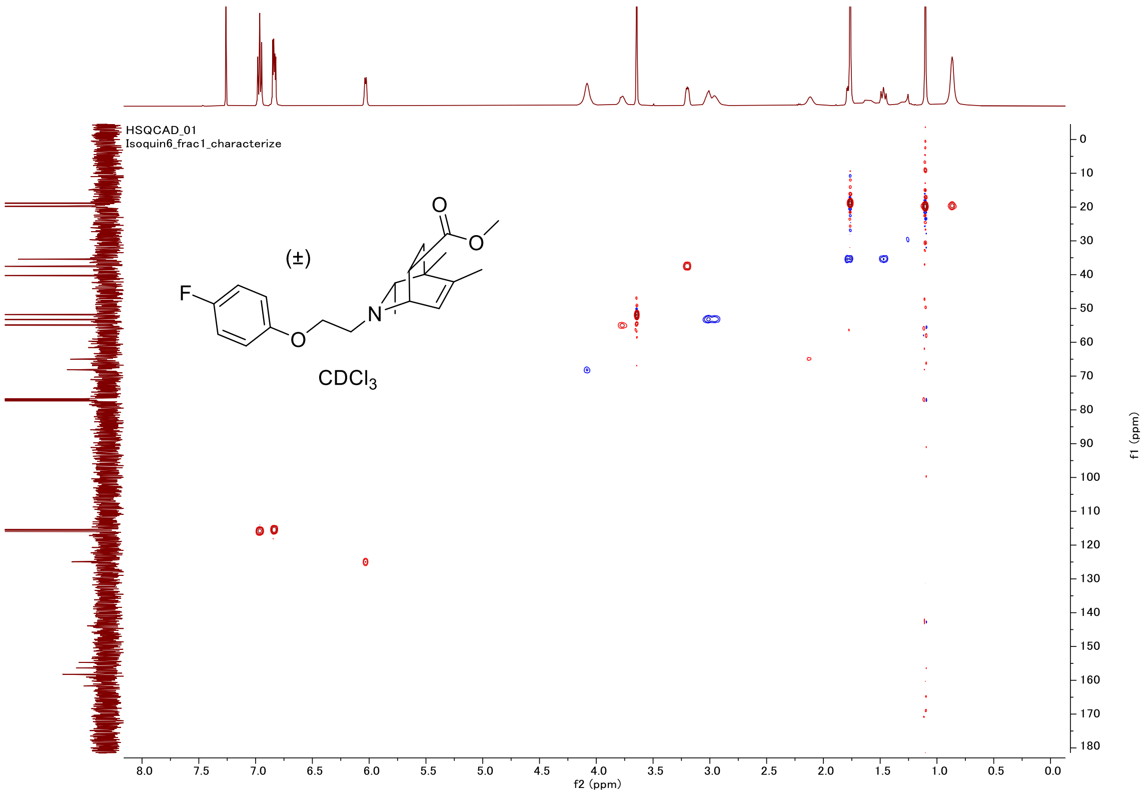

HSQCAD_01
Isoquin6_frac1_characterize
(±)
F
O
N
O
O
CDCl3
f2 (ppm)
f1 (ppm)

#### Proton (#09)

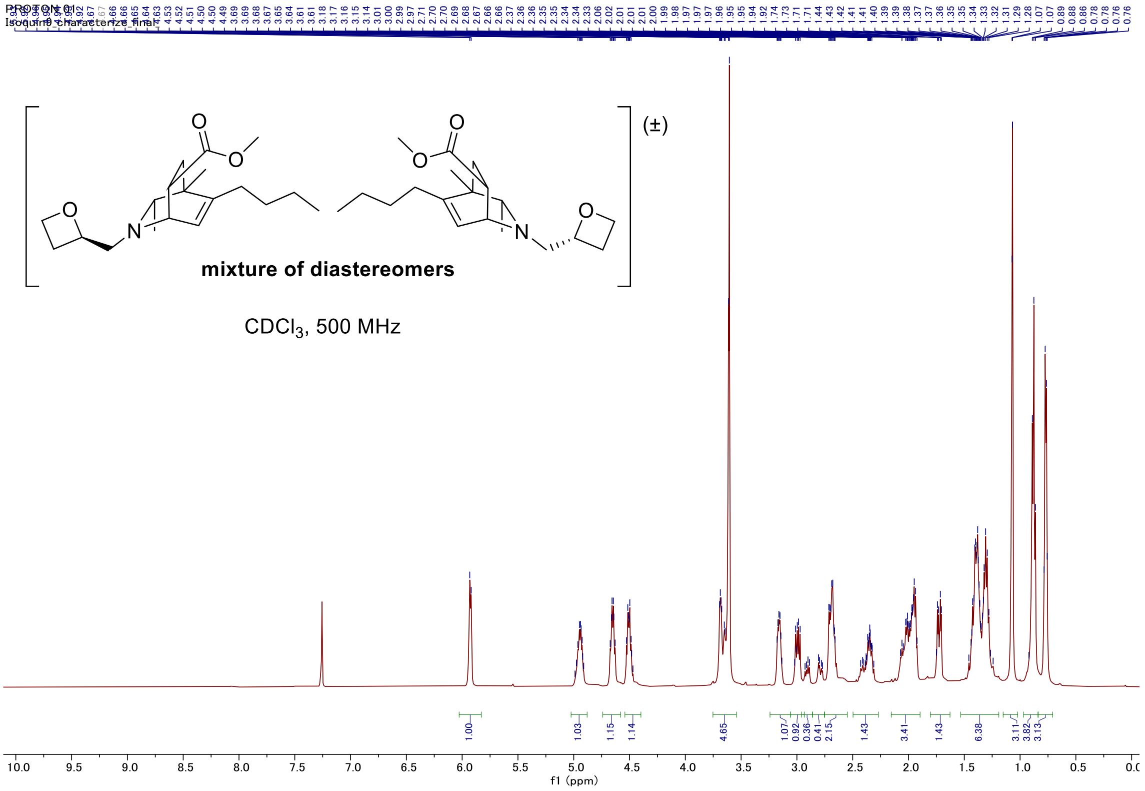


#### Carbon

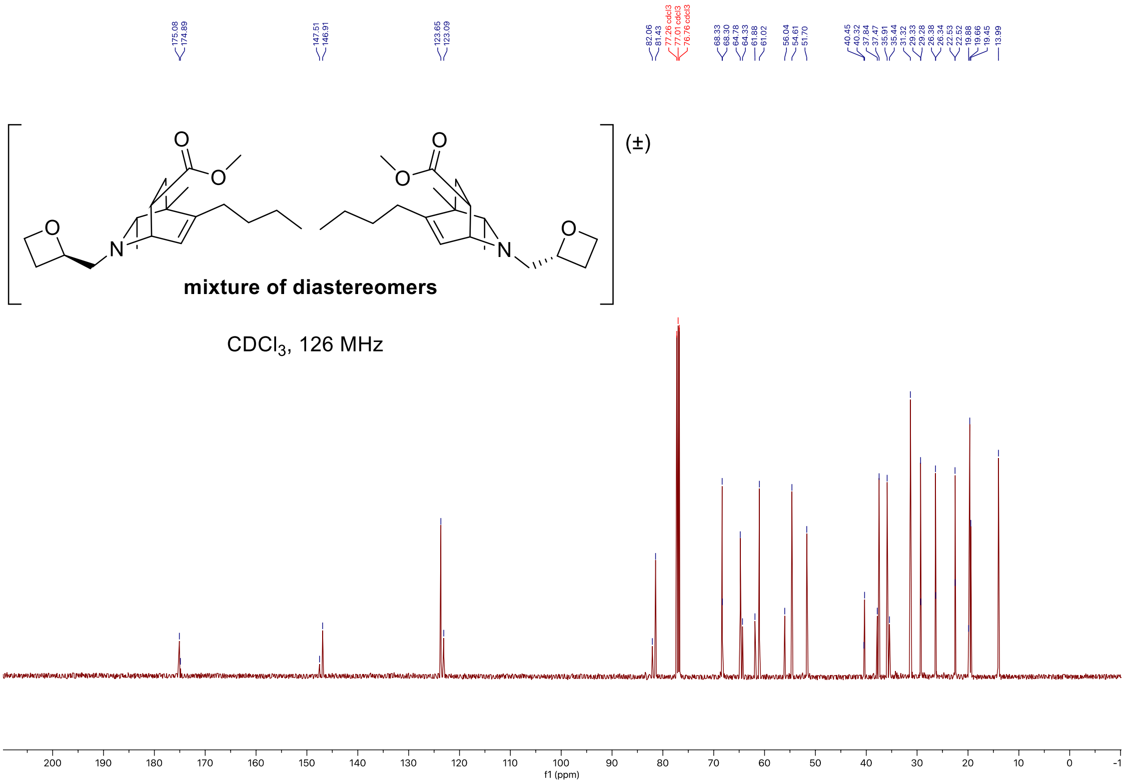

#### C-H COSY

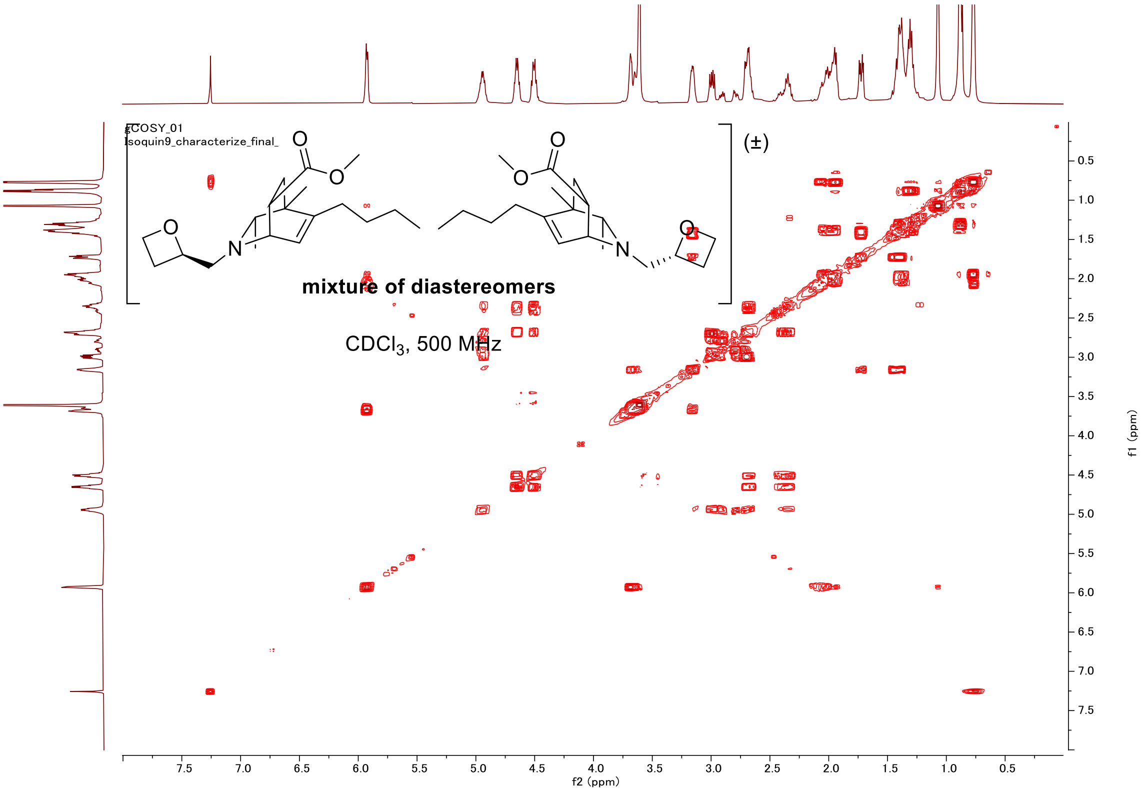
COSY_01
Isoquin9_characterize_final_
(±)
mixture of diastereomers
CDCl3, 500 MHz
f2 (ppm)
f1 (ppm)

#### HSQC

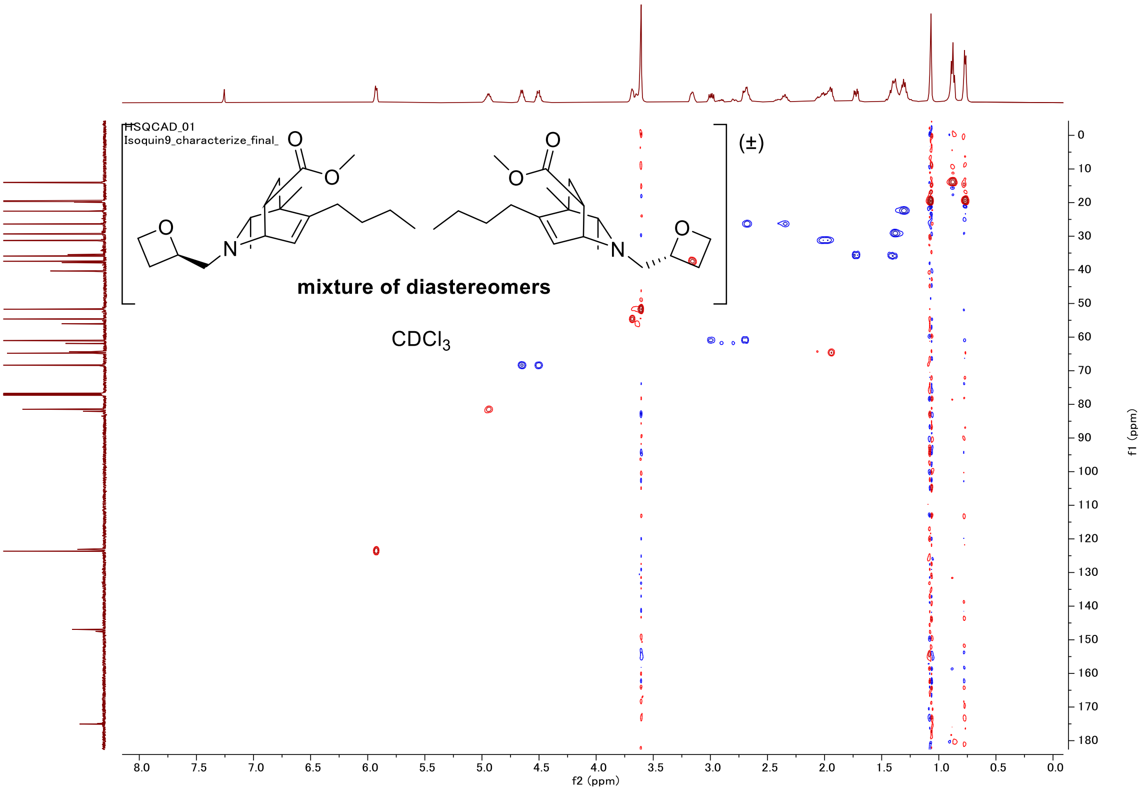
HSQCAD_01
Isoquin9_characterize_final_
(±)
mixture of diastereomers
CDCl3
f2 (ppm)
f1 (ppm)

#### Proton (#11, isomer1)

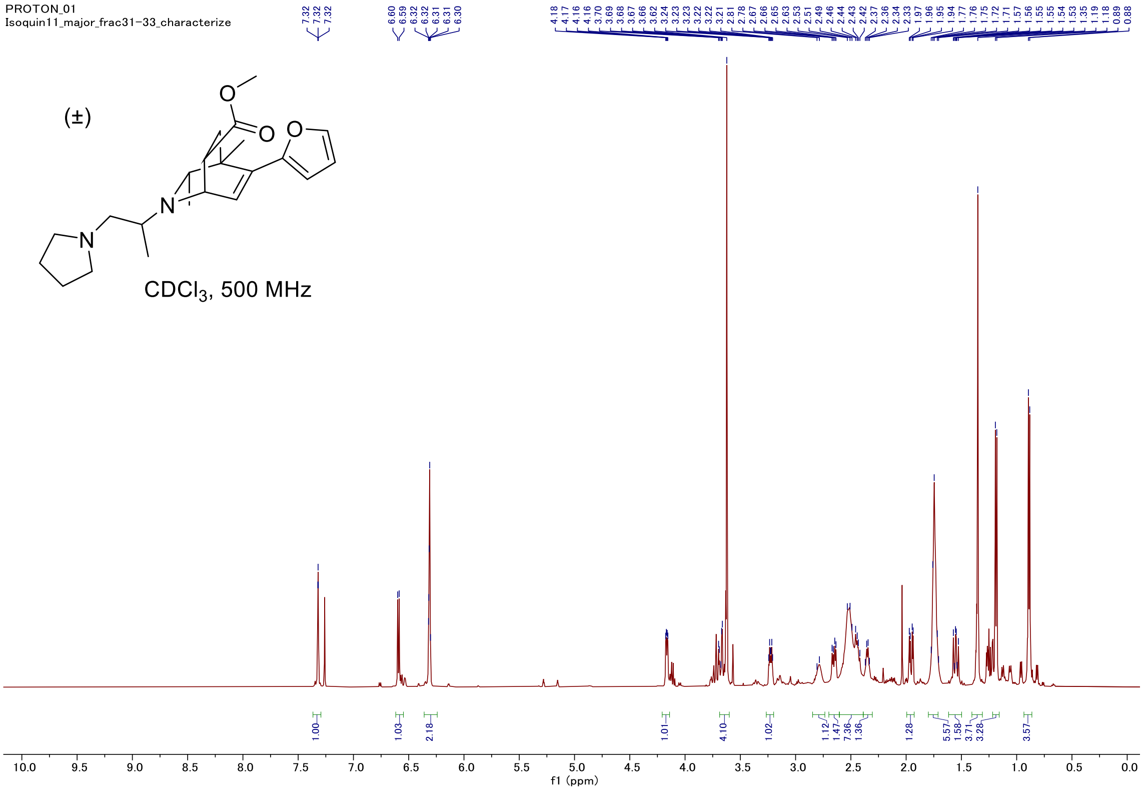


#### Carbon

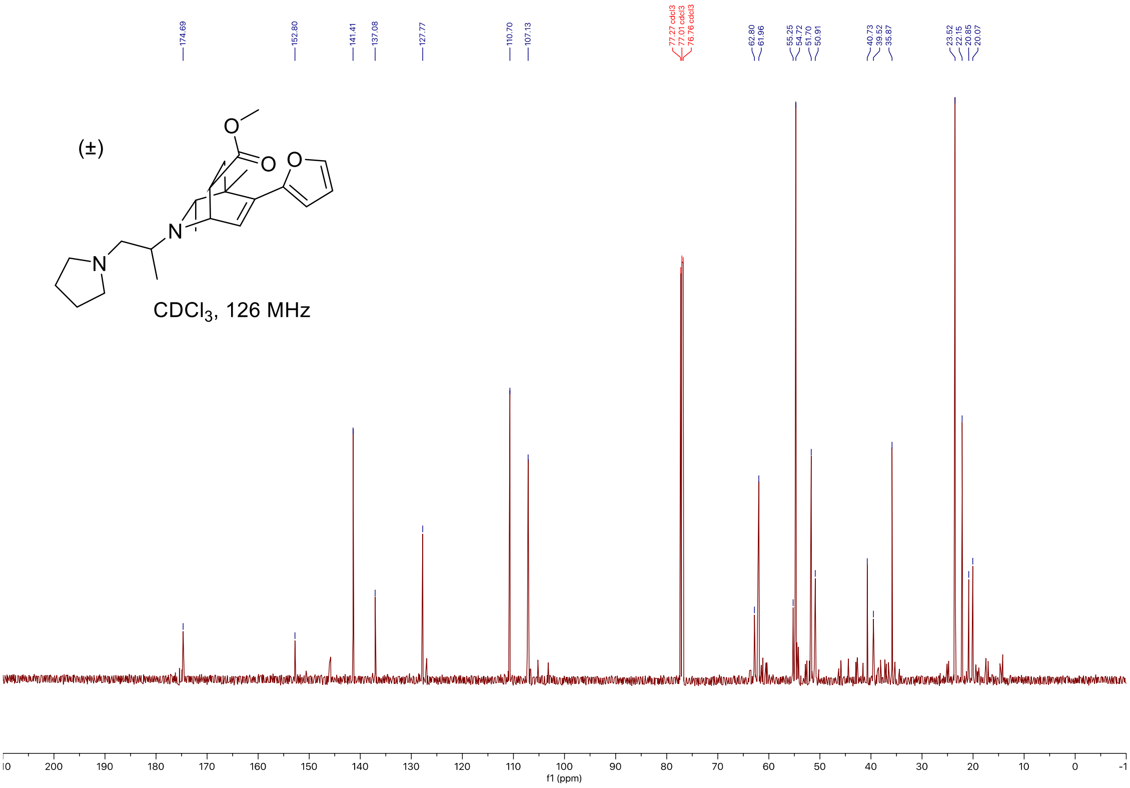

#### C-H COSY

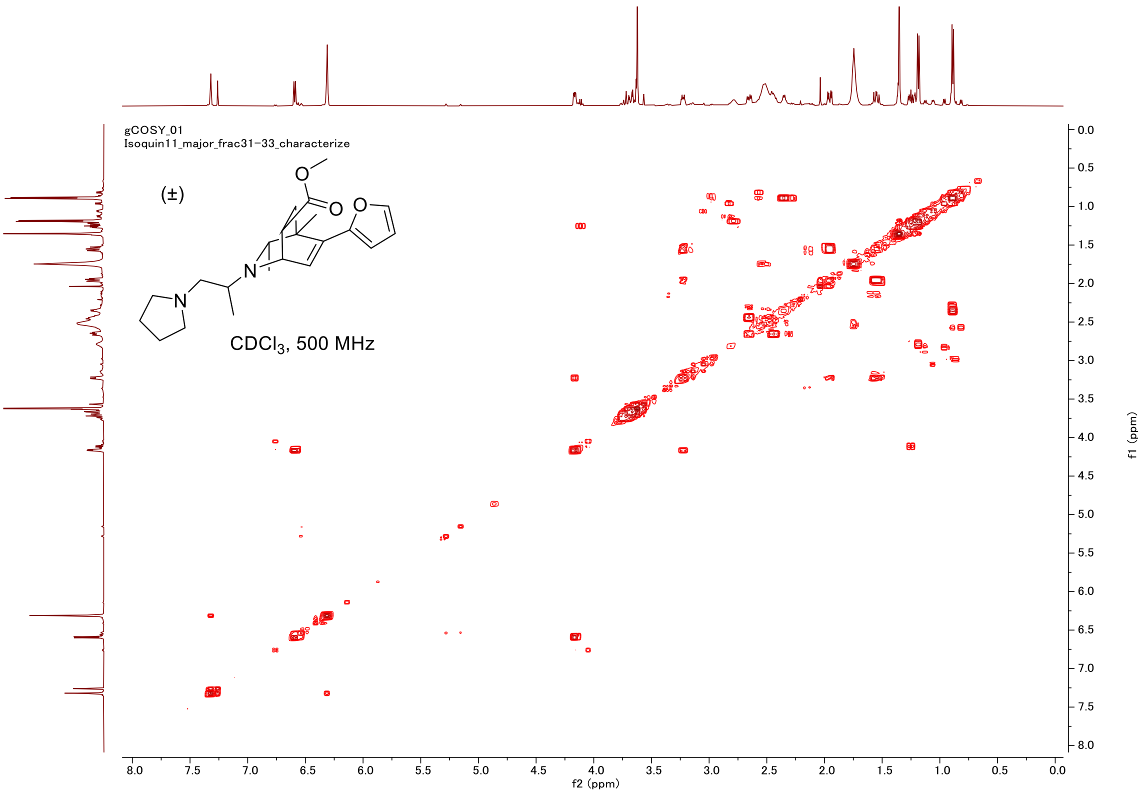

gCOSY_01
Isoquin11_major_frac31-33_characterize
(±)
O
O
O
N
N
CDCl3, 500 MHz
f2 (ppm)
f1 (ppm)


#### HSQC

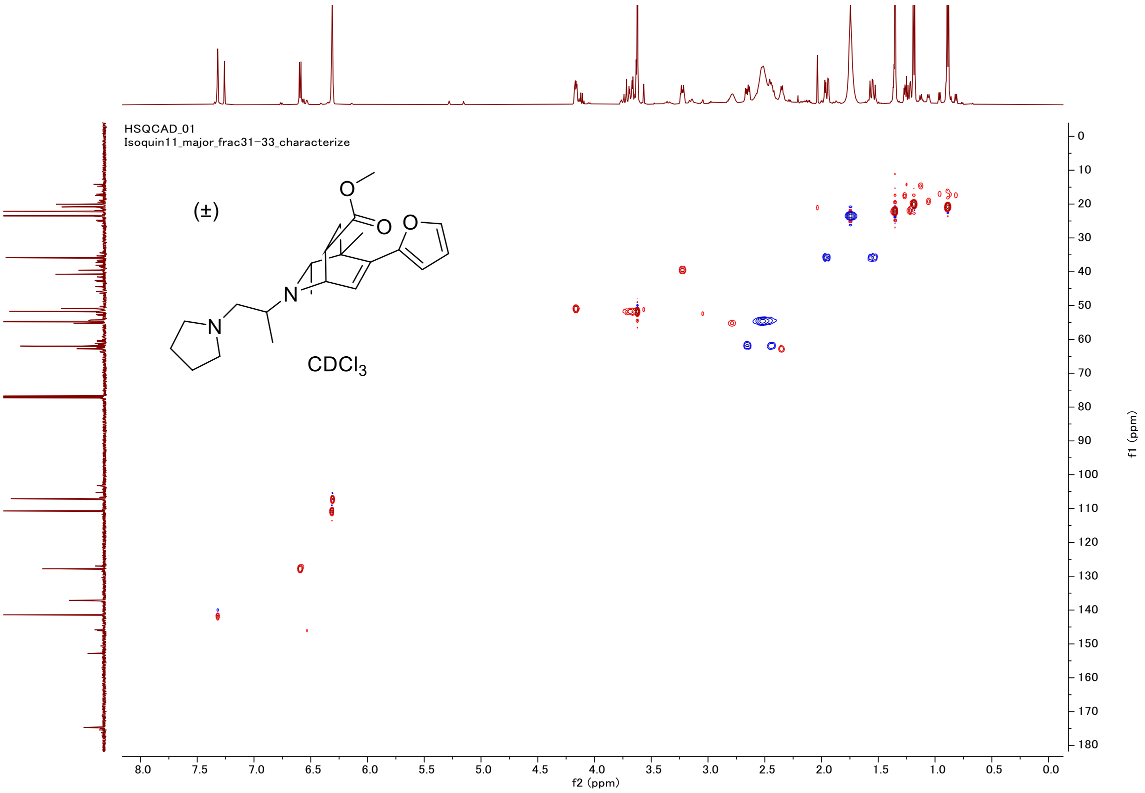

HSQCAD_01
Isoquin11_major_frac31-33_characterize
(±)
O
O
O
N
N
CDCl3
f2 (ppm)
f1 (ppm)

### Proton (#11, isomer2)

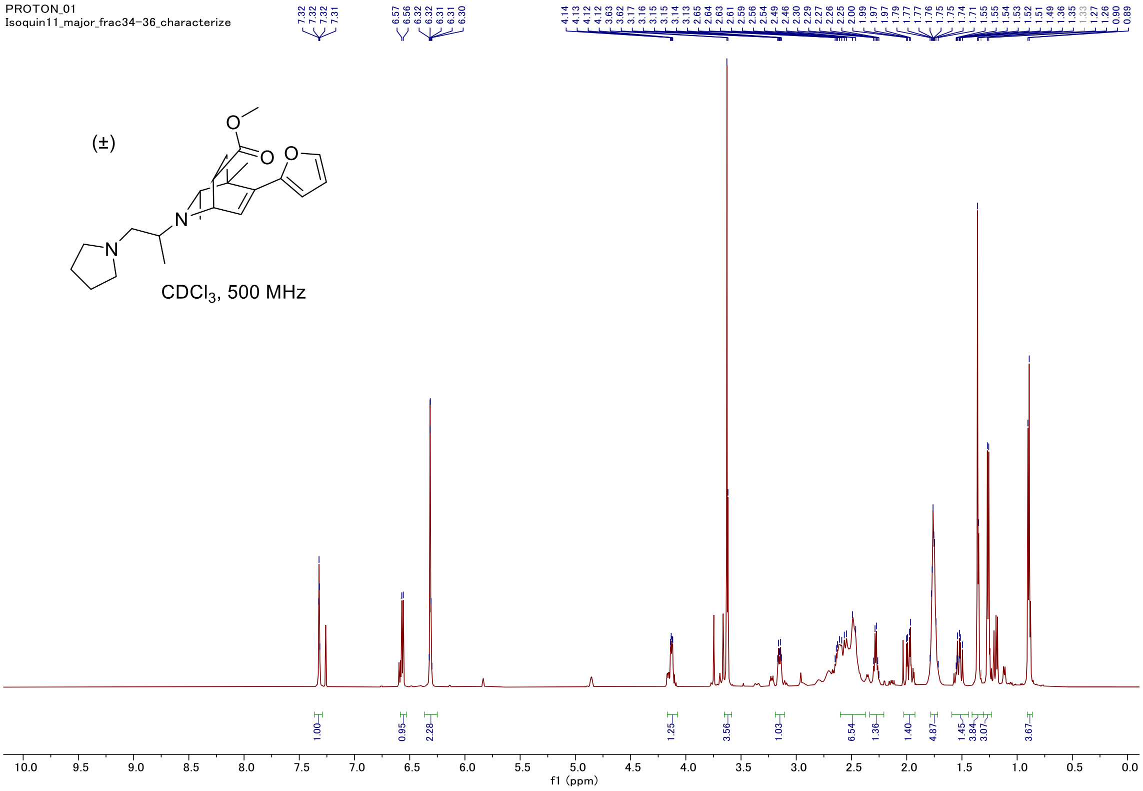


### Carbon

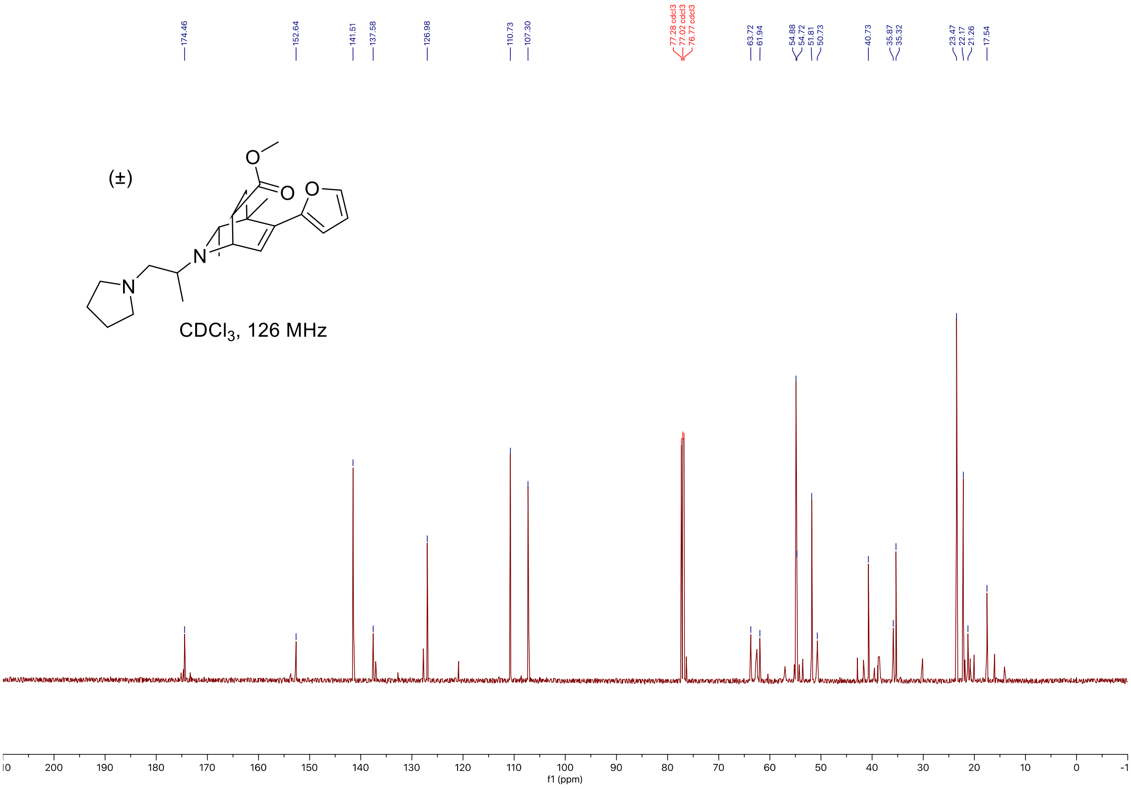

#### C-H COSY

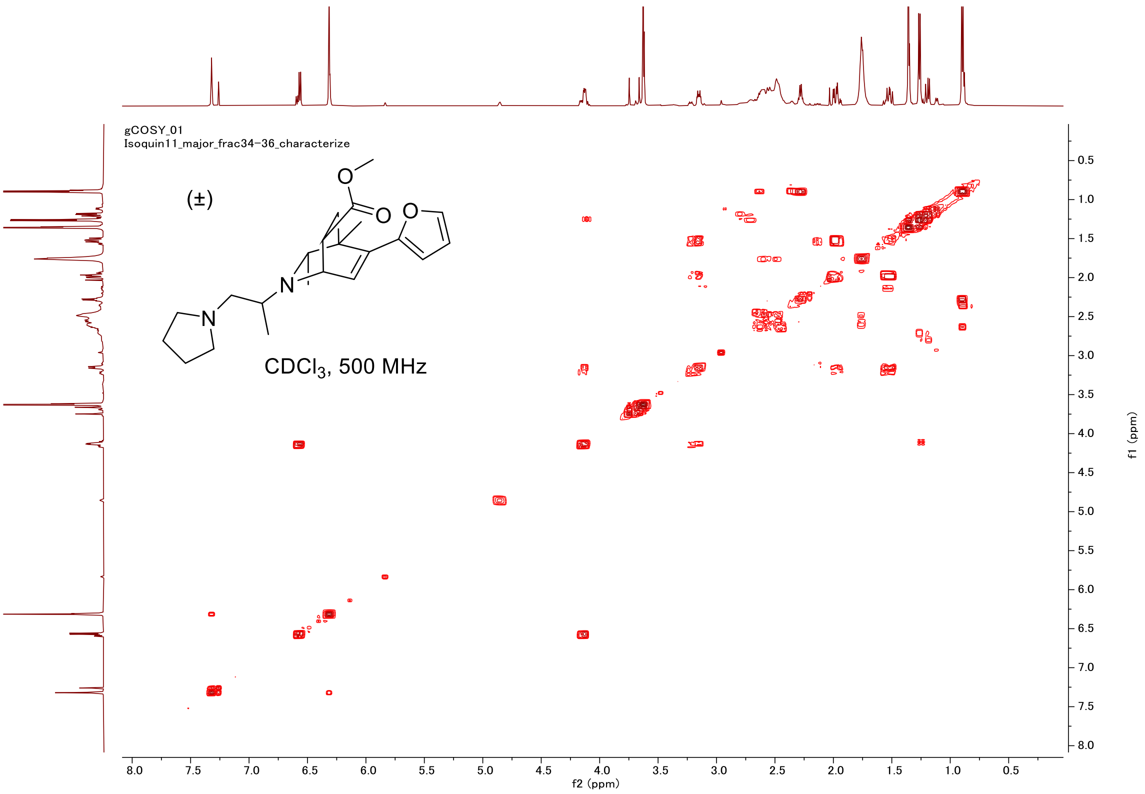
gCOSY_01
Isoquin11_major_frac34-36_characterize
(±)
CDCl3, 500 MHz
f2 (ppm)
f1 (ppm)

#### HSQC

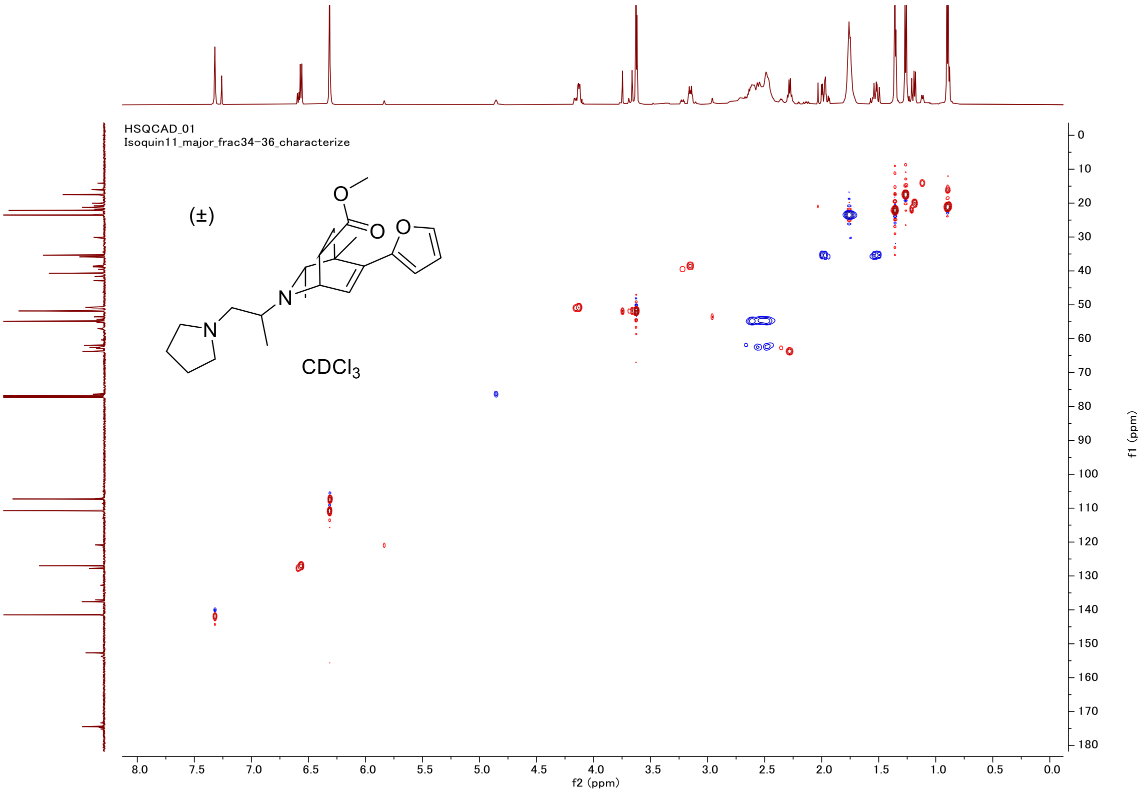
HSQCAD_01
Isoquin11_major_frac34-36_characterize
(±)
CDCl3
f2 (ppm)
f1 (ppm)

#### Proton (#11, isomer 3)

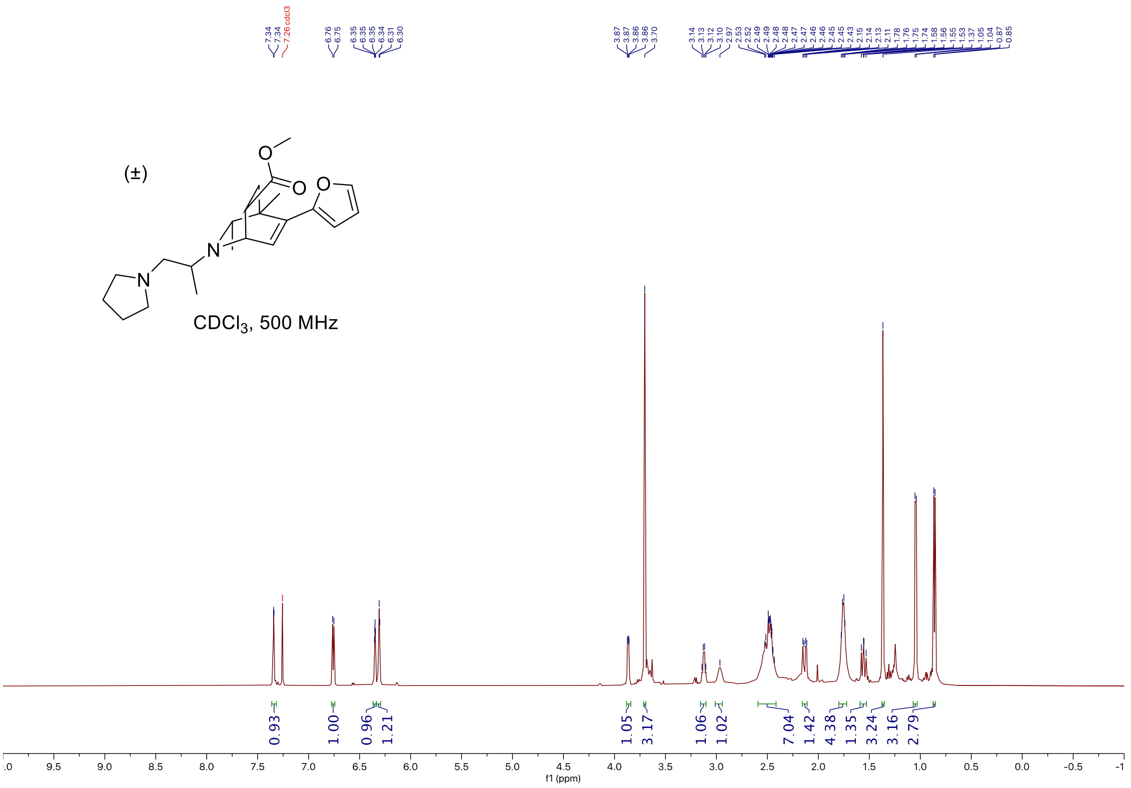


#### Carbon

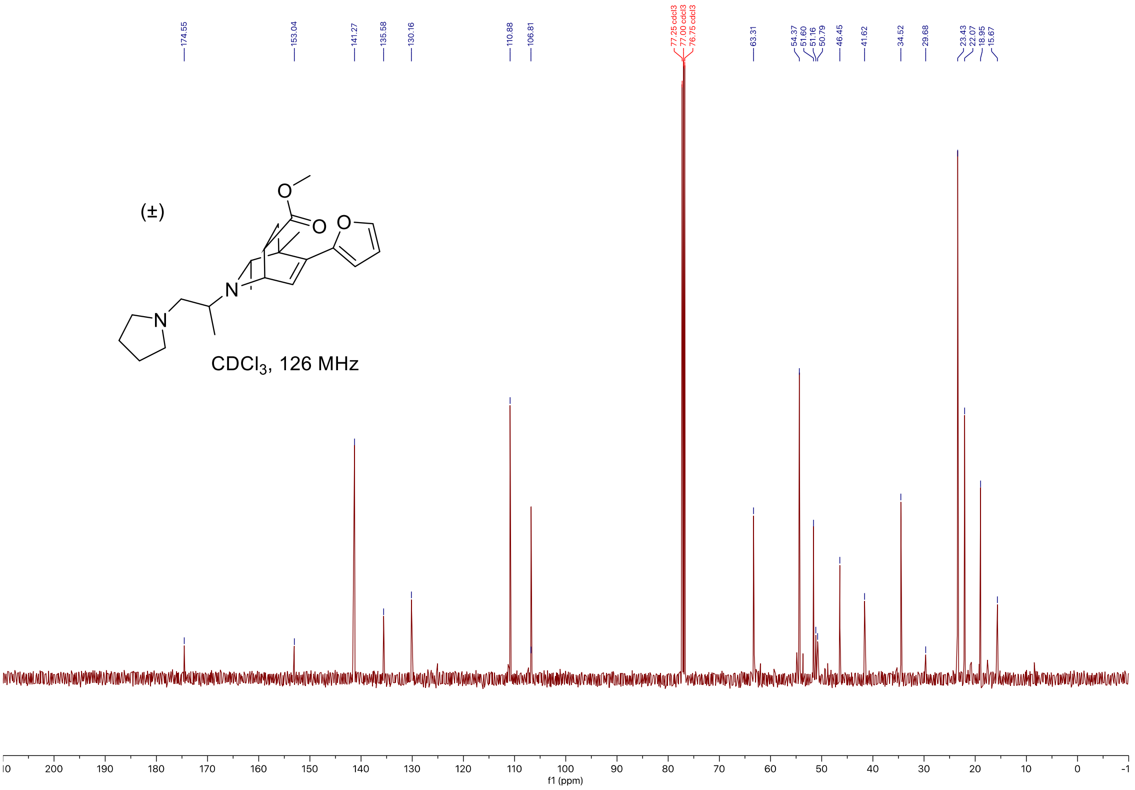

#### C-H COSY

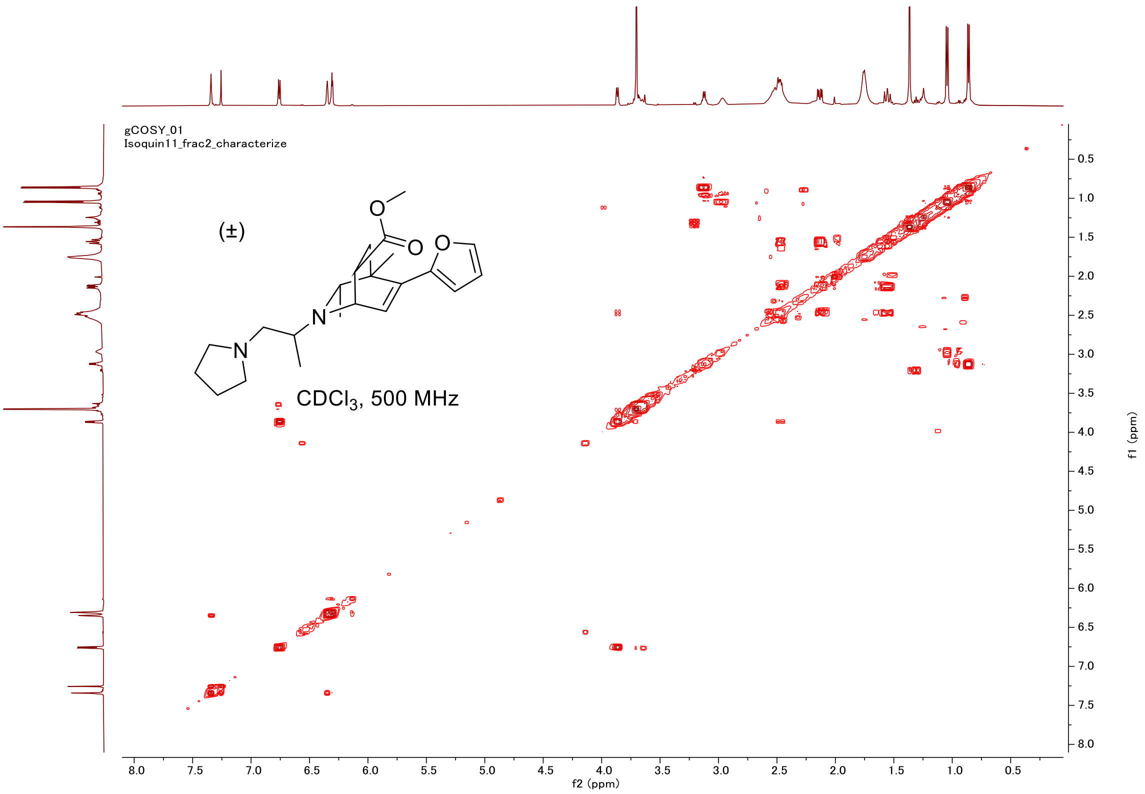

gCOSY_01
Isoquin11_frac2_characterize
(±)
CDCl3, 500 MHz
f2 (ppm)
f1 (ppm)


#### HSQC

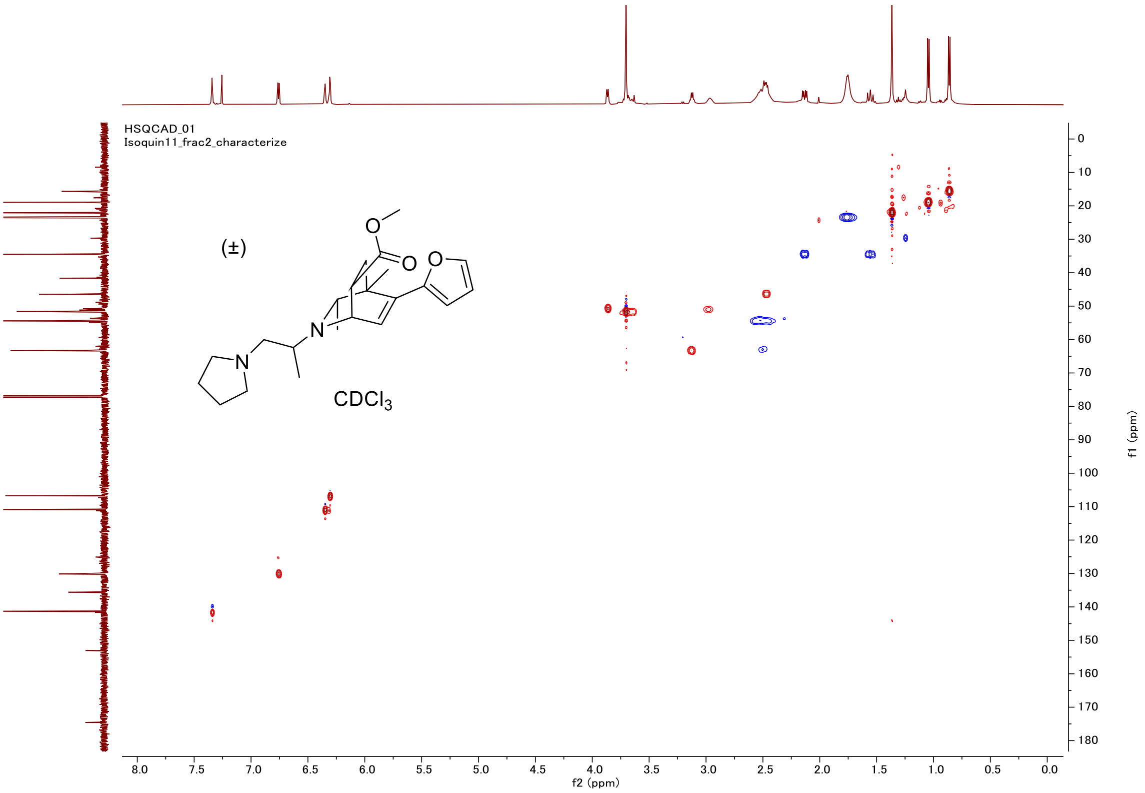

HSQCAD_01
Isoquin11_frac2_characterize
(±)
CDCl3
f2 (ppm)
f1 (ppm)

#### Proton (#12)

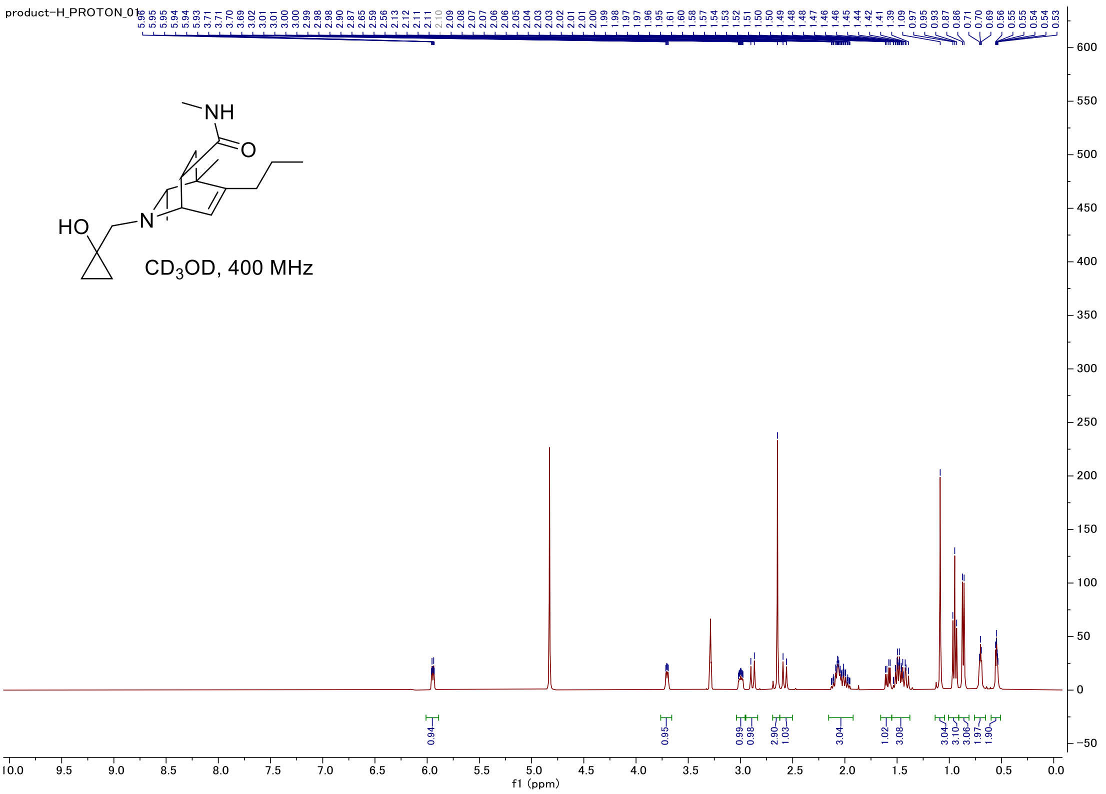


#### Carbon

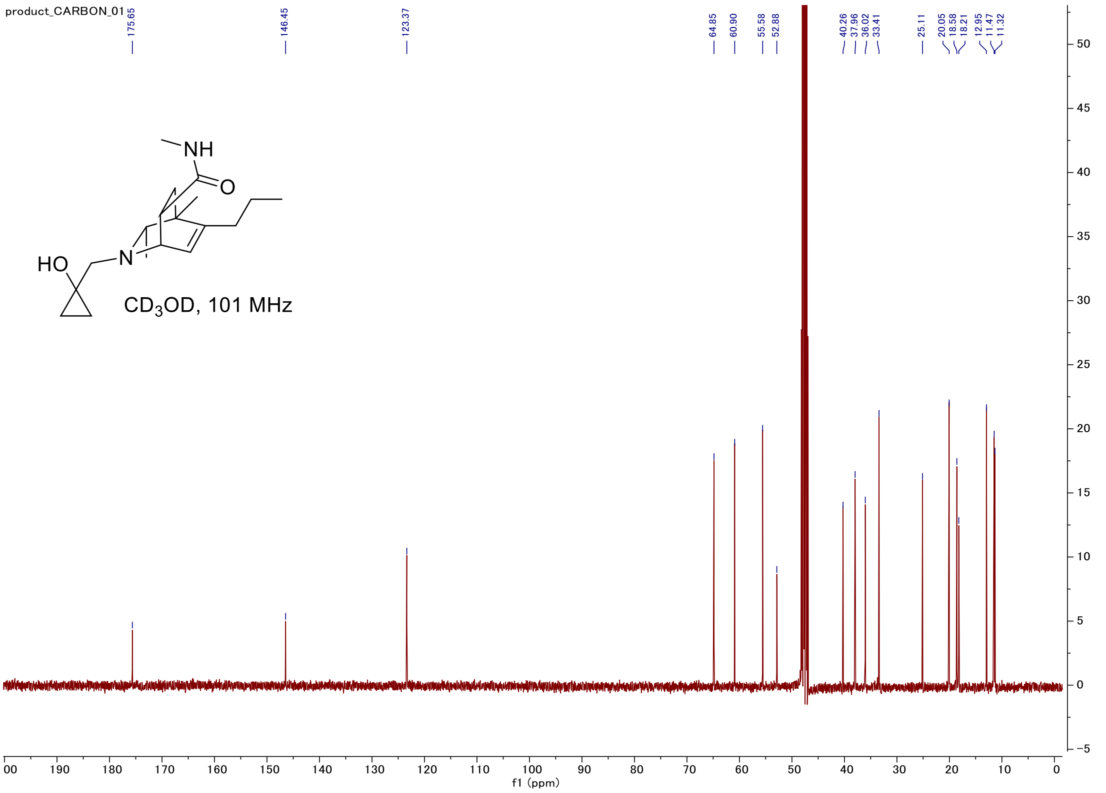

#### C-H COSY

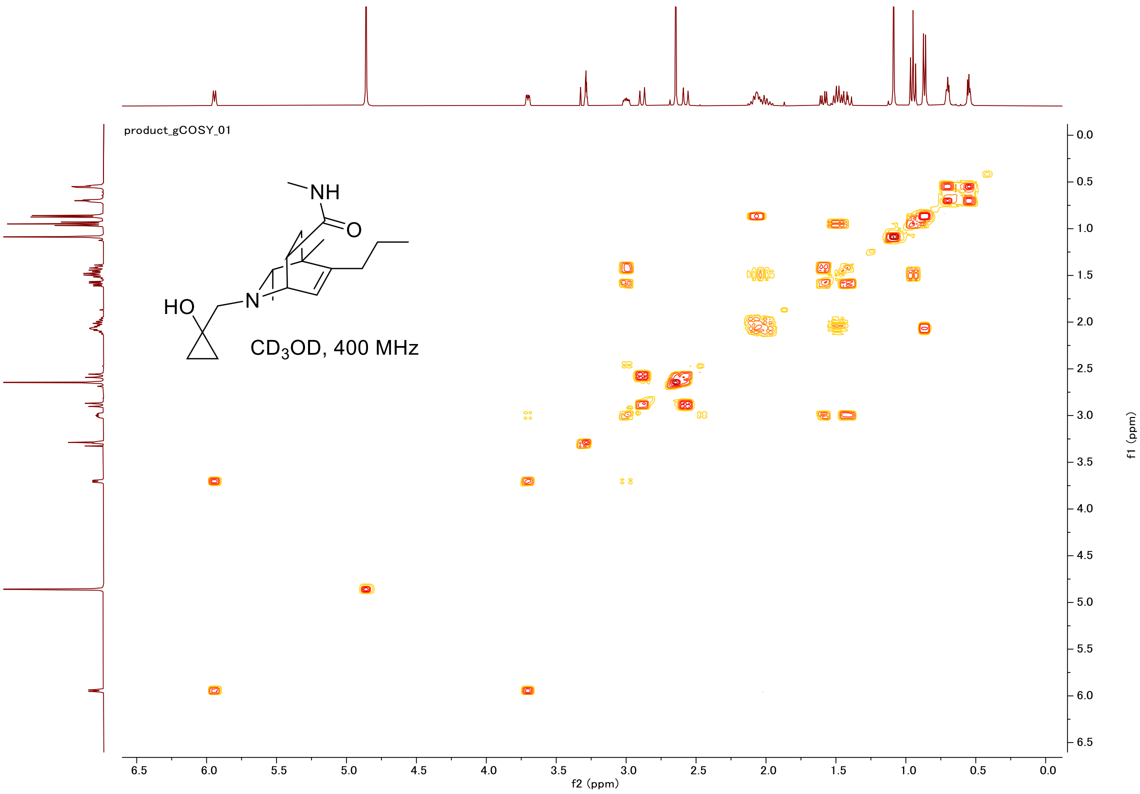
product_gCOSY_01
NH
O
HO
N
CD3OD, 400 MHz
f2 (ppm)
f1 (ppm)

#### HSQC

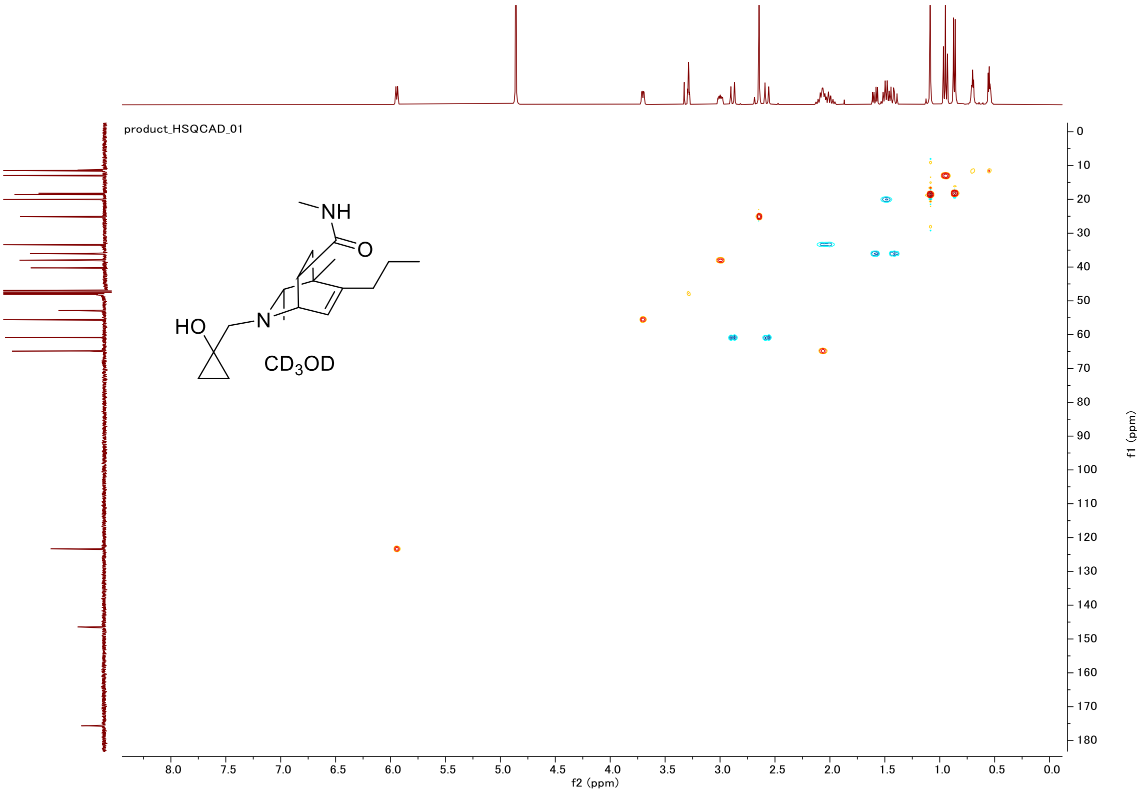
product_HSQCAD_01
NH
O
HO
N
CD3OD
f2 (ppm)
f1 (ppm)

#### Proton (#14)

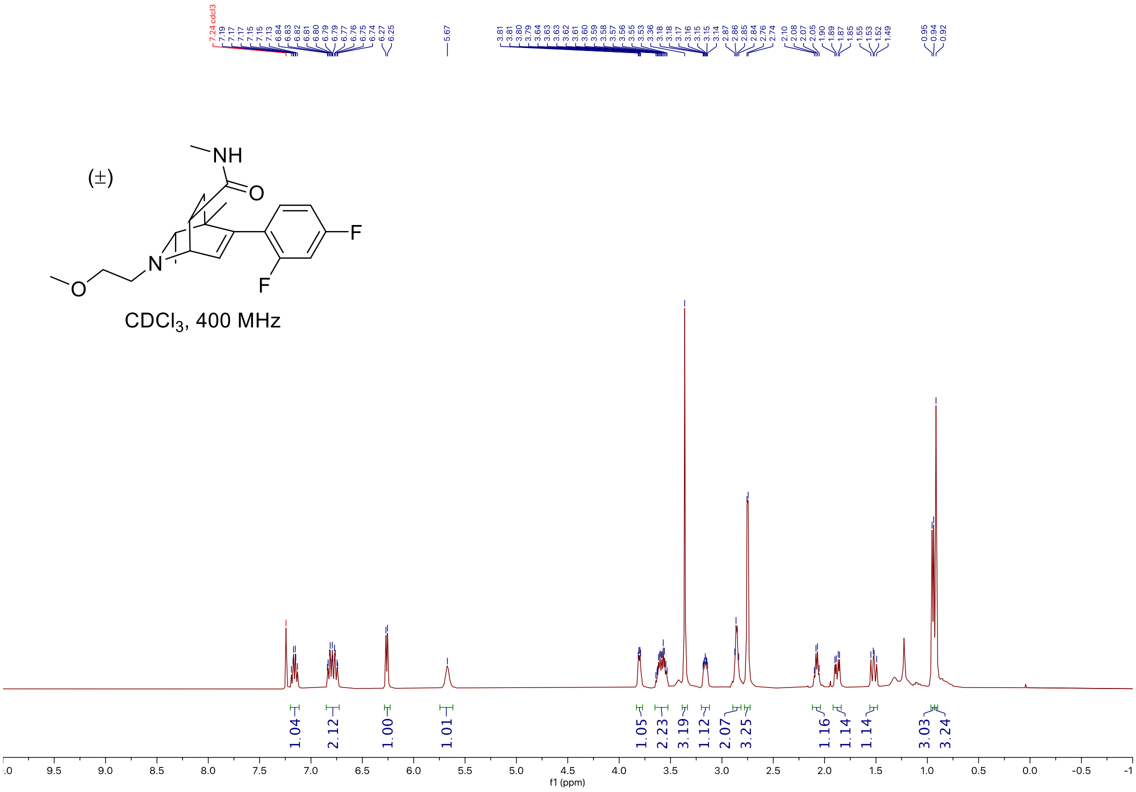


#### Carbon

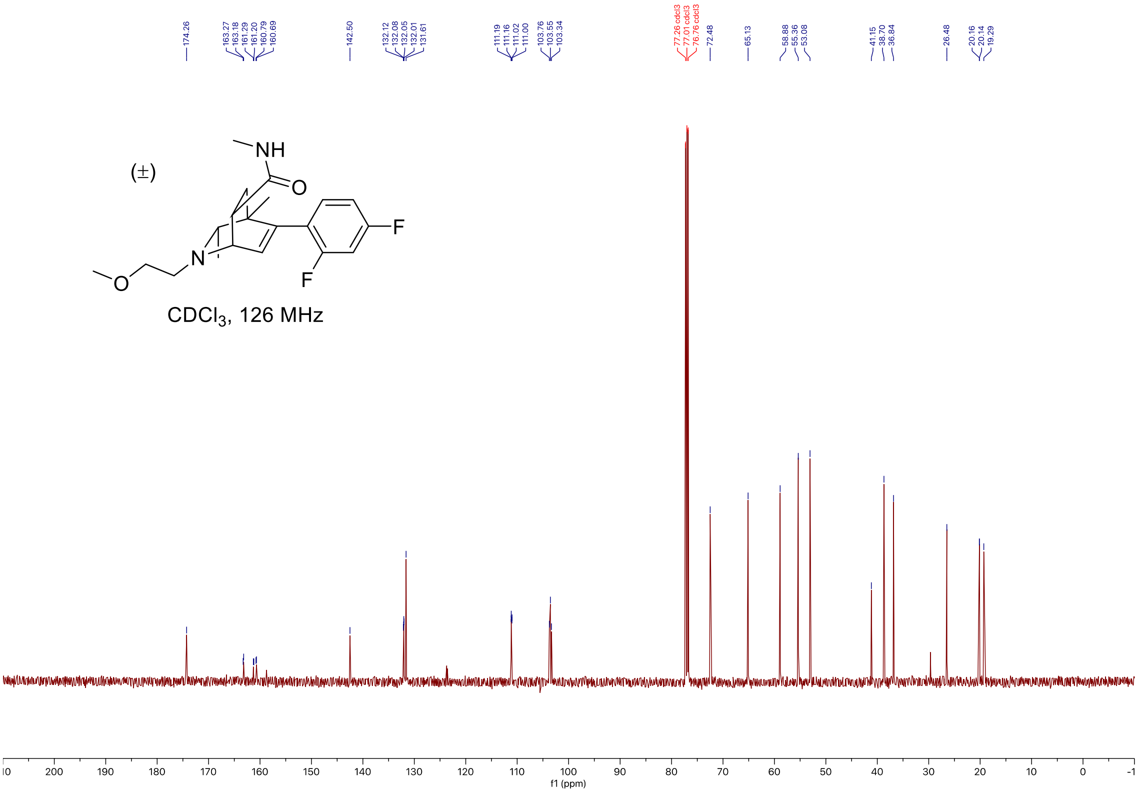

### Fluorine

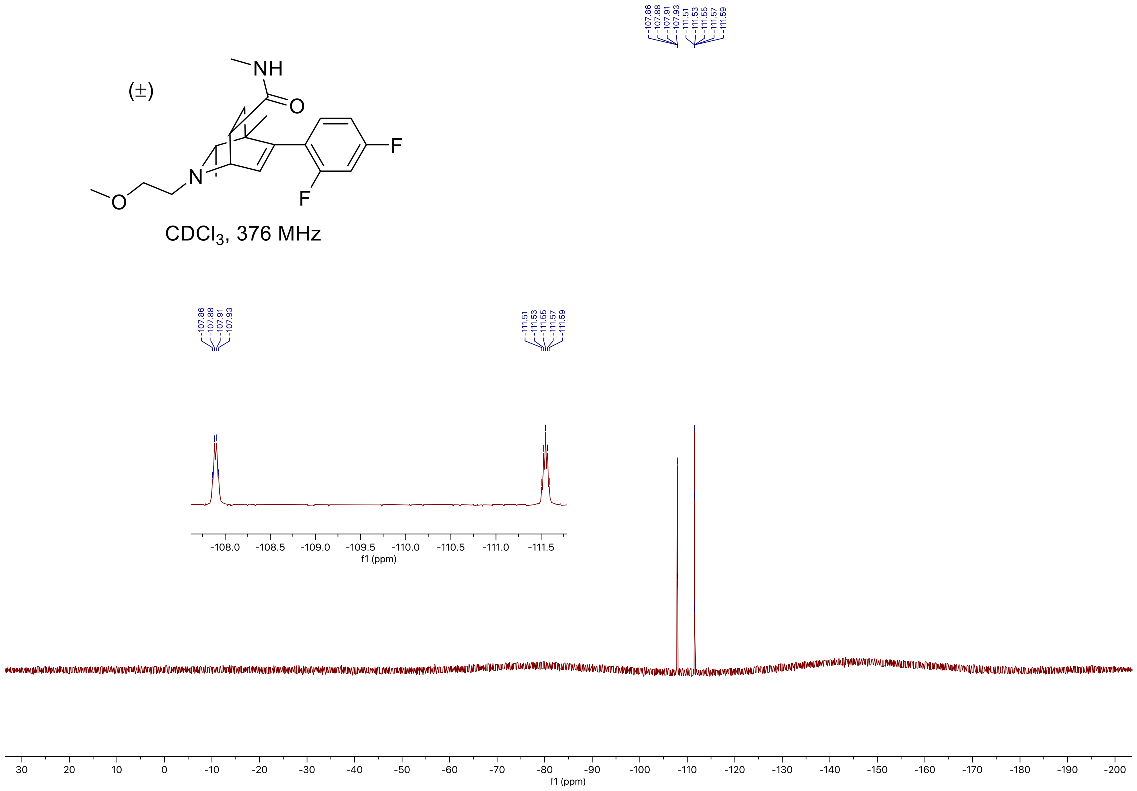


### C-H COSY

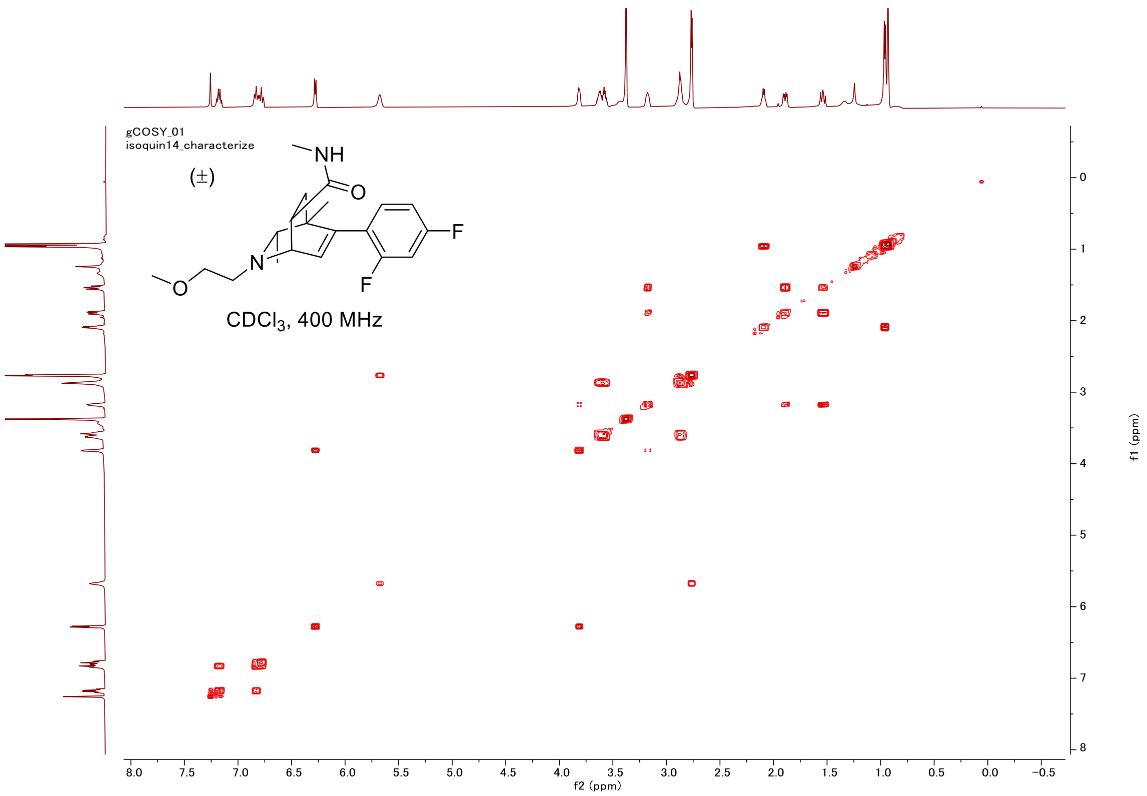

#### HSQC

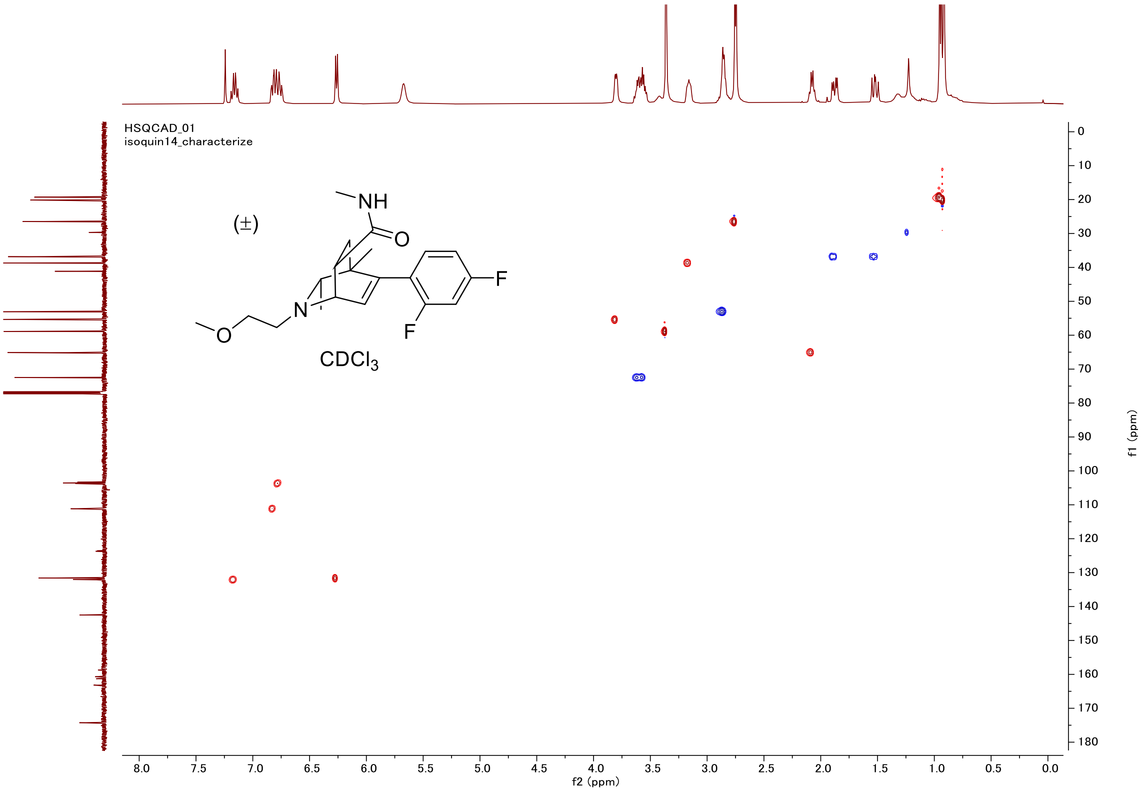

HSQCAD_01
isoquin14_characterize
(±)
NH
O
F
N
O
F
CDCl3
f2 (ppm)
f1 (ppm)

#### Proton (#17)

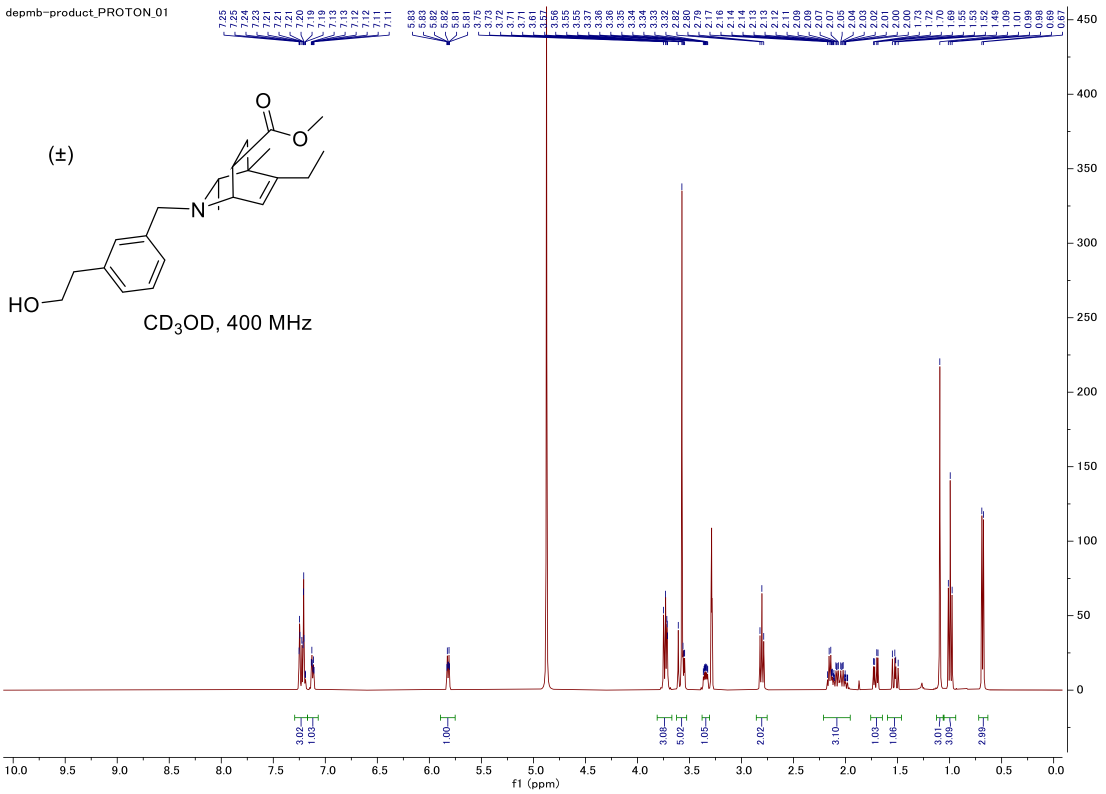


#### Carbon

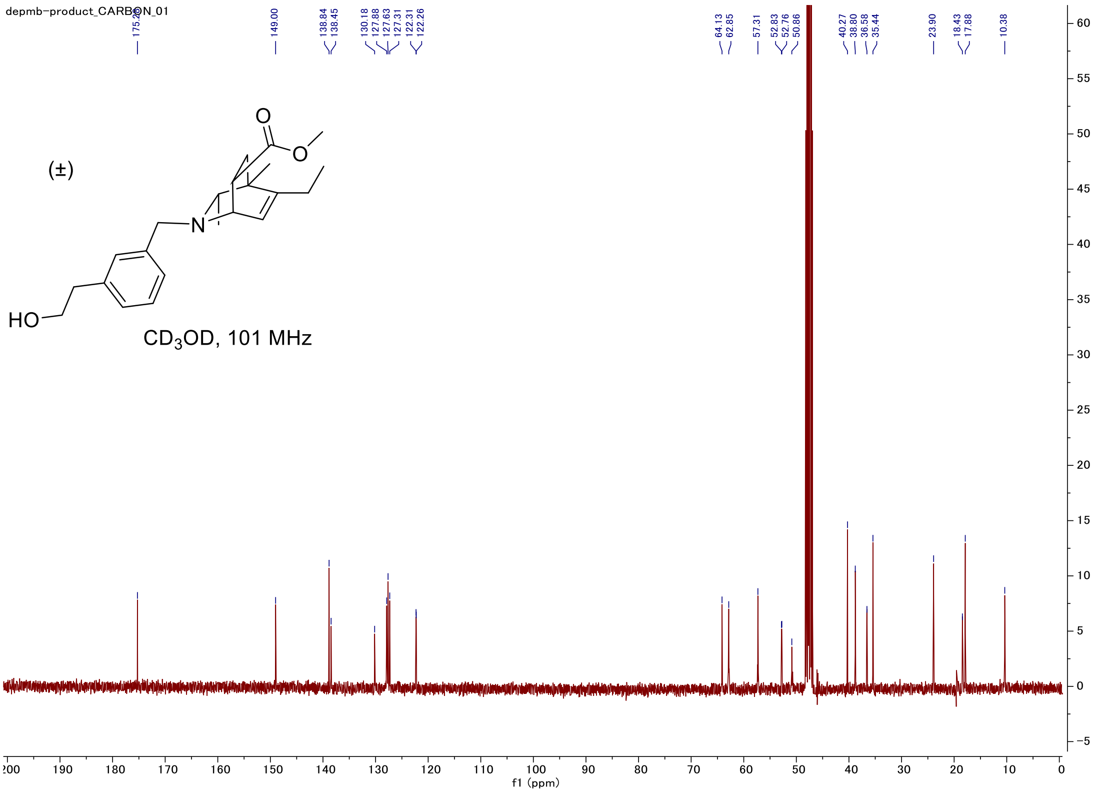

### C-H COSY

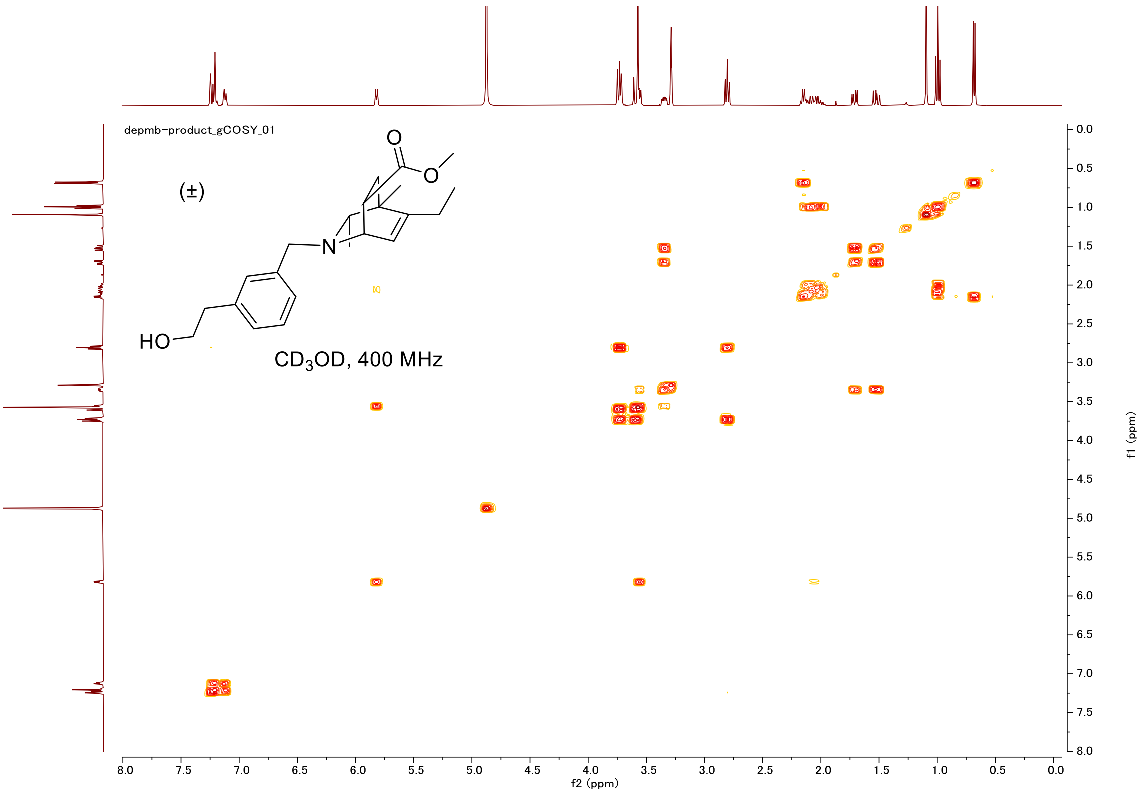
depmb-product_gCOSY_01
(±)
HO
N
O
CD3OD, 400 MHz
f2 (ppm)
f1 (ppm)

### HSQC

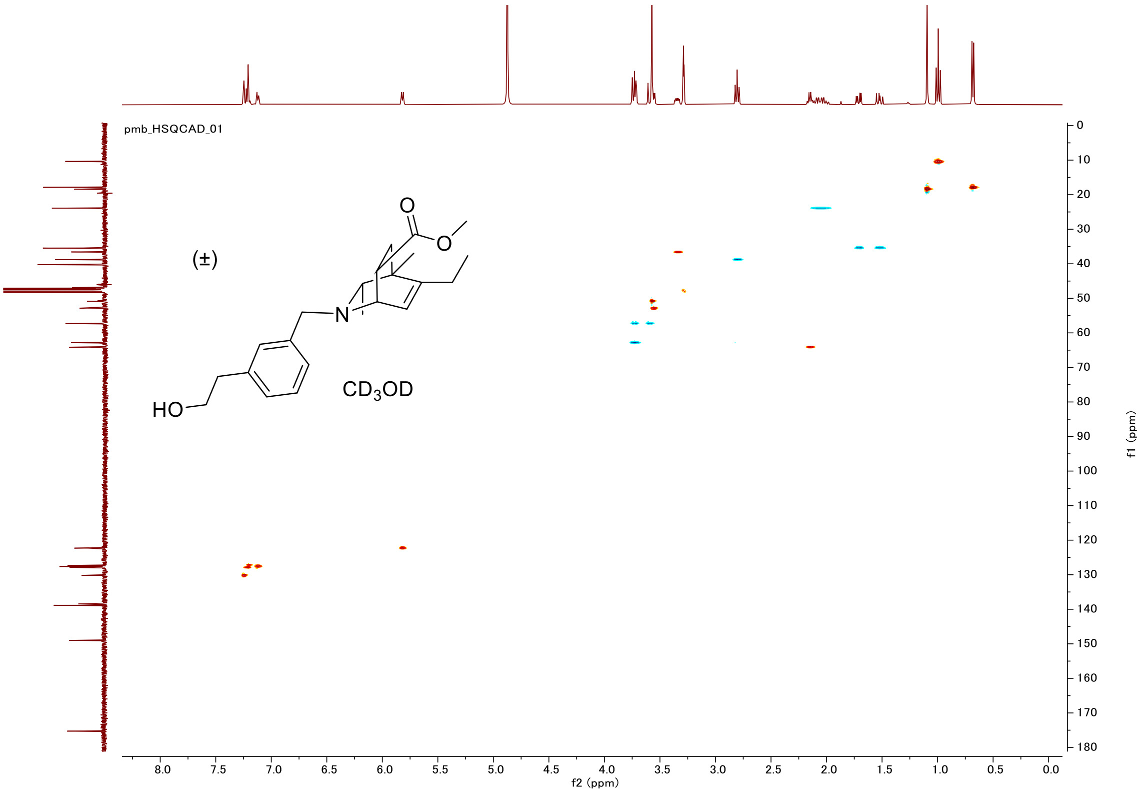
pmb_HSQCAD_01
(±)
HO
N
O
CD3OD
f2 (ppm)
f1 (ppm)

#### Proton (#19-major diastereomer)

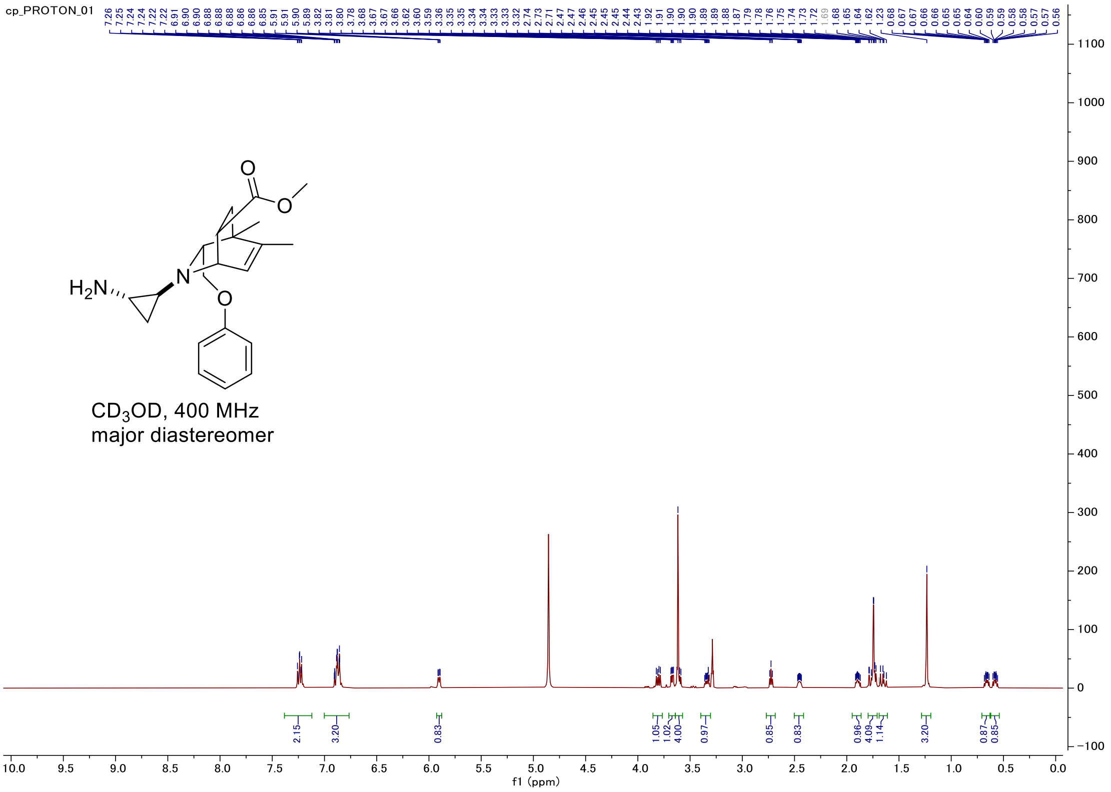


#### Carbon

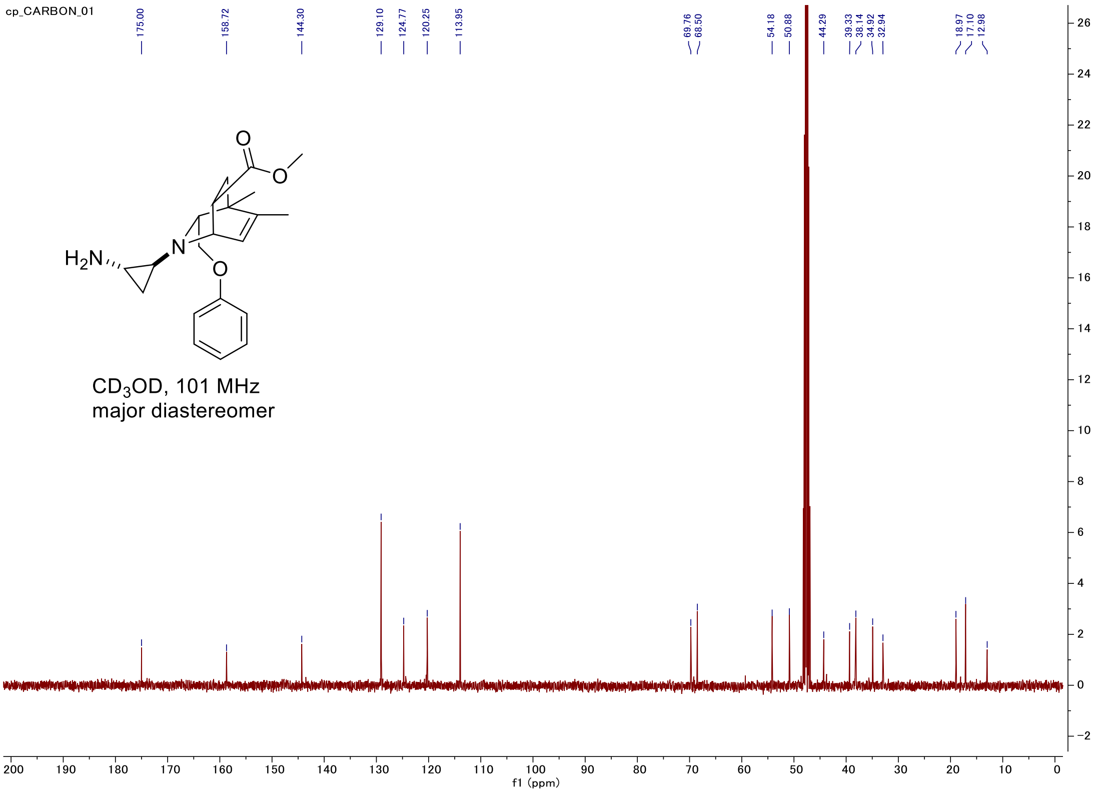

#### C-H COSY

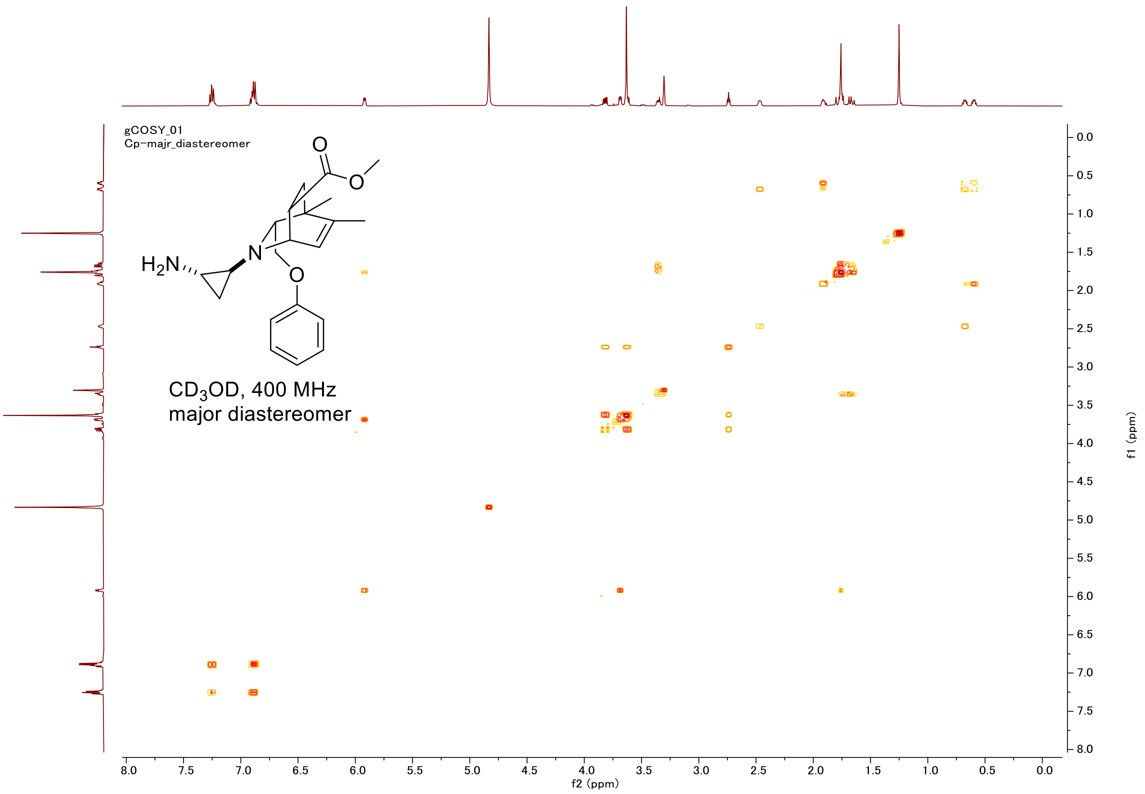

gCOSY_01
Cp−majr_diastereomer
H2N
N
O
O
O
CD3OD, 400 MHz
major diastereomer
f2 (ppm)
f1 (ppm)


#### HSQC

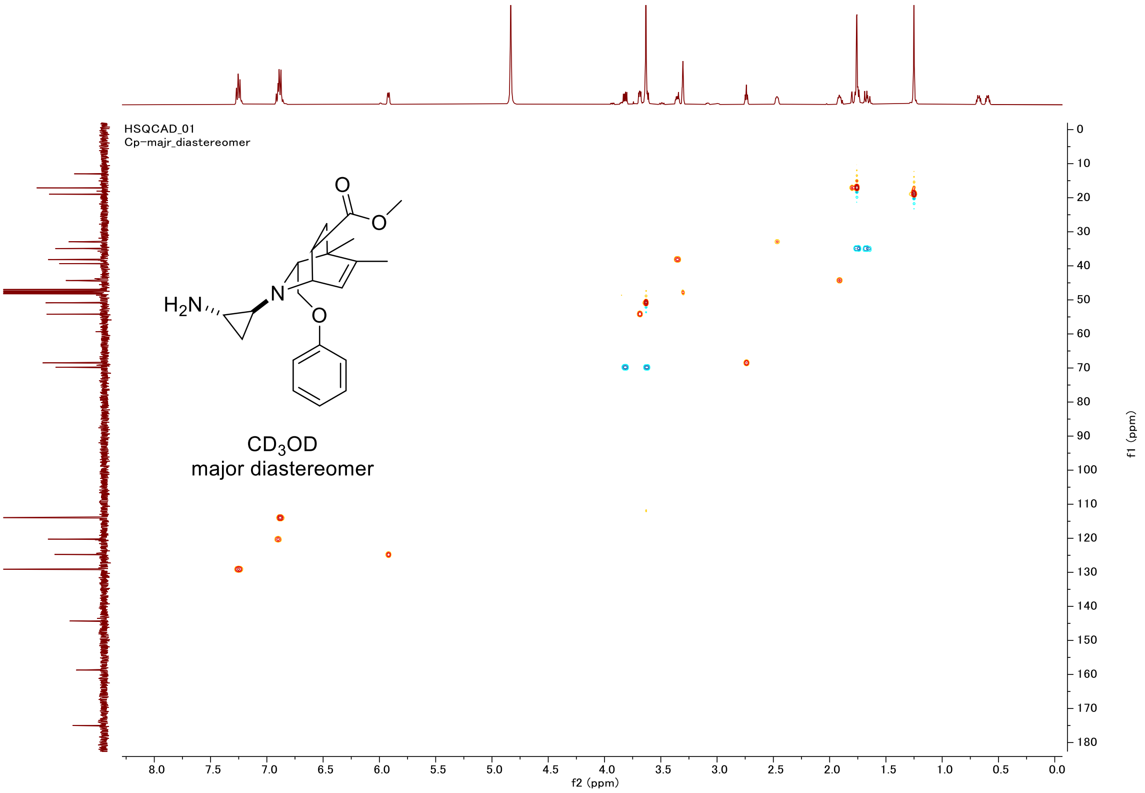

HSQCAD_01
Cp−majr_diastereomer
H2N
N
O
O
O
CD3OD
major diastereomer
f2 (ppm)
f1 (ppm)

#### Proton (#19-minor diastereomer)

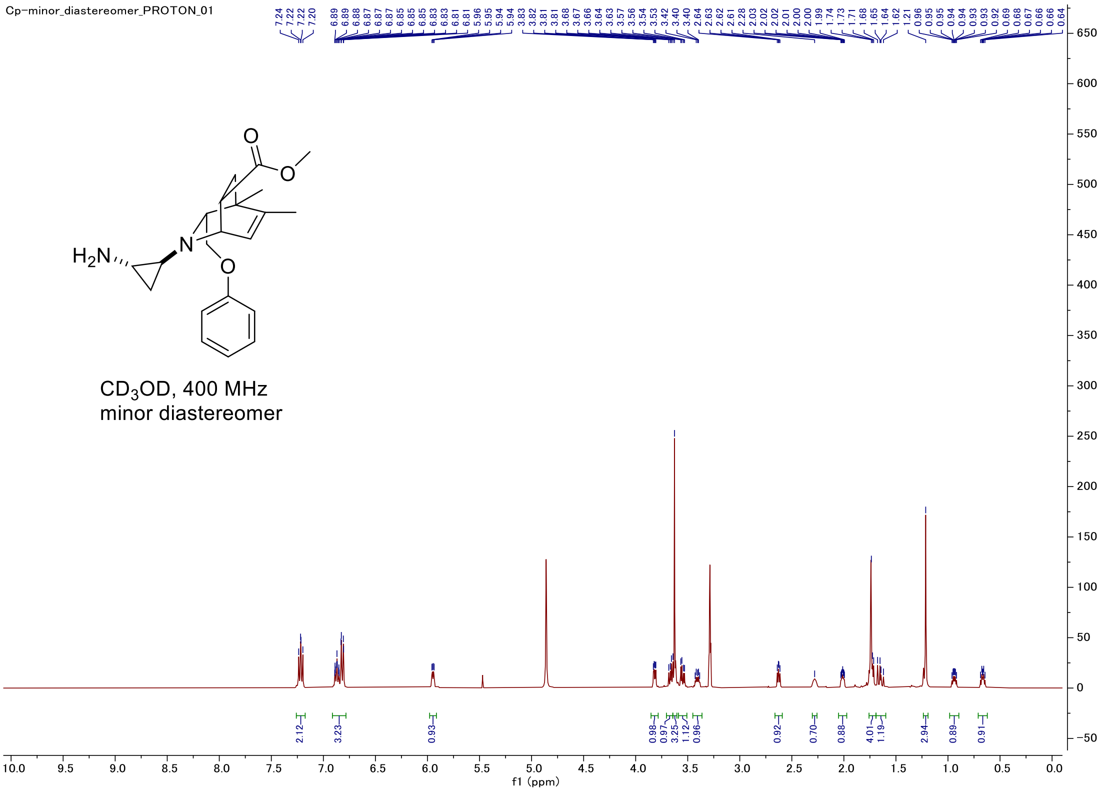


#### Carbon

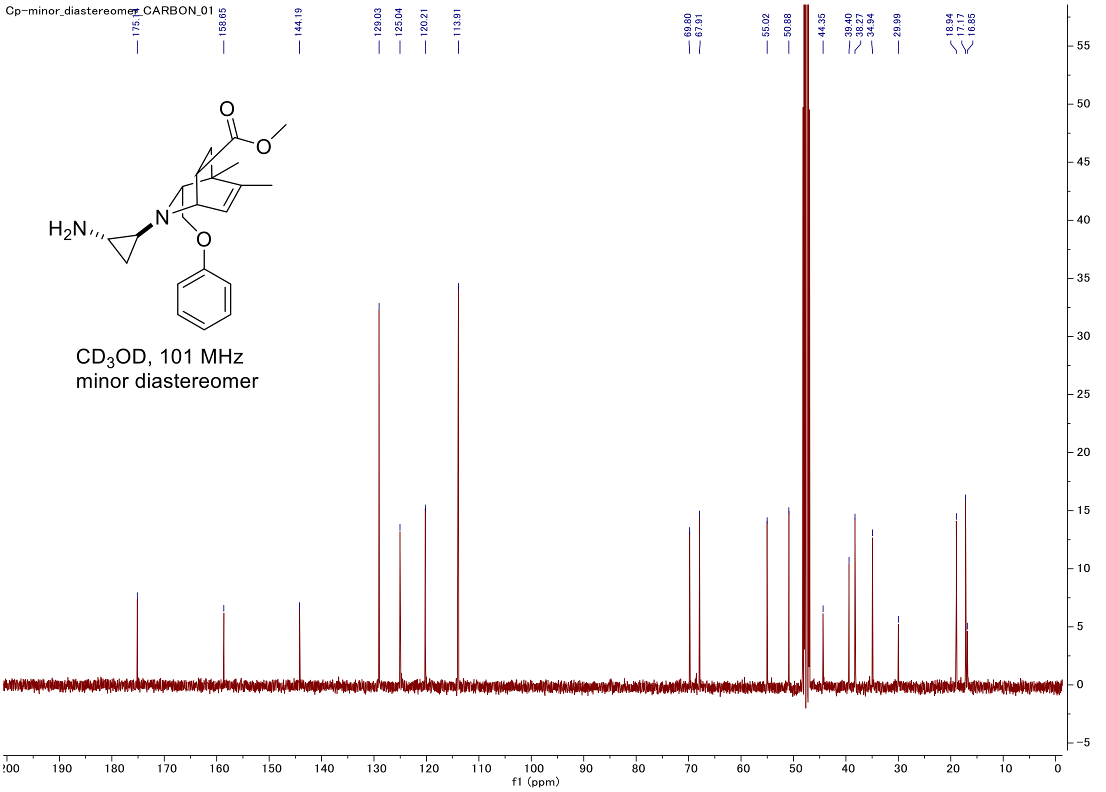

#### C-H COSY

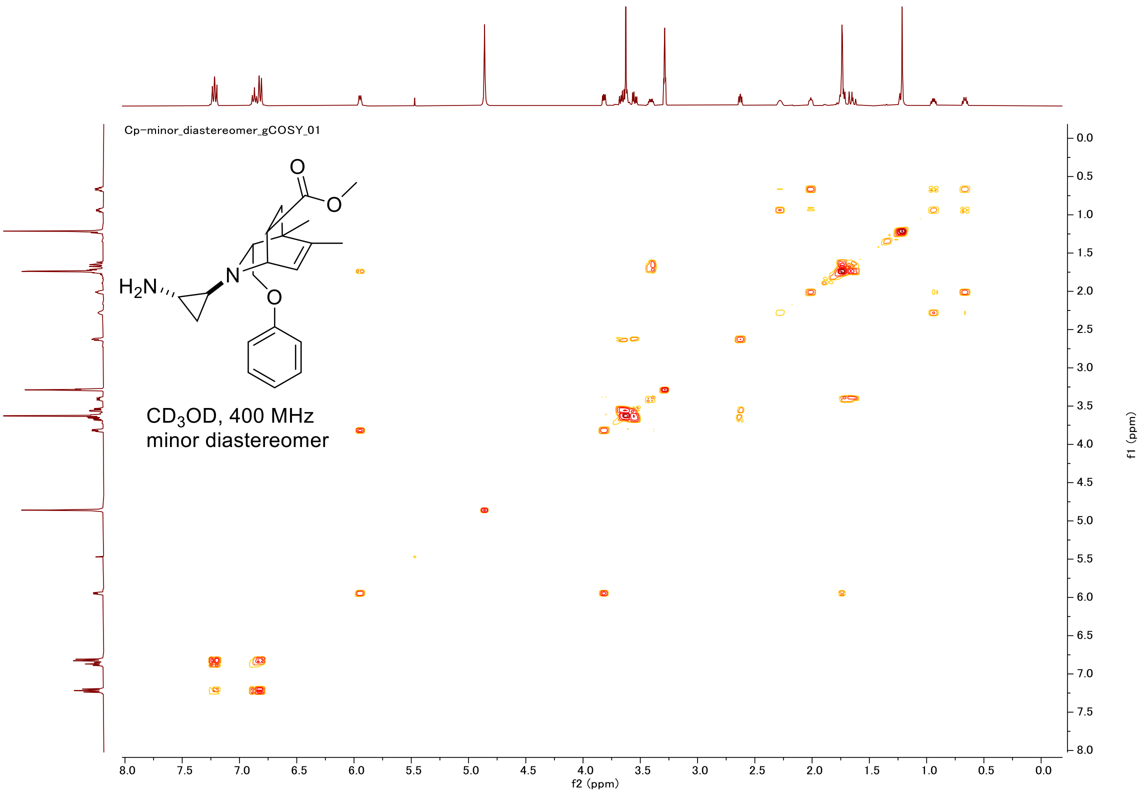

Cp−minor_diastereomer_gCOSY_01
H2N
N
O
O
O
CD3OD, 400 MHz
minor diastereomer
f2 (ppm)
f1 (ppm)


#### HSQC

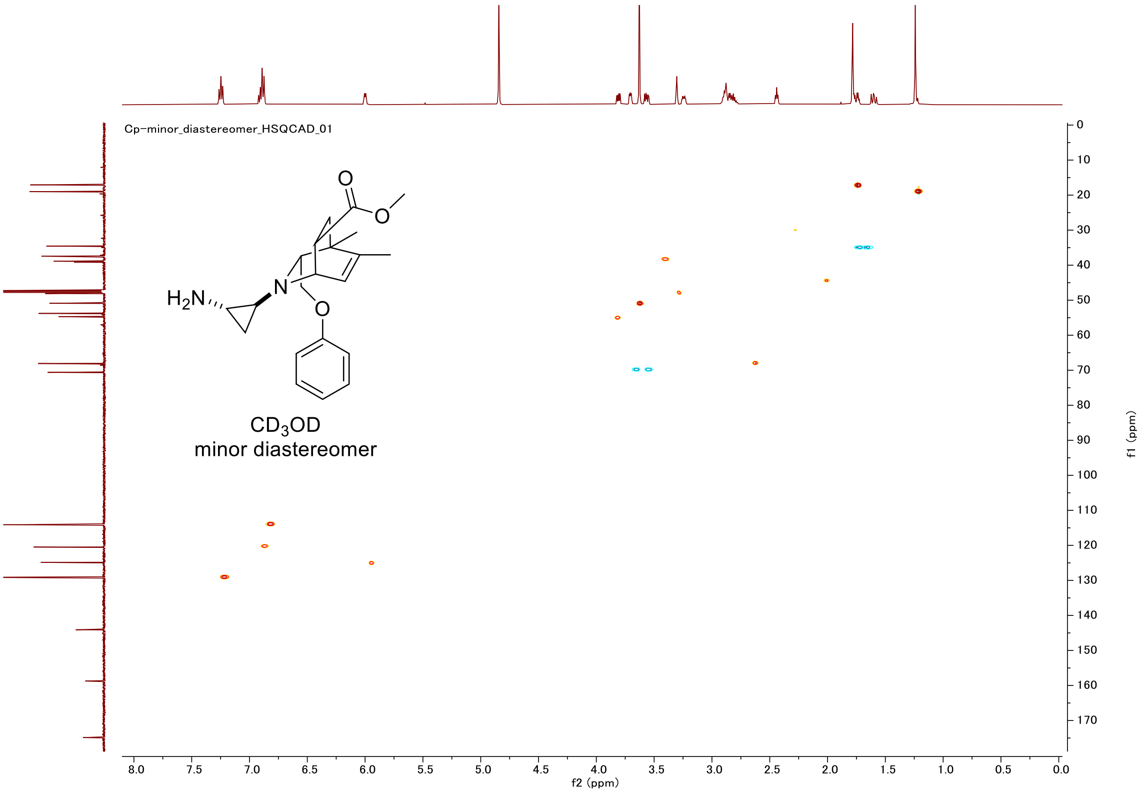

Cp−minor_diastereomer_HSQCAD_01
H2N
N
O
O
O
CD3OD
minor diastereomer
f2 (ppm)
f1 (ppm)

#### Proton (#21)

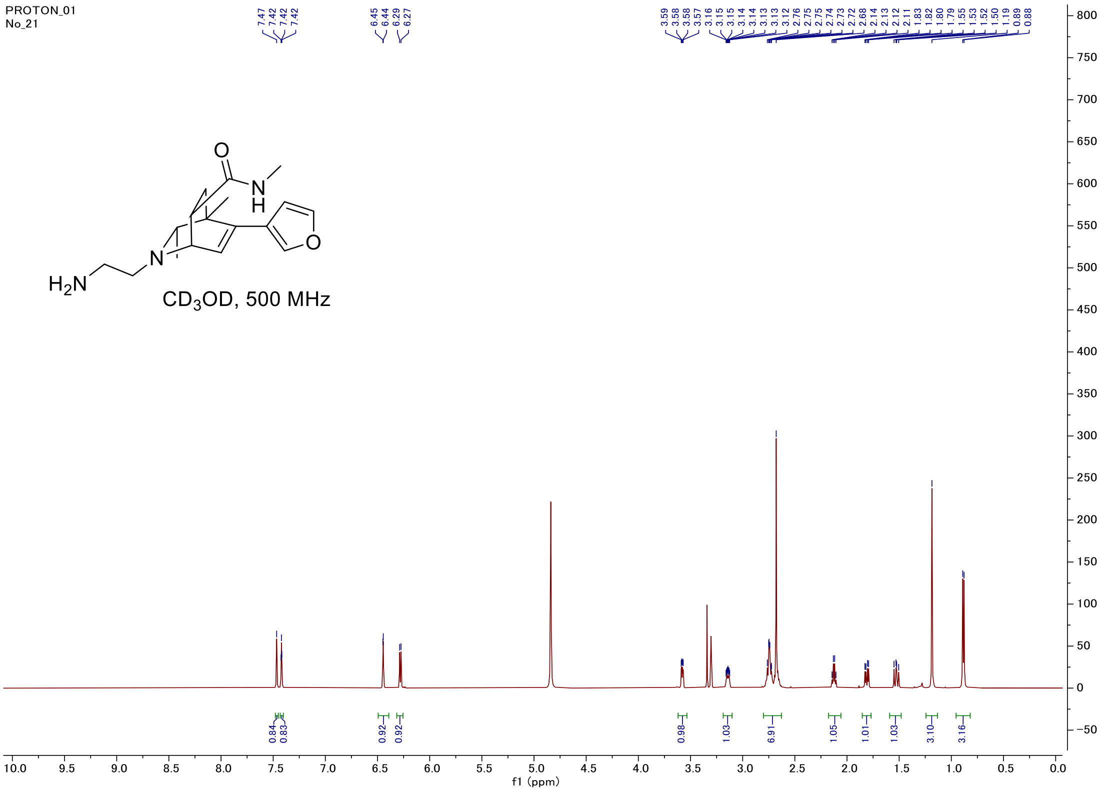


#### Carbon

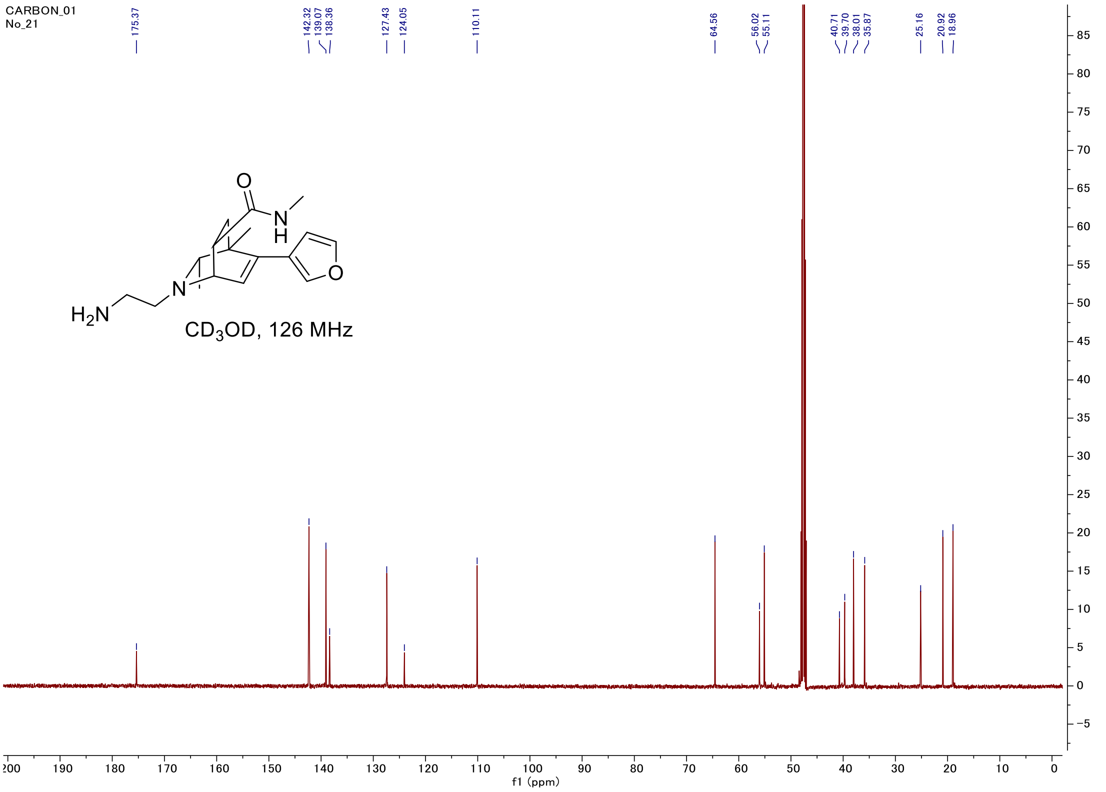

#### C-H COSY

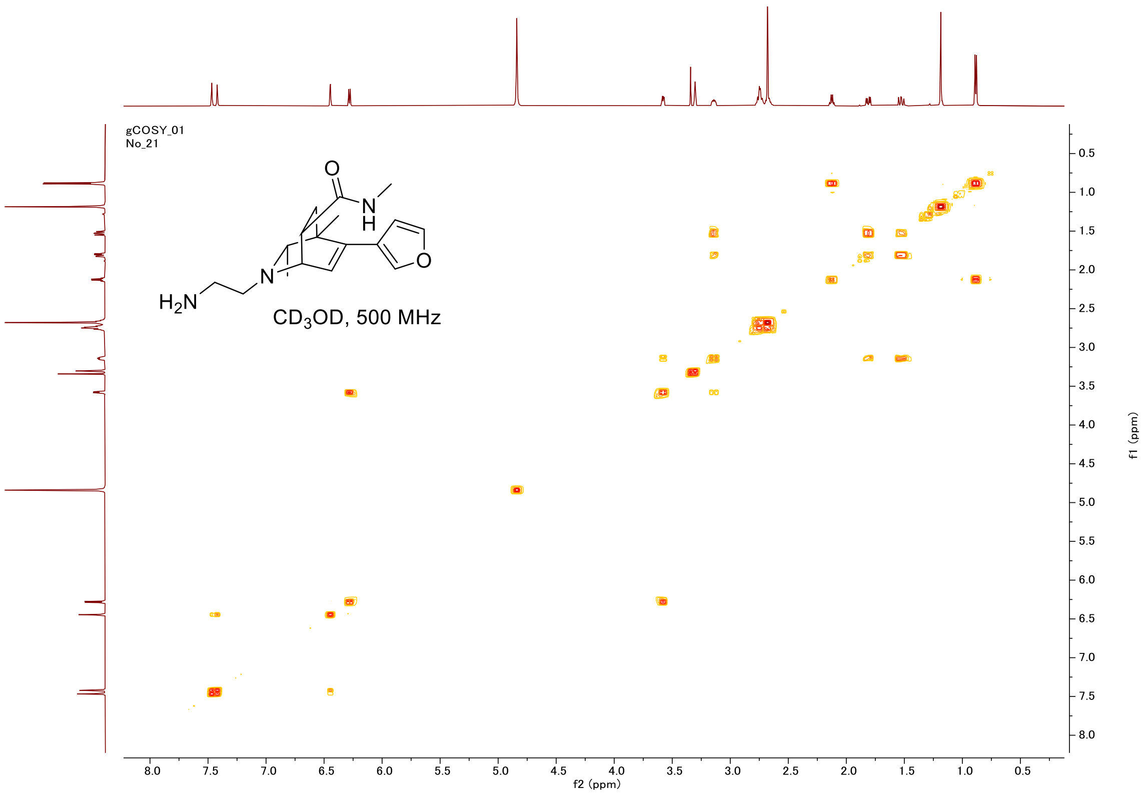

gCOSY_01
No_21
O
N
H
H2N
N
O
CD3OD, 500 MHz
f2 (ppm)
f1 (ppm)


#### HSQC

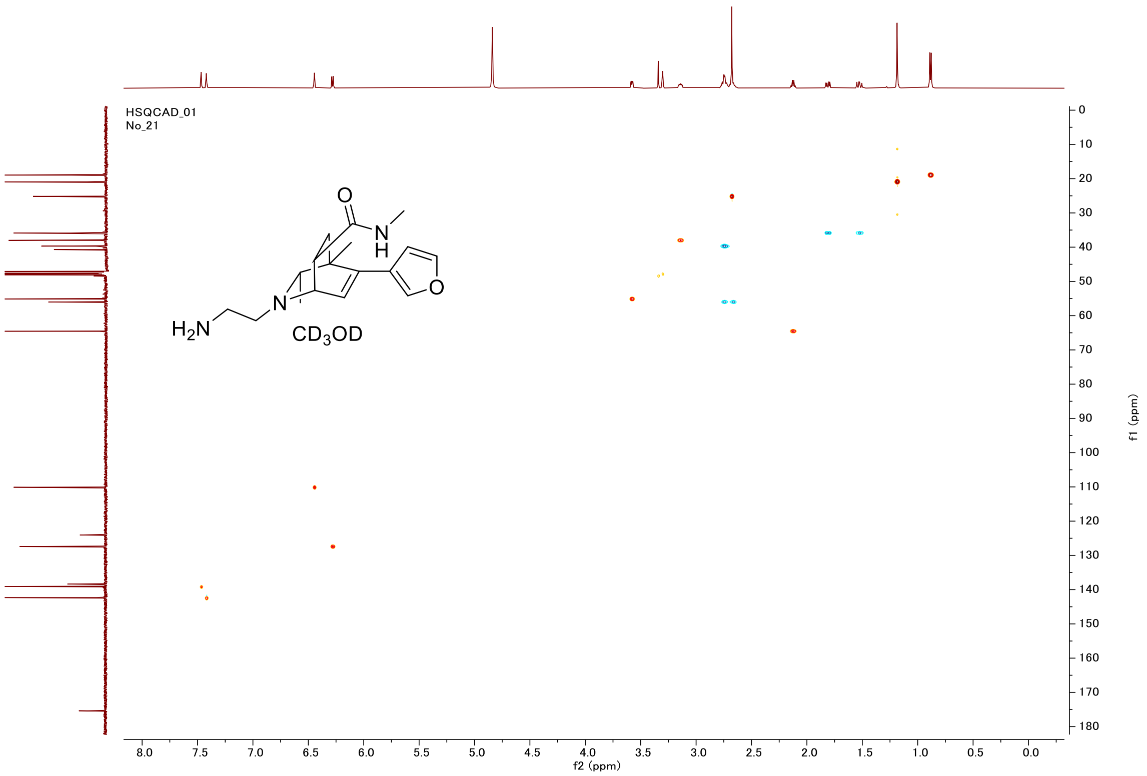

HSQCAD_01
No_21
O
N
H
H2N
N
O
CD3OD
f2 (ppm)
f1 (ppm)

### Proton (#33)

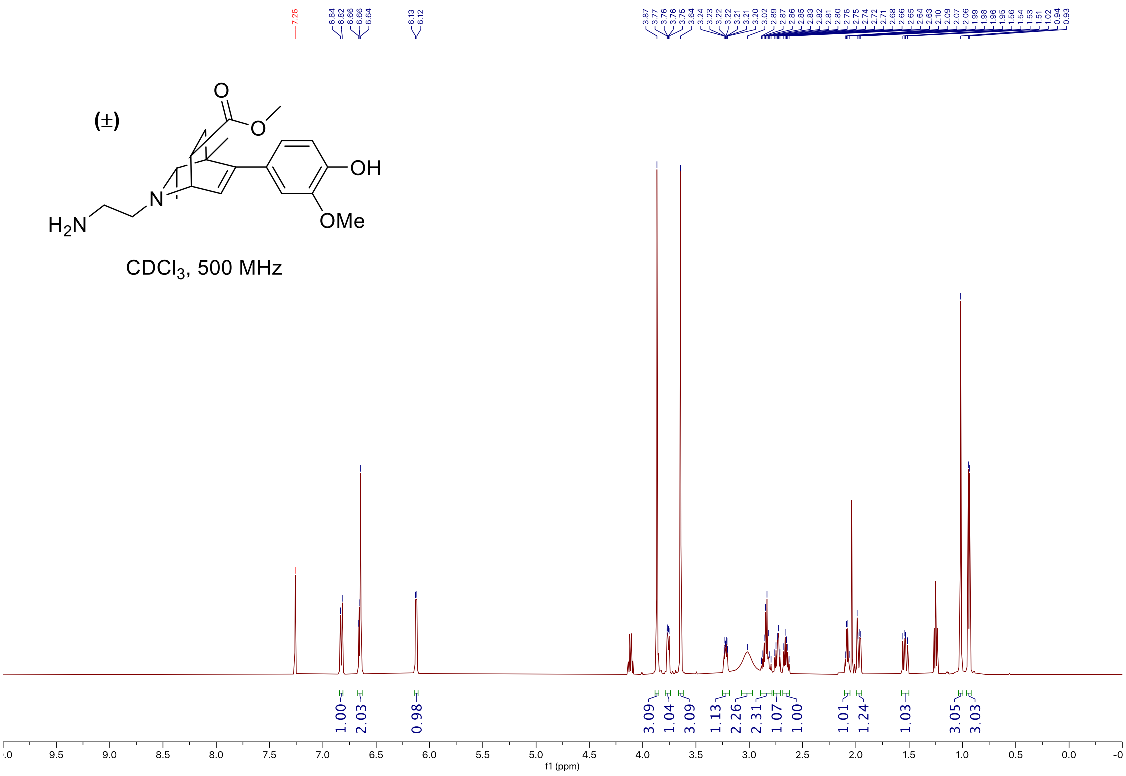


### Carbon

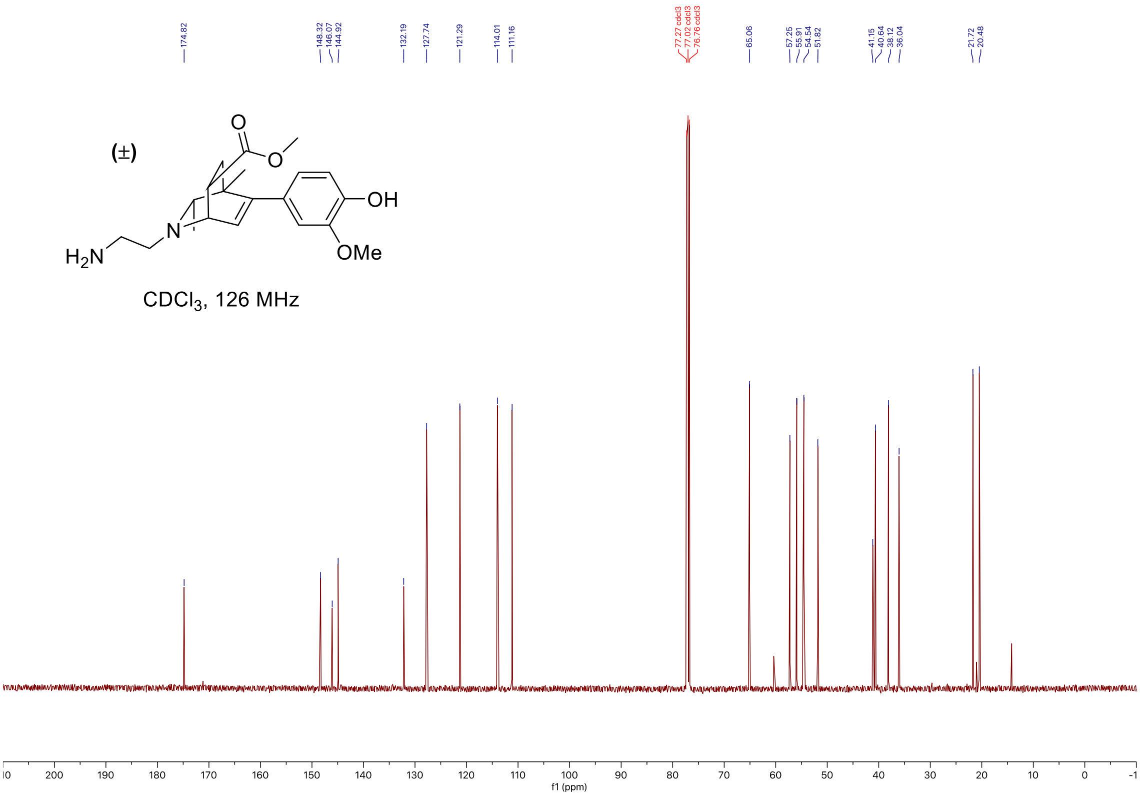

#### C-H COSY

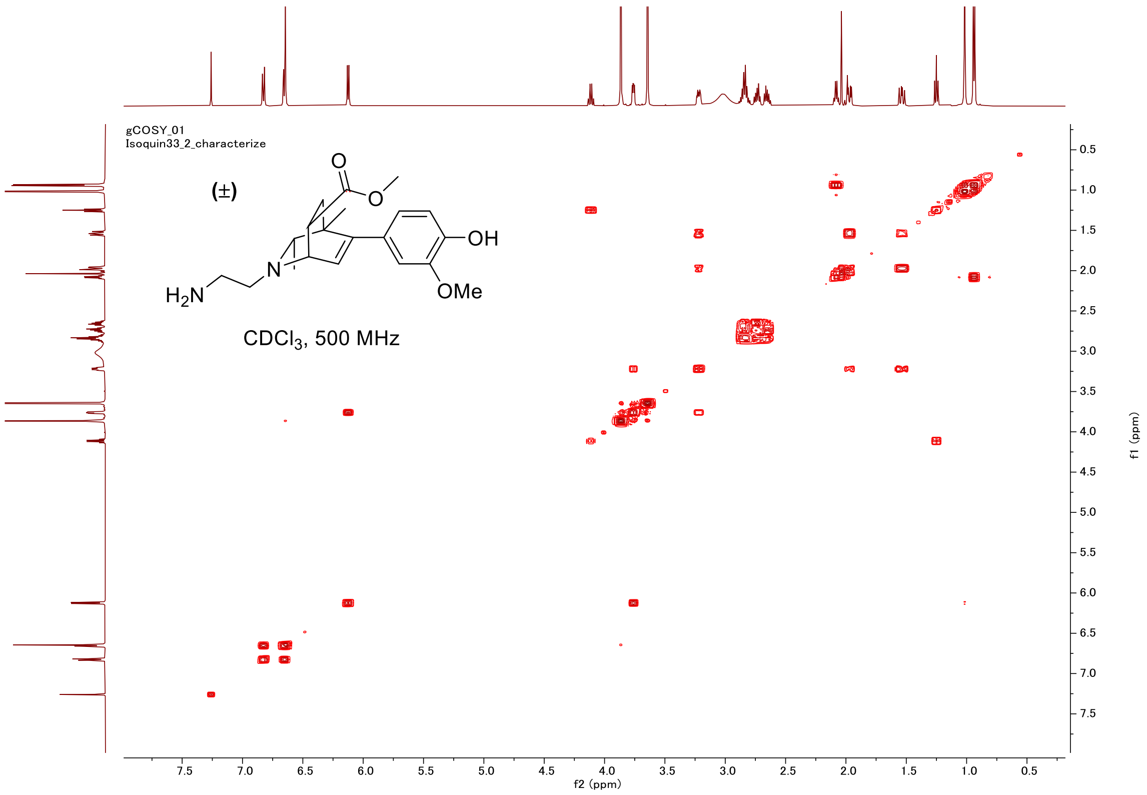

gCOSY_01
Isoquin33_2_characterize
(±)
OH
OMe
$H_2N$
N
CDCl$_3$, 500 MHz
f2 (ppm)
f1 (ppm)


#### HSQC

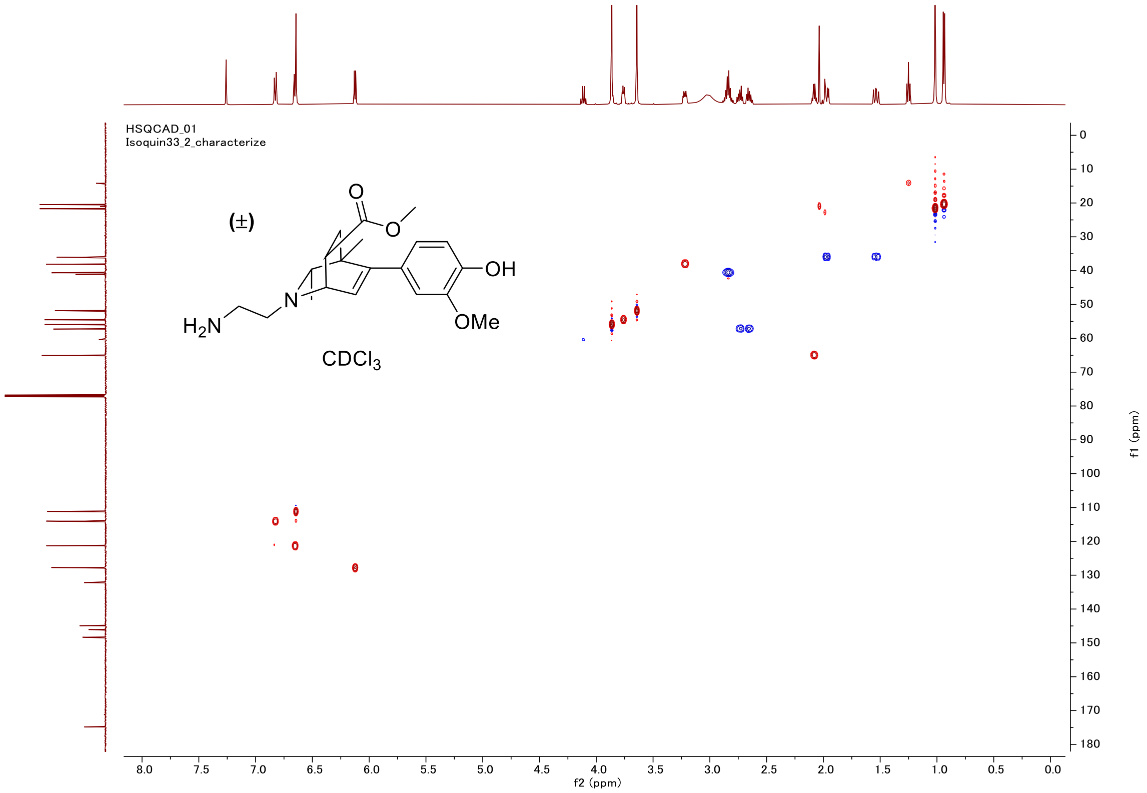

HSQCAD_01
Isoquin33_2_characterize
(±)
OH
OMe
$H_2N$
N
CDCl$_3$
f2 (ppm)
f1 (ppm)

#### Proton (#39)

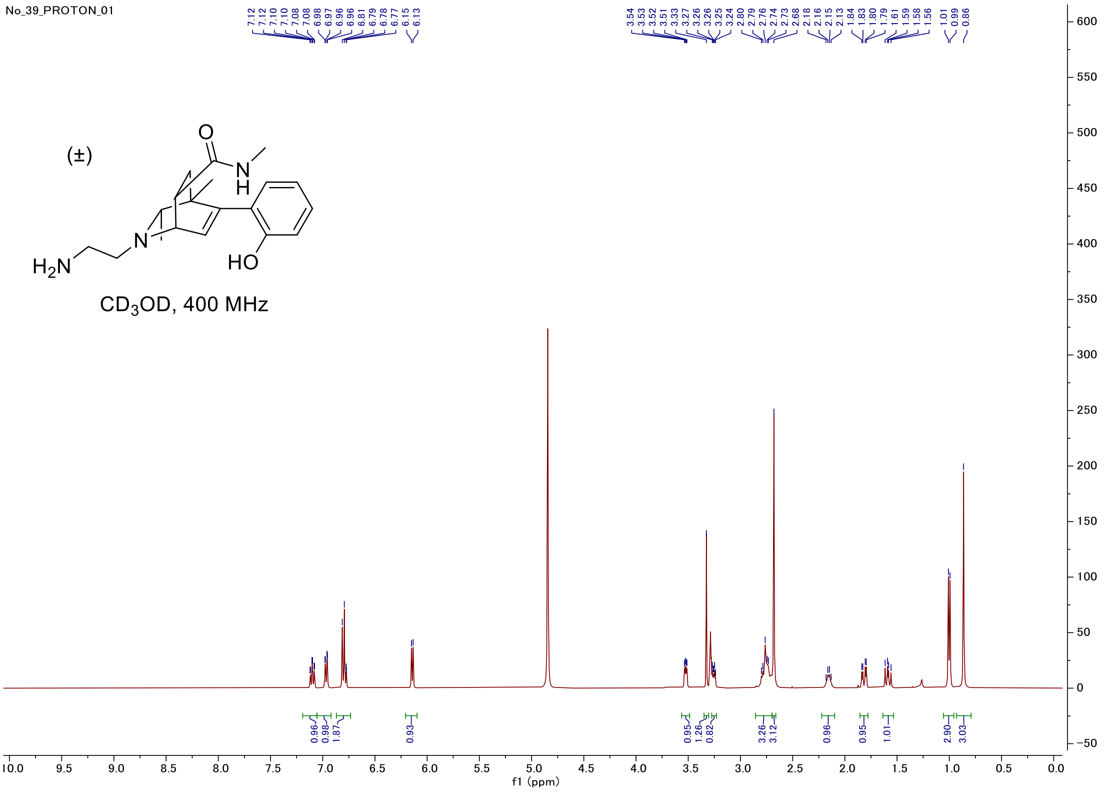


#### Carbon

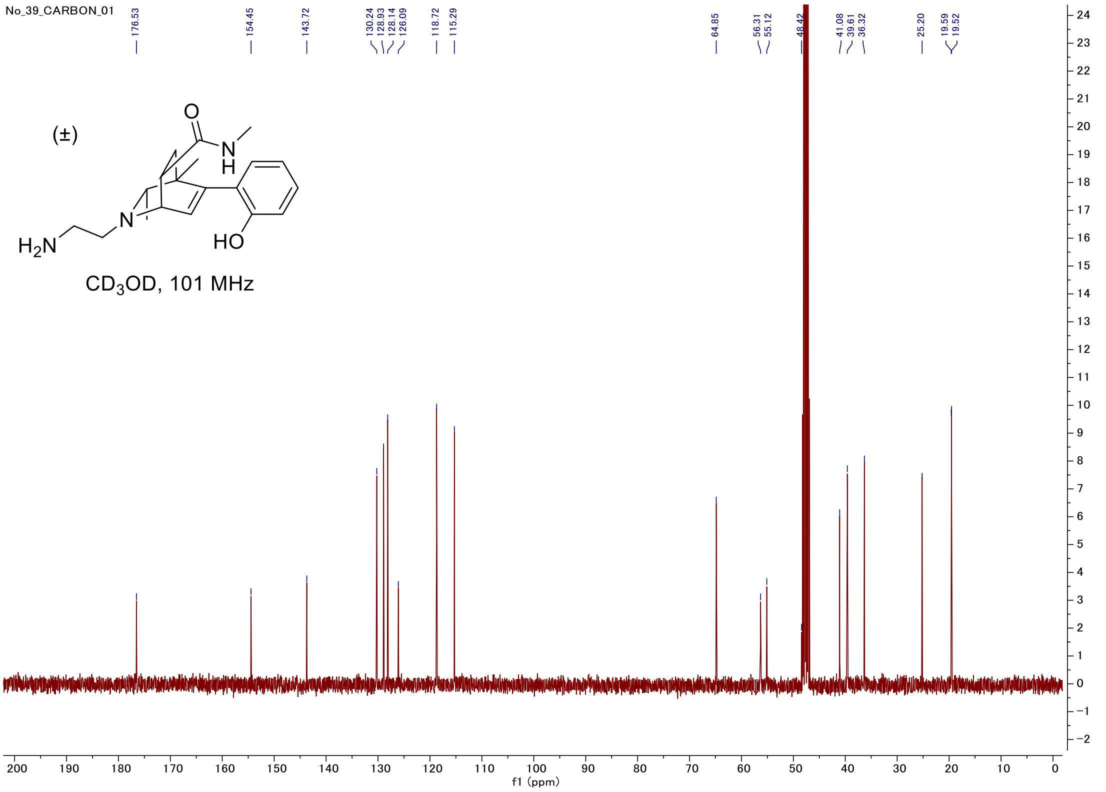

#### C-H COSY

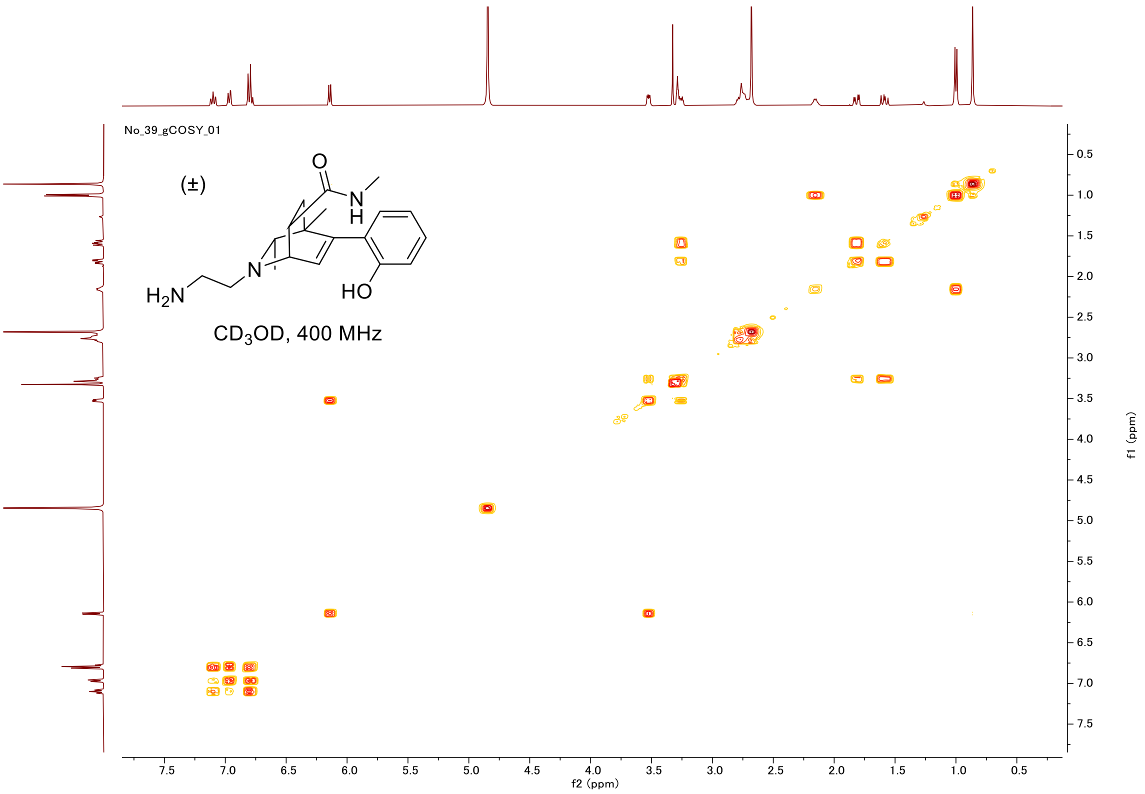
No_39_gCOSY_01
(±)
O
N
H
N
H2N
HO
CD3OD, 400 MHz
f2 (ppm)
f1 (ppm)

#### HSQC

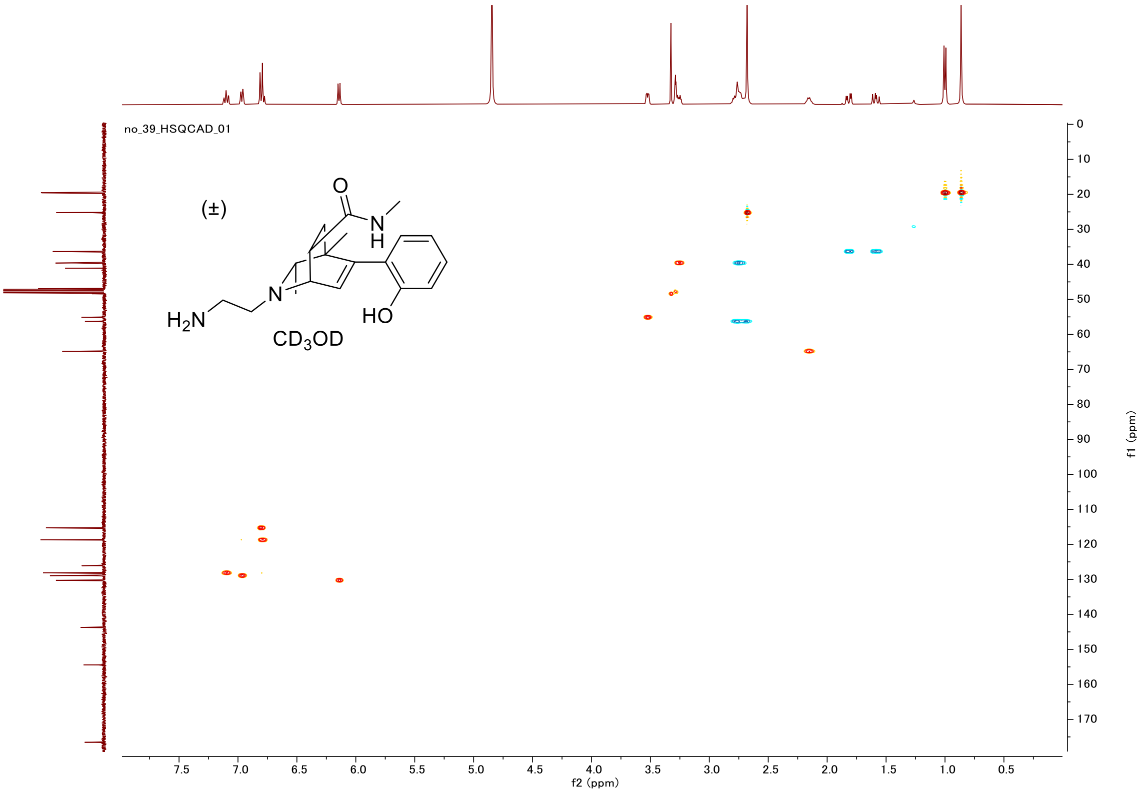
no_39_HSQCAD_01
(±)
O
N
H
N
H2N
HO
CD3OD
f2 (ppm)
f1 (ppm)

#### Proton (#53)

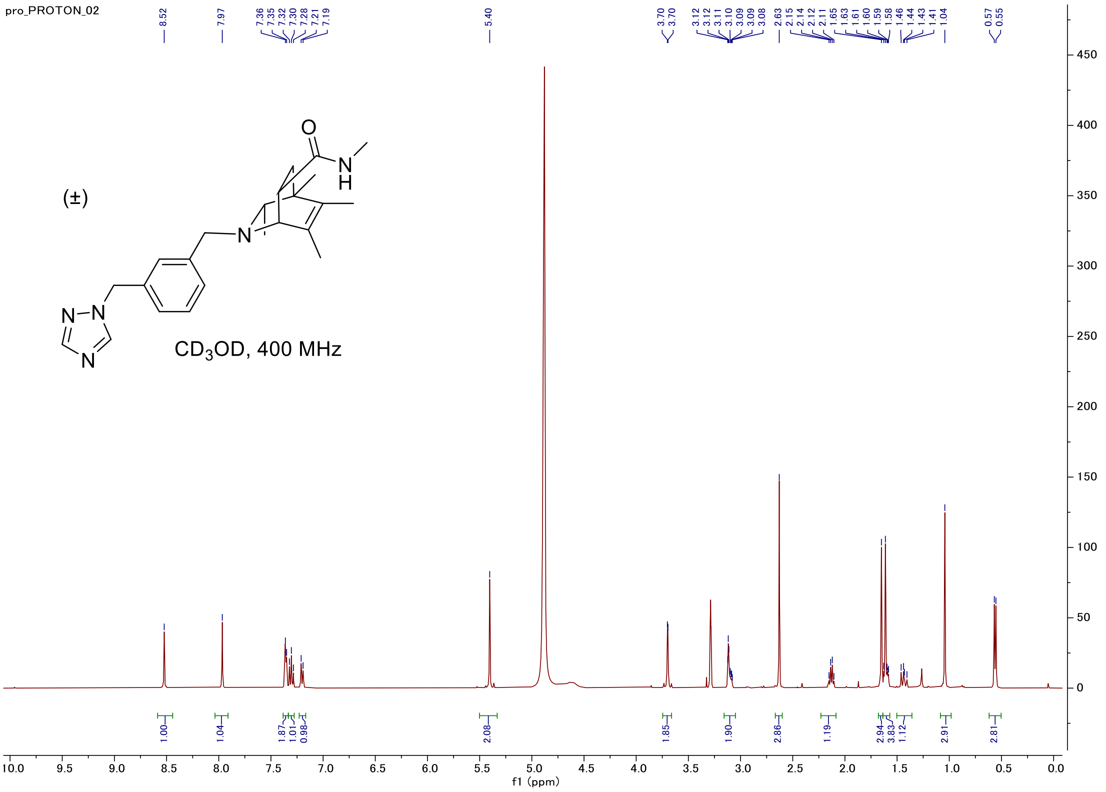


#### Carbon

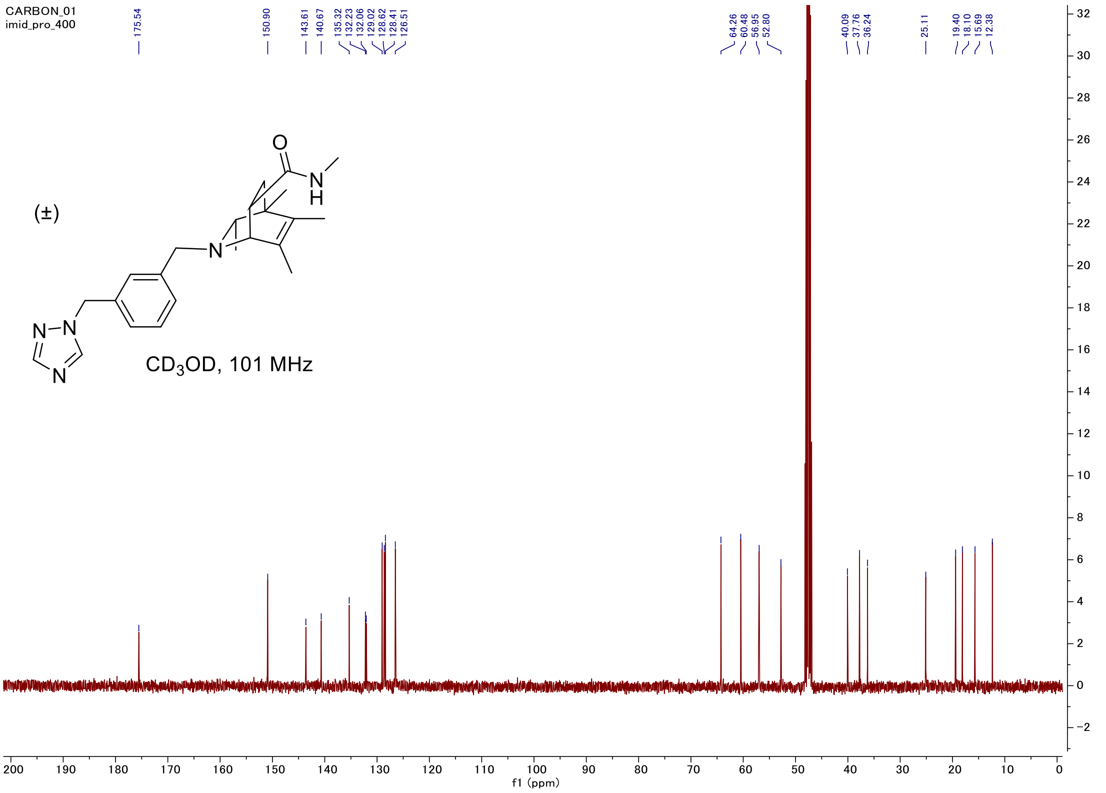

#### C-H COSY

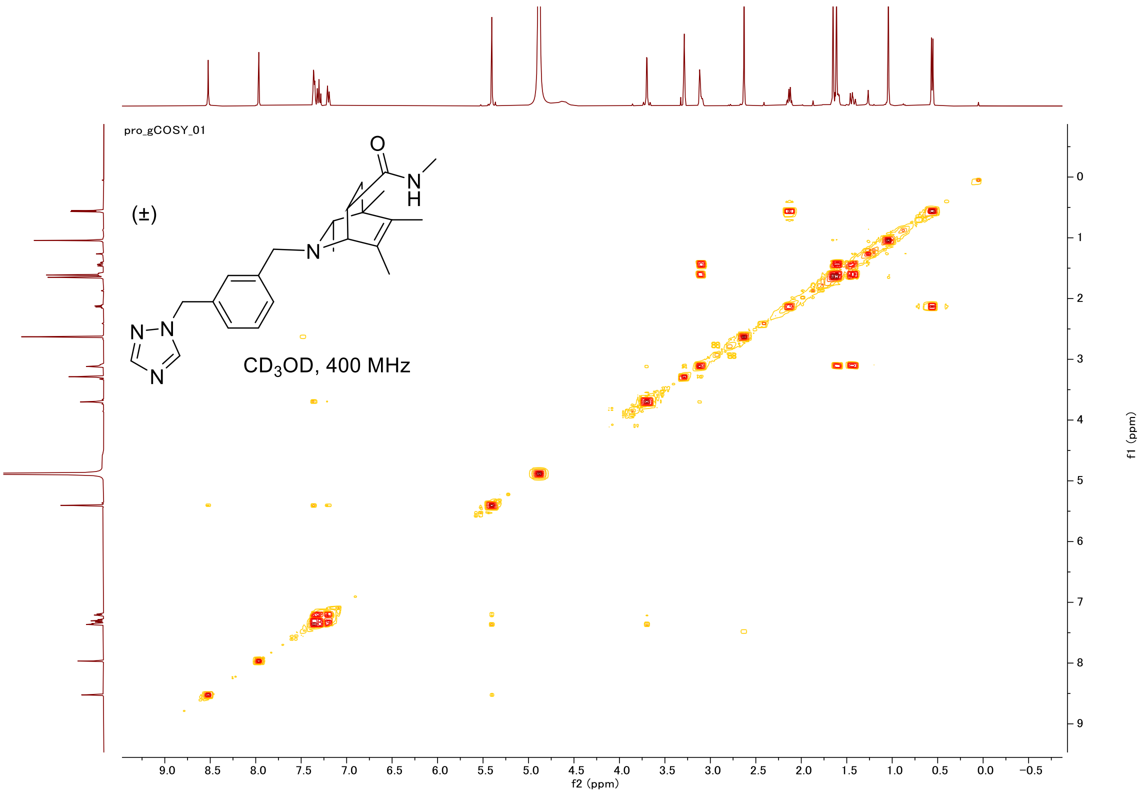
pro_gCOSY_01
(±)
CD3OD, 400 MHz
f2 (ppm)
f1 (ppm)

#### HSQC

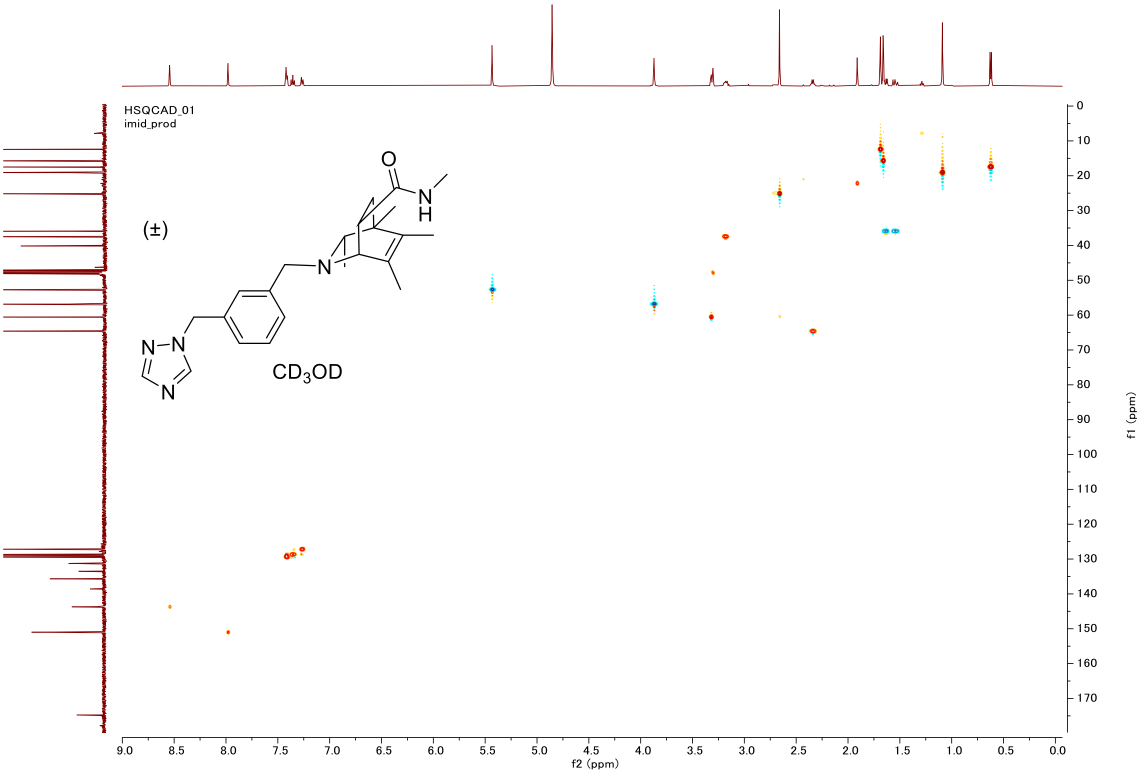
HSQCAD_01
imid_prod
(±)
CD3OD
f2 (ppm)
f1 (ppm)

#### Proton (#56)

PROTON_01
Isoquin56_characterize

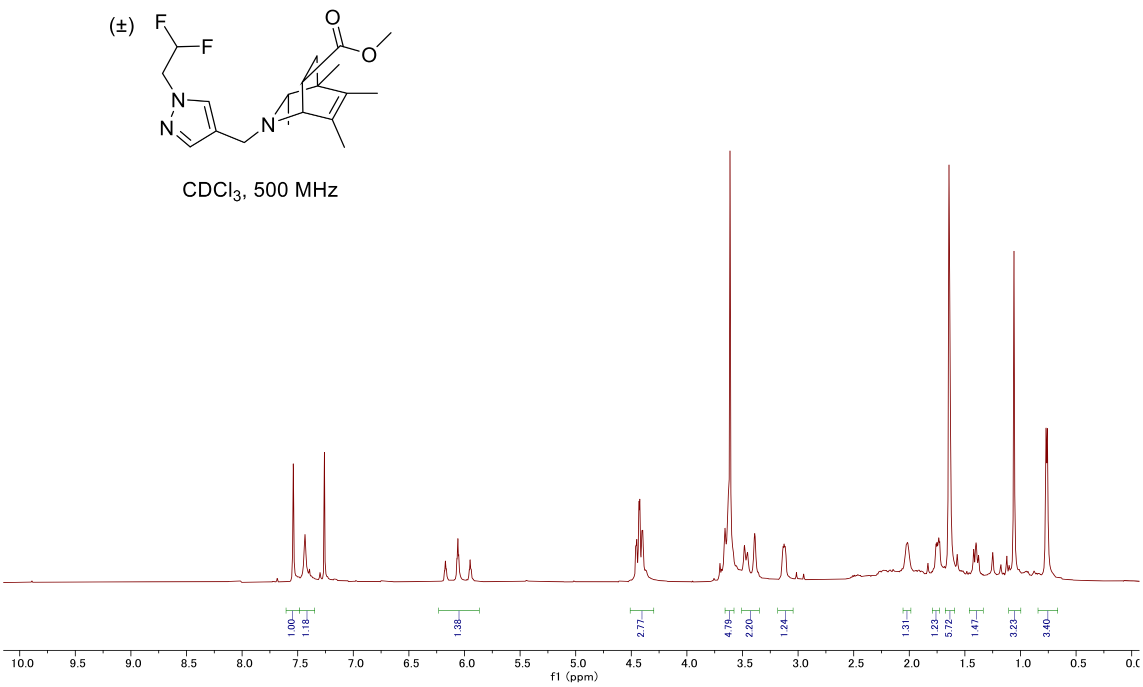


#### Carbon

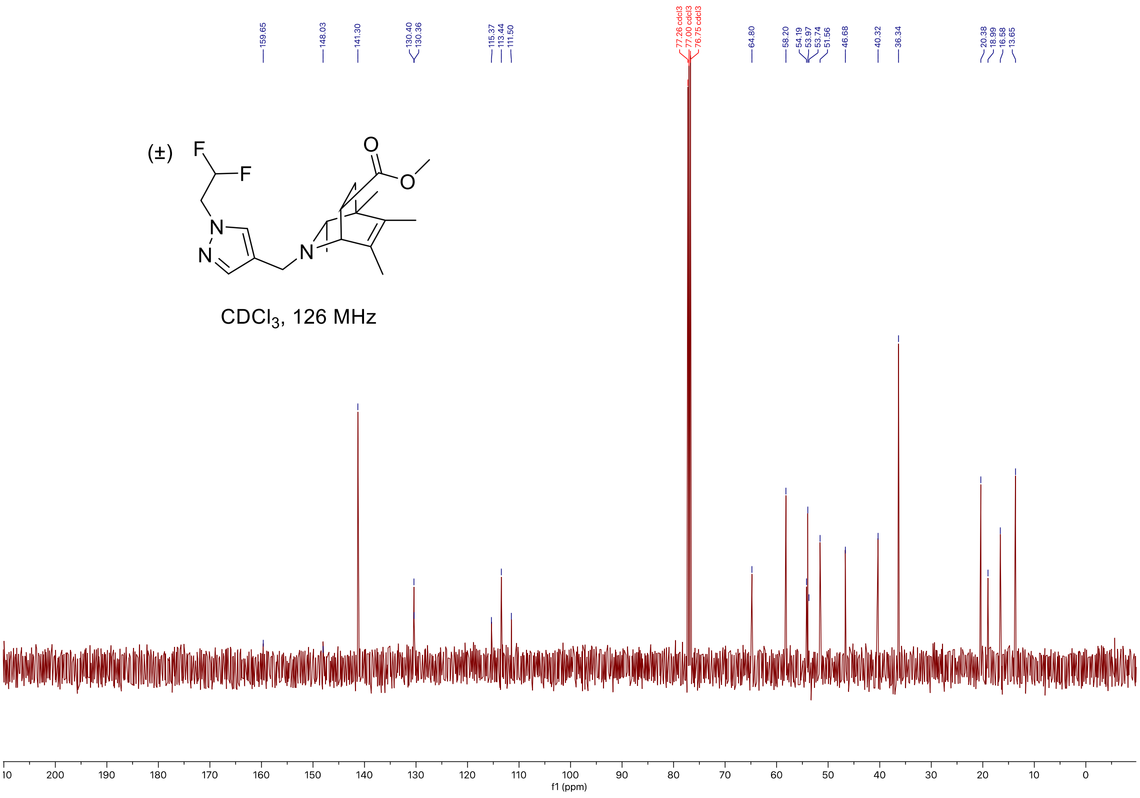

#### C-H COSY

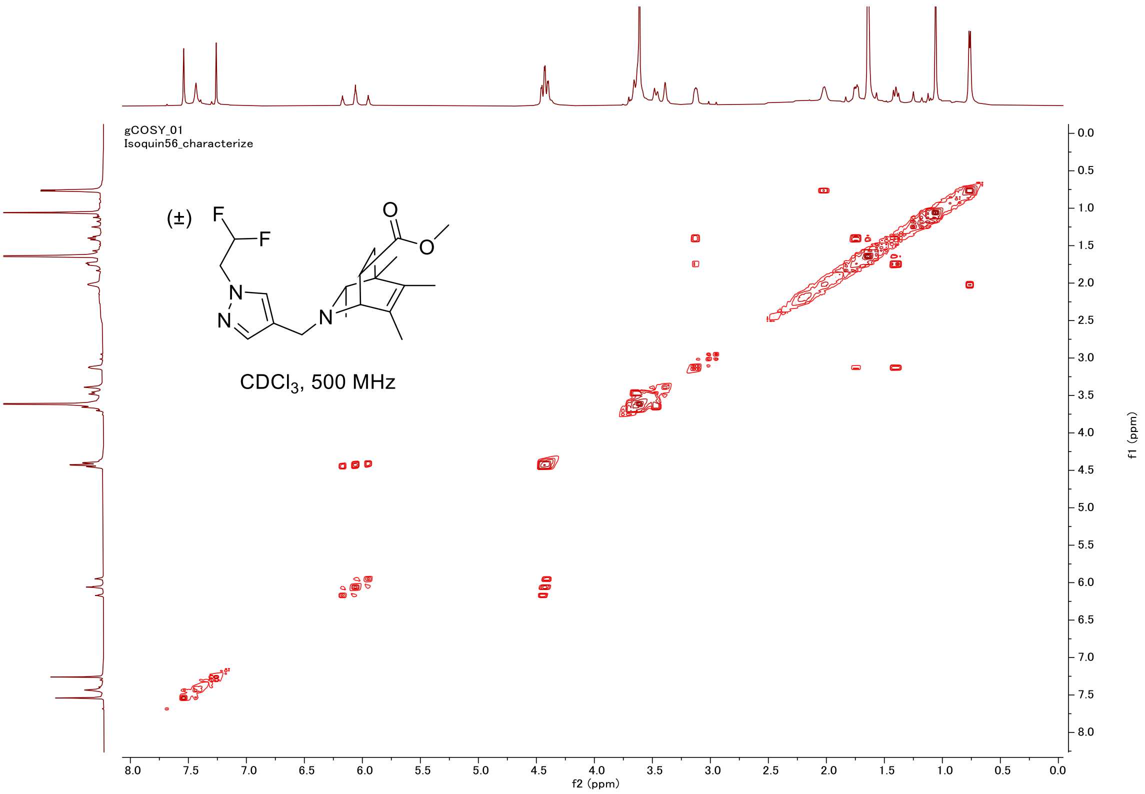
gCOSY_01
Isoquin56_characterize
(±)
CDCl3, 500 MHz
f2 (ppm)
f1 (ppm)

#### HSQC

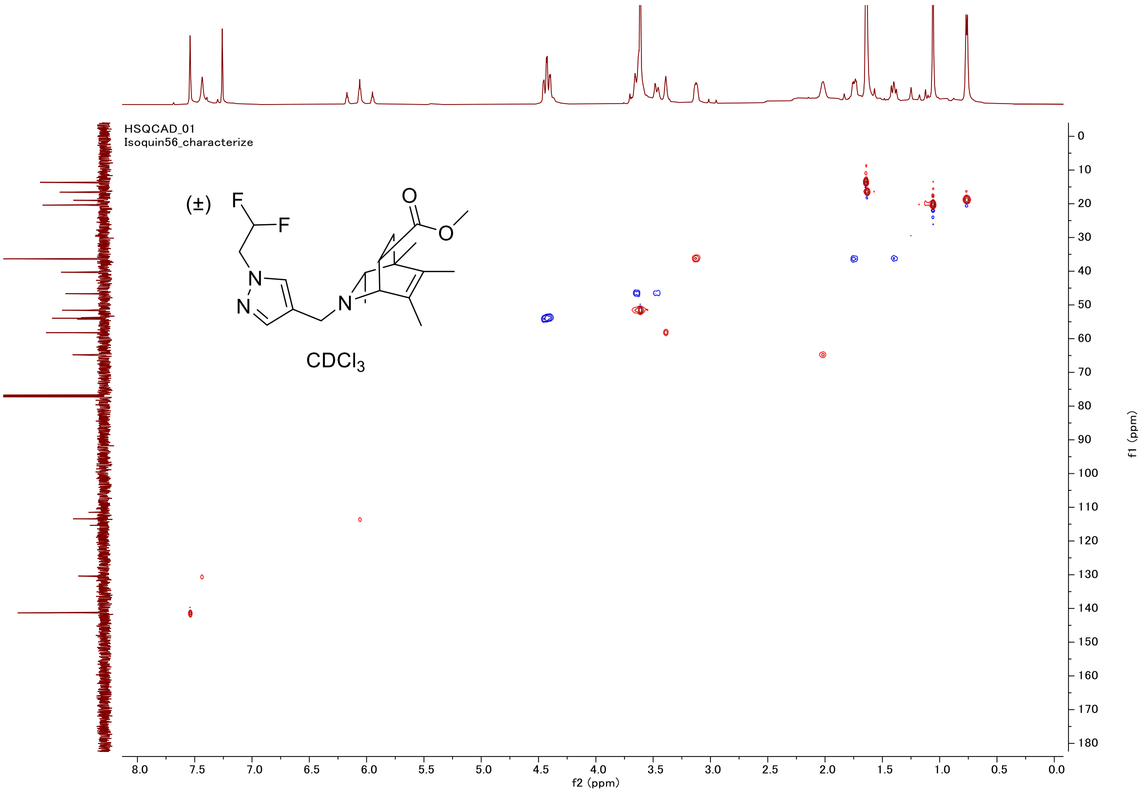
HSQCAD_01
Isoquin56_characterize
(±)
CDCl3
f2 (ppm)
f1 (ppm)

#### Proton (#57)

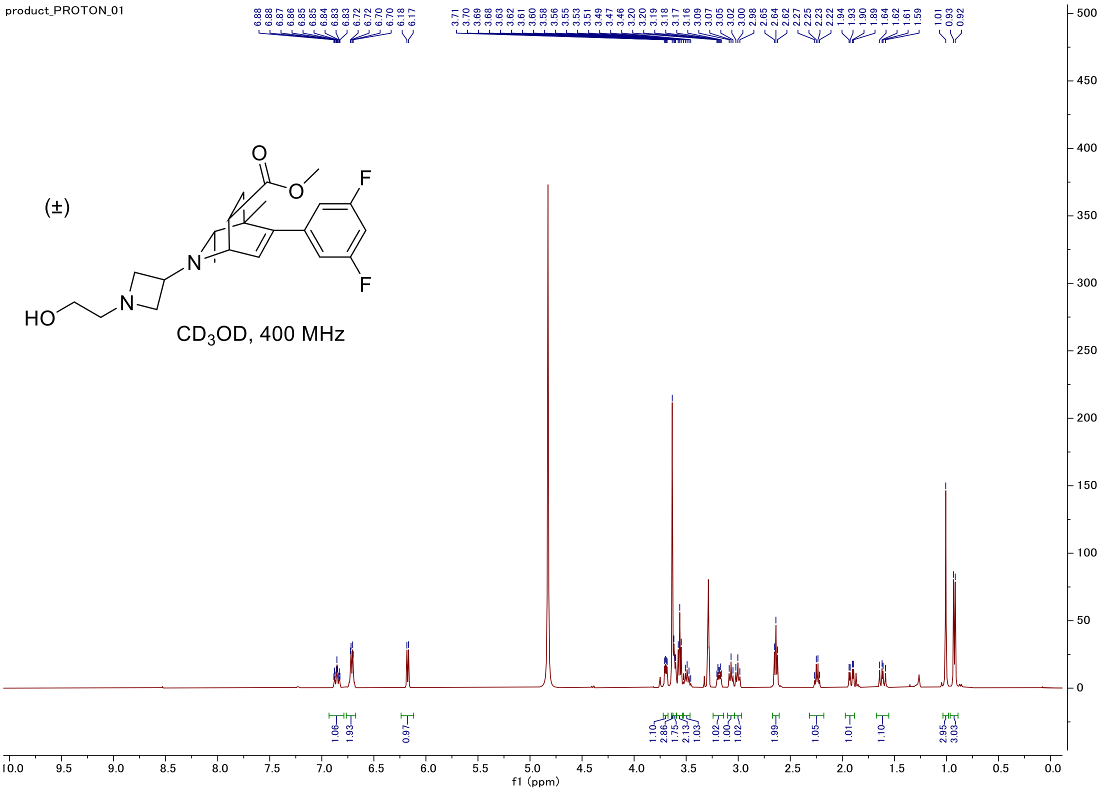


#### Carbon

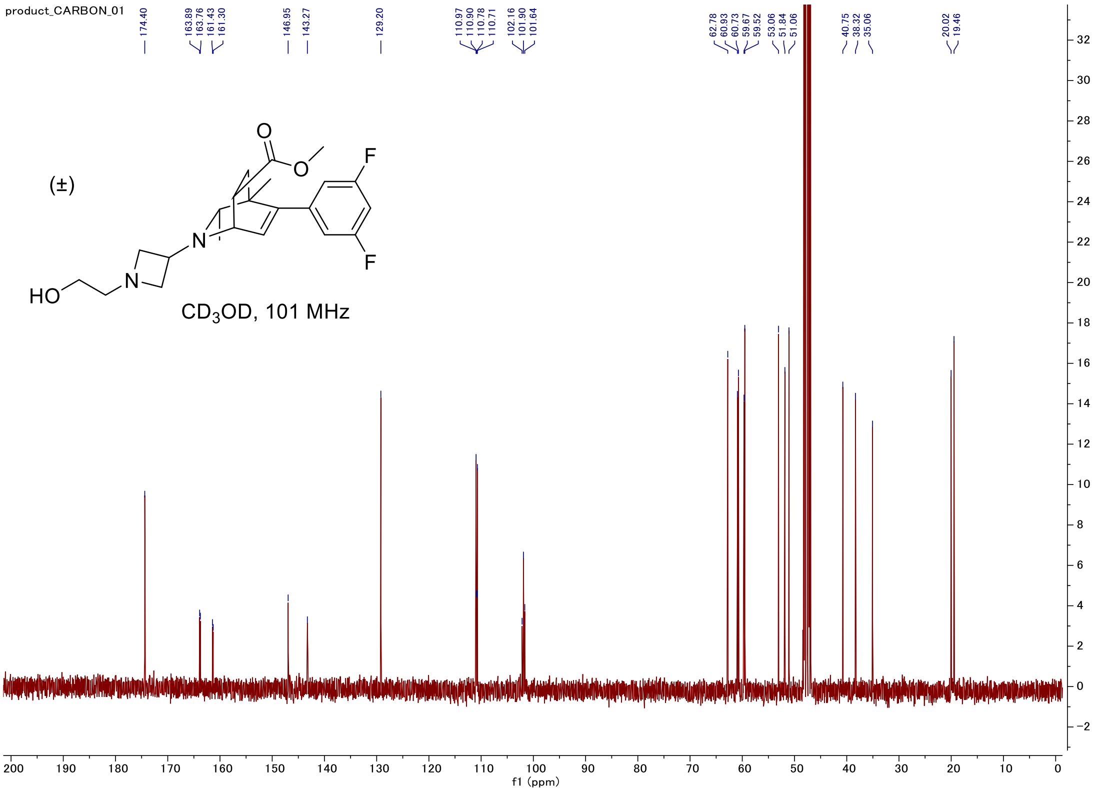

#### Fluorine

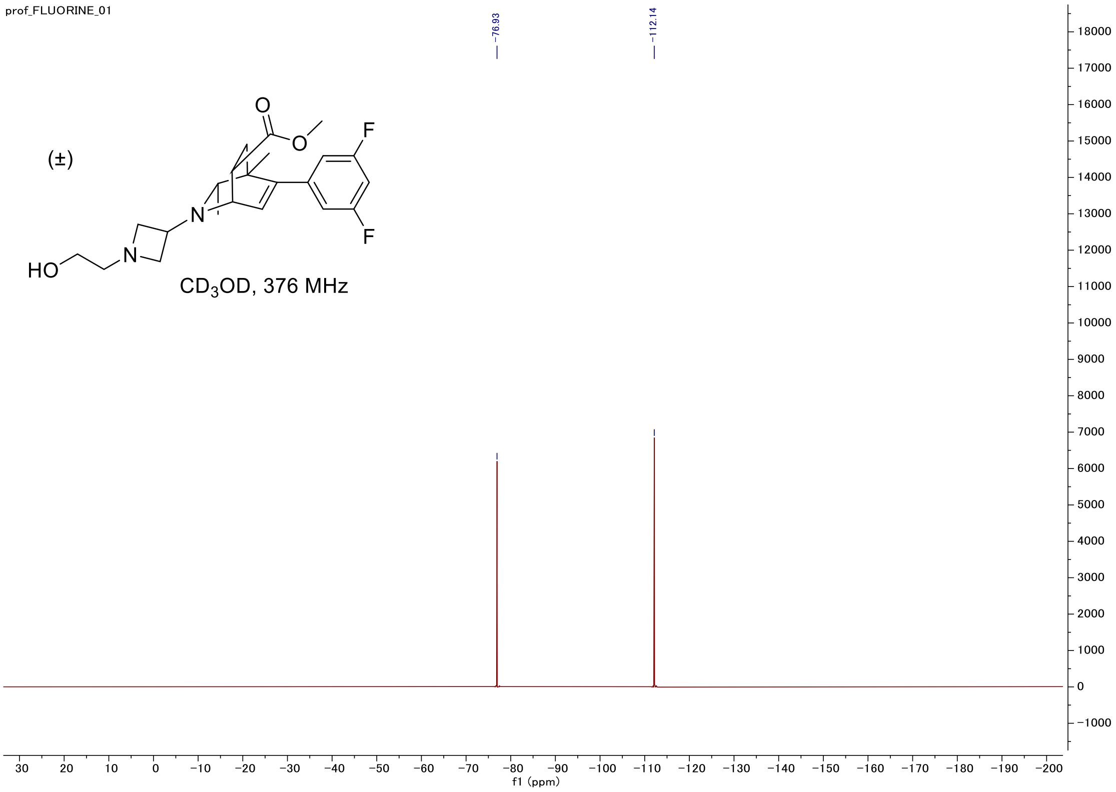


#### C-H COSY

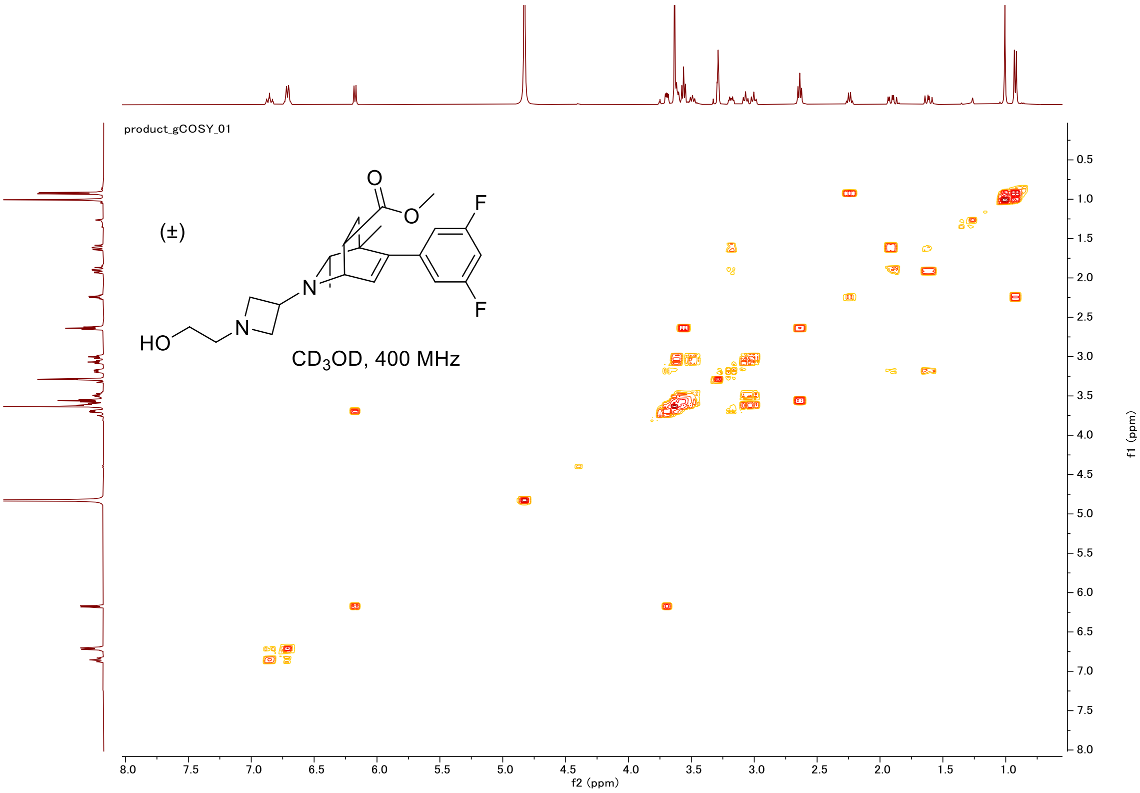

#### HSQC

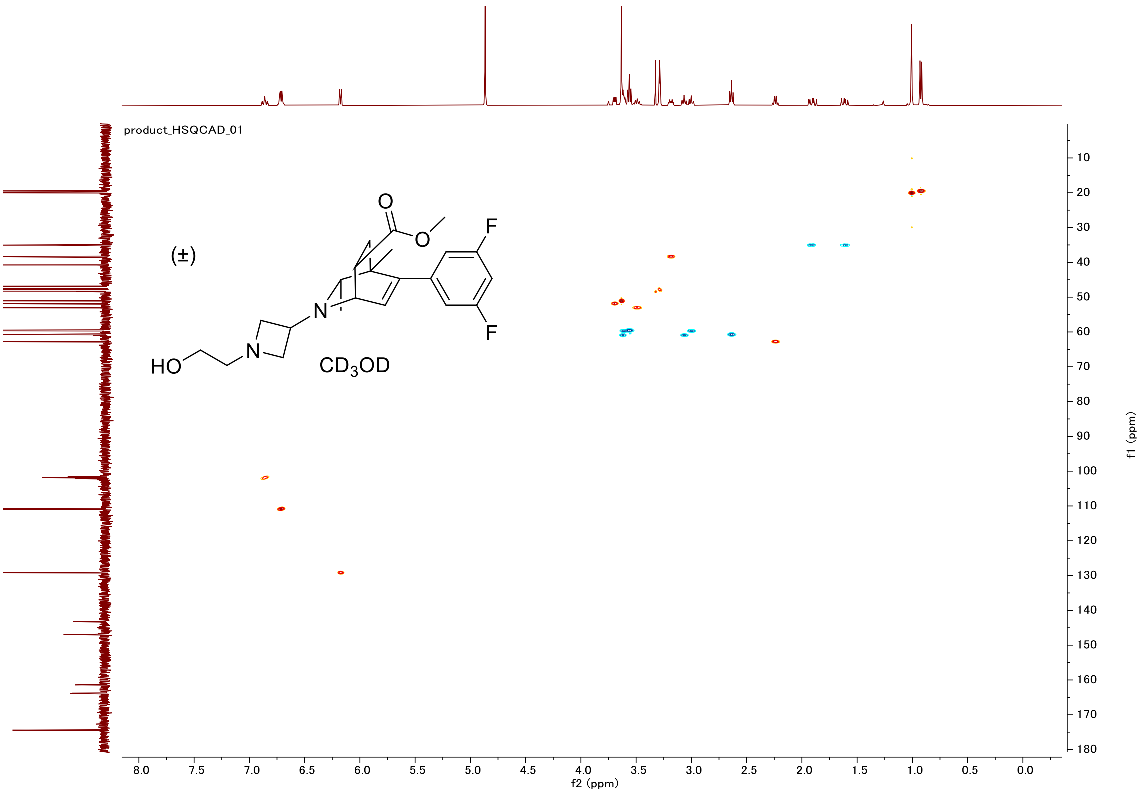

#### Proton (#58)

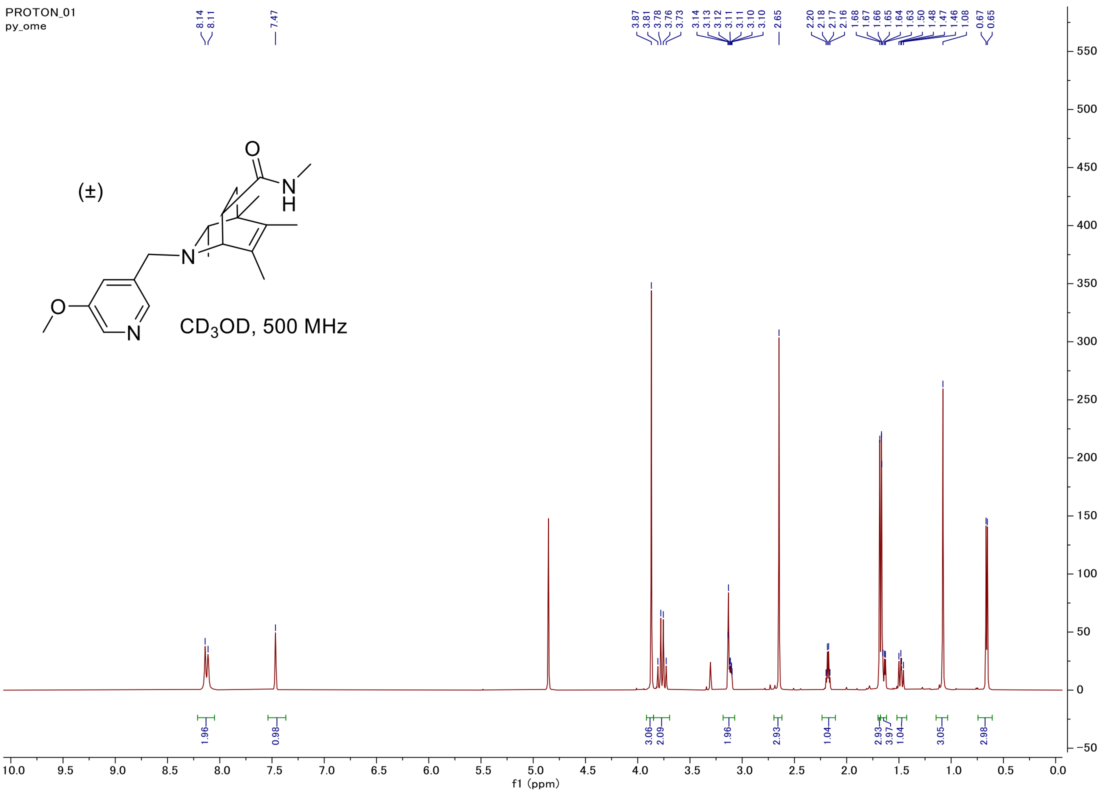

#### Carbon

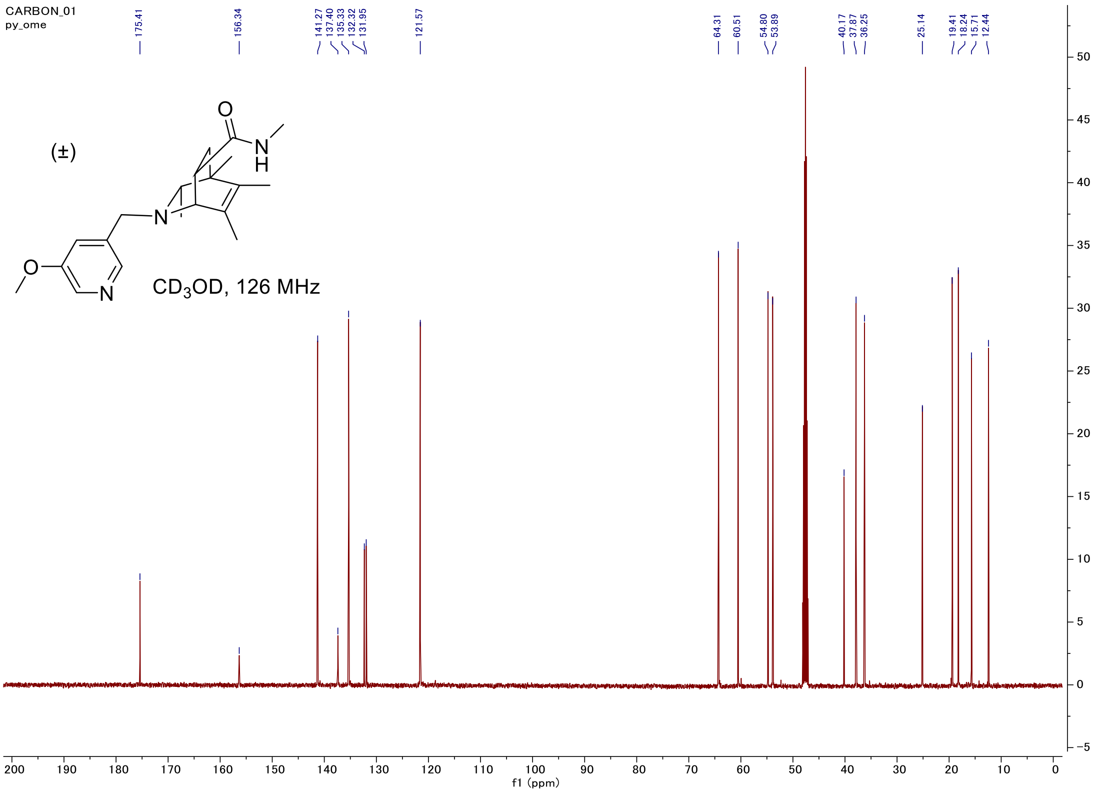

#### C-H COSY

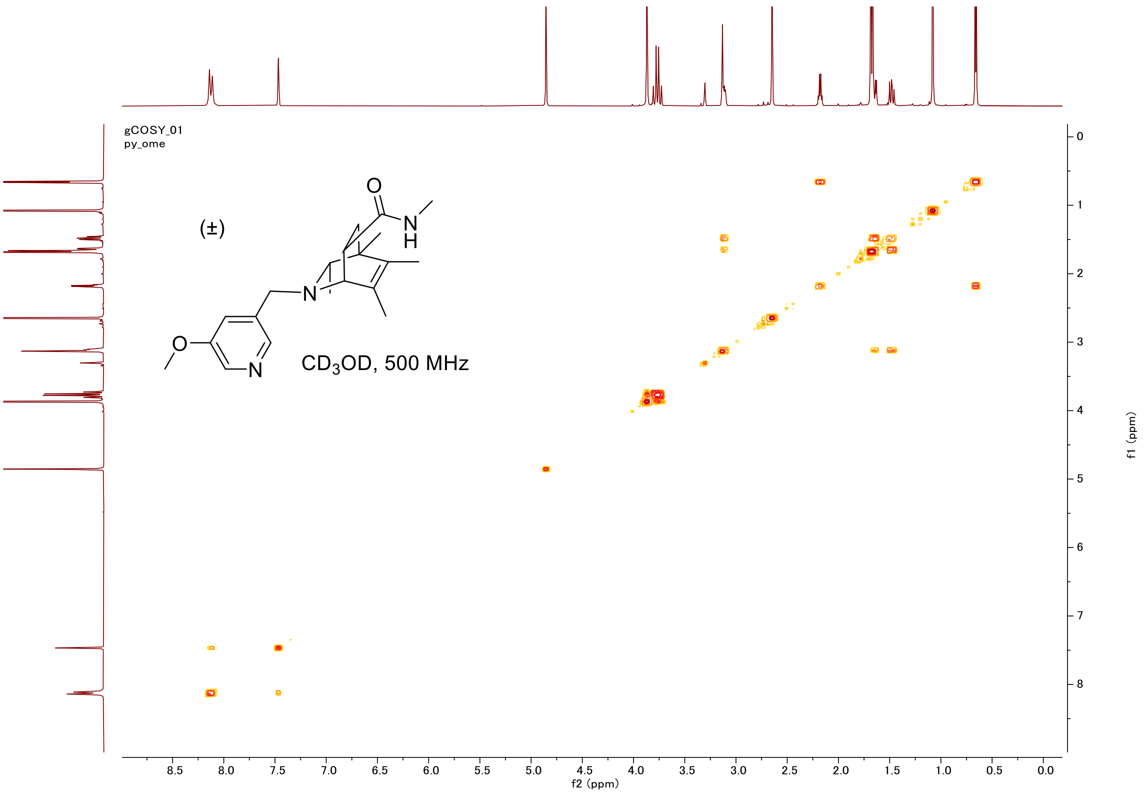
gCOSY_01
py_ome
(±)
O
N
H
N
O
N
CD3OD, 500 MHz
f2 (ppm)
f1 (ppm)

#### HSQC

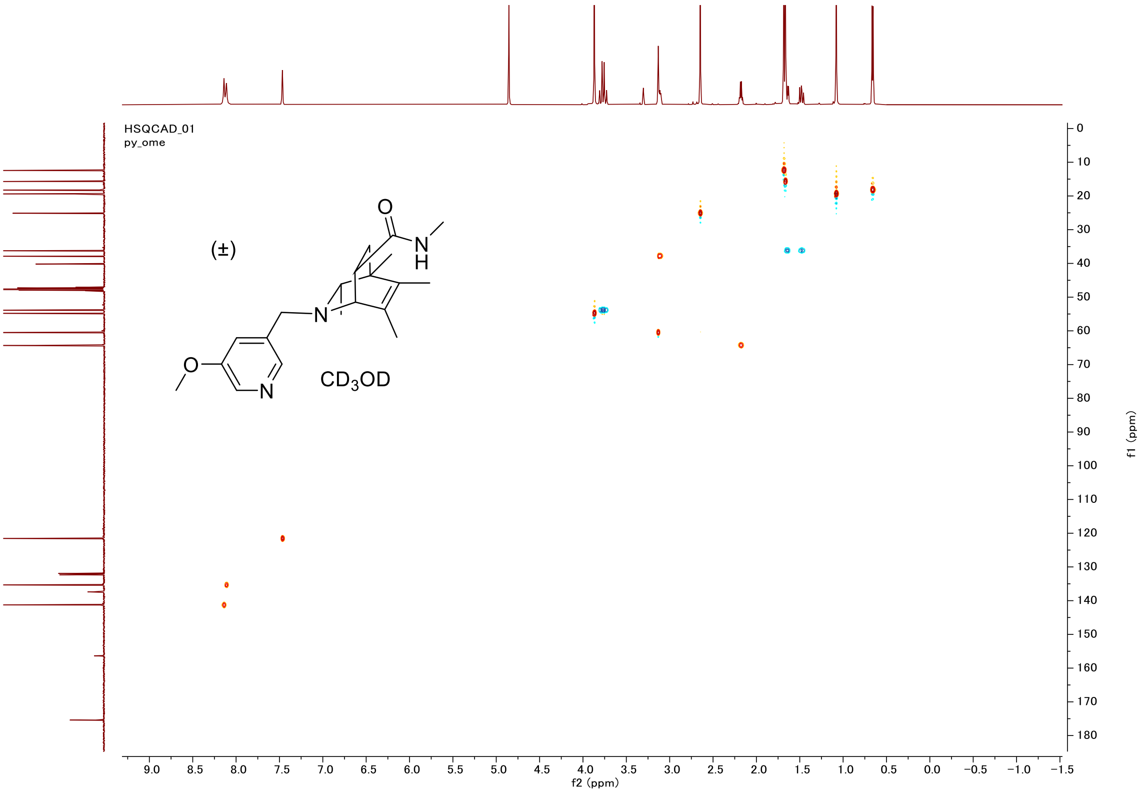
HSQCAD_01
py_ome
(±)
O
N
H
N
O
N
CD3OD
f2 (ppm)
f1 (ppm)

#### Proton (#61)

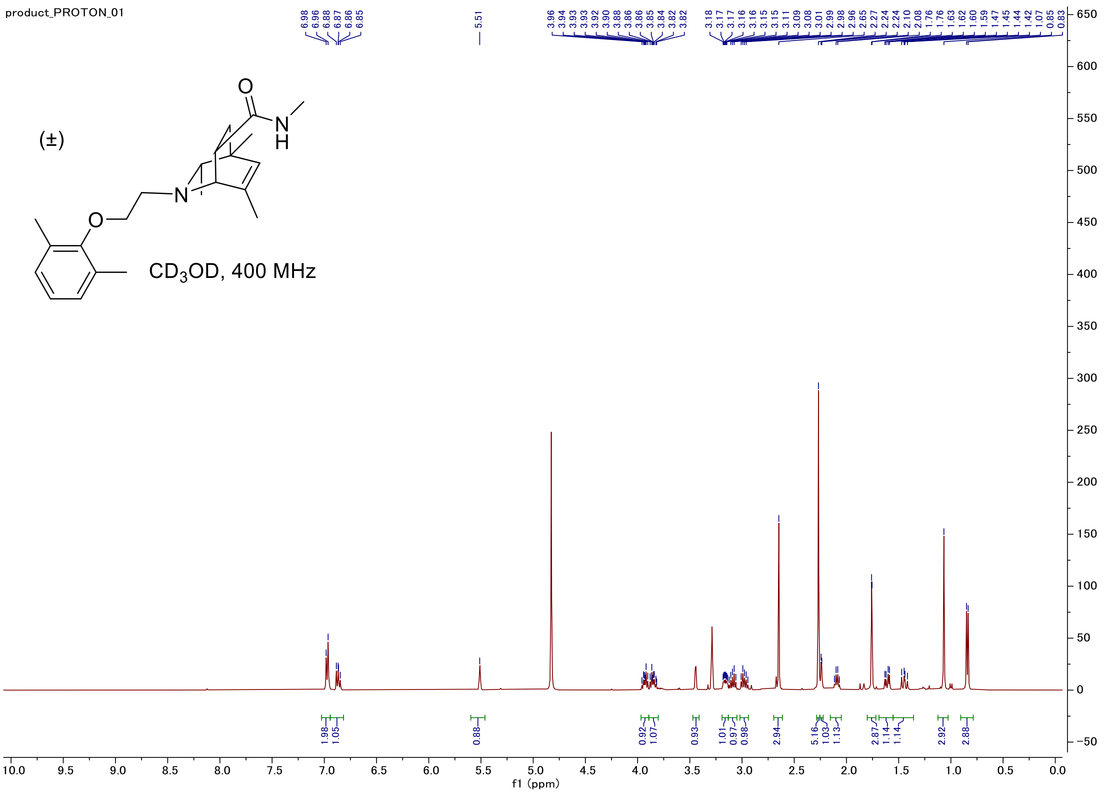


#### Carbon

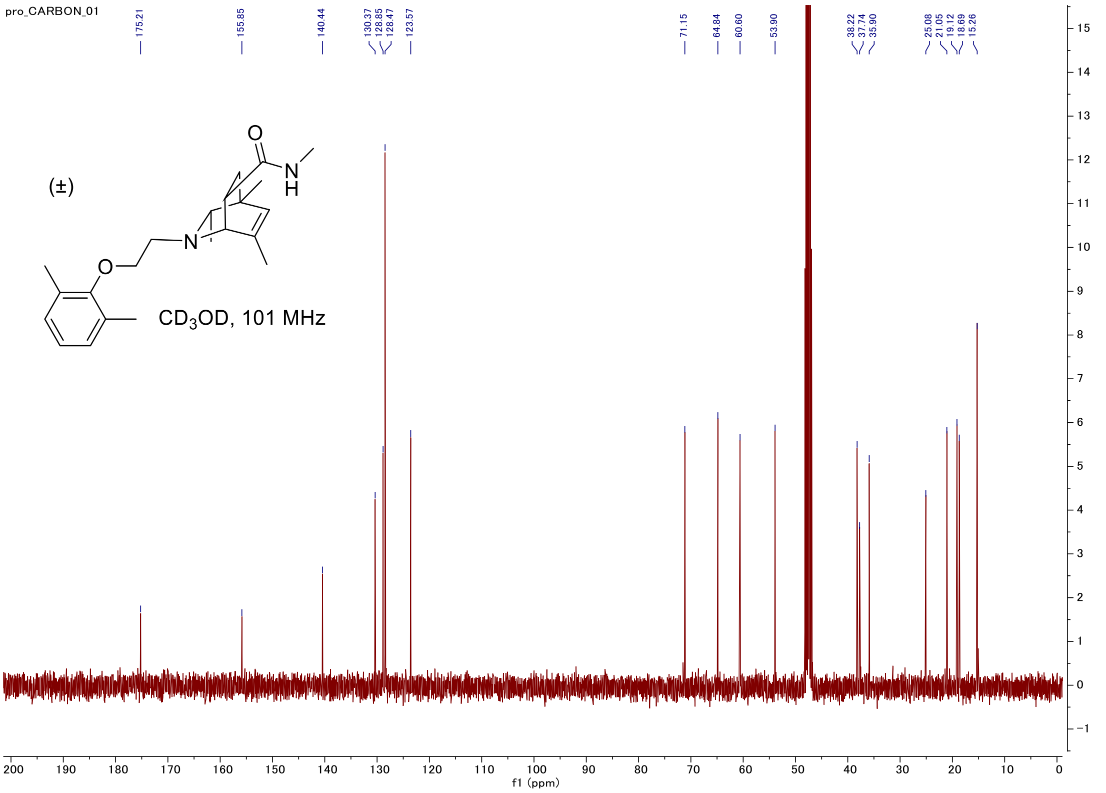

#### C-H COSY

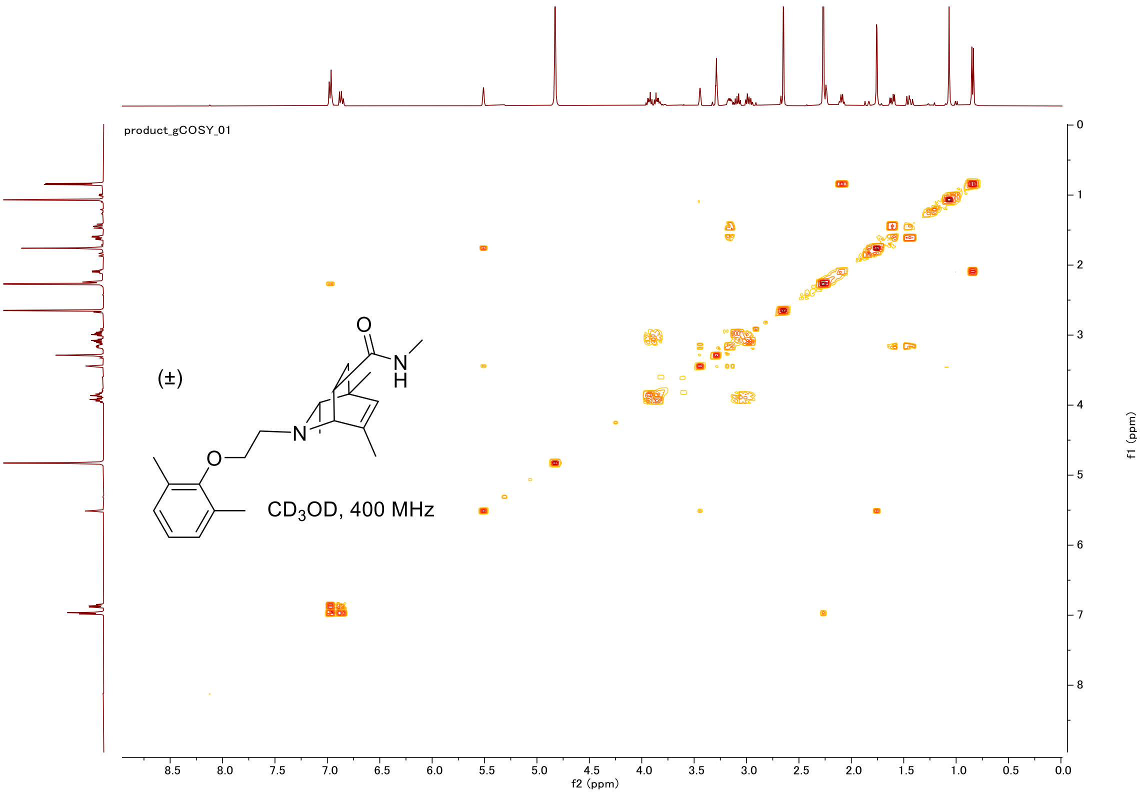

product_gCOSY_01
(±)
O
N
H
N
O
CD3OD, 400 MHz
f2 (ppm)
f1 (ppm)


#### HSQC

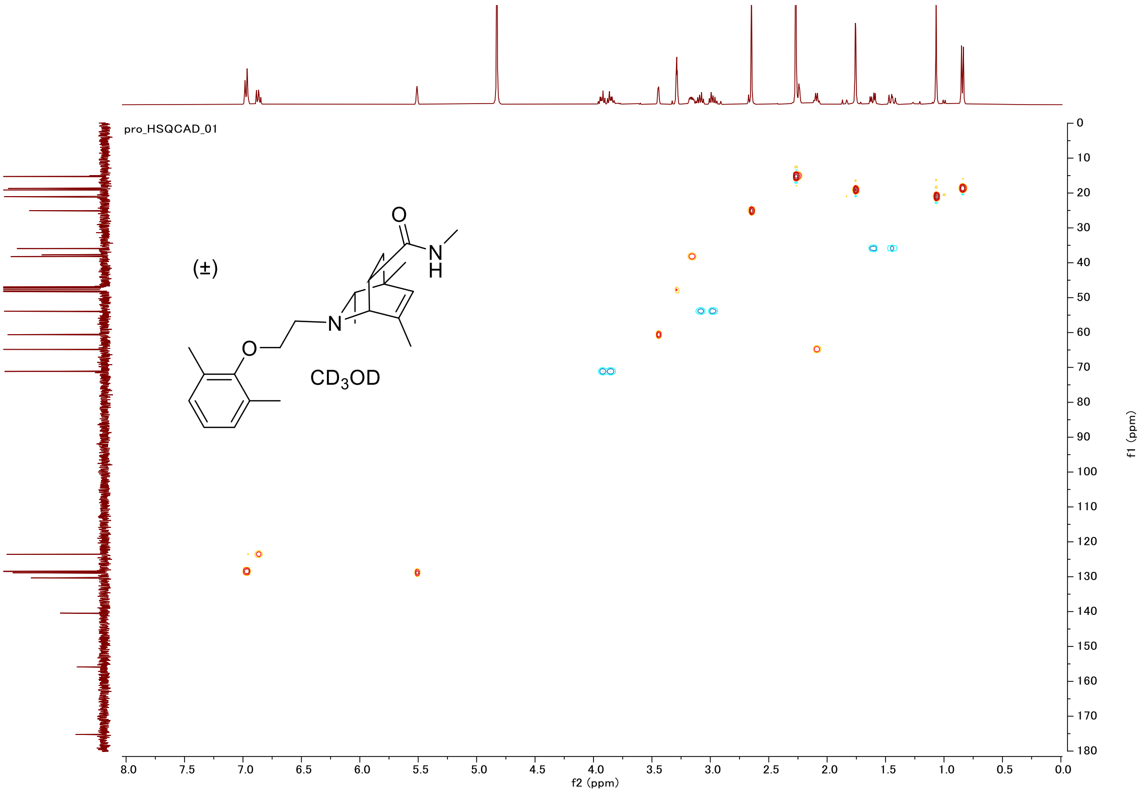

pro_HSQCAD_01
(±)
O
N
H
N
O
CD3OD
f2 (ppm)
f1 (ppm)

#### Proton (#64)

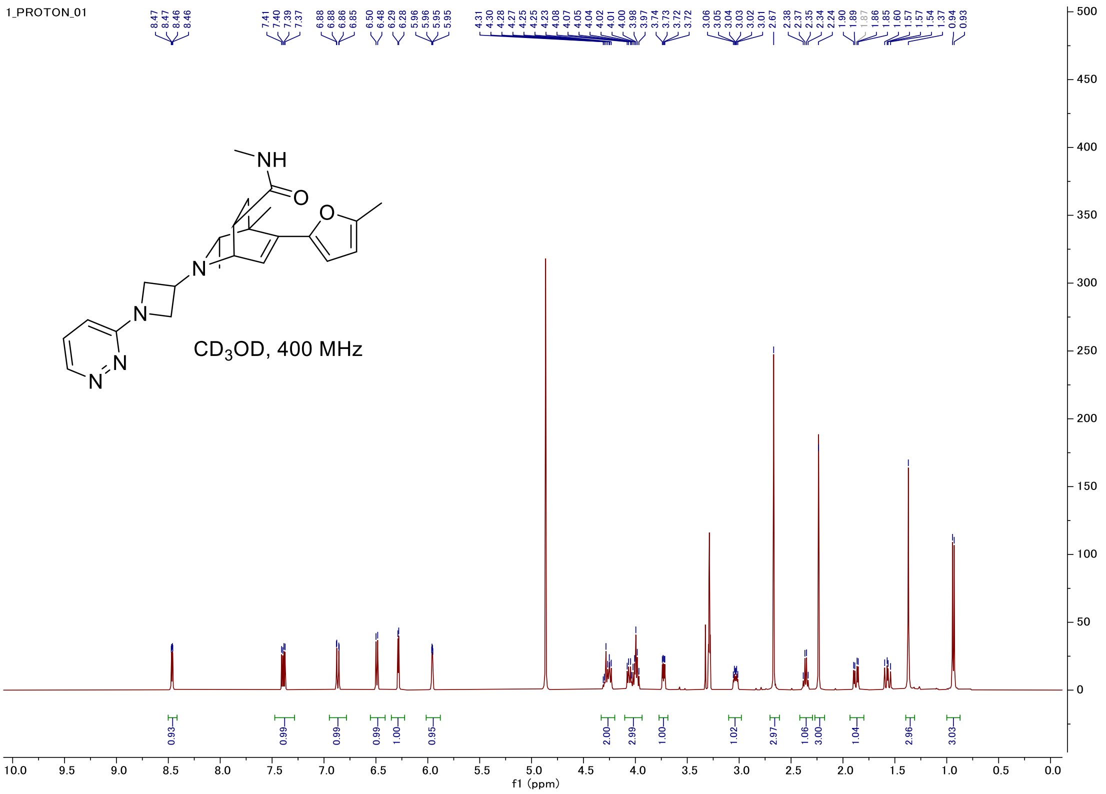


#### Carbon

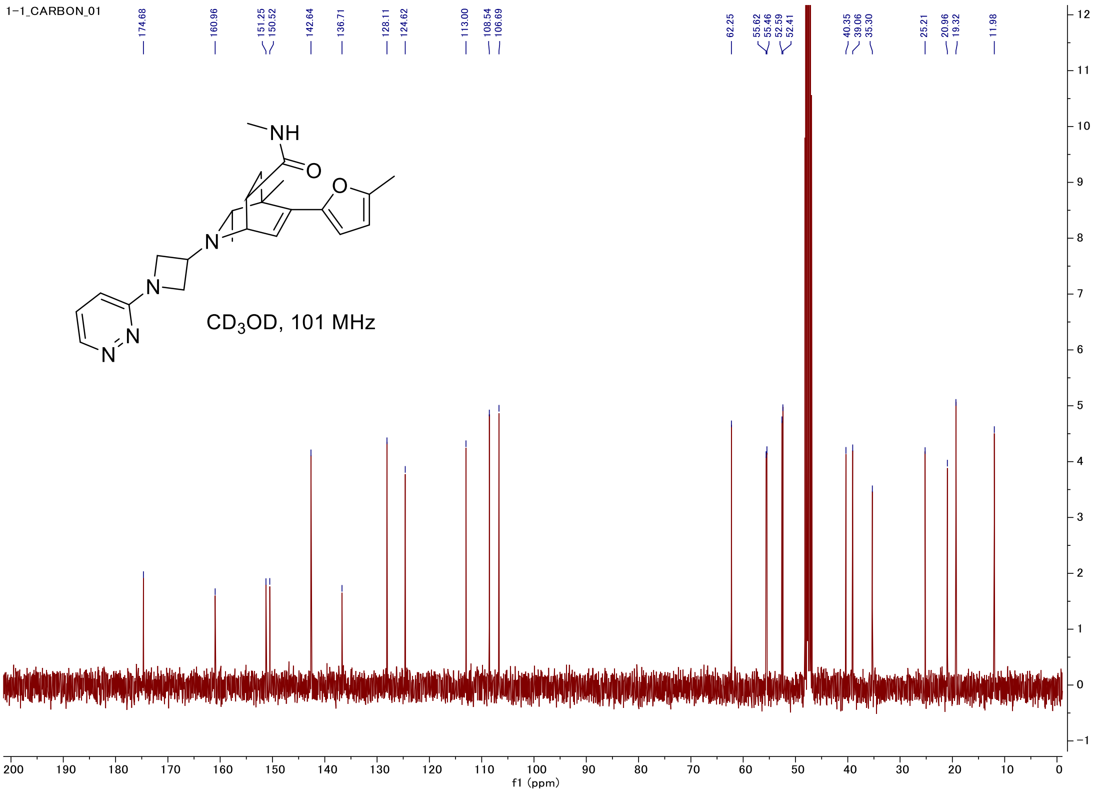

#### C-H COSY

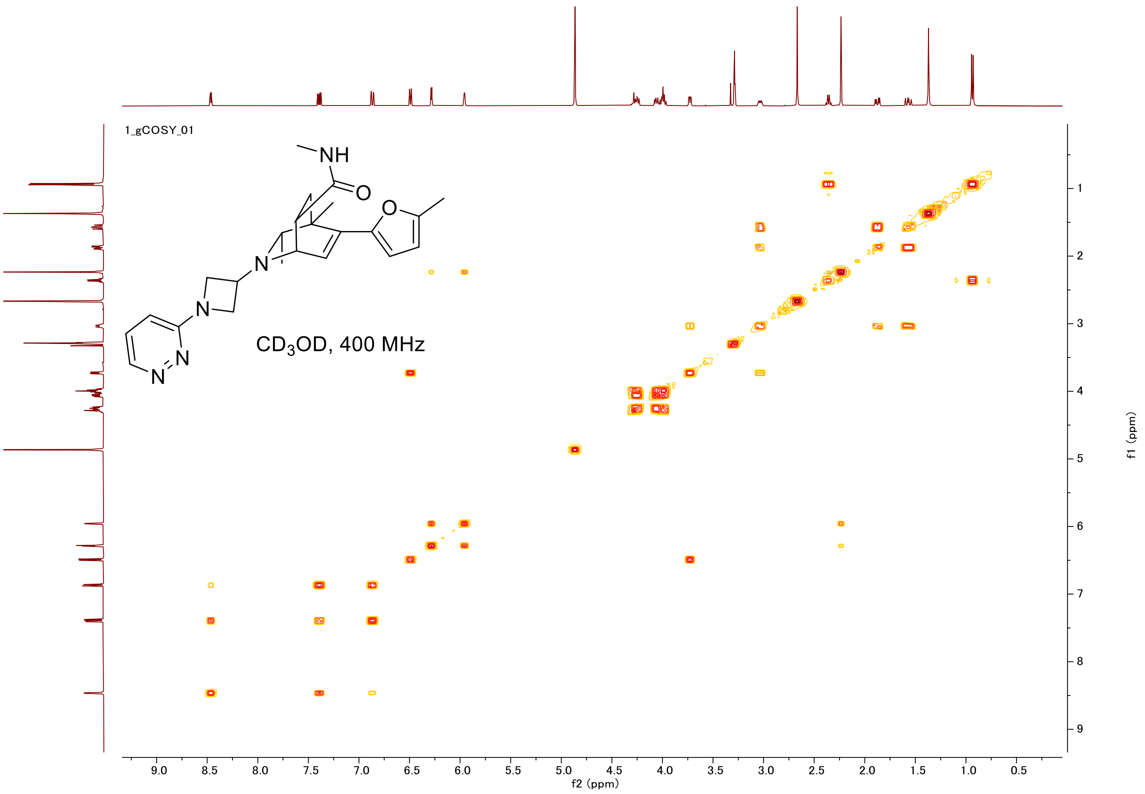
1_gCOSY_01
NH
O
O
N
N
N
N
CD3OD, 400 MHz
f2 (ppm)
f1 (ppm)

#### HSQC

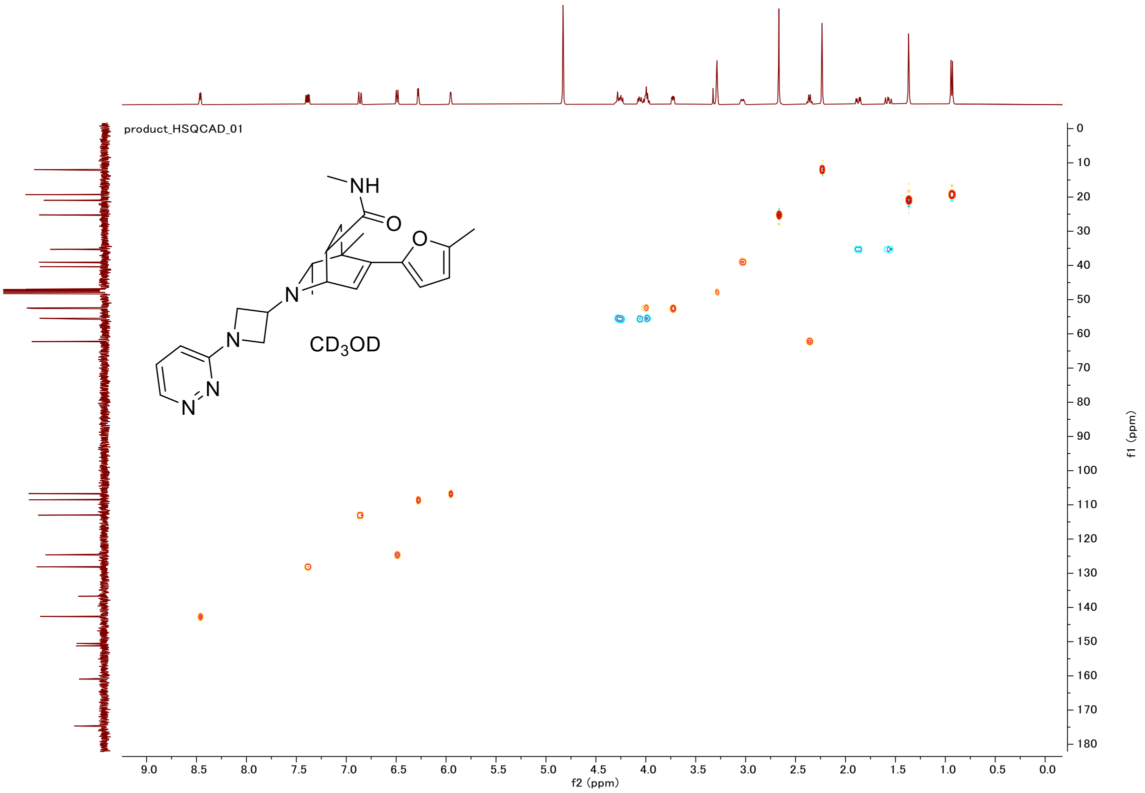
product_HSQCAD_01
NH
O
O
N
N
N
N
CD3OD
f2 (ppm)
f1 (ppm)

#### Proton (#001)

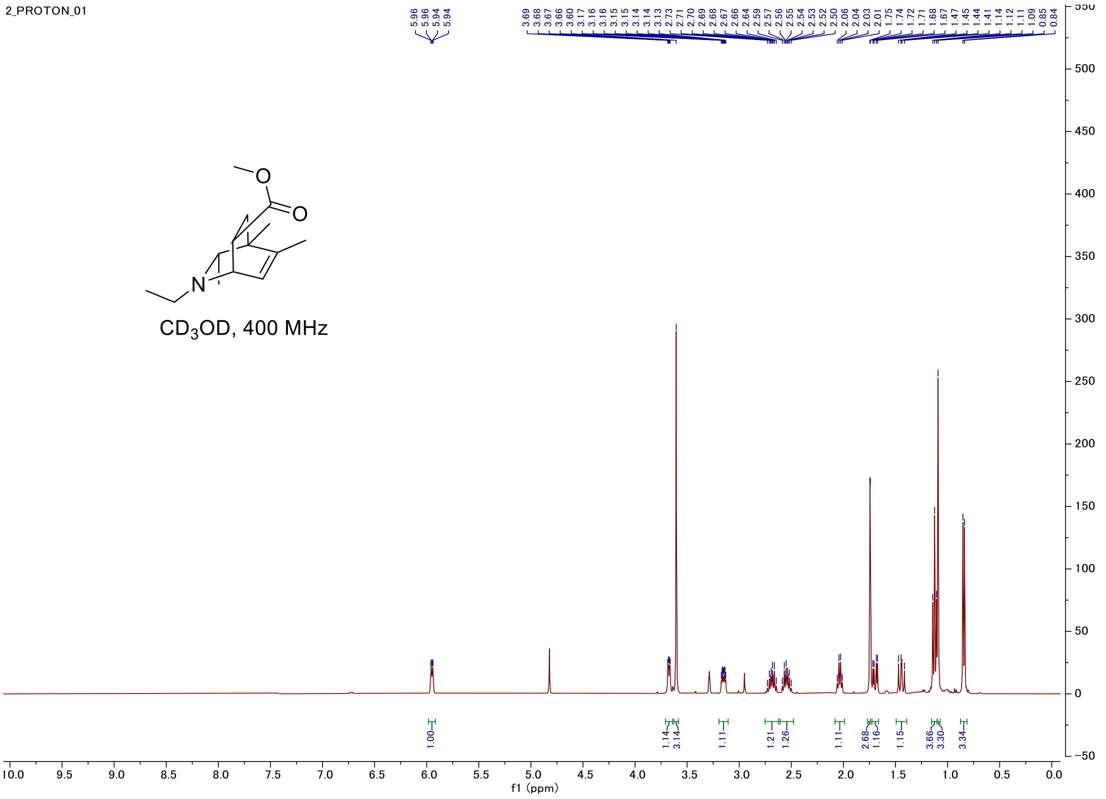


#### Carbon

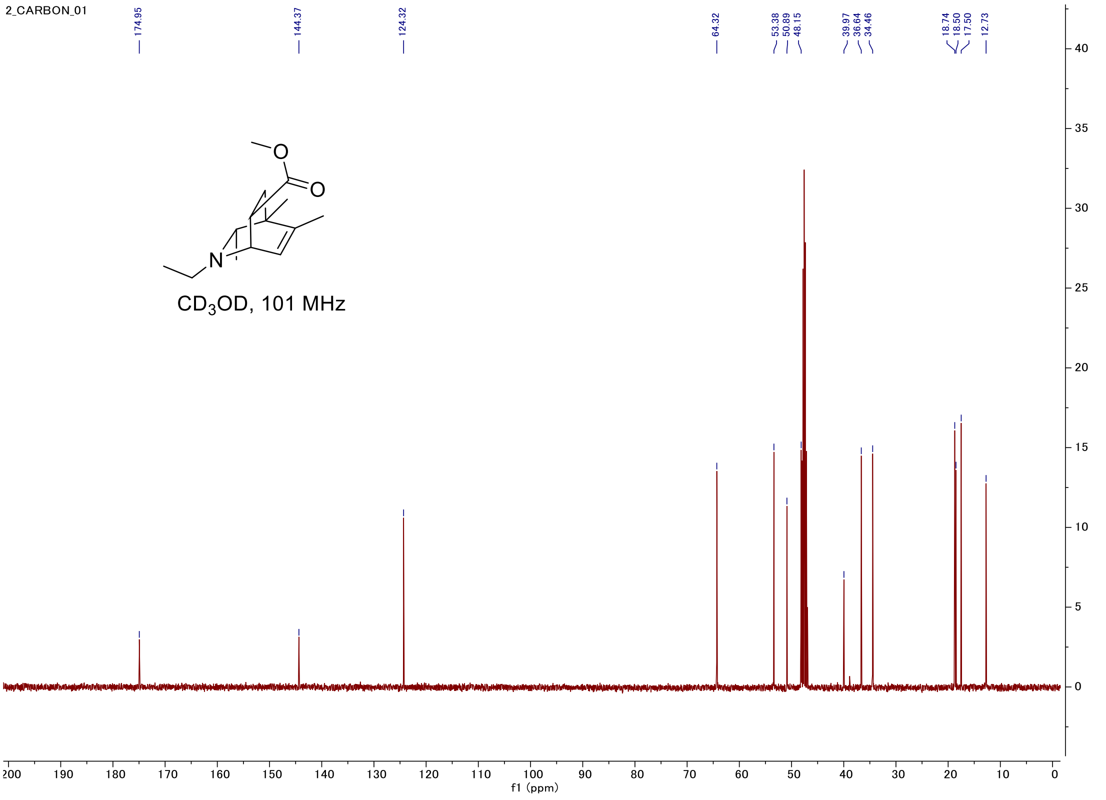

#### C-H COSY

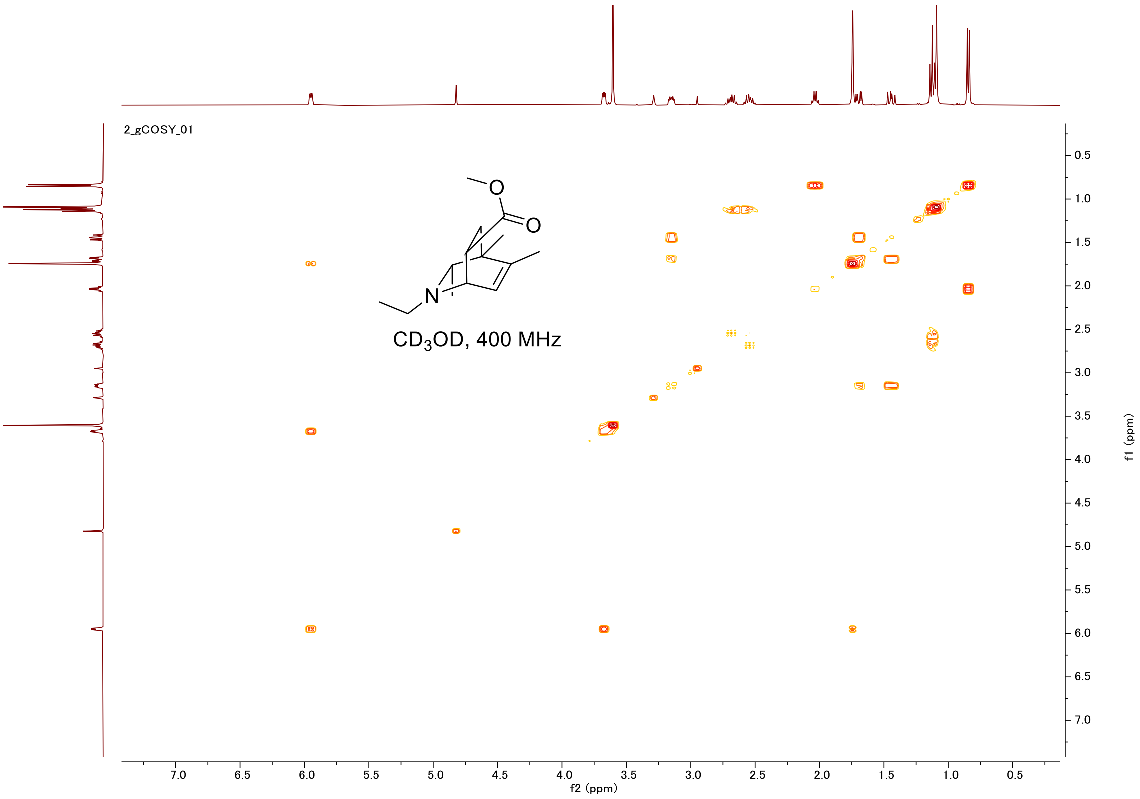
2_gCOSY_01
CD$_3$OD, 400 MHz
f2 (ppm)
f1 (ppm)

#### HSQC

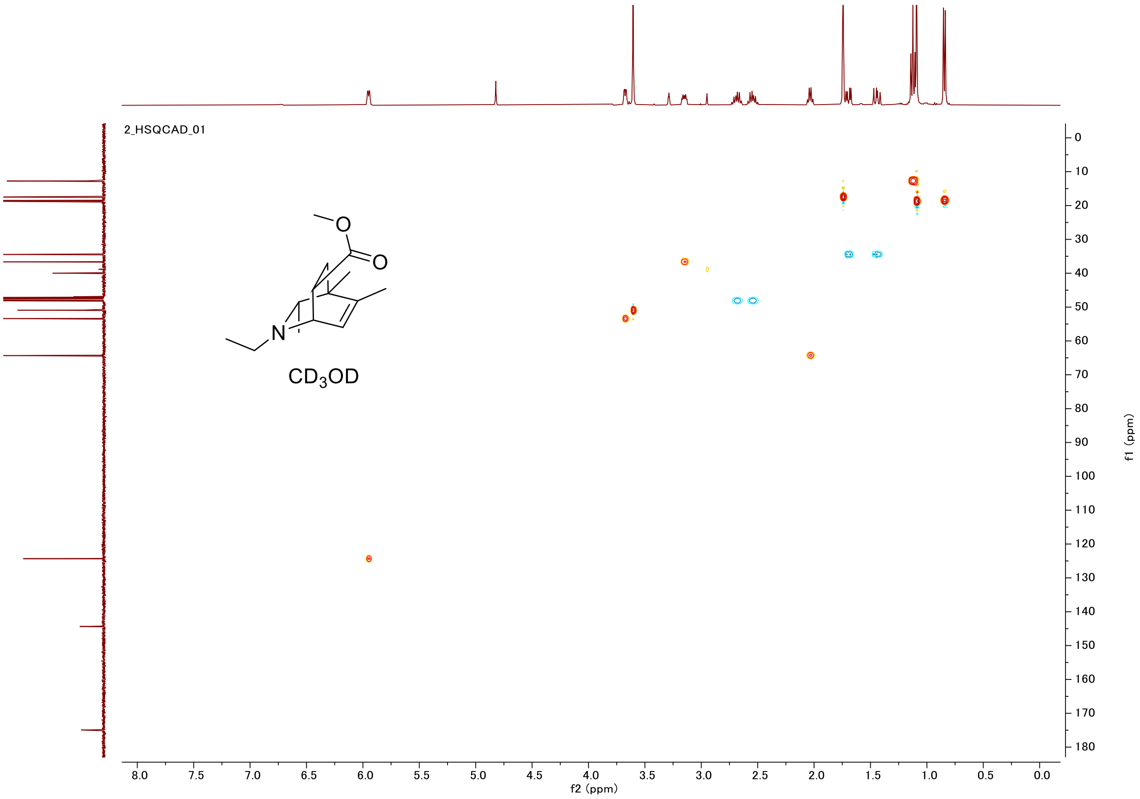
2_HSQCAD_01
CD$_3$OD
f2 (ppm)
f1 (ppm)

#### Proton (#002)

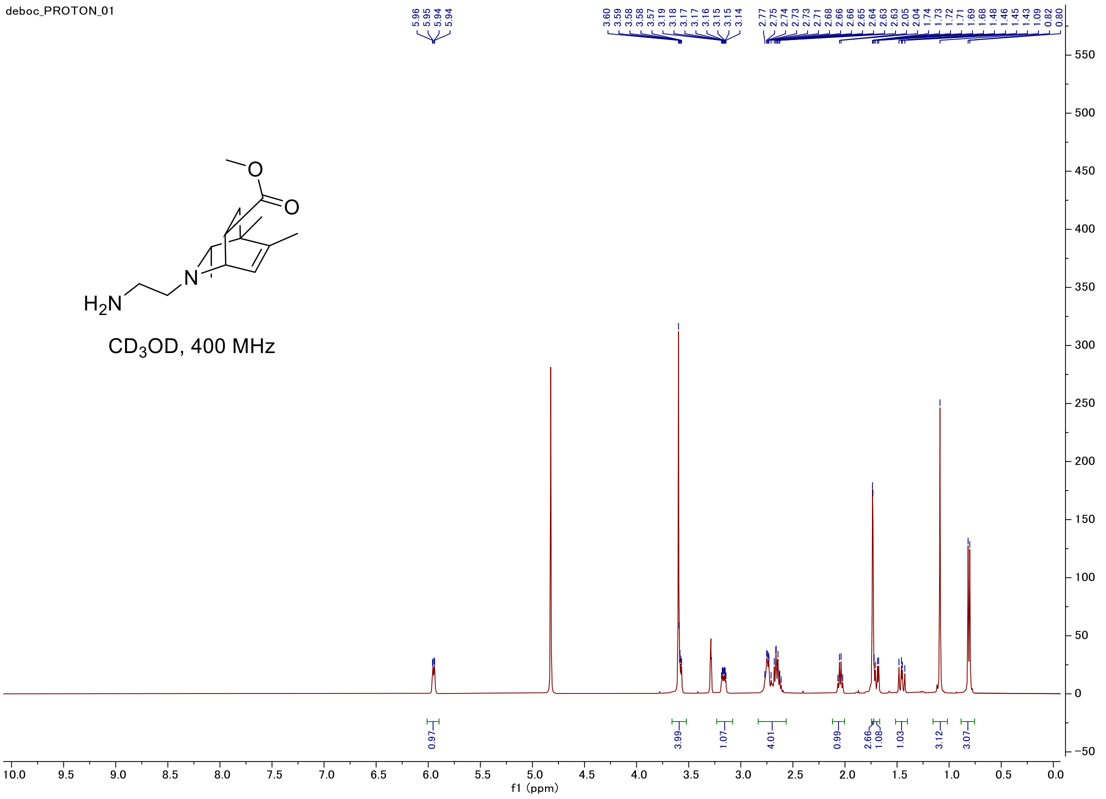


#### Carbon

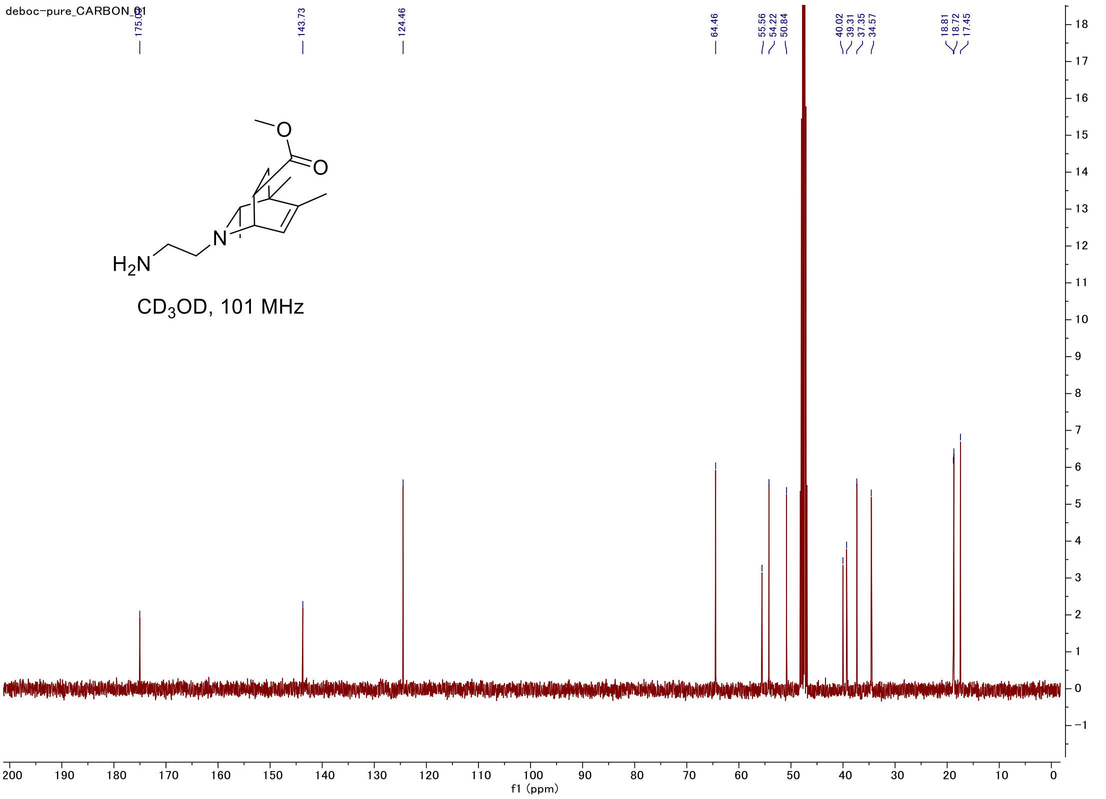

#### C-H COSY

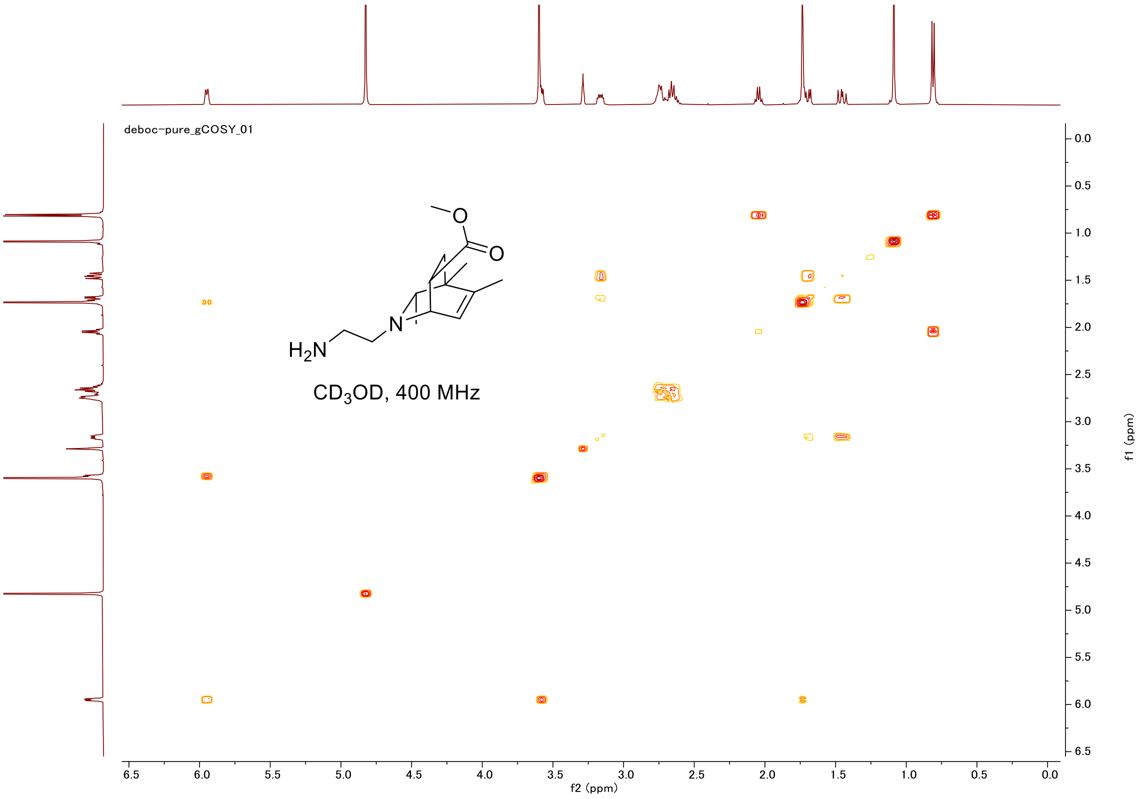

#### Proton (#003)

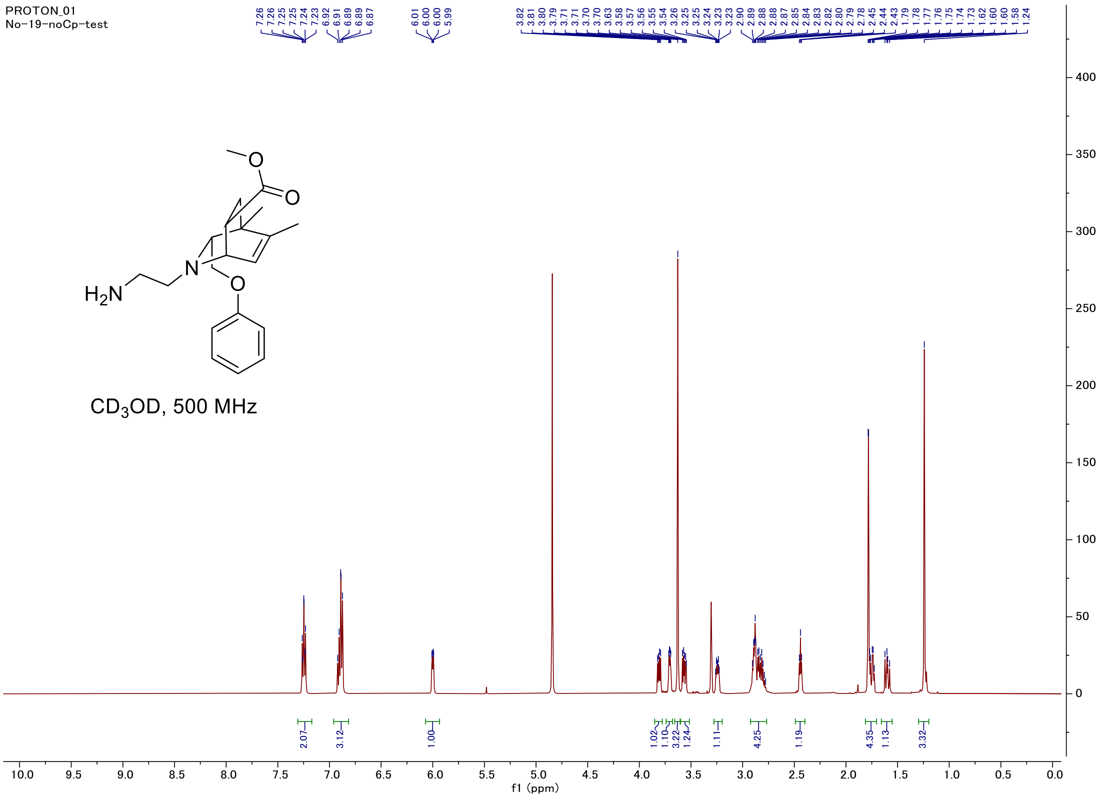


#### Carbon

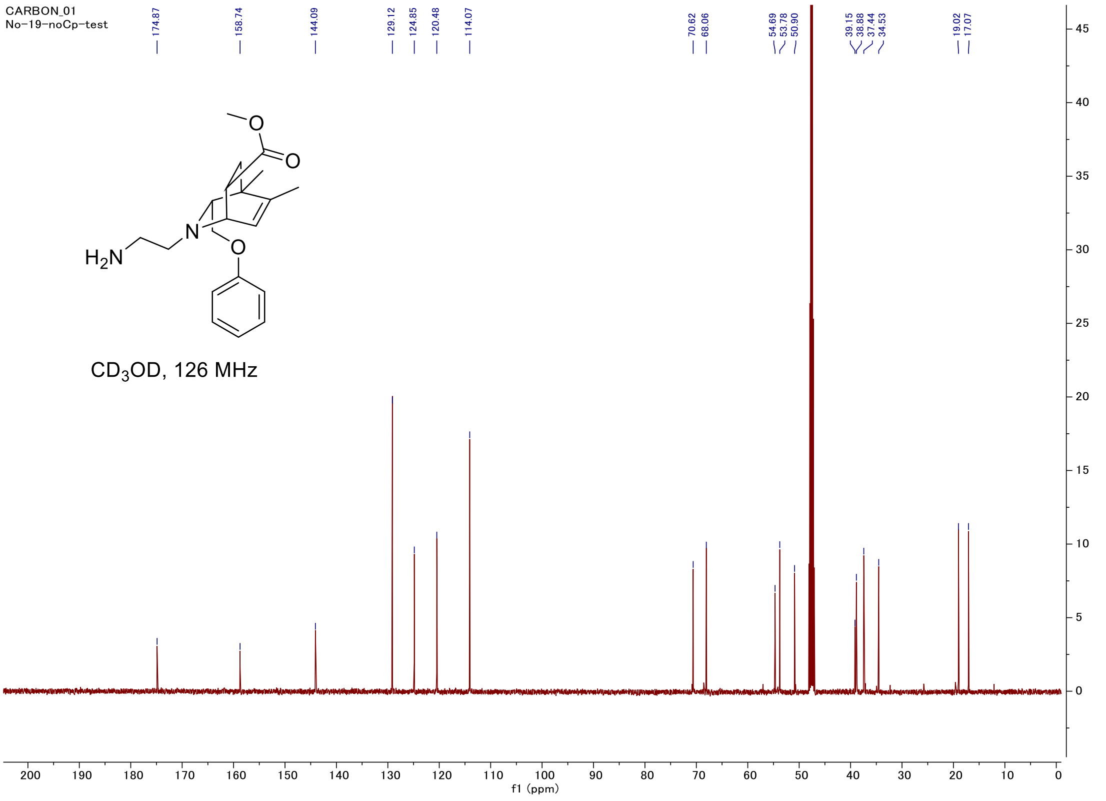

#### C-H COSY

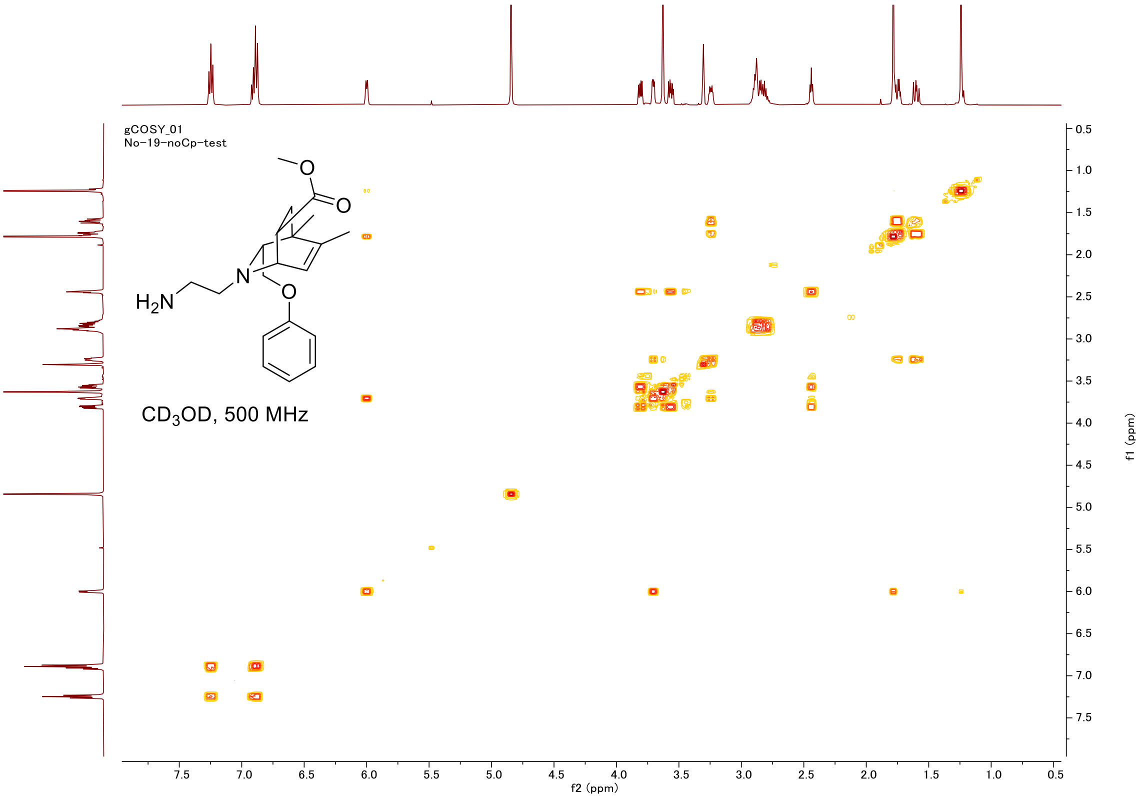
gCOSY_01
No-19-noCp-test
H2N
N
O
O
O
CD3OD, 500 MHz
f2 (ppm)
f1 (ppm)

#### HSQC

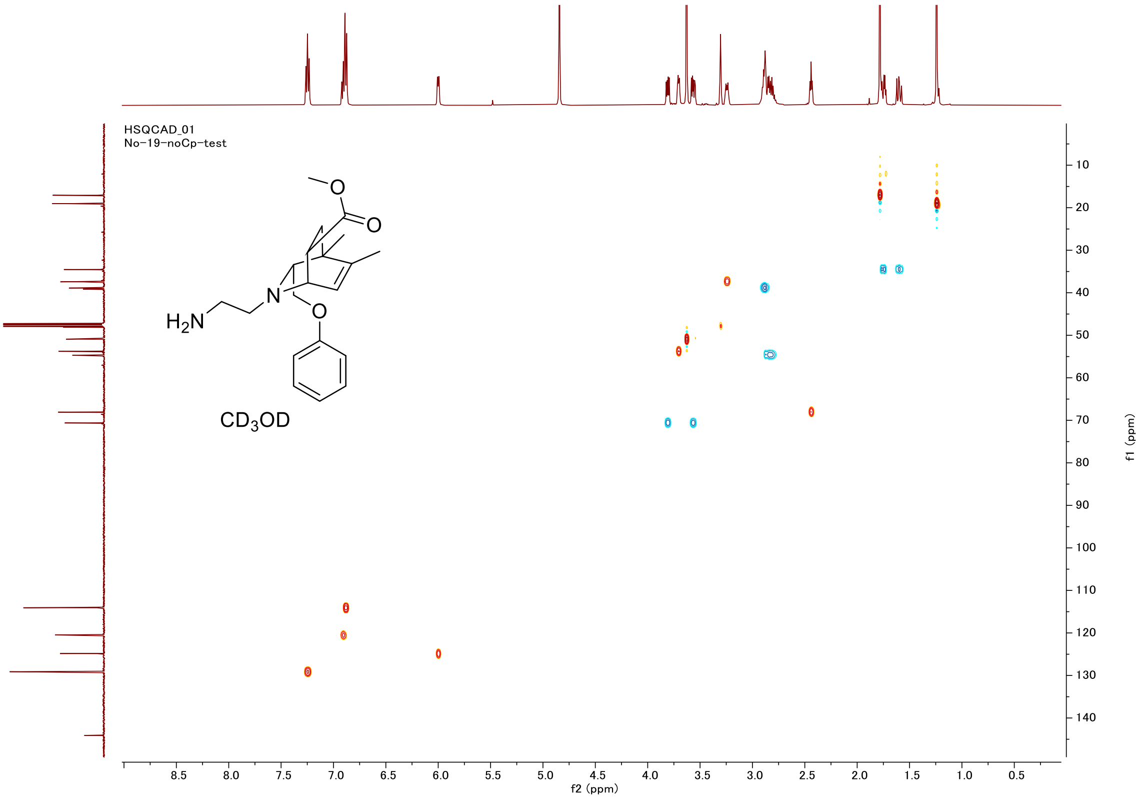
HSQCAD_01
No-19-noCp-test
H2N
N
O
O
O
CD3OD
f2 (ppm)
f1 (ppm)

#### Proton (#004)

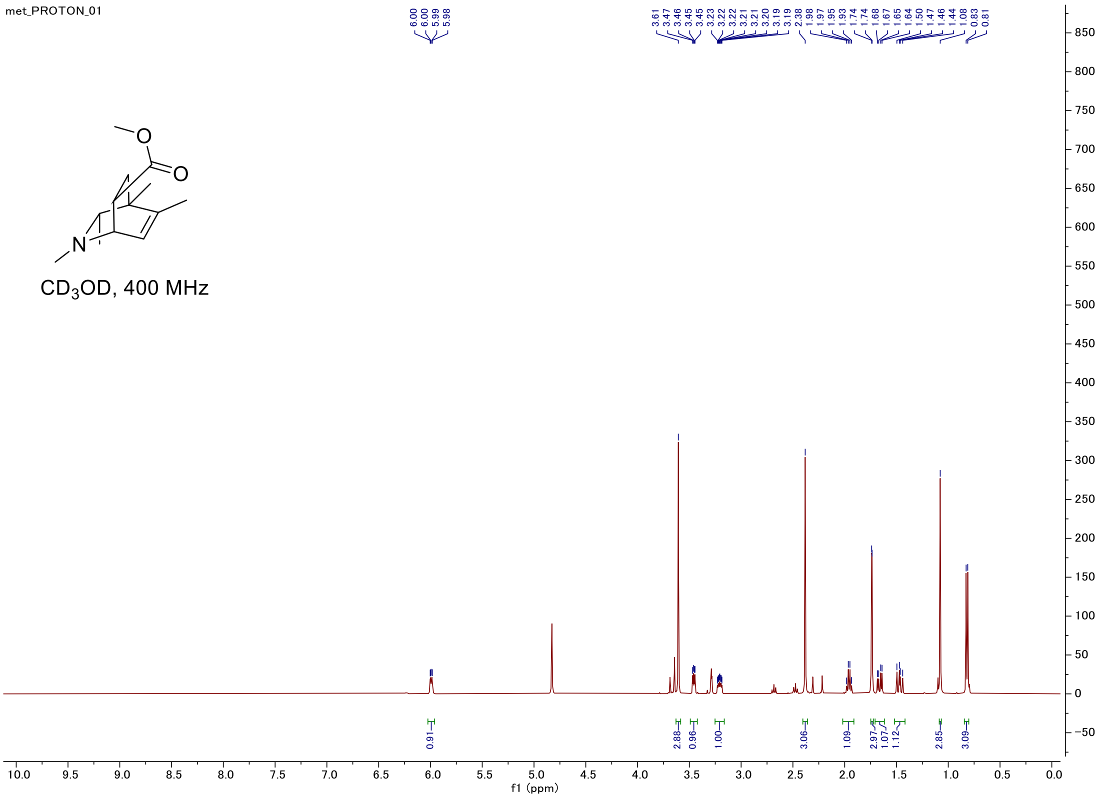


#### Carbon

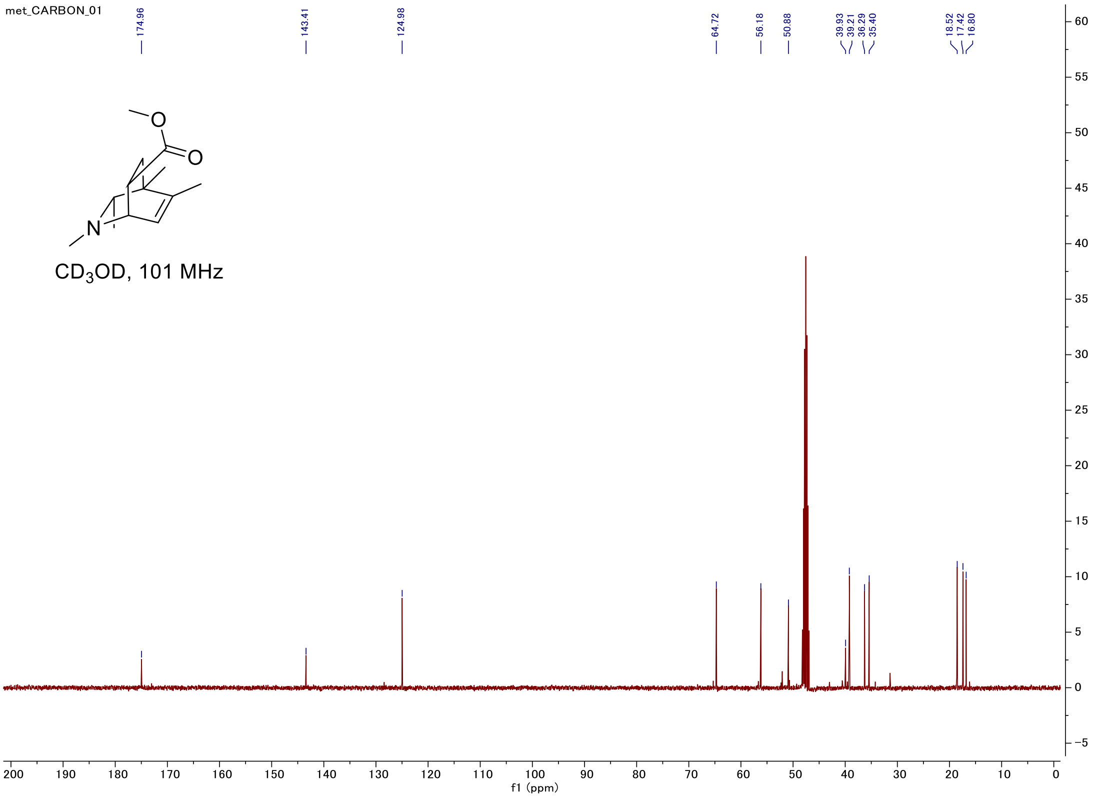

#### C-H COSY

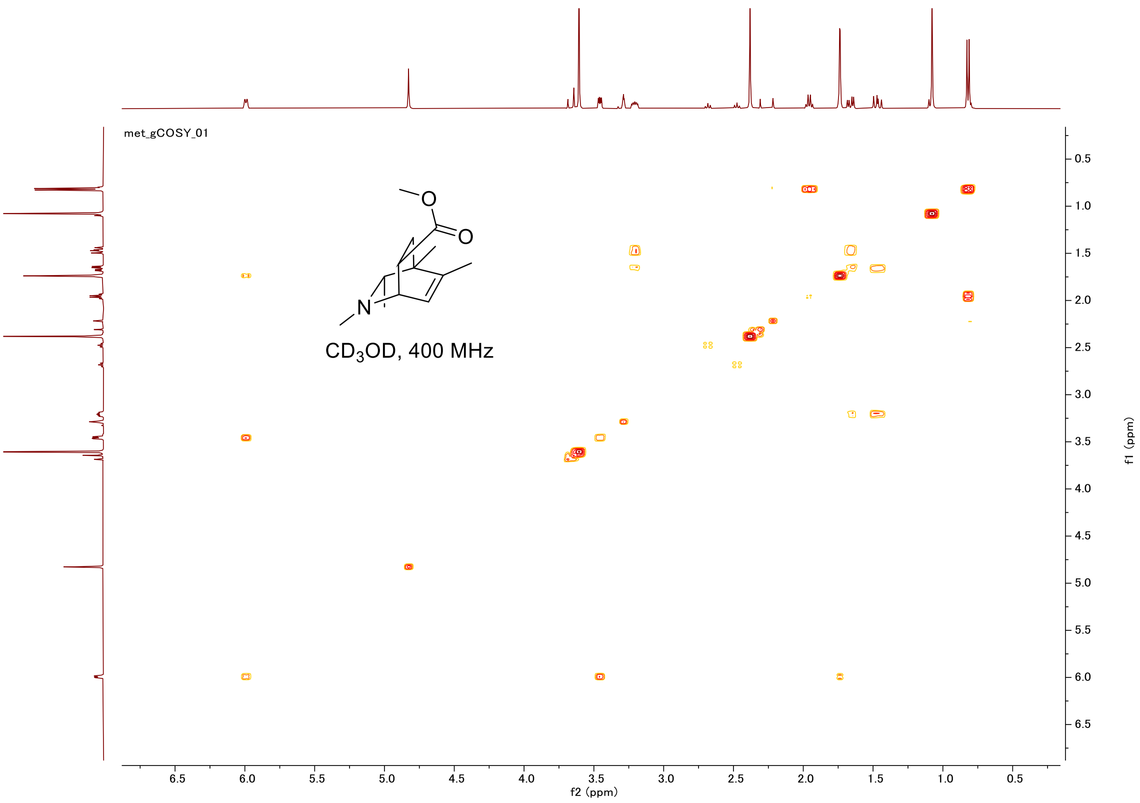
met_gCOSY_01
CD$_3$OD, 400 MHz
f2 (ppm)
f1 (ppm)

#### HSQC

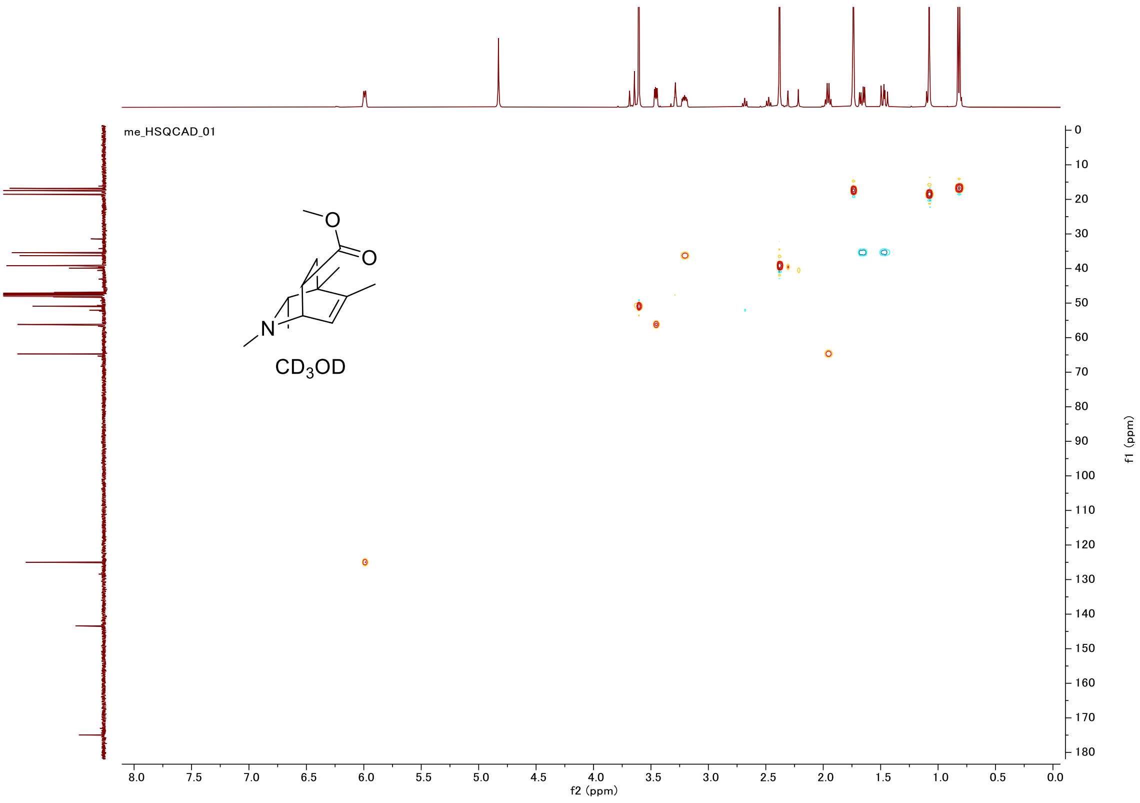
me_HSQCAD_01
CD$_3$OD
f2 (ppm)
f1 (ppm)

### Proton (#005)

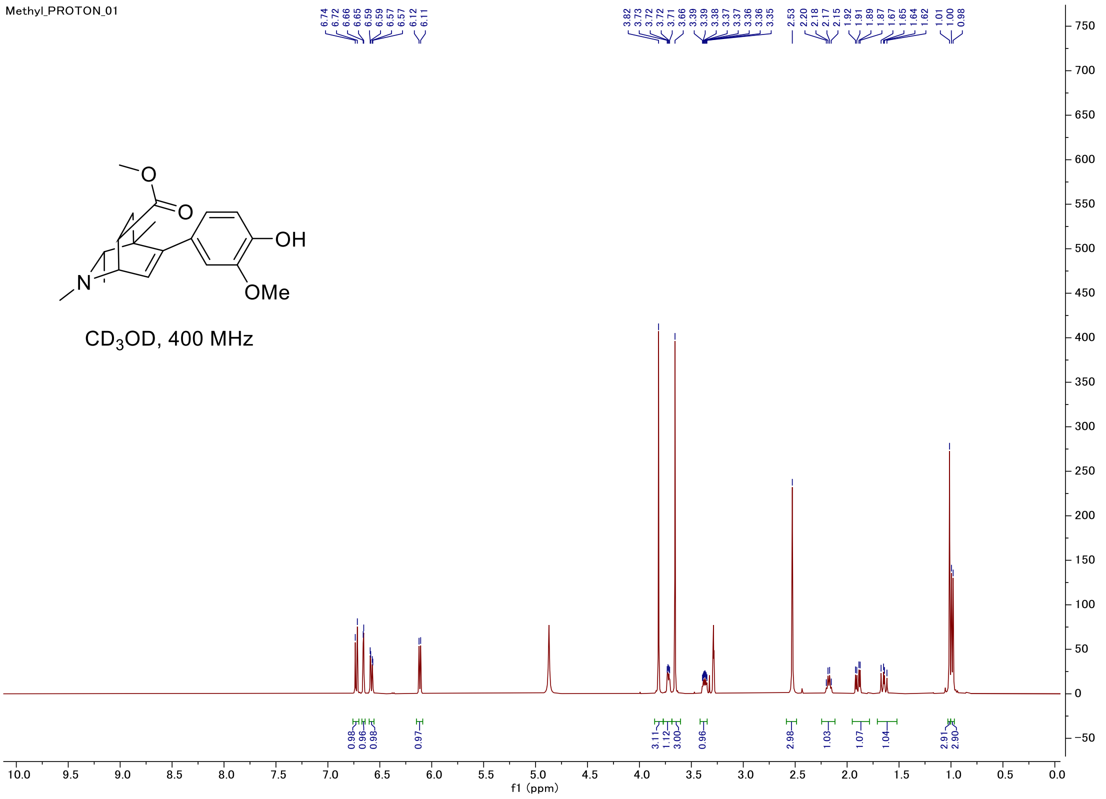


### Carbon

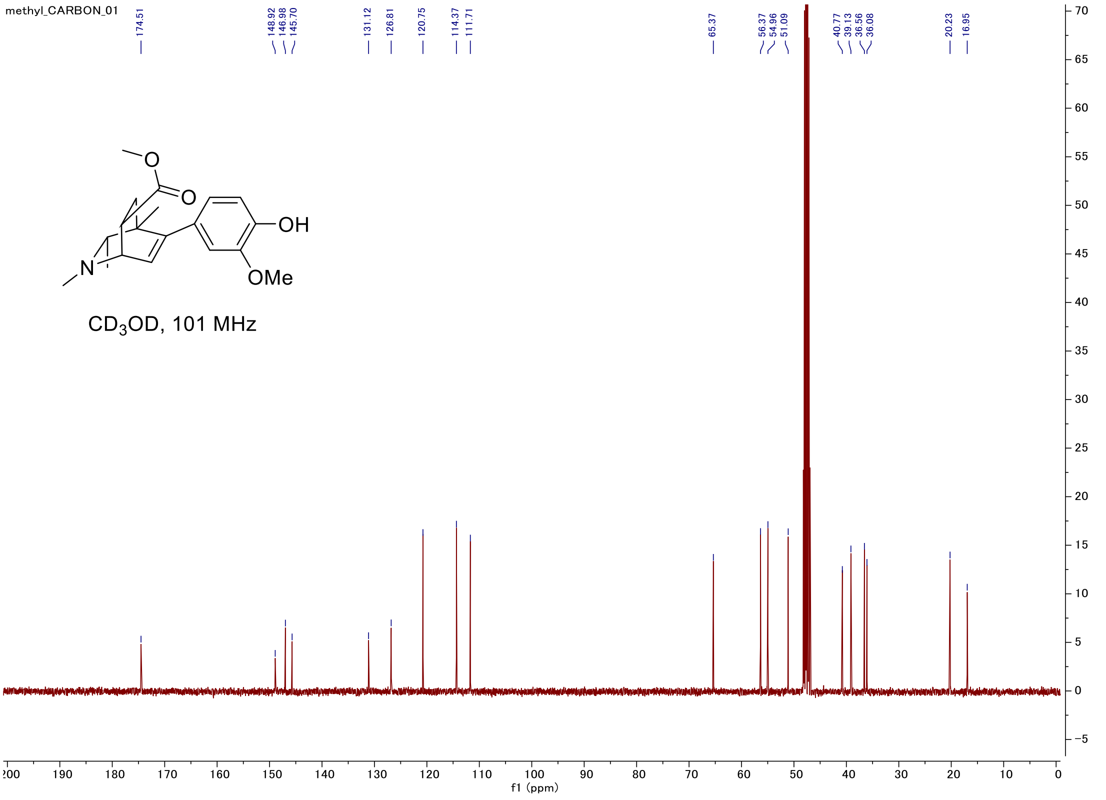

#### C-H COSY

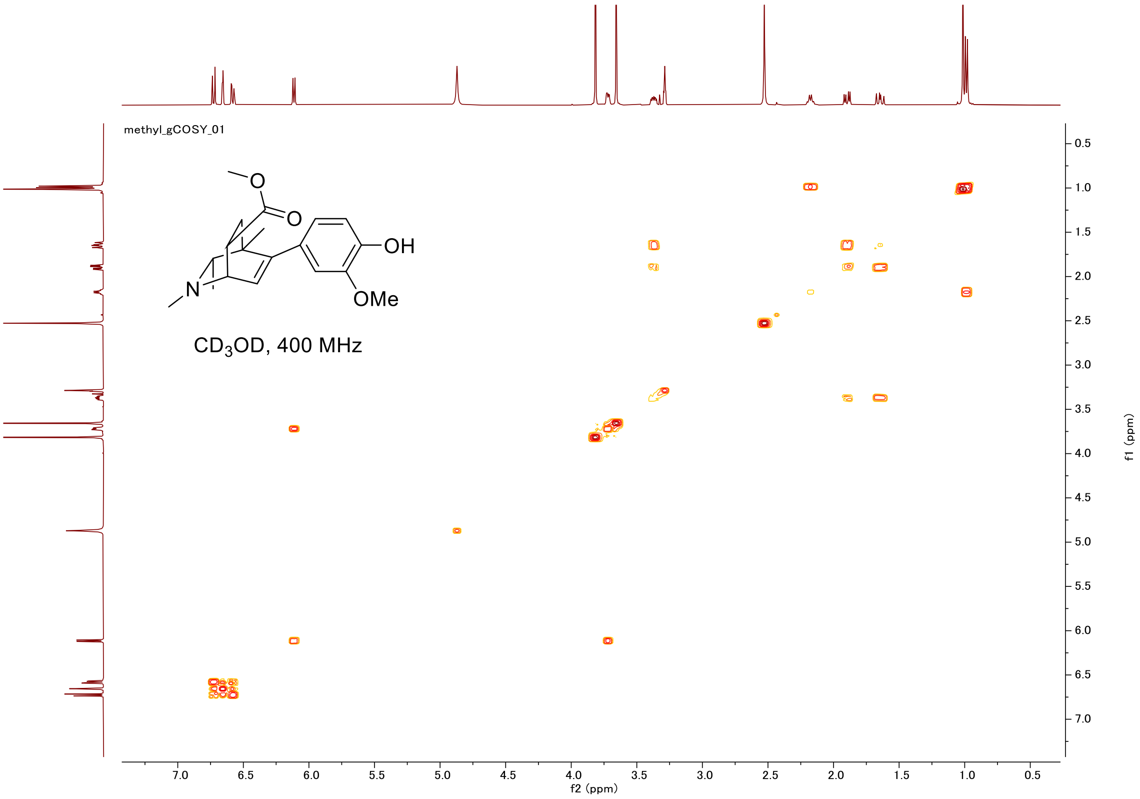
methyl_gCOSY_01
OH
OMe
N
O
CD$_3$OD, 400 MHz
f2 (ppm)
f1 (ppm)

#### HSQC

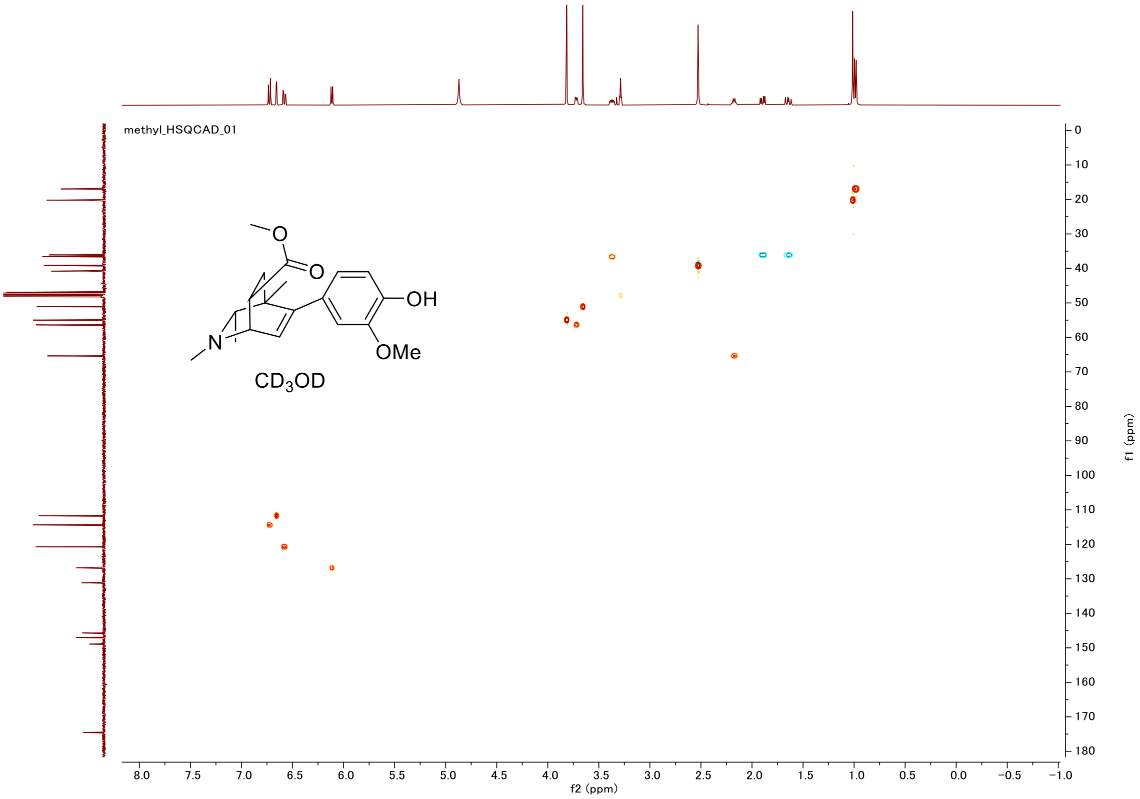
methyl_HSQCAD_01
OH
OMe
N
O
CD$_3$OD
f2 (ppm)
f1 (ppm)

### Proton (#006)

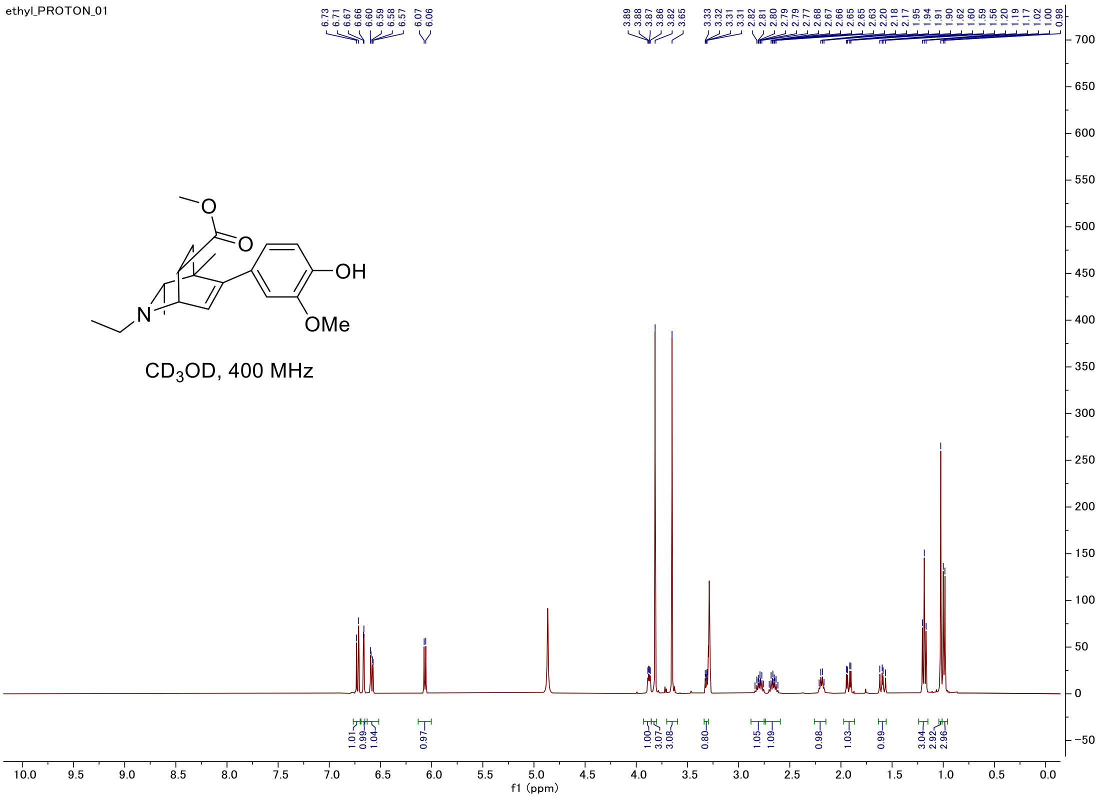


### Carbon

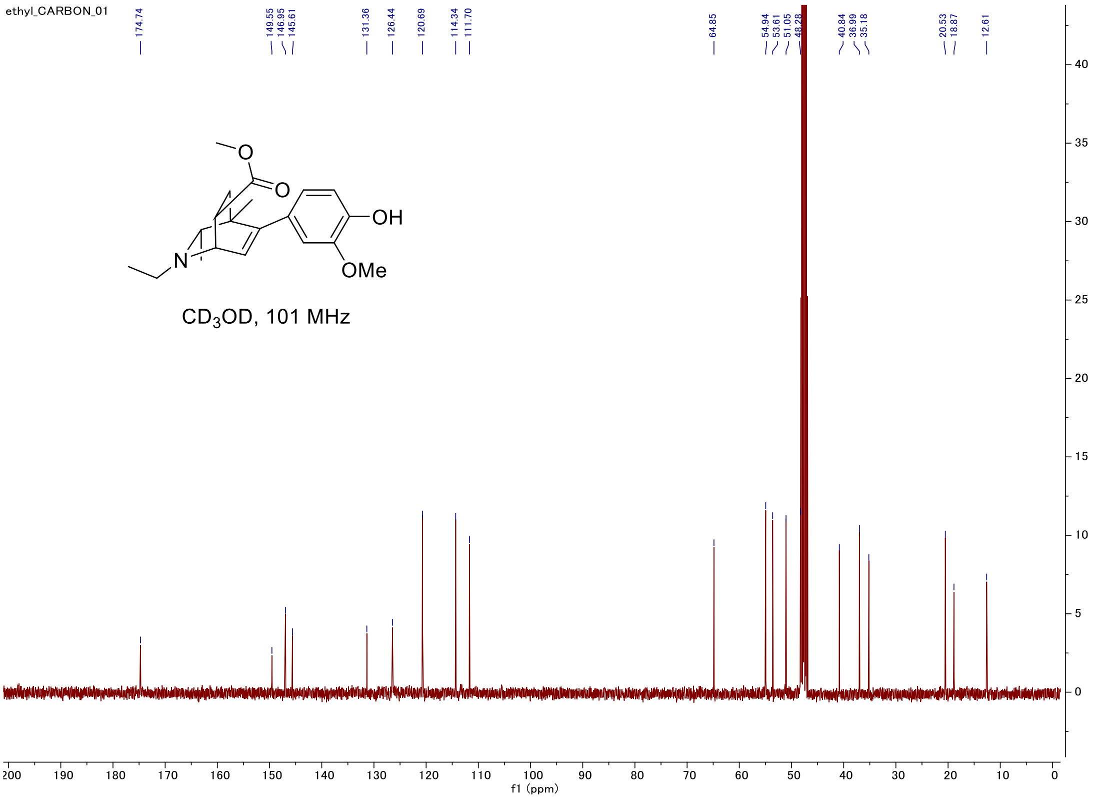

#### C-H COSY

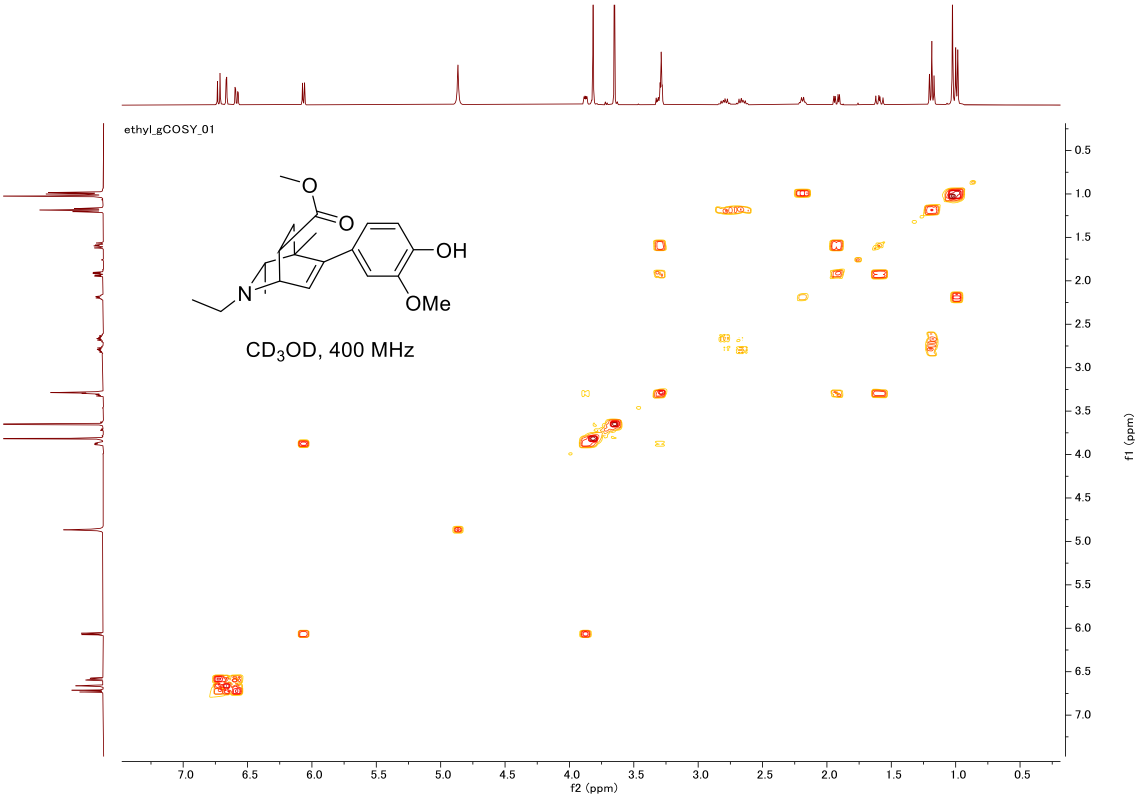
ethyl_gCOSY_01
OH
OMe
N
O
CD$_3$OD, 400 MHz
f2 (ppm)
f1 (ppm)

#### HSQC

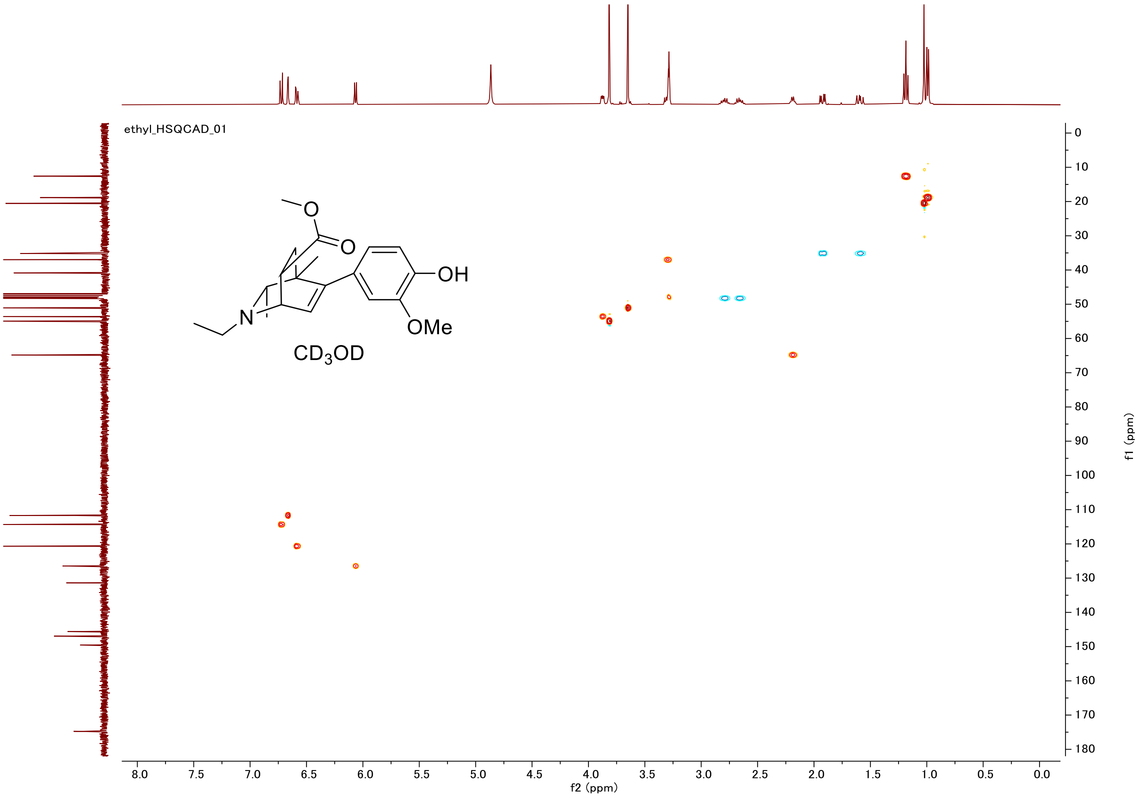
ethyl_HSQCAD_01
OH
OMe
N
O
CD$_3$OD
f2 (ppm)
f1 (ppm)

#### Proton (#007)

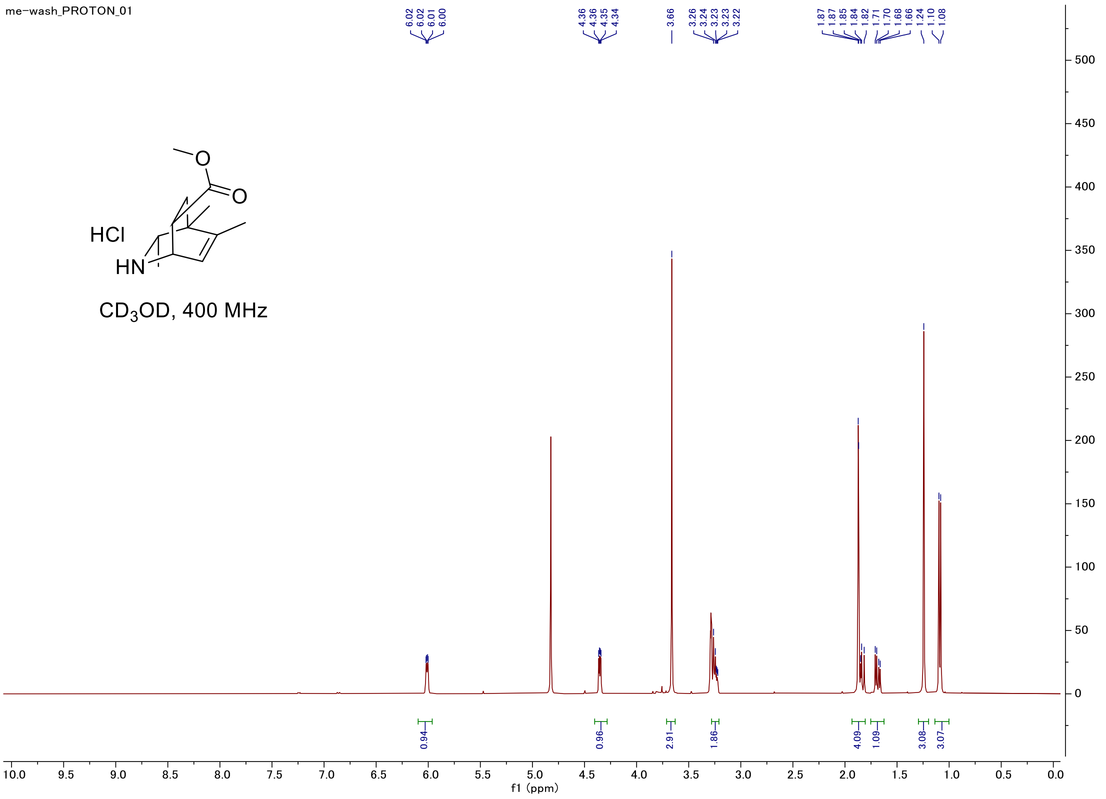


#### Carbon

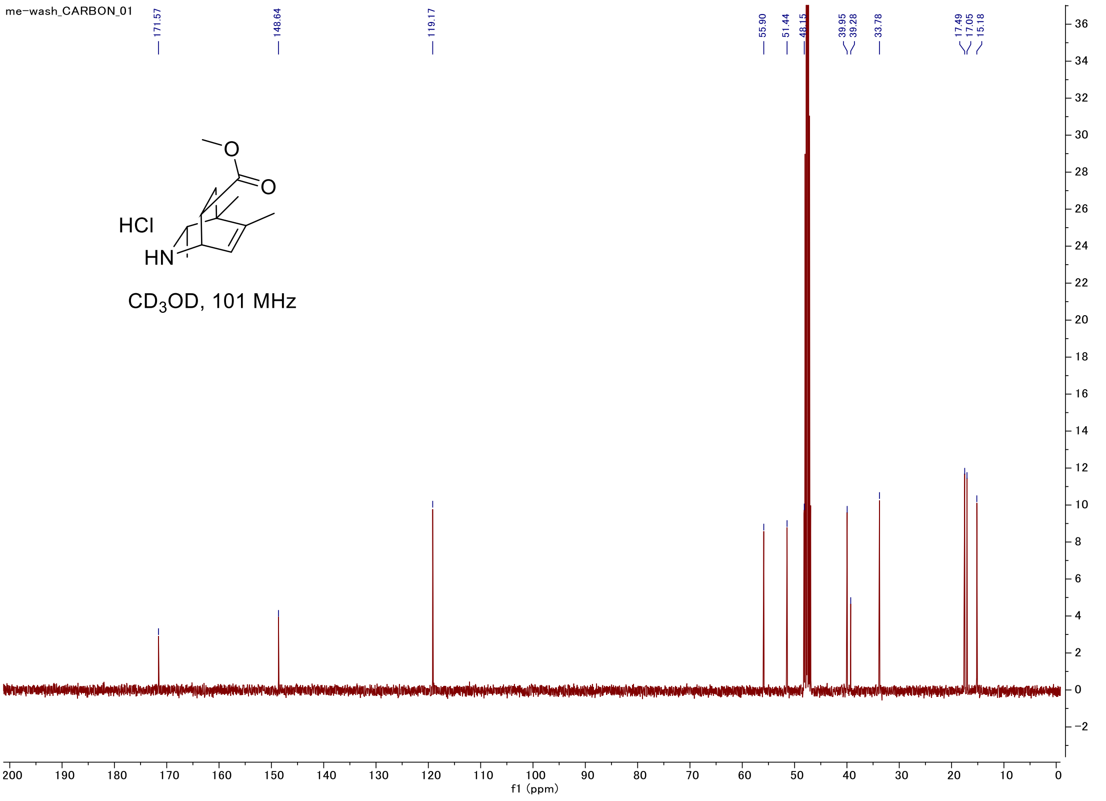

#### C-H COSY

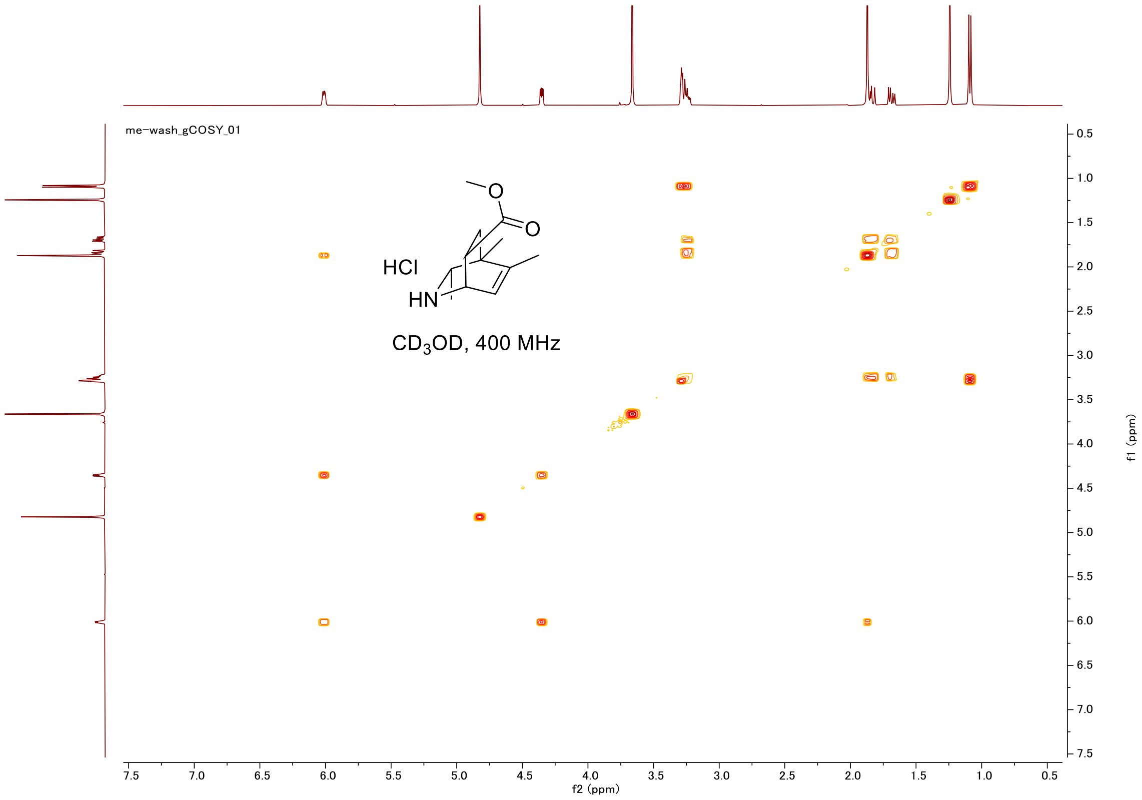
me-wash_gCOSY_01
HCl
HN
O
O
CD3OD, 400 MHz
f2 (ppm)
f1 (ppm)

#### HSQC

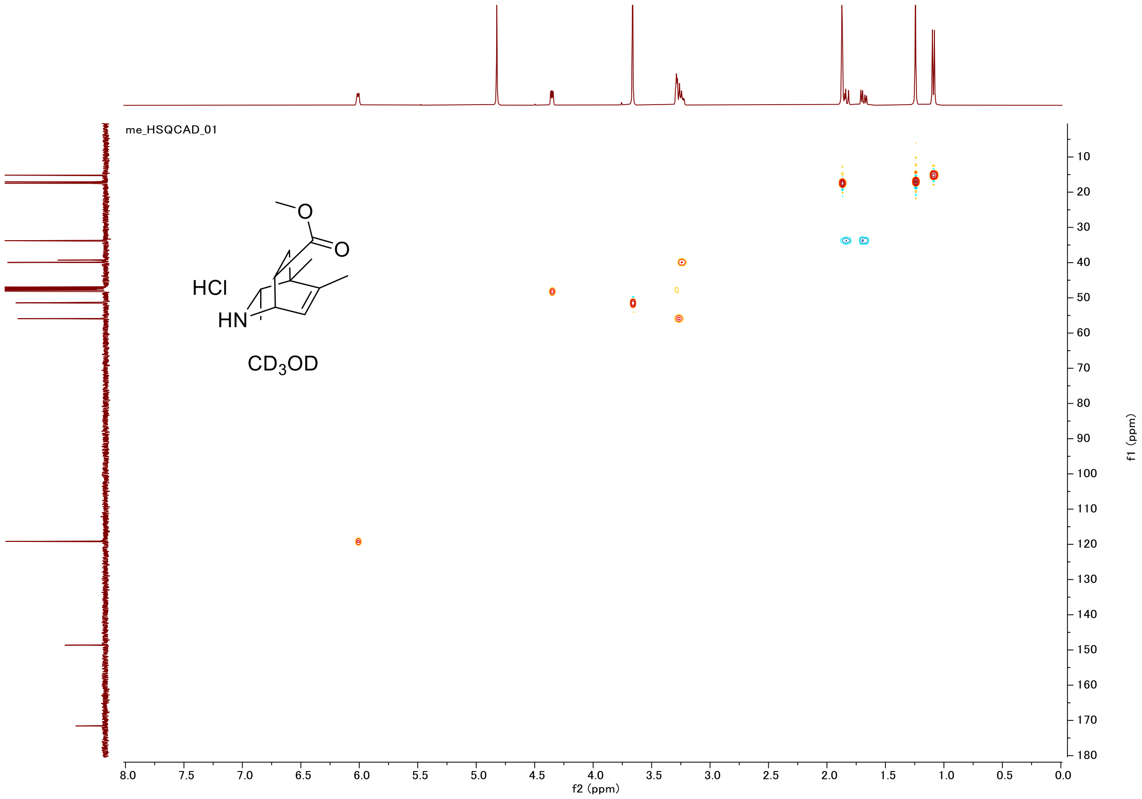
me_HSQCAD_01
HCl
HN
O
O
CD3OD
f2 (ppm)
f1 (ppm)

#### Proton (#008)

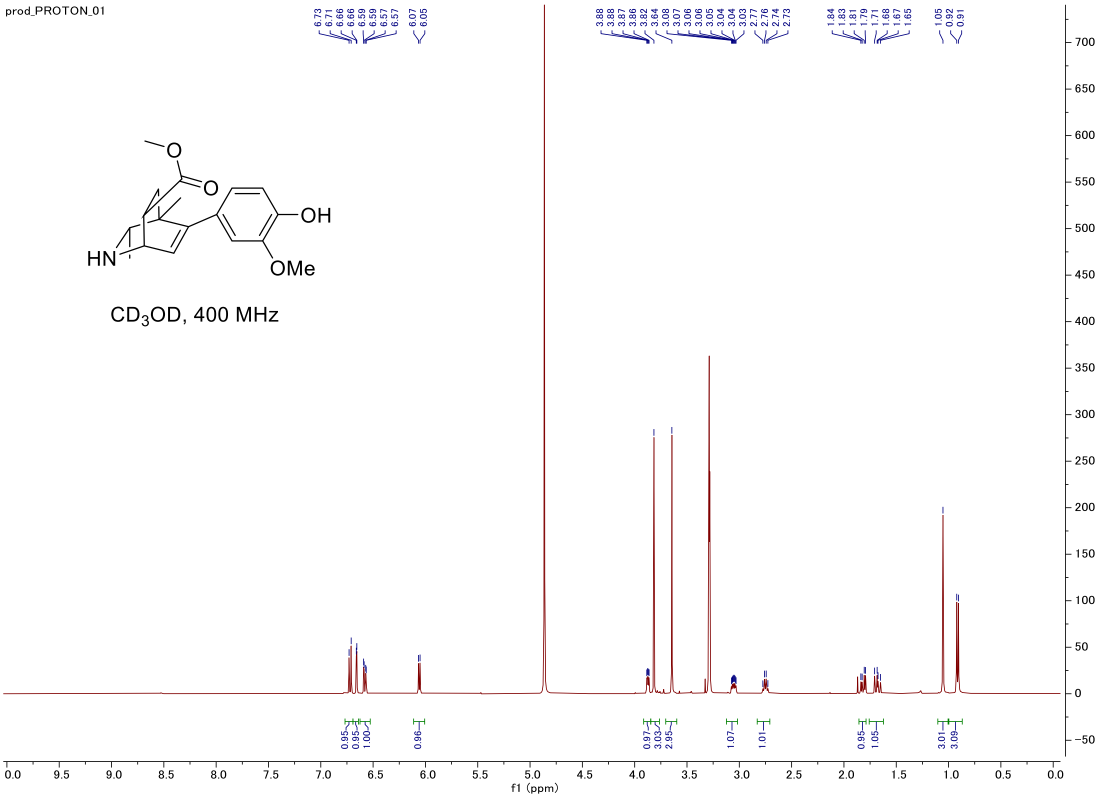


#### Carbon

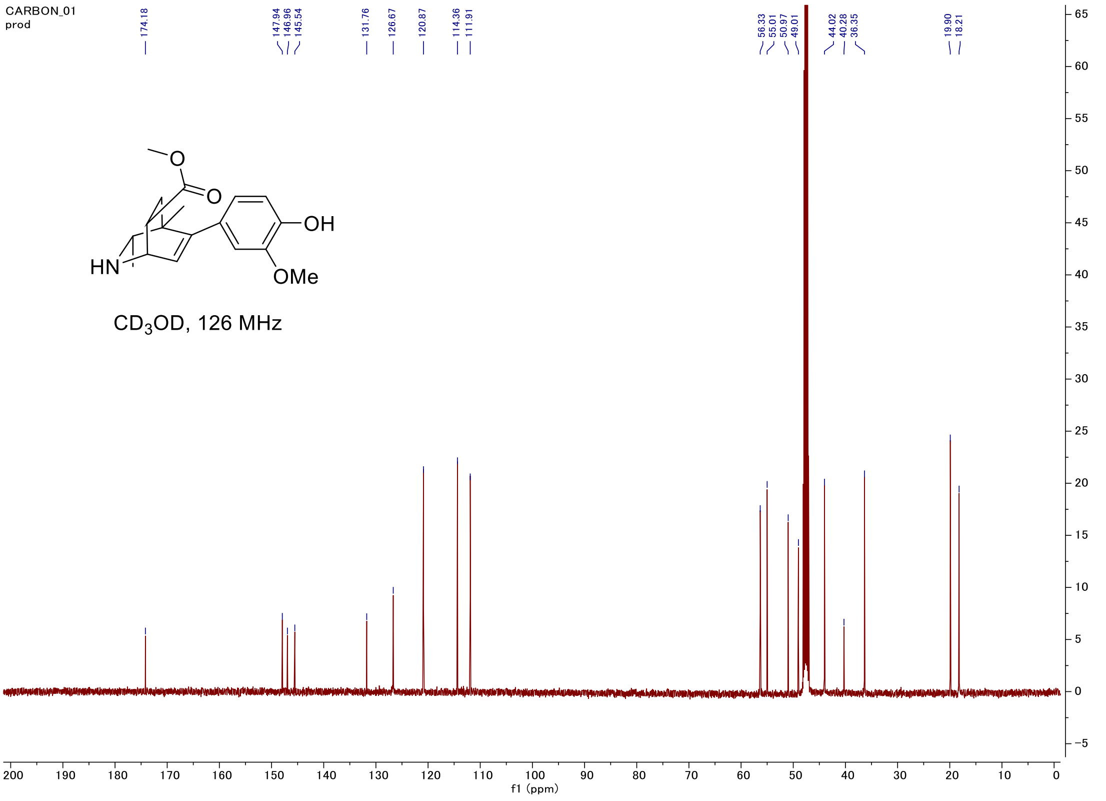

#### C-H COSY

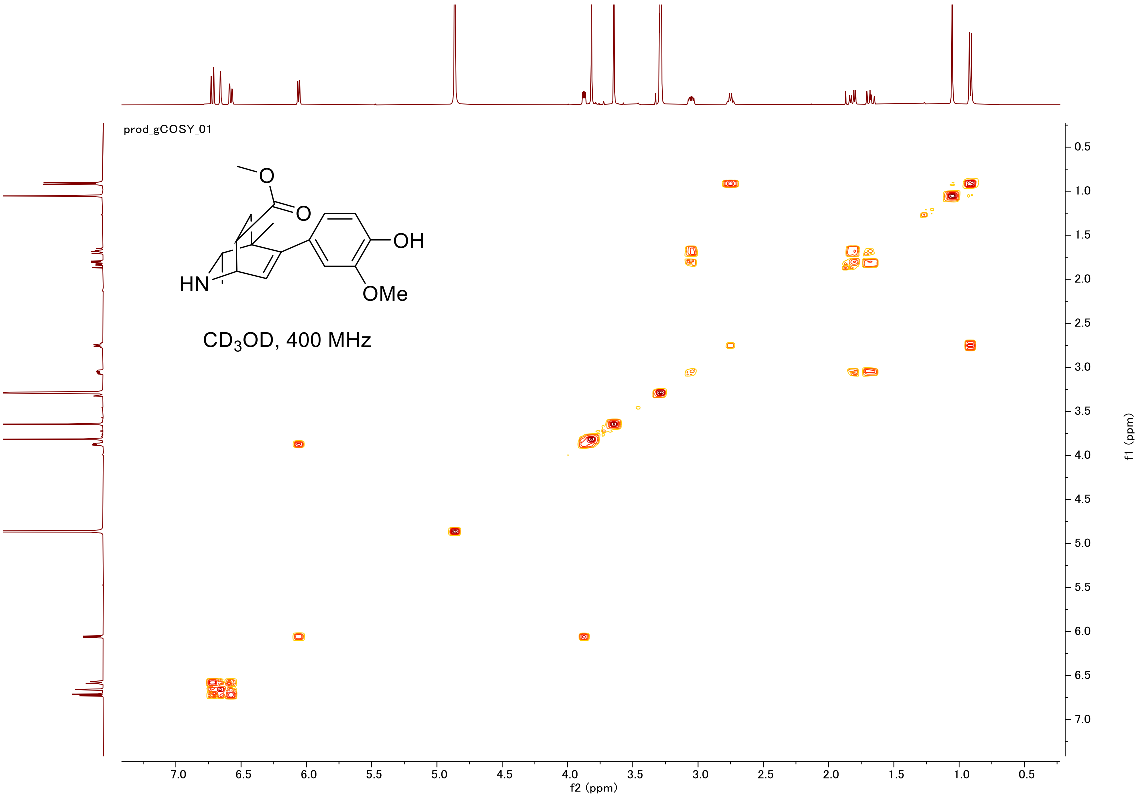
prod_gCOSY_01
OH
HN
OMe
CD3OD, 400 MHz
f2 (ppm)
f1 (ppm)

#### HSQC

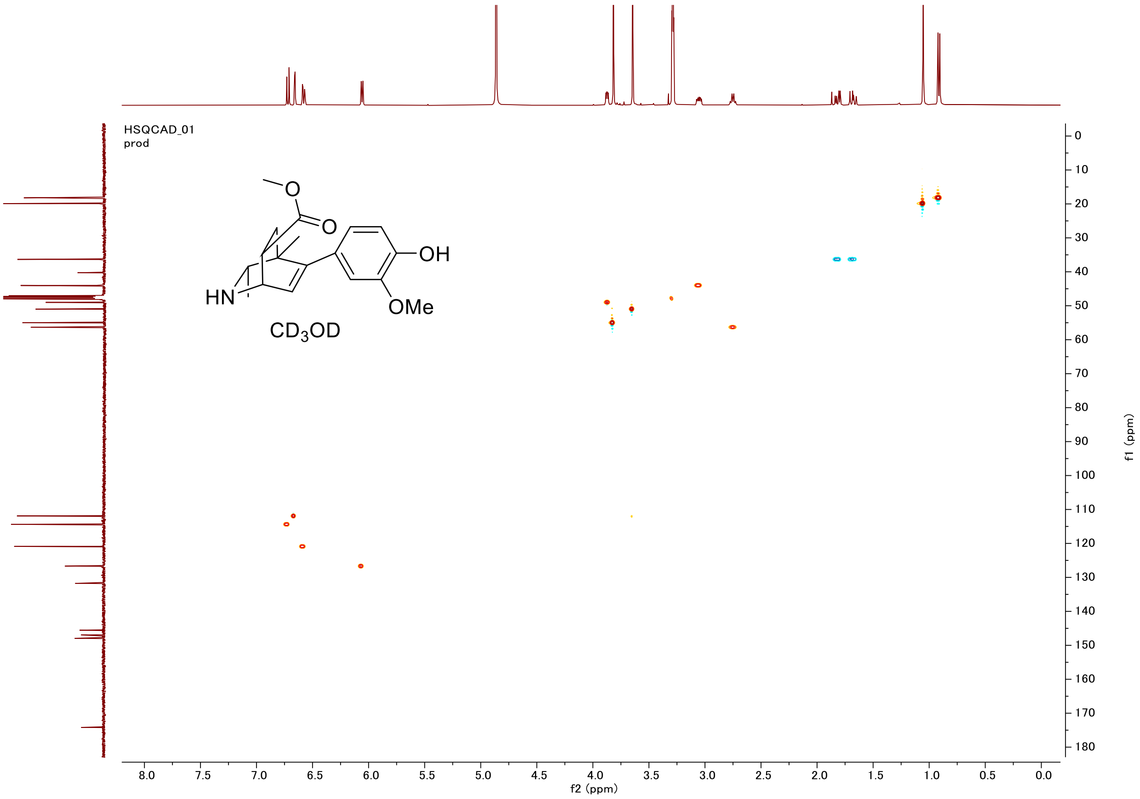
HSQCAD_01
prod
OH
HN
OMe
CD3OD
f2 (ppm)
f1 (ppm)

#### Proton (#009)

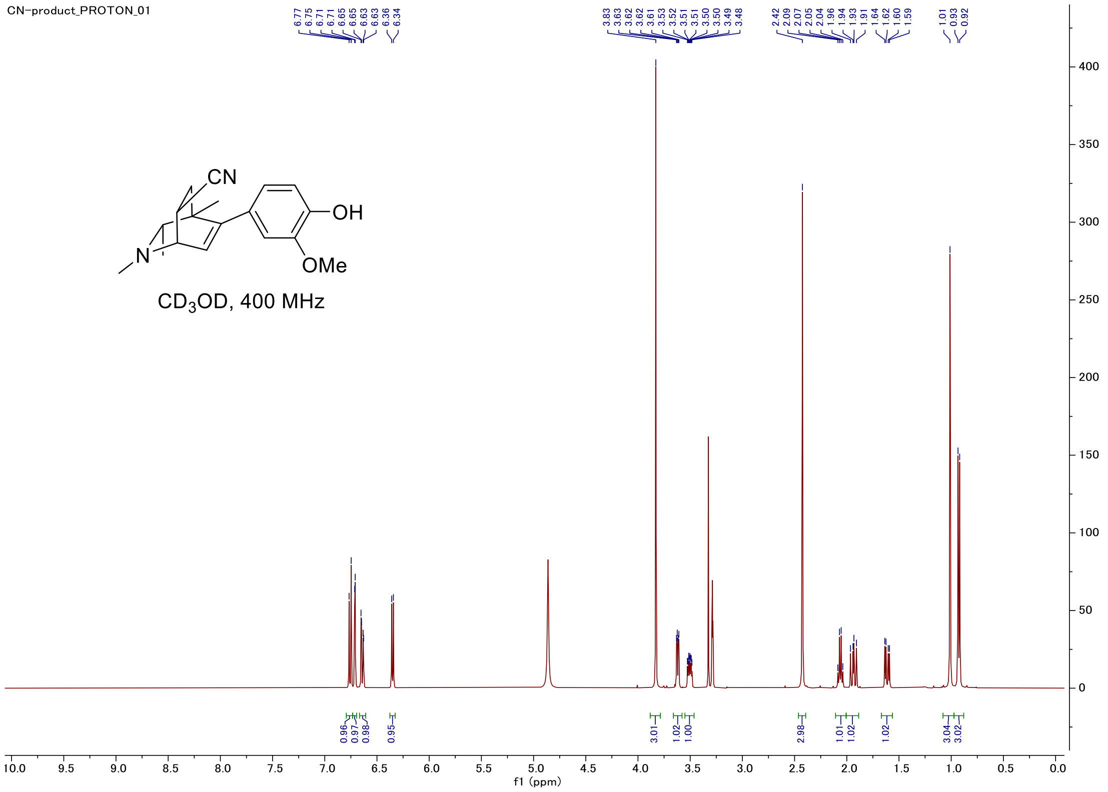


#### Carbon

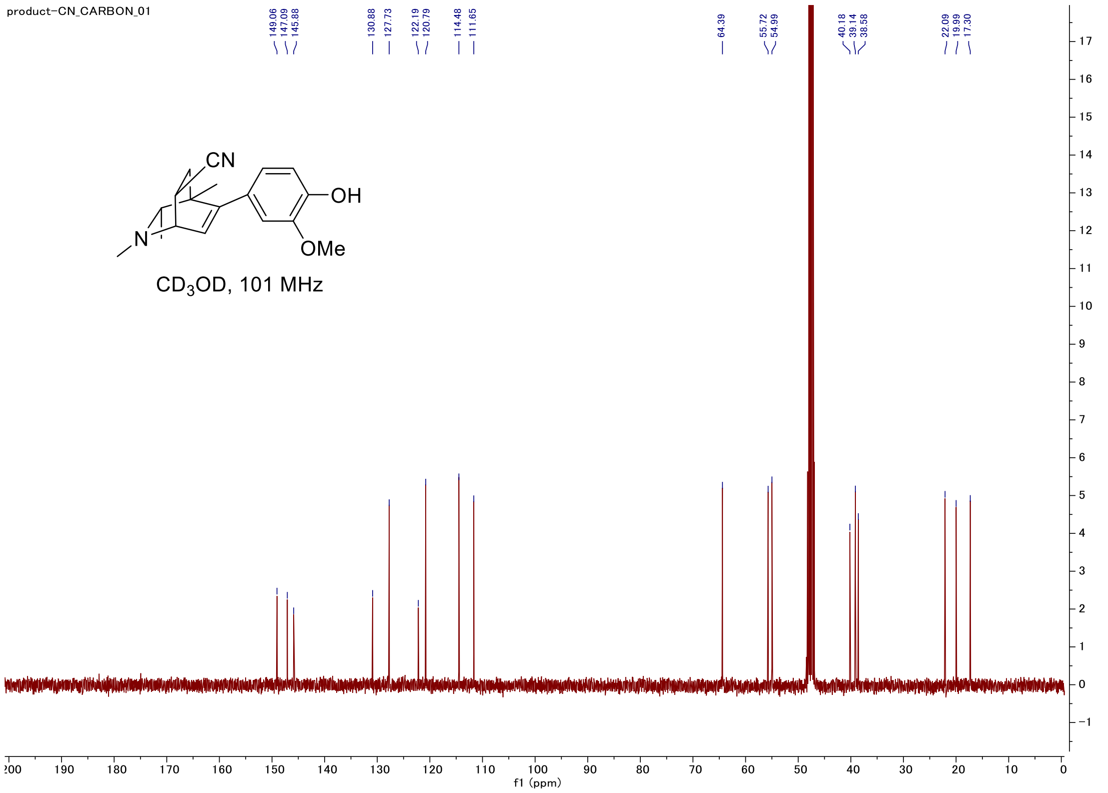

#### C-H COSY

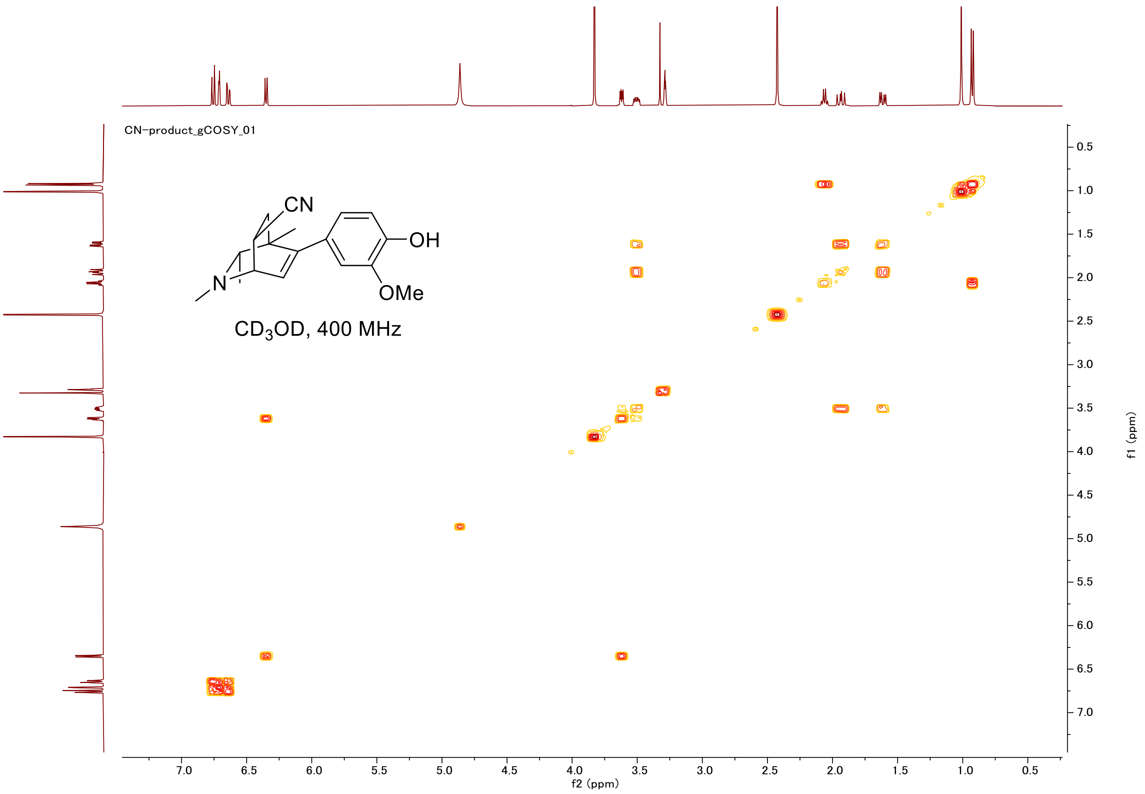
CN-product_gCOSY_01
CN
OH
N
OMe
CD3OD, 400 MHz
f2 (ppm)
f1 (ppm)

#### HSQC

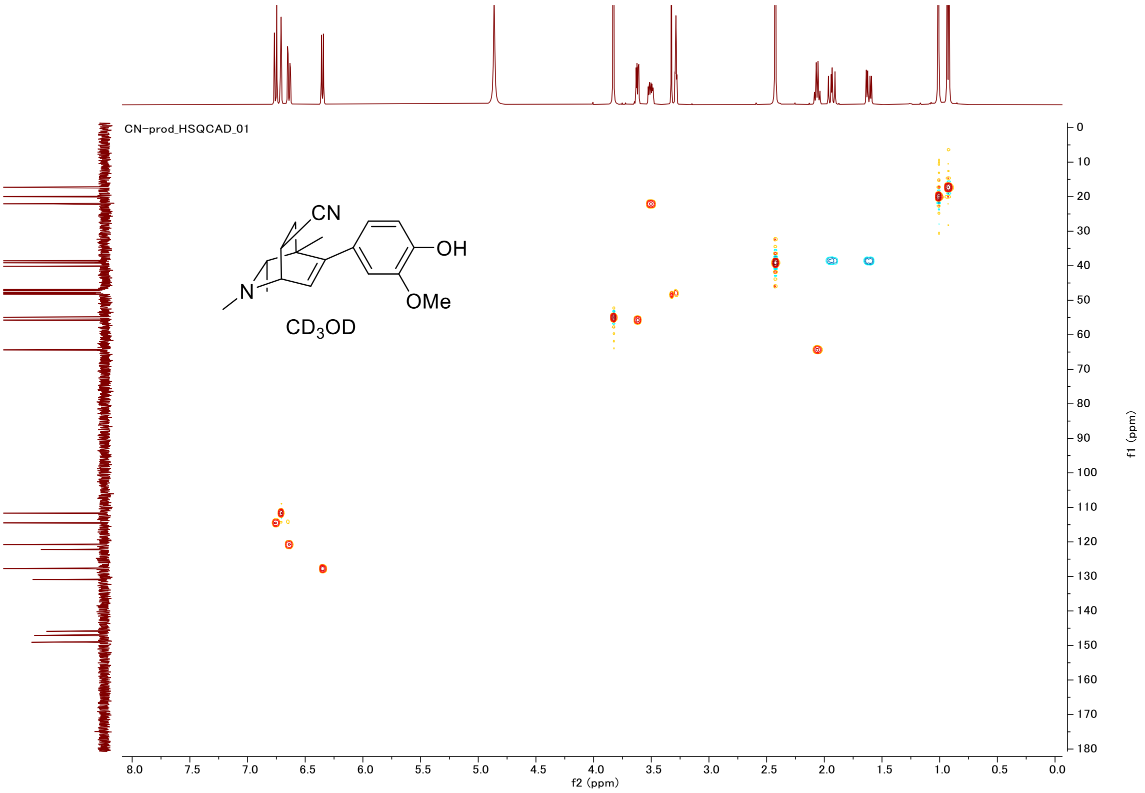
CN-prod_HSQCAD_01
CN
OH
N
OMe
CD3OD
f2 (ppm)
f1 (ppm)

#### Proton (#010)

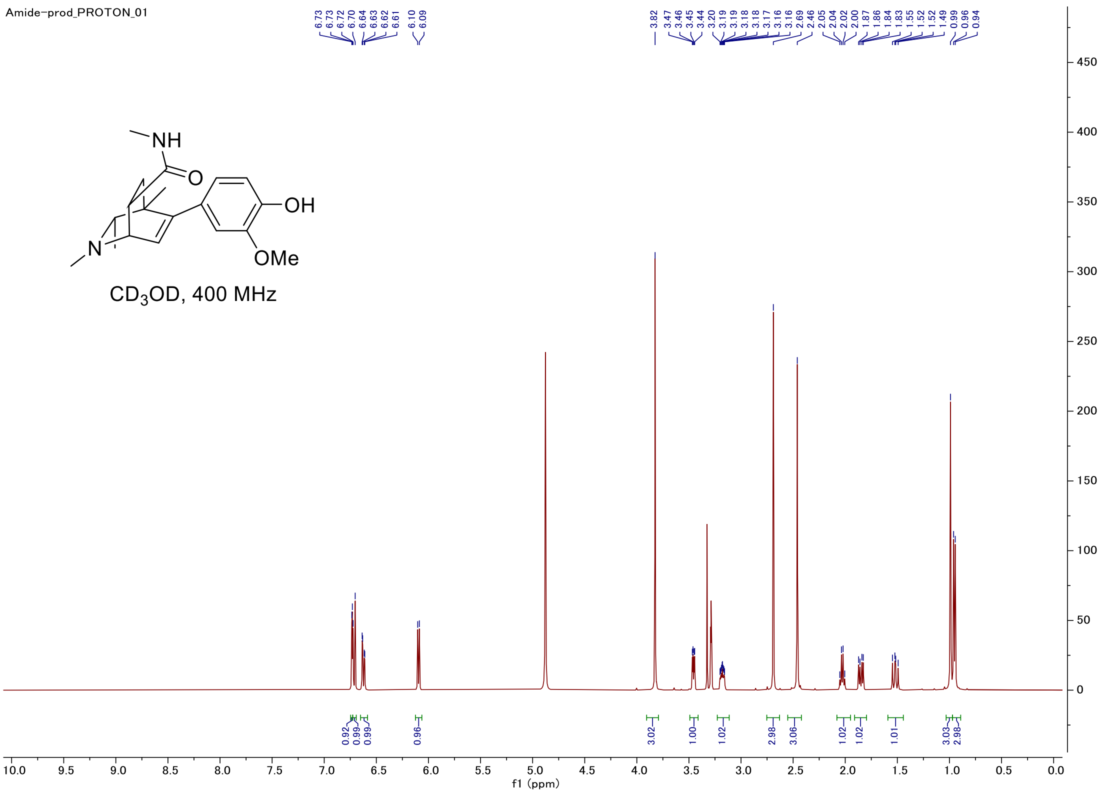


#### Carbon

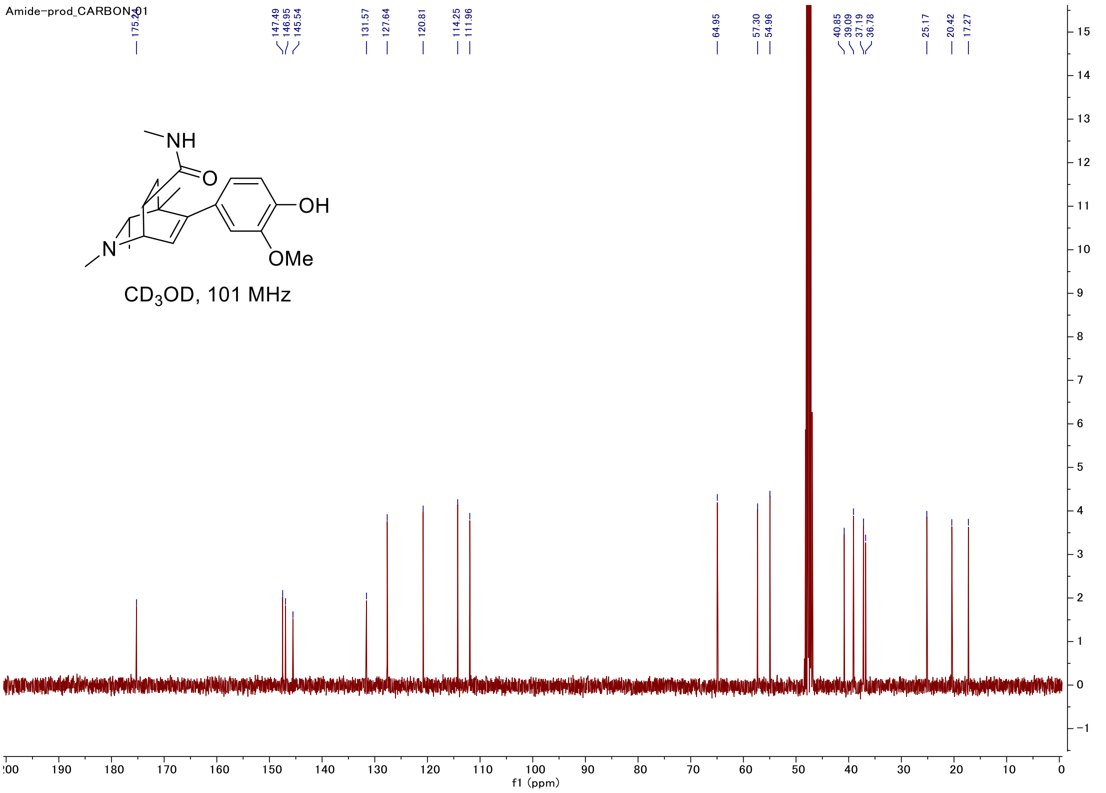

#### C-H COSY

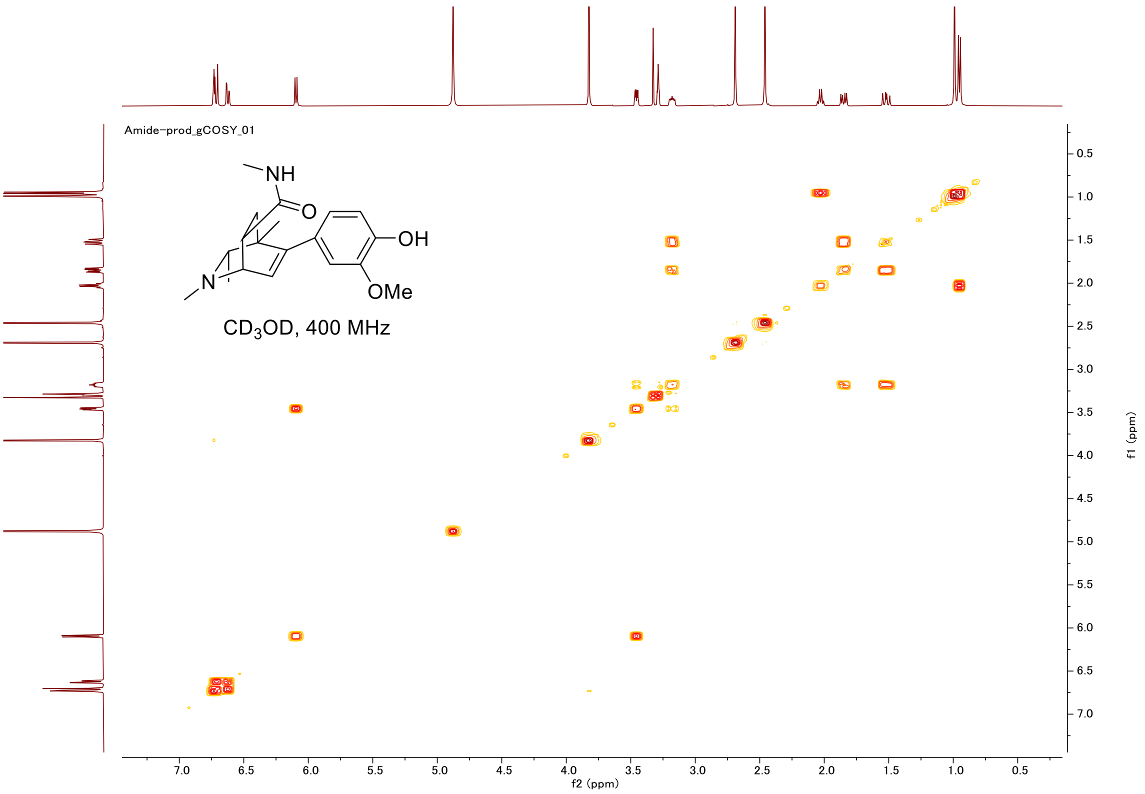

Amide-prod_gCOSY_01
NH
O
OH
N
OMe
CD3OD, 400 MHz
f2 (ppm)
f1 (ppm)


#### HSQC

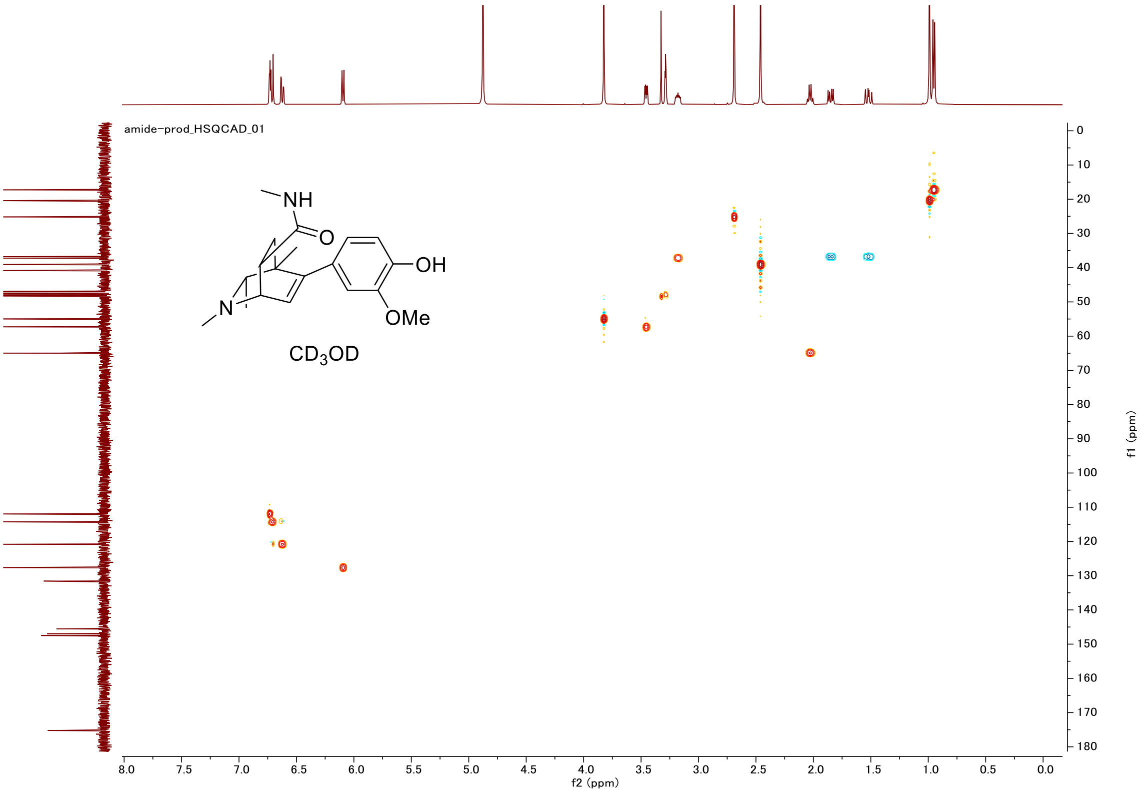

amide-prod_HSQCAD_01
NH
O
OH
N
OMe
CD3OD
f2 (ppm)
f1 (ppm)

#### Proton (#011)

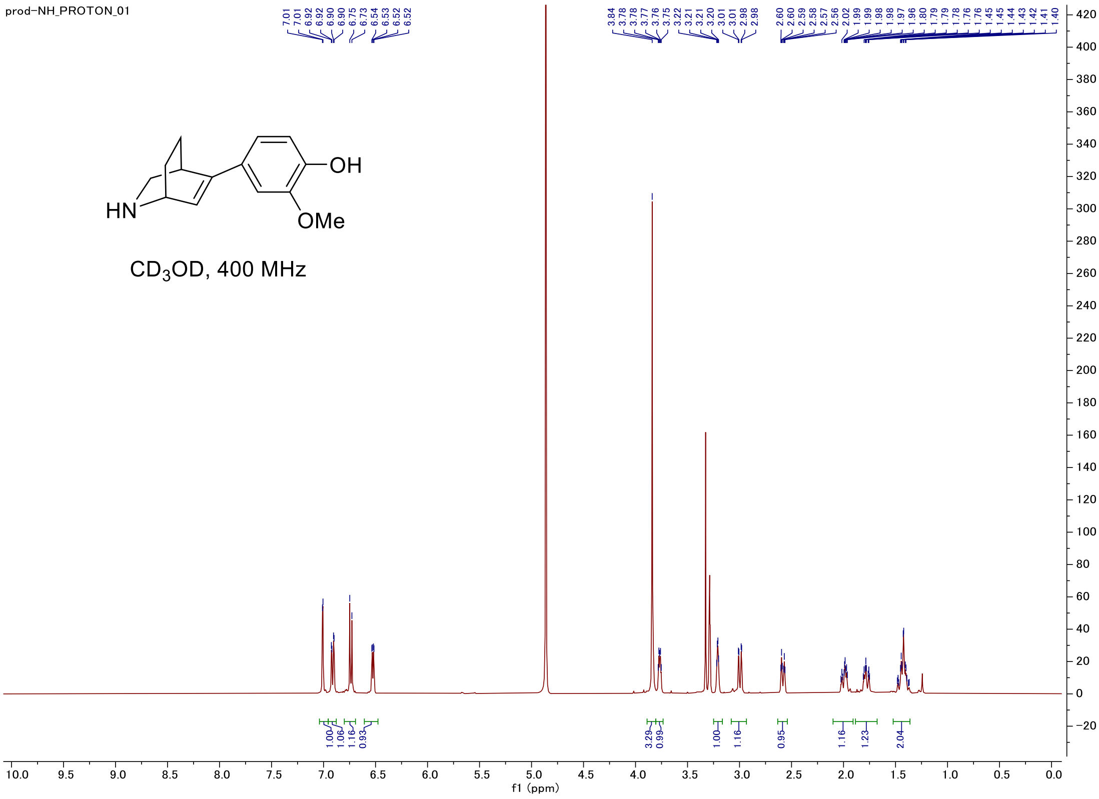


#### Carbon

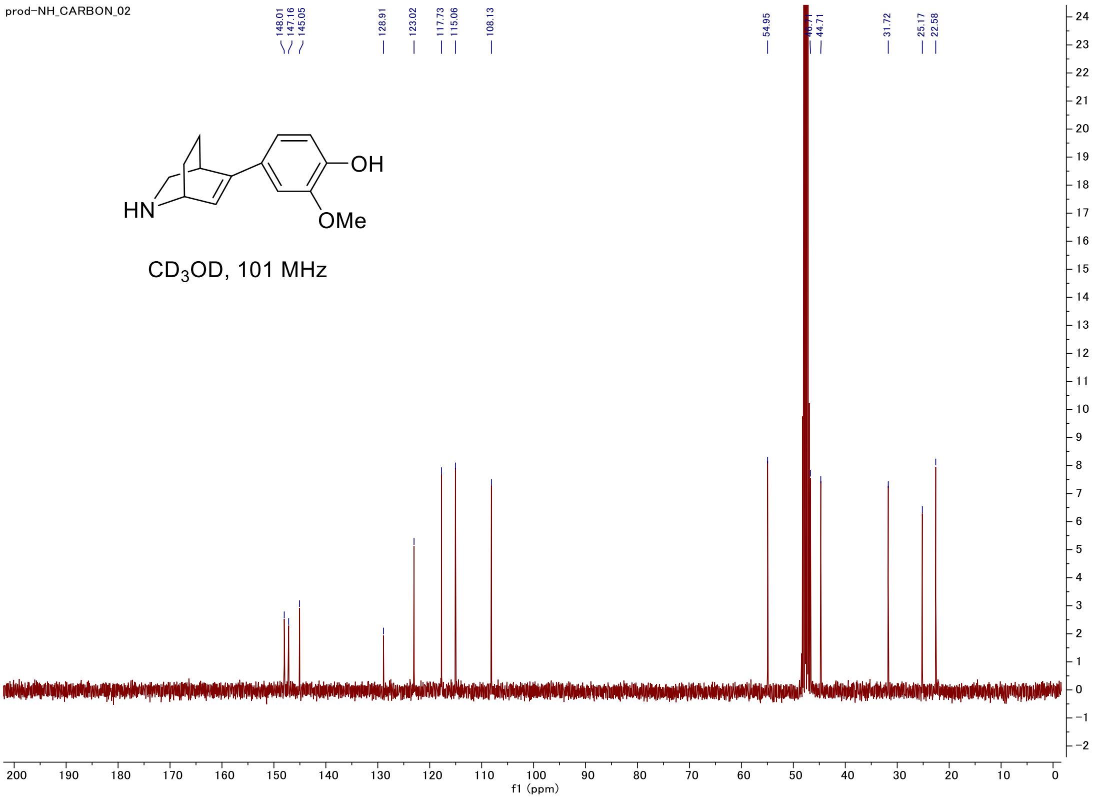

#### C-H COSY

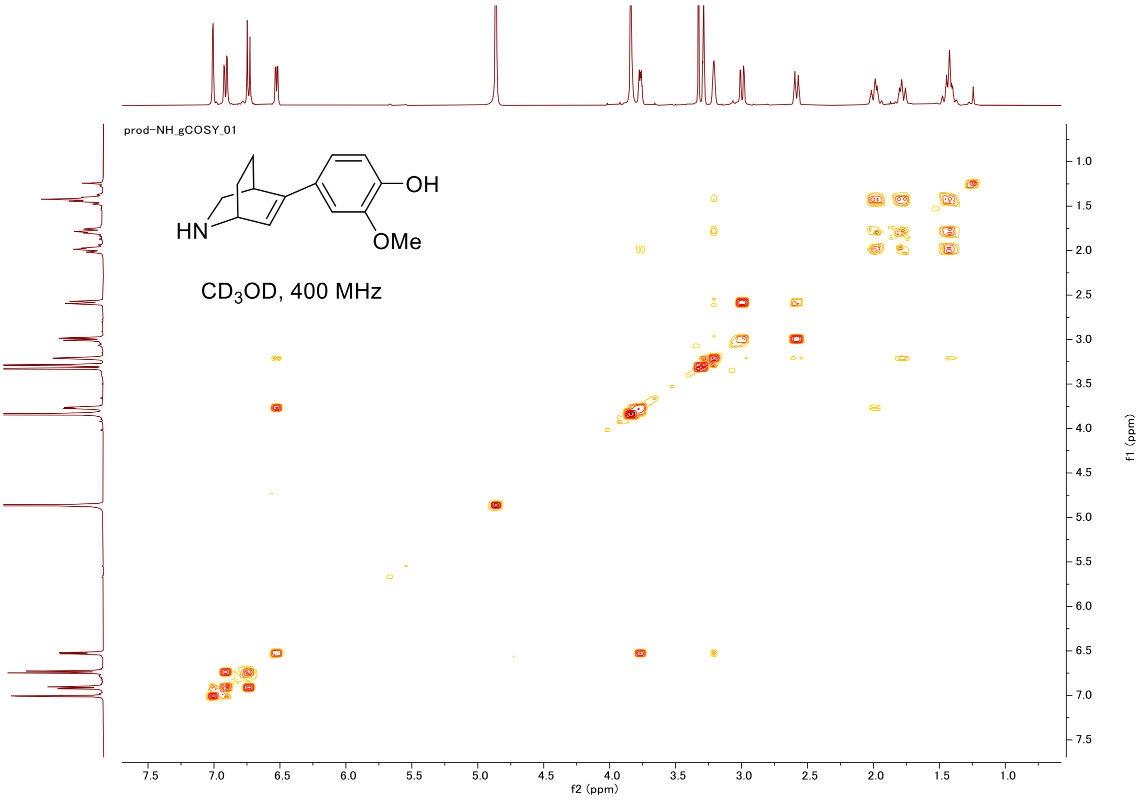

prod-NH_gCOSY_01
HN
OH
OMe
CD3OD, 400 MHz
f2 (ppm)
f1 (ppm)


#### HSQC

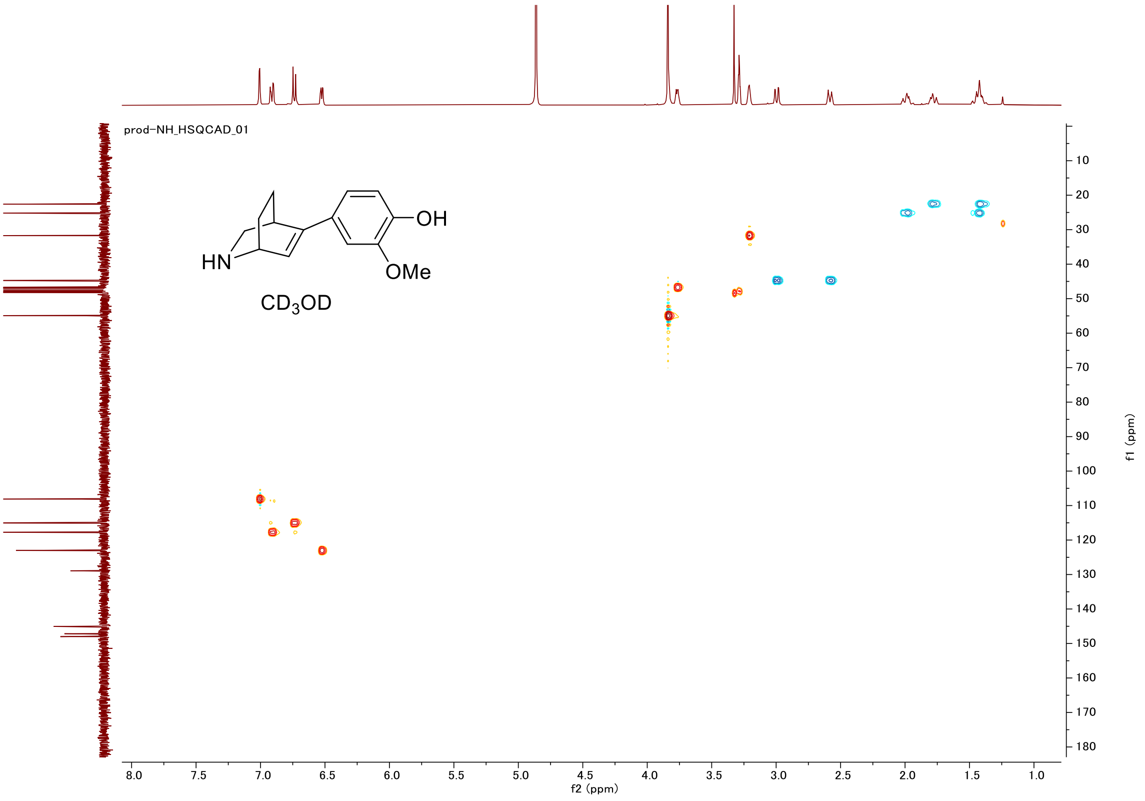

prod-NH_HSQCAD_01
HN
OH
OMe
CD3OD
f2 (ppm)
f1 (ppm)

#### Proton (#012)

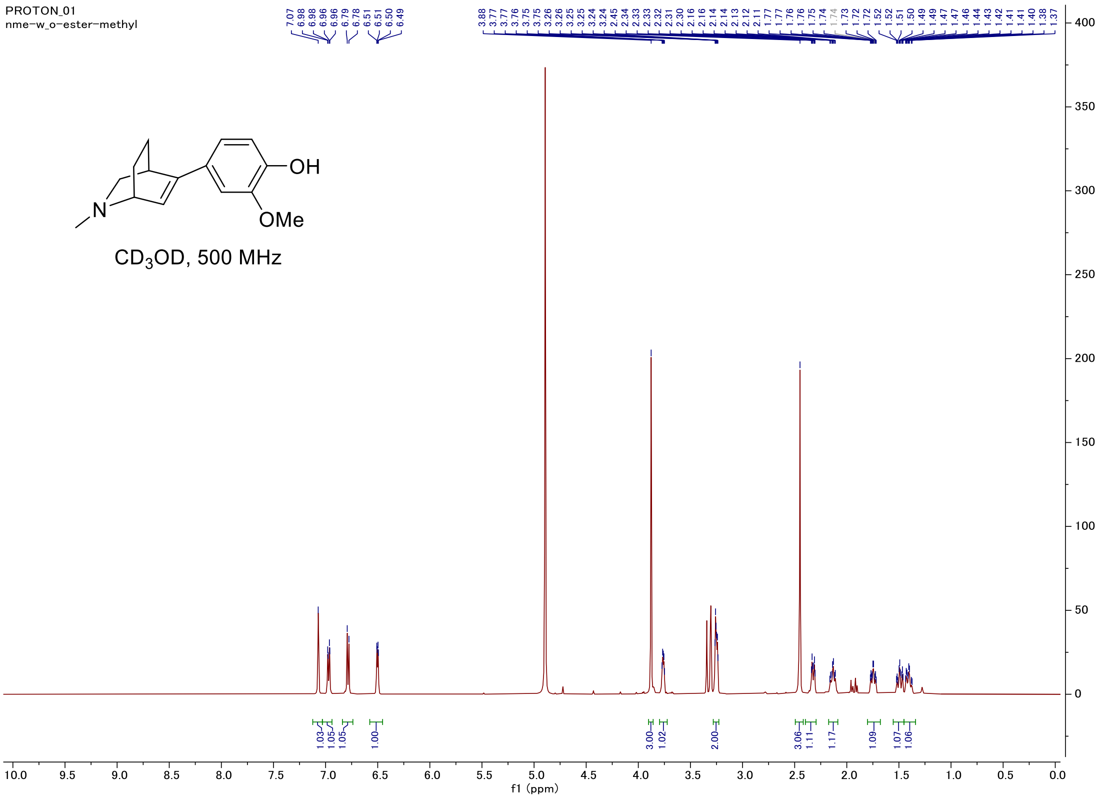


#### Carbon

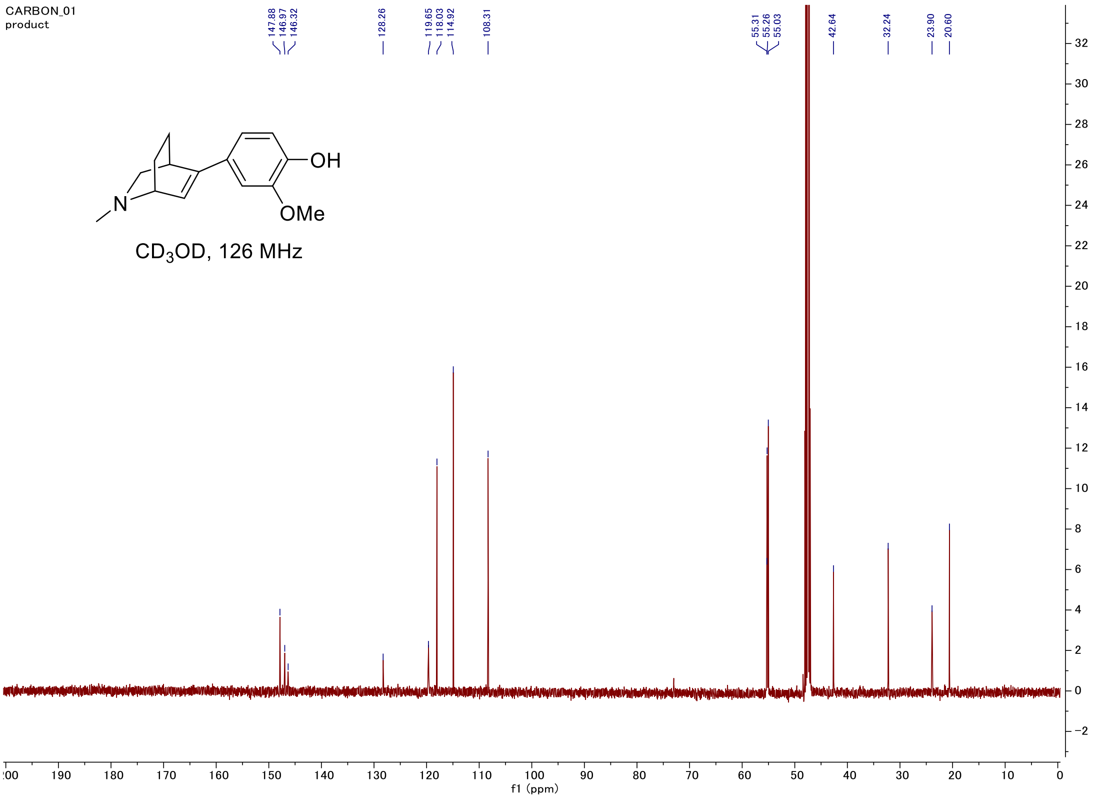

#### C-H COSY

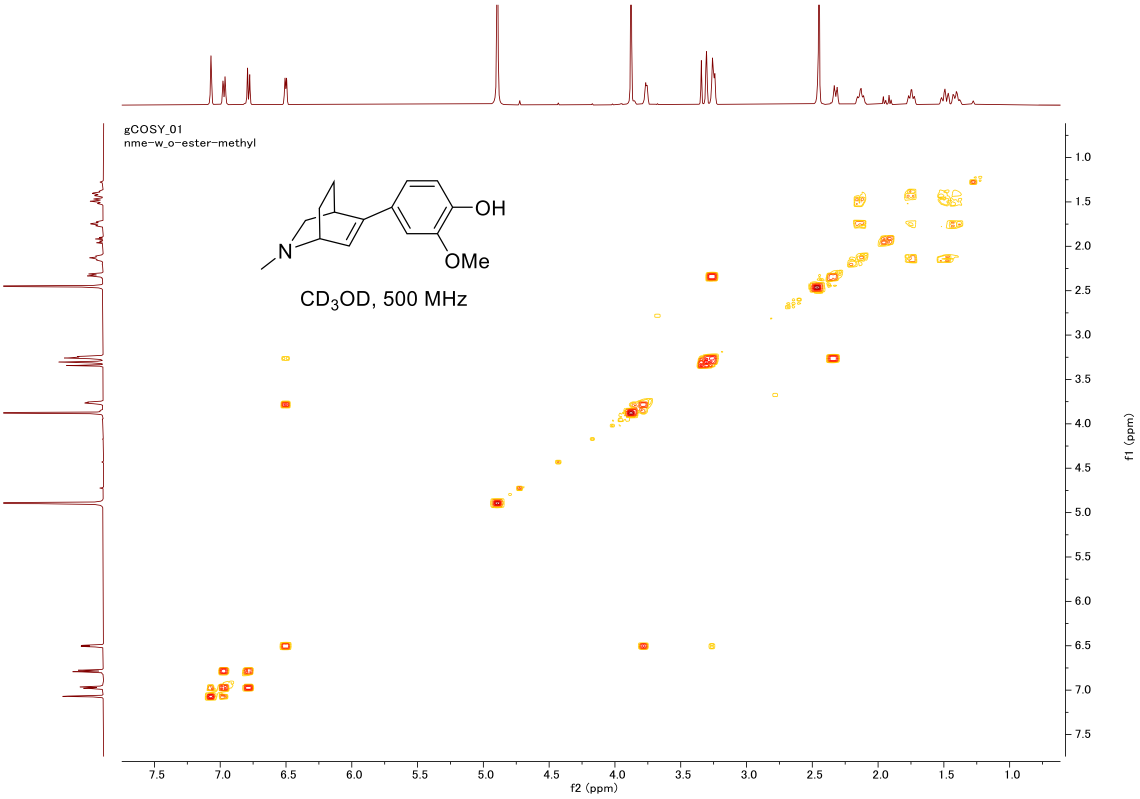

gCOSY_01
nme-w_o-ester-methyl
OH
N
OMe
CD3OD, 500 MHz
f2 (ppm)
f1 (ppm)


#### HSQC

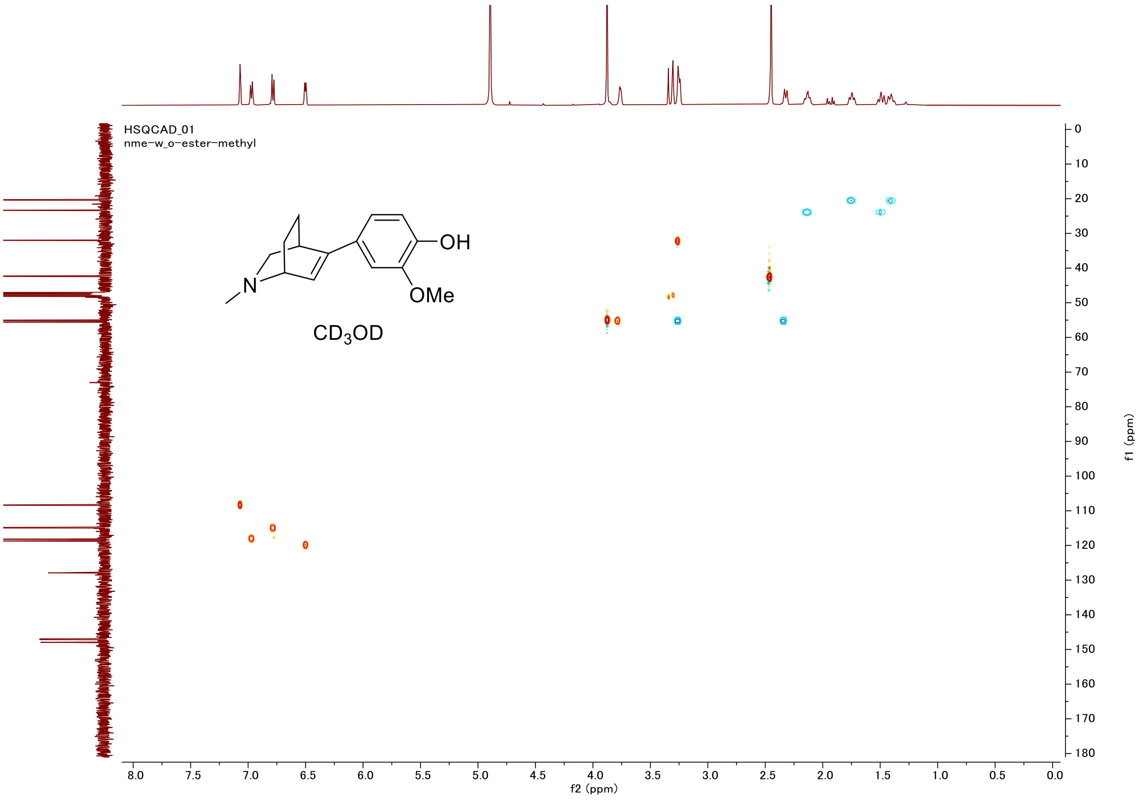

HSQCAD_01
nme-w_o-ester-methyl
OH
N
OMe
CD3OD
f2 (ppm)
f1 (ppm)

#### Proton (#013)

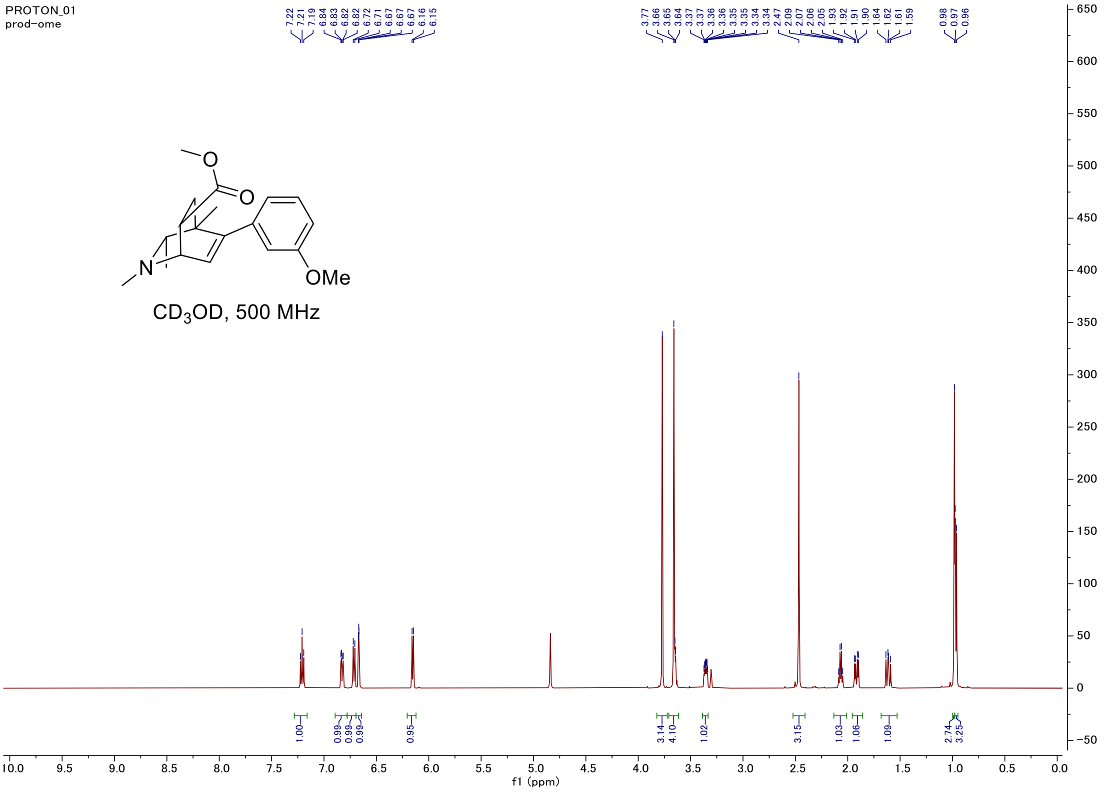


#### Carbon

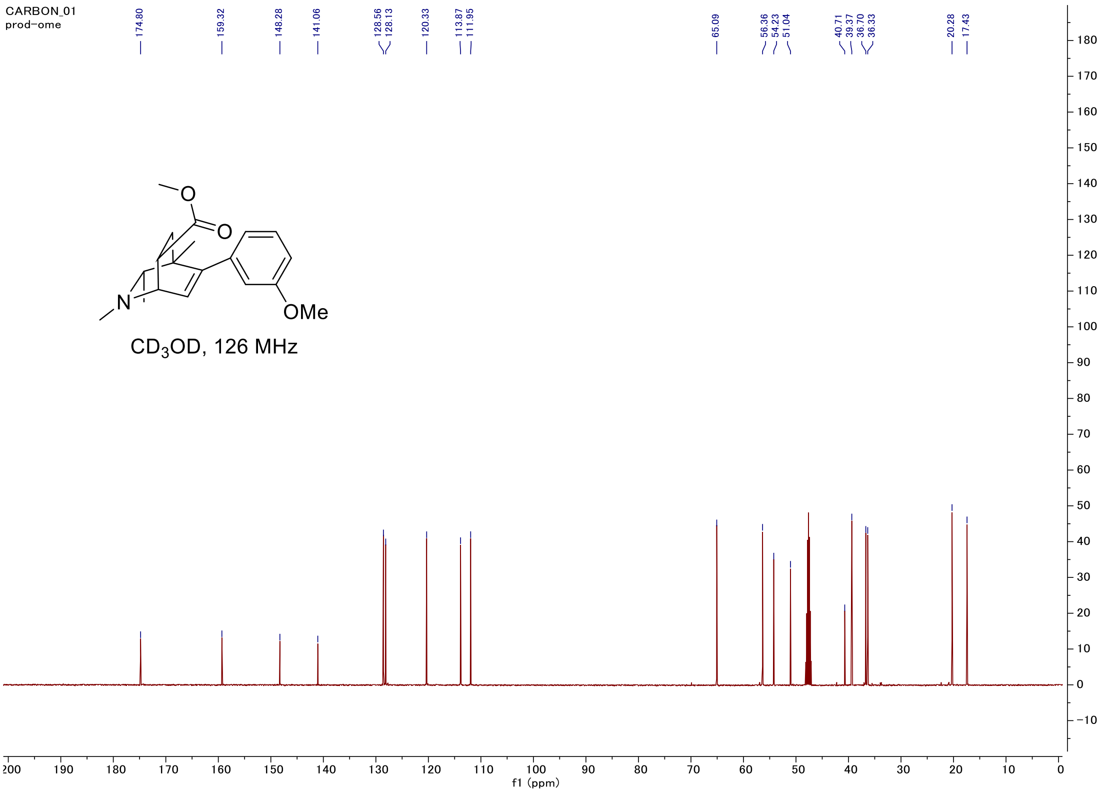

### C-H COSY

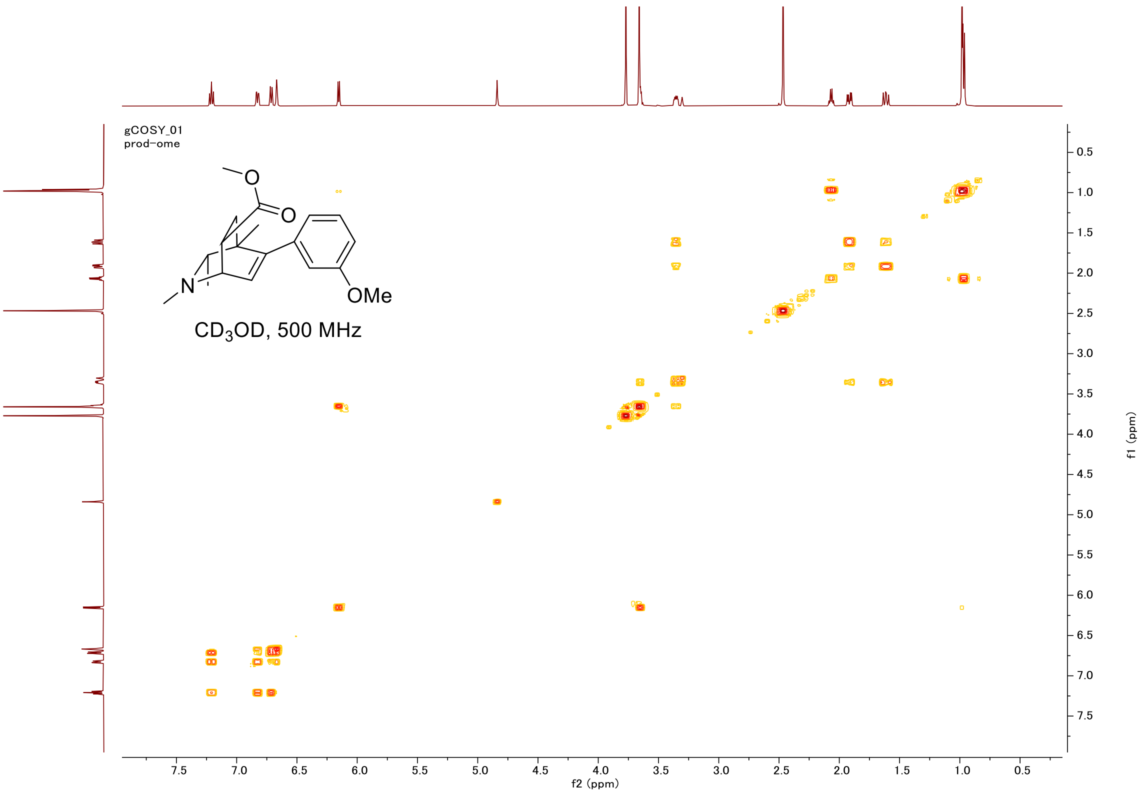
gCOSY_01
prod-ome
OMe
CD3OD, 500 MHz
f2 (ppm)
f1 (ppm)

### HSQC

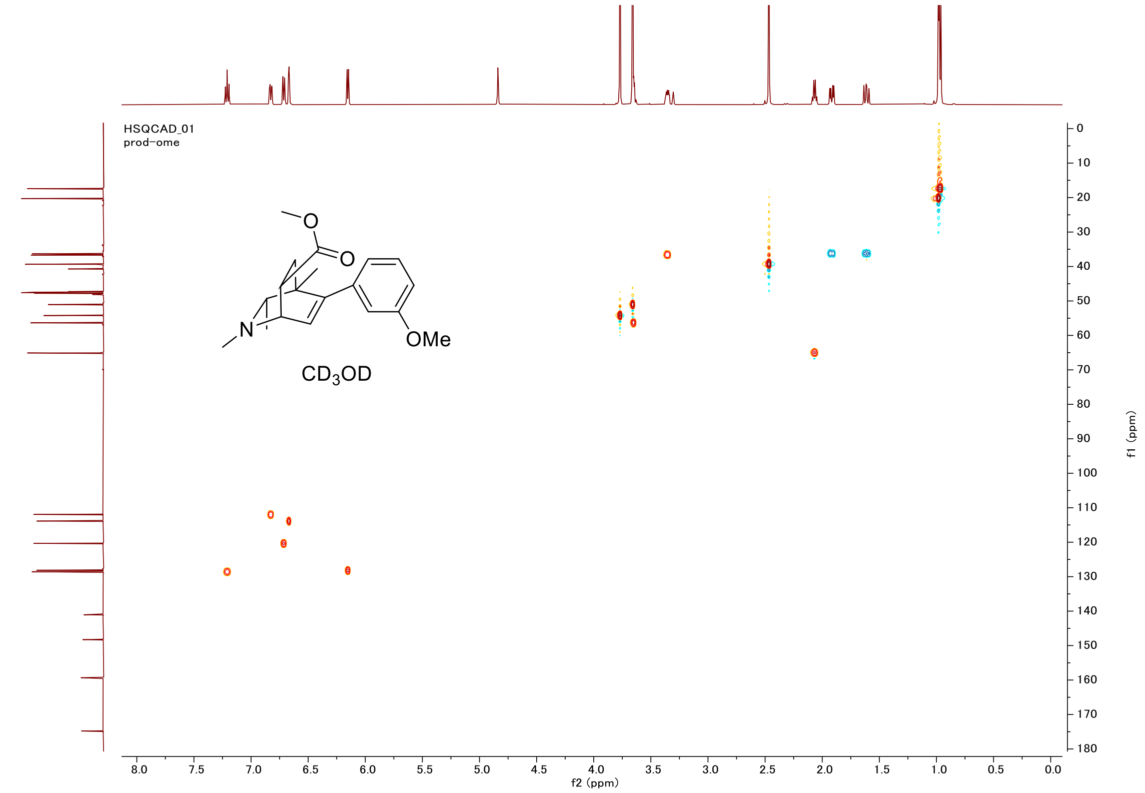
HSQCAD_01
prod-ome
OMe
CD3OD
f2 (ppm)
f1 (ppm)

### Proton (#014)

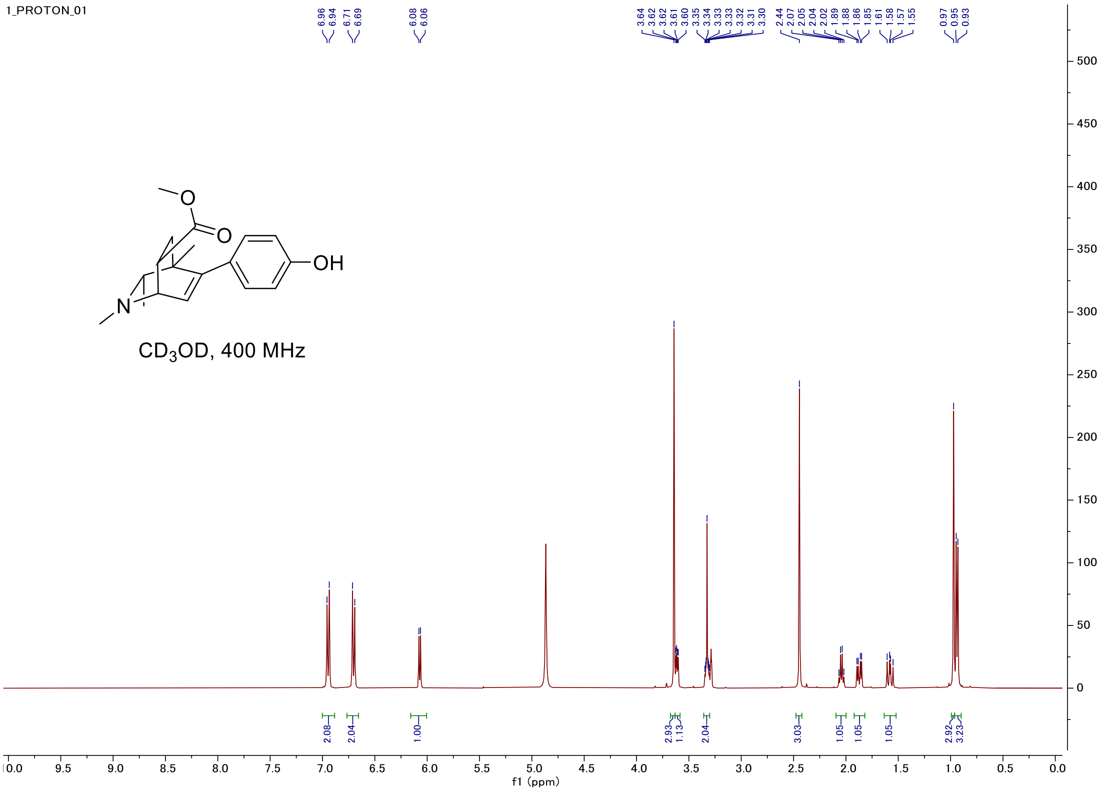


### Carbon

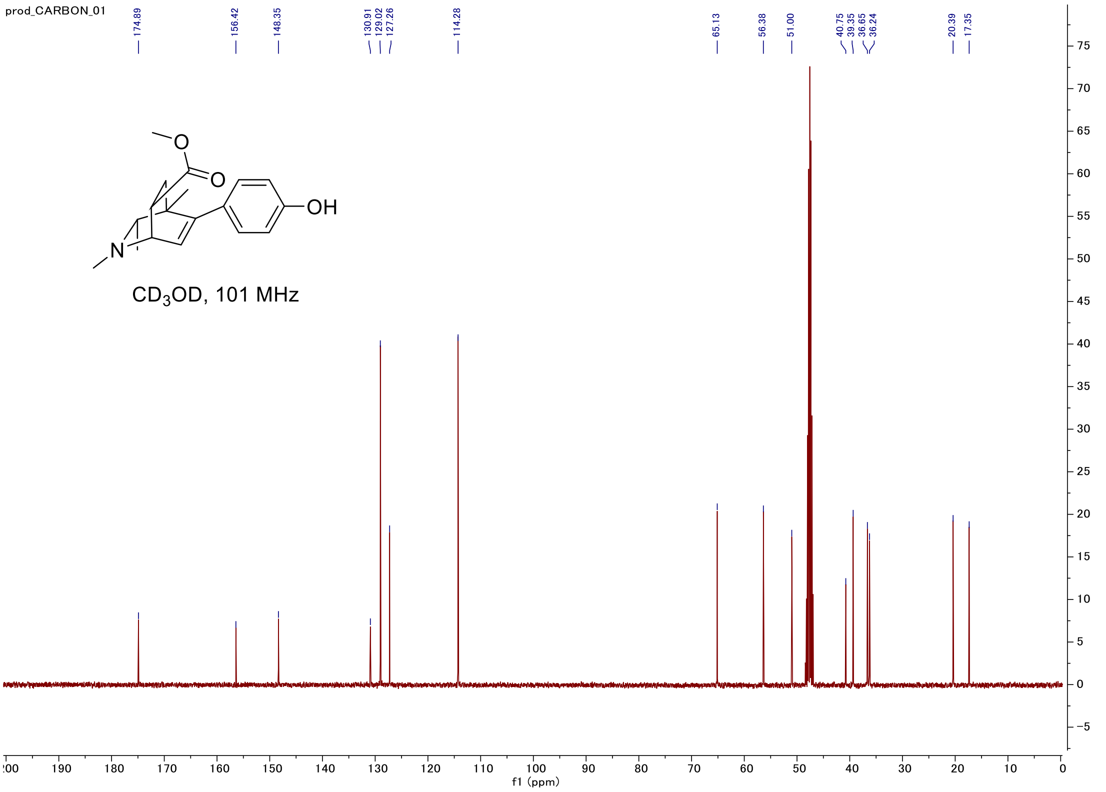

#### C-H COSY

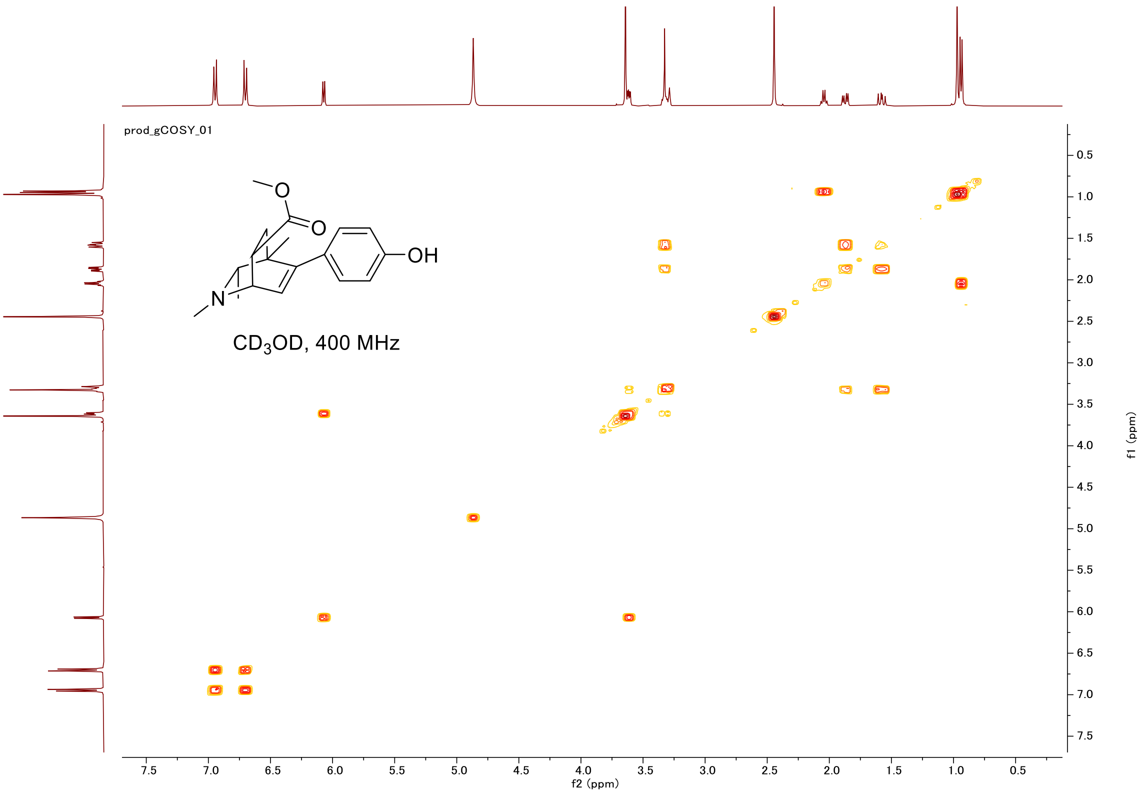
prod_gCOSY_01
OH
N
O
O
CD$_3$OD, 400 MHz
f2 (ppm)
f1 (ppm)

#### HSQC

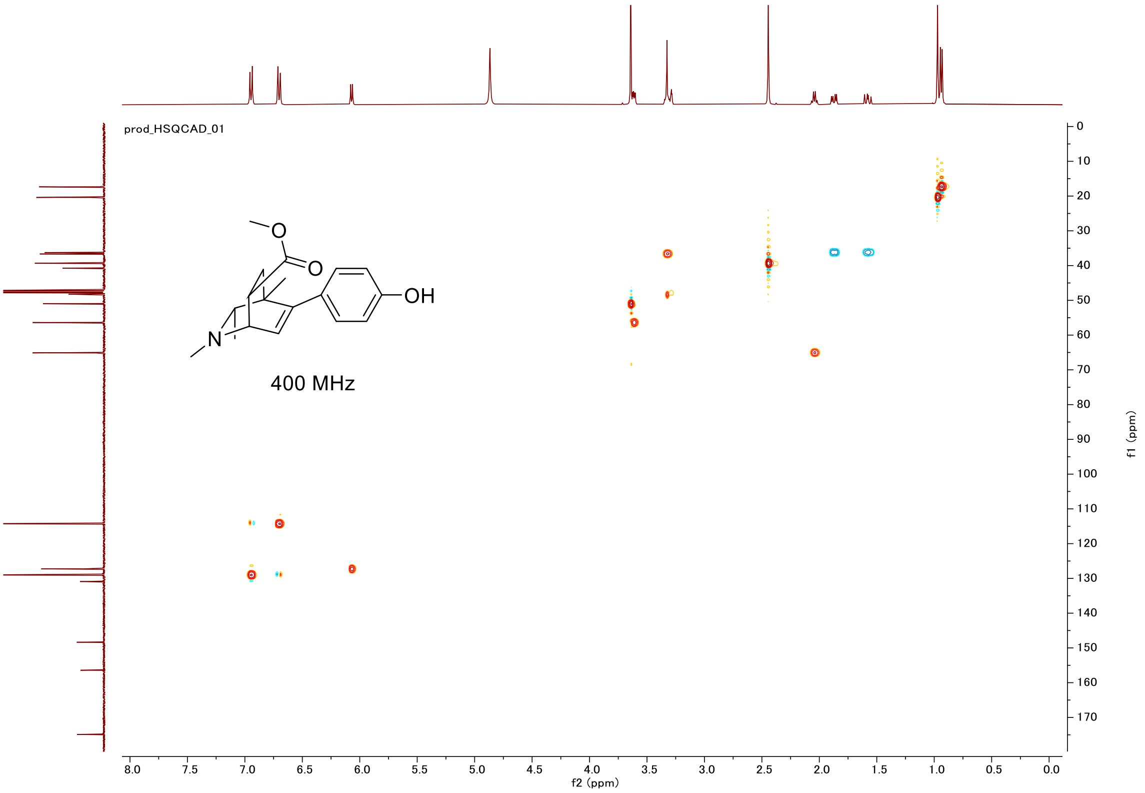
prod_HSQCAD_01
OH
N
O
O
400 MHz
f2 (ppm)
f1 (ppm)

#### Proton (#015)

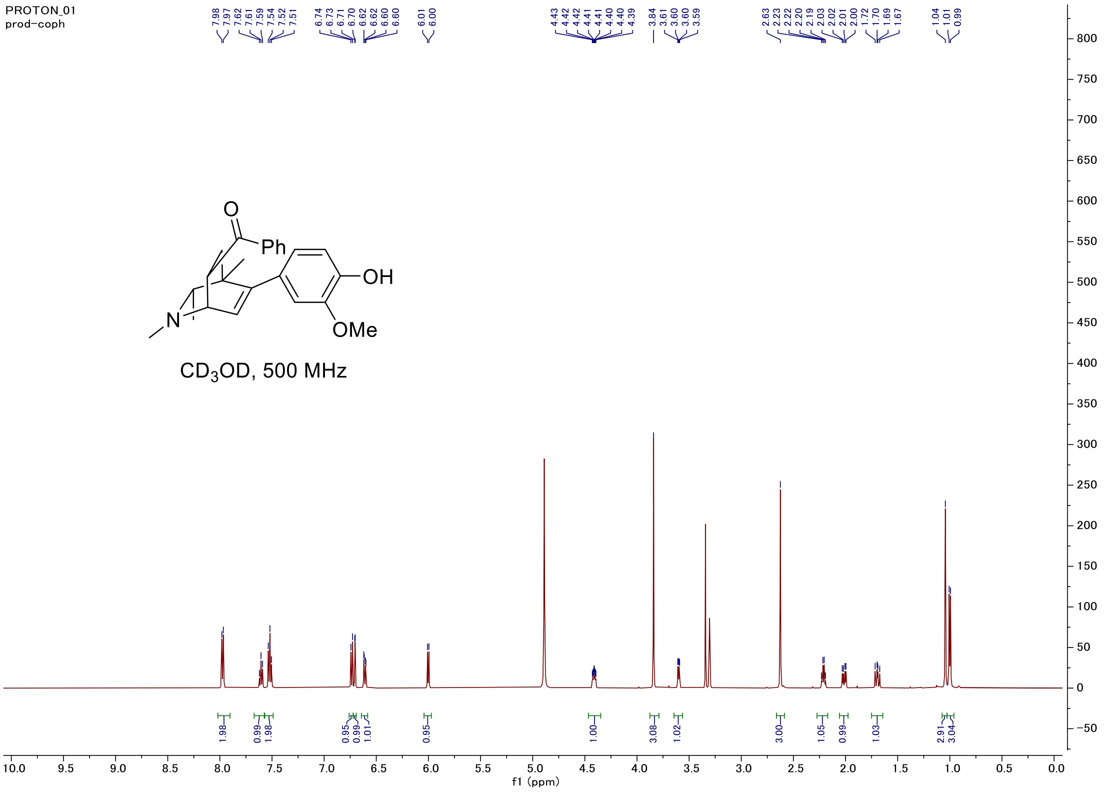


#### Carbon

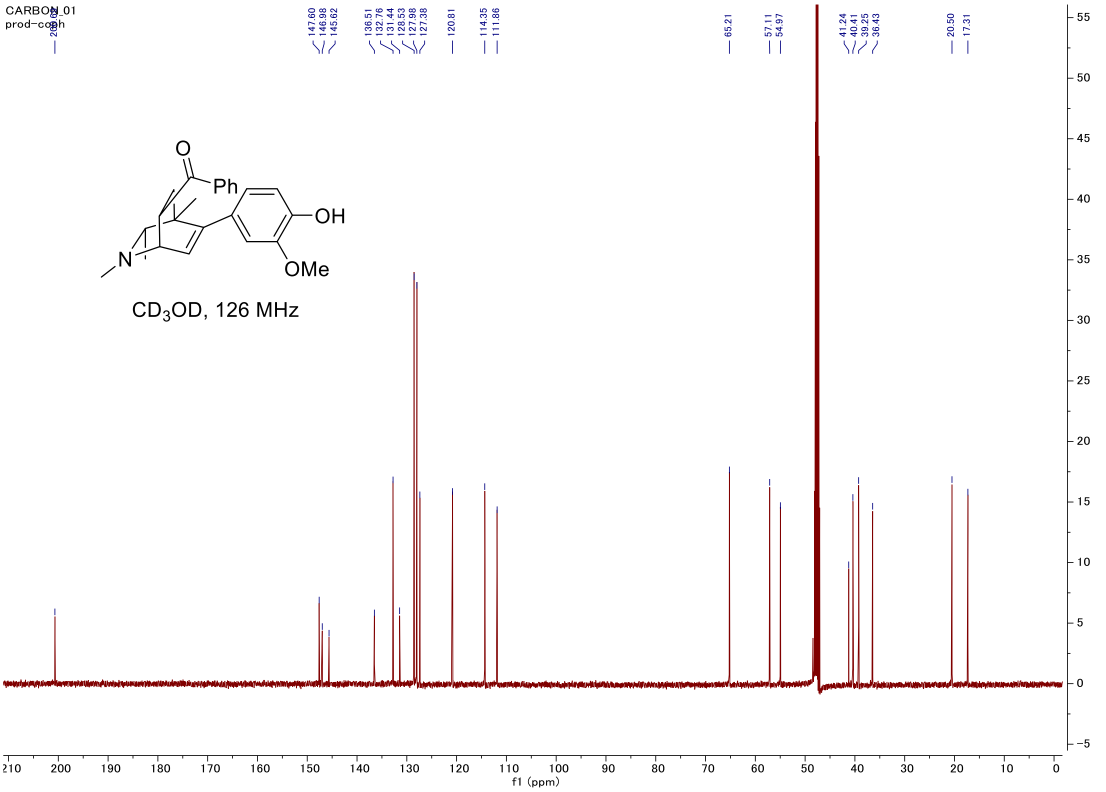

#### C-H COSY

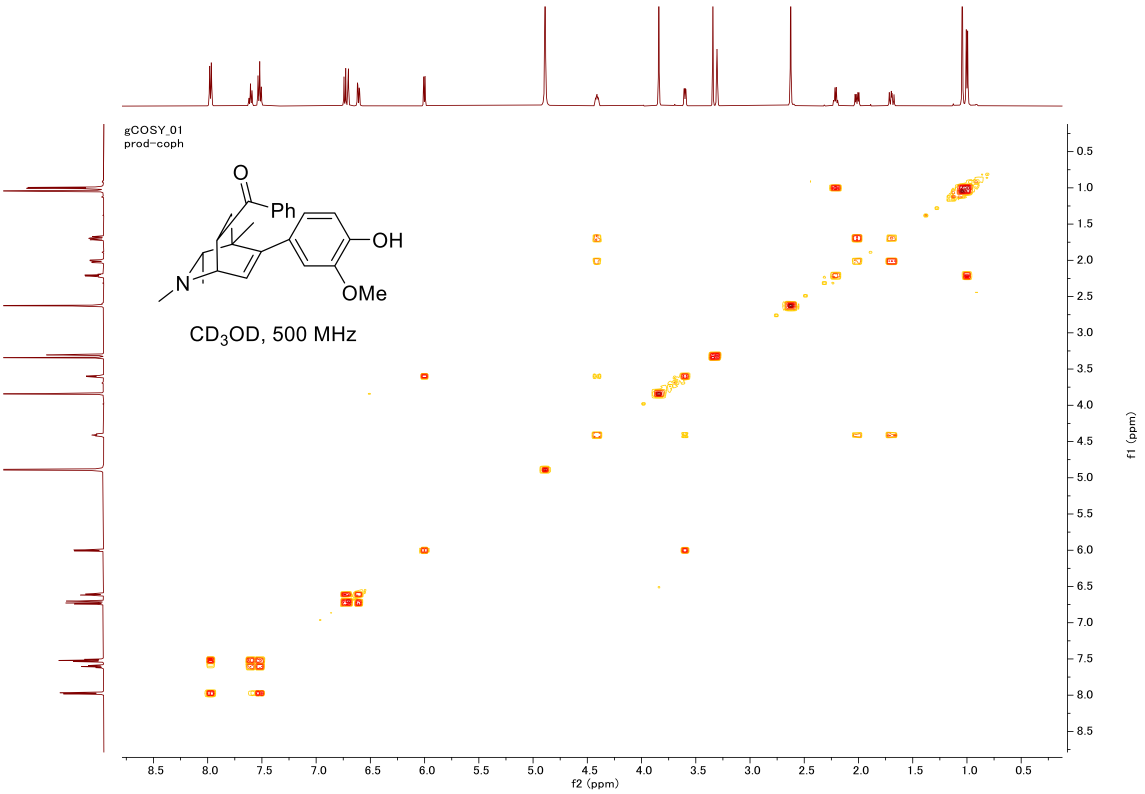
gCOSY_01
prod-coph
O
Ph
OH
N
OMe
CD3OD, 500 MHz
f2 (ppm)
f1 (ppm)

#### HSQC

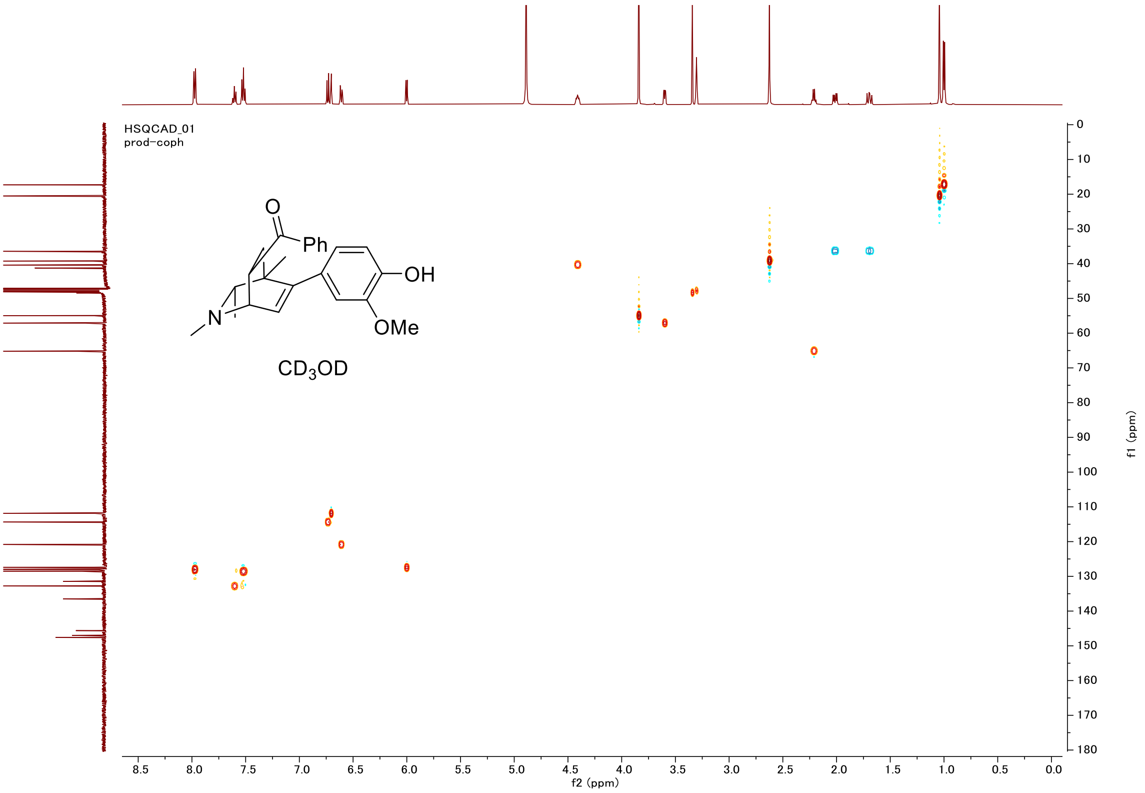
HSQCAD_01
prod-coph
O
Ph
OH
N
OMe
CD3OD
f2 (ppm)
f1 (ppm)

#### Proton (#016)

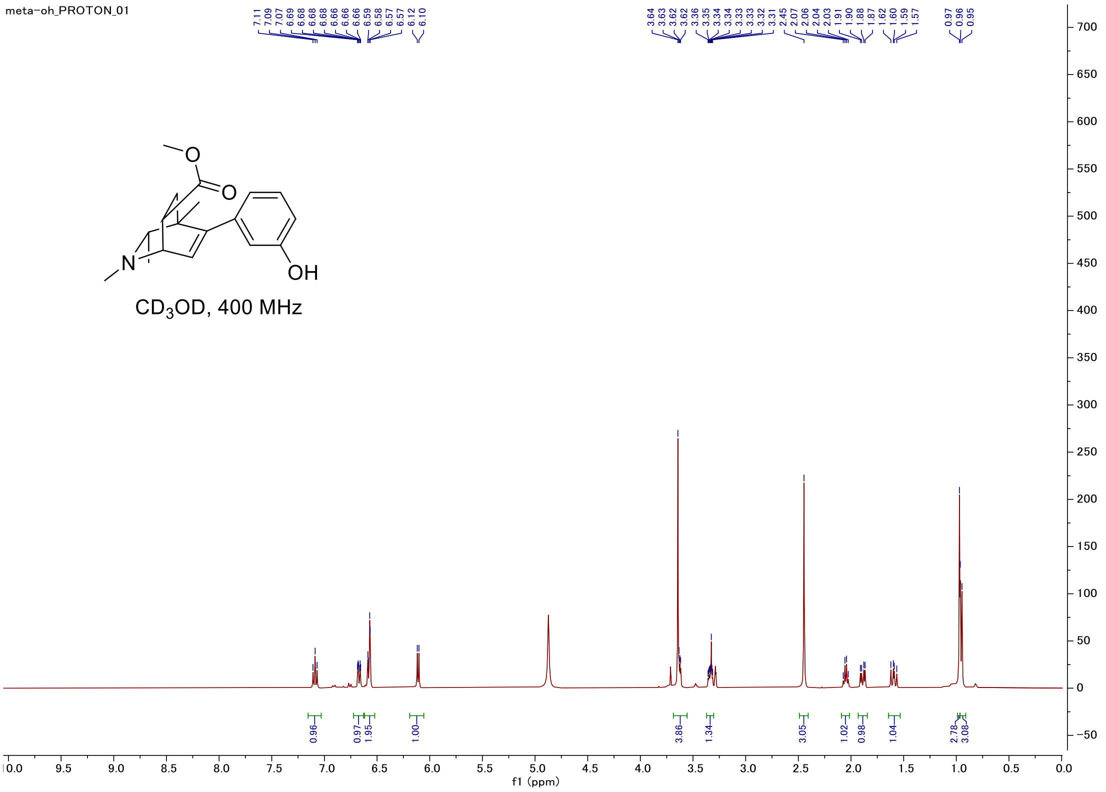


#### Carbon

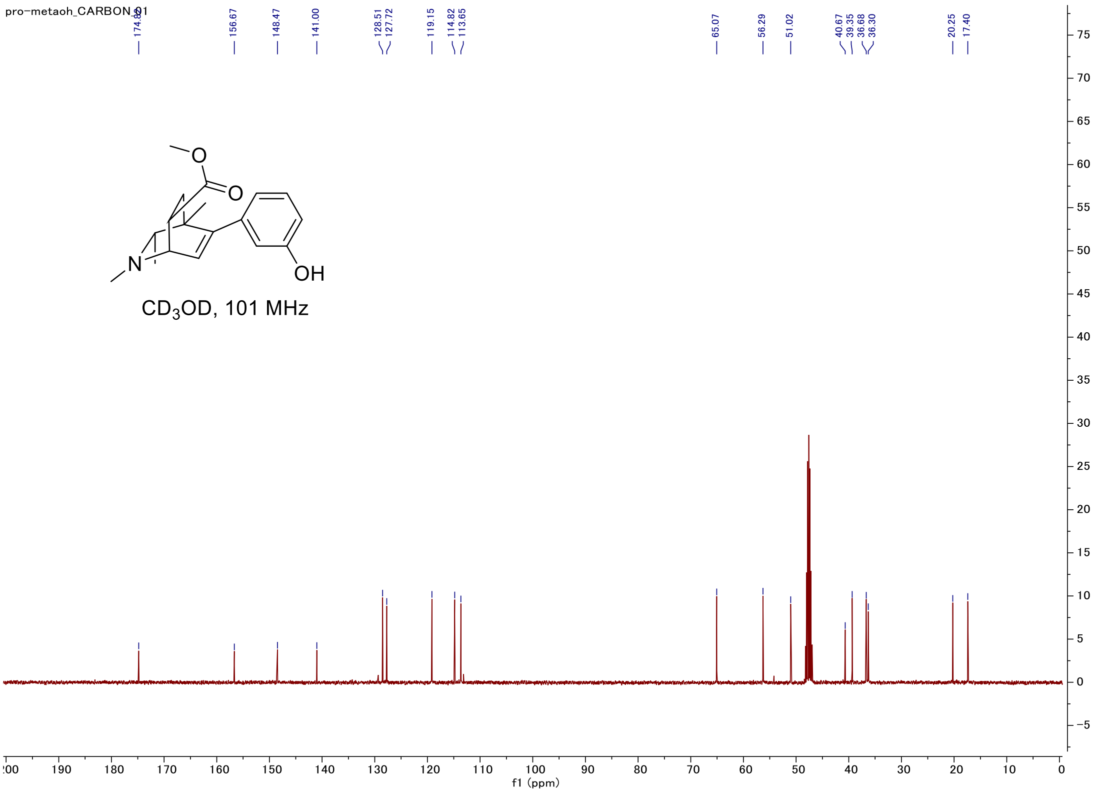

#### C-H COSY

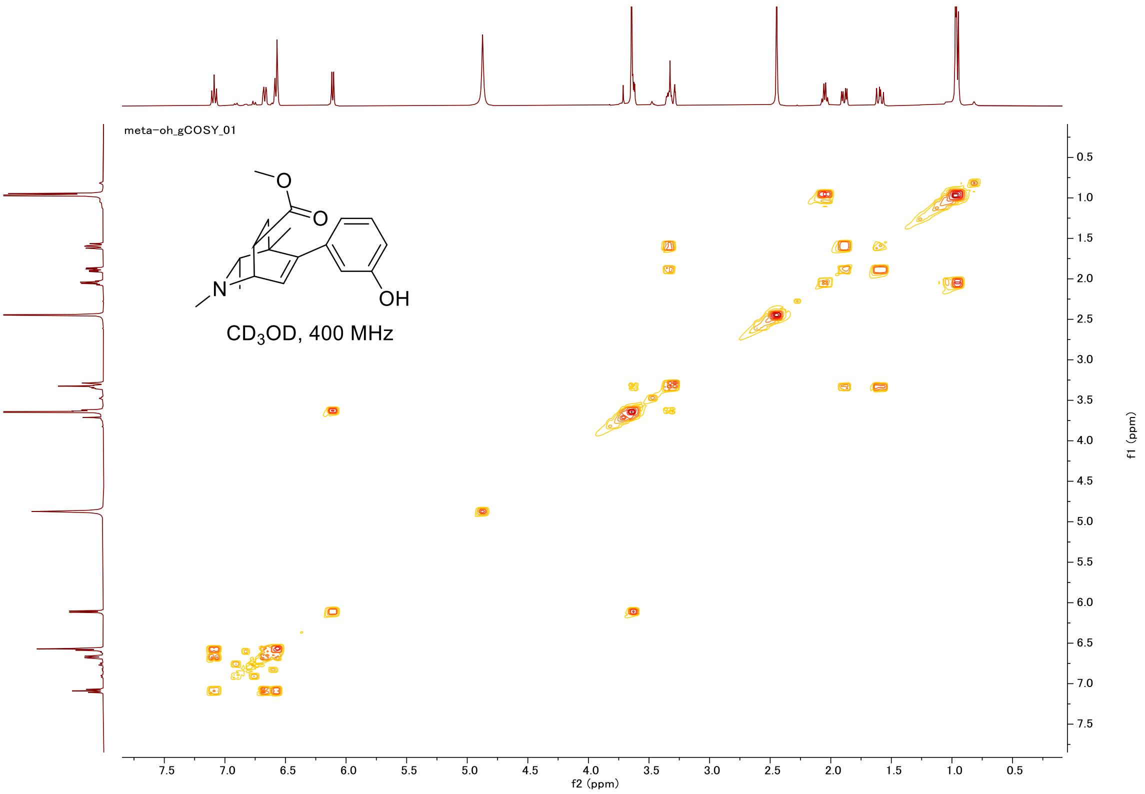
meta-oh_gCOSY_01
O
O
N
OH
CD3OD, 400 MHz
f2 (ppm)
f1 (ppm)

#### HSQC

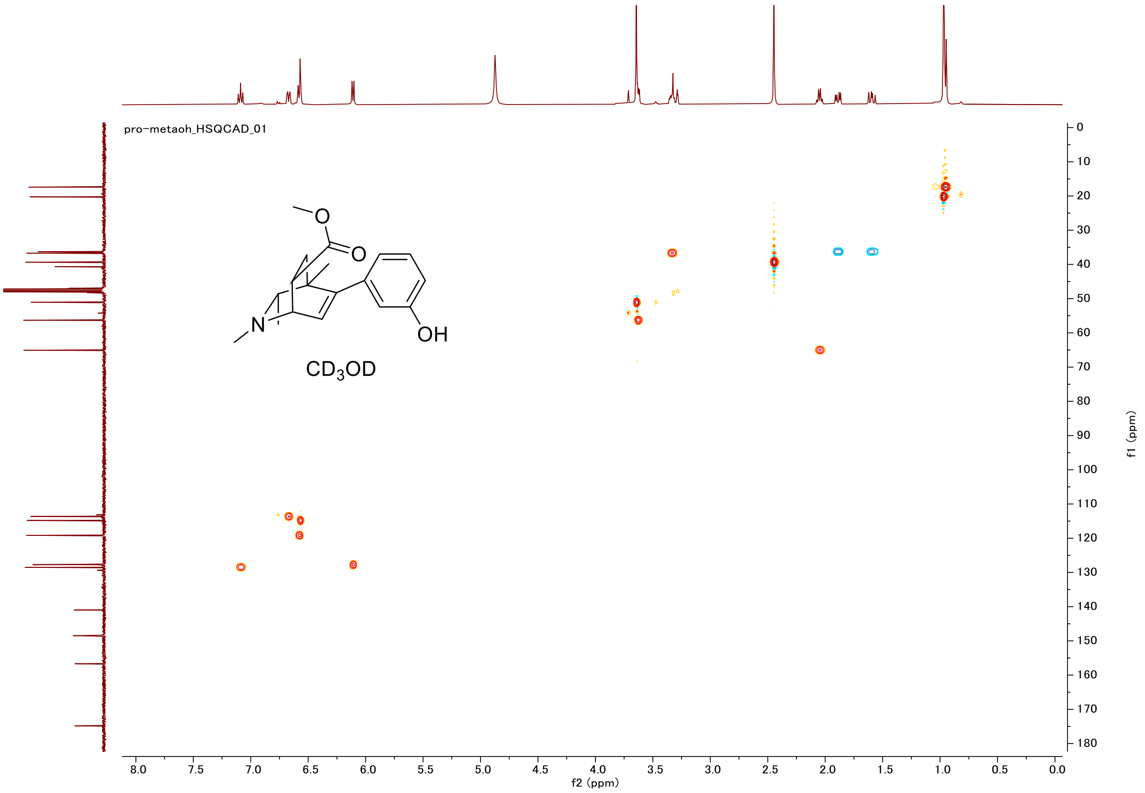
pro-metaoh_HSQCAD_01
O
O
N
OH
CD3OD
f2 (ppm)
f1 (ppm)

#### Proton (#017)

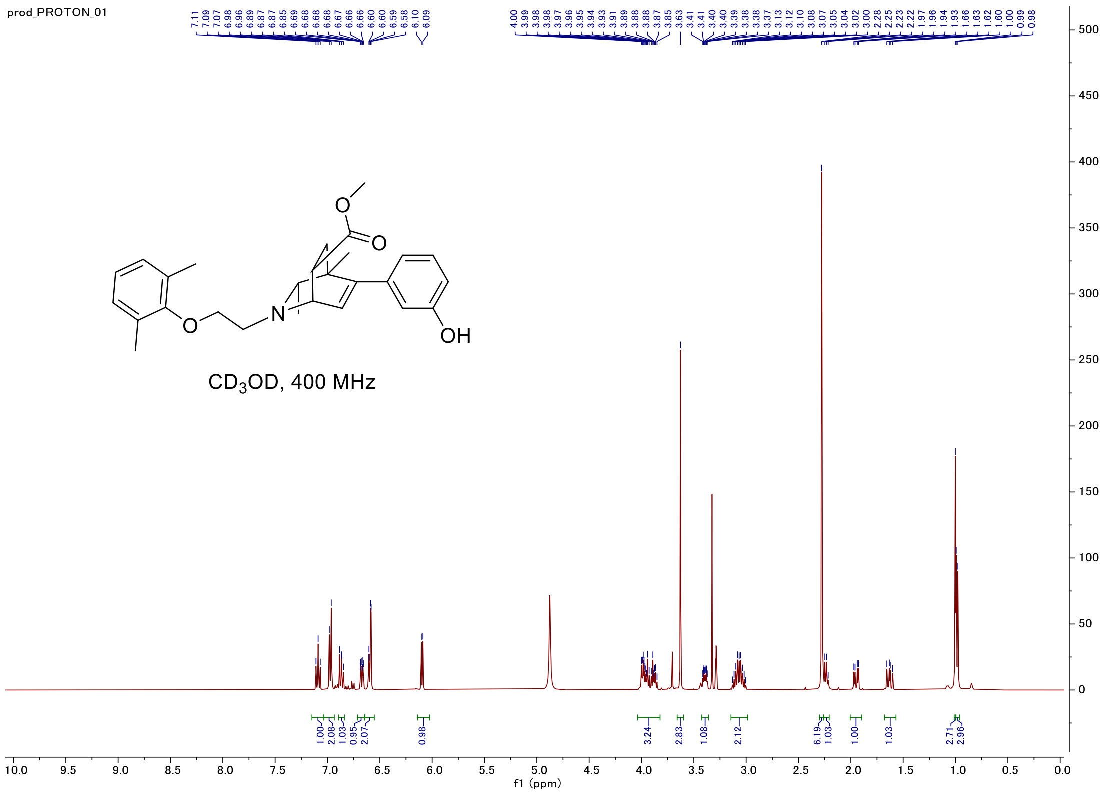


#### Carbon

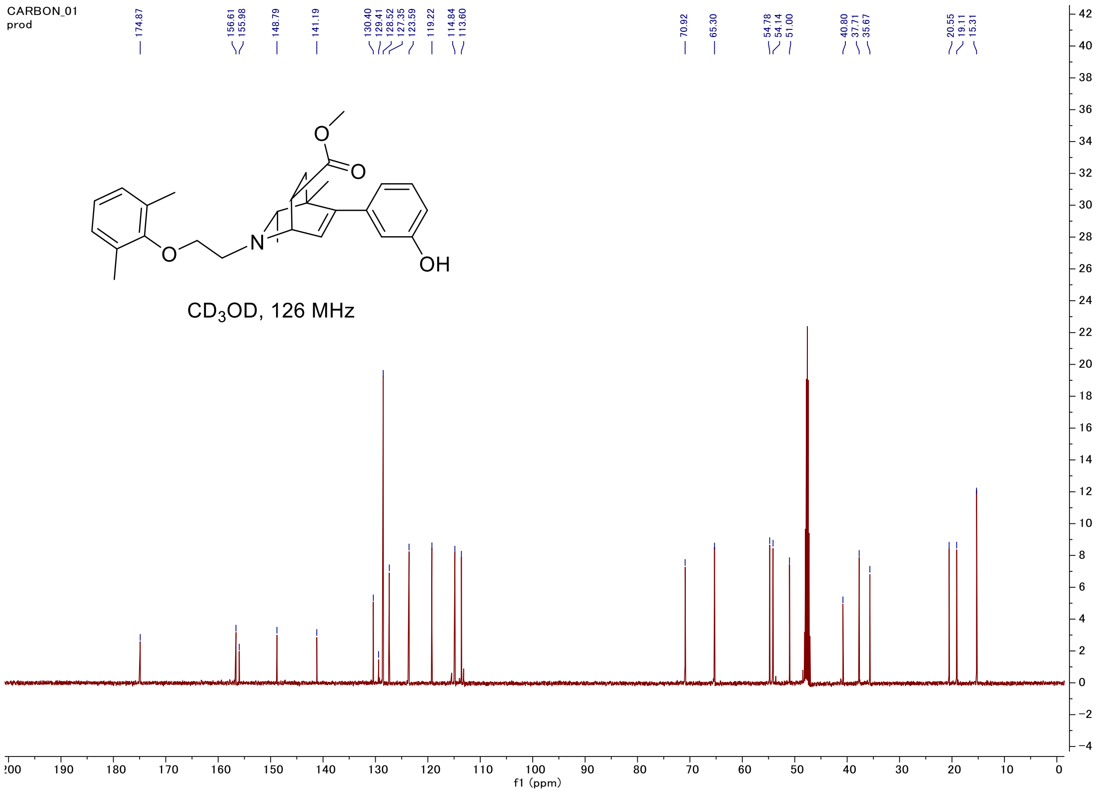

#### C-H COSY

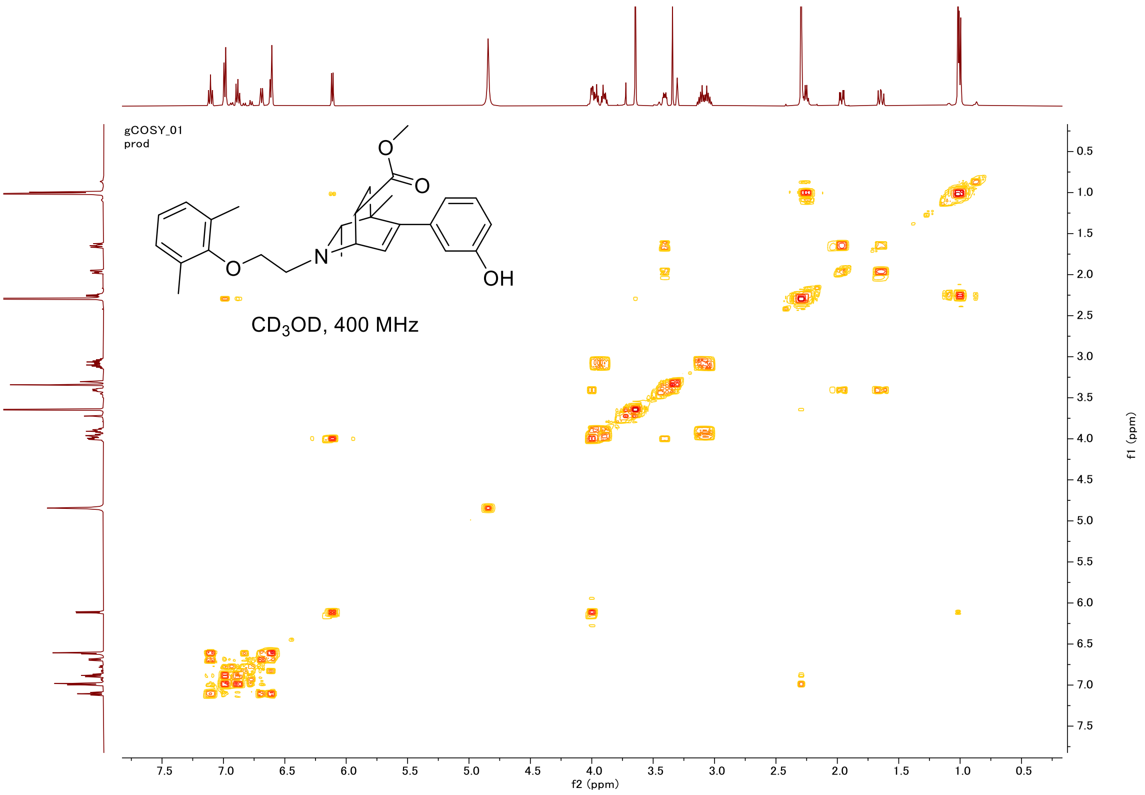

gCOSY_01
prod
OH
O
O
O
N
CD$_3$OD, 400 MHz
f2 (ppm)
f1 (ppm)


#### HSQC

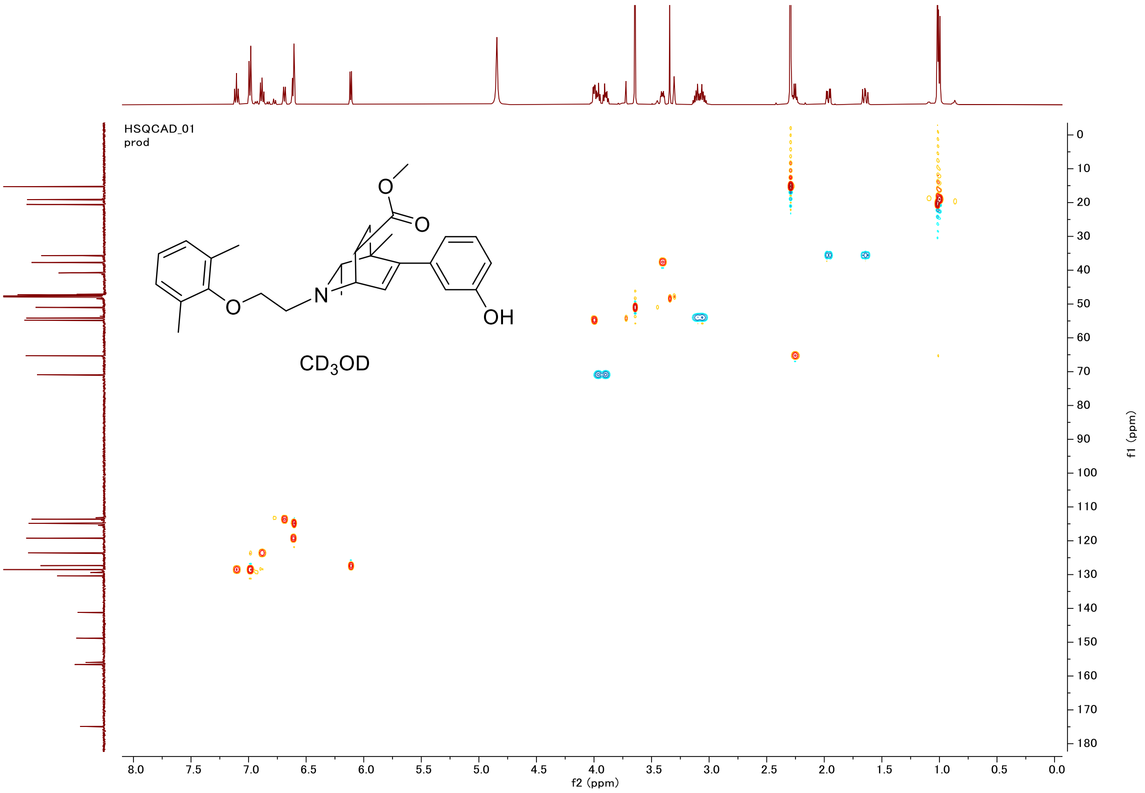

HSQCAD_01
prod
OH
O
O
O
N
CD$_3$OD
f2 (ppm)
f1 (ppm)

#### Proton (#018)

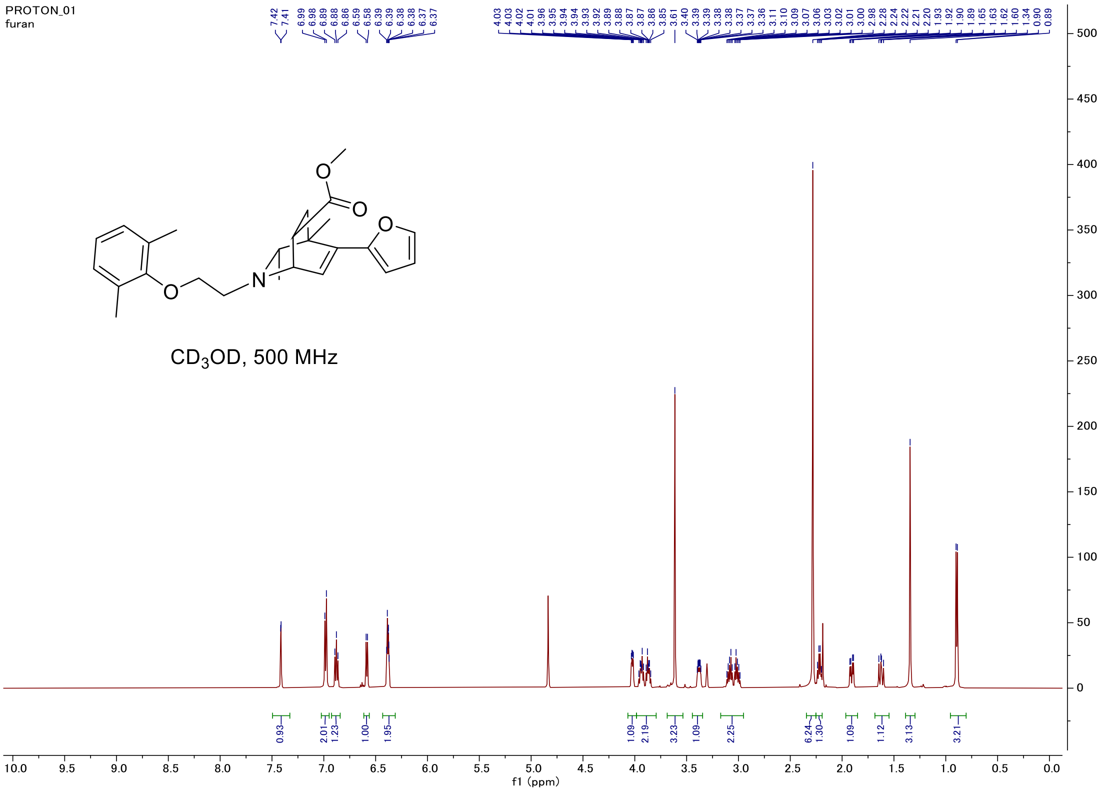


#### Carbon

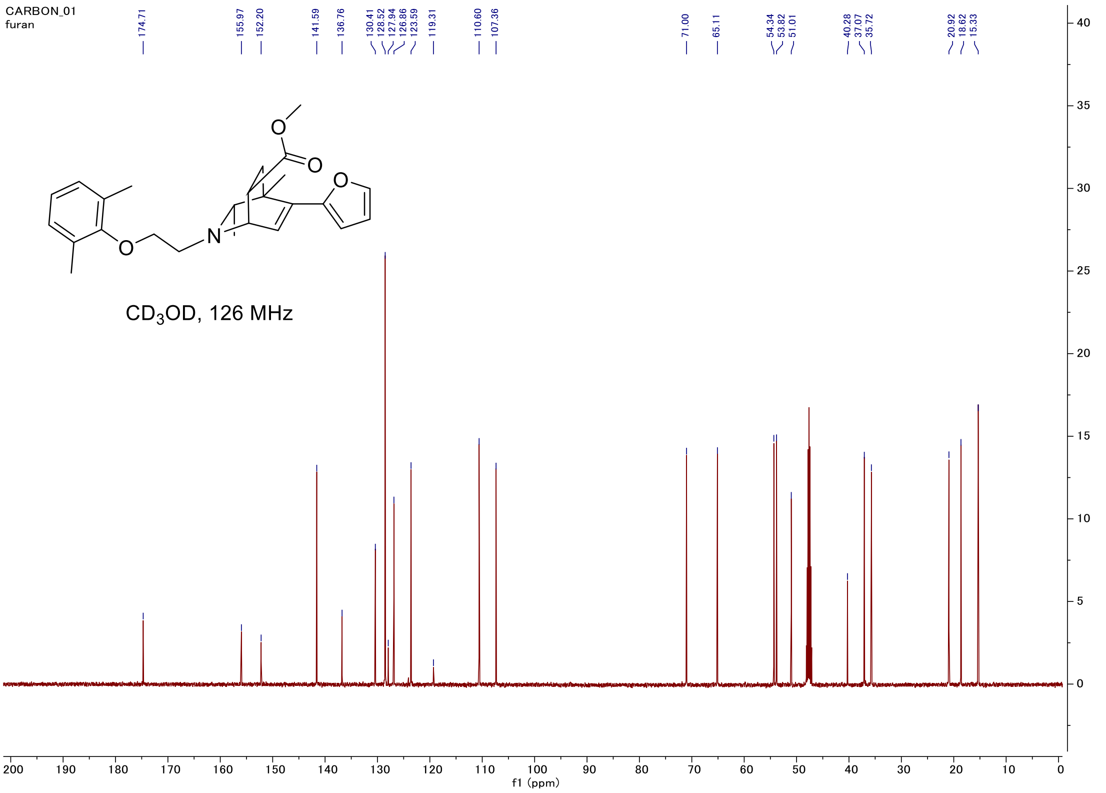

#### C-H COSY

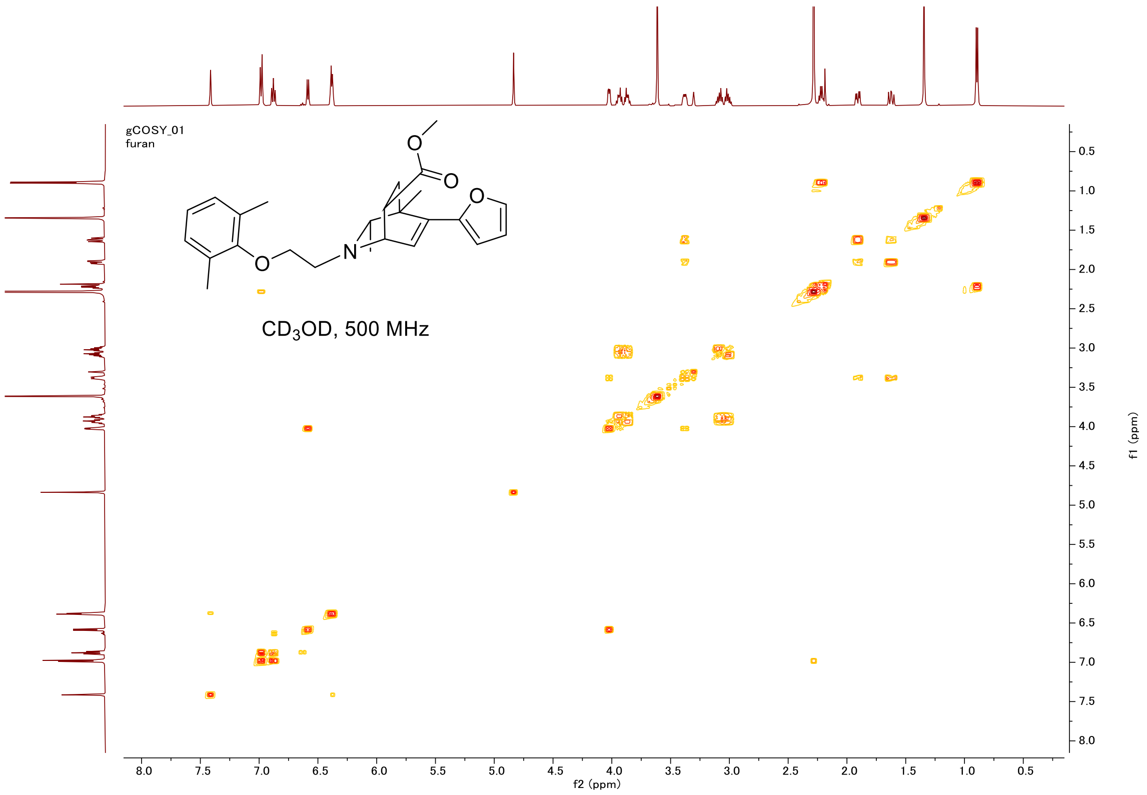
gCOSY_01
furan
CD3OD, 500 MHz
f2 (ppm)
f1 (ppm)

#### HSQC

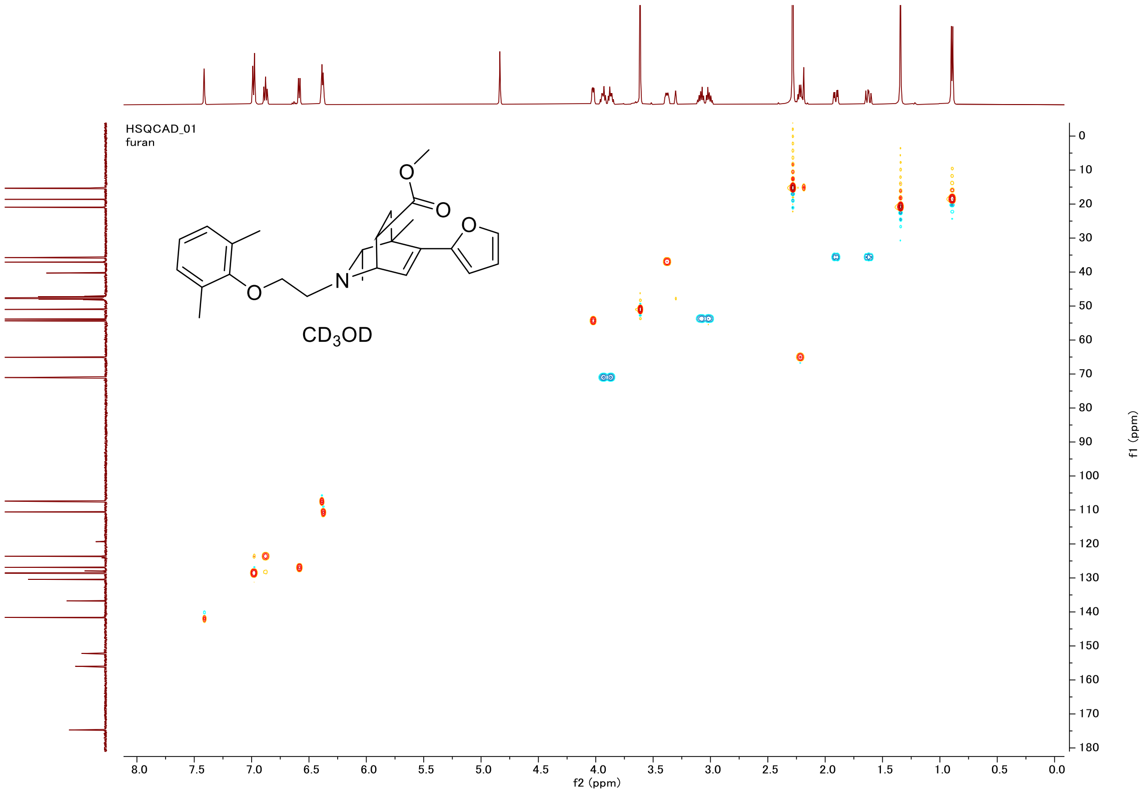
HSQCAD_01
furan
CD3OD
f2 (ppm)
f1 (ppm)

#### Proton (#019)

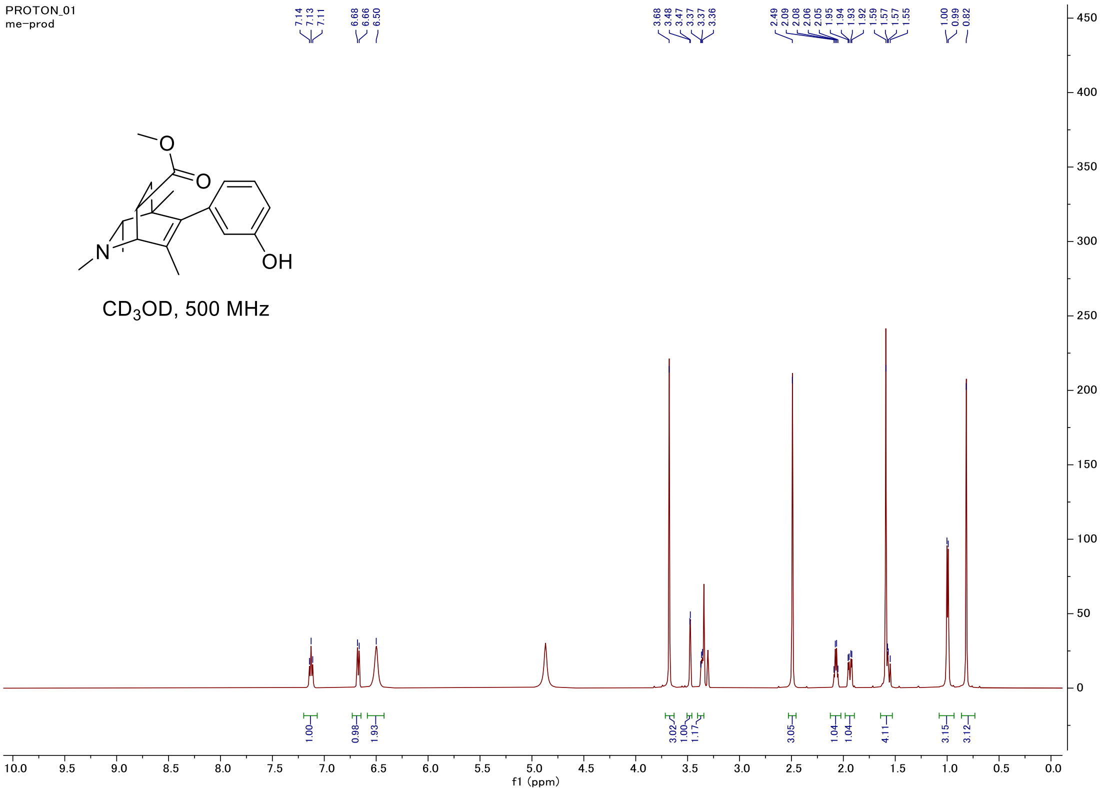


#### Carbon

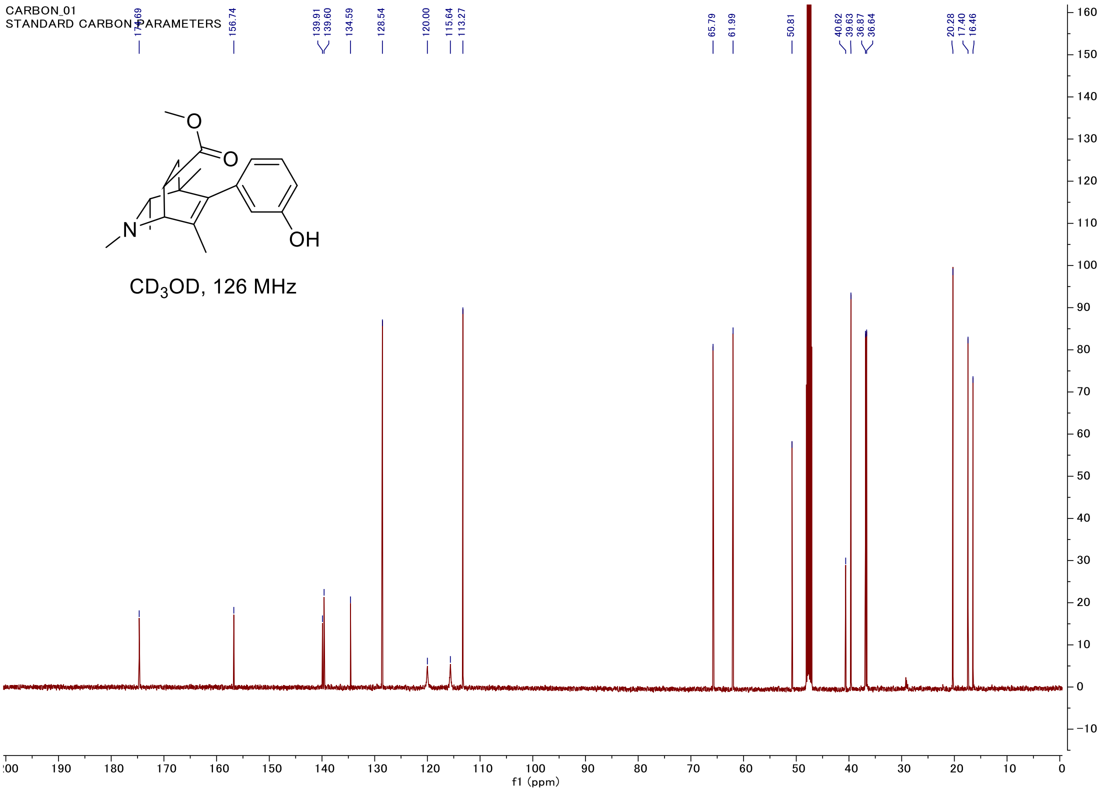

#### C-H COSY

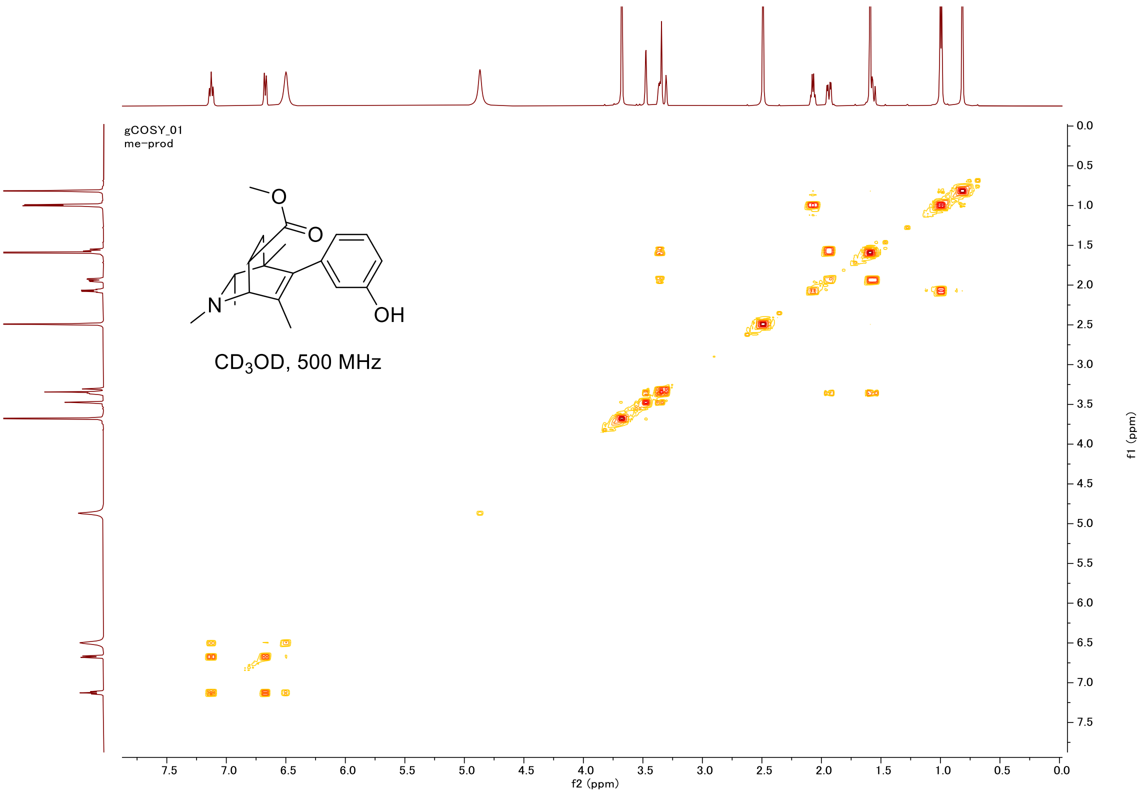
gCOSY_01
me-prod
O
O
N
OH
CD3OD, 500 MHz
f2 (ppm)
f1 (ppm)

#### HSQC

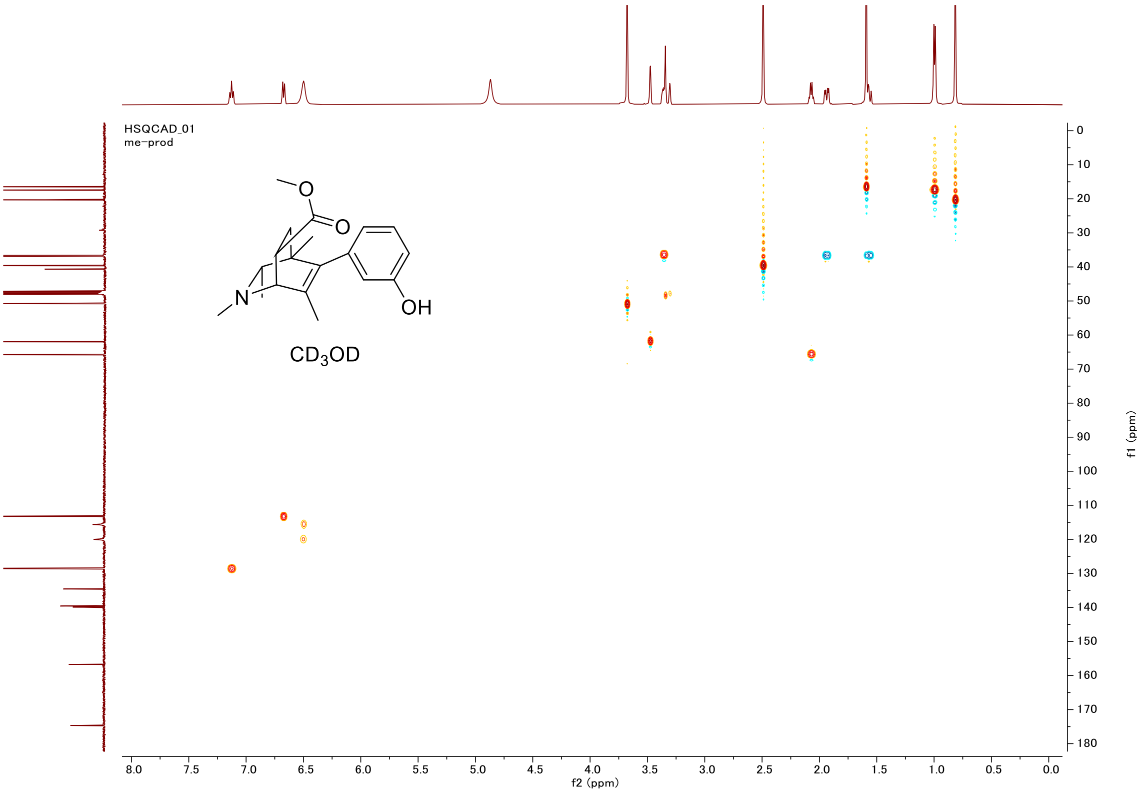
HSQCAD_01
me-prod
O
O
N
OH
CD3OD
f2 (ppm)
f1 (ppm)

#### Proton (#020)

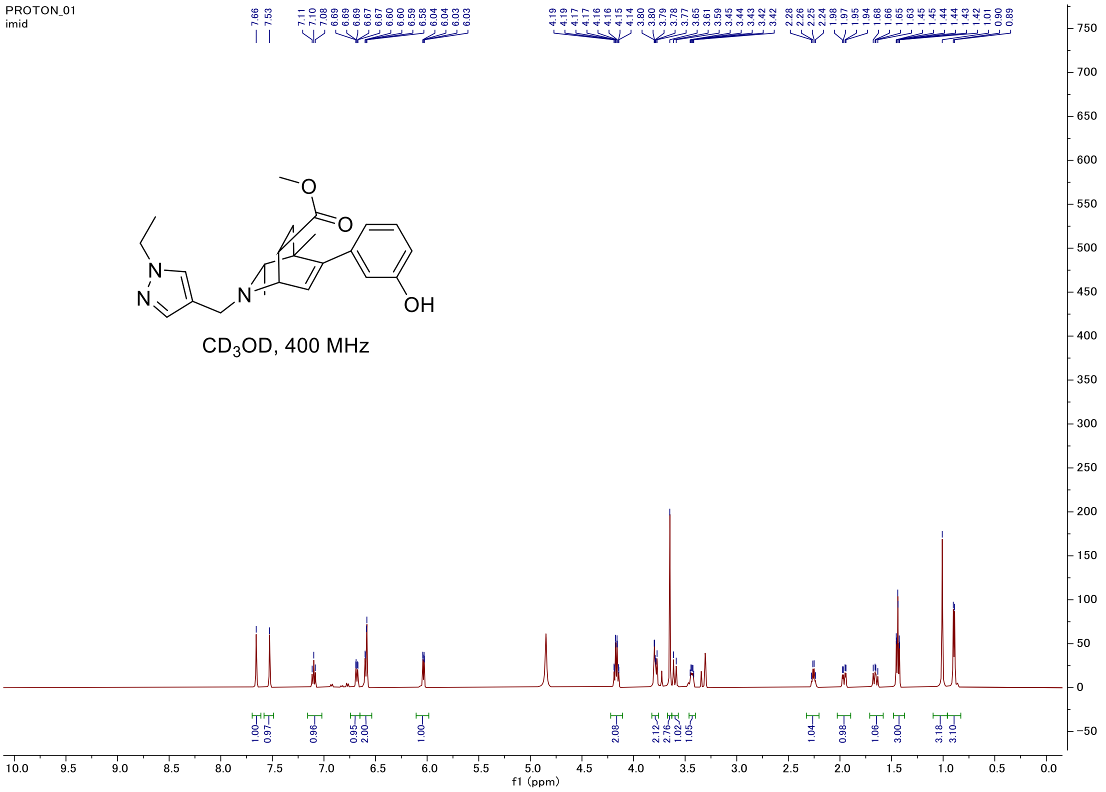


#### Carbon

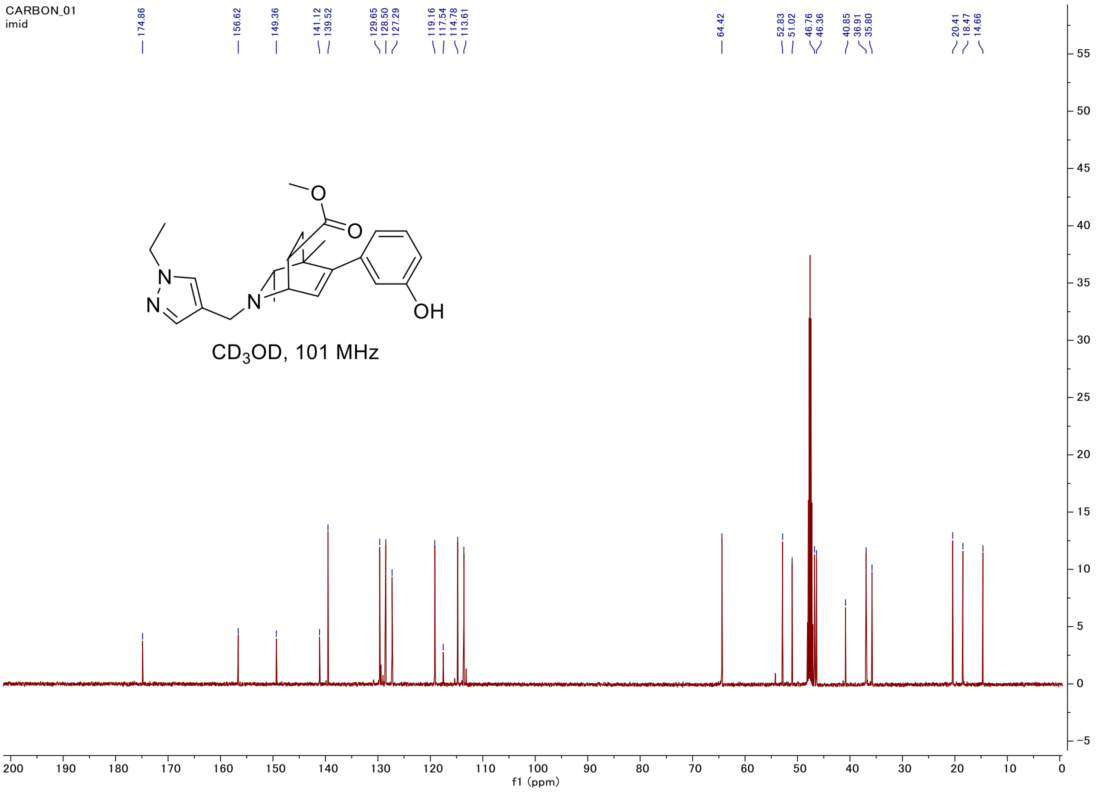

#### C-H COSY

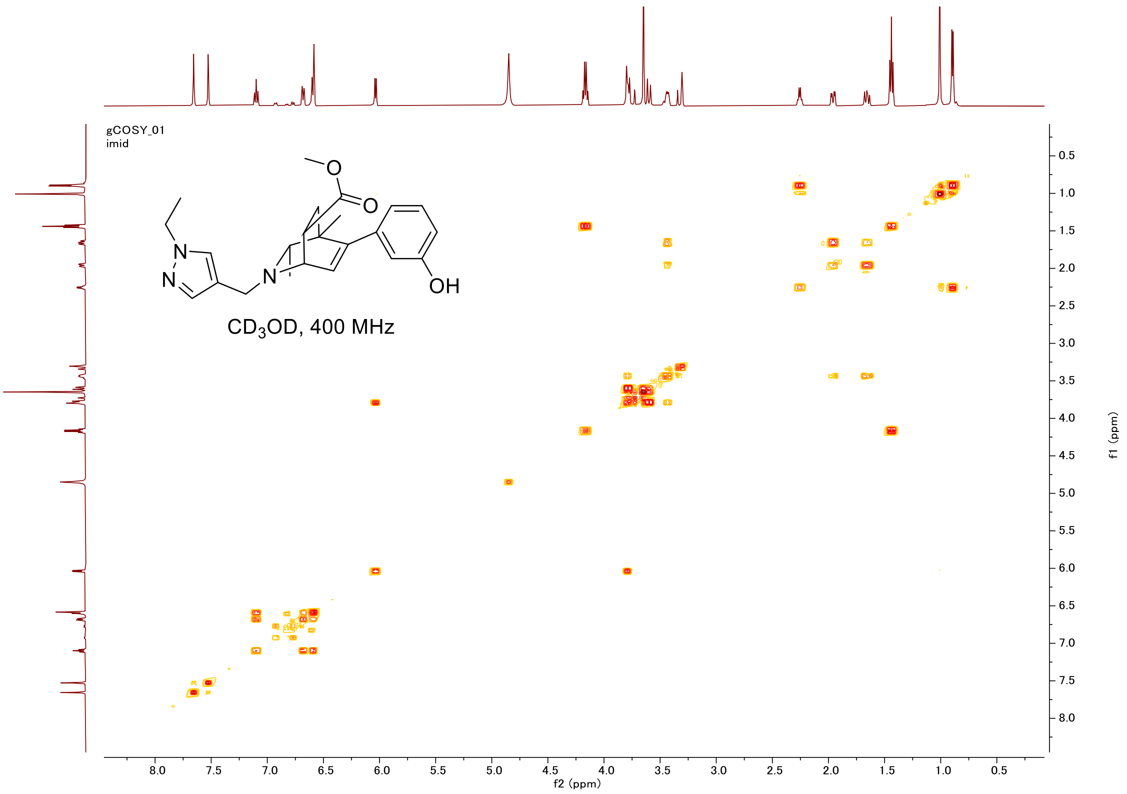

gCOSY_01
imid
O
O
N
N
N
OH
CD3OD, 400 MHz
f2 (ppm)
f1 (ppm)


#### HSQC

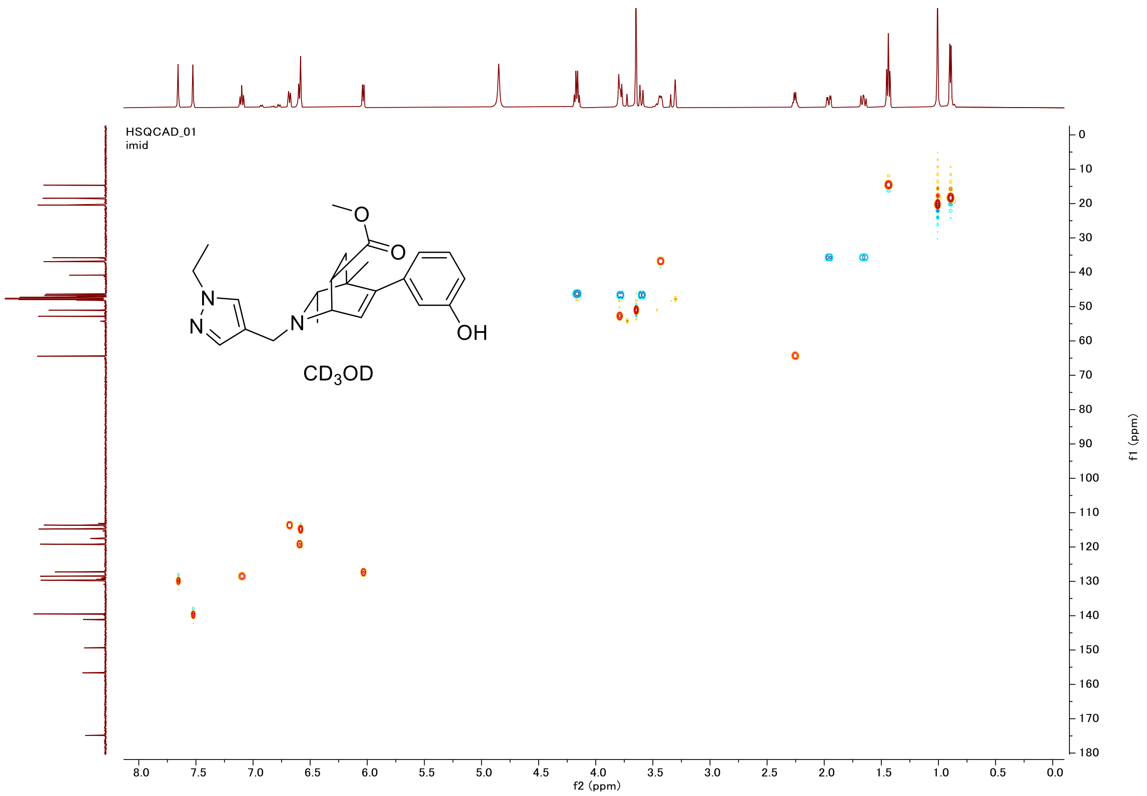

HSQCAD_01
imid
O
O
N
N
N
OH
CD3OD
f2 (ppm)
f1 (ppm)

#### Proton (#021)

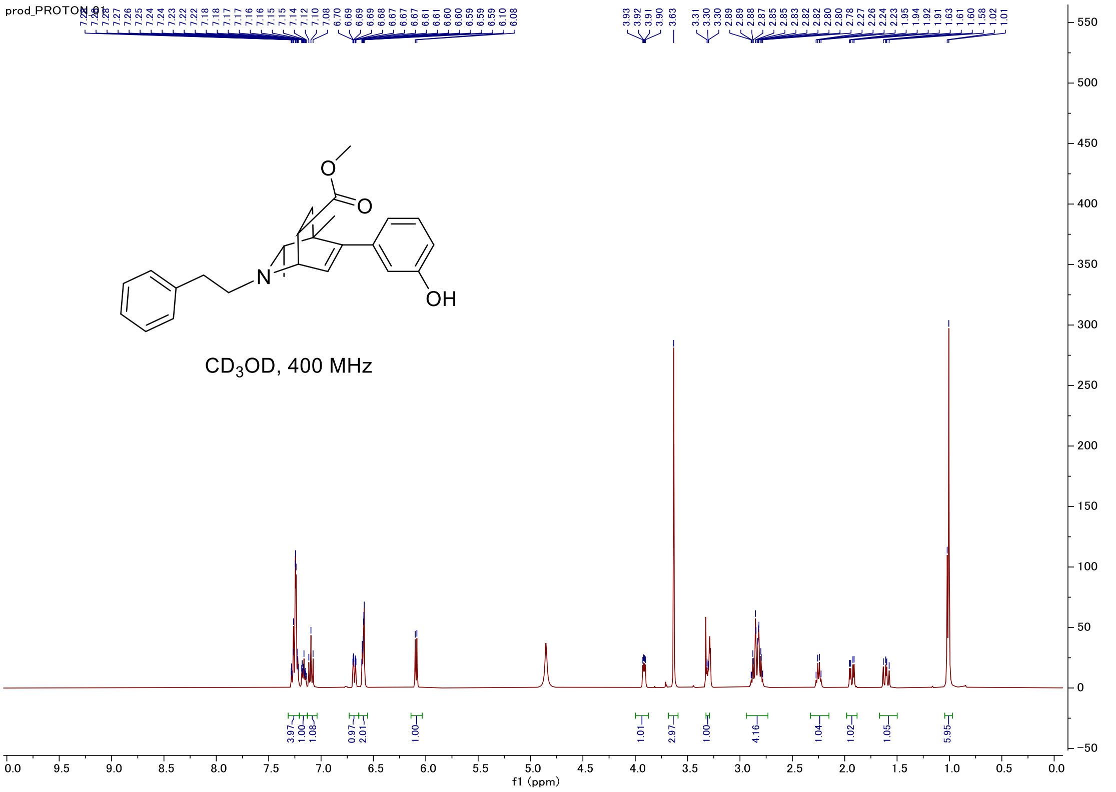


#### Carbon

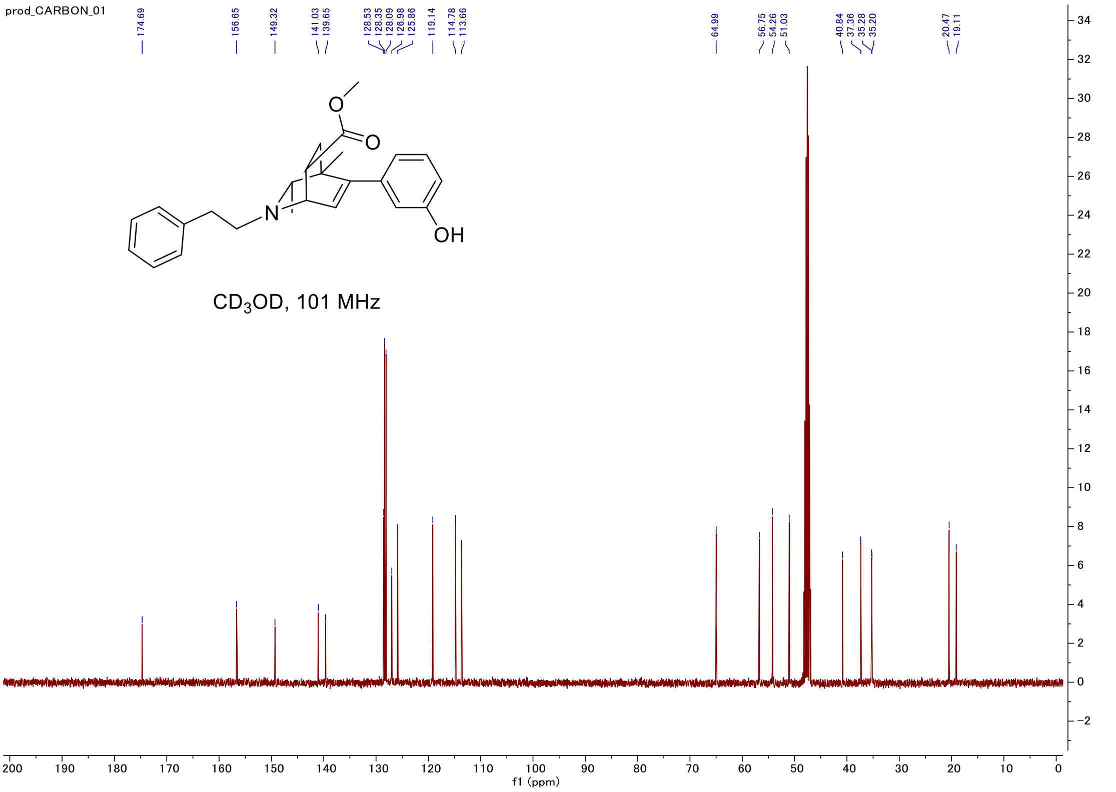

#### C-H COSY

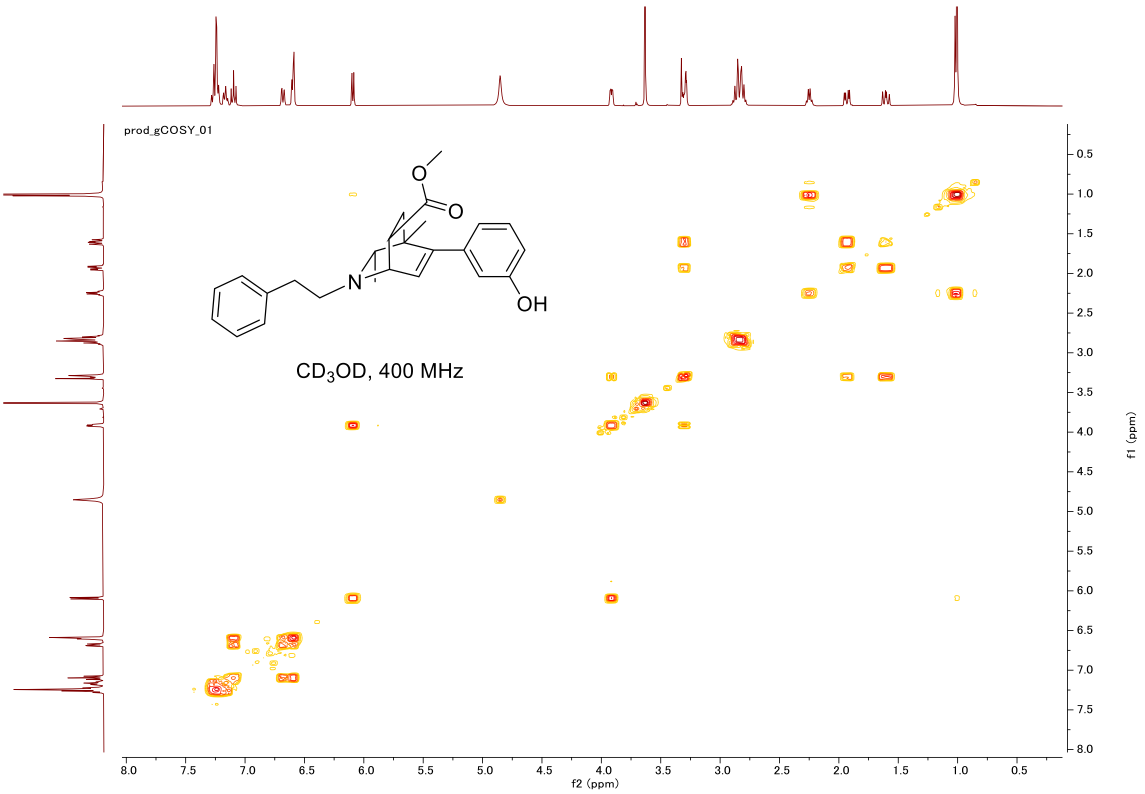
prod_gCOSY_01
O
O
N
OH
CD3OD, 400 MHz
f2 (ppm)
f1 (ppm)

#### HSQC

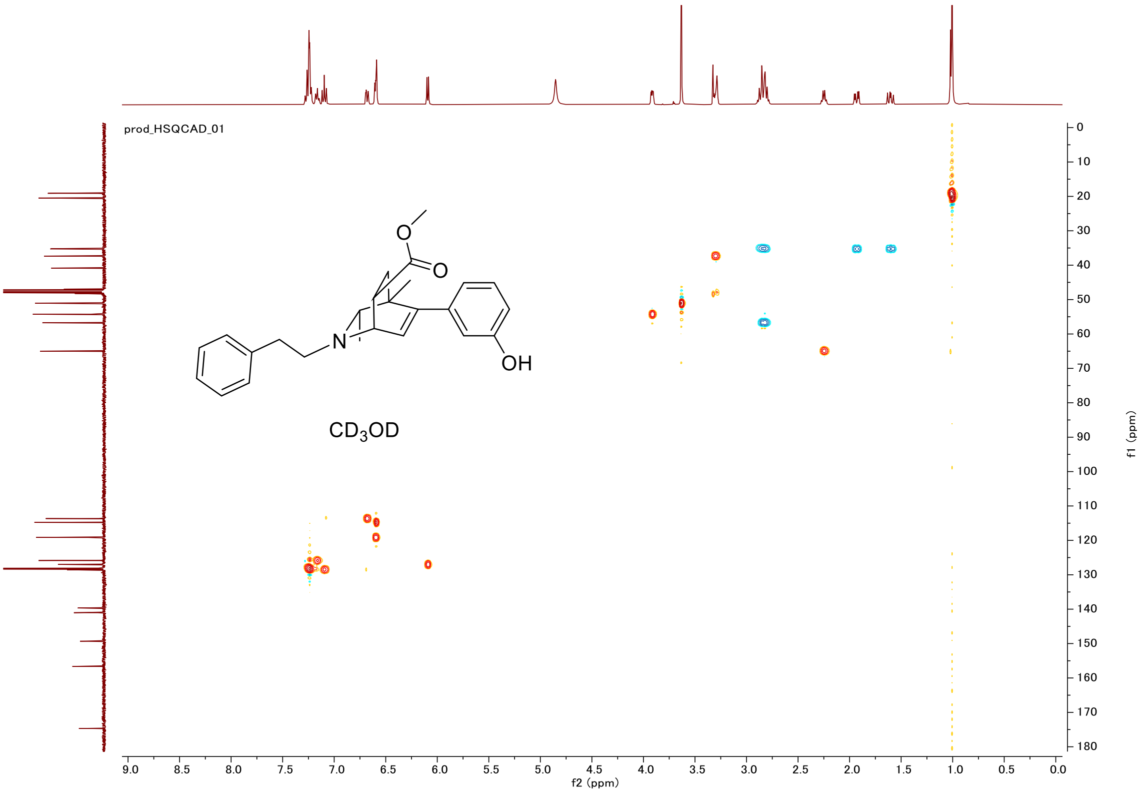
prod_HSQCAD_01
O
O
N
OH
CD3OD
f2 (ppm)
f1 (ppm)

#### Proton (#022-major)

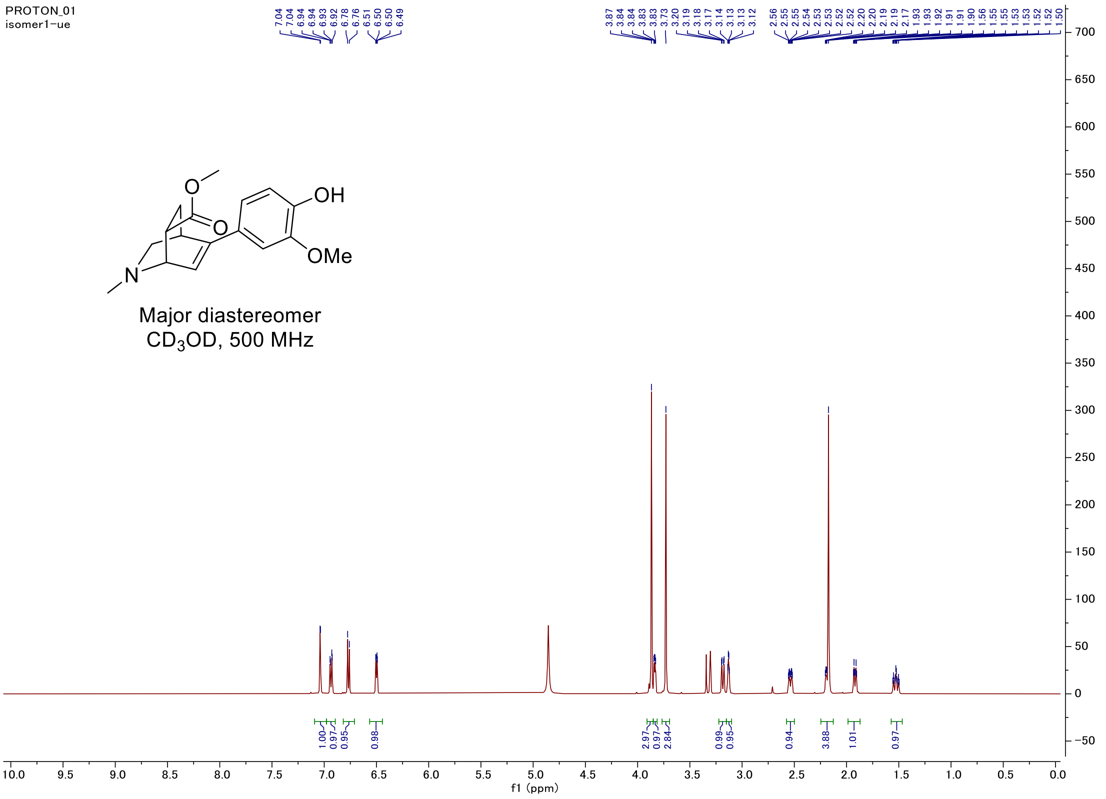


#### Carbon

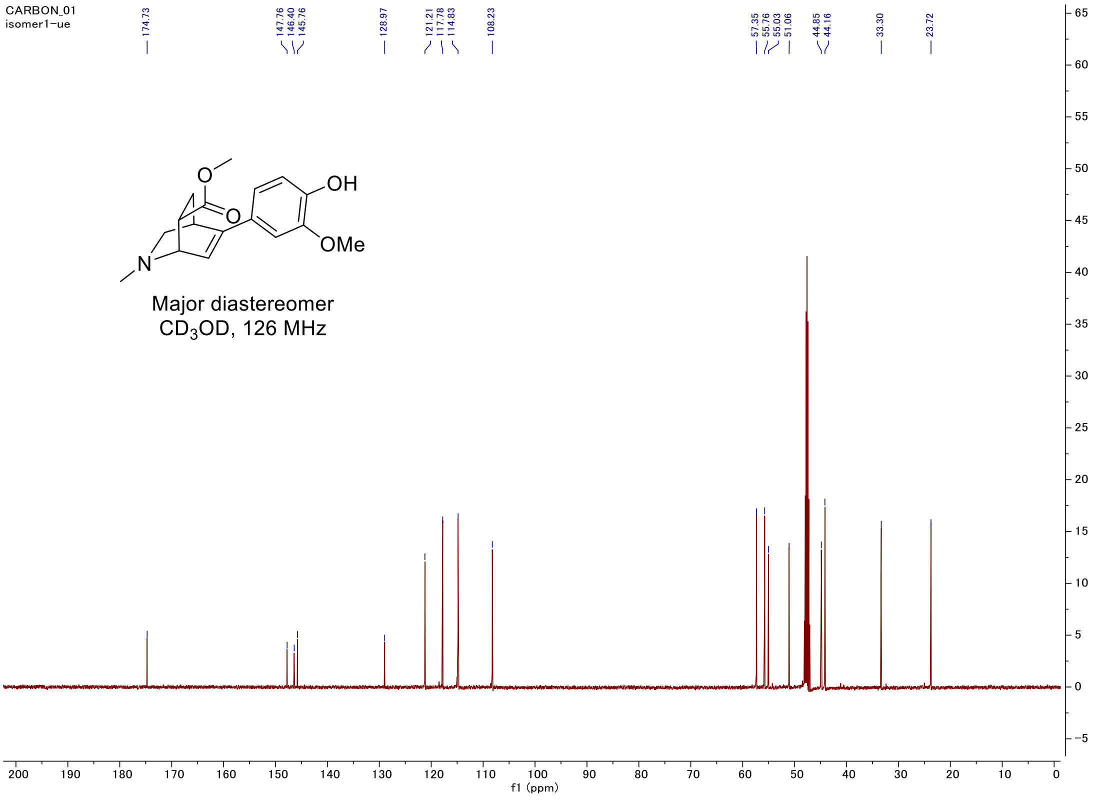

#### C-H COSY

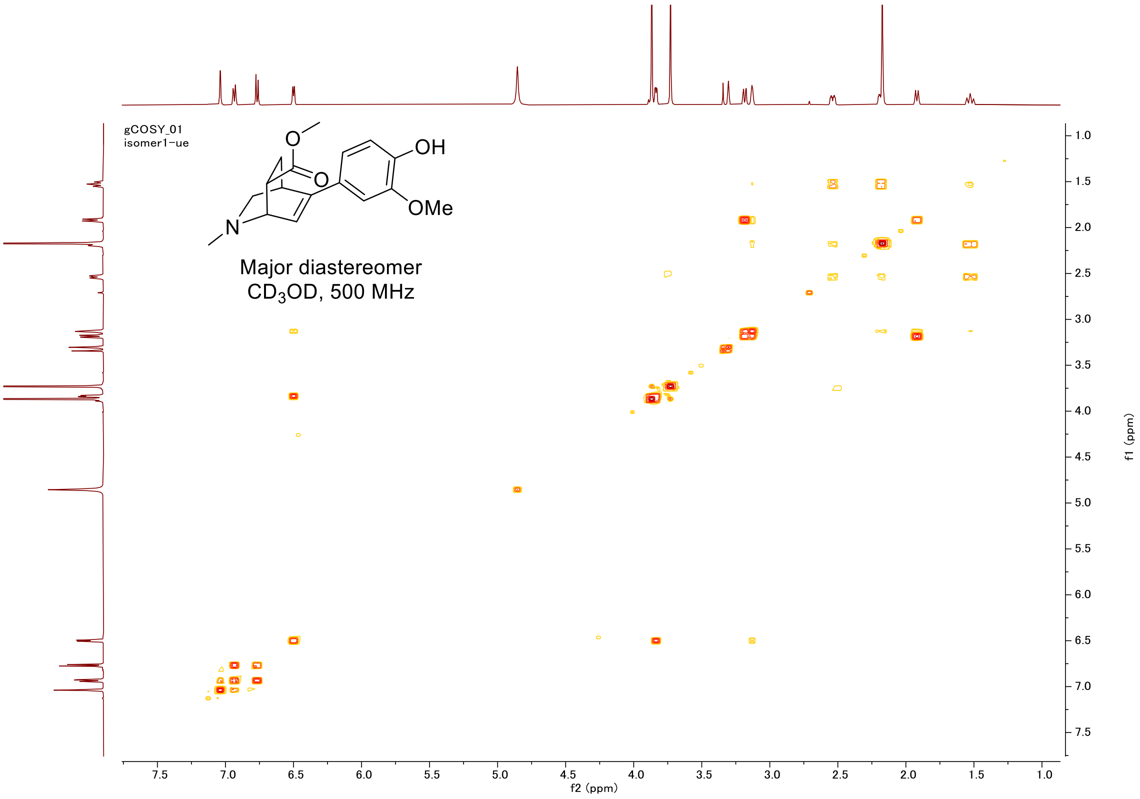
gCOSY_01
isomer1-ue
OH
OMe
N
O
Major diastereomer
CD3OD, 500 MHz
f2 (ppm)
f1 (ppm)

#### HSQC

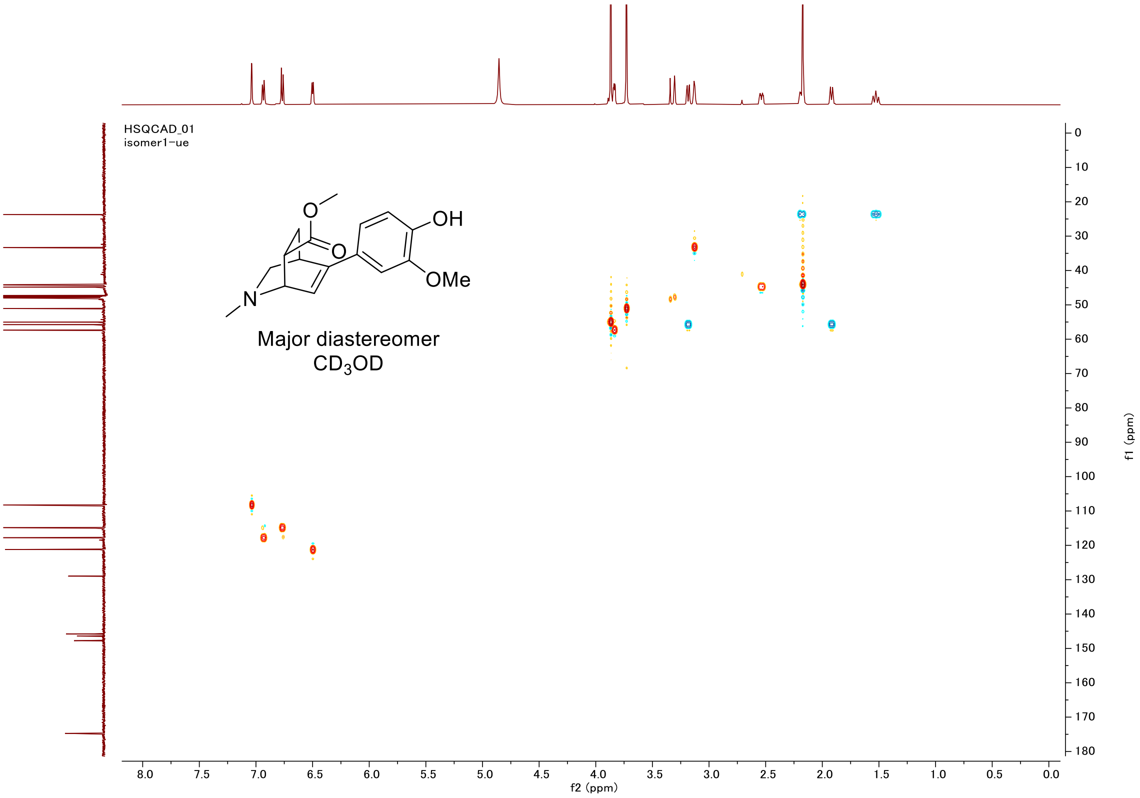
HSQCAD_01
isomer1-ue
OH
OMe
N
O
Major diastereomer
CD3OD
f2 (ppm)
f1 (ppm)

#### Proton (#022-minor)

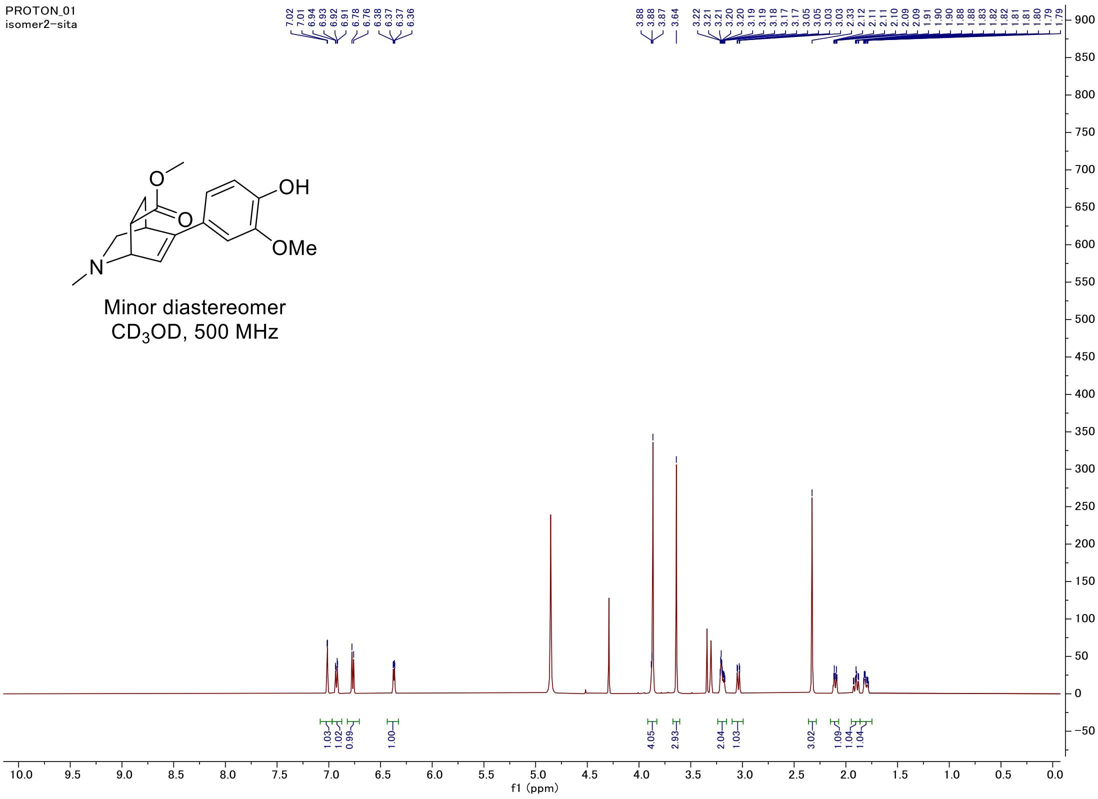


#### Carbon

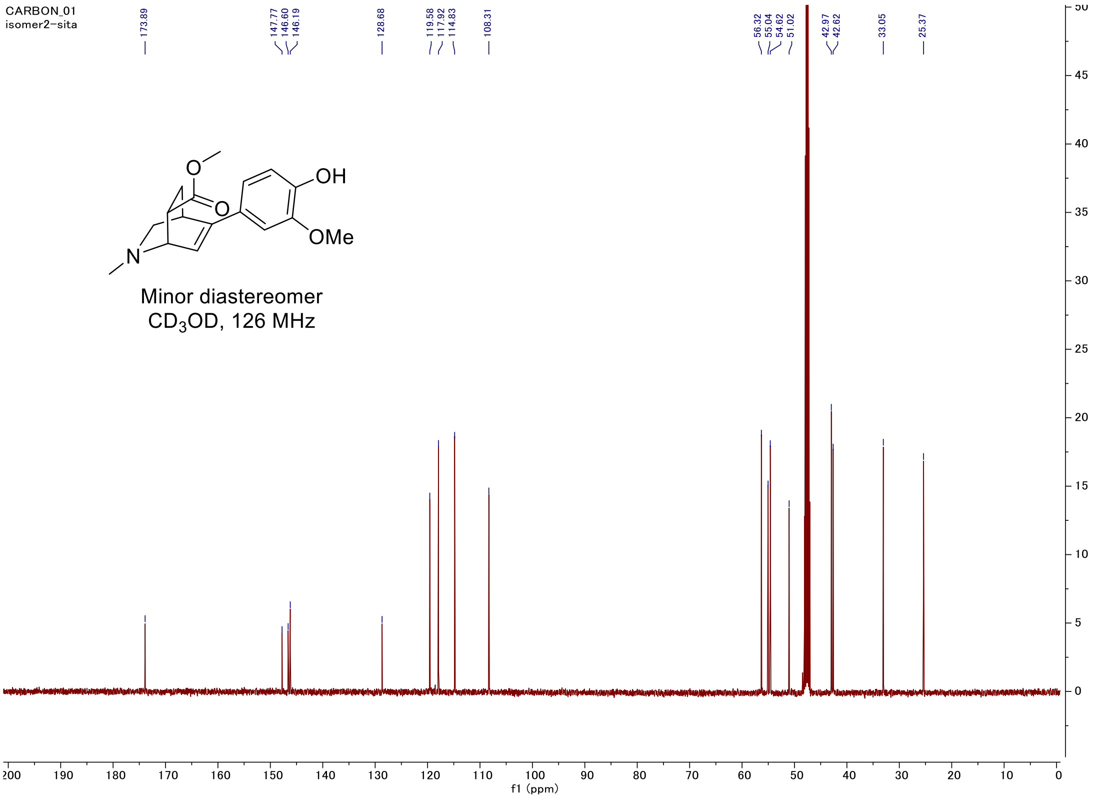

#### C-H COSY

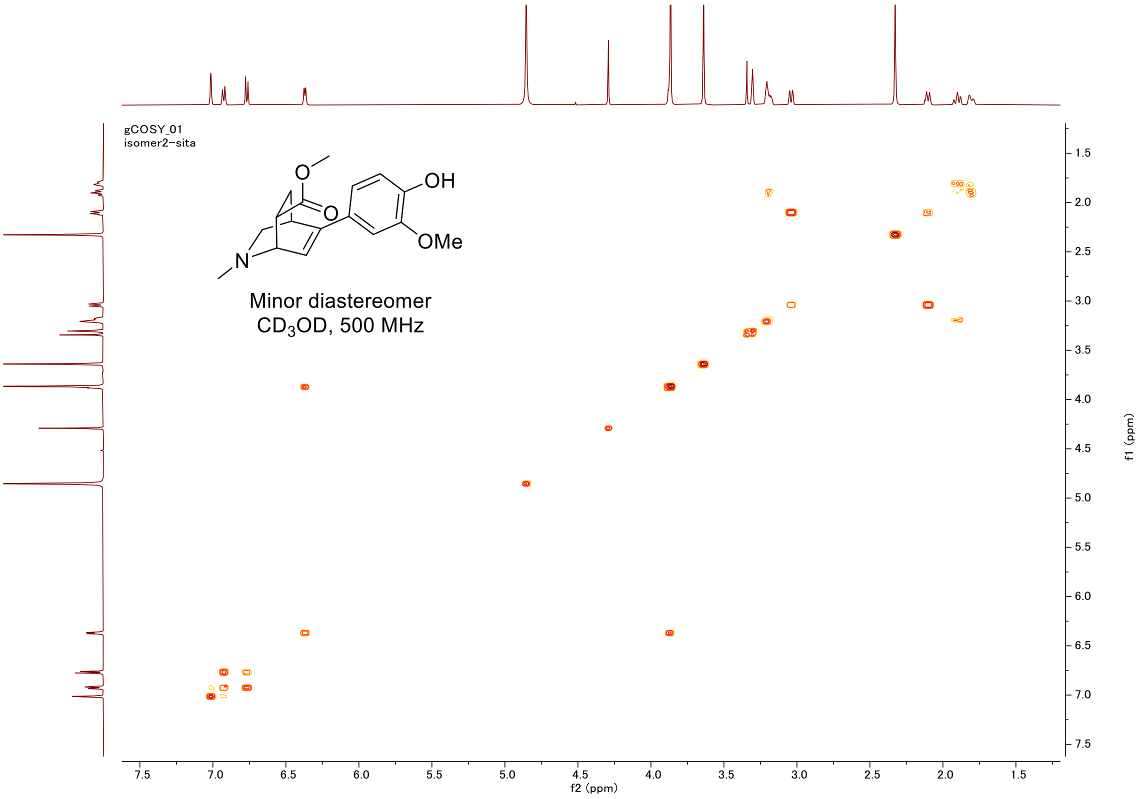
gCOSY_01
isomer2-sita
OH
OMe
O
O
N
Minor diastereomer
$CD_3OD$, 500 MHz
f2 (ppm)
f1 (ppm)

#### HSQC

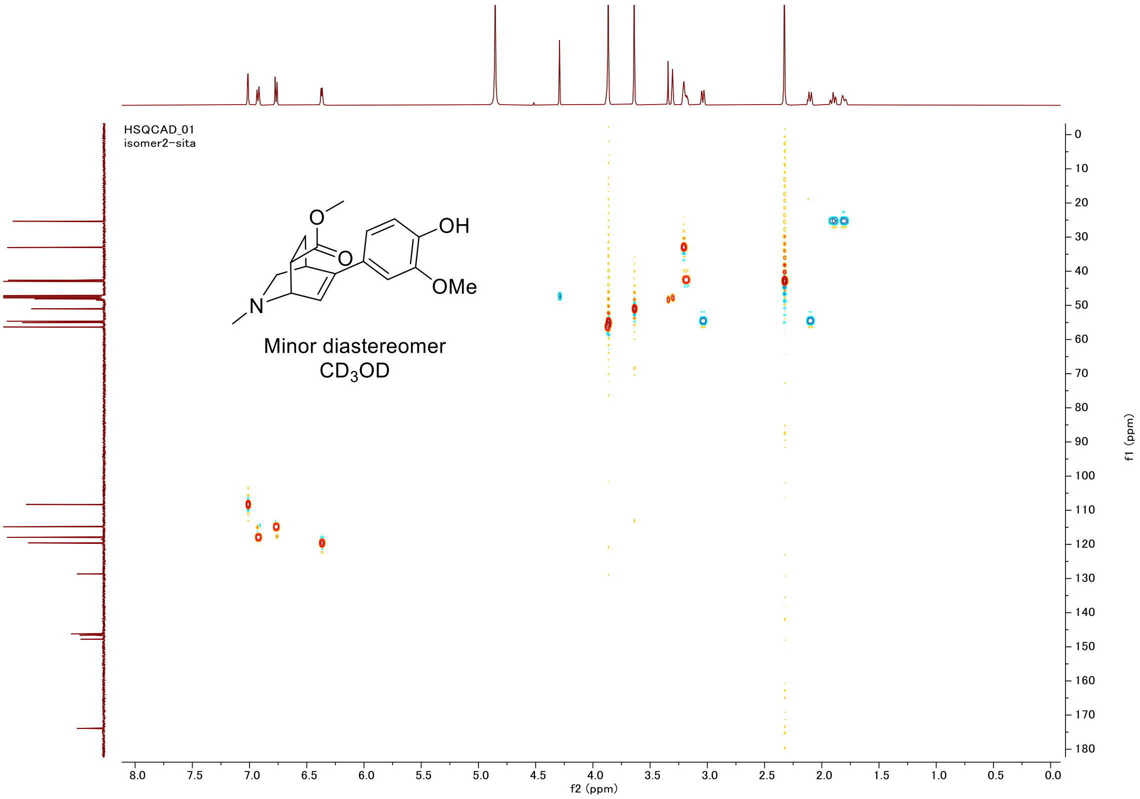
HSQCAD_01
isomer2-sita
OH
OMe
O
O
N
Minor diastereomer
$CD_3OD$
f2 (ppm)
f1 (ppm)

#### Proton (#023)

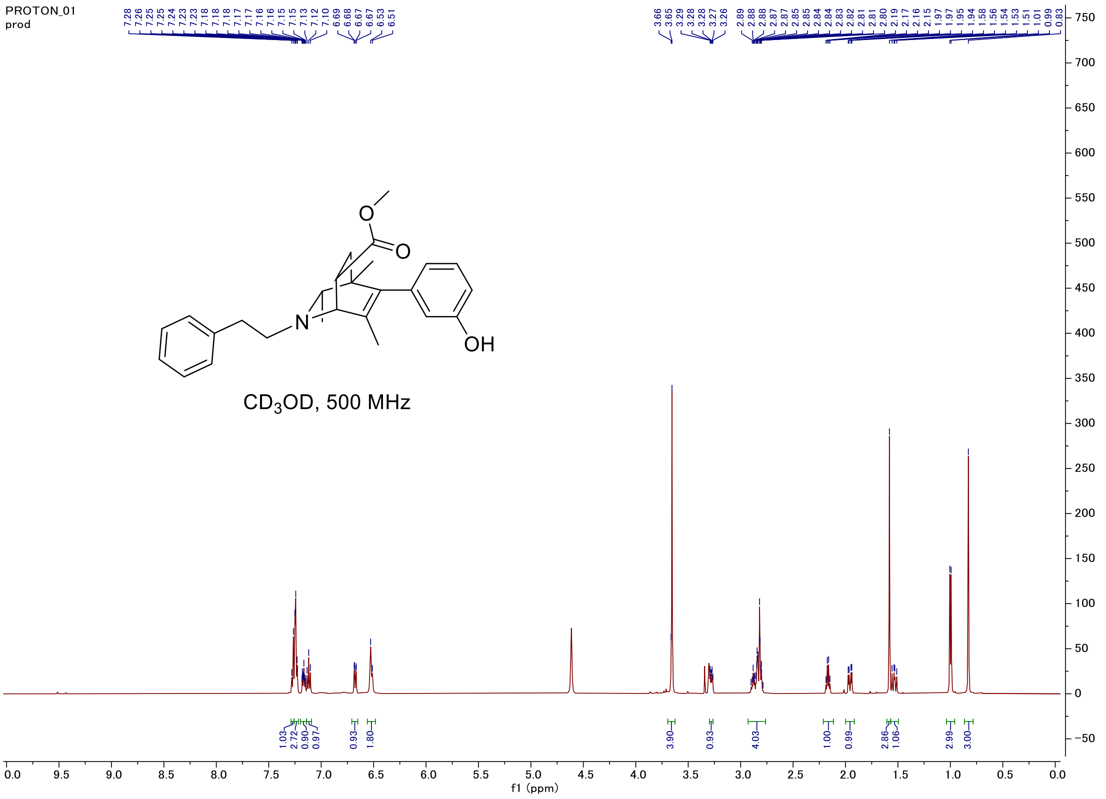


#### Carbon

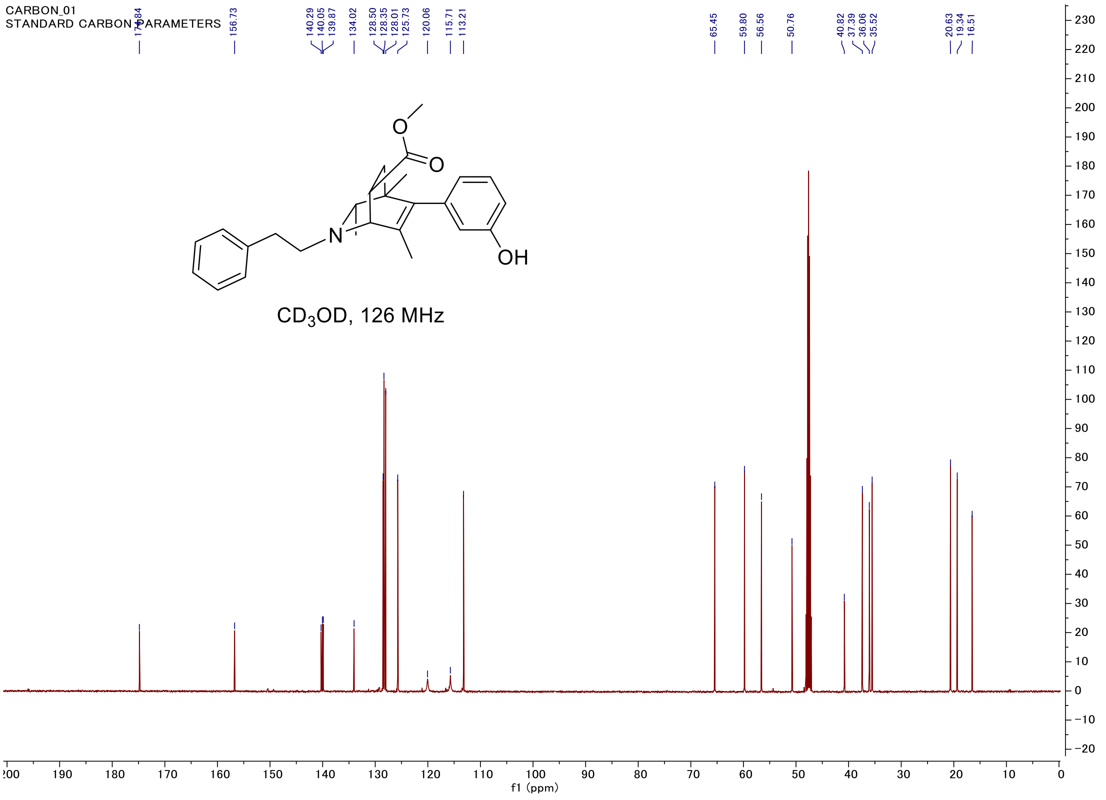

### C-H COSY

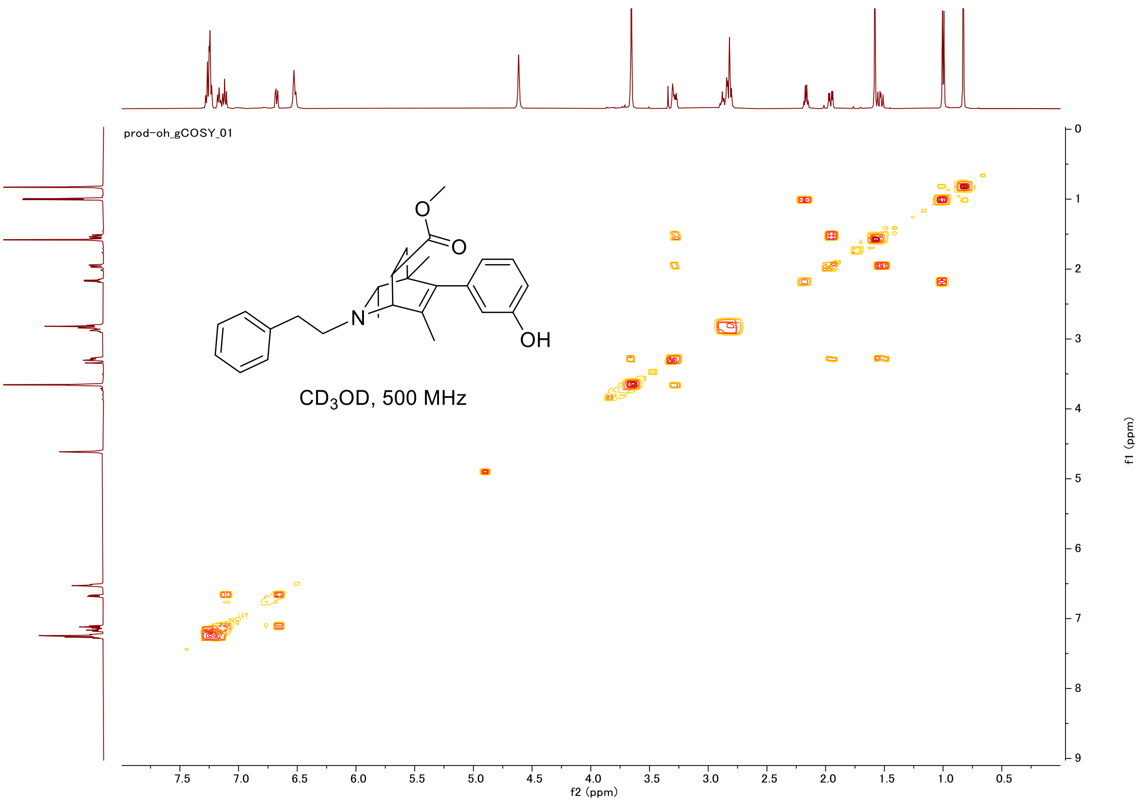
prod-oh_gCOSY_01
O
O
N
OH
CD$_3$OD, 500 MHz
f2 (ppm)
f1 (ppm)

### HSQC

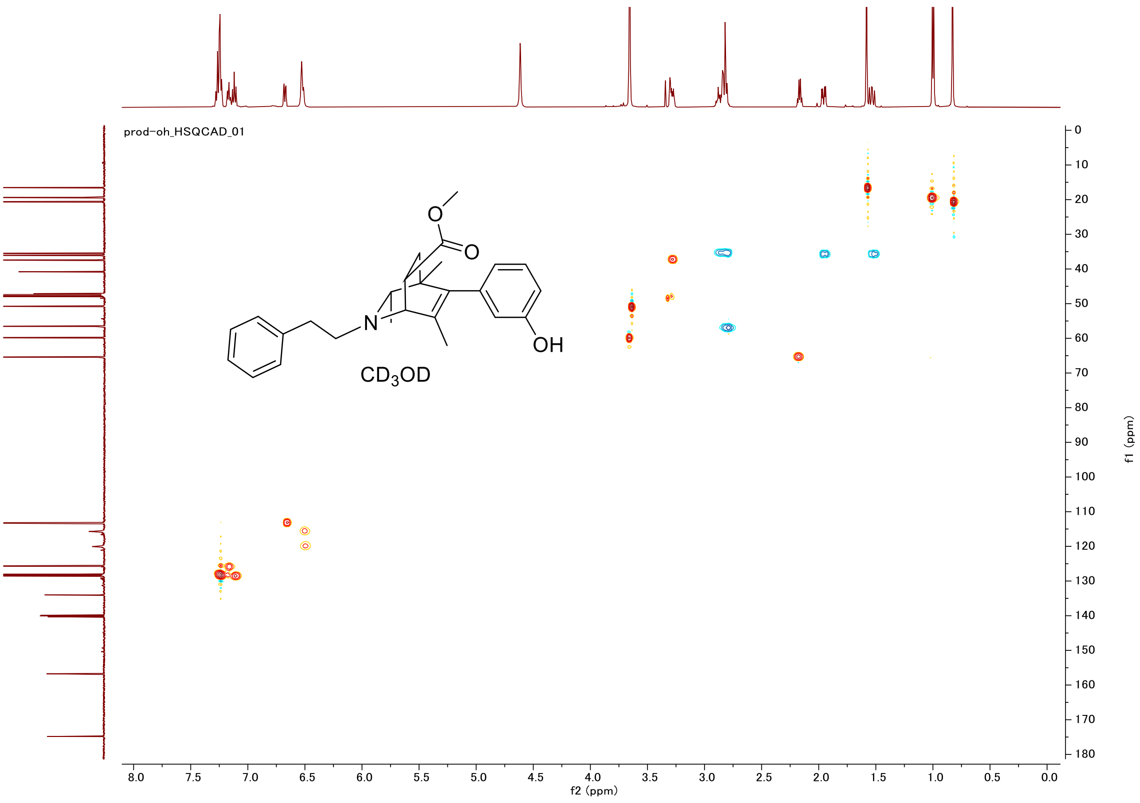
prod-oh_HSQCAD_01
O
O
N
OH
CD$_3$OD
f2 (ppm)
f1 (ppm)

#### Proton (#024)

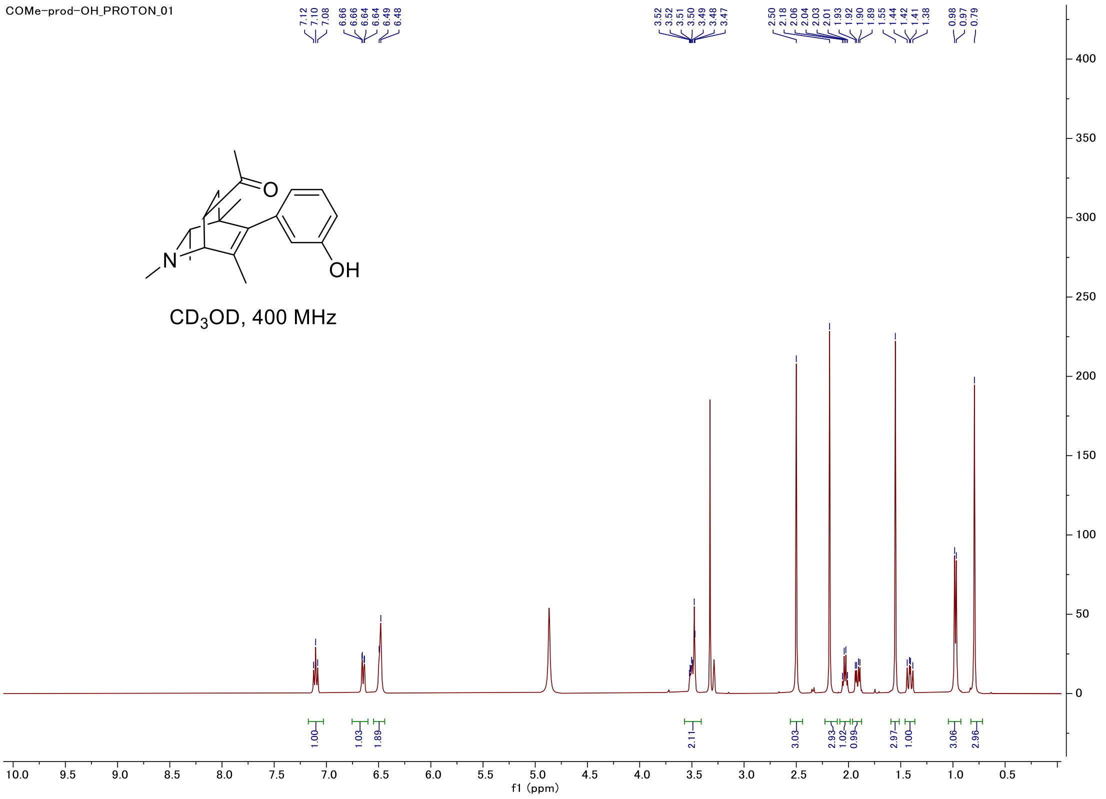


#### Carbon

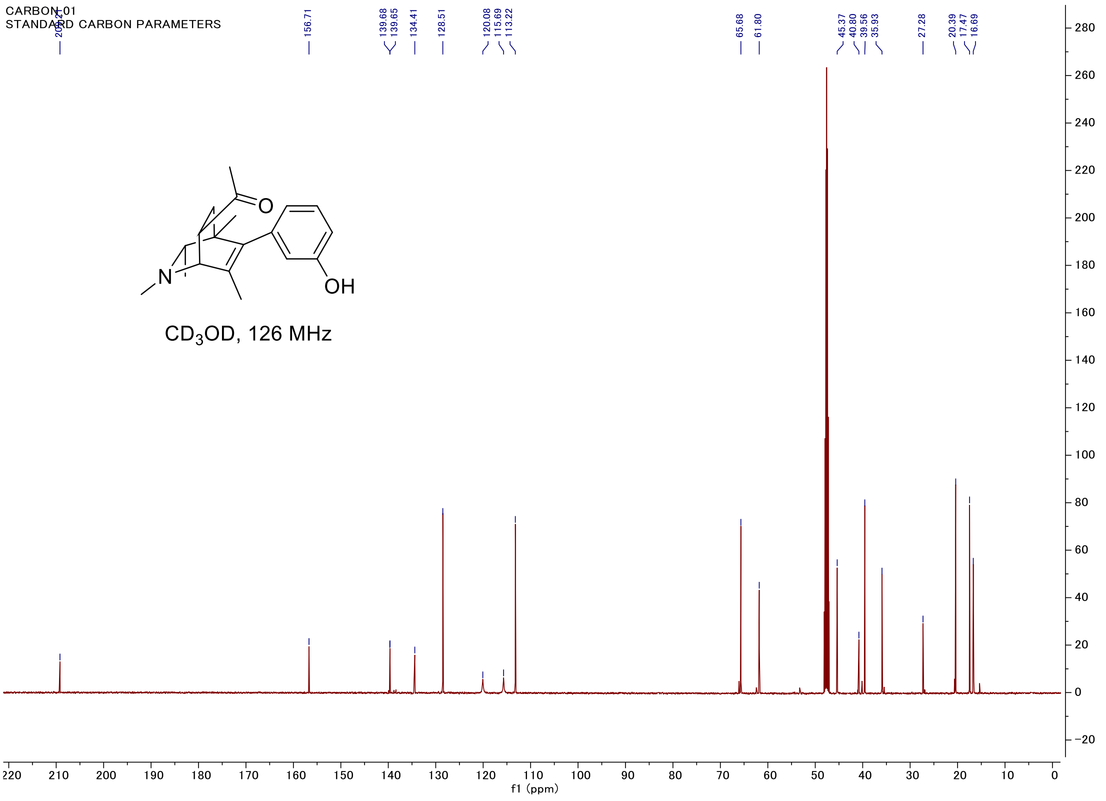

#### C-H COSY

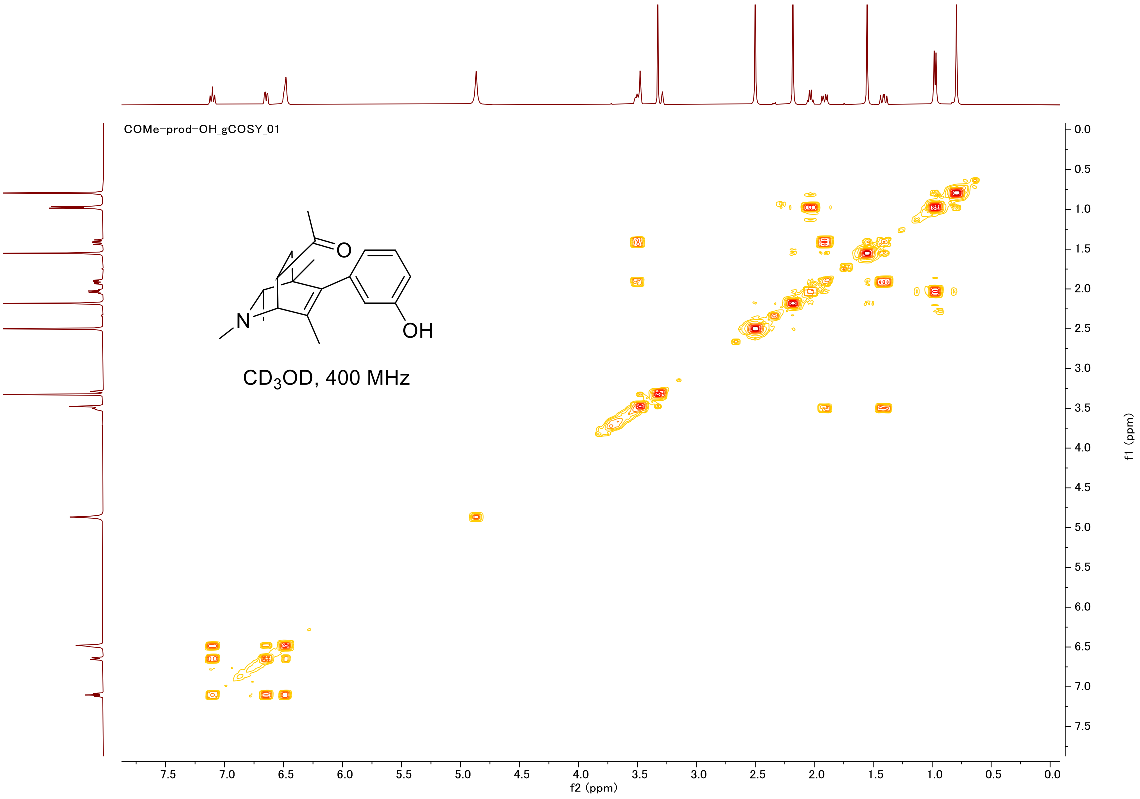
COMe-prod-OH_gCOSY_01
CD3OD, 400 MHz
OH
O
N
f2 (ppm)
f1 (ppm)

#### HSQC

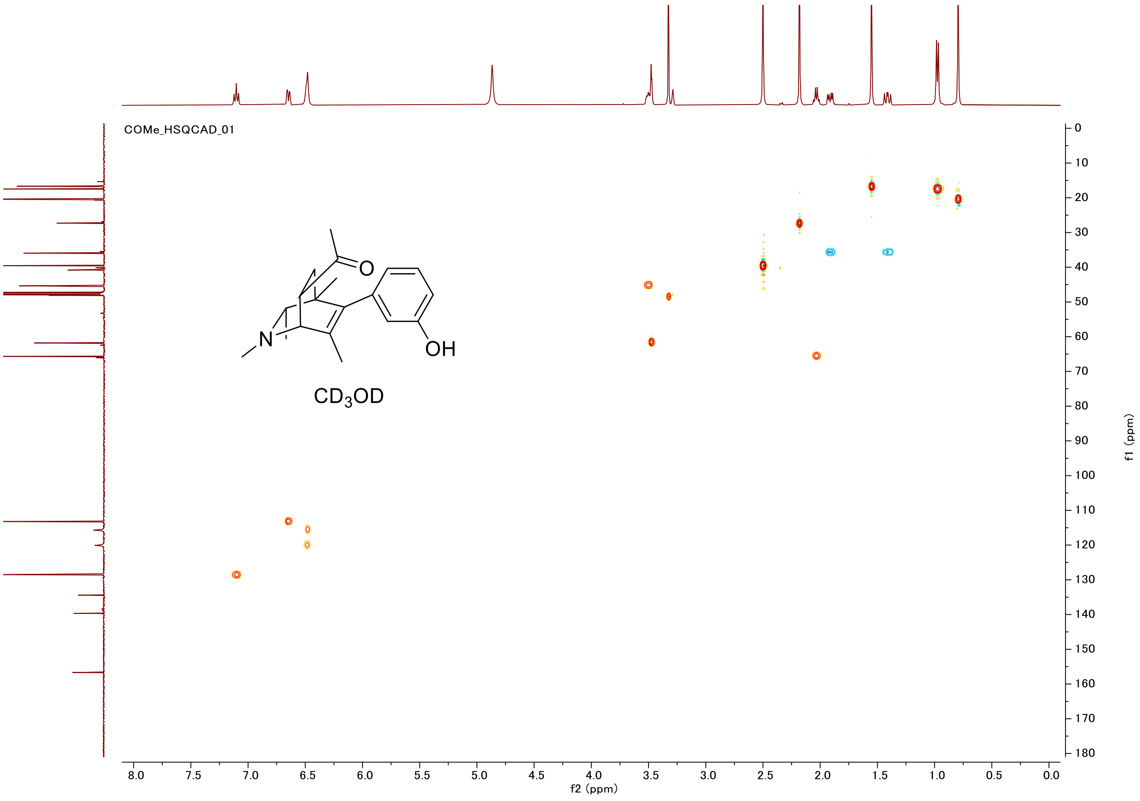
COMe_HSQCAD_01
CD3OD
OH
O
N
f2 (ppm)
f1 (ppm)

#### Proton (#025)

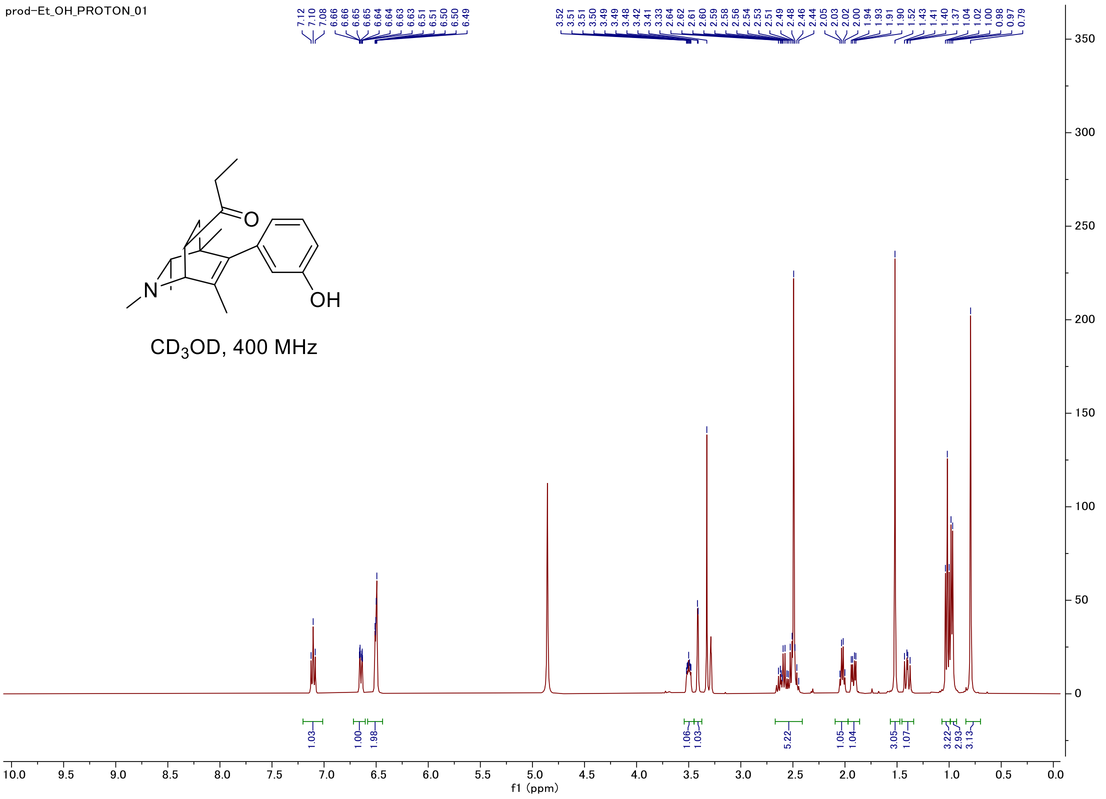


#### Carbon

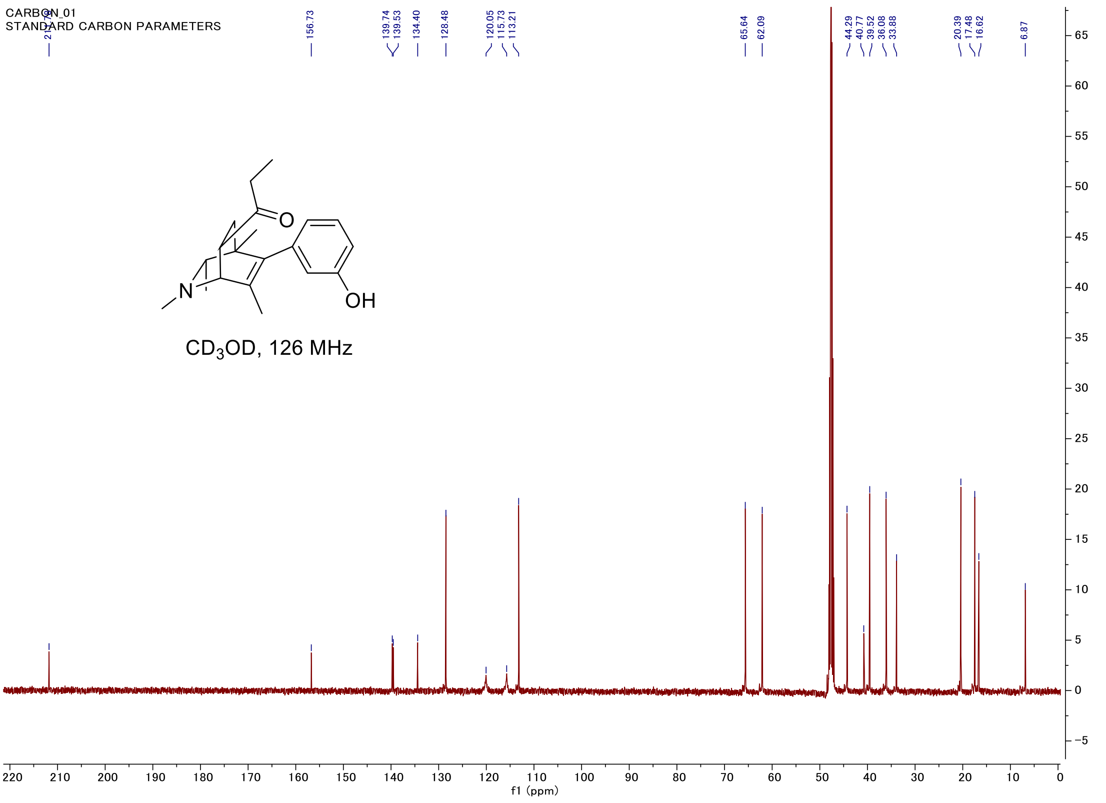

#### C-H COSY

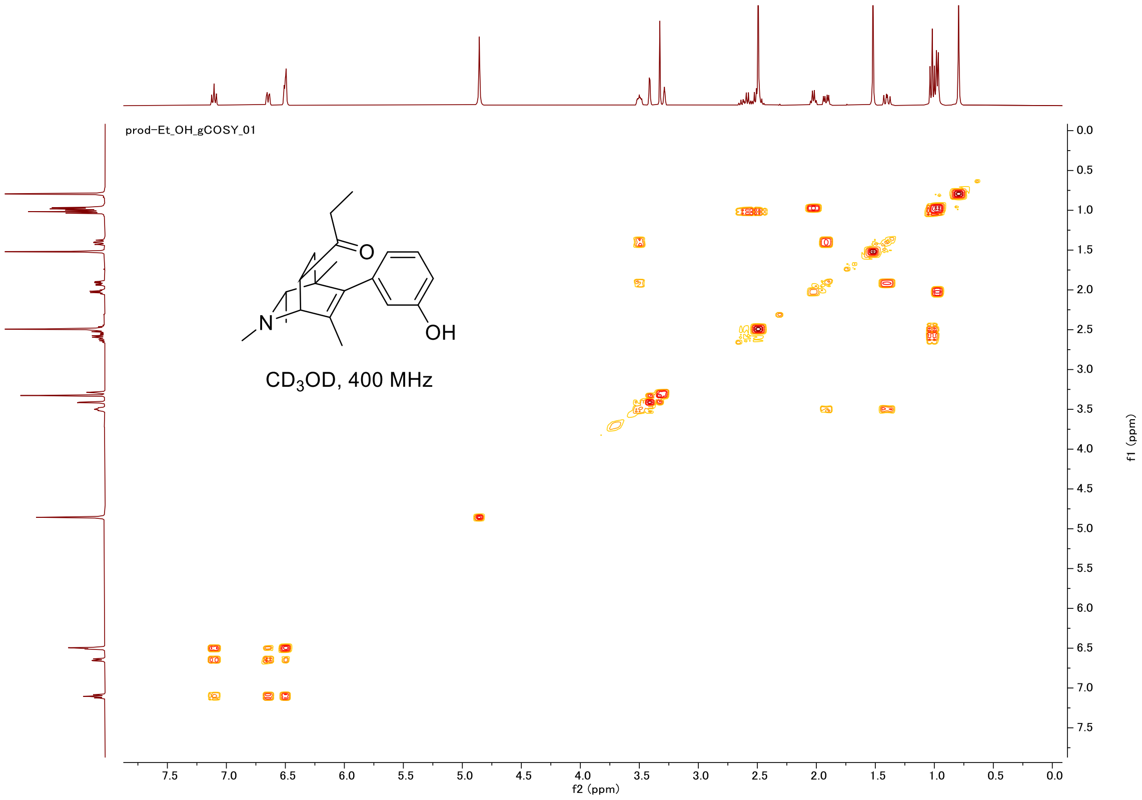
prod-Et_OH_gCOSY_01
N
O
OH
CD3OD, 400 MHz
f2 (ppm)
f1 (ppm)

#### HSQC

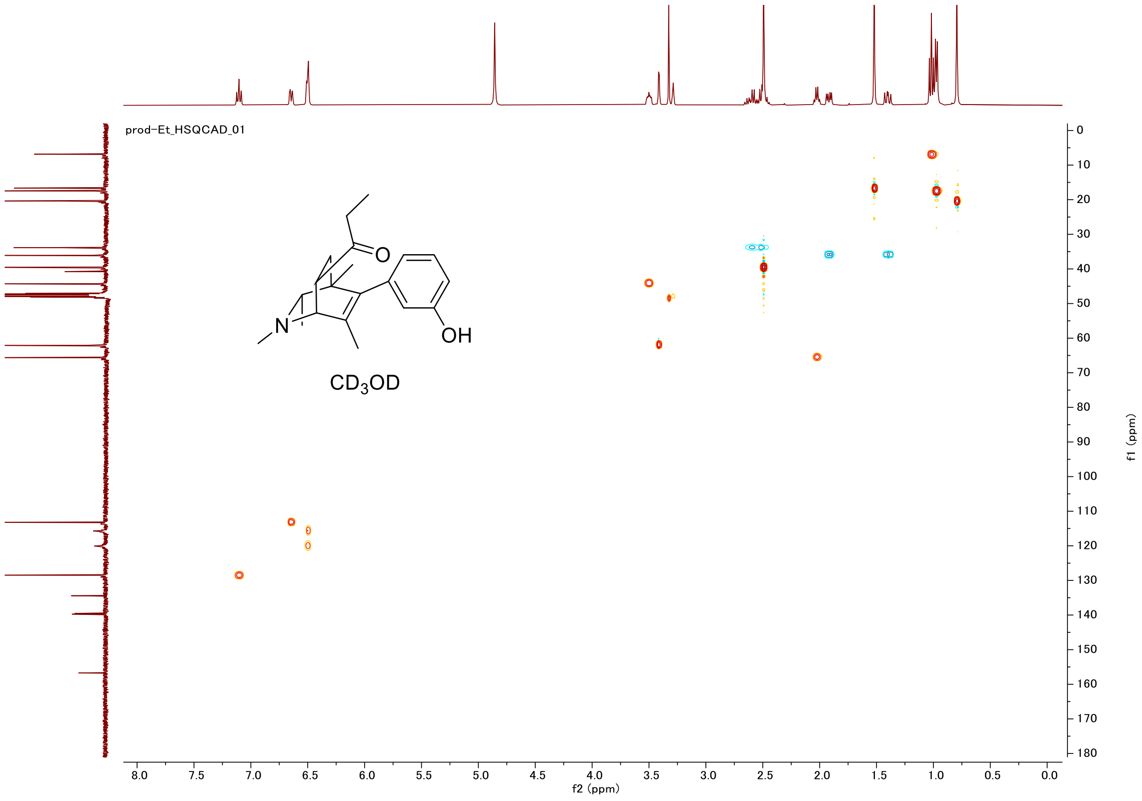
prod-Et_HSQCAD_01
N
O
OH
CD3OD
f2 (ppm)
f1 (ppm)

#### Proton (#026)

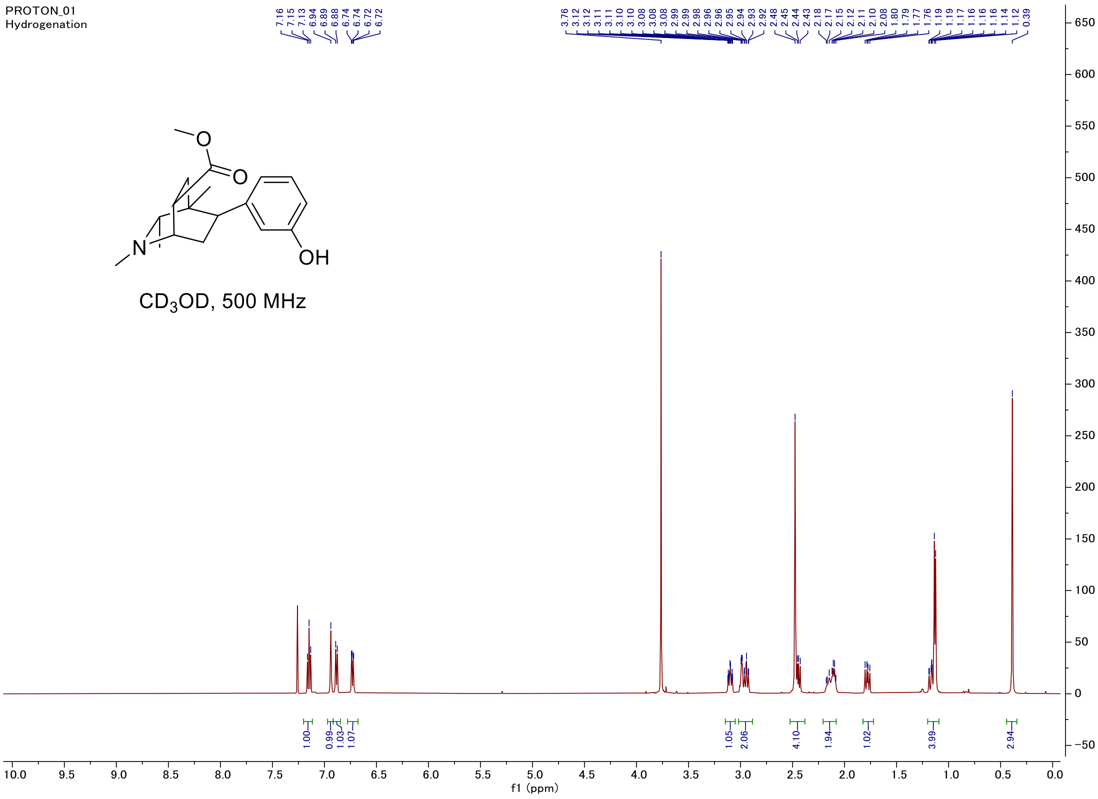


#### Carbon

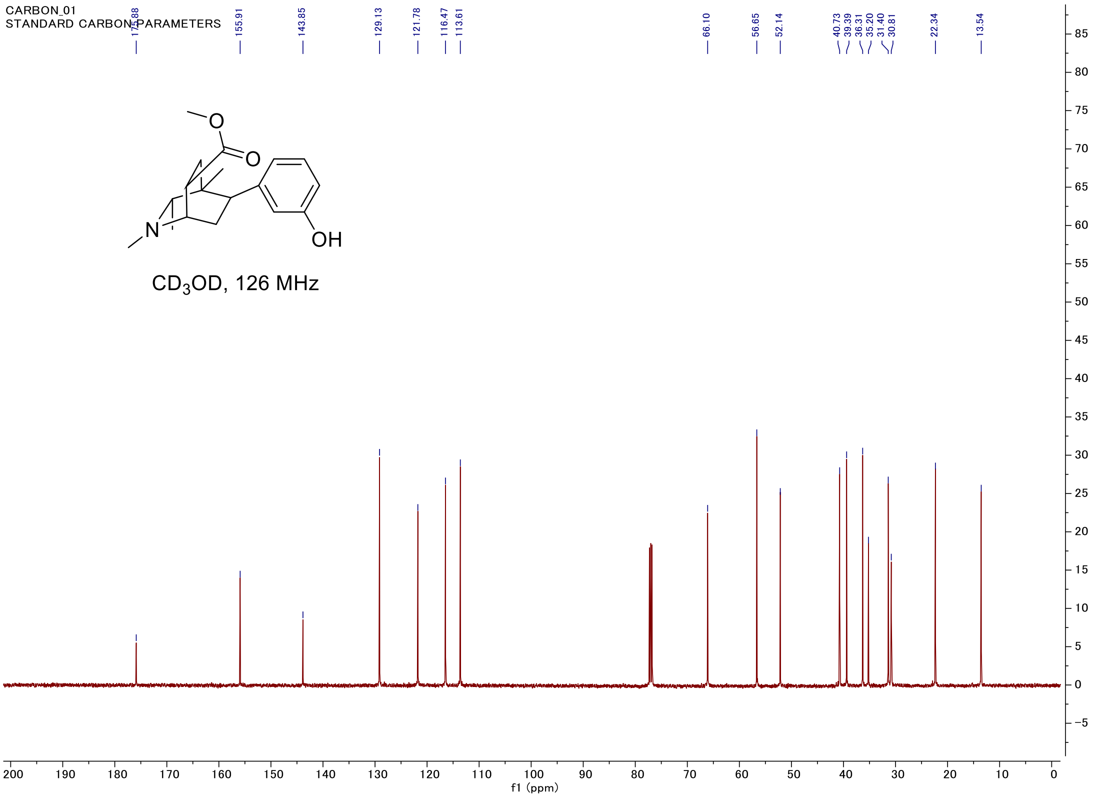

#### C-H COSY

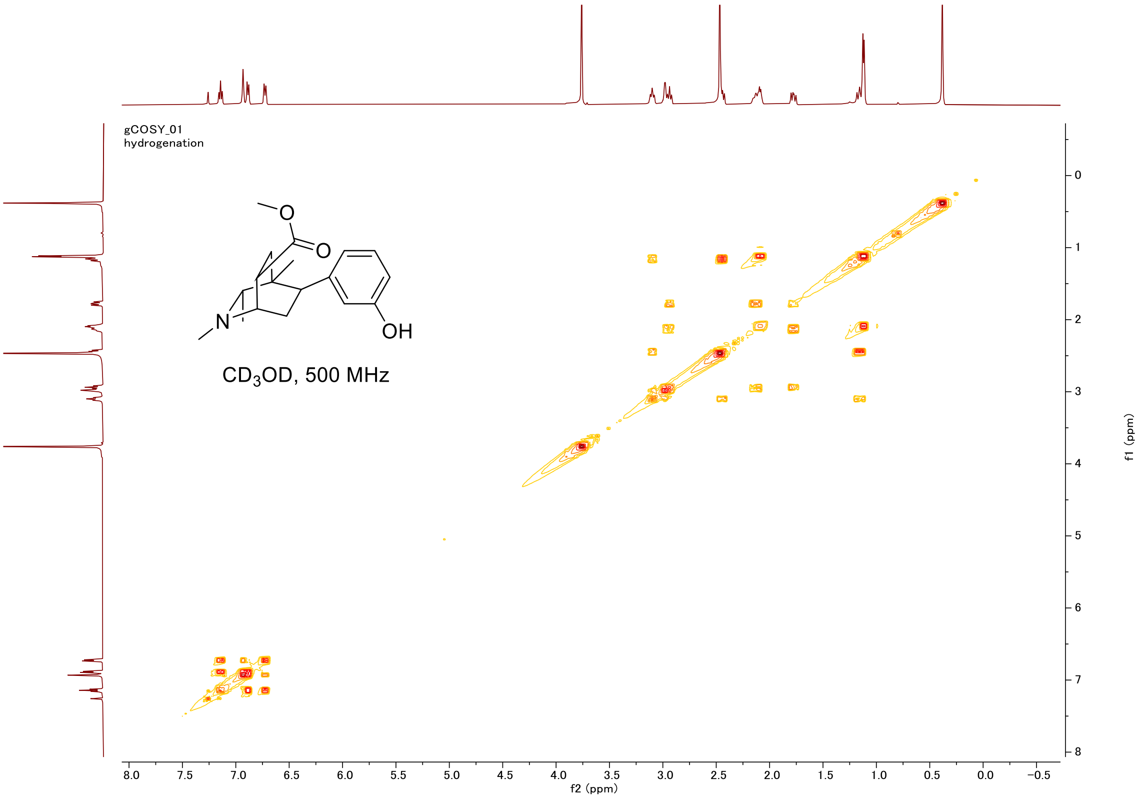

gCOSY_01
hydrogenation
CD3OD, 500 MHz
OH
N
O
f2 (ppm)
f1 (ppm)


#### HSQC

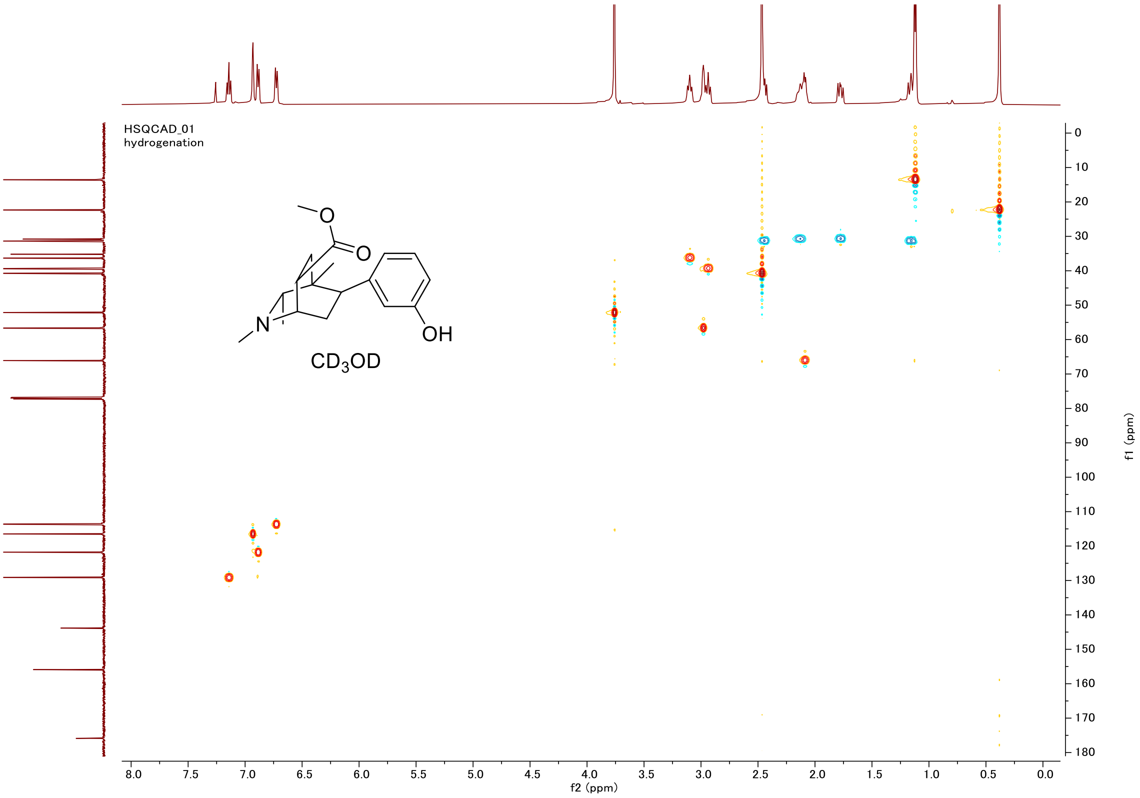

HSQCAD_01
hydrogenation
CD3OD
OH
N
O
f2 (ppm)
f1 (ppm)

#### Proton (#027)

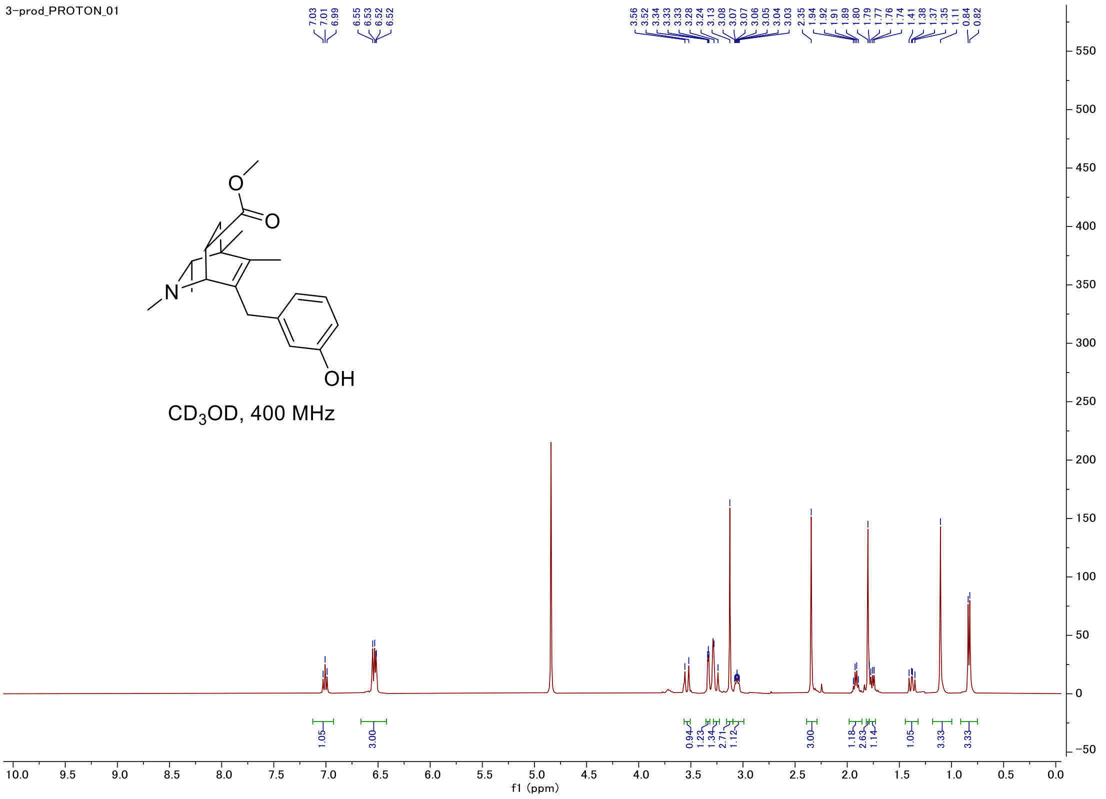


#### Carbon

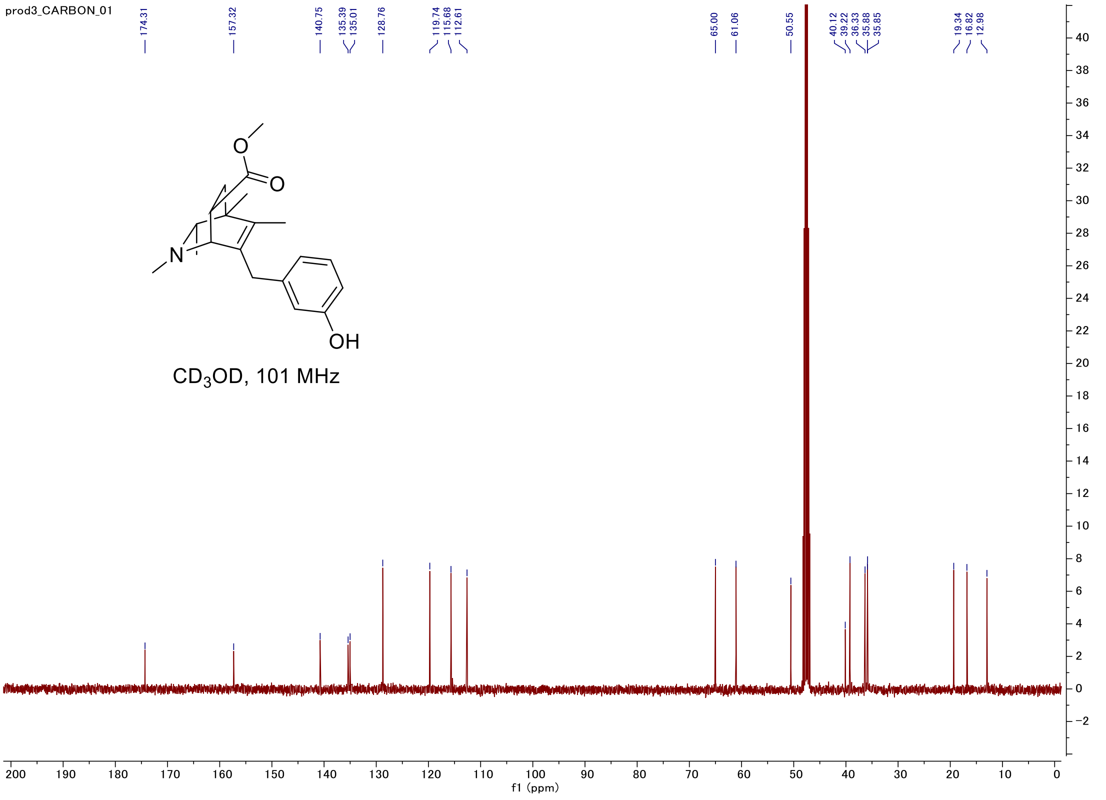

#### C-H COSY

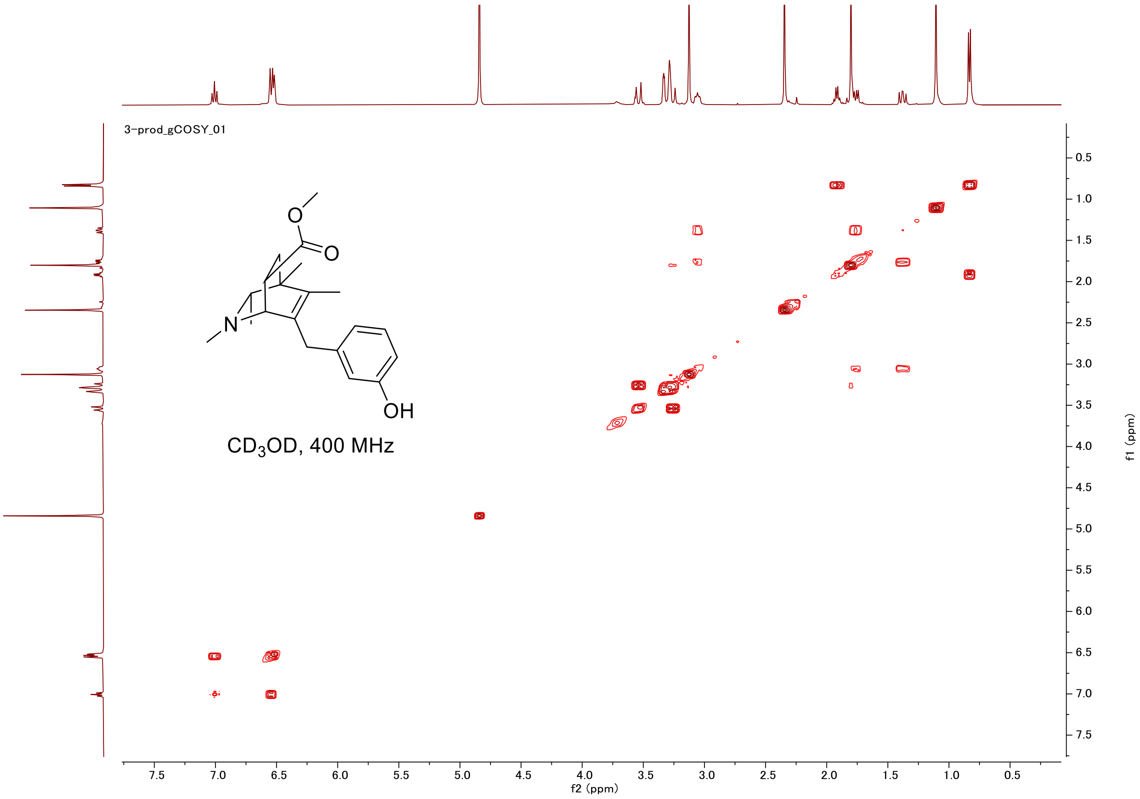
3-prod_gCOSY_01
O
O
N
OH
CD3OD, 400 MHz
f2 (ppm)
f1 (ppm)

#### HSQC

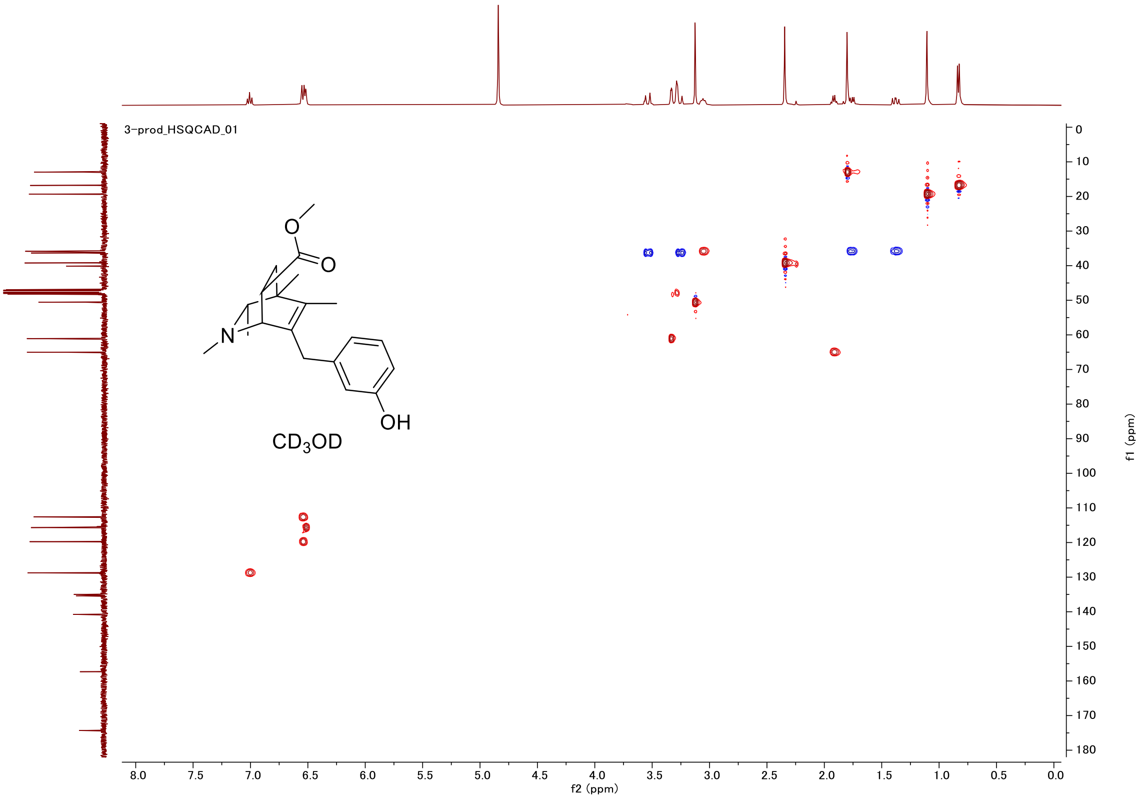
3-prod_HSQCAD_01
O
O
N
OH
CD3OD
f2 (ppm)
f1 (ppm)

#### Proton (#028)

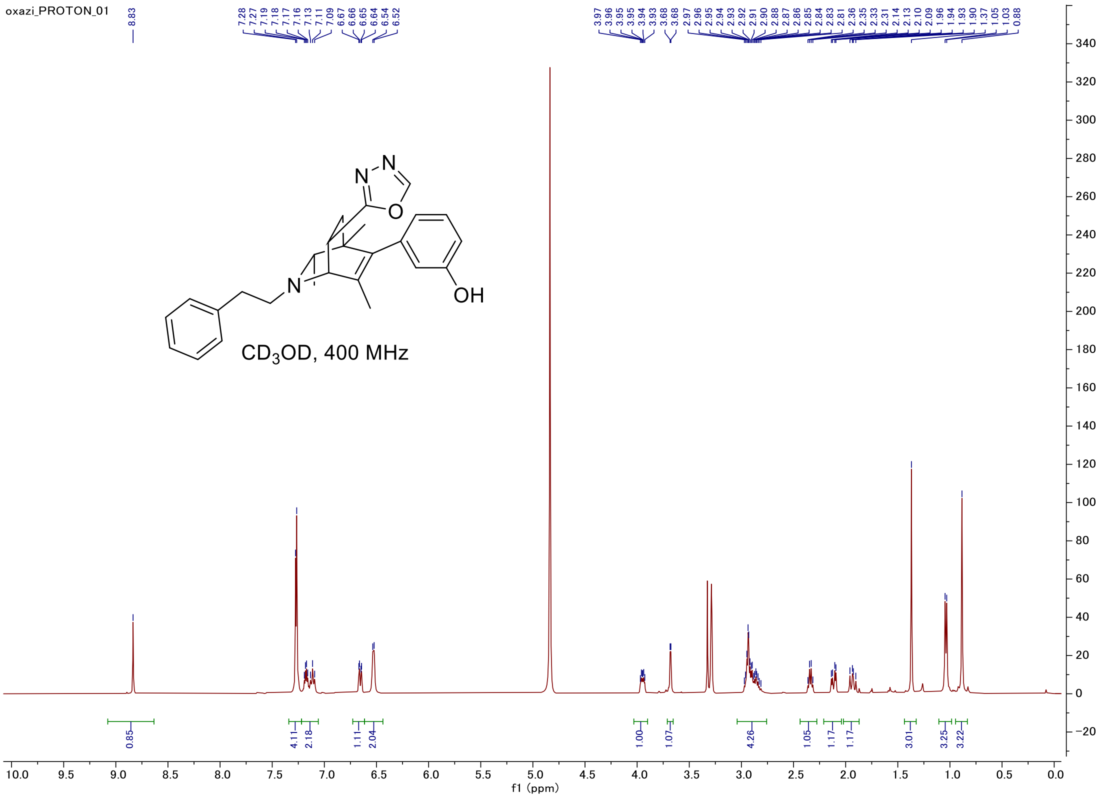


#### Carbon

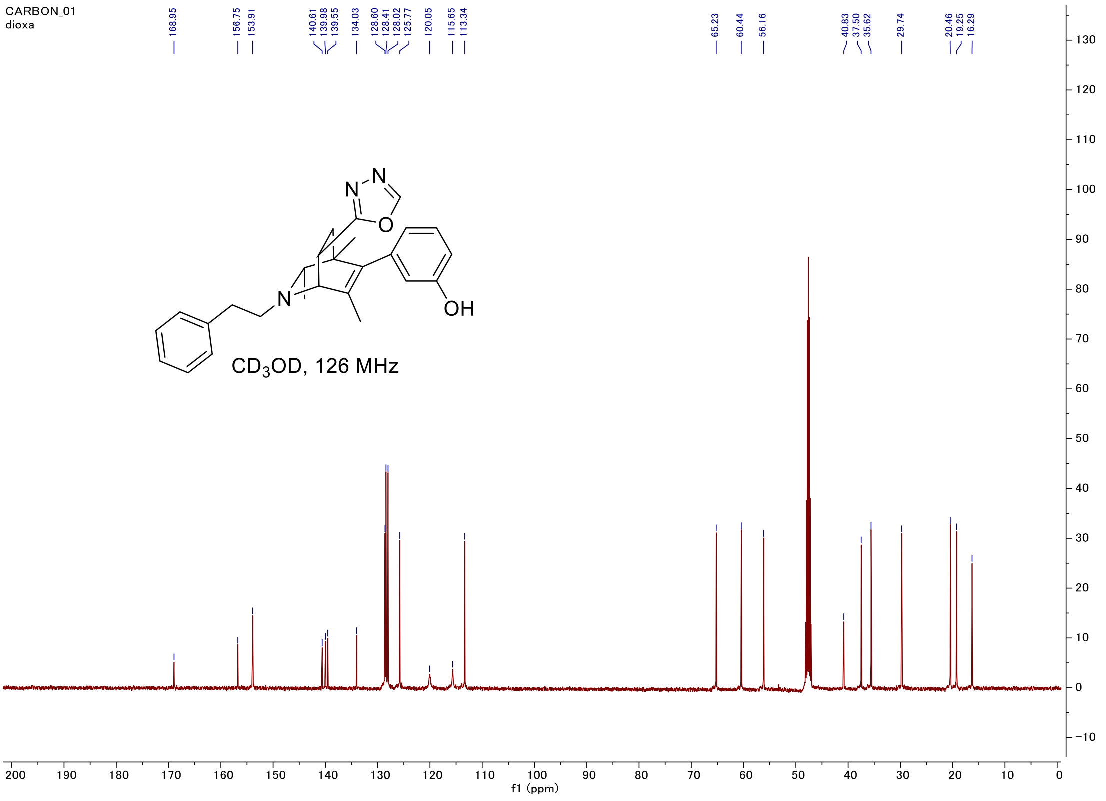

#### C-H COSY

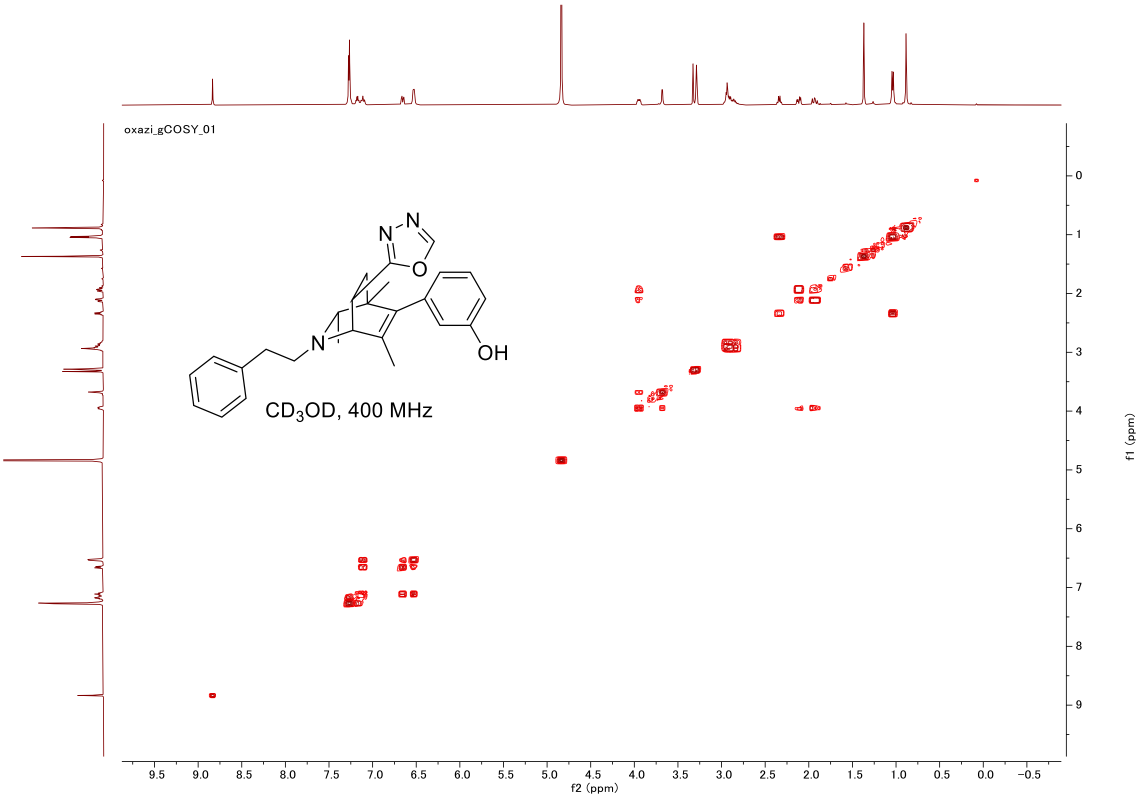
oxazi_gCOSY_01
CD3OD, 400 MHz
OH
f2 (ppm)
f1 (ppm)

#### HSQC

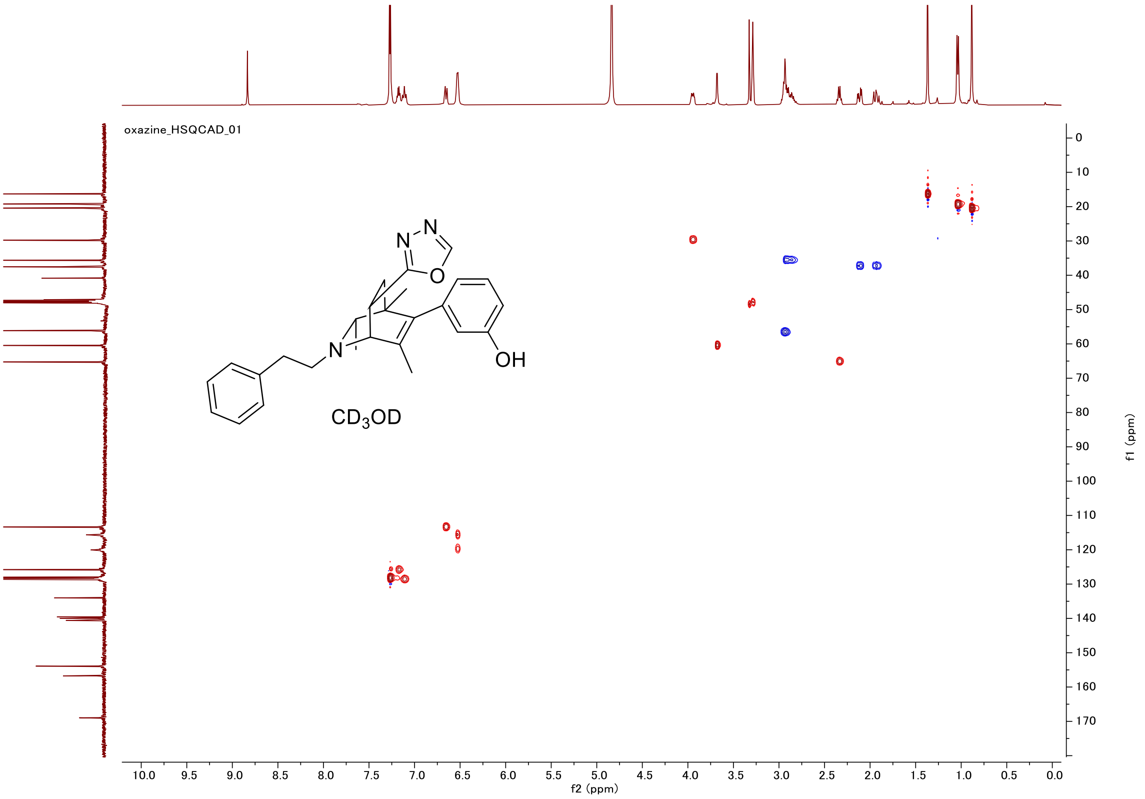
oxazine_HSQCAD_01
CD3OD
OH
f2 (ppm)
f1 (ppm)

### Proton (#029)

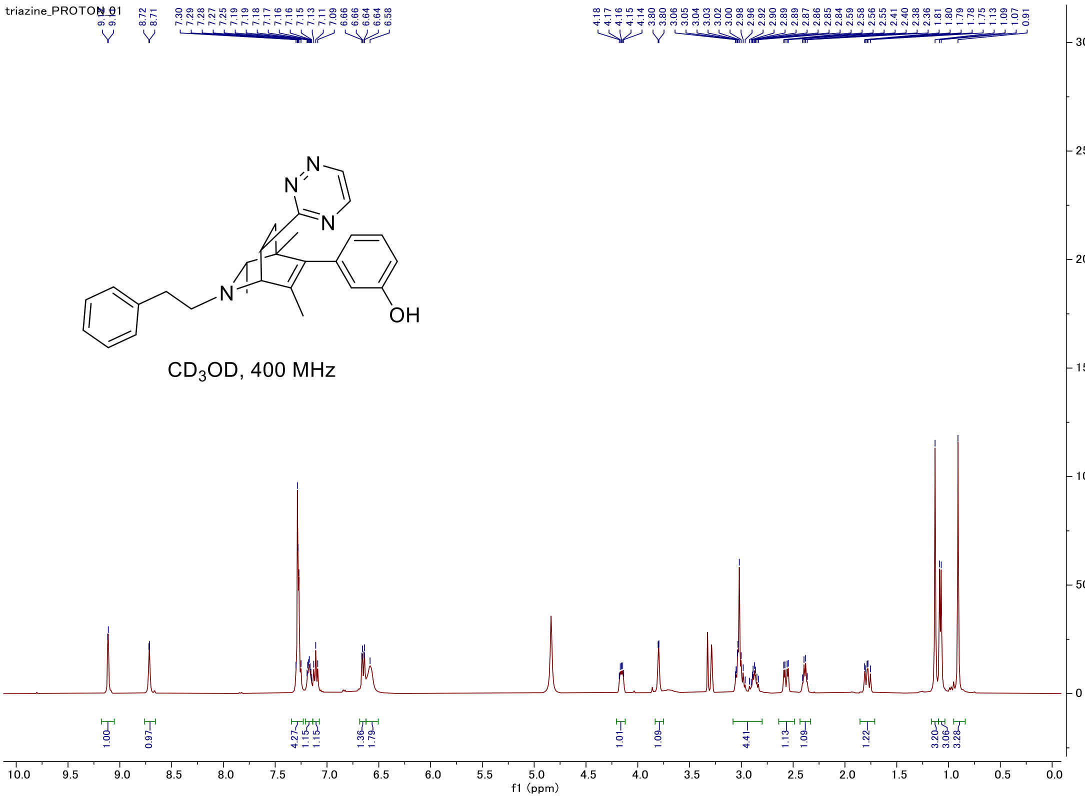


### Carbon

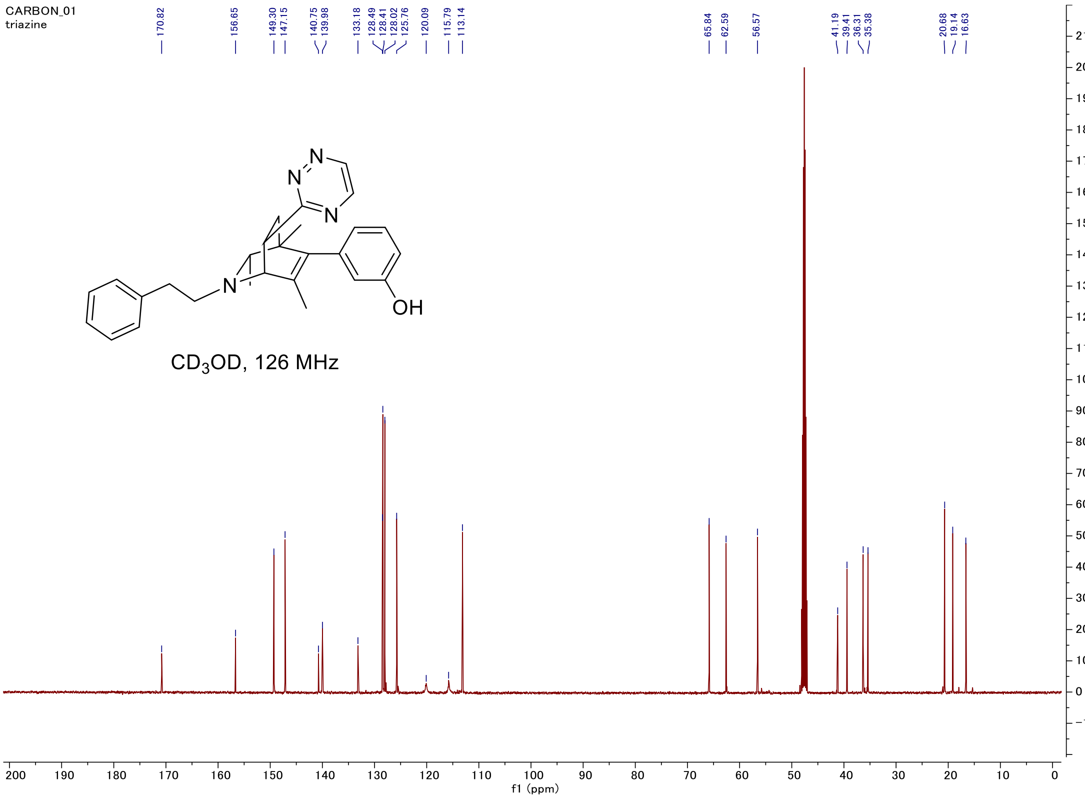

#### C-H COSY

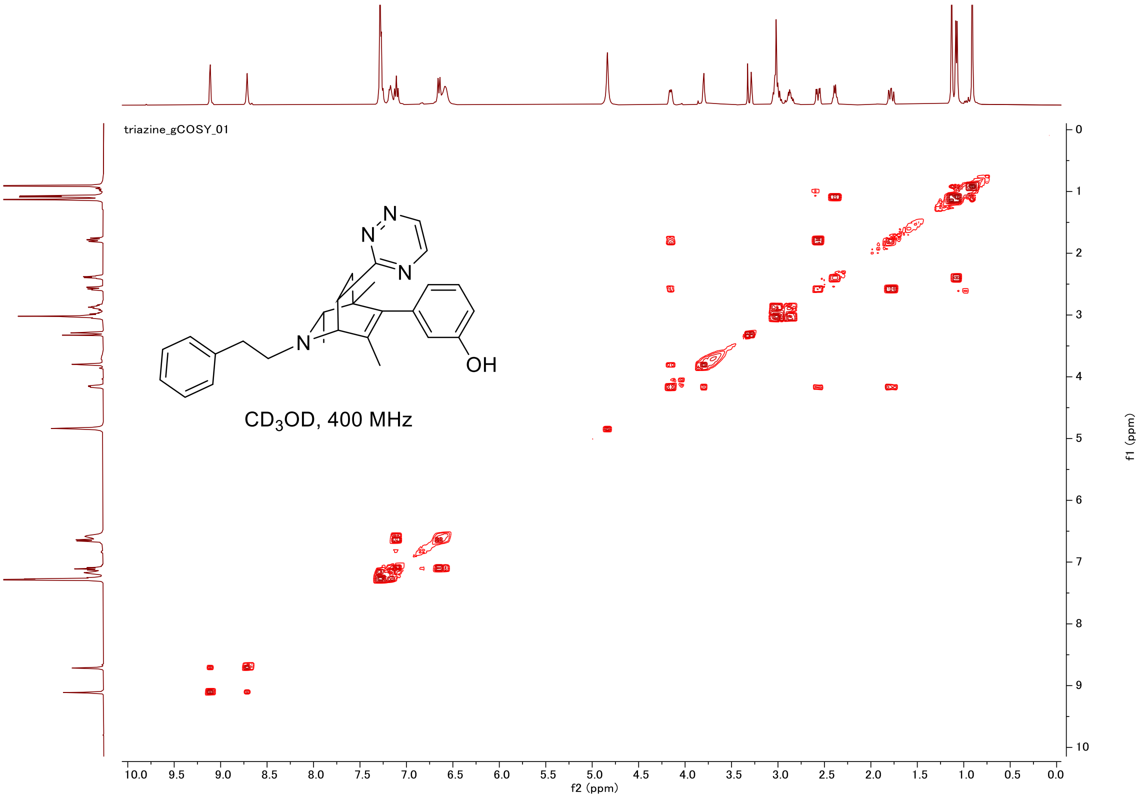

triazine_gCOSY_01
CD$_3$OD, 400 MHz
OH
f2 (ppm)
f1 (ppm)


#### HSQC

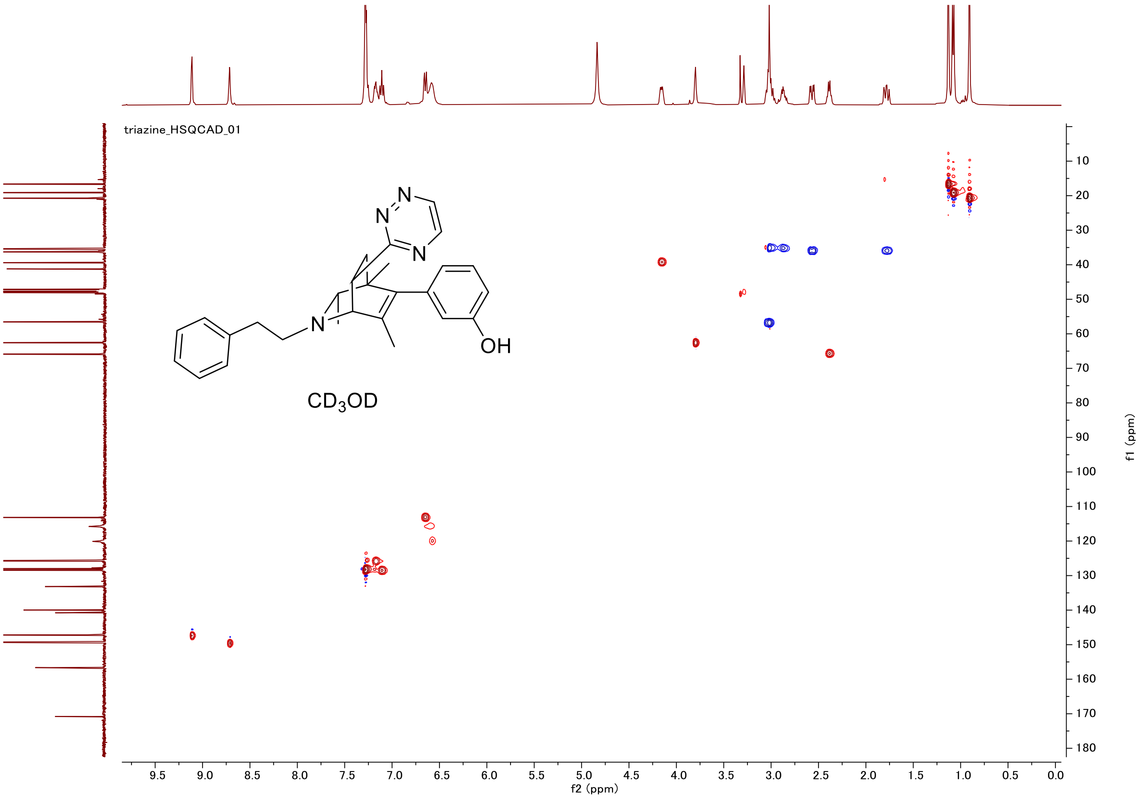

triazine_HSQCAD_01
CD$_3$OD
OH
f2 (ppm)
f1 (ppm)

#### Proton (#030)

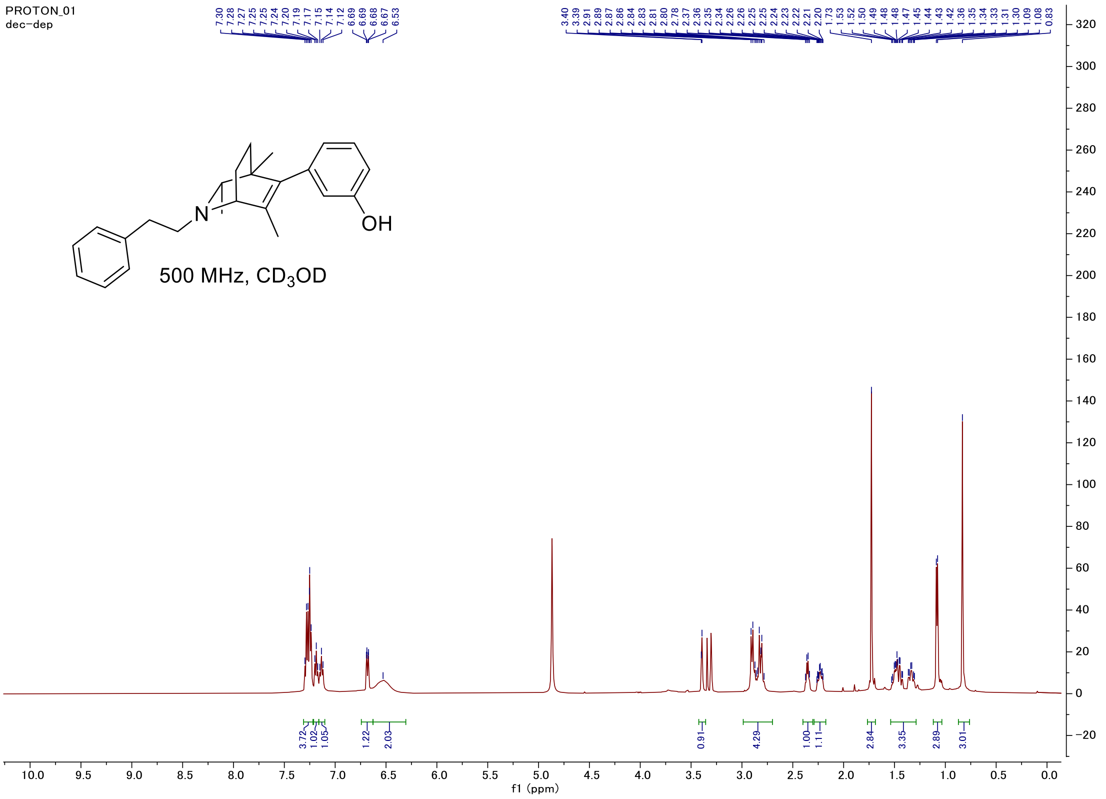


#### Carbon

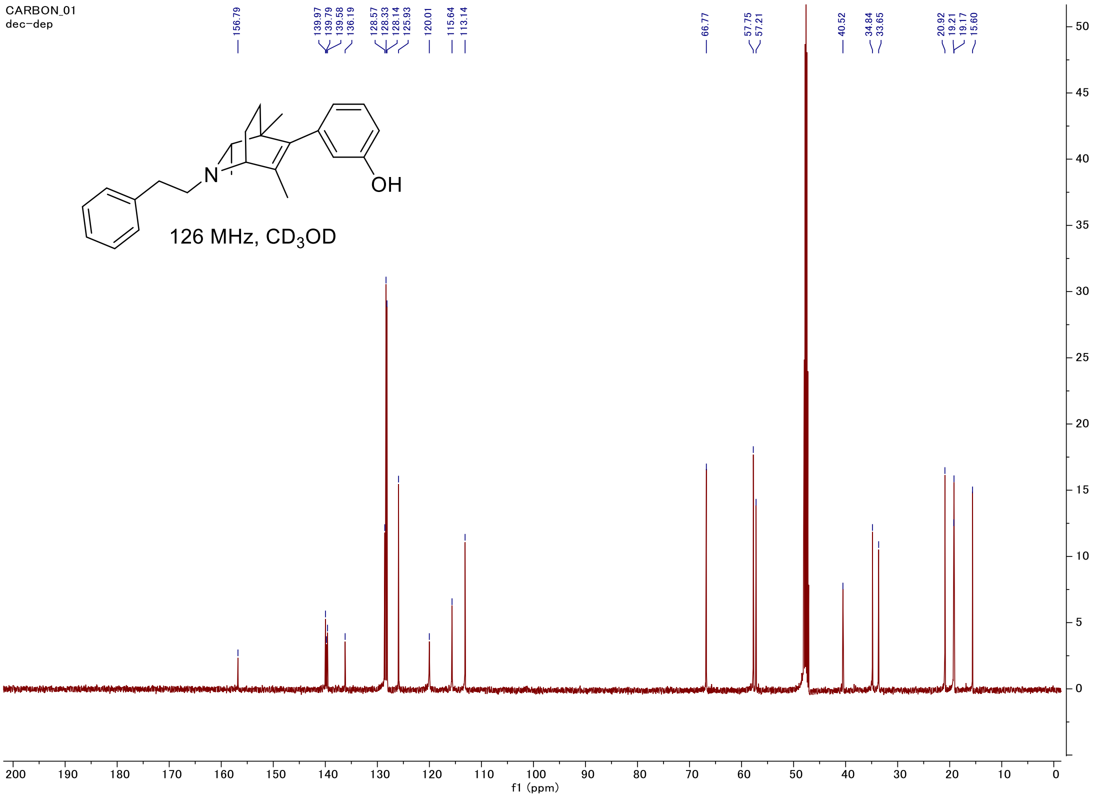

#### C-H COSY

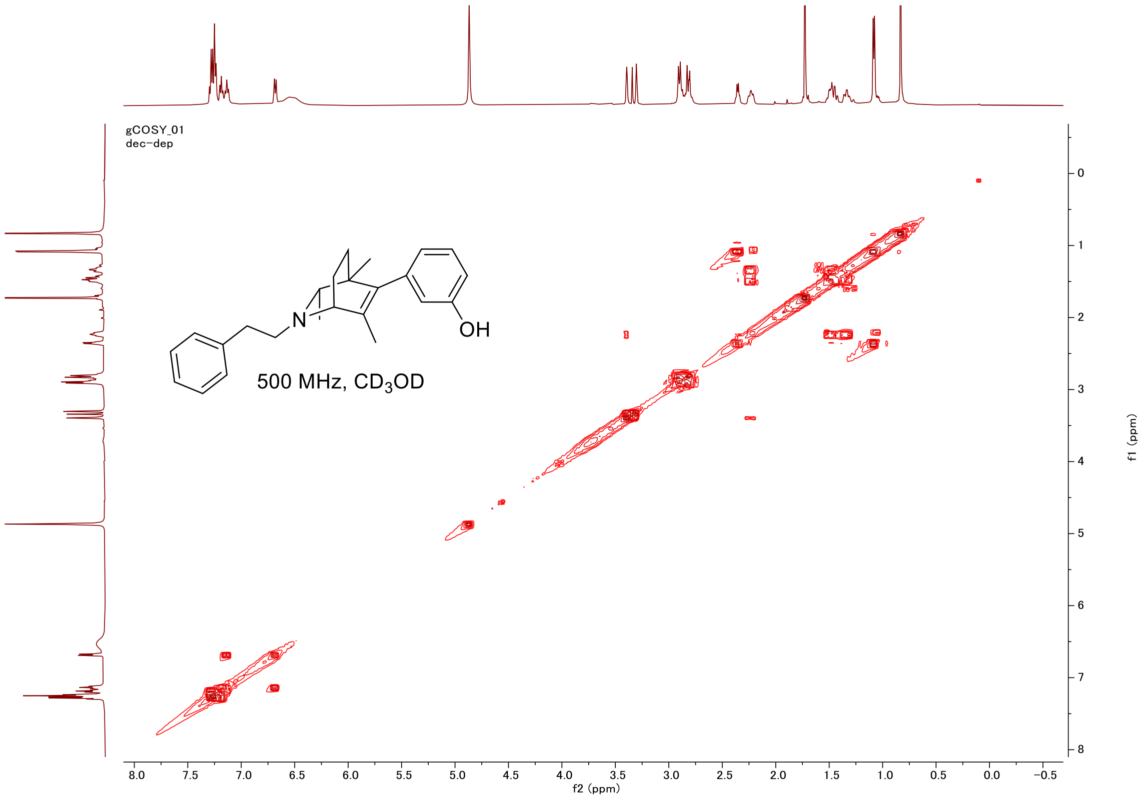
gCOSY_01
dec-dep
N
OH
500 MHz, CD3OD
f2 (ppm)
f1 (ppm)

#### HSQC

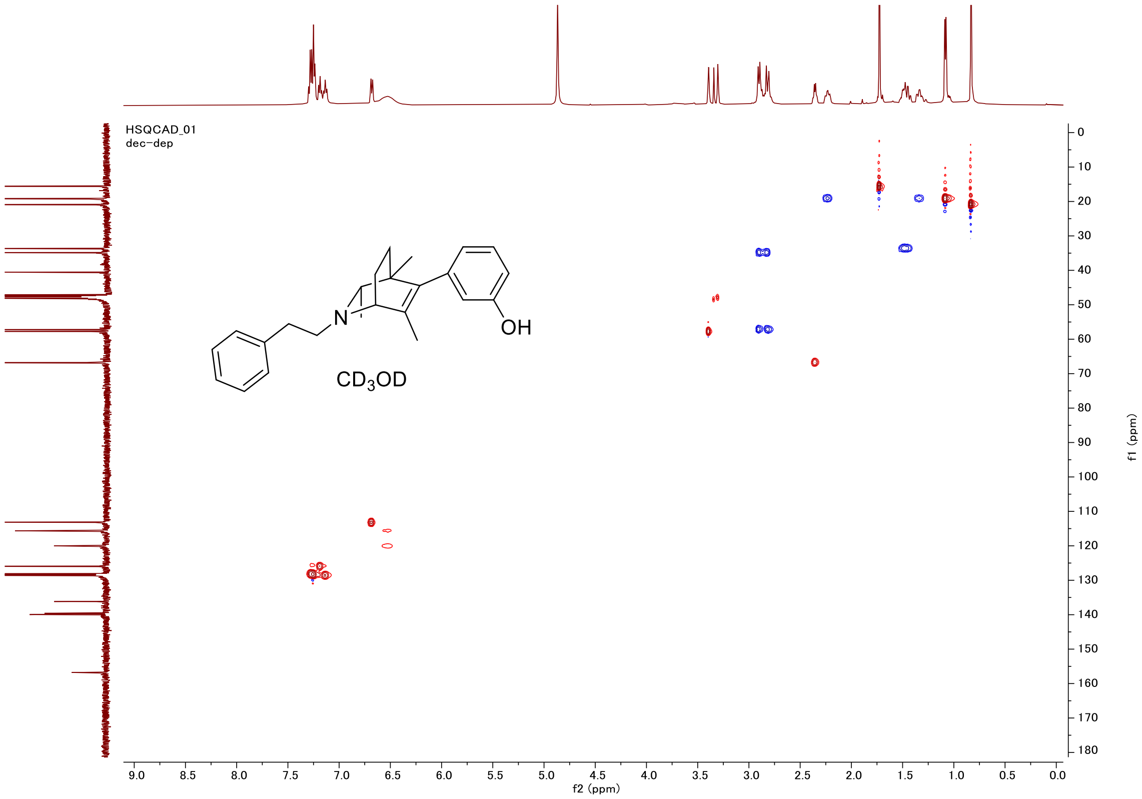
HSQCAD_01
dec-dep
N
OH
CD3OD
f2 (ppm)
f1 (ppm)

#### Proton (#031)

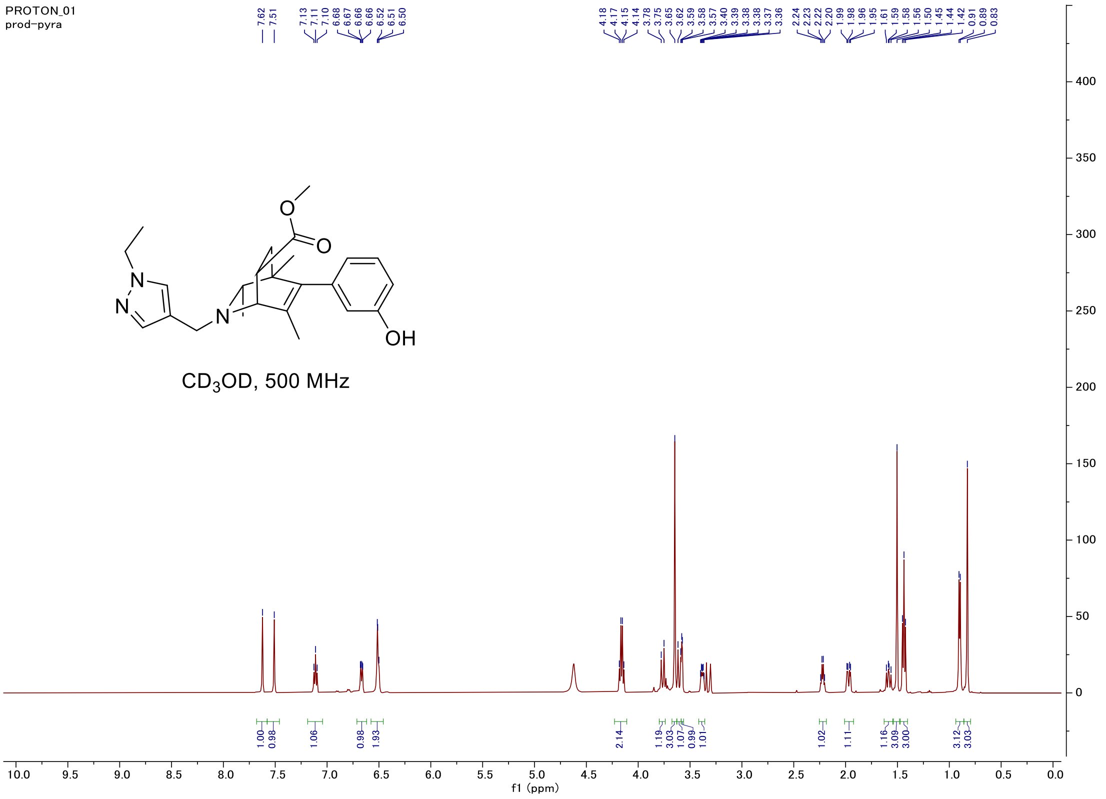


#### Carbon

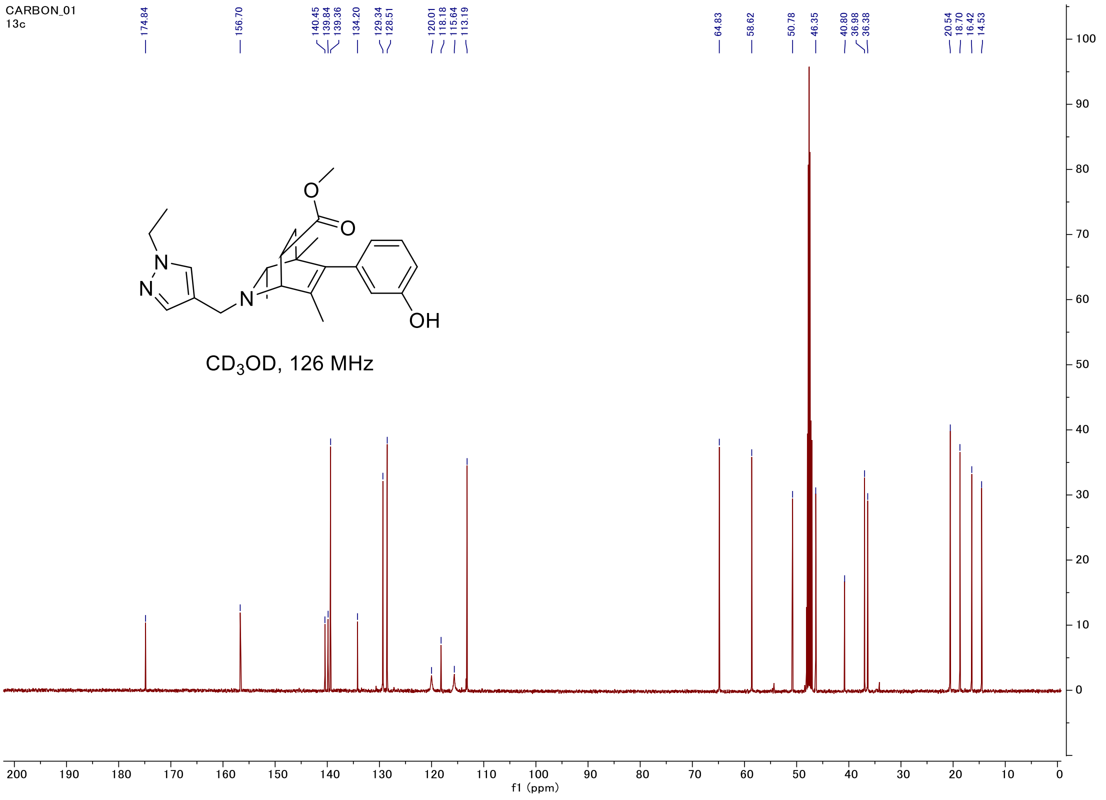

#### C-H COSY

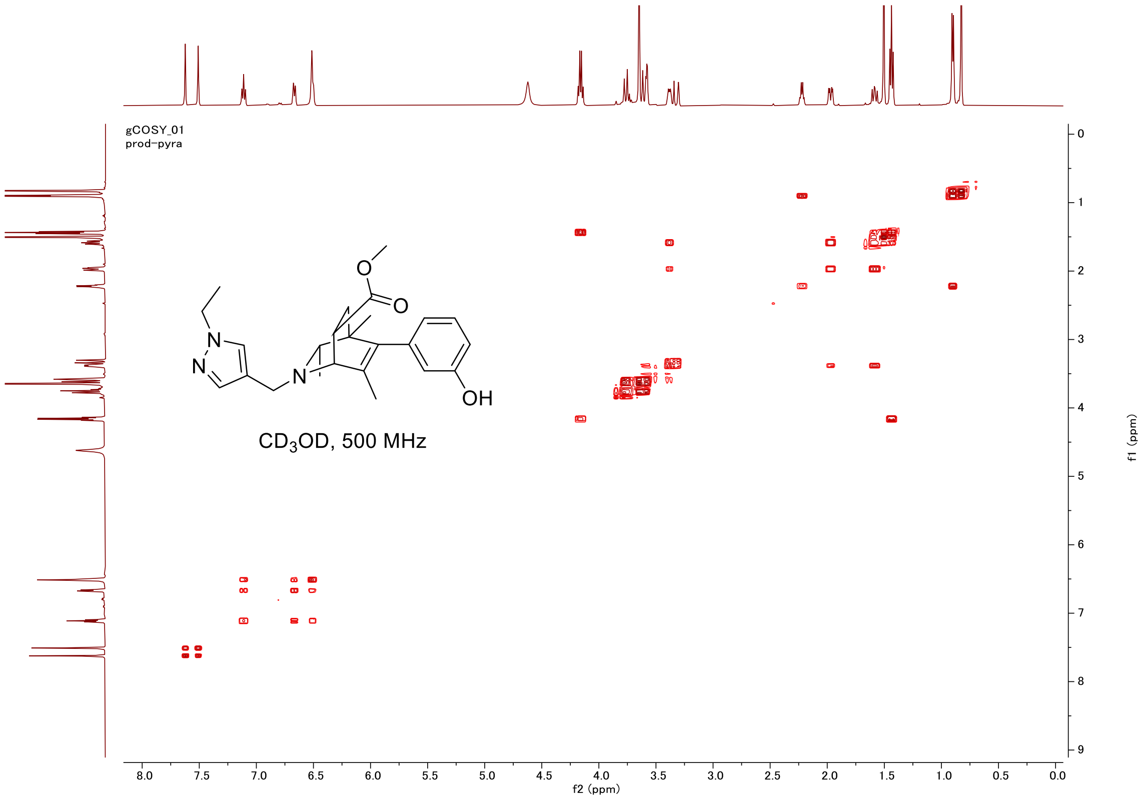
gCOSY_01
prod-pyra
CD$_3$OD, 500 MHz
OH
f2 (ppm)
f1 (ppm)

#### HSQC

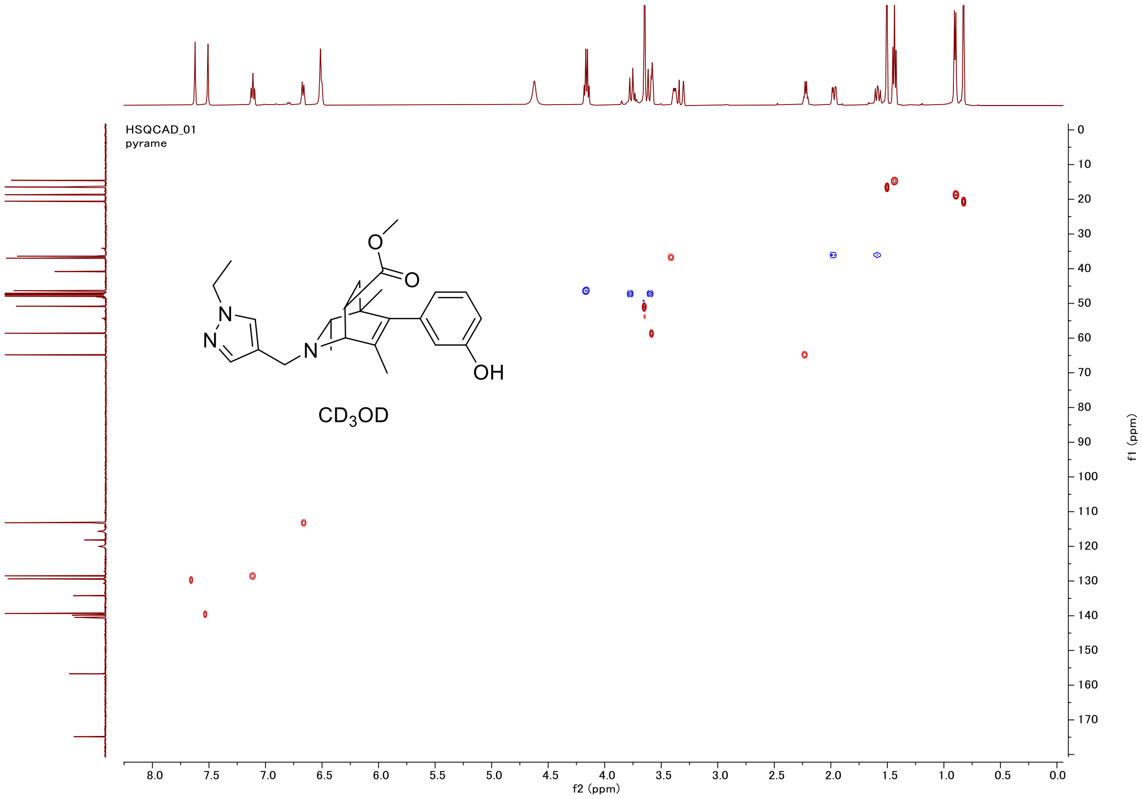
HSQCAD_01
pyrame
CD$_3$OD
OH
f2 (ppm)
f1 (ppm)

#### Proton (#036)

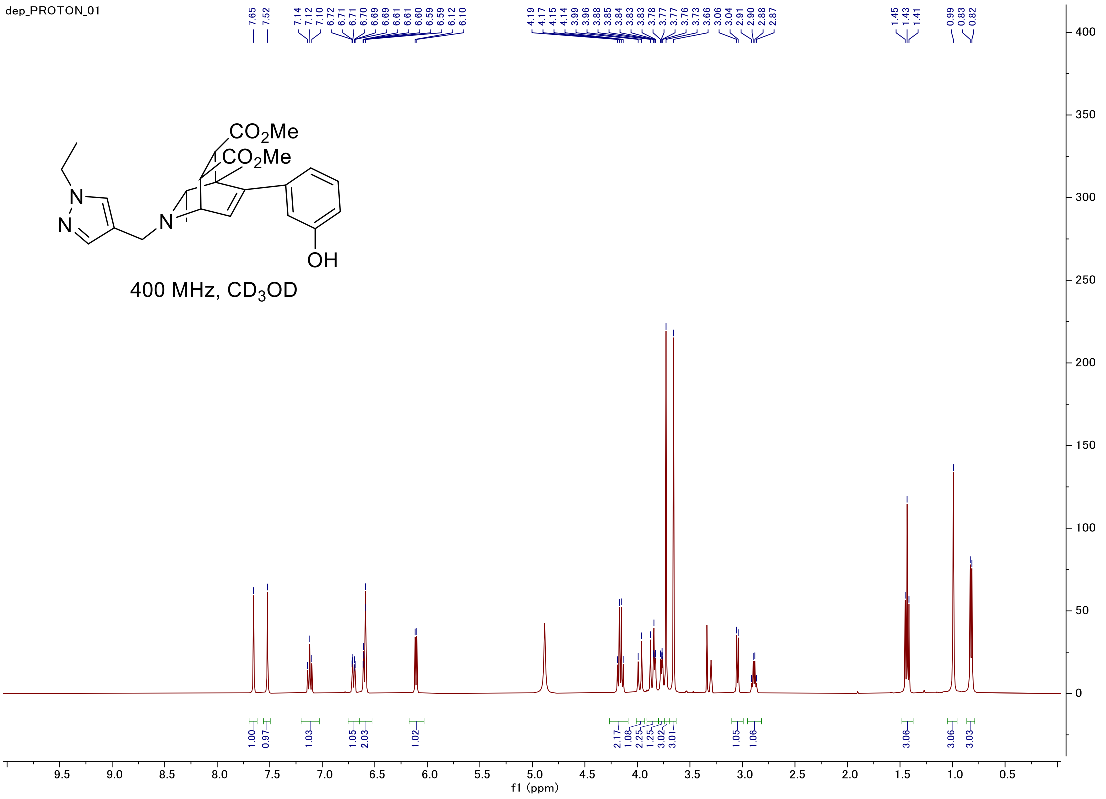


#### Carbon

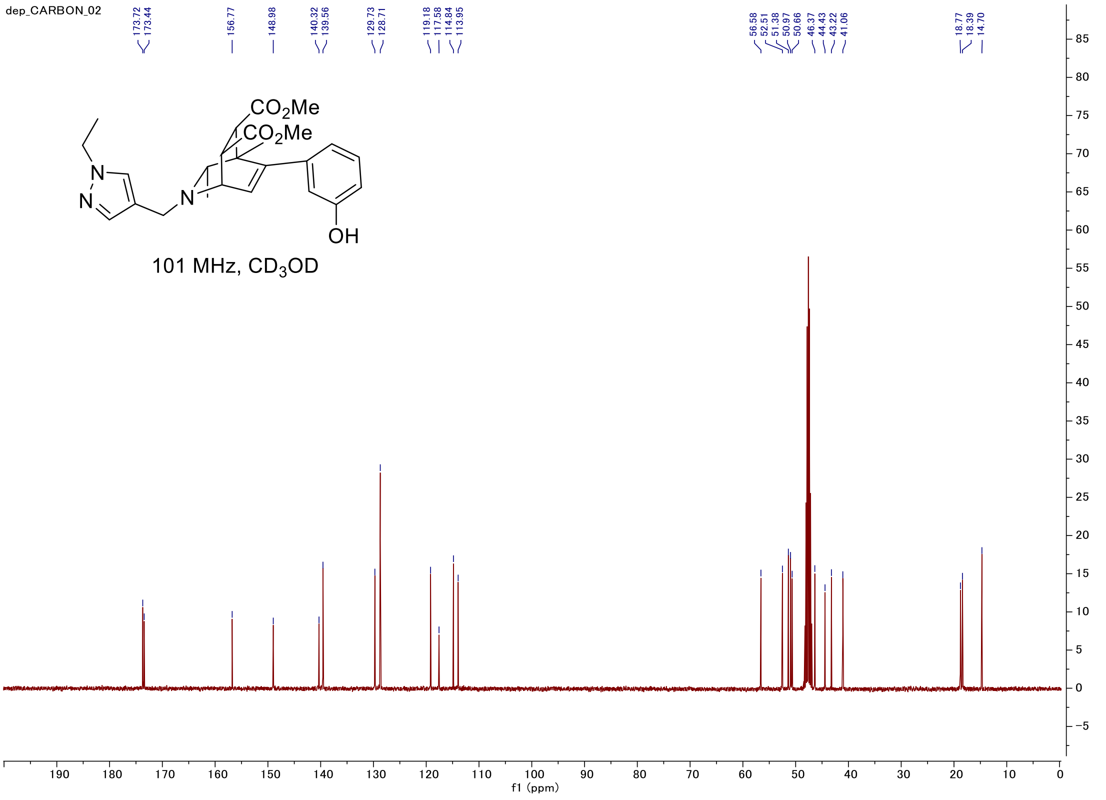

#### C-H COSY

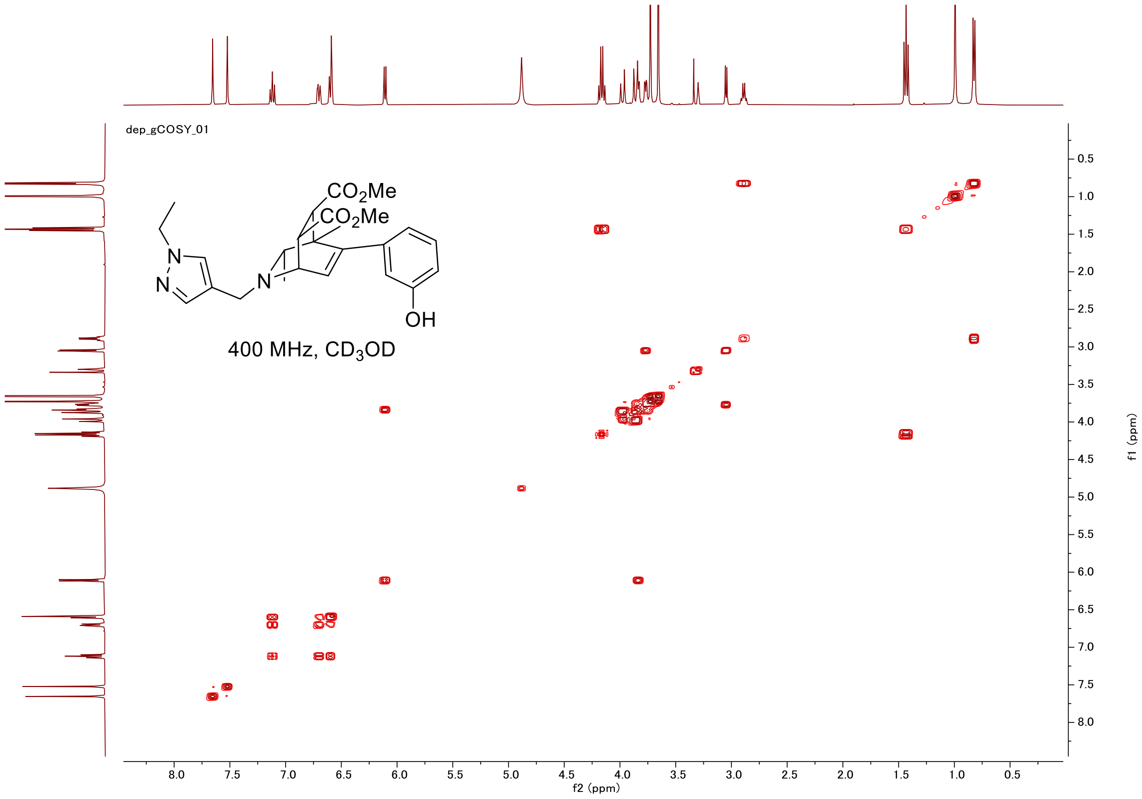
dep_gCOSY_01
$CO_2Me$
$CO_2Me$
N
N
N
OH
400 MHz, $CD_3OD$
f2 (ppm)
f1 (ppm)

#### HSQC

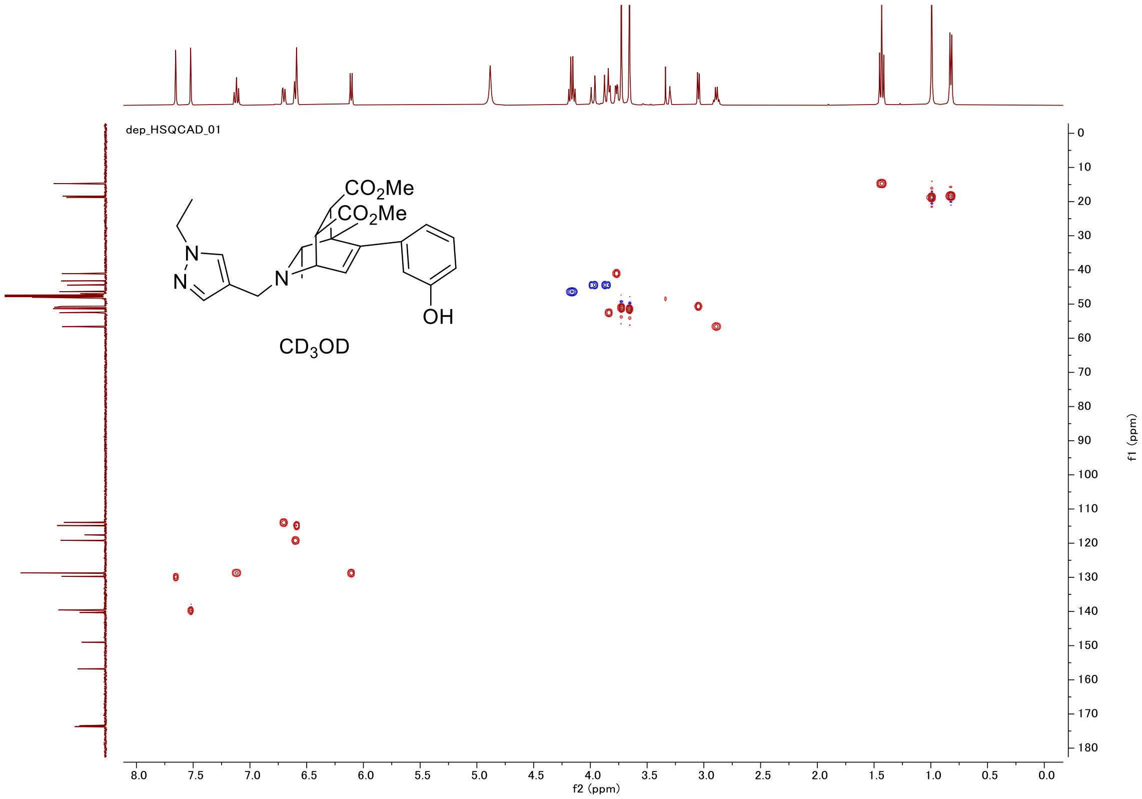
dep_HSQCAD_01
$CO_2Me$
$CO_2Me$
N
N
N
OH
$CD_3OD$
f2 (ppm)
f1 (ppm)
